# Uncoupling from transcription protects polyadenylation site cleavage from inhibition by DNA damage

**DOI:** 10.1101/2022.05.26.492040

**Authors:** Rym Sfaxi, Biswendu Biswas, Galina Boldina, Mandy Cadix, Nicolas Servant, Huimin Chen, Daniel R. Larson, Martin Dutertre, Caroline Robert, Stéphan Vagner

## Abstract

Pre-mRNA 3’-end processing by cleavage and polyadenylation (CPA) is a nuclear process in which RNA polymerase II (Pol II) transcripts are cleaved at the polyadenylation site (PAS cleavage) before addition of a poly(A) tail. While PAS cleavage is usually coupled to transcription termination, for some pre-mRNAs it occurs post-transcriptionally, *i.e*. after pre-mRNA release from chromatin to nucleoplasm through a downstream co-transcriptional cleavage (CoTC) event. DNA-damaging agents such as ultraviolet-C (UV) irradiation trigger rapid shutdown of pre-mRNA 3’-end processing. However, specific compensatory mechanisms exist to ensure efficient 3’-end processing for some pre-mRNAs encoding proteins involved in the DNA damage response (DDR), such as the p53 tumor suppressor protein. Here, we show that PAS cleavage of the *p53* pre-mRNA occurs in part post-transcriptionally, in a PCF11-independent manner, in the nucleoplasm, following a CoTC-type event. Upon UV-irradiation, cells with an engineered deletion of the *p53* CoTC site exhibit impaired 3’-end processing of the *p53* pre-mRNA, decreased mRNA and protein levels of p53 and its transcriptional target, p21, and altered cell cycle progression. Finally, using a transcriptome-wide analysis of PAS cleavage, we identified additional-including DDR related-pre-mRNAs whose PAS cleavage is maintained in response to UV and occurs post-transcriptionally. These findings indicate that CoTC-type cleavage of pre-mRNAs, followed by PAS cleavage in the nucleoplasm, allows specific pre-mRNAs to escape 3’-end processing inhibition in response to UV-induced DNA damage.

## Introduction

During UV-induced DNA damage and other genotoxic stresses, several steps in eukaryotic gene expression are repressed. Although global, this repression is associated with an upregulation of the expression of genes encoding proteins that are essential for the adaptation and response to stress. For instance, despite the inhibition of pre-mRNA 3’-end processing observed in UV-treated cells (Kim *et al*, 2006; Kleiman, 1999; Kleiman & Manley, 2001; Nazeer *et al*, 2011), pre-mRNA 3’-end processing of the pre-mRNA encoding the p53 tumor suppressor (TP53) protein is specifically maintained (Decorsière *et al*, 2011; Newman *et al*, 2017). This maintenance requires several RNA binding proteins, *i.e*. the heterogeneous nuclear ribonucleoprotein (hnRNP) F/H family of proteins that bind to an RNA G-quadruplex forming sequence located downstream of the p53 polyadenylation site (Decorsière *et al*, 2011), as well as the DHX36 RNA/DNA helicase (Newman *et al*, 2017).

The main mechanism of pre-mRNA 3’-end processing is cleavage and polyadenylation (CPA), which involves endonucleolytic cleavage of newly synthesized transcripts and addition of adenosine residues constituting the poly(A) tail to the generated 3’-end. CPA is crucial for mRNA stability, transport to the cytoplasm and translation (Millevoi & Vagner, 2010; Shi & Manley, 2015). This nuclear process involves the recognition of *cis*-acting elements in the pre-mRNA by a complex machinery comprising more than 80 proteins (Shi *et al*, 2009). The pre-mRNA sequences serving as the polyadenylation signal (PAS) include an hexameric sequence (most often AAUAAA) located 10–30 nucleotides (nt) upstream of the cleavage site (generally a CA dinucleotide) and a downstream sequence element (DSE) (U/GU-rich) located within 30 nt downstream of the cleavage site. Additional sequence elements located either upstream (upstream sequence element; USE) or downstream (auxiliary downstream sequence element; AuxDSE) of the cleavage site modulate the recognition of the PAS.

The cleavage reaction at the PAS (called thereafter PAS cleavage), which precedes the addition of the poly(A) tail, generally occurs in a co-transcriptional manner. PAS recognition is indeed tightly coupled to RNA polymerase II (Pol II) transcription termination (Proudfoot, 2016). Rpb1, the largest subunit of Pol II, contains a Carboxy-Terminal Domain (CTD) that is comprised of heptad repeats (consensus Tyr^1^-Ser^2^-Pro^3^-Thr^4^-Ser^5^-Pro^6^-Ser^7^) and plays a critical role in coupling pre-mRNA 3’-end processing and transcription termination, especially through its phosphorylated Ser^2^ (phospho-Ser^2^) residues (Ahn *et al*, 2004). Several components of the polyadenylation machinery, including PCF11, which is concentrated at the 3’-end of genes, preferentially bind the phospho-Ser^2^ CTD (Barilla *et al*, 2001; Licatalosi *et al*, 2002; Meinhart & Cramer, 2004). In human cells, PCF11 depletion leads to a transcription termination defect through a decrease in the degradation of the downstream RNA, generated after the PAS cleavage (West *et al*, 2008). In the Pause-Type model of transcription termination, Pol II pauses at a GC-rich region located a few nucleotides downstream of the PAS, stimulating the PAS cleavage of the pre-mRNA in a co-transcriptional manner *i.e*. when the pol II-bound pre-mRNA is on the chromatin (Nojima *et al*, 2013; Cortazar *et al*, 2019; Gromak *et al*, 2006).

Another model of transcription termination has been proposed (Nojima *et al*, 2013; West *et al*, 2008; Dye & Proudfoot, 2001). In this Co-Transcriptional Cleavage (CoTC)-type model, the pre-mRNA is released from chromatin to nucleoplasm through a cleavage event at a CoTC site located downstream of the PAS, and the PAS cleavage subsequently occurs in the nucleoplasm. This mechanism has been described in several human genes (Nojima et al., 2013). The CoTC-type termination model has also been observed in *Drosophilia* where a release of pre-mRNA from transcription sites to the nucleoplasm takes place prior to PAS cleavage (Sikes *et al*, 2002). The CoTC cleavage occurs a few kilobases downstream of the PAS, generally at an AT-rich sequence called CoTC element (White *et al*, 2013). Mutations in this element induce an inhibition of pre-mRNA 3’-end processing *in vitro* (Teixeira *et al*, 2004).

Here, we report that upon UV-irradiation, PAS cleavage of the *p53* pre-mRNA is independent from the cleavage/termination factor PCF11 and CTD Ser^2^ phosphorylation, and relies on a downstream CoTC site, thereby allowing 3’-end processing of the *p53* pre-mRNA to escape repression by DNA damage. We also identified several other pre-mRNAs that exhibit a CoTC-type mechanism of 3’-end processing in response to UV-induced DNA damage and that escape repression by DNA damage, like the *p53* pre-mRNA.

## Results

### PCF11 is dispensable for *p53* pre-mRNA 3’-end processing in UV-treated cells

To understand the contribution of the pre-mRNA 3’-end processing machinery in the response to UV treatment, we analyzed the abundance of 13 proteins constituting the different subcomplexes involved in 3’-end processing by Western blot (Figure 1A). To ascertain that the band observed in each Western blot corresponds to the expected protein, we used published siRNAs targeting each of the corresponding mRNAs (Masamha *et al*, 2014). The experiments were performed in A549 lung tumor cells irradiated with UV (254nm; 40J/m^2^) and harvested after 16 hours of recovery, conditions that we previously used to demonstrate the maintenance of *p53* pre-mRNA 3’-end processing following UV treatment (Decorsière *et al*, 2011; Newman *et al*, 2017). Consistent with previously reported data (Kleiman & Manley, 2001), we observed no changes in the levels of both CstF64 and CPSF160 in response to UV. The abundance of the other components of the CPSF, CstF and CFIm complexes was unchanged (Figure 1A). By contrast, we detected a significant decrease in the abundance of both PCF11 and CLP1 in UV-treated cells (Figure 1A). The UV-dependent reduction in PCF11 expression was confirmed in another set of experiments using 2 different siRNAs targeting PCF11 (Figure 1B) and was accompanied by a 5-fold decrease in *PCF11* mRNA level (Figure 1C). These observations suggest that PCF11 might be dispensable for *p53* pre-mRNA 3’-end processing following UV-induced DNA damage.

**Figure 1.**
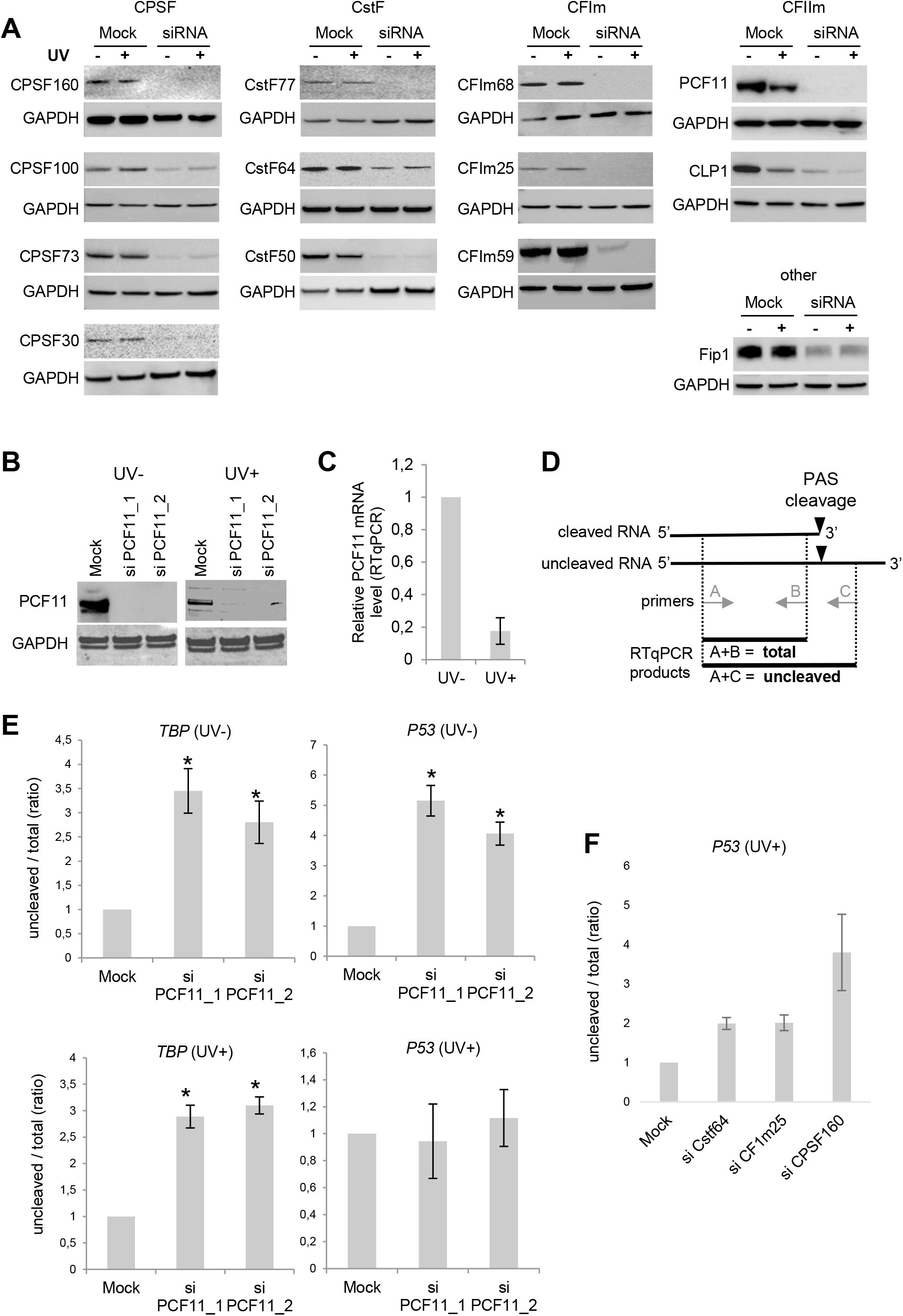
PCF11 is dispensable for *p53* pre-mRNA 3’-end processing in UV-treated cells. (A) Western blot of pre-mRNA 3’-end processing factors in response to UV treatment (40 J/m^2^) of A549 cells, followed by16 hours of recovery. GAPDH was used as a loading control. (B) Western blot analysis of PCF11 expression in A549 cells transfected for 48 hours with two different siRNA targeting PCF11 prior to exposure to UV. (C) RT-qPCR measuring relative PCF11 mRNA level in A549 cells in response to UV treatment (40 J/m^2^). The expression was normalized to RP18S. \**P* <0.05; unpaired t-test. (D) Scheme representing the RT-qPCR strategy for assessing pre-mRNA 3’-end processing efficiency. Primers for uncleaved pre-mRNAs are located downstream of the polyadenylation site, while primers that detect both processed (cleaved) and unprocessed RNAs (uncleaved) amplify upstream of the polyadenylation site. The ratio of uncleaved/total (cleaved+uncleaved) indicates the processing efficiency, where a greater uncleaved/total ratio represents a reduced processing. (E) RT-qPCR assay on nuclear RNA for assessing the uncleaved/total ratio of *p53* and TATA-binding protein (*TBP*) pre-mRNAs in A549 cells transfected for 48 hours with two different siRNA targeting PCF11 prior to exposure (UV+) or not (UV-) to UV irradiation (40 J/m^2^). (F) RT-qPCR assay on nuclear RNA for assessing the uncleaved/total ratio of *p53* pre-mRNA in A549 cells transfected for 48 hours with siRNAs targeting the Cstf64, CF1m25 and CPSF160 and exposed to UV irradiation (40 J/m^2^).

To confirm that PCF11 is not required for *p53* pre-mRNA 3’-end processing following UV, we evaluated the effect of siRNA-mediated depletion of PCF11 on the efficiency of PAS cleavage of the *p53* pre-mRNA by real-time quantitative PCR analysis (RT-qPCR) (Figure 1D). The *TBP* pre-mRNA was used as a control since it was previously shown to be inhibited at the level of PAS cleavage efficiency due to UV treatment (Decorsière *et al*, 2011; Newman *et al*, 2017). According to a previously described approach (Decorsière *et al*, 2011; Newman *et al*, 2017), we measured the ratio of uncleaved RNA to total RNA (that is the sum of cleaved and uncleaved RNA) in the nuclear pool of RNAs, by qPCR with antisense primers located either downstream or upstream of the PAS cleavage site, respectively (Figure 1D). In untreated cells, PAS cleavage of both the *TBP* and *p53* pre-mRNAs was inhibited by PCF11 depletion, as revealed by the increased ratio of uncleaved/total RNAs in PCF11-depleted cells (Figure 1E; top panels). This is consistent with the fact that this factor is essential for the co-transcriptional, Pol II-coupled PAS cleavage reaction (West *et al*, 2008). Following UV treatment, while *TBP* PAS cleavage was still inhibited by PCF11 depletion, *p53* PAS cleavage was no longer inhibited (Figure 1E; bottom panels). This effect was specific to PCF11 since siRNA-mediated depletion of CstF64, CFIm25 and CPSF160 all led to decreased *p53* PAS cleavage in UV-treated cells (Figure 1F). Altogether, these data indicate that PCF11, which exhibits reduced RNA and protein levels in UV-treated cells, is dispensable for *p53* (but not *TBP*) pre-mRNA 3’-end processing in UV-treated cells.

Previous reports showed that UV-induced DNA damage induces global changes in Pol II phosphorylation, including Ser^2^ phosphorylation (Muñoz et al., 2009; Rockx et al., 2000). Considering the link between PCF11 and the Pol II CTD phospho-Ser^2^ (PolII Ser2P), we sought to determine whether inhibition of Ser^2^ phosphorylation may mimic the effect of depleting PCF11 on *p53* 3’-end processing following UV. We treated cells with Ser^2^ kinase (CDK9) inhibitors (5,6-Dichlorobenzimidazole 1-β-D-ribofuranoside (DRB) or flavopiridol) and then assessed the efficiency of pre-mRNA 3’-end processing. Both DRB and flavopiridol reduced PolII Ser^2^ phosphorylation (Supplementary Figure 1A). DRB, as expected, inhibited both *TBP* and *p53* PAS cleavage in untreated cells (Supplementary Figure 1B). In UV-treated cells DRB inhibited *TBP*, but not *p53* PAS cleavage (Supplementary Figure 1C), and similar results were obtained with flavopiridol (Supplementary Figure 1D).

### PAS cleavage of the *p53* pre-mRNA occurs in the nucleoplasm following a CoTC event

The experiments above show that, in UV-treated cells, PAS cleavage of the *p53* pre-mRNA does not require PCF11 and Pol II CTD phospho-Ser^2^. This suggests that it might occur in a transcription termination uncoupled-manner, as described in the CoTC-type model, where PAS cleavage occurs post-transcriptionally, following a co-transcriptional cleavage at a downstream CoTC site (Nojima *et al*, 2013). In this case, a pre-mRNA that has not undergone PAS cleavage (PAS-uncleaved pre-mRNA) can be detected in the nucleoplasm, where it is released upon CoTC cleavage. We therefore analyzed the nuclear distribution of the PAS-uncleaved *p53* pre-mRNA by RT-qPCR in chromatin and nucleoplasm fractions. The quality of the fractionation was assessed by Western blot against histone H3 as a chromatin marker and toposiomerase IIα as a nucleoplasm marker (Figure 2A). The *GAPDH* and *WDR13* pre-mRNAs were used as controls as they were previously reported to be PAS-cleaved co-transcriptionally or post-transcriptionally (following a CoTC event) respectively (Nojima *et al*, 2013). Accordingly, the relative abundance of PAS-uncleaved pre-mRNA in the nucleoplasm, as compared to the chromatin, was much higher for *WDR13* than for *GAPDH* (Figure 2B). The nucleoplasm/chromatin ratio of PAS-uncleaved pre-mRNA of *p53* was similar to the one of *WDR13*, suggesting that the *p53* pre-mRNA may be released in the nucleoplasm following a CoTC event, while the *TBP* pre-mRNA behaved similarly to the *GAPDH* pre-mRNA (Figure 2B).

**Figure 2.**
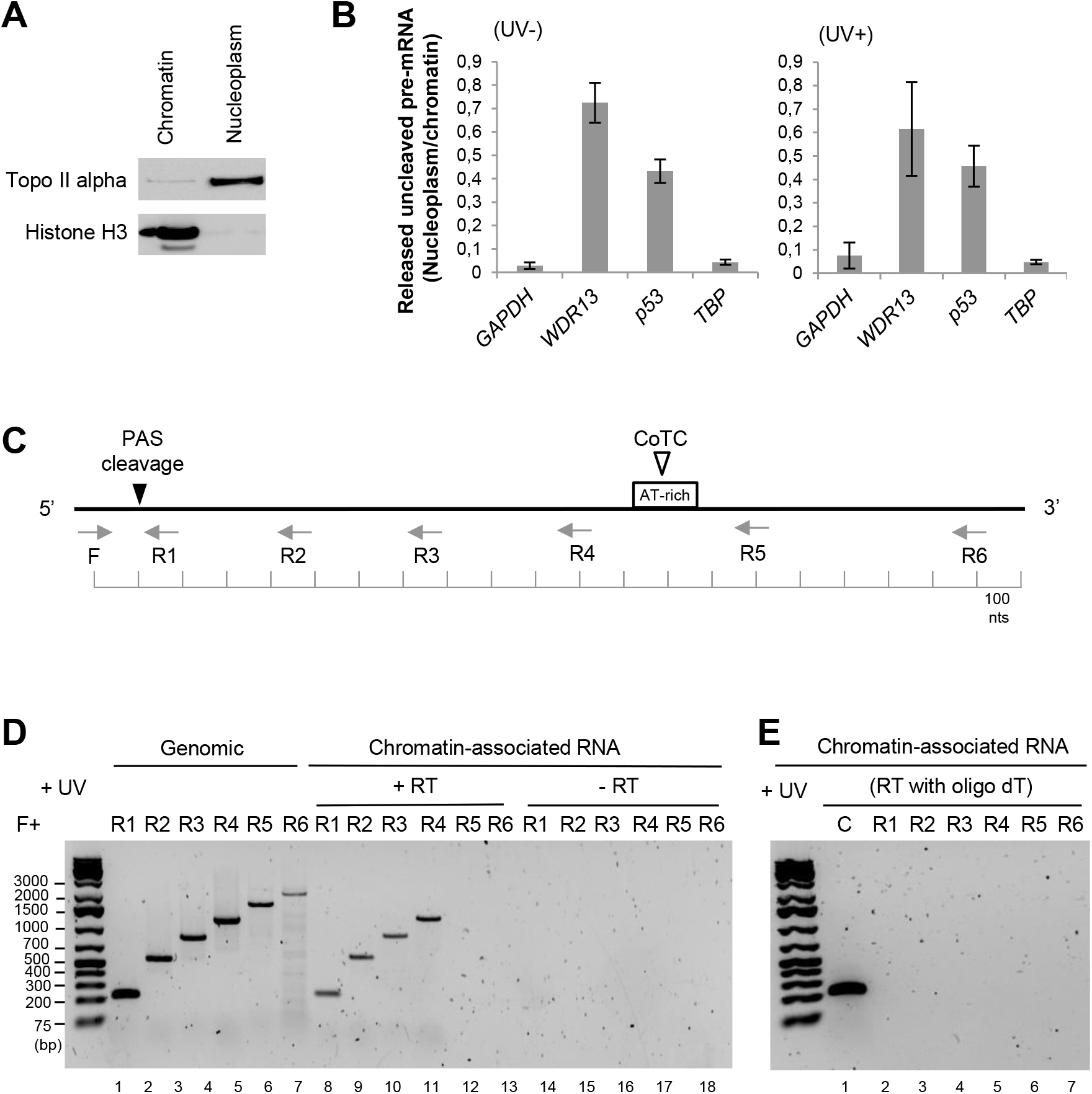
p53 pre-mRNA 3’-end cleavage occurs in the nucleoplasm following a CoTC. (A) Western blot analysis to verify the quality of the nuclear fractionation of UV-treated A549 cells (40 J/m^2^). The topoisomerase II alpha was used as a marker of the nucleoplasm compartment and the histone H3 for the chromatin fraction. (B) RT-qPCR analysis on RNA extracted from nucleoplasm and chromatin fractions. The ratio of uncleaved pre-mRNA (nucleoplasm/chromatin) was calculated to quantify the level of unprocessed *p53*, *TBP*, *WDR13* and *GAPDH* pre-mRNA released in the nucleoplasm compared to the chromatin-bound unprocessed pre-mRNA. *WDR13* and *GAPDH* were included as controls as they have been previously reported to be processed post- and co-transcriptionally, respectively. (C) Scheme representing the strategy to map the location of the CoTC site in the *p53* pre-mRNA. Forward primer (F) is located upstream of the PAS (poly(A) site) and reverse primers (R1-6) are located downstream at increasing distances from the PAS. (D) RT-PCR analysis of p53 3’ flanking region using primer pairs indicated (black arrows) in the scheme above the data panel. Lanes 1-6 correspond to amplified genomic DNA used as a PCR amplification control. Lanes 7-12 correspond to cDNA derived from chromatin-associated RNA reverse transcribed (+RT) using random primer. Lanes 13-18 are negative RT control samples (-RT). (E) RT-PCR analysis of *p53* 3’ flanking region using the same primers employed in the data panel above. Lane 1 is a control PCR amplification of cDNA derived from the reverse transcription of a control mRNA using oligo (dT). Lanes 2-7 are PCR amplification of reverse transcribed p53 chromatin-associated pre-mRNA using oligo oligo (dT).

An AT-rich sequence that could correspond to a potential CoTC sequence element is found around 1,200 nt downstream of the *p53* PAS (Figure 2C). To map the putative CoTC element, chromatin-associated RNA was reverse transcribed using random primers and the obtained cDNA was amplified by PCR using primers complementary to the 3’ flanking region of the *p53* gene (Figure 2C). PCR amplification was carried out using a single forward primer (F), located upstream of the *p53* PAS, in combination with reverse primers (R1-R6) located at increasing distance downstream of the *p53* PAS. The F/(R1-R6) primer pairs were used to amplify genomic DNA as an amplification control (Figure 2D). In cDNA derived from chromatin-bound RNA, the F-R1, F-R2, F-R3 and F-R4 primer pairs resulted in the detection of PCR products at the expected size of 229, 558, 852 and 1000 bp (Figure 2D; lanes 7-10). Of note, these PCR products precisely correspond to bands obtained with genomic DNA (lanes 1-4). In contrast, the F-R5 and F-R6 primer pairs did not yield detectable PCR products with cDNA samples from chromatin-bound RNA (lanes 11-12) even though PCR products were obtained with the genomic DNA control (lanes 5-6). These observations indicate that the *p53* pre-mRNA is cleaved in between approximately 1000 to 1400 nt downstream of the PAS, in the region where the AT-rich sequence is located. To ascertain that this cleavage event is not linked to the presence of an alternative PAS, we adopted the same mapping strategy using chromatin-bound pre-mRNA but reverse transcription was performed with an oligo-dT primer. A cDNA derived from an mRNA transcript was included as a control. No bands were detected with all primer pairs used previously, except for the control (Figure 2E). Altogether, these data indicate that the *p53* pre-mRNAs cleaved in the vicinity of the AT-rich region do not contain a poly(A) tail and are generated through a CoTC-type event in an UV-induced manner.

Consistently, using single-molecule fluorescence *in situ* hybridization (smFISH), we found that *p53* pre-mRNA regions downstream of the PAS (probe B) were detected following UV exposure (median number of smFISH spots: no UV = 11; with UV = 11) (Figure 3). This is not true for *GAPDH* (probe D), as expected for a pre-mRNA that undergoes efficient co-transcriptional PAS cleavage and no CoTC-type cleavage. Without UV exposure, *GAPDH* downstream regions targeted by probe D were visible and localized to the transcription sites (Figure 3). However, following UV exposure, the downstream regions targeted by probe D were no longer visible in the nucleus. In contrast, RNA regions upstream of the PAS were detected for both *p53* (probe A) (median number of smFISH spots: no UV = 18; with UV = 26) and *GAPDH* (probe C) (Figure 3), as expected for mature mRNAs. Thus, our smFISH data are consistent with our RT-qPCR data on chromatin (Figure 2) indicating that PAS cleavage of *p53* pre-mRNA occurs post-transcriptionally.

**Figure 3.**
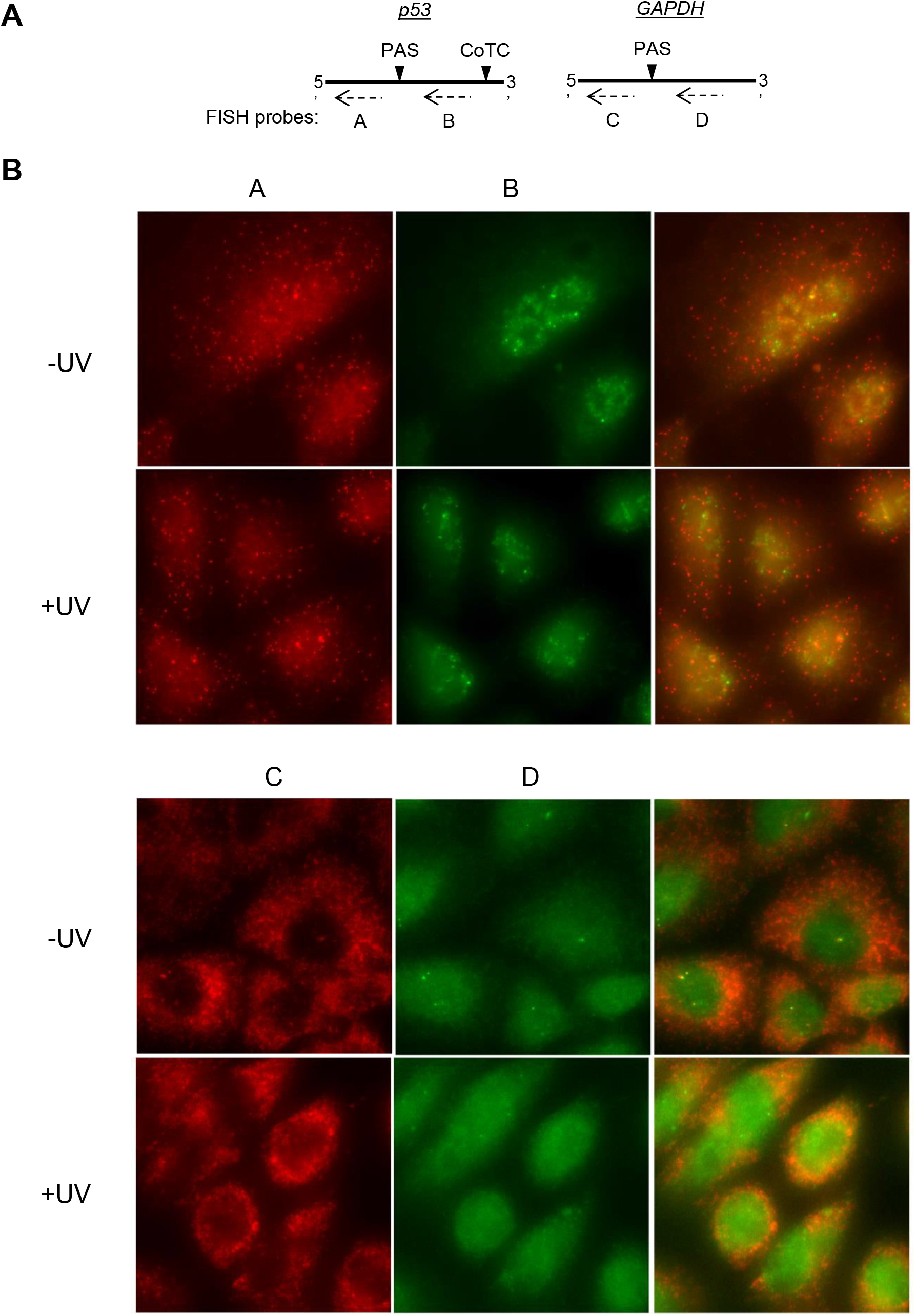
PAS cleavage of *p53* pre-mRNA occurs post-transcriptionally. (A) Probe design for single-molecule Fluorescence *in situ* hybridisation (smFISH) spanning regions upstream ‘A’ and downstream ‘B’ of *p53* PAS as well as those spanning regions upstream ‘C, and downstream ‘D’ of *GAPDH* PAS. (B) Representative images of smFISH in untreated (-UV) or UV-treated (+UV) A549 cells with the indicated probes (A, B, C and D).

We have previously shown that the two RBPs hnRNP H/F (Decorsière *et al*, 2011) and the RNA helicase DHX36 (Newman *et al*, 2017) are involved in the regulation of *p53* pre-mRNA 3’-end processing following UV-induced DNA damage. Consistent with *p53* pre-mRNA 3’-end processing mostly occurring in the nucleoplasm, the increased uncleaved/total ratio of *p53* pre-mRNA following depletion of DHX36 or hnRNP H/F was significantly higher in the nucleoplasm than in the chromatin (Supplementary Figure 2A-B).

### The CoTC site is implicated in the maintenance of *p53* pre-mRNA 3’-end processing in response to UV-induced DNA damage

In order to determine the importance of the CoTC site in *p53* pre-mRNA 3’-end processing following UV, the *p53* CoTC element was deleted using a CRISPR-based strategy in both A549 and A375 cells (Supplementary Figure 3A-B). We obtained an A549 clone with deletion of the CoTC site in all three *TP53* alleles existing in these cells (hereafter called ΔCoTC) and several A549 and A375 clones with deletion of only a subset of alleles (herafter called pΔCoTC) (Supplementary Figure 3). ΔCoTC, pΔCoTC and WT cells were then tested for *p53* pre-mRNA 3’-end processing efficiency in response to UV.

We observed an increase in the PAS-uncleaved to total ratio for *p53* in UV-treated ΔCoTC, but not WT cells (Figure 4A). Similar results were obtained with pΔCoTC A549 (Supplementary Figure 4A) and pΔCoTC A375 (Supplementary Figure 4B) cells. As a control, deletion of the *p53* CoTC region had no effect on the UV-dependent regulation of pre-mRNA 3’-end processing for *WDR13*, *GAPDH*, and *TBP* (Figure 4A and Supplementary Figure 4). Thus, the *p53* CoTC region is required for the maintenance of *p53* pre-mRNA 3’-end processing upon UV exposure. We also found a decrease in the nucleoplasm/chromatin ratio of the *p53* PAS-uncleaved pre-mRNA in ΔCoTC cells when compared to WT cells (Figure 4B). This effect was also observed in pΔCoTC A549 (Supplementary Figure 5A) and pΔCoTC A375 (Supplementary Figure 5B) cells and was not observed for the *WDR13*, *GAPDH* and *TBP* genes (Figure 4B and Supplementary Figure 5). This shows that the *p53* CoTC site is required for the release of the PAS-uncleaved *p53* pre-mRNA from chromatin to nucleoplasm in response to UV. Consistently, total *p53* mRNA levels were decreased in ΔCoTC and pΔCoTC cells, but not in WT cells, in response to UV (Figures 4C and Supplementary Figure 6A). In addition, the UV-dependent increase in p53 protein levels in WT cells was not observed in ΔCoTC and pΔCoTC cells (Figures 4D and Supplementary Figure 6B). Altogether, these data show that the CoTC site of *p53* is required to maintain *p53* PAS cleavage and promote *p53* expression following UV irradiation.

**Figure 4.**
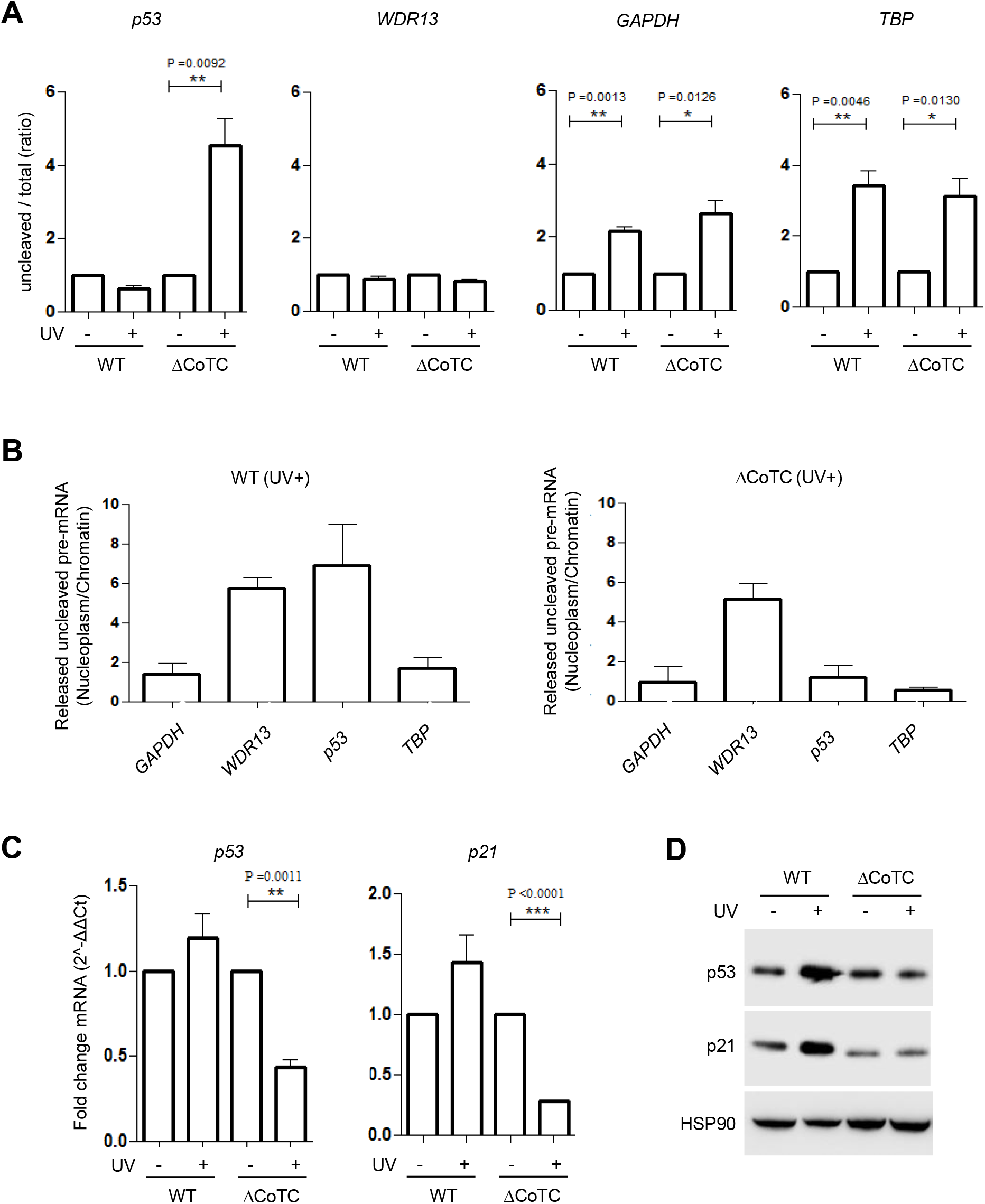
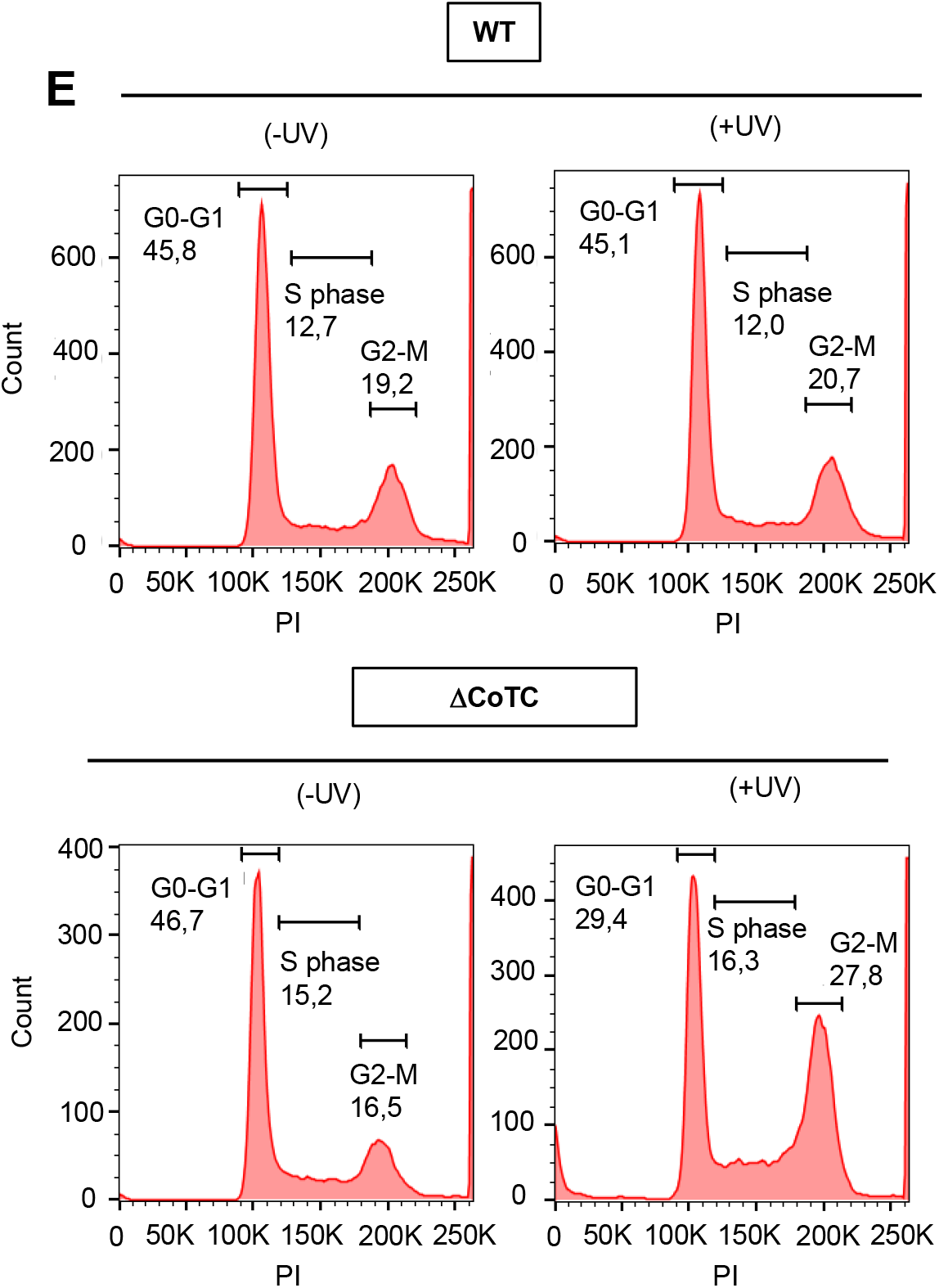
The CoTC is implicated in the maintenance of 3’-end processing of *p53* pre-mRNA in response to UV-induced DNA damage. (A) RT-qPCR assay on nuclear RNA for assessing the uncleaved/total ratio of *p53* pre-mRNA in wild type (WT) and CoTC deleted (ΔCoTC) A549 cells treated with or without UV irradiation (40 J/m^2^). *P* <0.05; unpaired t-test. (B) RT-qPCR analysis on RNA extracted from nucleoplasm and chromatin fractions. The ratio of uncleaved pre-mRNA (nucleoplasm/chromatin) was calculated to quantify the level of unprocessed *p53*, *WDR13*, *GAPDH* and *TBP* pre-mRNA released in the nucleoplasm compared to the chromatin-bound unprocessed pre-mRNA in wild type (WT) and CoTC deleted (ΔCoTC) A549 cells treated with or without UV irradiation (40 J/m^2^). *P* <0.05; unpaired t-test. (C) RT-qPCR measuring relative *p53* and *p21* mRNA levels in wild type (WT) and CoTC-deleted (ΔCoTC) cells in response to UV treatment (40 J/m^2^). The expression was normalized to HPRT. \**P* <0.05; unpaired t-test. (D) Western blot analysis of p53 and p21 expression in wild type (WT) and CoTC deleted (ΔCoTC) A549 cells treated with or without UV irradiation (40 J/m^2^). (E) Representative flow-cytometry analyses of the cell cycle (DNA content by Propidium Iodide; PI) in wild type (WT) and CoTC deleted (ΔCoTC) A549 cells treated with or without UV irradiation (40 J/m^2^). Indicated: percent of cells in the G0-G1, S and G2/M phases.

We then assessed potential consequences of CoTC site deletion on downstream functions of p53. A direct transcriptional target of the p53 protein is the *CDKN1A/p21* gene, which encodes an inhibitor of cell cycle progression from G1 to S phase (Galanos *et al*, 2016; Jeong *et al*, 2010; Matsuda *et al*, 2017). UV-induced up-regulation of *p21* mRNA and p21 protein levels was observed in WT cells but not in ΔCoTC and pΔCoTC cells (Figure 4C-D and Supplementary Figure 6A-B). Analysis of cell cycle distribution by FACS showed no effect of CoTC site deletion in the absence of UV (Figure 4E, left panels). However, a moderate UV treatment, which had no effect on cell cycle distribution in WT cells, led to a decrease in G0/G1 cells in ΔCoTC and pΔCoTC cells (Figures 4E and Supplementary Figure 7). Altogether, these data suggest that deletion of the *p53* CoTC site leads to impaired p21 induction and enhanced G1-S phase progression following a moderate UV irradiation.

### The 3’-end processing of several pre-mRNAs undergoing a CoTC cleavage event is maintained in response to UV-induced DNA damage

Our finding that the CoTC site of *p53* pre-mRNA is required for *p53* to escape 3’-end processing inhibition by UV prompted us to investigate whether CoTC-dependent cleavage may be linked to UV-resistant pre-mRNA 3’-end processing in other genes. Toward this aim, we first developed a strategy to analyze in a high-throughput manner the efficiency of pre-mRNA 3’-end cleavage by RNA sequencing (RNA-Seq). This strategy is based on the evaluation of the number of reads located in 500 nt-long windows either upstream (total RNA) or downstream of the PAS (PAS-uncleaved RNA; Figure 5A). An increase in the ratio of downstream reads to upstream reads indicates an inhibition of 3’-end cleavage, leading to read-through transcription (Vilborg *et al*, 2015). Focusing on 4208 expressed genes (Supplementary Table 1) with detectable reads downstream of the PAS and using a cut-off of p<0.05, this analysis identified 378 pre-mRNAs with UV-repressed 3’-end processing and 108 pre-mRNAs with a more efficient 3’end processing in UV-treated compared to untreated cells (UV-resistant 3’-end processing; Figure 5B). Example of the read distribution in the 500 nt-long windows located upstream and downstream of the PAS are illustrated in Figure 5C for the *ZRANB2* and the *HMGB1* genes. In the case of *ZRANB2* that belongs to the UV-repressed 3’-end processing group, the absence of reads downstream of the PAS in untreated cells indicates a very efficient 3’-end processing activity, while the presence of more reads in this window in UV-treated cells indicates a reduced 3’-end processing efficiency following UV treatment. In the case of *HMGB1* that belongs to the UV-resistant 3’-end processing group, we observed an opposite trend showing that there is an increase in pre-mRNA 3’-end processing following UV treatment (Figure 5C).

**Figure 5.**
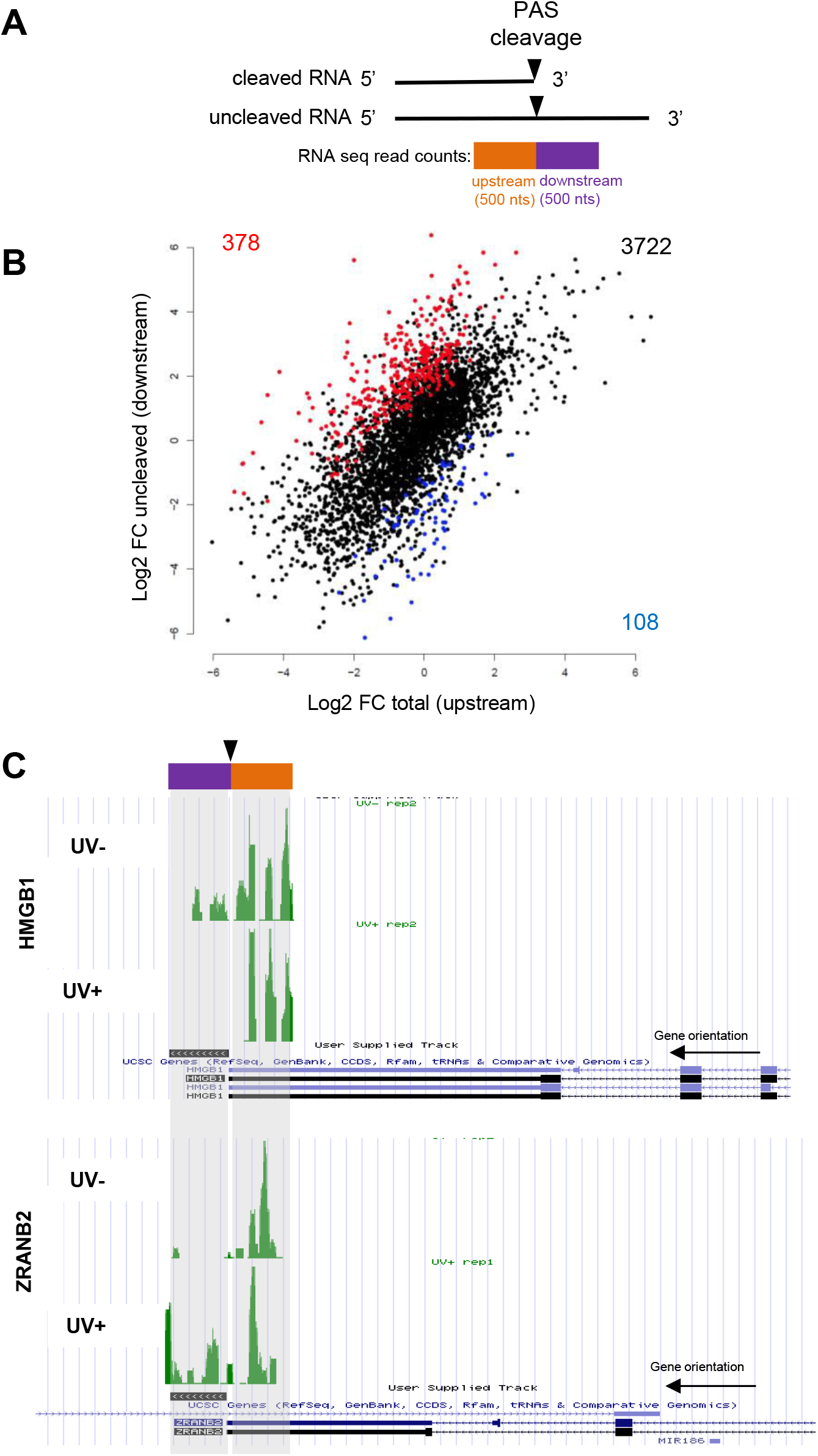
UV induces widespread regulation of pre-mRNA 3’-end processing. (A) Scheme representing the adopted strategy to study at genome wide level by RNA-sequencing the regulation of pre-mRNA 3’end processing in response to UV. Nuclear RNA from A549 cells UV-irradiated or unirradiated was extracted and cDNA library preparation was performed to assess RNA sequencing. The efficiency of 3’-end processing was studied by the quantification of the number of reads located in a window of 500 bp downstream (uncleaved RNA) and upstream (total RNA) of the poly (A) site. The ratio of reads downstream / reads upstream reflects the efficiency of pre-mRNA 3’end processing. (B) RNA-seq plot representing, for each gene, the log2 fold change (comparing UV-irradiated to non-irradiated cells) of the number of reads downstream of the poly(A) site (uncleaved RNA, *y* axis), and the log2 FC of the number of reads upstream of the poly(A) site (total RNA, *x* axis). Regulation events are considered significant if p-value is below 0.05. The 108 genes at bottom right have a decreased uncleaved/total RNA ratio, meaning that PAS cleavage is increased upon UV irradiation. In contrast, the 378 genes at top left have an increased uncleaved/total RNA ratio, meaning that PAS cleavage is decreased upon UV irradiation. (C) Visualization of reads distribution in a window of 500 nts upstream and downstream of the PAS of *HMGB1* and *ZRANB2* pre-mRNA.

We randomly chose 9 candidate genes from each group and validated the RNA-Seq data by RT-qPCR on PAS-uncleaved and total RNA using primers located downstream or upstream of the PAS, respectively (Figure 6A). These RT-qPCR analyses showed that 7 out of 9 candidate pre-mRNAs of the UV-repressed group indeed exhibited 3’-end processing inhibition by UV (88% validation rate; Figure 6A, red bars). In contrast, 9 out of 9 candidate pre-mRNAs of the UV-resistant group were indeed resistant to 3’-end processing inhibition by UV (100% validation rate; Figure 6A, blue bars). These include several DDR-related genes, namely XRCC5 and HMGB1. Because our RNA-seq approach is limited by sensitivity of read detection downstream of the PAS, we also analyzed 6 genes from the p53 pathway by RT-qPCR. One of them (*i.e*., *NOXA*) exhibited 3’-end processing inhibition by UV, while four genes escaped repression (*i.e*., *MDM2, PUMA, PTEN* and *FAS;* Figure 6A, right). Altogether, our RT-qPCR analyses identified 17 pre-mRNAs that were resistant to 3’-end processing inhibition by UV, and 11 pre-mRNAs that underwent inhibition.

**Figure 6.**
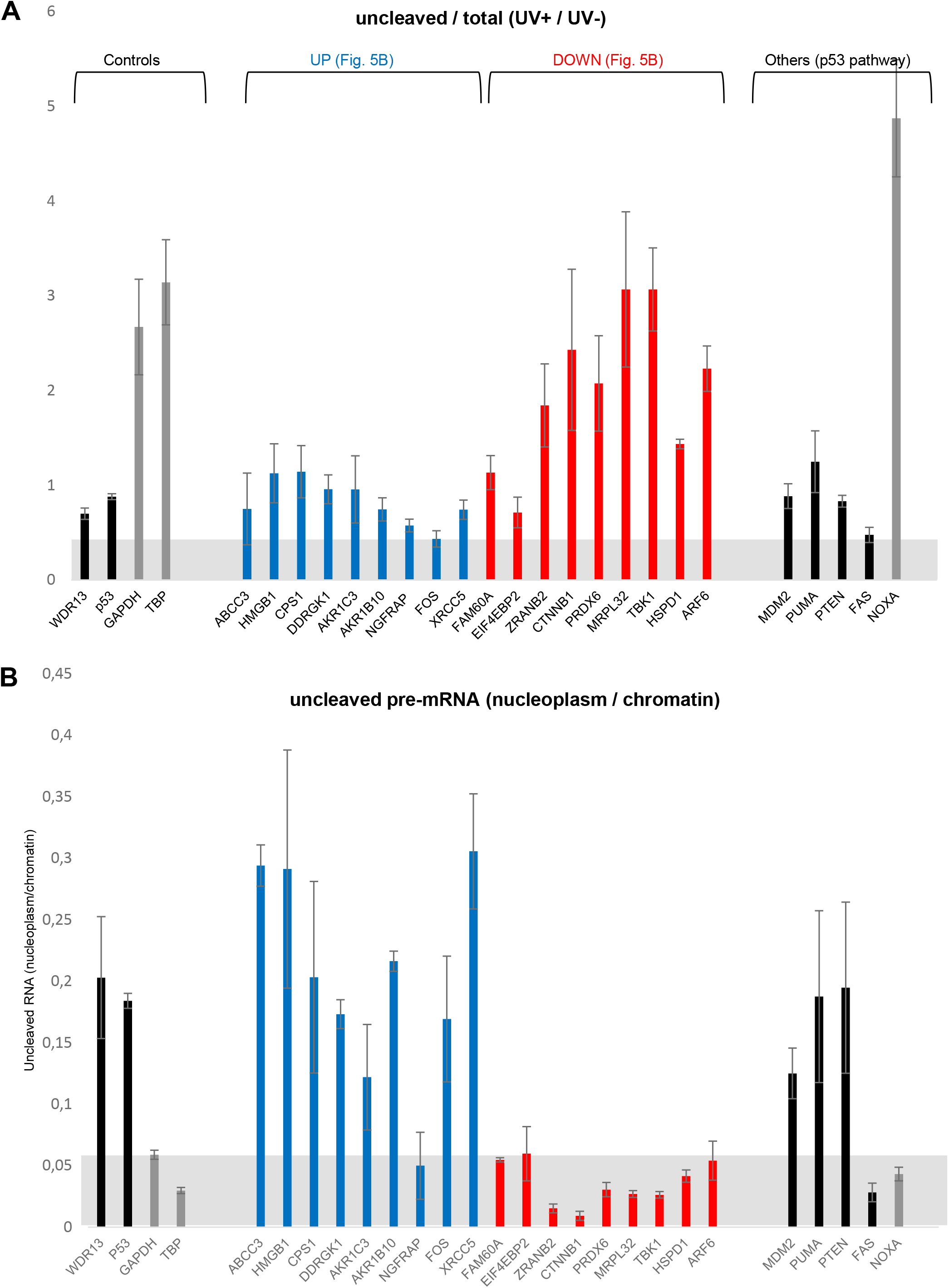
The 3’-end processing of diverse pre-mRNAs undergoing a CoTC cleavage event is maintained in response to UV-induced DNA damage. (A) RT-qPCR (uncleaved/total RNA) on nuclear RNA extracted from UV-treated or untreated A549 cells, to assess the regulation of 3’end processing of 20 pre-mRNA randomly selected from the previous RNA-sequencing data. (B) RT-qPCR (Released uncleaved pre-mRNA nucleoplasm/chromatin) on RNA extracted from the nucleoplasm and the chromatin fractions of A549 UV-treated cells.

Then, we measured by RT-qPCR the nucleoplasm/chromatin ratio of PAS-uncleaved pre-mRNA. Remarkably, 13 out of 17 (76%) UV-resistant pre-mRNAs had a high nucleoplasm/chromatin ratio, indicating that PAS cleavage occurs at least in part post-transcriptionally (Figure 6B). This proportion was even higher (8 out of 9 [89%]) when focusing on UV-resistant pre-mRNA candidates that we identified by RNA-seq (Figure 6B, blue bars). In contrast, none of the 11 UV-repressed pre-mRNAs (and none of the 9 candidates identified by RNA-seq) were abundant in the nucleoplasm (Figure 6B). These data show that resistance of PAS cleavage to UV correlates with its occurrence in the nucleoplasm, thus extending our findings on *p53*.

Because we found that the UV resistance and nucleoplasmic occurrence of the *p53* pre-mRNA PAS cleavage require a downstream CoTC cleavage site (Figure 3), we then used our PCR-based mapping strategy (Figure 2) to locate putative CoTC elements in 8 other UV-resistant pre-mRNAs (Supplementary Figure 8A-B). For the 8 tested genes, the presence of CoTC cleavage sites was validated by the sharp loss of amplification at a given distance (most often about 2.5 kb) downstream of the PAS (Supplementary Figure 8C). As for *p53* (Figure 2E), the CoTC-dependent cleavage event for the 8 tested genes is not linked to the presence of an alternative PAS (Supplementary Figure 8D). Gene Ontology (GO) analysis of pre-mRNAs with a more efficient 3’end processing in UV-treated compared to untreated cells shows an enrichment of genes involved in the inhibition of apoptosis and metabolism of a genotoxic stress inducing agent, doxorubicin (Supplementary Figure 9). Altogether, these results provide evidence for an association between maintained 3’-end processing following UV irradiation and the presence of a CoTC element in several DDR-related genes, including *XRCC5*, *PUMA*, *FAS*, *MDM2* and *TP53*, for the latter of which we showed that the CoTC site is required for 3’-end processing maintenance.

## Discussion

Pre-mRNA 3’-end processing by PAS cleavage and poly(A) tail addition mostly occurs in a co-transcriptional manner. We show here that PAS cleavage of the *p53* pre-mRNA occurs at least in part in a manner that is uncoupled from transcriptional termination. It involves a CoTC sequence that lies about 1.2 kb downstream of the PAS and allows a first 3’-end cleavage event, leading to dissociation of the pre-mRNA from chromatin. This is followed by a second 3’-end cleavage event (and polyadenylation) occurring at the PAS of the released RNA in the nucleoplasm.

One of the important factor in the coupling between 3’-end processing and transcription termination is PCF11, a 3’-end processing factor that mediates transcriptional termination in yeast (Grzechnik *et al*, 2015; Larochelle *et al*, 2018) as well as in vertebrates (Kamieniarz-Gdula *et al*, 2019). PCF11 recruits the yeast Rat1 or human Xrn2 exonucleases to exert a 5’-3’ exonucleolytic degradation on the nascent RNA leading Pol II to terminate transcription (Eaton *et al*, 2018; Luo, 2006; West & Proudfoot, 2007). We show that pre-mRNA 3’-end processing is inhibited in the absence of PCF11 (Figure 1). PCF11 interacts with CLP1 to target the cleavage site and modulates the binding and clevage efficiency of CFII. (Zhang *et al*, 2021) It also regulates polyadenylation site choice and plays a role in controlling the 3’ Untranslated Region (3’UTR) of transcripts. (Nourse *et al*, 2020; Ogorodnikov *et al*, 2018; Wang *et al*, 2019). However, *p53* pre-mRNA 3’-end processing is independent of PCF11 in UV-treated cells, allowing this pre-mRNA to escape from the decrease in PCF11 levels observed in UV-treated cells (Figure 1). Since PCF11 has a CTD Interacting Domain (CID) and binds preferentially the phosphorylated Ser2 of an elongating RNAP II (Meinhart & Cramer, 2004), our results clearly indicate an uncoupling of 3’ end processing and transcriptional termination in the p53 pre-mRNA following UV-induced DNA damage.

We demonstrate that *p53* pre-mRNA 3’-end processing does not require PCF11 because it is processed at a CoTC sequence (Figures 2–3).. CRISPR-based deletion of the p53 CoTC leads to inhibition of *p53* pre-mRNA 3’-end processing, decreased p53 and p21 protein levels and decreased G0/G1 cells in UV-treated cells (Figure 4). This is consistent with the fact that UV radiation induced cell cycle arrest is correlated to increase in p53 levels (Latonen *et al*, 2001) and causes retention of cells at the G2-M phase (Blackford & Jackson, 2017; Céraline *et al*, 1998; van Oosten *et al*, 2000; Pavey *et al*, 2001). CoTC-dependent cleavage therefore acts as a mechanism of escape from UV-induced global inhibition of pre-mRNA 3’-end processing. Beyond *p53*, the presence of CoTC elements in the 3’ flanking regions of a number of genes, including genes implicated in p53-mediated DNA damage response, was validated (Figures 5–6).

We thus propose a model where UV-induced DNA damage inhibits co-transcriptional PAS cleavage, that is chromatin-bound and PCF11-dependent, but not post-transcriptional PAS cleavage that is nucleoplasmic and PCF11-independent and occurs following a CoTC-dependent release of the pre-mRNA from chromatin. Our model thus explains the rescue of the 3’ end processing of specific mRNAs in a transcription-uncoupled manner, despite the global inhibition by UV induced DNA damage of the canonical chromatin-associated pre-mRNA processing that is tightly coordinated to transcriptional termination (Hamperl *et al*, 2017; Luna *et al*, 2005; Nilsson *et al*, 2018; Reimer *et al*, 2021; Teloni *et al*, 2019). Nucleoplasmic PAS-dependent 3’ cleavage occurs following a CoTC-dependent release of the pre-mRNA, thereby acting as a compensatory mechanism to maintain expression of genes involved in the p53 pathway and DNA damage response.

## Materials and Methods

### Cell culture, siRNA transfections and UV irradiation

A549 cells were cultured in DMEM (Eurobio) containing 10% FCS (Pan Biotech) and L-Glutamine (Eurobio) at 37°C in 5% CO_2_. siRNA reverse transfections were performed in 10 cm with Lipofectamine RNAiMAX (Thermo Scientific) at a final concentration of 20 nM siRNA (Eurogentec or Dharmacon; see Supplementary Table 2) as per the manufacturer’s instructions in OptiMEM reduced serum media (Thermo Scientific). After 48 h transfection, cells were washed with PBS and irradiated with 40 J/m^2^ UV (254 nm; Stratalinker), placed in fresh media and harvested on ice after 16h of recovery at 37°C.

### CRISPR-mediated deletion of CoTC element

CRISPR sgRNAs were designed for the CoTC element deletion of the p53 gene (Supplementary Figure 3A). sgRNAs were designed using the online tool http://crispr.mit.edu/. Guide sequences are identified that minimize identical genomic matches or near-matches to reduce risk of cleavage away from target sites (off-target effects). The guide sequences are constructed such that they consist of 20-mer protospacer sequence upstream of an NGG protospacer adjacent motif (PAM) at the genomic recognition site (Supplementary Table 3).

2 sgRNA oligos are constructed, each of 24-25 mer oligos and their associated reverse complement including additional nucleotides for cloning and expression purposes.

The two plasmids used are namely, pSpCas9 (BB) plasmid pX458 and pX459, which include GFP and puromycin as selectable markers, respectively.

1. First the sequences CACC and AAAC are added before the 20-mer guide sequence and the guide’s reverse complement for cloning into pX458/pX459 vectors using BbsI restriction enzyme (Supplementary Table 4).
2. A G nucleotide is added after the CACC sequence and before the 20-mer if the first position of the 20-mer is not G. sgRNA expression from the U6 promoter of the pX458/pX459 vector is enhanced by the inclusion of a G nucleotide after the CACC sequence.
3. A C nucleotide is added at the 3’-end of the reverse complement oligo. All resultant oligos are 25-mer oligos.

The sgRNA oligo sequences were cloned into the pX458 and pX459 plasmids using a Golden Gate assembly cloning strategy (Supplementary figure 3B). The plasmids were amplified followed by transfection of CRISPRS and selection of transfected cells were carried out in Puromycin containing medium. The cells are incubated for a total of 48–72 hours after transfection before harvesting for indel analysis.

Primers were designed surrounding the sgRNA cleavage sites for and screening for CRISPR/Cas9 screening deletion (Supplementary Figure 3). gDNA was isolated from control or transfected cells and PCR is performed to validate the primers and verify the presence of the intended genomic deletion.

### Cell fractionation

Cell pellets were resuspended in approximately 3 x cell pellet volume of lysis Buffer A (10 mM HEPES pH 7.9, 15 mM MgCl_2_, 10 mM KCl, 0.1% NP40, 1 mM DTT) containing RNAseOut (Thermo Scientific) and incubated on ice for 15 min. Cells were then pelleted at 3000 rpm for 5 min at 4°C and the supernatant retained for cytoplasmic RNA. Nuclear pellets were washed in 2 x 1 mL Lysis Buffer A at 3000 rpm for 5 min at 4°C, resuspended in 2 x pellet volume with Nuclear Lysis Buffer B (20 mM HEPES pH 7.9, 400 mM NaCl, 1.5 M MgCl_2_, 0.2 mM EDTA, 5 mM DTT) containing RNAseOut and incubated on ice for 30 min. Nuclear debris was pelleted at 13 000 rpm for 15 min at 4°C and supernatants placed in Trizol Reagent for RNA extraction.

### Nuclear fractionation

Cell nuclei was suspended in 1x nuclei pellet volume of buffer 1 (20 mM Tris pH 7.9, 75 mM NaCl, 0.5 mM EDTA, 0.85 mM DTT, 0.125 mM PMSF, 0.1 mg of yeast tRNA/mL, 50% glycerol) and 10x nuceli pellet volume of buffer 2 (20 mM HEPES, 300 mM NaCl, 1 mM DTT, 7.5 mM MgCl2, 0.2 mM EDTA, 1M Urea, 1% NP-40, 0.1 mg of yeast tRNA/mL). After vigorous agitation for 5 sec, nuclei pellet was incubated 10 min on ice. Chromatin fraction was then sedimented by full speed centrifugation for 5 min at 4°C. The supernatant corresponding to the nucleoplasmic fraction was transferred to a new tube, adjusted to 0.1% of SDS and trizol RNA extraction was proceeded. The insoluble fraction corresponding to the chromatin was resuspended in buffer 3 (10 mM Tris pH 7.5, 10 mM MgCl2, 500 mM NaCl) and 20U of DNAse were added before 30 min incubation at 37°C.RNA extraction was then proceeded.

### RT-qPCR and RT-PCR

cDNA was synthesized using Superscript III (Thermo Scientific). qPCR on cDNA derived from nuclear pre-mRNAs or cytoplasmic mRNAs was performed using 2 x Power Sybrgreen Master Mix (Thermo Scientific) and 0.4 μM oligonucleotide primers (Supplementary Table 5). The ratio of uncleaved/total RNA was calculated using 2^(total-uncleaved) and the ratio of released uncleaved RNA nucleoplasm/ chromatin-associated uncleaved pre-mRNA was calculated using 2^(Chromatin-nucleoplasm). For the RNA-IP, samples were normalised to the input using 2^(Input-IP). When conducting RT-PCR, cDNA amplification was performed using Go-Taq flexi DNA polymerase (Promega). PCR products were then applied to 1% agarose gel.

### Western Blot

Cells were harvested on ice and pelleted by centrifugation at 2000 rpm for 5 min at 4°C. Pellets were resuspended in RIPA Buffer containing complete protease inhibitors (EDTA-free) and sonicated (Bioruptor, Diagenode). Cellular debris was pelleted at 10 000 rpm for 10 min at 4°C and protein concentration determined. Primary antibodies used in this study from Bethyl Laboratories were: CPSF160, CPSF100, CPSF73, CPSF30, CstF77, CstF64, CstF50, CFIm68 and CFIm59. Other antibodies used include CFIm25 (PTGlabs), PCF11 (Santa-Cruz), CLP1 (Epitomics). GAPDH (Sigma), Ser2P (Millipore), Topoisomerase II α (Abcam), Histone H3 (Abcam), DHX36 (Abcam), hnRNP (H/F) (Abcam), p21 (Thermofisher) and p53 (Cell Signaling Technology).

### Single-molecule Fluorescence In-Situ Hybridization (FISH)

A549 cells were cultured on #1.5 cover glasses in 12-well plates. When cells were at approximately 50% confluency, the cover glass was washed once with PBS. Each cover glass was placed in one well of a 12-well plate that was filled with 200 μL PBS, which just barely covered the top of the cover glass. Half of the cover glasses were irradiated with UVC at 50 J/m^2^. After irradiation, cover glasses were returned into culture medium and placed in the incubator for 4 hours. After 4 hours, cells on cover glasses were washed 3 times with HBSS before fixation with 4% PFA in PBS. Fixed samples were washed with 1x PBS, and stored in 70% ethanol at 4°C overnight.

FISH probes were designed and ordered from Biosearch Stellaris using Quasar 570 and 670 fluorophores. Hybridization were performed according to the manufacturer’s protocol with minor modifications. Hybridized samples were mounted in Prolong Gold with DAPI and allowed to dry overnight.

Imaging of FISH was performed on a custom-built microscope. This microscope comprised an ASI (www.asiimaging.com) Rapid Automated Modular Microscope System (RAMM) base, a Hamamatsu ORCA-Flash4 V2 CMOS camera (https://www.hamamatsu.com/, C11440), Lumencore SpectraX (https://lumencor.com/), an ASI High Speed Filter Wheel (FW-1000), and an ASI MS-2000 Small XY stage. Excitation of DAPI, Quasar 570, and 670 was performed using SpectraX violet, red, and green respectively. Emission filters specific to these spectra were used. Image acquisition was performed through Micro-Manager. We obtained multiple z stacks at 0.5 μm intervals, spanning 3.5 μm. The maximum intensity projections were performed and used for transcript localization and analysis.

FISH analysis was performed with custom MATLAB software. Briefly, images of cells were segmented into nucleus and cytoplasm. Spots were localized with custom MATLAB software using an algorithm based on Thompson et al. The software outputs the number of nuclear and cytoplasmic spots per cell, and the distribution of spots per cell.

### Propidium Iodide Staining

Cells were harvested in ice and washed with PBS. They were fixed in 70% ethanol for 30 minutes at −20°C. They were washed twice in PBS pelleted by centrifugation at 850g for 5 min at 4°C. The cells were then resuspended in a solution containing 3.5mM Tris HCl pH 7.6 (Thermo Scientific), 10mM NaCl (Thermo Scientific), 50ug/mL Propidium Iodide (Sigma P4170), 0.1% IGEPAL (Thermo Scientific), 20ug/mL RnaseA (Sigma) and water. The acquisition of stained cells was performed using a LSRII flow cytometer (BD Biosciences). The acquired data was analyzed using FlowJo software.

### RNA-sequencing analysis

For RNA-seq, nuclear RNA from UV-irradiated and non-irradiated A549 cells (two biological replicates of each condition) was subjected to DNAse I treatment with TURBO DNase I (ThermoFisher Scientific), quantified and analyzed using an RNA 2100 Bioanalyzer (Agilent). 500 ng of good quality RNA (RIN > 9) were used for Illumina compatible library preparation using the TruSeq Stranded total RNA protocol allowing to take into account strand information. A first step of ribosomal RNA depletion was performed using the RiboZero Gold kit (Illumina). After fragmentation, cDNA synthesis was performed and resulting fragments were used for dA-tailing followed by ligation of TruSeq indexed adapters. PCR amplification was finally achieved to generate the final barcoded cDNA libraries. Libraries were equimolarly pooled. Sequencing was carried out on a HiSeq instrument (Illumina) to obtain around 40 million raw single-end reads of 100 nucleotides per sample.

Fastq files were generated using bcl2fastq. RNA-seq reads of good quality were trimmed in their 5’ and 3’-ends with the cutadapt software to remove uninformative nucleotides due to primer sequences. Trimmed reads of 100 bp or more were aligned on the Human genome (hg19) using Tophat2. Only reads with a mapping quality score of 20 or more were retained (samtools) for downstream analysis. Gene coordinates were obtained on the basis of overlapping Refseq transcripts with the same gene symbol. For each gene, two 500 bp regions located upstream and downstream the PAS at the end of the gene were defined (genes with downstream region overlapping another gene were discarded), and reads located in the upstream and downstream regions were counted in each sample. A table of counts was built with the featureCounts software (R version 3.4.0). Only genes with at least 10 reads in both regions in either condition were kept for further analysis. In total, 4,208 genes passed all these steps and were used for subsequent analysis. The differential analysis between the UV+ and UV-conditions was done using two independent biological replicates per condition. For each gene, the fold regulation of the downstream region (that is the ratio of normalized read counts between conditions) was compared to the fold regulation of the upstream region using a Wald test implemented in DESeq2 (Love *et al*, 2014).

Gene ontology (GO) analysis of genes was carried out by the functional enrichment analysis tool DAVID.

### Statistics

Statistical differences between experimental and control samples were assessed by unpaired t-test using GraphPad Prism, with significance achieved at *P* <0.05.

## Supplementary Figure and Table legends

**Supplementary Figure 1**

(A) Western Blot analysis of the CTD phospho-Ser2 expression in A549 cells treated with DRB (50 μM) for 24 hours or Flavopiridol (Fv) (1μM) for 6 hours prior to UV irradiation (40 J/m2).

(B) RT-qPCR to assess the efficiency of p53 and TBP pre-mRNA 3’-end processing in A549 cells treated with DRB (50 μM) for 24 hours without UV treatment.

(C) RT-qPCR to assess the efficiency of p53 and TBP pre-mRNA 3’-end processing in A549 cells treated with DRB (50 μM) for 24 hours prior to UV irradiation (40 J/m2).

(D) RT-qPCR to assess the efficiency of p53 and TBP pre-mRNA 3’-end processing in A549 cells treated with Flavopiridol (Fv) (1μM) for 6 hours prior to UV irradiation (40 J/m2).

**Supplementary Figure 2**

A549 cells were transfected with siRNA against hnRNP H/F 48 hours prior to UV irradiation (40 J/m2). The nucleus was fractionated into nucleoplasm and chromatin following 16 hours of recovery. (A) The depletion efficiency was verified by western blot. (B) RT-qPCR on RNA extracted from the nucleoplasm and the chromatin fraction to quantify the uncleaved/total ratio in both fractions.

**Supplementary Figure 3**

(A) CRISPR sgRNAs designed for the CoTC element deletion of the p53 gene.

(B) PCR band profile for ‘deletion’ bands for gDNA from wild type (WT) and CRISPR transfected cells. The profile shows bands for complete deletion (ΔCoTC) of the p53 CoTC element in A549 cells as opposed its partial deletion (pΔCoTC) in A549 and A375 cells.

**Supplementary Figure 4**

RT-qPCR assay on nuclear RNA for assessing the uncleaved/total ratio of p53 pre-mRNA in wild type (WT) and partial CoTC deleted (pΔCoTC) A549 (A) or A375 (B) cells treated with or without UV irradiation (40 J/m2). P <0.05; unpaired t-test.

**Supplementary Figure 5**

RT-qPCR analysis on RNA extracted from nucleoplasm and chromatin fractions. The ratio of uncleaved pre-mRNA (nucleoplasm/chromatin) was calculated to quantify the level of unprocessed *p53*, *WDR13*, *GAPDH* and *TBP* pre-mRNAs released in the nucleoplasm compared to the chromatin-bound unprocessed pre-mRNA in wild type (WT) and partial CoTC deleted (pΔCoTC) A549 or A375 cells treated with or without UV irradiation (40 J/m2). P <0.05; unpaired t-test.

**Supplementary Figure 6**

(A) RT-qPCR measuring relative p53 and p21 mRNA levels in wild type (WT) and partial CoTC-deleted (pΔCoTC) A549 cells in response to UV treatment (40 J/m2). The expression was normalized to HPRT. *P <0.05; unpaired t-test.

(B) Western blot analysis of p53 and p21 expression wild type (WT) and partial CoTC deleted (pΔCoTC) A549 cells treated with or without UV irradiation (40 J/m2).

**Supplementary Figure 7**

Representative flow-cytometry analyses of the cell cycle (DNA content by Propidium Iodide; PI) in wild type (WT) and partial CoTC deleted (pΔCoTC) A549 cells treated with or without UV irradiation (40 J/m2).. Indicated: percent of cells in the G0-G1, S and G2/M phases.

**Supplementary Figure 8**

(A) PCR analysis of RNA seq candidate gene 3’ flanking regions to map the location of CoTC elements.

(B) PCR analysis of 3’ flanking regions in candidate genes from the p53 signaling pathway to map the location of CoTC elements.

(C) Table to summarize the presence or absence of bands from PCR amplification using primer pairs F/(R1-R5) in 3’ flanking regions of candidate genes.

(D) PCR analysis of candidate gene 3’ flanking region using the same primers employed in the data panel above. Lane 1 is a control PCR amplification of cDNA derived from the reverse transcription of a control mRNA using oligo (dT). Lanes 2-6 are PCR amplification of reverse transcribed candidate gene chromatin-associated pre-mRNA using oligo oligo (dT). Likewise for all genes.

**Supplementary Figure 9**

Gene ontology (GO) analysis of the 108 pre-mRNAs with a more efficient 3’end processing in UV-treated compared to untreated cells (UV-resistant 3’-end processing) obtained from RNA Sequencing analyses. The bar chart shows the GO terms for biological processes, ranked by p-values, calculated by the functional enrichment analysis tool DAVID.

**Supplementary Table 1**

List of genes for pre-mRNAs with an upregulated or downregulated 3’end processing in UV-treated compared to untreated cells genes, as identified from RNA Sequencing analysis.

**Supplementary Table 2**

siRNA sequences for all genes tested.

**Supplementary Table 3**

20-mer protospacer sequences for two sgRNA and their reverse complement for the deletion of p53 CoTC

**Supplementary Table 4**

Protospacer sequences and their reverse complements with “CACC” and “AAAC” added for cloning into the pX458/pX459 vector using BbsI restriction enzyme

**Supplementary Table 5**

Oligonucleotides: Primer sequences used for all genes tested.

