## Supplementary figures and images for "Uncoupling from transcription protects polyadenylation site cleavage from inhibition by DNA damage"

### Supplementary Figure 1

#### Supplementary Figure 1

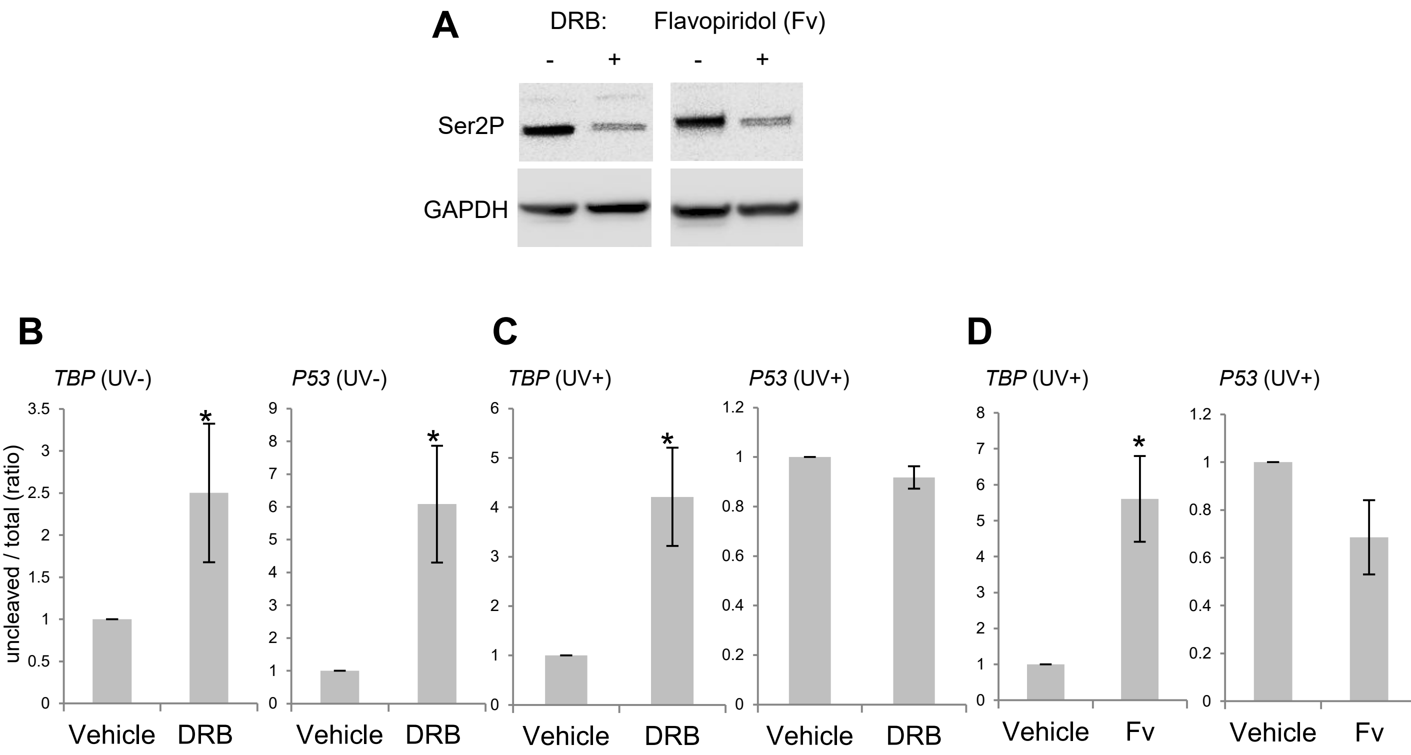

### Supplementary Figure 2

#### Supplementary Figure 2

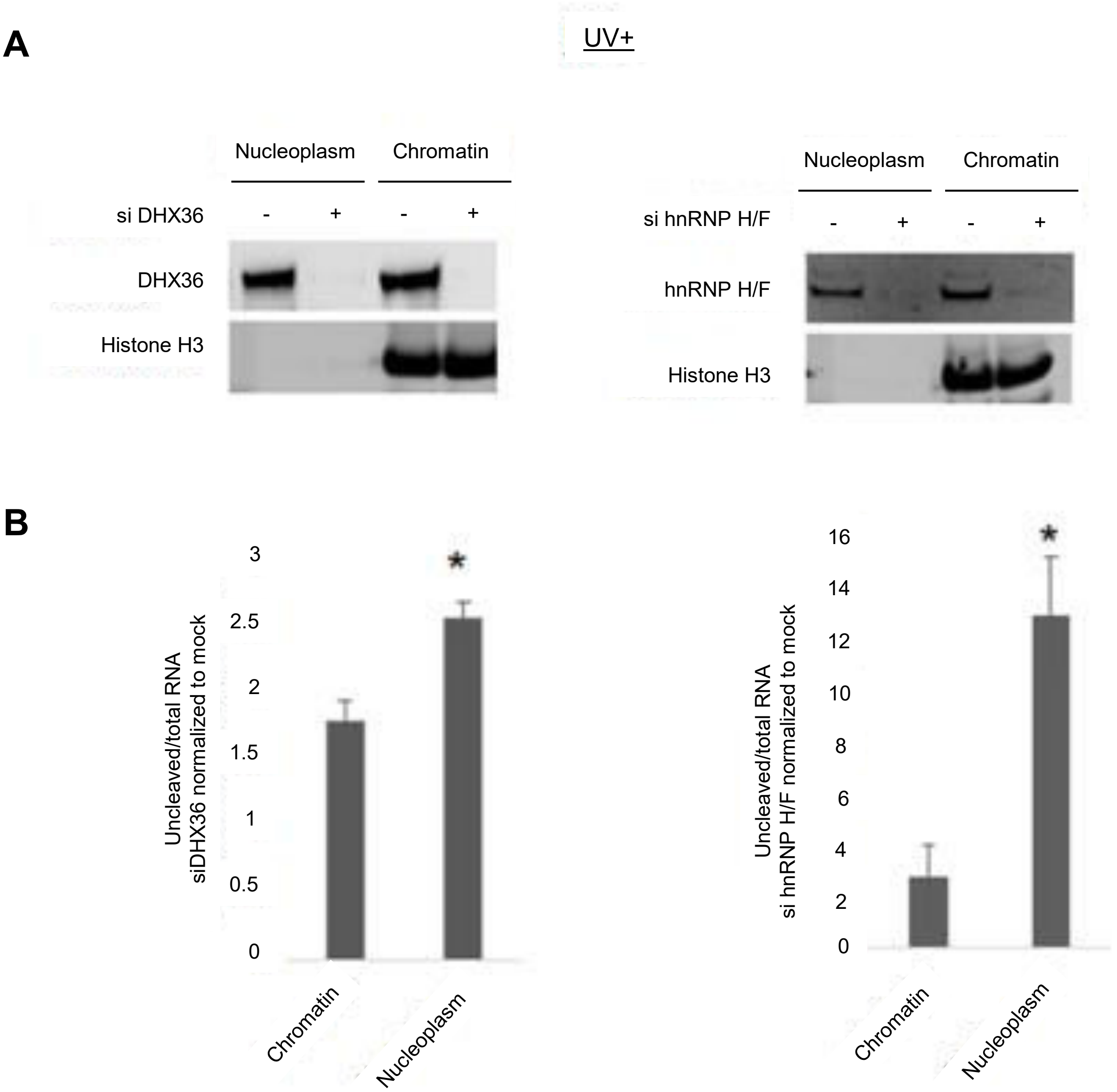

### Supplementary Figure 4

#### Supplementary Figure 4

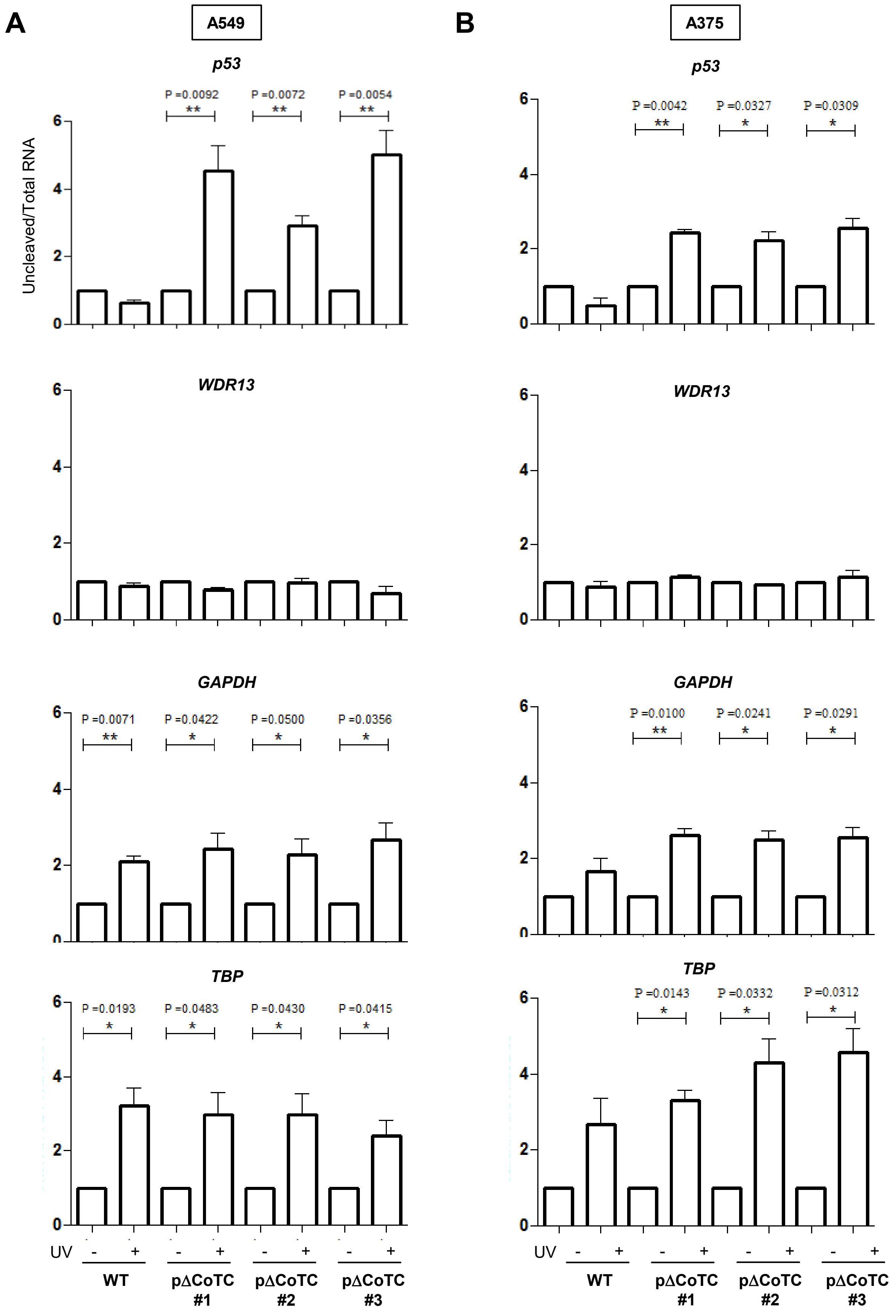

### Supplementary Figure 5

##### Supplementary Figure 5

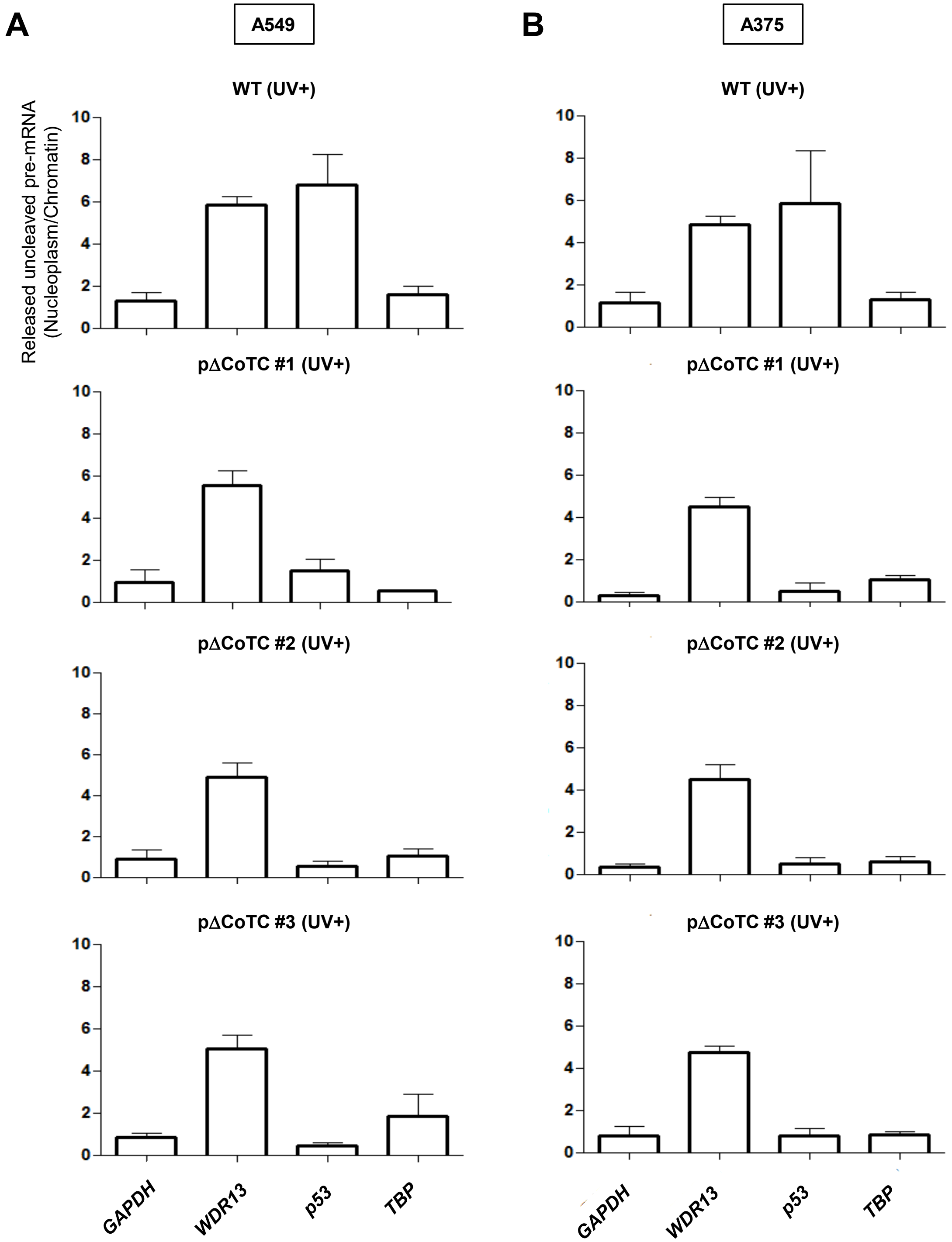

### Supplementary Figure 6

#### Supplementary Figure 6

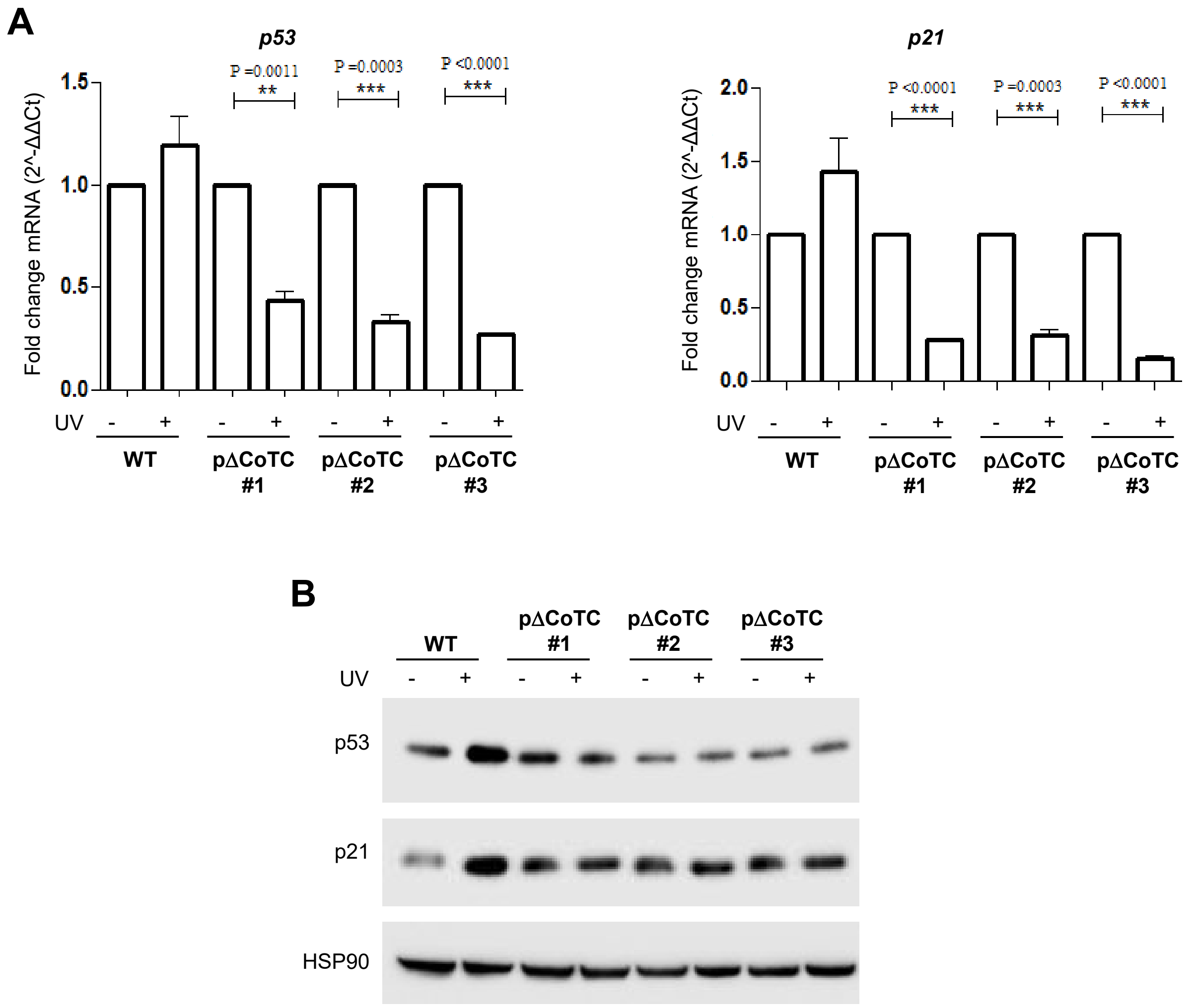

### Supplementary Figure 7

#### Supplementary Figure 7

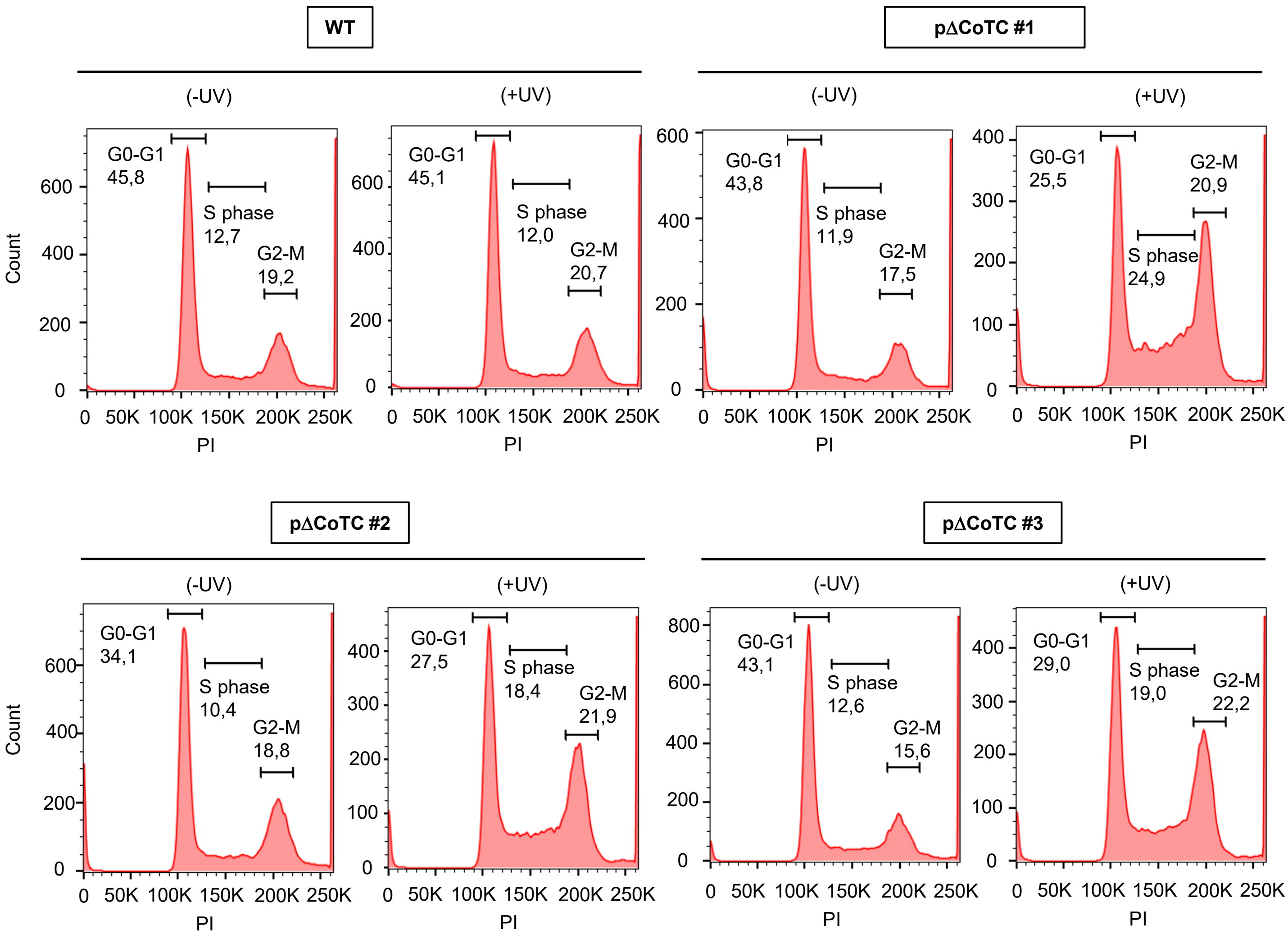

### Supplementary Figure 9

**Supplementary Figure 9**

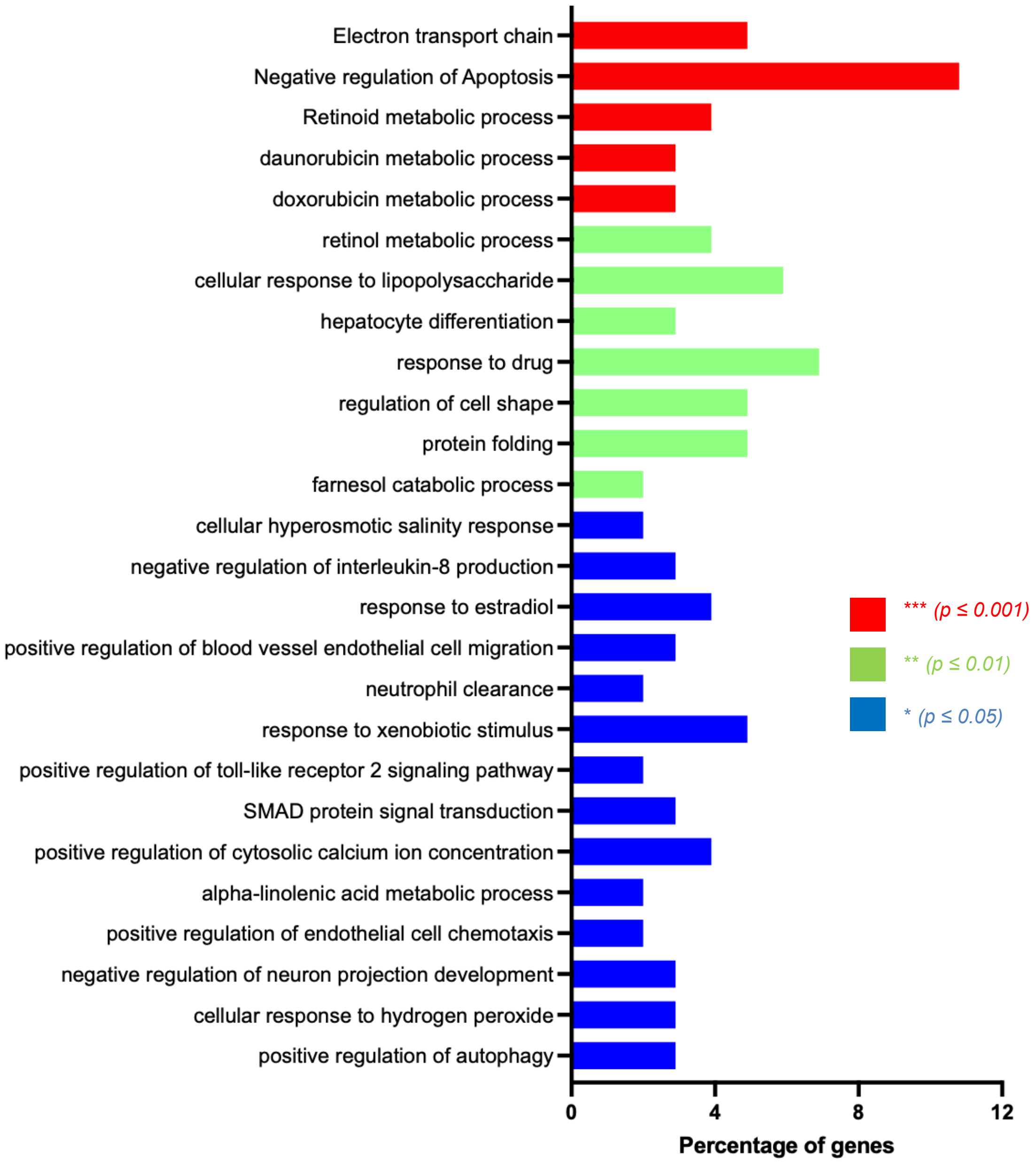
