## Supplementary Figure 3 for "Uncoupling from transcription protects polyadenylation site cleavage from inhibition by DNA damage"

**A**

Deletion screening primer (forward)

sgRNA A

5' GTCCCTA***CCCAGCAGGCAAACTAGAG***CTCCTGAAGCTCAGTCC**CTGTCCTTGCCTCTGTAGAC**AGGTCACCTTGA 3'

3' CAGGGATGGGTCGTCCGTTTGATCTCGAGGACTTCGAGTCAGGGACAGGAACGGAGACATCTGTCCAGTGGAACT 5'

p53 CoTC element

5' TGA**GCTTCCTTTTTTTTTTTTTTAATTTTTTTTTTATTTTAGGCTTTATT**GGGGCATAATTGATCCCCCAAAATTGCATACA 3'

3' ACTCGAAGGAAAAAAAAAAAAAATTAAAAAAAAAATAAAATCCGAAATAACCCCGTATAACTAGGGGGTTTTAACGTATGT 5'

5' TTCAAGGTATGCAGTGTGATGATTTGATATGGGGGTATATTGTGAAACCATTACCACAATCAAATTAATCAGCACGTCC 3'

3' AAGTTCCATACGTCACACTACTAAACTATACCCCCATATAACACTTTGGTAATGGT**GTTAGTTTAATTAGTCGTGC**AGG 5'

sgRNA B

5' ATCATCACACACAGTTACCATTTGTGTGTGTGCACGTGTGTTCACCTACGACGAGGACACTTGGACCTACTCTGCAGAT 3'

3' TAGTAGTGTGTGTCAAGGTAAACACACACACGTGCACACAAGTGGATGCTGCTCCTGTGAACCTGGATGAGACGTCTA 5'

5' CTCAAGTAAACAGAAAATCTCCCTTTTTGACAACCATCCTCCACCCTTTCAATCCC 3'

3' GAGTTCATTTGTCTTTTAGAGGGAA***AAACTGTTGGTAGGAGGTGG***GAAAGTTAGGG 5'

Deletion screening primer (reverse)

**B**

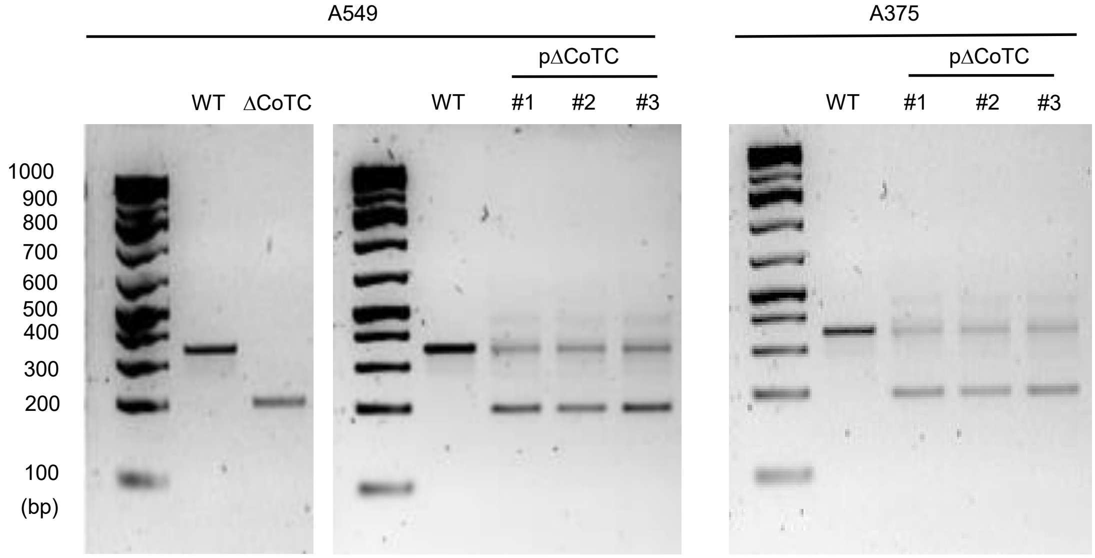
