## Supplementary Figure 8 for "Uncoupling from transcription protects polyadenylation site cleavage from inhibition by DNA damage"

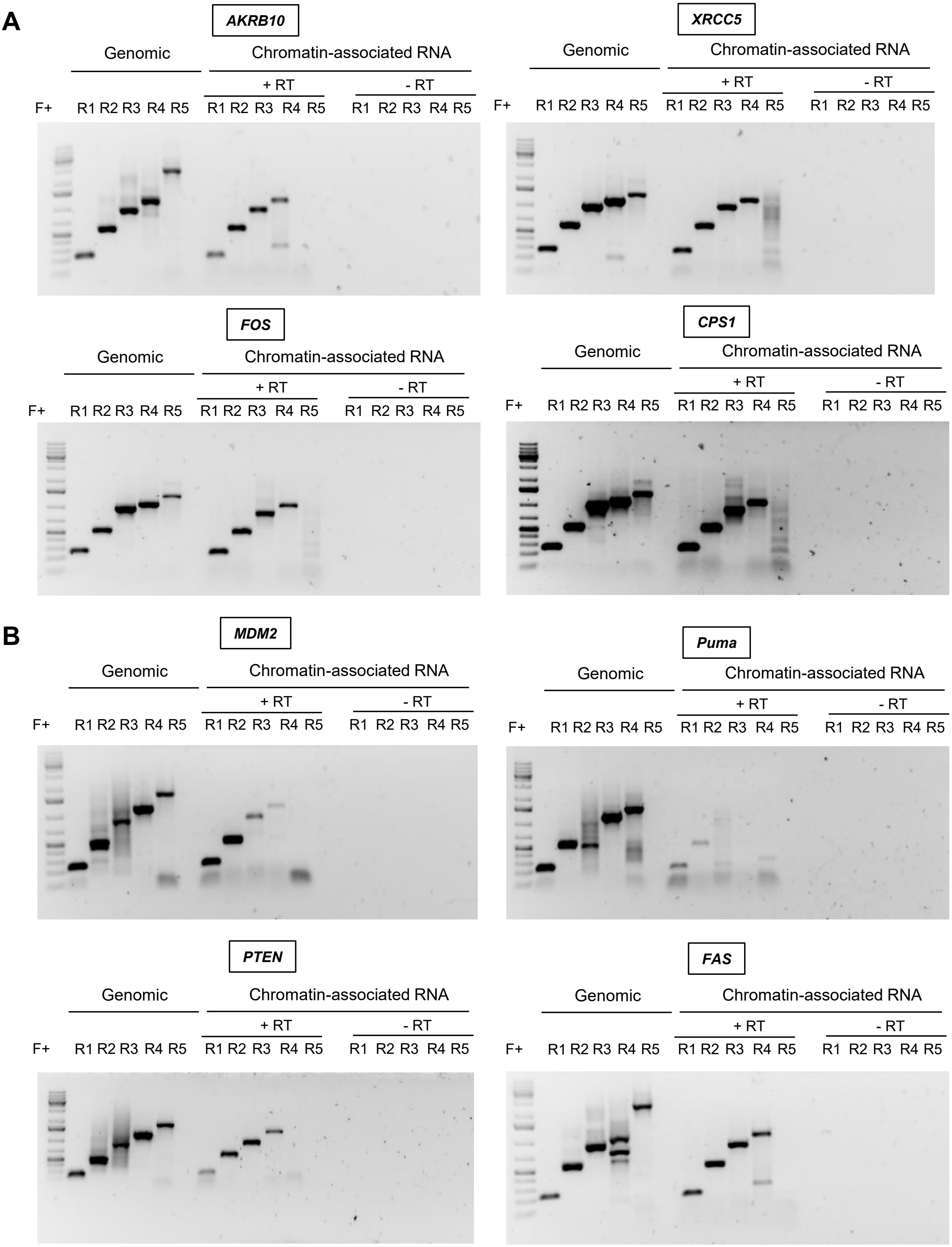

### Supplementary Figure 8 (continued)

**C**

| Basepairs downstream of polyA | 100-200 | 400-600 | 1000-1200 | 1500-2000 | 2500-3000 |
|---|---|---|---|---|---|
| MDM2 | + | + | + | + | - |
| Puma | + | + | - | - | - |
| PTEN | + | + | + | + | - |
| FAS | + | + | + | + | - |
| AKRB10 | + | + | + | + | - |
| XRCC5 | + | + | + | + | - |
| FOS | + | + | + | + | - |
| CPS1 | + | + | + | + | - |

COTC cleavage sites between 1 – 2.5 kb downstream to the PAS.

**D**

Chromatin-associated RNA
(RT with oligo dT)

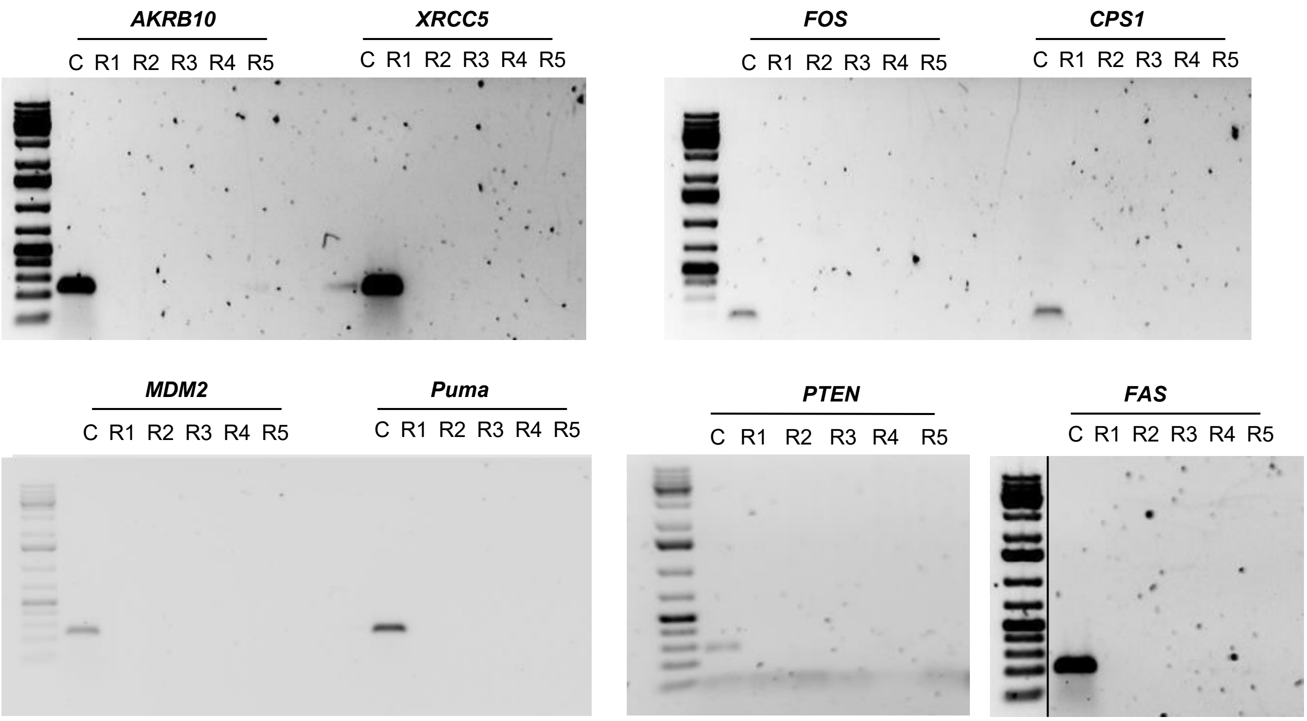
