## Supplementary Table 1 for "Uncoupling from transcription protects polyadenylation site cleavage from inhibition by DNA damage"

| downstream_region | Chromosome | Gene start | Gene end | Strand | Gene symbol | NM/NR | baseMean | log2FoldChange | lfcSE | stat | pvalue | padj | downstream_UV+_n1(99) | downstream_UV+_n2(138) | downstream_UV-_n1(95) | downstream_UV-_n2(96) | upstream_UV+_n1(99) | upstream_UV+_n2(138) | upstream_UV-_n1(95) | upstream_UV-_n2(96) | logFC_downstream | logFC_upstream | logFC_downstream-logFC_upstream |
|---|---|---|---|---|---|---|---|---|---|---|---|---|---|---|---|---|---|---|---|---|---|---|---|
| chr11:309410-309910:downstream_8969 | chr11 | 308106 | 309410 | + | IFITM2 | NM | 30 | -5,9 | 2 | 3 | 5,16E-04 | 2,61E-02 | 0 | 0 | 8 | 5 | 72 | 62 | 43 | 46 | -Inf | 0,6 | -Inf |
| chr9:107522403-107522903:downstream_7245 | chr9 | 107509968 | 107522403 | + | NIPSNAP3A | NM | 8 | -5,8 | 2 | 3 | 4,53E-03 | NA | 0 | 0 | 6 | 6 | 16 | 14 | 8 | 11 | -Inf | 0,6 | -Inf |
| chr6:18223899-18224399:downstream_24726 | chr6 | 18224399 | 18264799 | - | DEK | NM | 19 | -5,4 | 2 | 3 | 1,96E-03 | NA | 0 | 0 | 11 | 7 | 32 | 29 | 39 | 37 | -Inf | -0,3 | -Inf |
| chr4:174245118-174245618:downstream_3539 | chr4 | 174089903 | 174245118 | + | GALNT7 | NM | 48 | -5,4 | 2 | 3 | 1,05E-03 | 4,15E-02 | 0 | 0 | 7 | 9 | 98 | 68 | 95 | 107 | -Inf | -0,3 | -Inf |
| chr3:31679114-31679614:downstream_2598 | chr3 | 31573992 | 31679114 | + | STT3B | NM | 282 | -5,1 | 2 | 3 | 1,20E-03 | 4,15E-02 | 0 | 0 | 11 | 5 | 412 | 553 | 572 | 705 | -Inf | -0,4 | -Inf |
| chr16:89977792-89978292:downstream_12549 | chr16 | 89939993 | 89977792 | + | TCF25 | NM | 21 | -5,1 | 2 | 3 | 3,97E-03 | NA | 0 | 0 | 8 | 4 | 27 | 47 | 35 | 43 | -Inf | -0,1 | -Inf |
| chr7:27564558-27565058:downstream_5529 | chr7 | 27565058 | 27702620 | - | HIBADH | NM | 36 | -5,1 | 2 | 3 | 2,70E-03 | 6,93E-02 | 0 | 0 | 4 | 7 | 55 | 83 | 63 | 78 | -Inf | 0,0 | -Inf |
| chr3:148879696-148880196:downstream_2565 | chr3 | 148880196 | 148939832 | - | CP | NR | 130 | -5,0 | 2 | 3 | 1,55E-03 | 4,88E-02 | 0 | 0 | 7 | 6 | 230 | 247 | 303 | 250 | -Inf | -0,2 | -Inf |
| chr10:115348082-115348582:downstream_17674 | chr10 | 115348582 | 115423829 | - | NRAP | NM | 5 | -5,0 | 2 | 2 | 2,94E-02 | NA | 6 | 8 | 7 | 4 | 2 | 14 | 0 | 0 | 0,3 | Inf | -Inf |
| chr5:52979171-52979671:downstream_4016 | chr5 | 52856464 | 52979171 | + | NDUFS4 | NM | 19 | -4,8 | 2 | 3 | 7,71E-03 | NA | 0 | 0 | 4 | 11 | 31 | 18 | 60 | 30 | -Inf | -0,9 | -Inf |
| chr3:152183569-152184069:downstream_23265 | chr3 | 151985828 | 152183569 | + | MBNL1 | NM | 22 | -4,6 | 2 | 3 | 8,70E-03 | NA | 0 | 0 | 6 | 15 | 13 | 31 | 49 | 64 | -Inf | -1,4 | -Inf |
| chr3:42690233-42690733:downstream_2889 | chr3 | 42642146 | 42690233 | + | NKTR | NM | 11 | -4,6 | 2 | 2 | 1,79E-02 | NA | 0 | 0 | 3 | 8 | 14 | 18 | 14 | 32 | -Inf | -0,5 | -Inf |
| chrX:24234372-24234872:downstream_27125 | chrX | 24167761 | 24234372 | + | ZFX | NM | 7 | -4,6 | 2 | 2 | 3,28E-02 | NA | 0 | 0 | 21 | 12 | 3 | 1 | 10 | 8 | -Inf | -2,1 | -Inf |
| chr6:83919609-83920109:downstream_4758 | chr6 | 83920109 | 84140938 | - | ME1 | NM | 51 | -4,5 | 2 | 3 | 4,13E-03 | 8,70E-02 | 1 | 0 | 15 | 11 | 78 | 88 | 106 | 108 | -5,0 | -0,4 | -4,7 |
| chr7:80371353-80371853:downstream_5581 | chr7 | 80371853 | 80548667 | - | SEMA3C | NM | 80 | -4,5 | 2 | 3 | 5,50E-03 | 9,98E-02 | 0 | 0 | 7 | 6 | 91 | 141 | 207 | 185 | -Inf | -0,8 | -Inf |
| chr2:45615318-45615818:downstream_1615 | chr2 | 45615818 | 45838433 | - | SRBD1 | NM | 10 | -4,5 | 2 | 2 | 2,36E-02 | NA | 0 | 0 | 13 | 7 | 3 | 15 | 25 | 20 | -Inf | -1,3 | -Inf |
| chr11:16808706-16809206:downstream_8855 | chr11 | 16809206 | 17035963 | - | PLEKHA7 | NM | 10 | -4,5 | 2 | 2 | 2,22E-02 | NA | 1 | 0 | 18 | 20 | 2 | 12 | 17 | 11 | -5,5 | -1,0 | -4,6 |
| chr7:11409561-11410061:downstream_5470 | chr7 | 11410061 | 11871824 | - | THSD7A | NM | 54 | -4,4 | 2 | 3 | 5,98E-03 | 1,05E-01 | 0 | 0 | 29 | 14 | 26 | 31 | 174 | 162 | -Inf | -2,6 | -Inf |
| chr10:118642387-118642887:downstream_17768 | chr10 | 118642887 | 118886097 | - | SHTN1 | NM | 8 | -4,3 | 2 | 2 | 3,81E-02 | NA | 0 | 0 | 7 | 14 | 3 | 7 | 25 | 7 | -Inf | -1,7 | -Inf |
| chr5:145492088-145492588:downstream_4501 | chr5 | 145492588 | 145562294 | - | LARS | NM | 16 | -4,3 | 2 | 2 | 1,87E-02 | NA | 0 | 0 | 14 | 7 | 18 | 7 | 35 | 45 | -Inf | -1,7 | -Inf |
| chr13:26978569-26979069:downstream_10567 | chr13 | 26828755 | 26978569 | + | CDK8 | NM | 12 | -4,3 | 2 | 2 | 2,59E-02 | NA | 1 | 0 | 39 | 18 | 2 | 7 | 14 | 13 | -6,1 | -1,7 | -4,4 |
| chr17:58520009-58520509:downstream_20331 | chr17 | 58520509 | 58603601 | - | APPBP2 | NM | 9 | -4,2 | 2 | 2 | 2,98E-02 | NA | 0 | 0 | 11 | 7 | 7 | 8 | 21 | 22 | -Inf | -1,6 | -Inf |
| chr20:43061485-43061985:downstream_22439 | chr20 | 42984440 | 43061485 | + | HNF4A | NM | 14 | -4,2 | 2 | 2 | 2,28E-02 | NA | 0 | 0 | 14 | 8 | 7 | 13 | 32 | 37 | -Inf | -1,9 | -Inf |
| chr19:38228702-38229202:downstream_21680 | chr19 | 38229202 | 38270230 | - | ZNF573 | NM | 10 | -4,2 | 2 | 2 | 3,45E-02 | NA | 0 | 0 | 6 | 12 | 5 | 10 | 17 | 28 | -Inf | -1,6 | -Inf |
| chr7:94299006-94299506:downstream_25665 | chr7 | 94285636 | 94299006 | + | PEG10 | NM | 23 | -4,1 | 2 | 2 | 1,88E-02 | NA | 0 | 0 | 3 | 9 | 28 | 26 | 72 | 48 | -Inf | -1,2 | -Inf |
| chrX:48387104-48387604:downstream_7706 | chrX | 48380163 | 48387104 | + | EBP | NM | 245 | -4,1 | 2 | 3 | 8,25E-03 | 1,21E-01 | 1 | 0 | 10 | 5 | 505 | 509 | 472 | 459 | -4,2 | 0,1 | -4,3 |
| chr17:20902405-20902905:downstream_12907 | chr17 | 20902905 | 20946352 | - | USP22 | NM | 19 | -4,1 | 2 | 2 | 2,16E-02 | NA | 0 | 0 | 4 | 7 | 21 | 23 | 56 | 42 | -Inf | -1,1 | -Inf |
| chr2:187545629-187546129:downstream_21993 | chr2 | 187454789 | 187545629 | + | ITGAV | NM | 72 | -4,0 | 2 | 2 | 1,31E-02 | 1,49E-01 | 0 | 0 | 6 | 5 | 102 | 90 | 174 | 198 | -Inf | -0,9 | -Inf |
| chr9:136228041-136228541:downstream_26523 | chr9 | 136223420 | 136228041 | + | SURF2 | NM | 8 | -4,0 | 2 | 2 | 4,88E-02 | NA | 9 | 26 | 7 | 11 | 5 | 8 | 0 | 0 | 1,0 | Inf | -Inf |
| chr1:193223942-193224442:downstream_268 | chr1 | 193091087 | 193223942 | + | CDC73 | NM | 8 | -4,0 | 2 | 2 | 4,97E-02 | NA | 0 | 0 | 8 | 4 | 4 | 13 | 15 | 22 | -Inf | -1,2 | -Inf |
| chrX:118967488-118967988:downstream_27094 | chrX | 118967988 | 118986991 | - | UPF3B | NM | 15 | -4,0 | 2 | 2 | 2,81E-02 | NA | 0 | 0 | 8 | 10 | 14 | 9 | 36 | 45 | -Inf | -1,9 | -Inf |
| chr14:31359822-31360322:downstream_19186 | chr14 | 31343740 | 31359822 | + | COCH | NM | 12 | -3,9 | 2 | 2 | 3,77E-02 | NA | 1 | 0 | 10 | 2 | 15 | 32 | 22 | 17 | -3,9 | 0,2 | -4,1 |
| chrX:115033103-115033603:downstream_7546 | chrX | 115033603 | 115085444 | - | LOC642776 | NR | 10 | -3,9 | 2 | 2 | 3,75E-02 | NA | 1 | 0 | 6 | 11 | 13 | 15 | 22 | 13 | -4,3 | -0,3 | -4,0 |
| chr4:89010915-89011415:downstream_23535 | chr4 | 89011415 | 89152474 | - | ABCG2 | NM | 23 | -3,9 | 2 | 2 | 2,53E-02 | NA | 0 | 0 | 10 | 5 | 17 | 26 | 57 | 67 | -Inf | -1,5 | -Inf |
| chr17:16229573-16230073:downstream_12609 | chr17 | 16120508 | 16229573 | + | PIGL | NM | 21 | -3,9 | 1 | 3 | 6,62E-03 | NA | 2 | 0 | 22 | 7 | 37 | 30 | 38 | 34 | -4,1 | -0,1 | -4,1 |
| chr12:52679196-52679696:downstream_10174 | chr12 | 52679696 | 52685299 | - | KRT81 | NM | 379 | -3,9 | 1 | 5 | 1,96E-07 | 4,24E-05 | 3 | 3 | 35 | 28 | 936 | 827 | 507 | 690 | -3,3 | 0,6 | -3,9 |
| chr1:175913461-175913961:downstream_17175 | chr1 | 175913961 | 176176380 | - | RFWD2 | NM | 13 | -3,8 | 2 | 2 | 3,83E-02 | NA | 0 | 0 | 11 | 9 | 6 | 10 | 35 | 30 | -Inf | -2,1 | -Inf |
| chr22:30774302-30774802:downstream_15557 | chr22 | 30774802 | 30783302 | - | RNF215 | NM | 14 | -3,7 | 2 | 2 | 4,06E-02 | NA | 1 | 0 | 7 | 8 | 19 | 21 | 32 | 23 | -4,2 | -0,5 | -3,8 |
| chr12:96051082-96051582:downstream_9902 | chr12 | 96051582 | 96184536 | - | NTN4 | NM | 43 | -3,6 | 2 | 2 | 3,12E-02 | 2,19E-01 | 0 | 0 | 8 | 6 | 48 | 25 | 113 | 140 | -Inf | -1,8 | -Inf |
| chr1:52254365-52254865:downstream_17033 | chr1 | 52254865 | 52344609 | - | NRD1 | NM | 45 | -3,6 | 2 | 2 | 2,75E-02 | 2,09E-01 | 1 | 0 | 11 | 7 | 47 | 78 | 111 | 104 | -4,5 | -0,8 | -3,7 |
| chr15:62144089-62144589:downstream_19837 | chr15 | 62144589 | 62352664 | - | VPS13C | NM | 34 | -3,5 | 2 | 2 | 3,84E-02 | 2,39E-01 | 0 | 0 | 4 | 8 | 19 | 41 | 95 | 107 | -Inf | -1,8 | -Inf |
| chr9:75515077-75515577:downstream_7138 | chr9 | 75515577 | 75568233 | - | ALDH1A1 | NM | 3453 | -3,5 | 0 | 8 | 5,38E-15 | 4,08E-12 | 49 | 43 | 459 | 379 | 7082 | 7581 | 5239 | 6790 | -3,2 | 0,3 | -3,5 |
| chr1:161933860-161934360:downstream_229 | chr1 | 161736033 | 161933860 | + | ATF6 | NM | 20 | -3,4 | 2 | 2 | 4,98E-02 | NA | 0 | 0 | 7 | 8 | 11 | 17 | 65 | 55 | -Inf | -2,1 | -Inf |
| chr1:153515594-153516094:downstream_17210 | chr1 | 153516094 | 153518282 | - | S100A4 | NM | 374 | -3,4 | 1 | 4 | 1,84E-05 | 2,53E-03 | 1 | 6 | 31 | 16 | 953 | 811 | 469 | 710 | -2,8 | 0,6 | -3,4 |
| chrX:107940775-107941275:downstream_26671 | chrX | 107683073 | 107940775 | + | COL4A5 | NM | 37 | -3,4 | 2 | 2 | 4,10E-02 | 2,46E-01 | 0 | 0 | 7 | 7 | 28 | 25 | 121 | 103 | -Inf | -2,1 | -Inf |
| chr14:75748937-75749437:downstream_11234 | chr14 | 75745480 | 75748937 | + | FOS | NM | 22 | -3,4 | 1 | 3 | 9,17E-03 | NA | 3 | 0 | 7 | 4 | 55 | 67 | 26 | 11 | -1,8 | 1,7 | -3,5 |
| chr11:71713965-71714465:downstream_18062 | chr11 | 71709957 | 71713965 | + | IL18BP | NM | 9 | -3,4 | 1 | 2 | 2,03E-02 | NA | 7 | 2 | 10 | 17 | 12 | 15 | 6 | 3 | -1,7 | 1,6 | -3,3 |
| chr4:141309106-141309606:downstream_23593 | chr4 | 141309606 | 141348815 | - | CLGN | NM | 10 | -3,3 | 2 | 2 | 4,20E-02 | NA | 2 | 0 | 6 | 8 | 16 | 19 | 10 | 18 | -3,1 | 0,4 | -3,4 |
| chr3:188608460-188608960:downstream_23249 | chr3 | 187871660 | 188608460 | + | LPP | NM | 71 | -3,3 | 2 | 2 | 3,48E-02 | 2,29E-01 | 0 | 1 | 21 | 10 | 48 | 76 | 207 | 202 | -5,0 | -1,7 | -3,3 |
| chr19:38819649-38820149:downstream_14259 | chr19 | 38810483 | 38819649 | + | KCNK6 | NM | 11 | -3,3 | 1 | 2 | 2,44E-02 | NA | 3 | 0 | 7 | 6 | 32 | 20 | 8 | 11 | -2,0 | 1,5 | -3,4 |
| chr13:48516290-48516790:downstream_10746 | chr13 | 48516790 | 48575462 | - | SUCLA2 | NM | 13 | -3,3 | 2 | 2 | 3,28E-02 | NA | 2 | 0 | 6 | 11 | 19 | 25 | 15 | 29 | -3,4 | 0,0 | -3,4 |
| chr12:69159853-69160353:downstream_10308 | chr12 | 69139935 | 69159853 | + | SLC35E3 | NM | 13 | -3,2 | 1 | 2 | 1,59E-02 | NA | 2 | 2 | 7 | 14 | 30 | 19 | 19 | 13 | -2,6 | 0,6 | -3,2 |
| chrX:102633092-102633592:downstream_26939 | chrX | 102631250 | 102633092 | + | NGFRAP1 | NM | 119 | -3,2 | 1 | 3 | 1,08E-02 | 1,39E-01 | 2 | 0 | 4 | 7 | 236 | 319 | 153 | 229 | -2,7 | 0,5 | -3,3 |
| chr4:155524785-155525285:downstream_23655 | chr4 | 155525285 | 155533902 | - | FGG | NM | 554 | -3,1 | 1 | 3 | 9,69E-03 | 1,32E-01 | 1 | 1 | 3 | 11 | 981 | 1235 | 1212 | 988 | -2,9 | 0,0 | -2,9 |
| chr11:88026259-88026759:downstream_17975 | chr11 | 88026759 | 88070941 | - | CTSC | NM | 191 | -3,0 | 1 | 3 | 4,65E-03 | 9,28E-02 | 2 | 0 | 7 | 7 | 384 | 509 | 289 | 326 | -2,6 | 0,5 | -3,1 |
| chr5:76986493-76986993:downstream_24406 | chr5 | 76986993 | 77072185 | - | TBCA | NM | 32 | -3,0 | 1 | 2 | 1,77E-02 | 1,71E-01 | 7 | 5 | 6 | 10 | 17 | 179 | 21 | 14 | -0,4 | 2,5 | -2,9 |
| chr8:62412614-62413114:downstream_25856 | chr8 | 62413114 | 62627199 | - | ASPH | NM | 141 | -2,9 | 1 | 2 | 1,60E-02 | 1,67E-01 | 2 | 0 | 4 | 17 | 234 | 171 | 337 | 363 | -3,7 | -0,8 | -2,9 |
| chr7:94214035-94214535:downstream_25724 | chr7 | 94214535 | 94285521 | - | SGCE | NM | 31 | -2,9 | 1 | 2 | 3,34E-02 | 2,25E-01 | 2 | 0 | 7 | 10 | 36 | 66 | 49 | 83 | -3,4 | -0,4 | -3,0 |
| chr14:24608176-24608676:downstream_19352 | chr14 | 24605366 | 24608176 | + | PSME1 | NM | 52 | -2,9 | 1 | 3 | 8,46E-03 | 1,23E-01 | 2 | 1 | 1 | 11 | 134 | 125 | 56 | 84 | -1,9 | 0,9 | -2,8 |
| chr7:75933614-75934114:downstream_5914 | chr7 | 75931874 | 75933614 | + | HSPB1 | NM | 44 | -2,8 | 1 | 3 | 7,67E-03 | 1,18E-01 | 2 | 2 | 6 | 11 | 92 | 104 | 60 | 72 | -2,2 | 0,6 | -2,8 |
| chr19:54955493-54955993:downstream_14642 | chr19 | 54955993 | 54960216 | - | LENG8-AS1 | NR | 32 | -2,8 | 1 | 3 | 7,73E-03 | 1,18E-01 | 6 | 0 | 19 | 14 | 76 | 43 | 53 | 43 | -2,6 | 0,3 | -2,9 |
| chr4:95589378-95589878:downstream_23794 | chr4 | 95373007 | 95589378 | + | PDLIM5 | NM | 235 | -2,8 | 1 | 3 | 1,19E-03 | 4,15E-02 | 4 | 0 | 29 | 47 | 212 | 283 | 616 | 684 | -4,2 | -1,4 | -2,8 |
| chr10:75879514-75880014:downstream_17475 | chr10 | 75880014 | 75910826 | - | AP3M1 | NM | 29 | -2,7 | 1 | 2 | 1,26E-02 | 1,46E-01 | 29 | 41 | 63 | 81 | 9 | 7 | 1 | 3 | -1,0 | 1,7 | -2,8 |
| chr1:236648008-236648508:downstream_16679 | chr1 | 236557679 | 236648008 | + | EDARADD | NM | 202 | -2,7 | 1 | 3 | 7,80E-04 | 3,29E-02 | 6 | 4 | 21 | 22 | 303 | 621 | 213 | 427 | -2,2 | 0,5 | -2,7 |
| chr19:6585349-6585849:downstream_13929 | chr19 | 6585849 | 6591163 | - | CD70 | NM | 46 | -2,7 | 1 | 2 | 2,16E-02 | 1,90E-01 | 2 | 0 | 6 | 7 | 87 | 110 | 72 | 85 | -2,4 | 0,3 | -2,7 |
| chr3:105295757-105296257:downstream_22992 | chr3 | 105085556 | 105295757 | + | ALCAM | NM | 371 | -2,5 | 1 | 2 | 2,84E-02 | 2,09E-01 | 2 | 0 | 10 | 12 | 455 | 469 | 1003 | 1016 | -3,8 | -1,1 | -2,6 |
| chr15:25325381-25325881:downstream_11588 | chr15 | 25325287 | 25325381 | + | SNORD116-14 | NR | 45 | -2,4 | 1 | 3 | 5,23E-03 | 9,95E-02 | 6 | 3 | 10 | 12 | 106 | 119 | 54 | 52 | -1,4 | 1,1 | -2,4 |
| chr4:74860858-74861358:downstream_3742 | chr4 | 74861358 | 74864446 | - | CXCL5 | NM | 26 | -2,4 | 1 | 2 | 2,19E-02 | NA | 4 | 2 | 15 | 7 | 51 | 61 | 31 | 41 | -1,9 | 0,6 | -2,5 |
| chr2:70053596-70054096:downstream_1643 | chr2 | 69969126 | 70053596 | + | ANXA4 | NM | 136 | -2,3 | 1 | 2 | 1,45E-02 | 1,59E-01 | 2 | 2 | 11 | 8 | 253 | 268 | 228 | 316 | -2,4 | -0,1 | -2,4 |
| chr13:31032378-31032878:downstream_10693 | chr13 | 31032878 | 31040081 | - | HMGB1 | NM | 30 | -2,3 | 1 | 2 | 3,12E-02 | 2,19E-01 | 5 | 0 | 6 | 11 | 63 | 71 | 47 | 39 | -1,7 | 0,6 | -2,4 |
| chr6:111590261-111590761:downstream_5301 | chr6 | 111580481 | 111590261 | + | KIAA1919 | NM | 17 | -2,3 | 1 | 2 | 3,87E-02 | NA | 13 | 6 | 32 | 16 | 20 | 27 | 10 | 14 | -1,3 | 1,0 | -2,3 |
| chr9:113635553-113636053:downstream_26386 | chr9 | 113636053 | 113800365 | - | LPAR1 | NM | 36 | -2,2 | 1 | 2 | 2,22E-02 | 1,91E-01 | 4 | 1 | 77 | 57 | 11 | 12 | 63 | 61 | -4,7 | -2,4 | -2,3 |
| chr5:74017113-74017613:downstream_24160 | chr5 | 73935847 | 74017113 | + | HEXB | NM | 187 | -2,2 | 1 | 3 | 9,53E-03 | 1,31E-01 | 3 | 2 | 14 | 11 | 298 | 437 | 344 | 391 | -2,2 | 0,0 | -2,2 |
| chr7:134226166-134226666:downstream_5466 | chr7 | 134212343 | 134226166 | + | AKR1B10 | NM | 1513 | -2,1 | 1 | 4 | 4,20E-04 | 2,20E-02 | 7 | 4 | 35 | 33 | 2627 | 2532 | 3280 | 3589 | -2,6 | -0,4 | -2,2 |
| chr3:149221181-149221681:downstream_3226 | chr3 | 149192367 | 149221181 | + | TM4SF4 | NM | 446 | -2,1 | 1 | 2 | 2,72E-02 | 2,09E-01 | 3 | 0 | 6 | 7 | 951 | 1011 | 731 | 861 | -1,9 | 0,3 | -2,2 |
| chr1:159887396-159887896:downstream_17309 | chr1 | 159887896 | 159895332 | - | TAGLN2 | NM | 108 | -2,1 | 1 | 2 | 3,22E-02 | 2,22E-01 | 3 | 2 | 11 | 2 | 261 | 309 | 107 | 173 | -1,3 | 1,0 | -2,3 |
| chr1:53517289-53517789:downstream_17216 | chr1 | 53392900 | 53517289 | + | SCP2 | NM | 72 | -2,1 | 1 | 2 | 2,84E-02 | 2,09E-01 | 2 | 2 | 10 | 7 | 132 | 159 | 139 | 122 | -2,0 | 0,2 | -2,1 |
| chr9:75785310-75785810:downstream_7139 | chr9 | 75766646 | 75785310 | + | ANXA1 | NM | 753 | -2,1 | 1 | 3 | 9,48E-03 | 1,31E-01 | 4 | 1 | 18 | 11 | 1367 | 1301 | 1532 | 1794 | -2,5 | -0,3 | -2,2 |
| chr6:86205509-86206009:downstream_25042 | chr6 | 86159301 | 86205509 | + | NT5E | NM | 107 | -2,0 | 1 | 2 | 4,69E-02 | 2,60E-01 | 3 | 0 | 11 | 7 | 229 | 137 | 205 | 263 | -2,5 | -0,4 | -2,2 |
| chrX:37697588-37698088:downstream_7663 | chrX | 37698088 | 37706889 | - | DYNLT3 | NM | 53 | -2,0 | 1 | 2 | 3,12E-02 | 2,19E-01 | 4 | 4 | 7 | 7 | 98 | 175 | 57 | 67 | -0,9 | 1,1 | -2,0 |
| chr17:48769063-48769563:downstream_20308 | chr17 | 48712217 | 48769063 | + | ABCC3 | NM | 284 | -2,0 | 1 | 3 | 8,96E-03 | 1,26E-01 | 3 | 4 | 18 | 14 | 450 | 590 | 569 | 621 | -2,2 | -0,2 | -2,0 |
| chr4:15850706-15851206:downstream_3630 | chr4 | 15779930 | 15850706 | + | CD38 | NM | 32 | -1,9 | 1 | 2 | 3,51E-02 | 2,29E-01 | 7 | 3 | 11 | 21 | 53 | 60 | 47 | 52 | -1,7 | 0,2 | -1,9 |
| chr2:188328457-188328957:downstream_22273 | chr2 | 188328957 | 188419219 | - | TFPI | NM | 173 | -1,9 | 1 | 2 | 4,26E-02 | 2,47E-01 | 2 | 1 | 10 | 7 | 305 | 326 | 348 | 387 | -2,3 | -0,2 | -2,0 |
| chr2:211543832-211544332:downstream_21847 | chr2 | 211342405 | 211543832 | + | CPS1 | NM | 453 | -1,9 | 1 | 2 | 4,22E-02 | 2,47E-01 | 2 | 1 | 8 | 28 | 382 | 525 | 1266 | 1407 | -3,4 | -1,6 | -1,9 |
| chr3:155655520-155656020:downstream_3247 | chr3 | 155588324 | 155655520 | + | GMPS | NM | 38 | -1,9 | 1 | 2 | 4,60E-02 | 2,58E-01 | 26 | 8 | 127 | 93 | 10 | 8 | 19 | 12 | -2,7 | -0,9 | -1,8 |
| chr11:61566596-61567096:downstream_9381 | chr11 | 61567096 | 61584529 | - | FADS1 | NM | 261 | -1,8 | 1 | 3 | 4,45E-03 | 9,24E-02 | 7 | 6 | 50 | 30 | 380 | 367 | 580 | 669 | -2,6 | -0,7 | -1,9 |
| chr8:30535079-30535579:downstream_25956 | chr8 | 30535579 | 30585486 | - | GSR | NM | 71 | -1,8 | 1 | 2 | 2,05E-02 | 1,85E-01 | 5 | 5 | 33 | 26 | 99 | 81 | 189 | 131 | -2,6 | -0,8 | -1,8 |
| chrX:77081361-77081861:downstream_7815 | chrX | 77081861 | 77151065 | - | MAGT1 | NM | 72 | -1,8 | 1 | 2 | 4,22E-02 | 2,47E-01 | 8 | 0 | 10 | 16 | 131 | 152 | 106 | 151 | -1,7 | 0,1 | -1,8 |
| chr1:109643234-109643734:downstream_870 | chr1 | 109642814 | 109643234 | + | SCARNA2 | NR | 654 | -1,7 | 1 | 2 | 1,86E-02 | 1,75E-01 | 16 | 2 | 6 | 11 | 2368 | 1721 | 508 | 602 | 0,2 | 1,9 | -1,7 |
| chr5:174955621-174956121:downstream_4591 | chr5 | 174905513 | 174955621 | + | SFXN1 | NM | 77 | -1,7 | 1 | 2 | 1,87E-02 | 1,75E-01 | 21 | 13 | 89 | 89 | 47 | 105 | 128 | 126 | -2,4 | -0,7 | -1,7 |
| chr20:3170511-3171011:downstream_14863 | chr20 | 3171011 | 3185295 | - | DDRGK1 | NM | 47 | -1,7 | 1 | 2 | 3,28E-02 | 2,24E-01 | 8 | 15 | 33 | 20 | 97 | 80 | 56 | 70 | -1,2 | 0,5 | -1,7 |
| chr22:43011399-43011899:downstream_15660 | chr22 | 43011250 | 43011399 | + | RNU12 | NR | 652 | -1,6 | 1 | 3 | 1,16E-02 | 1,43E-01 | 7 | 11 | 6 | 14 | 1856 | 1850 | 711 | 763 | -0,2 | 1,3 | -1,5 |
| chr8:8641498-8641998:downstream_6258 | chr8 | 8641998 | 8751131 | - | MFHAS1 | NM | 54 | -1,6 | 1 | 2 | 4,43E-02 | 2,52E-01 | 10 | 8 | 128 | 84 | 12 | 30 | 72 | 91 | -3,6 | -2,0 | -1,6 |
| chr10:17279592-17280092:downstream_8430 | chr10 | 17270257 | 17279592 | + | VIM | NM | 158 | -1,6 | 1 | 2 | 3,21E-02 | 2,22E-01 | 7 | 3 | 8 | 17 | 310 | 352 | 250 | 313 | -1,3 | 0,2 | -1,6 |
| chr8:17721399-17721899:downstream_25944 | chr8 | 17721899 | 17753047 | - | FGL1 | NM | 336 | -1,5 | 1 | 3 | 5,25E-03 | 9,95E-02 | 174 | 144 | 345 | 192 | 562 | 577 | 289 | 403 | -0,8 | 0,7 | -1,5 |
| chr10:5149878-5150378:downstream_17461 | chr10 | 5090957 | 5149878 | + | AKR1C3 | NM | 1831 | -1,4 | 0 | 3 | 1,11E-03 | 4,15E-02 | 48 | 44 | 277 | 235 | 2369 | 2190 | 4411 | 5076 | -2,5 | -1,1 | -1,4 |
| chr2:85133799-85134299:downstream_2071 | chr2 | 85132762 | 85133799 | + | TMSB10 | NM | 1544 | -1,4 | 1 | 2 | 1,63E-02 | 1,67E-01 | 34 | 9 | 38 | 34 | 3730 | 3888 | 2220 | 2404 | -0,7 | 0,7 | -1,5 |
| chr12:104341708-104342208:downstream_9911 | chr12 | 104324111 | 104341708 | + | HSP90B1 | NM | 704 | -1,4 | 1 | 2 | 1,39E-02 | 1,56E-01 | 13 | 12 | 36 | 28 | 1610 | 1208 | 1172 | 1553 | -1,4 | 0,0 | -1,4 |
| chr12:6994950-6995450:downstream_10004 | chr12 | 6993845 | 6994950 | + | DSTNP2 | NR | 240 | -1,4 | 1 | 2 | 1,63E-02 | 1,67E-01 | 17 | 15 | 29 | 27 | 485 | 622 | 314 | 413 | -0,8 | 0,6 | -1,4 |
| chrX:12995346-12995846:downstream_7601 | chrX | 12993225 | 12995346 | + | TMSB4X | NM | 845 | -1,3 | 1 | 2 | 1,69E-02 | 1,69E-01 | 18 | 14 | 26 | 27 | 1830 | 2111 | 1233 | 1504 | -0,8 | 0,5 | -1,3 |
| chr16:69742803-69743303:downstream_20153 | chr16 | 69743303 | 69760571 | - | NQO1 | NM | 460 | -1,3 | 1 | 2 | 2,07E-02 | 1,85E-01 | 16 | 10 | 36 | 29 | 870 | 920 | 856 | 943 | -1,3 | 0,0 | -1,3 |
| chr7:134126606-134127106:downstream_6098 | chr7 | 134127106 | 134143888 | - | AKR1B1 | NM | 168 | -1,2 | 1 | 2 | 4,64E-02 | 2,59E-01 | 11 | 15 | 21 | 21 | 342 | 410 | 227 | 297 | -0,7 | 0,5 | -1,2 |
| chr3:169481897-169482397:downstream_3267 | chr3 | 169482397 | 169482848 | - | TERC | NR | 440 | -1,2 | 1 | 2 | 3,95E-02 | 2,41E-01 | 20 | 11 | 19 | 16 | 1312 | 1009 | 603 | 526 | -0,2 | 1,0 | -1,3 |
| chr5:57749309-57749809:downstream_24303 | chr5 | 57749809 | 57755966 | - | PLK2 | NM | 401 | -1,2 | 1 | 2 | 2,83E-02 | 2,09E-01 | 26 | 36 | 19 | 37 | 1024 | 1165 | 490 | 411 | 0,1 | 1,3 | -1,1 |
| chr11:93463180-93463680:downstream_9583 | chr11 | 93463680 | 93463812 | - | SNORA25 | NR | 194 | -1,2 | 1 | 2 | 2,22E-02 | 1,91E-01 | 56 | 68 | 78 | 60 | 391 | 475 | 235 | 185 | -0,2 | 1,0 | -1,2 |
| chr2:217071016-217071516:downstream_1555 | chr2 | 216974019 | 217071016 | + | XRCC5 | NM | 104 | -1,2 | 1 | 2 | 3,95E-02 | 2,41E-01 | 31 | 46 | 118 | 135 | 102 | 104 | 127 | 169 | -1,7 | -0,5 | -1,2 |
| chr11:43878169-43878669:downstream_8873 | chr11 | 43702142 | 43878169 | + | HSD17B12 | NM | 297 | -1,1 | 1 | 2 | 3,59E-02 | 2,29E-01 | 20 | 27 | 61 | 49 | 517 | 542 | 619 | 542 | -1,2 | -0,1 | -1,1 |
| chr17:57918698-57919198:downstream_13277 | chr17 | 57918626 | 57918698 | + | MIR21 | NR | 1355 | -1,0 | 0 | 2 | 1,40E-02 | 1,56E-01 | 434 | 491 | 2093 | 1991 | 820 | 1005 | 1884 | 2121 | -2,1 | -1,1 | -1,0 |
| chr15:66791152-66791652:downstream_11773 | chr15 | 66791652 | 66797193 | - | RPL4 | NM | 685 | 0,9 | 0 | -2 | 3,36E-02 | 2,25E-01 | 414 | 374 | 114 | 125 | 1544 | 1292 | 810 | 805 | 1,7 | 0,8 | 0,9 |
| chr9:19375753-19376253:downstream_6993 | chr9 | 19376253 | 19380235 | - | RPS6 | NM | 778 | 1,1 | 0 | -2 | 2,83E-02 | 2,09E-01 | 181 | 113 | 43 | 41 | 1865 | 1799 | 1060 | 1117 | 1,8 | 0,8 | 1,1 |
| chr15:60638849-60639349:downstream_19461 | chr15 | 60639349 | 60690185 | - | ANXA2 | NM | 1520 | 1,2 | 1 | -2 | 3,84E-02 | 2,39E-01 | 41 | 23 | 10 | 15 | 3441 | 2979 | 2643 | 3006 | 1,4 | 0,2 | 1,2 |
| chr17:8280333-8280833:downstream_12830 | chr17 | 8280833 | 8286565 | - | RPL26 | NM | 559 | 1,2 | 1 | -2 | 4,37E-02 | 2,52E-01 | 49 | 44 | 8 | 8 | 1618 | 1485 | 566 | 692 | 2,5 | 1,3 | 1,2 |
| chr3:52742197-52742697:downstream_3003 | chr3 | 52739856 | 52742197 | + | SPCS1 | NM | 432 | 1,2 | 1 | -2 | 2,11E-02 | 1,87E-01 | 62 | 76 | 19 | 26 | 814 | 1035 | 686 | 737 | 1,6 | 0,4 | 1,2 |
| chr14:45584804-45585304:downstream_11125 | chr14 | 45553301 | 45584804 | + | PRPF39 | NM | 108 | 1,2 | 1 | -2 | 3,50E-02 | 2,29E-01 | 50 | 62 | 36 | 29 | 139 | 159 | 196 | 197 | 0,8 | -0,4 | 1,2 |
| chr12:131362223-131362723:downstream_18785 | chr12 | 131356538 | 131362223 | + | RAN | NM | 177 | 1,3 | 1 | -2 | 4,65E-02 | 2,59E-01 | 26 | 41 | 10 | 11 | 350 | 410 | 321 | 244 | 1,7 | 0,4 | 1,3 |
| chr11:17095438-17095938:downstream_9159 | chr11 | 17095938 | 17099220 | - | RPS13 | NM | 80 | 1,3 | 1 | -2 | 3,25E-02 | 2,23E-01 | 161 | 154 | 43 | 41 | 68 | 80 | 47 | 48 | 1,9 | 0,6 | 1,3 |
| chr12:46751470-46751970:downstream_18832 | chr12 | 46751970 | 46766645 | - | SLC38A2 | NM | 348 | 1,3 | 0 | -3 | 7,53E-03 | 1,18E-01 | 38 | 40 | 77 | 69 | 233 | 234 | 1072 | 1023 | -0,9 | -2,2 | 1,3 |
| chr3:98514313-98514813:downstream_2554 | chr3 | 98514813 | 98620533 | - | DCBLD2 | NM | 209 | 1,4 | 1 | -2 | 3,79E-02 | 2,38E-01 | 19 | 6 | 24 | 15 | 161 | 172 | 576 | 703 | -0,6 | -1,9 | 1,3 |
| chr20:2441780-2442280:downstream_22566 | chr20 | 2442280 | 2451499 | - | SNRPB | NM | 83 | 1,4 | 1 | -2 | 2,78E-02 | 2,09E-01 | 135 | 93 | 33 | 24 | 119 | 114 | 63 | 86 | 2,0 | 0,6 | 1,3 |
| chr22:29190047-29190547:downstream_22966 | chr22 | 29190547 | 29196560 | - | XBP1 | NM | 163 | 1,4 | 1 | -3 | 8,09E-03 | 1,21E-01 | 51 | 69 | 114 | 137 | 80 | 64 | 390 | 403 | -1,1 | -2,5 | 1,4 |
| chr11:10818092-10818592:downstream_18003 | chr11 | 10818592 | 10830582 | - | EIF4G2 | NM | 298 | 1,4 | 1 | -2 | 1,51E-02 | 1,64E-01 | 25 | 40 | 10 | 23 | 444 | 494 | 669 | 677 | 1,0 | -0,5 | 1,5 |
| chr17:41154971-41155471:downstream_13139 | chr17 | 41150445 | 41154971 | + | RPL27 | NM | 116 | 1,4 | 1 | -2 | 4,48E-02 | 2,54E-01 | 24 | 31 | 4 | 8 | 234 | 288 | 138 | 203 | 2,1 | 0,6 | 1,5 |
| chr19:36487220-36487720:downstream_14239 | chr19 | 36486089 | 36487220 | + | SDHAF1 | NM | 81 | 1,5 | 1 | -2 | 2,23E-02 | 1,91E-01 | 155 | 144 | 56 | 28 | 66 | 83 | 57 | 55 | 1,8 | 0,4 | 1,4 |
| chr14:52471420-52471920:downstream_11140 | chr14 | 52456227 | 52471420 | + | C14orf166 | NM | 289 | 1,5 | 1 | -3 | 1,01E-02 | 1,35E-01 | 37 | 36 | 8 | 15 | 625 | 529 | 519 | 542 | 1,7 | 0,1 | 1,5 |
| chr1:172410096-172410596:downstream_17081 | chr1 | 172410596 | 172413230 | - | PIGC | NM | 52 | 1,5 | 1 | -2 | 4,56E-02 | 2,57E-01 | 33 | 32 | 15 | 17 | 46 | 87 | 83 | 103 | 1,0 | -0,5 | 1,5 |
| chr9:74297781-74298281:downstream_26538 | chr9 | 74298281 | 74383800 | - | TMEM2 | NM | 84 | 1,5 | 1 | -2 | 3,86E-02 | 2,39E-01 | 8 | 11 | 13 | 17 | 46 | 67 | 276 | 236 | -0,7 | -2,2 | 1,5 |
| chr6:26124918-26125418:downstream_4912 | chr6 | 26124372 | 26124918 | + | HIST1H2AC | NM | 274 | 1,5 | 1 | -3 | 3,20E-03 | 7,50E-02 | 63 | 51 | 79 | 60 | 252 | 186 | 781 | 719 | -0,3 | -1,8 | 1,5 |
| chr20:5524579-5525079:downstream_14878 | chr20 | 5525079 | 5591672 | - | GPCPD1 | NM | 48 | 1,5 | 1 | -2 | 3,35E-02 | 2,25E-01 | 20 | 26 | 53 | 43 | 19 | 16 | 103 | 103 | -1,0 | -2,6 | 1,5 |
| chr11:2418649-2419149:downstream_17941 | chr11 | 2397406 | 2418649 | + | CD81 | NM | 62 | 1,5 | 1 | -2 | 4,86E-02 | 2,69E-01 | 20 | 26 | 4 | 7 | 132 | 119 | 85 | 107 | 2,0 | 0,4 | 1,6 |
| chr6:72112753-72113253:downstream_5222 | chr6 | 72113253 | 72113324 | - | MIR30A | NR | 42 | 1,5 | 1 | -2 | 4,25E-02 | 2,47E-01 | 33 | 23 | 60 | 61 | 9 | 14 | 70 | 70 | -1,1 | -2,6 | 1,5 |
| chr5:180663427-180663927:downstream_4639 | chr5 | 180663927 | 180670906 | - | GNB2L1 | NM | 427 | 1,6 | 1 | -3 | 2,87E-03 | 7,20E-02 | 53 | 80 | 22 | 25 | 725 | 851 | 770 | 886 | 1,5 | -0,1 | 1,6 |
| chr3:52728510-52729010:downstream_23172 | chr3 | 52719935 | 52728510 | + | GNL3 | NM | 60 | 1,6 | 1 | -2 | 3,17E-02 | 2,21E-01 | 46 | 41 | 8 | 13 | 127 | 83 | 65 | 94 | 2,0 | 0,4 | 1,6 |
| chr14:23240998-23241498:downstream_11053 | chr14 | 23235730 | 23240998 | + | OXA1L | NM | 115 | 1,6 | 1 | -2 | 2,20E-02 | 1,91E-01 | 31 | 22 | 7 | 7 | 221 | 265 | 175 | 191 | 2,0 | 0,4 | 1,6 |
| chr1:235272157-235272657:downstream_311 | chr1 | 235272657 | 235292256 | - | TOMM20 | NM | 184 | 1,6 | 1 | -2 | 1,55E-02 | 1,64E-01 | 14 | 25 | 7 | 9 | 285 | 331 | 424 | 373 | 1,3 | -0,4 | 1,7 |
| chr11:10328949-10329449:downstream_9141 | chr11 | 10326526 | 10328949 | + | ADM | NM | 104 | 1,6 | 1 | -2 | 2,70E-02 | 2,09E-01 | 20 | 16 | 14 | 3 | 177 | 161 | 206 | 237 | 1,1 | -0,4 | 1,5 |
| chr12:27909237-27909737:downstream_18690 | chr12 | 27863705 | 27909237 | + | MRPS35 | NM | 47 | 1,6 | 1 | -2 | 4,13E-02 | 2,46E-01 | 20 | 36 | 14 | 12 | 69 | 53 | 61 | 113 | 1,1 | -0,5 | 1,6 |
| chr6:24704589-24705089:downstream_4893 | chr6 | 24705089 | 24719403 | - | C6orf62 | NM | 48 | 1,6 | 1 | -2 | 2,68E-02 | 2,09E-01 | 63 | 44 | 18 | 11 | 67 | 66 | 49 | 62 | 1,8 | 0,3 | 1,6 |
| chr4:100868743-100869243:downstream_3794 | chr4 | 100869243 | 100871512 | - | H2AFZ | NM | 103 | 1,6 | 1 | -2 | 1,73E-02 | 1,69E-01 | 20 | 20 | 10 | 9 | 180 | 135 | 192 | 254 | 1,1 | -0,5 | 1,6 |
| chr17:26673153-26673653:downstream_20644 | chr17 | 26673653 | 26684612 | - | POLDIP2 | NM | 44 | 1,6 | 1 | -2 | 3,76E-02 | 2,37E-01 | 64 | 70 | 17 | 7 | 55 | 72 | 40 | 28 | 2,5 | 0,9 | 1,6 |
| chr10:75008100-75008600:downstream_8567 | chr10 | 75008600 | 75012451 | - | MRPS16 | NM | 98 | 1,7 | 1 | -3 | 3,24E-03 | 7,50E-02 | 157 | 116 | 49 | 50 | 91 | 102 | 104 | 116 | 1,5 | -0,2 | 1,7 |
| chr6:26205249-26205749:downstream_4909 | chr6 | 26204872 | 26205249 | + | HIST1H4E | NM | 205 | 1,7 | 1 | -3 | 3,32E-03 | 7,50E-02 | 37 | 37 | 32 | 49 | 198 | 130 | 518 | 637 | -0,1 | -1,8 | 1,7 |
| chr4:6708928-6709428:downstream_3595 | chr4 | 6709428 | 6711606 | - | MRFAP1L1 | NM | 93 | 1,7 | 1 | -2 | 1,92E-02 | 1,78E-01 | 28 | 22 | 7 | 7 | 194 | 172 | 178 | 132 | 1,9 | 0,2 | 1,7 |
| chr15:55473011-55473511:downstream_11730 | chr15 | 55473511 | 55489231 | - | RSL24D1 | NM | 111 | 1,7 | 1 | -3 | 6,15E-03 | 1,06E-01 | 59 | 60 | 10 | 21 | 207 | 187 | 167 | 179 | 1,9 | 0,2 | 1,7 |
| chr6:26284853-26285353:downstream_4922 | chr6 | 26285353 | 26285727 | - | HIST1H4H | NM | 79 | 1,7 | 1 | -3 | 8,66E-03 | 1,24E-01 | 54 | 41 | 24 | 39 | 75 | 75 | 132 | 191 | 0,6 | -1,1 | 1,7 |
| chr17:65740324-65740824:downstream_20612 | chr17 | 65713948 | 65740324 | + | NOL11 | NM | 80 | 1,7 | 1 | -2 | 4,12E-02 | 2,46E-01 | 8 | 14 | 3 | 6 | 108 | 148 | 199 | 154 | 1,3 | -0,5 | 1,8 |
| chr2:198350807-198351307:downstream_21971 | chr2 | 198351307 | 198364998 | - | HSPD1 | NM | 68 | 1,7 | 1 | -3 | 7,60E-03 | 1,18E-01 | 40 | 38 | 19 | 18 | 81 | 88 | 132 | 131 | 1,0 | -0,6 | 1,7 |
| chr2:24299802-24300302:downstream_22293 | chr2 | 24300302 | 24308085 | - | TP53I3 | NM | 78 | 1,7 | 1 | -3 | 1,07E-02 | 1,39E-01 | 69 | 81 | 7 | 11 | 163 | 157 | 68 | 66 | 3,0 | 1,3 | 1,8 |
| chr2:38970240-38970740:downstream_22253 | chr2 | 38970740 | 38978636 | - | SRSF7 | NM | 52 | 1,7 | 1 | -2 | 3,12E-02 | 2,19E-01 | 18 | 18 | 7 | 5 | 91 | 87 | 107 | 81 | 1,6 | -0,1 | 1,7 |
| chr14:69846339-69846839:downstream_10956 | chr14 | 69846839 | 69865021 | - | ERH | NM | 137 | 1,7 | 1 | -2 | 1,85E-02 | 1,75E-01 | 13 | 11 | 7 | 5 | 235 | 172 | 324 | 332 | 1,0 | -0,7 | 1,7 |
| chr9:136218280-136218780:downstream_7378 | chr9 | 136215068 | 136218280 | + | RPL7A | NM | 27 | 1,7 | 1 | -2 | 4,36E-02 | NA | 47 | 37 | 11 | 9 | 32 | 33 | 22 | 29 | 2,1 | 0,3 | 1,7 |
| chr7:106842974-106843474:downstream_25552 | chr7 | 106809405 | 106842974 | + | HBP1 | NM | 51 | 1,8 | 1 | -2 | 4,02E-02 | 2,44E-01 | 7 | 15 | 8 | 8 | 35 | 67 | 159 | 107 | 0,4 | -1,4 | 1,8 |
| chr13:26786404-26786904:downstream_19080 | chr13 | 26786904 | 26796508 | - | RNF6 | NM | 66 | 1,8 | 1 | -3 | 1,14E-02 | 1,43E-01 | 123 | 80 | 21 | 16 | 92 | 84 | 38 | 71 | 2,4 | 0,7 | 1,8 |
| chr3:139075887-139076387:downstream_2720 | chr3 | 139062860 | 139075887 | + | MRPS22 | NM | 25 | 1,8 | 1 | -2 | 4,65E-02 | NA | 16 | 14 | 10 | 11 | 23 | 21 | 67 | 41 | 0,5 | -1,3 | 1,8 |
| chr16:3072383-3072883:downstream_12139 | chr16 | 3070312 | 3072383 | + | TNFRSF12A | NM | 318 | 1,8 | 1 | -3 | 6,63E-04 | 2,87E-02 | 189 | 264 | 17 | 16 | 851 | 793 | 218 | 198 | 3,8 | 2,0 | 1,8 |
| chr5:7901235-7901735:downstream_24237 | chr5 | 7869216 | 7901235 | + | MTRR | NM | 39 | 1,8 | 1 | -2 | 3,57E-02 | 2,29E-01 | 17 | 19 | 3 | 8 | 77 | 46 | 67 | 78 | 1,7 | -0,2 | 2,0 |

| | | | | | | | | | | | | | | | | | | | | | | | |
|---|---|---|---|---|---|---|---|---|---|---|---|---|---|---|---|---|---|---|---|---|---|---|---|
| chr11:62557286-62557786:downstream_9404 | chr11 | 62557786 | 62559486 | - | TMEM223 | NM | 69 | 1,9 | 1 | -3 | 1,00E-02 | 1,35E-01 | 50 | 77 | 45 | 22 | 81 | 44 | 102 | 130 | 0,9 | -0,9 | 1,8 |
| chr19:34960798-34961298:downstream_14203 | chr19 | 34919267 | 34960798 | + | UBA2 | NM | 45 | 1,9 | 1 | -2 | 2,94E-02 | 2,13E-01 | 7 | 15 | 4 | 8 | 46 | 56 | 106 | 118 | 0,8 | -1,1 | 2,0 |
| chr22:29185289-29185789:downstream_22766 | chr22 | 29168661 | 29185289 | + | CCDC117 | NM | 22 | 1,9 | 1 | -2 | 4,48E-02 | NA | 24 | 29 | 13 | 12 | 24 | 13 | 35 | 28 | 1,1 | -0,8 | 1,9 |
| chr6:27806888-27807388:downstream_4959 | chr6 | 27806439 | 27806888 | + | HIST1H2BN | NM | 45 | 1,9 | 1 | -2 | 2,34E-02 | 1,93E-01 | 11 | 10 | 18 | 14 | 33 | 14 | 132 | 130 | -0,6 | -2,5 | 1,9 |
| chr17:4636905-4637405:downstream_20572 | chr17 | 4634702 | 4636905 | + | MED11 | NM | 54 | 1,9 | 1 | -2 | 2,81E-02 | 2,09E-01 | 12 | 6 | 6 | 5 | 57 | 74 | 148 | 127 | 0,8 | -1,1 | 1,9 |
| chr10:70102953-70103453:downstream_17598 | chr10 | 70091767 | 70102953 | + | HNRNPH3 | NM | 47 | 1,9 | 1 | -2 | 1,93E-02 | 1,78E-01 | 17 | 15 | 4 | 7 | 73 | 66 | 95 | 98 | 1,5 | -0,5 | 2,0 |
| chr16:57505369-57505869:downstream_12384 | chr16 | 57505869 | 57520385 | - | DOK4 | NM | 68 | 1,9 | 1 | -2 | 1,95E-02 | 1,79E-01 | 13 | 62 | 28 | 33 | 52 | 47 | 150 | 155 | 0,3 | -1,6 | 1,9 |
| chr17:56048409-56048909:downstream_13259 | chr17 | 56048909 | 56065615 | - | VEZF1 | NM | 35 | 1,9 | 1 | -2 | 3,80E-02 | 2,38E-01 | 9 | 7 | 6 | 7 | 28 | 37 | 114 | 76 | 0,4 | -1,5 | 1,9 |
| chr10:104238485-104238985:downstream_8697 | chr10 | 104238985 | 104262512 | - | ACTR1A | NM | 43 | 1,9 | 1 | -2 | 4,93E-02 | 2,72E-01 | 10 | 6 | 3 | 3 | 76 | 52 | 89 | 103 | 1,4 | -0,6 | 1,9 |
| chr4:6644470-6644970:downstream_23760 | chr4 | 6641817 | 6644470 | + | MRFAP1 | NM | 206 | 1,9 | 1 | -2 | 3,71E-02 | 2,36E-01 | 10 | 11 | 3 | 2 | 397 | 517 | 342 | 367 | 2,2 | 0,4 | 1,8 |
| chr3:190022989-190023489:downstream_3326 | chr3 | 190023489 | 190040235 | - | CLDN1 | NM | 26 | 1,9 | 1 | -2 | 4,54E-02 | NA | 30 | 14 | 11 | 5 | 27 | 36 | 39 | 44 | 1,4 | -0,4 | 1,9 |
| chr1:109780804-109781304:downstream_873 | chr1 | 109756514 | 109780804 | + | SARS | NM | 76 | 1,9 | 1 | -2 | 1,85E-02 | 1,75E-01 | 23 | 25 | 4 | 3 | 152 | 199 | 93 | 111 | 2,7 | 0,8 | 1,9 |
| chr11:82896835-82897335:downstream_9555 | chr11 | 82868136 | 82896835 | + | PCF11 | NM | 40 | 1,9 | 1 | -2 | 3,46E-02 | 2,29E-01 | 13 | 11 | 3 | 4 | 63 | 73 | 79 | 78 | 1,8 | -0,2 | 2,0 |
| chr10:70744825-70745325:downstream_17539 | chr10 | 70715878 | 70744825 | + | DDX21 | NM | 55 | 2,0 | 1 | -2 | 1,74E-02 | 1,69E-01 | 21 | 24 | 6 | 5 | 98 | 106 | 67 | 113 | 2,1 | 0,2 | 1,9 |
| chr19:36038429-36038929:downstream_21556 | chr19 | 36036501 | 36038429 | + | TMEM147 | NM | 103 | 2,0 | 1 | -2 | 1,63E-02 | 1,67E-01 | 16 | 31 | 6 | 2 | 254 | 221 | 143 | 148 | 2,6 | 0,7 | 1,9 |
| chr6:160177352-160177852:downstream_25354 | chr6 | 160148029 | 160177352 | + | WTAP | NM | 24 | 2,0 | 1 | -2 | 4,84E-02 | NA | 15 | 15 | 4 | 3 | 41 | 41 | 40 | 35 | 2,1 | 0,1 | 1,9 |
| chrX:154468122-154468622:downstream_27102 | chrX | 154425283 | 154468122 | + | VBP1 | NM | 165 | 2,0 | 1 | -2 | 1,54E-02 | 1,64E-01 | 15 | 16 | 3 | 3 | 313 | 405 | 282 | 280 | 2,4 | 0,4 | 2,0 |
| chr6:26188437-26188937:downstream_4917 | chr6 | 26188937 | 26189304 | - | HIST1H4D | NM | 63 | 2,0 | 1 | -2 | 3,50E-02 | 2,29E-01 | 5 | 4 | 7 | 6 | 52 | 20 | 192 | 216 | -0,5 | -2,5 | 2,0 |
| chr12:122691708-122692208:downstream_18543 | chr12 | 122692208 | 122712081 | - | DIABLO | NM | 32 | 2,0 | 1 | -2 | 3,99E-02 | 2,43E-01 | 15 | 30 | 3 | 8 | 67 | 33 | 47 | 56 | 2,0 | 0,0 | 2,1 |
| chr1:202909452-202909952:downstream_16450 | chr1 | 202909952 | 202927700 | - | ADIPOR1 | NM | 115 | 2,0 | 1 | -2 | 2,63E-02 | 2,08E-01 | 13 | 19 | 1 | 3 | 263 | 277 | 202 | 138 | 2,8 | 0,7 | 2,1 |
| chr1:44439043-44439543:downstream_16667 | chr1 | 44435652 | 44439043 | + | DPH2 | NM | 48 | 2,0 | 1 | -3 | 1,24E-02 | 1,46E-01 | 69 | 36 | 10 | 7 | 82 | 80 | 50 | 48 | 2,7 | 0,7 | 2,0 |
| chr1:173457946-173458446:downstream_1228 | chr1 | 173446485 | 173457946 | + | PRDX6 | NM | 238 | 2,0 | 1 | -3 | 2,43E-03 | 6,58E-02 | 33 | 21 | 8 | 6 | 490 | 430 | 508 | 409 | 2,0 | 0,0 | 1,9 |
| chr17:79215098-79215598:downstream_13421 | chr17 | 79213110 | 79215098 | + | C17orf89 | NM | 108 | 2,0 | 1 | -3 | 1,10E-03 | 4,15E-02 | 85 | 77 | 8 | 15 | 217 | 205 | 134 | 121 | 2,8 | 0,7 | 2,1 |
| chr22:38878942-38879442:downstream_22789 | chr22 | 38879442 | 38902345 | - | DDX17 | NM | 53 | 2,0 | 1 | -3 | 1,12E-02 | 1,42E-01 | 15 | 30 | 10 | 8 | 67 | 73 | 85 | 137 | 1,3 | -0,7 | 2,0 |
| chr17:4848517-4849017:downstream_12767 | chr17 | 4843629 | 4848517 | + | RNF167 | NM | 49 | 2,0 | 1 | -2 | 2,89E-02 | 2,11E-01 | 12 | 30 | 3 | 4 | 93 | 109 | 64 | 75 | 2,6 | 0,5 | 2,1 |
| chr12:99006681-99007181:downstream_18605 | chr12 | 99007181 | 99038829 | - | IKBIP | NM | 217 | 2,0 | 1 | -2 | 1,63E-02 | 1,67E-01 | 19 | 14 | 1 | 3 | 512 | 523 | 333 | 331 | 2,8 | 0,6 | 2,1 |
| chr14:70238722-70239222:downstream_19405 | chr14 | 70233828 | 70238722 | + | SRSF5 | NM | 30 | 2,0 | 1 | -2 | 3,39E-02 | 2,26E-01 | 24 | 41 | 6 | 7 | 64 | 28 | 39 | 34 | 2,4 | 0,3 | 2,1 |
| chr19:41767672-41768172:downstream_21018 | chr19 | 41725103 | 41767672 | + | AXL | NM | 32 | 2,0 | 1 | -2 | 1,72E-02 | 1,69E-01 | 57 | 61 | 8 | 7 | 41 | 42 | 18 | 24 | 3,0 | 1,0 | 2,0 |
| chr20:5079983-5080483:downstream_22584 | chr20 | 5080483 | 5093733 | - | TMEM230 | NM | 53 | 2,0 | 1 | -2 | 2,00E-02 | 1,83E-01 | 27 | 15 | 10 | 2 | 98 | 89 | 82 | 103 | 1,9 | 0,0 | 1,8 |
| chr17:15932723-15933223:downstream_20795 | chr17 | 15902693 | 15932723 | + | TTC19 | NM | 21 | 2,0 | 1 | -2 | 3,63E-02 | NA | 20 | 39 | 19 | 19 | 9 | 10 | 21 | 29 | 0,6 | -1,4 | 2,0 |
| chr12:93801587-93802087:downstream_10356 | chr12 | 93802087 | 93836026 | - | UBE2N | NM | 20 | 2,0 | 1 | -2 | 4,53E-02 | NA | 15 | 15 | 4 | 13 | 14 | 17 | 36 | 42 | 0,8 | -1,3 | 2,1 |
| chr12:49046494-49046994:downstream_10143 | chr12 | 49046994 | 49076035 | - | KANSL2 | NM | 26 | 2,1 | 1 | -2 | 4,89E-02 | NA | 63 | 52 | 8 | 9 | 39 | 9 | 13 | 17 | 2,7 | 0,7 | 2,1 |
| chr2:29171086-29171586:downstream_1923 | chr2 | 29117508 | 29171086 | + | WDR43 | NM | 79 | 2,1 | 1 | -2 | 2,95E-02 | 2,13E-01 | 16 | 3 | 1 | 5 | 146 | 98 | 171 | 191 | 1,6 | -0,6 | 2,2 |
| chr1:71528473-71528973:downstream_17442 | chr1 | 71528973 | 71546972 | - | ZRANB2 | NM | 29 | 2,1 | 1 | -2 | 2,83E-02 | 2,09E-01 | 32 | 32 | 13 | 3 | 30 | 46 | 32 | 41 | 2,0 | 0,1 | 1,9 |
| chr11:57470686-57471186:downstream_9323 | chr11 | 57471186 | 57479673 | - | MED19 | NM | 30 | 2,1 | 1 | -2 | 2,32E-02 | 1,92E-01 | 31 | 45 | 3 | 9 | 42 | 45 | 32 | 30 | 2,7 | 0,5 | 2,2 |
| chr2:176040485-176040985:downstream_21760 | chr2 | 176040985 | 176046490 | - | ATP5G3 | NM | 25 | 2,1 | 1 | -2 | 3,11E-02 | NA | 47 | 57 | 11 | 11 | 28 | 12 | 22 | 13 | 2,3 | 0,1 | 2,1 |
| chr7:42948371-42948871:downstream_25412 | chr7 | 42948871 | 42951689 | - | C7orf25 | NM | 49 | 2,1 | 1 | -2 | 2,90E-02 | 2,11E-01 | 28 | 14 | 10 | 1 | 103 | 73 | 90 | 70 | 2,0 | 0,1 | 1,9 |
| chr12:76418726-76419226:downstream_10326 | chr12 | 76419226 | 76425556 | - | PHLDA1 | NM | 55 | 2,1 | 1 | -2 | 1,53E-02 | 1,64E-01 | 27 | 31 | 3 | 4 | 105 | 140 | 68 | 61 | 3,1 | 0,9 | 2,1 |
| chr12:6979599-6980099:downstream_18850 | chr12 | 6980099 | 6982521 | - | SPSB2 | NM | 71 | 2,1 | 1 | -3 | 1,07E-02 | 1,39E-01 | 94 | 157 | 114 | 98 | 3 | 20 | 35 | 48 | 0,2 | -1,8 | 2,0 |
| chr11:59573107-59573607:downstream_9349 | chr11 | 59573607 | 59578345 | - | MRPL16 | NM | 73 | 2,1 | 1 | -2 | 3,56E-02 | 2,29E-01 | 3 | 14 | 3 | 2 | 97 | 142 | 174 | 148 | 1,7 | -0,4 | 2,1 |
| chr2:85772403-85772903:downstream_2076 | chr2 | 85766100 | 85772403 | + | MAT2A | NM | 65 | 2,1 | 1 | -3 | 3,31E-03 | 7,50E-02 | 50 | 37 | 7 | 13 | 112 | 93 | 118 | 94 | 2,1 | 0,0 | 2,2 |
| chr5:72847524-72848024:downstream_4298 | chr5 | 72848024 | 72861511 | - | ANKRA2 | NM | 33 | 2,1 | 1 | -2 | 4,52E-02 | 2,56E-01 | 76 | 75 | 3 | 4 | 35 | 55 | 15 | 4 | 4,5 | 2,2 | 2,2 |
| chr8:42263455-42263955:downstream_26175 | chr8 | 42249278 | 42263455 | + | VDAC3 | NM | 77 | 2,1 | 1 | -3 | 2,54E-03 | 6,76E-02 | 37 | 28 | 10 | 7 | 136 | 117 | 128 | 150 | 1,9 | -0,1 | 2,1 |
| chr11:94232744-94233244:downstream_9589 | chr11 | 94227152 | 94232744 | + | ANKRD49 | NM | 30 | 2,1 | 1 | -2 | 3,36E-02 | 2,25E-01 | 8 | 15 | 3 | 4 | 50 | 41 | 63 | 57 | 1,8 | -0,4 | 2,2 |
| chr1:234519795-234520295:downstream_16614 | chr1 | 234509182 | 234519795 | + | COA6 | NM | 54 | 2,1 | 1 | -2 | 2,42E-02 | 1,94E-01 | 21 | 9 | 4 | 2 | 120 | 87 | 90 | 97 | 2,2 | 0,1 | 2,0 |
| chr16:75661121-75661621:downstream_20038 | chr16 | 75661621 | 75681585 | - | KARS | NM | 172 | 2,1 | 1 | -4 | 3,37E-04 | 1,96E-02 | 99 | 66 | 18 | 11 | 338 | 337 | 273 | 235 | 2,5 | 0,4 | 2,1 |
| chr9:127639696-127640196:downstream_7314 | chr9 | 127631483 | 127639696 | + | ARPC5L | NM | 38 | 2,1 | 1 | -2 | 4,32E-02 | 2,50E-01 | 11 | 16 | 1 | 2 | 82 | 83 | 50 | 59 | 2,8 | 0,6 | 2,2 |
| chr20:60963576-60964076:downstream_15131 | chr20 | 60962120 | 60963576 | + | RPS21 | NM | 425 | 2,1 | 1 | -4 | 3,70E-04 | 2,00E-02 | 55 | 52 | 11 | 6 | 947 | 1035 | 568 | 726 | 2,7 | 0,6 | 2,1 |
| chr2:198339851-198340351:downstream_2365 | chr2 | 198318230 | 198339851 | + | COQ10B | NM | 44 | 2,2 | 1 | -2 | 2,05E-02 | 1,85E-01 | 6 | 21 | 10 | 4 | 49 | 49 | 89 | 125 | 1,0 | -1,1 | 2,1 |
| chr14:102546574-102547074:downstream_19255 | chr14 | 102547074 | 102606086 | - | HSP90AA1 | NM | 70 | 2,2 | 1 | -2 | 3,04E-02 | 2,18E-01 | 10 | 16 | 3 | 2 | 158 | 149 | 83 | 135 | 2,6 | 0,5 | 2,1 |
| chr3:88100599-88101099:downstream_23073 | chr3 | 88101099 | 88199016 | - | CGGBP1 | NM | 29 | 2,2 | 1 | -2 | 3,92E-02 | 2,41E-01 | 5 | 14 | 1 | 5 | 38 | 39 | 71 | 61 | 1,5 | -0,8 | 2,3 |
| chr19:35233774-35234274:downstream_21612 | chr19 | 35225479 | 35233774 | + | ZNF181 | NM | 33 | 2,2 | 1 | -2 | 1,28E-02 | 1,48E-01 | 45 | 38 | 10 | 15 | 43 | 22 | 57 | 33 | 1,8 | -0,5 | 2,2 |
| chr15:55608881-55609381:downstream_19666 | chr15 | 55609381 | 55611392 | - | PIGBOS1 | NM | 47 | 2,2 | 1 | -3 | 7,64E-03 | 1,18E-01 | 53 | 133 | 43 | 40 | 20 | 15 | 45 | 29 | 1,2 | -1,0 | 2,2 |
| chr16:57274387-57274887:downstream_20201 | chr16 | 57220195 | 57274387 | + | RSPRY1 | NM | 18 | 2,2 | 1 | -2 | 4,45E-02 | NA | 19 | 13 | 7 | 9 | 21 | 8 | 32 | 35 | 1,0 | -1,2 | 2,2 |
| chr14:23947402-23947902:downstream_19320 | chr14 | 23938897 | 23947402 | + | NGDN | NM | 35 | 2,2 | 1 | -2 | 1,71E-02 | 1,69E-01 | 28 | 18 | 4 | 4 | 63 | 60 | 50 | 48 | 2,5 | 0,3 | 2,2 |
| chr3:41281939-41282439:downstream_23103 | chr3 | 41240941 | 41281939 | + | CTNNB1 | NM | 106 | 2,2 | 1 | -3 | 4,10E-03 | 8,70E-02 | 12 | 38 | 6 | 12 | 148 | 140 | 228 | 267 | 1,5 | -0,8 | 2,3 |
| chr1:19586622-19587122:downstream_510 | chr1 | 19578074 | 19586622 | + | MRTO4 | NM | 34 | 2,2 | 1 | -3 | 1,08E-02 | 1,39E-01 | 28 | 26 | 4 | 7 | 51 | 53 | 47 | 58 | 2,2 | 0,0 | 2,2 |
| chr11:77326695-77327195:downstream_8926 | chr11 | 77327195 | 77348851 | - | CLNS1A | NM | 175 | 2,2 | 1 | -4 | 3,41E-04 | 1,96E-02 | 41 | 42 | 10 | 12 | 246 | 332 | 394 | 323 | 1,9 | -0,3 | 2,2 |
| chr7:99174076-99174576:downstream_25826 | chr7 | 99156044 | 99174076 | + | ZNF655 | NM | 20 | 2,2 | 1 | -2 | 4,51E-02 | NA | 13 | 16 | 3 | 3 | 34 | 28 | 31 | 29 | 2,3 | 0,1 | 2,2 |
| chr1:207974696-207975196:downstream_1345 | chr1 | 207975196 | 207975284 | - | MIR29C | NR | 95 | 2,2 | 1 | -3 | 1,48E-03 | 4,77E-02 | 219 | 121 | 141 | 107 | 27 | 13 | 77 | 56 | 0,5 | -1,7 | 2,2 |
| chr7:66452189-66452689:downstream_5565 | chr7 | 66452689 | 66460588 | - | SBDS | NM | 46 | 2,2 | 1 | -2 | 2,07E-02 | 1,85E-01 | 28 | 22 | 3 | 2 | 118 | 97 | 49 | 51 | 3,3 | 1,1 | 2,2 |
| chr12:111157112-111157612:downstream_18755 | chr12 | 111157612 | 111180783 | - | PPP1CC | NM | 65 | 2,2 | 1 | -3 | 1,84E-03 | 5,58E-02 | 43 | 64 | 24 | 12 | 64 | 89 | 100 | 127 | 1,6 | -0,6 | 2,1 |
| chr5:137890070-137890570:downstream_4445 | chr5 | 137890570 | 137911318 | - | HSPA9 | NM | 52 | 2,2 | 1 | -3 | 6,34E-03 | 1,07E-01 | 22 | 29 | 4 | 8 | 98 | 66 | 85 | 107 | 2,0 | -0,2 | 2,3 |
| chr12:6437422-6437922:downstream_9990 | chr12 | 6437922 | 6451283 | - | TNFRSF1A | NM | 46 | 2,2 | 1 | -3 | 5,34E-03 | 9,96E-02 | 11 | 12 | 13 | 14 | 29 | 21 | 152 | 118 | -0,2 | -2,4 | 2,2 |
| chr14:78183443-78183943:downstream_11257 | chr14 | 78183943 | 78227497 | - | SNW1 | NM | 49 | 2,2 | 1 | -3 | 1,14E-02 | 1,43E-01 | 10 | 15 | 6 | 3 | 75 | 65 | 104 | 113 | 1,5 | -0,6 | 2,2 |
| chr6:27101314-27101814:downstream_4941 | chr6 | 27100816 | 27101314 | + | HIST1H2AG | NM | 218 | 2,2 | 1 | -4 | 3,36E-04 | 1,96E-02 | 15 | 14 | 28 | 14 | 123 | 97 | 687 | 767 | -0,6 | -2,7 | 2,2 |
| chr3:184427654-184428154:downstream_3300 | chr3 | 184428154 | 184429836 | - | MAGEF1 | NM | 49 | 2,2 | 1 | -3 | 5,39E-03 | 9,96E-02 | 17 | 14 | 7 | 6 | 60 | 61 | 121 | 110 | 1,3 | -0,9 | 2,2 |
| chr3:133309118-133309618:downstream_23068 | chr3 | 133292433 | 133309118 | + | CDV3 | NM | 61 | 2,2 | 1 | -3 | 8,14E-03 | 1,21E-01 | 12 | 26 | 3 | 6 | 96 | 109 | 104 | 128 | 2,2 | -0,2 | 2,3 |
| chr12:31433019-31433519:downstream_18562 | chr12 | 31433519 | 31479159 | - | FAM60A | NM | 31 | 2,2 | 1 | -3 | 6,00E-03 | 1,05E-01 | 36 | 48 | 14 | 13 | 25 | 28 | 45 | 37 | 1,6 | -0,6 | 2,3 |
| chr3:149679925-149680425:downstream_23386 | chr3 | 149530474 | 149679925 | + | RNF13 | NM | 53 | 2,3 | 1 | -2 | 2,55E-02 | 2,03E-01 | 4 | 6 | 4 | 2 | 54 | 61 | 152 | 145 | 0,8 | -1,4 | 2,1 |
| chr6:57035098-57035598:downstream_25361 | chr6 | 56954807 | 57035098 | + | ZNF451 | NM | 19 | 2,3 | 1 | -2 | 4,83E-02 | NA | 16 | 12 | 10 | 6 | 24 | 6 | 54 | 26 | 0,8 | -1,4 | 2,3 |
| chr1:33151812-33152312:downstream_17161 | chr1 | 33116748 | 33151812 | + | RBBP4 | NM | 28 | 2,3 | 1 | -2 | 1,81E-02 | NA | 11 | 9 | 15 | 6 | 18 | 12 | 86 | 64 | -0,1 | -2,3 | 2,2 |
| chr16:46988773-46989273:downstream_12341 | chr16 | 46989273 | 47007625 | - | DNAJA2 | NM | 27 | 2,3 | 1 | -2 | 3,79E-02 | NA | 7 | 7 | 0 | 6 | 20 | 38 | 68 | 75 | 1,2 | -1,3 | 2,5 |
| chr1:156706593-156707093:downstream_16983 | chr1 | 156707093 | 156710923 | - | MRPL24 | NM | 203 | 2,3 | 1 | -3 | 1,95E-03 | 5,70E-02 | 29 | 18 | 7 | 2 | 431 | 408 | 381 | 345 | 2,3 | 0,2 | 2,1 |
| chr7:99751833-99752333:downstream_5990 | chr7 | 99746529 | 99751833 | + | LAMTOR4 | NM | 30 | 2,3 | 1 | -2 | 1,55E-02 | 1,64E-01 | 25 | 32 | 4 | 3 | 50 | 60 | 33 | 35 | 2,9 | 0,7 | 2,3 |
| chrX:135950852-135951352:downstream_27003 | chrX | 135951352 | 135962939 | - | RBMX | NM | 109 | 2,3 | 1 | -4 | 2,30E-04 | 1,84E-02 | 35 | 36 | 15 | 9 | 150 | 144 | 244 | 240 | 1,5 | -0,7 | 2,3 |
| chr17:71202991-71203491:downstream_20473 | chr17 | 71203491 | 71228533 | - | FAM104A | NM | 34 | 2,3 | 1 | -2 | 2,81E-02 | 2,09E-01 | 20 | 15 | 3 | 2 | 85 | 62 | 50 | 39 | 3,0 | 0,7 | 2,3 |
| chr19:39891203-39891703:downstream_14286 | chr19 | 39881962 | 39891203 | + | MED29 | NM | 58 | 2,3 | 1 | -2 | 1,29E-02 | 1,48E-01 | 16 | 15 | 1 | 4 | 137 | 81 | 128 | 82 | 2,5 | 0,1 | 2,5 |
| chr2:101623189-101623689:downstream_1527 | chr2 | 101623689 | 101767846 | - | TBC1D8 | NM | 64 | 2,4 | 1 | -3 | 1,16E-03 | 4,15E-02 | 12 | 15 | 13 | 11 | 40 | 45 | 209 | 170 | 0,2 | -2,1 | 2,4 |
| chr3:63995724-63996224:downstream_23365 | chr3 | 63996224 | 64009686 | - | PSMD6 | NM | 106 | 2,4 | 1 | -4 | 1,95E-04 | 1,64E-02 | 60 | 54 | 14 | 17 | 181 | 107 | 207 | 203 | 1,9 | -0,5 | 2,4 |
| chr1:233808258-233808758:downstream_1429 | chr1 | 233749749 | 233808258 | + | KCNK1 | NM | 17 | 2,4 | 1 | -2 | 3,84E-02 | NA | 9 | 4 | 4 | 6 | 13 | 10 | 47 | 44 | 0,4 | -2,0 | 2,4 |
| chr8:101254132-101254632:downstream_26131 | chr8 | 101170262 | 101254132 | + | SPAG1 | NM | 22 | 2,4 | 1 | -2 | 1,96E-02 | NA | 19 | 27 | 3 | 6 | 33 | 26 | 32 | 29 | 2,4 | 0,0 | 2,5 |
| chr14:67853233-67853733:downstream_11192 | chr14 | 67827033 | 67853233 | + | EIF2S1 | NM | 30 | 2,4 | 1 | -2 | 2,85E-02 | 2,09E-01 | 14 | 14 | 1 | 2 | 69 | 50 | 47 | 45 | 2,8 | 0,4 | 2,5 |
| chr10:71962085-71962585:downstream_8552 | chr10 | 71962585 | 71993190 | - | PPA1 | NM | 69 | 2,4 | 1 | -2 | 3,54E-02 | 2,29E-01 | 4 | 13 | 1 | 2 | 97 | 172 | 138 | 128 | 2,5 | 0,0 | 2,4 |
| chr6:53361639-53362139:downstream_24777 | chr6 | 53362139 | 53409927 | - | GCLC | NM | 18 | 2,4 | 1 | -2 | 4,51E-02 | NA | 2 | 9 | 3 | 4 | 16 | 13 | 57 | 43 | 0,7 | -1,8 | 2,5 |
| chr6:44080872-44081372:downstream_5158 | chr6 | 44081372 | 44095191 | - | MRPL14 | NM | 18 | 2,4 | 1 | -2 | 4,43E-02 | NA | 8 | 17 | 6 | 3 | 31 | 9 | 40 | 31 | 1,5 | -0,9 | 2,4 |
| chr5:177023099-177023599:downstream_4609 | chr5 | 177019212 | 177023099 | + | TMED9 | NM | 65 | 2,4 | 1 | -2 | 2,41E-02 | 1,94E-01 | 14 | 19 | 4 | 0 | 165 | 151 | 67 | 97 | 3,0 | 0,9 | 2,0 |
| chr17:7146405-7146905:downstream_20424 | chr17 | 7146905 | 7155259 | - | CTDNEP1 | NM | 27 | 2,4 | 1 | -3 | 6,64E-03 | NA | 24 | 28 | 10 | 6 | 33 | 27 | 43 | 48 | 1,8 | -0,6 | 2,4 |
| chr11:66051550-66052050:downstream_18396 | chr11 | 66052050 | 66056638 | - | YIF1A | NM | 39 | 2,4 | 1 | -2 | 3,11E-02 | 2,19E-01 | 12 | 20 | 4 | 0 | 97 | 82 | 50 | 48 | 3,0 | 0,9 | 2,1 |
| chrX:69505710-69506210:downstream_7779 | chrX | 69506210 | 69509798 | - | PDZD11 | NM | 95 | 2,4 | 1 | -2 | 4,24E-02 | 2,47E-01 | 10 | 15 | 1 | 1 | 228 | 281 | 117 | 111 | 3,5 | 1,2 | 2,4 |
| chrX:13727944-13728444:downstream_26996 | chrX | 13707239 | 13727944 | + | RAB9A | NM | 41 | 2,4 | 1 | -2 | 1,87E-02 | 1,75E-01 | 8 | 15 | 4 | 2 | 45 | 86 | 96 | 72 | 2,0 | -0,4 | 2,3 |
| chr3:150259279-150259779:downstream_3233 | chr3 | 150259779 | 150264428 | - | SERP1 | NM | 46 | 2,5 | 1 | -3 | 1,65E-03 | 5,10E-02 | 51 | 44 | 13 | 5 | 60 | 70 | 65 | 57 | 2,5 | 0,1 | 2,4 |
| chr1:241660356-241660856:downstream_1449 | chr1 | 241660856 | 241683085 | - | FH | NM | 31 | 2,5 | 1 | -2 | 2,38E-02 | 1,94E-01 | 14 | 9 | 4 | 1 | 60 | 46 | 58 | 54 | 2,2 | -0,1 | 2,3 |
| chr12:110872194-110872694:downstream_18442 | chr12 | 110872694 | 110888216 | - | ARPC3 | NM | 220 | 2,5 | 1 | -2 | 4,21E-02 | 2,47E-01 | 4 | 7 | 0 | 2 | 455 | 393 | 505 | 397 | 2,7 | -0,1 | 2,8 |
| chr1:115246584-115247084:downstream_916 | chr1 | 115247084 | 115259515 | - | NRAS | NM | 70 | 2,5 | 1 | -3 | 1,99E-03 | 5,70E-02 | 28 | 15 | 7 | 4 | 121 | 96 | 160 | 128 | 2,0 | -0,4 | 2,4 |
| chr20:3763997-3764497:downstream_14865 | chr20 | 3764497 | 3767337 | - | CENPB | NM | 20 | 2,5 | 1 | -2 | 1,48E-02 | NA | 24 | 37 | 4 | 6 | 23 | 25 | 19 | 25 | 2,6 | 0,1 | 2,5 |
| chr6:28097856-28098356:downstream_4966 | chr6 | 28092386 | 28097856 | + | ZSCAN16 | NM | 21 | 2,5 | 1 | -2 | 3,60E-02 | NA | 17 | 15 | 7 | 1 | 45 | 15 | 36 | 36 | 2,0 | -0,3 | 2,3 |
| chr20:5095098-5095598:downstream_22500 | chr20 | 5095598 | 5107268 | - | PCNA | NM | 313 | 2,5 | 1 | -4 | 7,47E-05 | 8,08E-03 | 77 | 74 | 15 | 4 | 629 | 786 | 401 | 517 | 3,0 | 0,6 | 2,3 |
| chr12:47468989-47469489:downstream_18423 | chr12 | 47469489 | 47473734 | - | AMIGO2 | NM | 27 | 2,5 | 1 | -2 | 2,70E-02 | NA | 11 | 8 | 39 | 29 | 6 | 0 | 86 | 39 | -1,9 | -4,5 | 2,6 |
| chr15:78462884-78463384:downstream_11833 | chr15 | 78441718 | 78462884 | + | IDH3A | NM | 23 | 2,5 | 1 | -2 | 4,55E-02 | NA | 7 | 10 | 4 | 0 | 46 | 30 | 45 | 40 | 2,0 | -0,2 | 2,2 |
| chr7:99090353-99090853:downstream_5973 | chr7 | 99090853 | 99097877 | - | ZNF394 | NM | 22 | 2,5 | 1 | -2 | 2,02E-02 | NA | 11 | 15 | 1 | 4 | 27 | 35 | 47 | 35 | 2,2 | -0,4 | 2,7 |
| chr12:21671337-21671837:downstream_10099 | chr12 | 21654698 | 21671337 | + | GOLT1B | NM | 23 | 2,5 | 1 | -3 | 7,36E-03 | NA | 39 | 24 | 11 | 7 | 18 | 21 | 28 | 36 | 1,8 | -0,7 | 2,5 |
| chr5:1797998-1798498:downstream_4166 | chr5 | 1798498 | 1799956 | - | MRPL36 | NM | 51 | 2,5 | 1 | -3 | 1,96E-03 | 5,70E-02 | 33 | 32 | 6 | 4 | 85 | 96 | 89 | 65 | 2,7 | 0,2 | 2,5 |
| chr10:114189547-114190047:downstream_17824 | chr10 | 114190047 | 114206717 | - | ZDHHC6 | NM | 43 | 2,5 | 1 | -2 | 2,27E-02 | 1,92E-01 | 9 | 2 | 4 | 1 | 44 | 59 | 104 | 125 | 1,1 | -1,2 | 2,3 |
| chr17:73262457-73262957:downstream_13355 | chr17 | 73257748 | 73262457 | + | MRPS7 | NM | 53 | 2,5 | 1 | -3 | 1,05E-03 | 4,15E-02 | 51 | 31 | 6 | 8 | 76 | 87 | 75 | 89 | 2,6 | 0,0 | 2,6 |
| chr7:35671769-35672269:downstream_5790 | chr7 | 35672269 | 35734772 | - | HERPUD2 | NM | 15 | 2,6 | 1 | -2 | 4,10E-02 | NA | 4 | 9 | 6 | 2 | 9 | 13 | 47 | 29 | 0,7 | -1,8 | 2,5 |
| chr6:144415517-144416017:downstream_5384 | chr6 | 144416017 | 144416754 | - | SF3B5 | NM | 36 | 2,6 | 1 | -3 | 8,77E-03 | 1,24E-01 | 21 | 25 | 3 | 4 | 82 | 40 | 65 | 46 | 2,7 | 0,1 | 2,6 |
| chr6:160219461-160219961:downstream_5424 | chr6 | 160211491 | 160219461 | + | MRPL18 | NM | 157 | 2,6 | 1 | -4 | 1,61E-05 | 2,44E-03 | 84 | 71 | 14 | 11 | 275 | 282 | 295 | 221 | 2,7 | 0,1 | 2,5 |
| chr2:234204320-234204820:downstream_21757 | chr2 | 234160216 | 234204320 | + | ATG16L1 | NM | 28 | 2,6 | 1 | -2 | 2,70E-02 | NA | 4 | 11 | 1 | 2 | 46 | 29 | 64 | 63 | 1,9 | -0,8 | 2,7 |
| chr3:93776265-93776765:downstream_23112 | chr3 | 93776765 | 93782067 | - | DHFRL1 | NM | 17 | 2,6 | 1 | -2 | 2,08E-02 | NA | 44 | 31 | 6 | 6 | 16 | 11 | 18 | 7 | 2,7 | 0,1 | 2,6 |
| chr5:145891069-145891569:downstream_24407 | chr5 | 145826872 | 145891069 | + | TCERG1 | NM | 15 | 2,6 | 1 | -2 | 2,69E-02 | NA | 8 | 5 | 4 | 4 | 11 | 10 | 42 | 39 | 0,6 | -1,9 | 2,6 |
| chr8:144656454-144656954:downstream_26023 | chr8 | 144656954 | 144660521 | - | NAPRT | NM | 10 | 2,6 | 1 | -2 | 4,87E-02 | NA | 15 | 16 | 3 | 3 | 12 | 9 | 11 | 14 | 2,4 | -0,3 | 2,6 |
| chr9:101992901-101993401:downstream_7225 | chr9 | 101984569 | 101992901 | + | SEC61B | NM | 38 | 2,6 | 1 | -3 | 3,48E-03 | 7,72E-02 | 28 | 28 | 4 | 5 | 68 | 53 | 43 | 77 | 2,6 | 0,0 | 2,6 |
| chr6:27783267-27783767:downstream_4949 | chr6 | 27782821 | 27783267 | + | HIST1H2BM | NM | 47 | 2,6 | 1 | -3 | 1,16E-03 | 4,15E-02 | 10 | 21 | 13 | 15 | 24 | 25 | 111 | 155 | 0,2 | -2,5 | 2,6 |
| chr5:153437014-153437514:downstream_24227 | chr5 | 153418518 | 153437014 | + | MFAP3 | NM | 48 | 2,6 | 1 | -3 | 4,60E-03 | 9,28E-02 | 17 | 35 | 0 | 7 | 71 | 88 | 88 | 75 | 2,8 | 0,0 | 2,9 |
| chr1:101447311-101447811:downstream_17248 | chr1 | 101361631 | 101447311 | + | SLC30A7 | NM | 17 | 2,6 | 1 | -2 | 4,32E-02 | NA | 7 | 1 | 1 | 6 | 7 | 12 | 45 | 60 | 0,2 | -2,5 | 2,7 |
| chr12:49395554-49396054:downstream_18761 | chr12 | 49396054 | 49412629 | - | PRKAG1 | NM | 32 | 2,6 | 1 | -2 | 4,16E-02 | 2,47E-01 | 9 | 1 | 0 | 2 | 37 | 50 | 90 | 66 | 2,0 | -0,8 | 2,8 |
| chr2:206857944-206858444:downstream_2383 | chr2 | 206858444 | 206950906 | - | INO80D | NM | 18 | 2,6 | 1 | -2 | 4,89E-02 | NA | 3 | 3 | 0 | 8 | 4 | 9 | 68 | 48 | -0,4 | -3,2 | 2,8 |
| chr19:58269527-58270027:downstream_14706 | chr19 | 58258163 | 58269527 | + | ZNF776 | NM | 13 | 2,6 | 1 | -2 | 3,57E-02 | NA | 27 | 14 | 4 | 2 | 16 | 13 | 13 | 16 | 2,6 | 0,0 | 2,6 |
| chr12:122215159-122215659:downstream_10464 | chr12 | 122215659 | 122231594 | - | RHOF | NM | 11 | 2,7 | 1 | -2 | 3,87E-02 | NA | 19 | 21 | 7 | 7 | 7 | 3 | 13 | 10 | 1,6 | -1,1 | 2,7 |
| chr12:10851175-10851675:downstream_18920 | chr12 | 10851675 | 10875953 | - | YBX3 | NM | 70 | 2,7 | 1 | -3 | 6,43E-04 | 2,87E-02 | 70 | 64 | 4 | 5 | 161 | 128 | 53 | 73 | 3,9 | 1,2 | 2,7 |
| chr18:23595716-23596216:downstream_20949 | chr18 | 23596216 | 23671181 | - | SS18 | NM | 17 | 2,7 | 1 | -2 | 2,63E-02 | NA | 2 | 8 | 3 | 6 | 9 | 9 | 50 | 51 | 0,3 | -2,5 | 2,8 |
| chr19:48957164-48957664:downstream_14446 | chr19 | 48949029 | 48957164 | + | GRWD1 | NM | 43 | 2,7 | 1 | -2 | 3,76E-02 | 2,37E-01 | 7 | 7 | 1 | 1 | 67 | 100 | 83 | 74 | 2,7 | 0,1 | 2,6 |
| chr1:151374412-151374912:downstream_997 | chr1 | 151372040 | 151374412 | + | PSMB4 | NM | 140 | 2,7 | 1 | -3 | 7,49E-03 | 1,18E-01 | 17 | 24 | 4 | 0 | 363 | 355 | 166 | 192 | 3,3 | 1,0 | 2,3 |
| chr13:103331523-103332023:downstream_10625 | chr13 | 103249285 | 103331523 | + | TPP2 | NM | 26 | 2,7 | 1 | -2 | 2,63E-02 | NA | 3 | 9 | 4 | 2 | 12 | 37 | 83 | 60 | 1,0 | -1,6 | 2,6 |
| chr1:38031912-38032412:downstream_646 | chr1 | 38032412 | 38061586 | - | GNL2 | NM | 31 | 2,7 | 1 | -2 | 1,42E-02 | 1,57E-01 | 14 | 15 | 3 | 2 | 68 | 41 | 61 | 43 | 2,7 | 0,1 | 2,6 |
| chr11:62443469-62443969:downstream_18380 | chr11 | 62443969 | 62446589 | - | UBXN1 | NM | 41 | 2,7 | 1 | -3 | 1,26E-03 | 4,16E-02 | 20 | 36 | 11 | 6 | 46 | 42 | 90 | 75 | 1,7 | -0,9 | 2,6 |
| chr14:90874619-90875119:downstream_11268 | chr14 | 90863326 | 90874619 | + | CALM1 | NM | 49 | 2,7 | 1 | -3 | 8,18E-03 | 1,21E-01 | 8 | 28 | 3 | 2 | 83 | 99 | 93 | 77 | 2,8 | 0,1 | 2,7 |
| chr6:27778314-27778814:downstream_4951 | chr6 | 27777841 | 27778314 | + | HIST1H3H | NM | 59 | 2,7 | 1 | -4 | 2,78E-04 | 1,92E-02 | 19 | 21 | 15 | 22 | 33 | 22 | 131 | 208 | 0,1 | -2,6 | 2,7 |
| chr5:135518422-135518922:downstream_24377 | chr5 | 135468533 | 135518422 | + | SMAD5 | NM | 20 | 2,7 | 1 | -2 | 3,99E-02 | NA | 7 | 3 | 3 | 1 | 26 | 20 | 42 | 59 | 1,4 | -1,1 | 2,5 |
| chr19:41271297-41271797:downstream_14304 | chr19 | 41256758 | 41271297 | + | SNRPA | NM | 71 | 2,7 | 1 | -3 | 5,53E-04 | 2,71E-02 | 31 | 33 | 4 | 5 | 107 | 147 | 113 | 126 | 2,8 | 0,1 | 2,7 |
| chr19:45406946-45407446:downstream_21568 | chr19 | 45394476 | 45406946 | + | TOMM40 | NM | 70 | 2,7 | 1 | -3 | 5,53E-03 | 9,98E-02 | 15 | 21 | 0 | 3 | 149 | 155 | 118 | 100 | 3,5 | 0,5 | 3,0 |
| chr10:112271302-112271802:downstream_8719 | chr10 | 112257624 | 112271302 | + | DUSP5 | NM | 45 | 2,7 | 1 | -2 | 1,34E-02 | 1,52E-01 | 29 | 34 | 3 | 1 | 133 | 87 | 39 | 34 | 4,1 | 1,6 | 2,5 |
| chr3:183901879-183902379:downstream_23003 | chr3 | 183892633 | 183901879 | + | AP2M1 | NM | 29 | 2,7 | 1 | -3 | 6,23E-03 | 1,06E-01 | 21 | 18 | 3 | 3 | 54 | 39 | 45 | 51 | 2,7 | 0,0 | 2,7 |
| chr16:67967906-67968406:downstream_12425 | chr16 | 67968406 | 67970780 | - | PSMB10 | NM | 15 | 2,7 | 1 | -2 | 2,61E-02 | NA | 13 | 23 | 3 | 2 | 25 | 16 | 24 | 16 | 2,8 | 0,1 | 2,7 |
| chr10:103910090-103910590:downstream_17722 | chr10 | 103892750 | 103910090 | + | PPRC1 | NM | 40 | 2,7 | 1 | -3 | 1,06E-02 | 1,39E-01 | 9 | 14 | 3 | 2 | 82 | 49 | 70 | 94 | 2,3 | -0,3 | 2,7 |
| chr11:105969419-105969919:downstream_9625 | chr11 | 105948291 | 105969419 | + | AASDHPPT | NM | 26 | 2,7 | 1 | -2 | 2,35E-02 | NA | 6 | 22 | 4 | 1 | 34 | 52 | 39 | 52 | 2,5 | -0,1 | 2,6 |
| chr17:80197375-80197875:downstream_20711 | chr17 | 80186281 | 80197375 | + | SLC16A3 | NM | 14 | 2,7 | 1 | -2 | 4,43E-02 | NA | 2 | 7 | 1 | 2 | 13 | 12 | 38 | 36 | 1,3 | -1,6 | 2,8 |
| chr19:38874464-38874964:downstream_14264 | chr19 | 38865189 | 38874464 | + | PSMD8 | NM | 84 | 2,7 | 1 | -2 | 2,30E-02 | 1,92E-01 | 11 | 9 | 1 | 1 | 191 | 186 | 138 | 132 | 3,1 | 0,5 | 2,6 |
| chr6:26157343-26157843:downstream_4913 | chr6 | 26156558 | 26157343 | + | HIST1H1E | NM | 417 | 2,7 | 1 | -4 | 3,49E-04 | 1,96E-02 | 42 | 8 | 10 | 11 | 542 | 311 | 1343 | 1073 | 1,2 | -1,5 | 2,7 |
| chr1:2335740-2336240:downstream_17070 | chr1 | 2336240 | 2344010 | - | PEX10 | NM | 14 | 2,8 | 1 | -2 | 2,93E-02 | NA | 13 | 13 | 3 | 2 | 20 | 19 | 21 | 22 | 2,5 | -0,1 | 2,7 |
| chr13:41302931-41303431:downstream_10716 | chr13 | 41303431 | 41345347 | - | MRPS31 | NM | 19 | 2,8 | 1 | -2 | 3,08E-02 | NA | 4 | 11 | 1 | 2 | 30 | 15 | 49 | 37 | 1,9 | -0,9 | 2,9 |
| chr1:29096287-29096787:downstream_17412 | chr1 | 29063132 | 29096287 | + | YTHDF2 | NM | 33 | 2,8 | 1 | -2 | 2,52E-02 | 2,01E-01 | 5 | 14 | 3 | 1 | 65 | 45 | 93 | 41 | 2,4 | -0,3 | 2,6 |
| chrX:44382384-44382884:downstream_7682 | chrX | 44382884 | 44402221 | - | FUNDC1 | NM | 28 | 2,8 | 1 | -2 | 4,38E-02 | 2,52E-01 | 2 | 8 | 1 | 1 | 48 | 39 | 67 | 58 | 2,2 | -0,5 | 2,7 |
| chr20:44440066-44440566:downstream_14804 | chr20 | 44420575 | 44440066 | + | DNTTIP1 | NM | 15 | 2,8 | 1 | -2 | 4,72E-02 | NA | 8 | 6 | 0 | 2 | 28 | 15 | 25 | 37 | 2,5 | -0,5 | 3,0 |
| chr15:44854393-44854893:downstream_19790 | chr15 | 44854893 | 44955876 | - | SPG11 | NM | 18 | 2,8 | 1 | -2 | 1,86E-02 | NA | 4 | 10 | 4 | 2 | 15 | 13 | 47 | 44 | 1,0 | -1,7 | 2,8 |
| chr4:2742886-2743386:downstream_23893 | chr4 | 2743386 | 2758103 | - | TNIP2 | NM | 23 | 2,8 | 1 | -2 | 1,44E-02 | NA | 4 | 22 | 8 | 2 | 20 | 20 | 50 | 60 | 1,3 | -1,4 | 2,7 |
| chr2:27264565-27265065:downstream_22290 | chr2 | 27255773 | 27264565 | + | TMEM214 | NM | 32 | 2,8 | 1 | -2 | 1,68E-02 | 1,69E-01 | 9 | 9 | 4 | 0 | 46 | 56 | 57 | 75 | 2,1 | -0,4 | 2,5 |
| chr16:14763093-14763593:downstream_12204 | chr16 | 14726667 | 14763093 | + | BFAR | NM | 47 | 2,8 | 1 | -2 | 1,26E-02 | 1,46E-01 | 9 | 1 | 0 | 3 | 52 | 44 | 134 | 133 | 1,6 | -1,5 | 3,1 |
| chr15:63434264-63434764:downstream_19589 | chr15 | 63413998 | 63434264 | + | LACTB | NM | 22 | 2,8 | 1 | -2 | 1,49E-02 | NA | 4 | 9 | 0 | 6 | 18 | 15 | 53 | 74 | 1,2 | -1,9 | 3,1 |
| chr9:35099272-35099772:downstream_26519 | chr9 | 35099772 | 35103192 | - | STOML2 | NM | 27 | 2,9 | 1 | -2 | 2,14E-02 | NA | 6 | 17 | 1 | 2 | 54 | 40 | 39 | 53 | 2,9 | 0,0 | 2,9 |
| chr14:50249856-50250356:downstream_11131 | chr14 | 50234786 | 50249856 | + | KLHDC2 | NM | 21 | 2,9 | 1 | -3 | 3,41E-03 | NA | 35 | 31 | 7 | 8 | 20 | 12 | 25 | 28 | 2,1 | -0,8 | 2,9 |
| chr10:99477909-99478409:downstream_8664 | chr10 | 99473464 | 99477909 | + | MARVELD1 | NM | 25 | 2,9 | 1 | -3 | 7,43E-03 | NA | 25 | 17 | 4 | 2 | 43 | 35 | 29 | 42 | 2,9 | 0,1 | 2,7 |
| chr3:119243125-119243625:downstream_3125 | chr3 | 119217367 | 119243125 | + | TIMMDC1 | NM | 99 | 2,9 | 1 | -2 | 2,41E-02 | 1,94E-01 | 5 | 4 | 3 | 0 | 186 | 90 | 295 | 206 | 1,7 | -0,9 | 2,5 |
| chr14:53194716-53195216:downstream_11143 | chr14 | 53173895 | 53194716 | + | PSMC6 | NM | 31 | 2,9 | 1 | -3 | 5,17E-03 | 9,95E-02 | 11 | 17 | 3 | 2 | 41 | 48 | 75 | 48 | 2,5 | -0,5 | 2,9 |
| chr17:48167849-48168349:downstream_20524 | chr17 | 48133339 | 48167849 | + | ITGA3 | NM | 76 | 2,9 | 1 | -3 | 4,62E-03 | 9,28E-02 | 15 | 14 | 3 | 1 | 164 | 150 | 136 | 126 | 3,0 | 0,3 | 2,8 |

| | | | | | | | | | | | | | | | | | | | | | | | |
|---|---|---|---|---|---|---|---|---|---|---|---|---|---|---|---|---|---|---|---|---|---|---|---|
| chr19:48985571-48986071:downstream_21090 | chr19 | 48972464 | 48985571 | + | CYTH2 | NM | 38 | 3,0 | 1 | -2 | 2,38E-02 | 1,94E-01 | 4 | 7 | 1 | 1 | 58 | 59 | 99 | 79 | 2,3 | -0,6 | 2,9 |
| chr9:136227824-136228324:downstream_26524 | chr9 | 136228324 | 136244820 | - | SURF4 | NM | 150 | 3,0 | 1 | -3 | 1,24E-02 | 1,46E-01 | 9 | 8 | 1 | 1 | 281 | 319 | 276 | 305 | 2,9 | 0,0 | 2,9 |
| chr4:99991629-99992129:downstream_3479 | chr4 | 99992129 | 100009939 | - | ADH5 | NM | 12 | 3,0 | 1 | -2 | 2,53E-02 | NA | 10 | 9 | 4 | 2 | 7 | 15 | 25 | 25 | 1,7 | -1,2 | 2,9 |
| chr22:30825041-30825541:downstream_22856 | chr22 | 30821610 | 30825041 | + | MTFP1 | NM | 12 | 3,0 | 1 | -2 | 3,98E-02 | NA | 6 | 4 | 3 | 2 | 14 | 4 | 32 | 30 | 1,1 | -1,8 | 2,9 |
| chr7:87833931-87834431:downstream_25742 | chr7 | 87834431 | 87856308 | - | SRI | NM | 29 | 3,0 | 1 | -3 | 6,43E-04 | 2,87E-02 | 56 | 53 | 8 | 6 | 26 | 29 | 28 | 27 | 3,0 | 0,0 | 3,0 |
| chr2:235401185-235401685:downstream_21748 | chr2 | 235401685 | 235405697 | - | ARL4C | NM | 14 | 3,0 | 1 | -2 | 2,11E-02 | NA | 8 | 16 | 7 | 2 | 15 | 4 | 35 | 23 | 1,4 | -1,6 | 3,0 |
| chr19:36641255-36641755:downstream_21041 | chr19 | 36630827 | 36641255 | + | CAPNS1 | NM | 503 | 3,0 | 1 | -2 | 4,12E-02 | 2,46E-01 | 8 | 4 | 0 | 1 | 1082 | 1288 | 831 | 814 | 3,9 | 0,5 | 3,3 |
| chr3:49053386-49053886:downstream_2949 | chr3 | 49044636 | 49053386 | + | WDR6 | NM | 38 | 3,0 | 1 | -2 | 2,17E-02 | 1,90E-01 | 6 | 6 | 0 | 2 | 68 | 51 | 79 | 93 | 2,8 | -0,5 | 3,3 |
| chr1:150937220-150937720:downstream_17227 | chr1 | 150898814 | 150937220 | + | SETDB1 | NM | 8 | 3,0 | 2 | -2 | 4,77E-02 | NA | 10 | 15 | 7 | 3 | 5 | 3 | 4 | 21 | 1,3 | -1,7 | 3,0 |
| chr14:96011055-96011555:downstream_11293 | chr14 | 96001322 | 96011055 | + | GLRX5 | NM | 17 | 3,0 | 2 | -2 | 4,82E-02 | NA | 3 | 10 | 0 | 2 | 15 | 39 | 42 | 28 | 3,0 | -0,4 | 3,3 |
| chr11:93463522-93464022:downstream_8933 | chr11 | 93394815 | 93463522 | + | CEP295 | NM | 18 | 3,0 | 1 | -2 | 1,80E-02 | NA | 4 | 38 | 10 | 8 | 13 | 5 | 32 | 32 | 1,2 | -1,8 | 3,1 |
| chr6:27805157-27805657:downstream_4958 | chr6 | 27805657 | 27806117 | - | HIST1H2AK | NM | 50 | 3,1 | 1 | -4 | 2,10E-05 | 2,65E-03 | 64 | 44 | 26 | 34 | 25 | 15 | 96 | 94 | 0,9 | -2,2 | 3,1 |
| chr16:8943194-8943694:downstream_11964 | chr16 | 8891669 | 8943194 | + | PMM2 | NM | 38 | 3,1 | 1 | -3 | 3,14E-03 | 7,50E-02 | 10 | 10 | 6 | 1 | 52 | 30 | 107 | 88 | 1,6 | -1,2 | 2,9 |
| chr16:30966259-30966759:downstream_12307 | chr16 | 30960404 | 30966259 | + | ORAI3 | NM | 15 | 3,1 | 1 | -2 | 2,28E-02 | NA | 7 | 28 | 3 | 2 | 17 | 21 | 22 | 18 | 3,0 | -0,1 | 3,0 |
| chr9:135136326-135136826:downstream_7369 | chr9 | 135136826 | 135230372 | - | SETX | NM | 25 | 3,1 | 1 | -3 | 4,97E-03 | NA | 7 | 4 | 4 | 3 | 9 | 17 | 81 | 76 | 0,5 | -2,6 | 3,1 |
| chr16:75691341-75691841:downstream_12484 | chr16 | 75681634 | 75691341 | + | TERF2IP | NM | 64 | 3,1 | 1 | -3 | 3,51E-03 | 7,72E-02 | 27 | 15 | 4 | 0 | 155 | 126 | 100 | 89 | 3,3 | 0,6 | 2,7 |
| chr4:37864559-37865059:downstream_3652 | chr4 | 37828281 | 37864559 | + | PGM2 | NM | 9 | 3,1 | 1 | -2 | 3,32E-02 | NA | 9 | 15 | 1 | 7 | 7 | 3 | 18 | 12 | 1,6 | -1,7 | 3,3 |
| chr2:219119071-219119571:downstream_21751 | chr2 | 219081816 | 219119071 | + | ARPC2 | NM | 45 | 3,1 | 1 | -3 | 4,93E-03 | 9,71E-02 | 15 | 18 | 0 | 2 | 108 | 73 | 75 | 71 | 3,7 | 0,3 | 3,4 |
| chr10:119763926-119764426:downstream_8352 | chr10 | 119764426 | 119806114 | - | RAB11FIP2 | NM | 26 | 3,1 | 1 | -3 | 6,39E-03 | NA | 5 | 15 | 1 | 3 | 37 | 20 | 67 | 60 | 2,1 | -1,2 | 3,3 |
| chr12:4596395-4596895:downstream_18460 | chr12 | 4596895 | 4647674 | - | C12orf4 | NM | 19 | 3,2 | 1 | -3 | 6,01E-03 | NA | 11 | 8 | 7 | 6 | 14 | 3 | 50 | 53 | 0,5 | -2,6 | 3,2 |
| chr22:32894818-32895318:downstream_22806 | chr22 | 32870706 | 32894818 | + | FBXO7 | NM | 37 | 3,2 | 1 | -2 | 1,70E-02 | 1,69E-01 | 7 | 4 | 3 | 0 | 41 | 65 | 86 | 92 | 2,0 | -0,7 | 2,8 |
| chr4:107270381-107270881:downstream_23550 | chr4 | 107236766 | 107270381 | + | AIMP1 | NM | 36 | 3,2 | 2 | -2 | 4,41E-02 | 2,52E-01 | 11 | 8 | 0 | 1 | 106 | 74 | 47 | 41 | 4,5 | 1,0 | 3,5 |
| chr4:100816906-100817406:downstream_23619 | chr4 | 100817406 | 100867883 | - | DNAJB14 | NM | 27 | 3,2 | 1 | -3 | 4,15E-03 | NA | 11 | 3 | 3 | 3 | 17 | 21 | 71 | 84 | 1,2 | -2,0 | 3,2 |
| chr6:27781579-27782079:downstream_4952 | chr6 | 27782079 | 27782518 | - | HIST1H2AJ | NM | 182 | 3,2 | 1 | -4 | 1,08E-04 | 1,03E-02 | 7 | 6 | 4 | 5 | 105 | 76 | 757 | 495 | 0,4 | -2,8 | 3,2 |
| chr3:112250853-112251353:downstream_23011 | chr3 | 112251353 | 112280810 | - | ATG3 | NM | 13 | 3,2 | 1 | -2 | 1,80E-02 | NA | 10 | 4 | 1 | 4 | 11 | 6 | 39 | 26 | 1,3 | -2,0 | 3,3 |
| chr9:123616651-123617151:downstream_7287 | chr9 | 123605319 | 123616651 | + | PSMD5-AS1 | NR | 11 | 3,2 | 2 | -2 | 4,10E-02 | NA | 7 | 14 | 1 | 1 | 17 | 15 | 14 | 16 | 3,2 | 0,1 | 3,1 |
| chr7:108215294-108215794:downstream_6041 | chr7 | 108210188 | 108215294 | + | DNAJB9 | NM | 13 | 3,2 | 1 | -2 | 2,17E-02 | NA | 7 | 13 | 3 | 1 | 11 | 21 | 28 | 24 | 2,4 | -0,7 | 3,1 |
| chr1:151131723-151132223:downstream_16928 | chr1 | 151132223 | 151138424 | - | LYSMD1 | NM | 13 | 3,2 | 1 | -2 | 1,51E-02 | NA | 3 | 12 | 3 | 2 | 9 | 11 | 35 | 31 | 1,5 | -1,8 | 3,3 |
| chr12:49223038-49223538:downstream_10145 | chr12 | 49223538 | 49245957 | - | DDX23 | NM | 47 | 3,2 | 1 | -3 | 1,26E-03 | 4,16E-02 | 8 | 27 | 6 | 2 | 49 | 72 | 90 | 122 | 2,3 | -0,8 | 3,1 |
| chr1:201986315-201986815:downstream_16688 | chr1 | 201979689 | 201986315 | + | ELF3 | NM | 24 | 3,3 | 1 | -3 | 6,74E-03 | NA | 34 | 7 | 6 | 2 | 39 | 14 | 45 | 48 | 2,3 | -0,8 | 3,2 |
| chr1:113243248-113243748:downstream_17187 | chr1 | 113243748 | 113250025 | - | RHOC | NM | 52 | 3,3 | 2 | -2 | 3,56E-02 | 2,29E-01 | 12 | 4 | 0 | 1 | 120 | 127 | 70 | 81 | 4,3 | 0,7 | 3,6 |
| chr1:43629344-43629844:downstream_16675 | chr1 | 43629844 | 43638241 | - | EBNA1BP2 | NM | 122 | 3,3 | 2 | -2 | 3,20E-02 | 2,22E-01 | 4 | 20 | 0 | 1 | 332 | 334 | 121 | 162 | 4,9 | 1,2 | 3,7 |
| chr13:37583751-37584251:downstream_10704 | chr13 | 37574677 | 37583751 | + | EXOSC8 | NM | 13 | 3,3 | 1 | -2 | 1,42E-02 | NA | 11 | 3 | 4 | 2 | 9 | 6 | 31 | 37 | 1,0 | -2,2 | 3,2 |
| chr2:128618706-128619206:downstream_21737 | chr2 | 128619206 | 128643514 | - | AMMECR1L | NM | 9 | 3,3 | 1 | -2 | 2,58E-02 | NA | 11 | 11 | 6 | 2 | 7 | 3 | 11 | 20 | 1,6 | -1,6 | 3,2 |
| chr15:72533021-72533521:downstream_11789 | chr15 | 72533521 | 72563628 | - | PARP6 | NM | 22 | 3,3 | 1 | -2 | 1,89E-02 | NA | 3 | 8 | 1 | 1 | 26 | 29 | 50 | 55 | 2,3 | -0,9 | 3,2 |
| chr9:130628258-130628758:downstream_7330 | chr9 | 130628758 | 130640022 | - | AK1 | NM | 19 | 3,3 | 2 | -2 | 3,93E-02 | NA | 68 | 57 | 1 | 1 | 19 | 5 | 1 | 2 | 5,8 | 2,6 | 3,2 |
| chr6:43479064-43479564:downstream_5153 | chr6 | 43479564 | 43484728 | - | YIPF3 | NM | 25 | 3,3 | 2 | -2 | 3,88E-02 | NA | 11 | 7 | 0 | 1 | 52 | 63 | 38 | 30 | 4,5 | 0,8 | 3,7 |
| chr12:53458162-53458662:downstream_18886 | chr12 | 53440809 | 53458162 | + | TNS2 | NM | 8 | 3,4 | 2 | -2 | 3,43E-02 | NA | 3 | 10 | 8 | 2 | 3 | 2 | 18 | 21 | 0,4 | -2,9 | 3,3 |
| chr19:59066578-59067078:downstream_14731 | chr19 | 59067078 | 59070343 | - | UBE2M | NM | 18 | 3,4 | 1 | -3 | 1,23E-02 | NA | 29 | 8 | 1 | 2 | 26 | 29 | 25 | 22 | 3,6 | 0,2 | 3,4 |
| chr15:23004183-23004683:downstream_19647 | chr15 | 23004683 | 23034427 | - | NIPA2 | NM | 17 | 3,4 | 1 | -3 | 6,37E-03 | NA | 19 | 10 | 3 | 2 | 24 | 11 | 42 | 25 | 2,4 | -0,9 | 3,4 |
| chr6:99989762-99990262:downstream_24550 | chr6 | 99990262 | 100016690 | - | CCNC | NM | 25 | 3,4 | 1 | -3 | 1,27E-03 | NA | 18 | 20 | 4 | 2 | 23 | 32 | 61 | 35 | 2,5 | -0,8 | 3,3 |
| chr4:174252026-174252526:downstream_23692 | chr4 | 174252526 | 174255595 | - | HMGB2 | NM | 14 | 3,4 | 1 | -3 | 1,18E-02 | NA | 23 | 28 | 7 | 7 | 11 | 0 | 18 | 15 | 1,8 | -1,6 | 3,5 |
| chr17:28574712-28575212:downstream_12952 | chr17 | 28575212 | 28619184 | - | BLMH | NM | 34 | 3,4 | 1 | -4 | 2,71E-04 | 1,92E-02 | 11 | 9 | 4 | 7 | 22 | 13 | 95 | 111 | 0,9 | -2,6 | 3,5 |
| chr8:95937699-95938199:downstream_26159 | chr8 | 95938199 | 95961615 | - | TP53INP1 | NM | 24 | 3,5 | 2 | -2 | 4,52E-02 | NA | 11 | 11 | 0 | 0 | 81 | 60 | 19 | 10 | Inf | 2,3 | Inf |
| chr19:39305561-39306061:downstream_13772 | chr19 | 39306061 | 39322497 | - | ECH1 | NM | 104 | 3,5 | 2 | -2 | 2,73E-02 | 2,09E-01 | 1 | 11 | 1 | 0 | 251 | 147 | 206 | 217 | 3,0 | -0,1 | 3,1 |
| chr6:27859976-27860476:downstream_4956 | chr6 | 27860476 | 27860963 | - | HIST1H2AM | NM | 597 | 3,5 | 1 | -6 | 2,33E-08 | 6,34E-06 | 22 | 16 | 7 | 14 | 385 | 253 | 2197 | 1879 | 0,9 | -2,7 | 3,6 |
| chr17:57936340-57936840:downstream_20798 | chr17 | 57936840 | 57970306 | - | TUBD1 | NM | 10 | 3,5 | 2 | -2 | 2,07E-02 | NA | 8 | 14 | 3 | 2 | 2 | 15 | 18 | 20 | 2,3 | -1,2 | 3,4 |
| chr10:1085463-1085963:downstream_8378 | chr10 | 1085963 | 1095061 | - | IDI1 | NM | 85 | 3,5 | 1 | -3 | 3,81E-03 | 8,26E-02 | 14 | 6 | 3 | 0 | 168 | 130 | 188 | 176 | 2,8 | -0,3 | 3,1 |
| chr1:115109680-115110180:downstream_915 | chr1 | 115110180 | 115124265 | - | BCAS2 | NM | 20 | 3,5 | 2 | -2 | 3,31E-02 | NA | 12 | 12 | 0 | 1 | 35 | 55 | 18 | 27 | 4,9 | 1,0 | 3,9 |
| chrX:119755016-119755516:downstream_26912 | chrX | 119737743 | 119755016 | + | MCTS1 | NM | 10 | 3,6 | 2 | -2 | 2,57E-02 | NA | 8 | 10 | 1 | 1 | 9 | 15 | 19 | 15 | 3,0 | -0,5 | 3,5 |
| chr12:12268460-12268960:downstream_9821 | chr12 | 12268960 | 12419811 | - | LRP6 | NM | 60 | 3,6 | 1 | -3 | 2,04E-03 | 5,72E-02 | 1 | 3 | 8 | 3 | 3 | 10 | 209 | 244 | -1,6 | -5,1 | 3,5 |
| chr9:123577831-123578331:downstream_26459 | chr9 | 123578331 | 123605299 | - | PSMD5 | NM | 12 | 3,6 | 2 | -2 | 2,04E-02 | NA | 10 | 10 | 3 | 0 | 12 | 20 | 29 | 16 | 2,8 | -0,5 | 3,3 |
| chr7:42977453-42977953:downstream_5803 | chr7 | 42971938 | 42977453 | + | MRPL32 | NM | 306 | 3,6 | 1 | -4 | 2,54E-04 | 1,92E-02 | 23 | 37 | 3 | 1 | 781 | 675 | 504 | 422 | 4,0 | 0,7 | 3,4 |
| chr1:154527504-154528004:downstream_1083 | chr1 | 154526084 | 154527504 | + | UBE2Q1-AS1 | NR | 7 | 3,6 | 2 | -2 | 4,30E-02 | NA | 3 | 10 | 10 | 3 | 2 | 0 | 18 | 12 | 0,0 | -3,6 | 3,6 |
| chr12:6665535-6666035:downstream_18709 | chr12 | 6666035 | 6677498 | - | NOP2 | NM | 119 | 3,6 | 2 | -2 | 2,30E-02 | 1,92E-01 | 7 | 6 | 0 | 0 | 377 | 313 | 117 | 133 | Inf | 1,5 | Inf |
| chr6:27105571-27106071:downstream_4938 | chr6 | 27106071 | 27114637 | - | HIST1H2BK | NM | 59 | 3,6 | 2 | -2 | 1,67E-02 | 1,69E-01 | 11 | 12 | 0 | 1 | 146 | 141 | 82 | 79 | 4,8 | 0,8 | 4,0 |
| chrX:40465888-40466388:downstream_7672 | chrX | 40440215 | 40465888 | + | ATP6AP2 | NM | 264 | 3,6 | 1 | -4 | 3,04E-04 | 1,96E-02 | 15 | 35 | 1 | 2 | 559 | 593 | 476 | 431 | 4,0 | 0,3 | 3,7 |
| chr1:24121588-24122088:downstream_16759 | chr1 | 24122088 | 24127294 | - | GALE | NM | 14 | 3,6 | 1 | -2 | 1,45E-02 | NA | 16 | 18 | 0 | 2 | 20 | 22 | 15 | 15 | 4,4 | 0,5 | 3,9 |
| chr5:128449719-128450219:downstream_4393 | chr5 | 128430441 | 128449719 | + | ISOC1 | NM | 11 | 3,7 | 2 | -2 | 4,28E-02 | NA | 12 | 14 | 0 | 1 | 24 | 16 | 13 | 8 | 5,0 | 0,9 | 4,0 |
| chr7:104756320-104756820:downstream_25743 | chr7 | 104756820 | 105029377 | - | SRPK2 | NM | 11 | 3,7 | 2 | -2 | 3,23E-02 | NA | 2 | 3 | 6 | 8 | 2 | 0 | 29 | 39 | -1,6 | -5,4 | 3,8 |
| chr9:72999012-72999512:downstream_6870 | chr9 | 72999512 | 73029573 | - | KLF9 | NM | 12 | 3,7 | 2 | -2 | 1,79E-02 | NA | 1 | 9 | 3 | 1 | 8 | 8 | 39 | 32 | 1,4 | -2,2 | 3,6 |
| chr14:34901643-34902143:downstream_11103 | chr14 | 34902143 | 34931468 | - | SPTSSA | NM | 19 | 3,7 | 2 | -2 | 3,67E-02 | NA | 7 | 1 | 0 | 1 | 37 | 19 | 47 | 40 | 3,3 | -0,6 | 4,0 |
| chr12:64895899-64896399:downstream_9872 | chr12 | 64845839 | 64895899 | + | TBK1 | NM | 84 | 3,7 | 1 | -3 | 6,17E-04 | 2,87E-02 | 7 | 16 | 1 | 2 | 109 | 125 | 221 | 189 | 2,9 | -0,8 | 3,7 |
| chr1:24114722-24115222:downstream_547 | chr1 | 24104875 | 24114722 | + | PITHD1 | NM | 19 | 3,7 | 2 | -2 | 2,87E-02 | NA | 9 | 21 | 0 | 1 | 44 | 39 | 15 | 21 | 5,2 | 1,2 | 4,0 |
| chr3:195775654-195776154:downstream_23464 | chr3 | 195776154 | 195809032 | - | TFRC | NM | 150 | 3,7 | 1 | -3 | 2,38E-03 | 6,57E-02 | 11 | 4 | 0 | 2 | 234 | 192 | 417 | 344 | 3,1 | -0,8 | 4,0 |
| chr10:103923628-103924128:downstream_17672 | chr10 | 103911932 | 103923628 | + | NOLC1 | NM | 119 | 3,7 | 1 | -3 | 1,23E-02 | 1,46E-01 | 14 | 10 | 1 | 0 | 320 | 266 | 175 | 165 | 4,1 | 0,8 | 3,3 |
| chr1:224572344-224572844:downstream_17402 | chr1 | 224572844 | 224622001 | - | WDR26 | NM | 15 | 3,7 | 2 | -2 | 1,86E-02 | NA | 2 | 12 | 0 | 2 | 17 | 18 | 25 | 41 | 3,1 | -0,9 | 4,0 |
| chr11:47606115-47606615:downstream_9239 | chr11 | 47600561 | 47606115 | + | NDUFS3 | NM | 25 | 3,7 | 1 | -2 | 1,26E-02 | NA | 2 | 12 | 0 | 2 | 34 | 26 | 85 | 39 | 3,0 | -1,0 | 4,1 |
| chr3:169716161-169716661:downstream_3270 | chr3 | 169684579 | 169716161 | + | SEC62 | NM | 15 | 3,8 | 1 | -3 | 4,84E-03 | NA | 7 | 12 | 1 | 2 | 15 | 15 | 35 | 34 | 2,6 | -1,1 | 3,8 |
| chr11:67921829-67922329:downstream_18323 | chr11 | 67922329 | 67981239 | - | SUV420H1 | NM | 12 | 3,8 | 2 | -2 | 4,40E-02 | NA | 2 | 9 | 1 | 0 | 24 | 11 | 24 | 27 | 2,9 | -0,5 | 3,4 |
| chr1:169432648-169433148:downstream_1212 | chr1 | 169433148 | 169455208 | - | SLC19A2 | NM | 13 | 3,8 | 2 | -2 | 1,29E-02 | NA | 8 | 8 | 3 | 0 | 13 | 15 | 25 | 29 | 2,5 | -0,9 | 3,4 |
| chr1:200829835-200830335:downstream_16548 | chr1 | 200708685 | 200829835 | + | CAMSAP2 | NM | 7 | 3,8 | 2 | -2 | 2,96E-02 | NA | 9 | 9 | 4 | 7 | 2 | 0 | 8 | 15 | 0,6 | -3,2 | 3,9 |
| chr7:151819432-151819932:downstream_25528 | chr7 | 151722758 | 151819432 | + | GALNT11 | NM | 20 | 3,8 | 1 | -3 | 1,99E-03 | NA | 5 | 18 | 3 | 2 | 15 | 17 | 65 | 37 | 2,1 | -1,6 | 3,8 |
| chr6:49460820-49461320:downstream_5173 | chr6 | 49431095 | 49460820 | + | CENPQ | NM | 6 | 3,8 | 2 | -2 | 4,17E-02 | NA | 3 | 8 | 1 | 2 | 3 | 3 | 18 | 7 | 1,9 | -2,0 | 3,9 |
| chr15:40618916-40619416:downstream_11663 | chr15 | 40616256 | 40618916 | + | INAFM2 | NM | 12 | 3,8 | 2 | -2 | 3,63E-02 | NA | 19 | 28 | 0 | 1 | 23 | 12 | 4 | 7 | 5,8 | 1,7 | 4,2 |
| chr3:186303589-186304089:downstream_3311 | chr3 | 186288464 | 186303589 | + | DNAJB11 | NM | 71 | 3,8 | 2 | -3 | 1,16E-02 | 1,43E-01 | 8 | 6 | 0 | 1 | 141 | 129 | 129 | 157 | 4,1 | -0,1 | 4,2 |
| chr17:1605583-1606083:downstream_12728 | chr17 | 1606083 | 1613662 | - | TLCD2 | NM | 14 | 3,8 | 2 | -2 | 3,99E-02 | NA | 2 | 8 | 0 | 0 | 33 | 31 | 21 | 19 | Inf | 0,7 | Inf |
| chr11:65689048-65689548:downstream_9462 | chr11 | 65686727 | 65689048 | + | DRAP1 | NM | 38 | 3,8 | 2 | -2 | 2,28E-02 | 1,92E-01 | 7 | 3 | 0 | 0 | 114 | 72 | 63 | 48 | Inf | 0,8 | Inf |
| chr5:159855751-159856251:downstream_24320 | chr5 | 159848813 | 159855751 | + | PTTG1 | NM | 28 | 3,9 | 1 | -3 | 7,69E-03 | 1,18E-01 | 65 | 57 | 3 | 0 | 18 | 62 | 4 | 16 | 5,5 | 2,0 | 3,4 |
| chr16:28548161-28548661:downstream_20160 | chr16 | 28548661 | 28550495 | - | NUPR1 | NM | 11 | 3,9 | 2 | -2 | 4,94E-02 | NA | 2 | 8 | 0 | 0 | 29 | 17 | 15 | 14 | Inf | 0,7 | Inf |
| chr10:70706603-70707103:downstream_8543 | chr10 | 70661033 | 70706603 | + | DDX50 | NM | 78 | 3,9 | 2 | -3 | 1,17E-02 | 1,44E-01 | 3 | 9 | 1 | 0 | 136 | 126 | 150 | 195 | 3,1 | -0,4 | 3,5 |
| chr14:23389219-23389719:downstream_19344 | chr14 | 23389719 | 23398794 | - | PRMT5 | NM | 20 | 3,9 | 2 | -2 | 3,22E-02 | NA | 6 | 3 | 0 | 0 | 64 | 27 | 26 | 34 | Inf | 0,6 | Inf |
| chr10:3179418-3179918:downstream_17709 | chr10 | 3179918 | 3215033 | - | PITRM1 | NM | 38 | 3,9 | 2 | -2 | 2,30E-02 | 1,92E-01 | 3 | 6 | 0 | 0 | 61 | 119 | 53 | 62 | Inf | 0,6 | Inf |
| chr16:4932402-4932902:downstream_20265 | chr16 | 4897631 | 4932402 | + | UBN1 | NM | 15 | 3,9 | 1 | -3 | 7,33E-03 | NA | 9 | 13 | 1 | 1 | 22 | 19 | 25 | 34 | 3,3 | -0,5 | 3,8 |
| chr4:159629778-159630278:downstream_3884 | chr4 | 159630278 | 159644552 | - | PPID | NM | 28 | 3,9 | 1 | -3 | 1,36E-03 | NA | 10 | 18 | 0 | 2 | 34 | 40 | 67 | 50 | 3,5 | -0,7 | 4,2 |
| chr12:104510357-104510857:downstream_10392 | chr12 | 104510857 | 104532040 | - | NFYB | NM | 6 | 3,9 | 2 | -2 | 2,98E-02 | NA | 11 | 15 | 0 | 5 | 2 | 3 | 7 | 7 | 2,4 | -1,6 | 4,0 |
| chr22:31500262-31500762:downstream_15564 | chr22 | 31500762 | 31503551 | - | SELM | NM | 11 | 3,9 | 2 | -2 | 4,27E-02 | NA | 3 | 7 | 0 | 0 | 26 | 21 | 10 | 18 | Inf | 0,8 | Inf |
| chrX:107326934-107327434:downstream_26995 | chrX | 107327434 | 107334874 | - | PSMD10 | NM | 27 | 4,0 | 2 | -2 | 1,33E-02 | NA | 8 | 10 | 0 | 1 | 46 | 56 | 54 | 40 | 4,4 | 0,1 | 4,3 |
| chr19:58853712-58854212:downstream_13810 | chr19 | 58838384 | 58853712 | + | ZSCAN22 | NM | 11 | 4,0 | 2 | -3 | 8,23E-03 | NA | 20 | 11 | 1 | 2 | 17 | 4 | 15 | 20 | 3,3 | -0,7 | 4,1 |
| chr19:19820780-19821280:downstream_14149 | chr19 | 19821280 | 19843921 | - | ZNF14 | NM | 9 | 4,0 | 2 | -2 | 3,03E-02 | NA | 16 | 18 | 0 | 1 | 13 | 13 | 7 | 6 | 5,4 | 1,0 | 4,4 |
| chr10:120793854-120794354:downstream_8745 | chr10 | 120789227 | 120793854 | + | NANOS1 | NM | 7 | 4,0 | 2 | -2 | 4,05E-02 | NA | 13 | 12 | 1 | 0 | 11 | 7 | 7 | 6 | 4,1 | 0,4 | 3,7 |
| chr8:21940038-21940538:downstream_25909 | chr8 | 21911065 | 21940038 | + | DMTN | NM | 8 | 4,0 | 2 | -2 | 4,83E-02 | NA | 0 | 10 | 1 | 1 | 11 | 1 | 28 | 16 | 2,1 | -1,9 | 4,0 |
| chrX:55034306-55034806:downstream_26613 | chrX | 55026755 | 55034306 | + | APEX2 | NM | 14 | 4,0 | 2 | -2 | 1,96E-02 | NA | 17 | 12 | 1 | 0 | 24 | 25 | 11 | 19 | 4,4 | 0,7 | 3,6 |
| chr17:40276493-40276993:downstream_20664 | chr17 | 40276993 | 40307062 | - | RAB5C | NM | 69 | 4,1 | 2 | -3 | 7,18E-03 | 1,18E-01 | 10 | 9 | 1 | 0 | 124 | 151 | 134 | 127 | 3,7 | 0,1 | 3,7 |
| chr1:36646451-36646951:downstream_16936 | chr1 | 36621565 | 36646451 | + | MAP7D1 | NM | 7 | 4,1 | 2 | -3 | 1,20E-02 | NA | 7 | 15 | 4 | 2 | 4 | 2 | 10 | 16 | 1,9 | -2,1 | 4,0 |
| chr4:25121126-25121626:downstream_3644 | chr4 | 25121626 | 25162204 | - | SEPSECS | NM | 7 | 4,1 | 2 | -2 | 2,04E-02 | NA | 8 | 14 | 3 | 1 | 7 | 2 | 17 | 6 | 2,6 | -1,4 | 4,0 |
| chr14:70181861-70182361:downstream_10958 | chr14 | 70078309 | 70181861 | + | SUSD6 | NM | 11 | 4,1 | 2 | -2 | 3,02E-02 | NA | 7 | 3 | 1 | 0 | 20 | 7 | 25 | 23 | 2,9 | -0,8 | 3,7 |
| chr12:108963160-108963660:downstream_18611 | chr12 | 108955238 | 108963160 | + | ISCU | NM | 205 | 4,1 | 2 | -3 | 8,54E-03 | 1,23E-01 | 10 | 21 | 0 | 0 | 711 | 607 | 152 | 138 | Inf | 2,2 | Inf |
| chr8:99172069-99172569:downstream_26059 | chr8 | 99129520 | 99172069 | + | POP1 | NM | 8 | 4,1 | 2 | -2 | 4,20E-02 | NA | 3 | 6 | 0 | 0 | 15 | 15 | 13 | 11 | Inf | 0,4 | Inf |
| chr15:74922542-74923042:downstream_19514 | chr15 | 74900712 | 74922542 | + | CLK3 | NM | 19 | 4,1 | 2 | -2 | 1,43E-02 | NA | 16 | 8 | 1 | 0 | 36 | 36 | 21 | 33 | 4,1 | 0,4 | 3,7 |
| chr17:38154213-38154713:downstream_12636 | chr17 | 38137020 | 38154213 | + | PSMD3 | NM | 32 | 4,1 | 2 | -3 | 1,23E-02 | 1,46E-01 | 3 | 14 | 1 | 0 | 38 | 73 | 56 | 68 | 3,6 | -0,2 | 3,8 |
| chr19:58006048-58006548:downstream_21641 | chr19 | 57999078 | 58006048 | + | ZNF419 | NM | 14 | 4,1 | 2 | -2 | 2,25E-02 | NA | 7 | 6 | 0 | 0 | 34 | 33 | 17 | 18 | Inf | 0,9 | Inf |
| chr19:44234800-44235300:downstream_14353 | chr19 | 44235300 | 44259142 | - | SMG9 | NM | 22 | 4,1 | 2 | -2 | 1,35E-02 | NA | 2 | 10 | 1 | 0 | 33 | 30 | 47 | 52 | 3,1 | -0,7 | 3,8 |
| chr9:130872949-130873449:downstream_7335 | chr9 | 130873449 | 130881013 | - | SLC25A25-AS1 | NR | 5 | 4,2 | 2 | -2 | 4,03E-02 | NA | 4 | 5 | 3 | 2 | 2 | 0 | 10 | 12 | 1,0 | -3,2 | 4,2 |
| chr20:32675614-32676114:downstream_14979 | chr20 | 32676114 | 32700085 | - | EIF2S2 | NM | 14 | 4,2 | 2 | -2 | 1,68E-02 | NA | 7 | 17 | 1 | 0 | 25 | 23 | 18 | 21 | 4,1 | 0,3 | 3,8 |
| chr5:81568638-81569138:downstream_4328 | chr5 | 81569138 | 81574235 | - | RPS23 | NM | 67 | 4,2 | 1 | -4 | 1,29E-04 | 1,15E-02 | 89 | 43 | 3 | 1 | 171 | 107 | 70 | 52 | 5,2 | 1,2 | 4,0 |
| chr1:44433694-44434194:downstream_123 | chr1 | 44412477 | 44433694 | + | IPO13 | NM | 7 | 4,2 | 2 | -2 | 4,21E-02 | NA | 6 | 5 | 0 | 0 | 20 | 9 | 7 | 12 | Inf | 0,6 | Inf |
| chr22:42069436-42069936:downstream_22861 | chr22 | 42069936 | 42084913 | - | NHP2L1 | NM | 8 | 4,2 | 2 | -2 | 3,70E-02 | NA | 5 | 10 | 0 | 0 | 15 | 17 | 6 | 11 | Inf | 1,0 | Inf |
| chr7:30407308-30407808:downstream_5772 | chr7 | 30323922 | 30407308 | + | ZNRF2 | NM | 7 | 4,2 | 2 | -2 | 4,82E-02 | NA | 2 | 7 | 4 | 10 | 1 | 0 | 19 | 10 | -0,7 | -5,2 | 4,4 |
| chr19:11663357-11663857:downstream_13730 | chr19 | 11663857 | 11670051 | - | ELOF1 | NM | 26 | 4,3 | 2 | -2 | 1,33E-02 | NA | 3 | 10 | 0 | 0 | 57 | 64 | 38 | 34 | Inf | 0,7 | Inf |
| chr20:56222947-56223447:downstream_22511 | chr20 | 56223447 | 56286592 | - | PMEPA1 | NM | 13 | 4,3 | 2 | -2 | 1,44E-02 | NA | 9 | 8 | 0 | 1 | 18 | 21 | 22 | 25 | 4,3 | -0,3 | 4,6 |
| chr16:30667734-30668234:downstream_12301 | chr16 | 30662240 | 30667734 | + | PRR14 | NM | 9 | 4,3 | 2 | -2 | 2,29E-02 | NA | 5 | 15 | 1 | 0 | 11 | 15 | 13 | 15 | 3,9 | -0,1 | 3,9 |
| chrX:153733989-153734489:downstream_26810 | chrX | 153734489 | 153744566 | - | FAM3A | NM | 12 | 4,3 | 2 | -2 | 2,12E-02 | NA | 7 | 5 | 0 | 0 | 24 | 28 | 13 | 22 | Inf | 0,6 | Inf |
| chr9:27546043-27546543:downstream_26241 | chr9 | 27546543 | 27573864 | - | C9orf72 | NM | 6 | 4,3 | 2 | -2 | 1,60E-02 | NA | 9 | 11 | 1 | 1 | 6 | 4 | 10 | 11 | 3,1 | -1,1 | 4,2 |
| chr6:26026623-26027123:downstream_4899 | chr6 | 26027123 | 26027480 | - | HIST1H4B | NM | 65 | 4,3 | 1 | -6 | 2,51E-08 | 6,34E-06 | 37 | 59 | 6 | 13 | 45 | 35 | 121 | 208 | 2,4 | -2,0 | 4,4 |
| chr6:27099594-27100094:downstream_4939 | chr6 | 27100094 | 27100575 | - | HIST1H2BJ | NM | 265 | 4,3 | 1 | -7 | 1,98E-12 | 9,98E-10 | 28 | 35 | 7 | 12 | 158 | 115 | 829 | 932 | 1,7 | -2,7 | 4,4 |
| chr16:53361414-53361914:downstream_19928 | chr16 | 53088944 | 53361414 | + | CHD9 | NM | 11 | 4,3 | 2 | -3 | 8,59E-03 | NA | 2 | 10 | 13 | 7 | 2 | 0 | 26 | 31 | -0,7 | -5,1 | 4,4 |
| chr6:26233939-26234439:downstream_4926 | chr6 | 26234439 | 26235216 | - | HIST1H1D | NM | 134 | 4,3 | 1 | -4 | 1,20E-05 | 2,02E-03 | 7 | 2 | 1 | 3 | 51 | 31 | 448 | 530 | 0,9 | -3,6 | 4,4 |
| chr19:18688270-18688770:downstream_21580 | chr19 | 18682613 | 18688270 | + | UBA52 | NM | 12 | 4,4 | 2 | -2 | 1,93E-02 | NA | 9 | 11 | 0 | 0 | 23 | 30 | 14 | 8 | Inf | 1,3 | Inf |
| chr14:62263146-62263646:downstream_10950 | chr14 | 62229074 | 62263146 | + | SNAPC1 | NM | 8 | 4,4 | 2 | -2 | 2,86E-02 | NA | 11 | 5 | 10 | 11 | 1 | 0 | 14 | 10 | -0,4 | -4,9 | 4,5 |
| chr7:112130943-112131443:downstream_25620 | chr7 | 112120907 | 112130943 | + | LSMEM1 | NM | 16 | 4,4 | 2 | -2 | 1,58E-02 | NA | 7 | 16 | 0 | 0 | 39 | 37 | 11 | 16 | Inf | 1,5 | Inf |
| chr12:56538460-56538960:downstream_18558 | chr12 | 56521985 | 56538460 | + | ESYT1 | NM | 37 | 4,4 | 2 | -3 | 6,00E-03 | 1,05E-01 | 15 | 4 | 1 | 0 | 65 | 62 | 72 | 75 | 3,7 | -0,2 | 4,0 |
| chrX:102864855-102865355:downstream_27067 | chrX | 102862833 | 102864855 | + | TCEAL3 | NM | 8 | 4,4 | 2 | -2 | 3,24E-02 | NA | 5 | 15 | 0 | 0 | 11 | 18 | 7 | 6 | Inf | 1,2 | Inf |
| chr1:16450331-16450831:downstream_489 | chr1 | 16450831 | 16482582 | - | EPHA2 | NM | 44 | 4,5 | 2 | -3 | 7,10E-03 | 1,18E-01 | 9 | 9 | 0 | 0 | 101 | 123 | 35 | 73 | Inf | 1,1 | Inf |
| chr5:139948689-139949189:downstream_4457 | chr5 | 139944149 | 139948689 | + | SLC35A4 | NM | 8 | 4,5 | 2 | -2 | 3,49E-02 | NA | 7 | 3 | 0 | 0 | 24 | 5 | 8 | 16 | Inf | 0,2 | Inf |
| chrX:62566606-62567106:downstream_7763 | chrX | 62567106 | 62571218 | - | SPIN4 | NM | 6 | 4,5 | 2 | -2 | 4,17E-02 | NA | 7 | 10 | 0 | 0 | 7 | 13 | 8 | 2 | Inf | 0,8 | Inf |
| chr7:100399686-100400186:downstream_5596 | chr7 | 100400186 | 100425143 | - | EPHB4 | NM | 7 | 4,5 | 2 | -2 | 2,75E-02 | NA | 6 | 5 | 0 | 0 | 16 | 11 | 7 | 16 | Inf | 0,3 | Inf |
| chr11:65142162-65142662:downstream_18298 | chr11 | 65142662 | 65151172 | - | SLC25A45 | NM | 18 | 4,5 | 2 | -2 | 1,37E-02 | NA | 4 | 20 | 0 | 0 | 33 | 51 | 15 | 16 | Inf | 1,4 | Inf |
| chr12:10551105-10551605:downstream_10047 | chr12 | 10516367 | 10551105 | + | LOC101928100 | NR | 9 | 4,5 | 2 | -2 | 2,08E-02 | NA | 9 | 6 | 0 | 0 | 20 | 14 | 11 | 10 | Inf | 0,7 | Inf |
| chr10:101991552-101992052:downstream_17532 | chr10 | 101992052 | 102027438 | - | CWF19L1 | NM | 7 | 4,6 | 2 | -3 | 1,06E-02 | NA | 7 | 6 | 3 | 2 | 3 | 0 | 13 | 19 | 1,3 | -3,3 | 4,6 |
| chr11:114321000-114321500:downstream_9660 | chr11 | 114310107 | 114321000 | + | REXO2 | NM | 61 | 4,6 | 2 | -3 | 2,61E-03 | 6,83E-02 | 18 | 11 | 0 | 1 | 134 | 113 | 121 | 93 | 5,1 | 0,2 | 4,9 |
| chr16:30367921-30368421:downstream_12296 | chr16 | 30368421 | 30381522 | - | TBC1D10B | NM | 12 | 4,6 | 2 | -2 | 1,65E-02 | NA | 4 | 8 | 0 | 0 | 13 | 33 | 19 | 17 | Inf | 0,3 | Inf |
| chr7:6628610-6629110:downstream_25816 | chr7 | 6617064 | 6628610 | + | ZDHHC4 | NM | 24 | 4,6 | 2 | -3 | 8,65E-03 | NA | 4 | 8 | 0 | 0 | 63 | 38 | 51 | 30 | Inf | 0,3 | Inf |
| chr17:79640936-79641436:downstream_13430 | chr17 | 79633760 | 79640936 | + | CCDC137 | NM | 23 | 4,6 | 2 | -3 | 6,46E-03 | NA | 9 | 10 | 0 | 0 | 52 | 55 | 25 | 31 | Inf | 0,9 | Inf |
| chr11:117075286-117075786:downstream_9668 | chr11 | 117075786 | 117103241 | - | PCSK7 | NM | 11 | 4,6 | 2 | -3 | 1,06E-02 | NA | 6 | 11 | 1 | 0 | 13 | 13 | 17 | 25 | 3,6 | -0,7 | 4,3 |
| chr18:68019695-68020195:downstream_13664 | chr18 | 68002674 | 68019695 | + | LOC101927481 | NR | 9 | 4,7 | 2 | -2 | 2,37E-02 | NA | 8 | 6 | 29 | 8 | 0 | 0 | 4 | 17 | -1,4 | -Inf | Inf |
| chr6:49397572-49398072:downstream_4737 | chr6 | 49398072 | 49431041 | - | MUT | NM | 5 | 4,7 | 2 | -2 | 2,90E-02 | NA | 5 | 9 | 1 | 0 | 7 | 2 | 10 | 10 | 3,3 | -1,1 | 4,4 |
| chr8:20079207-20079707:downstream_6475 | chr8 | 20054703 | 20079207 | + | ATP6V1B2 | NM | 53 | 4,8 | 2 | -3 | 2,90E-03 | 7,20E-02 | 12 | 10 | 0 | 0 | 135 | 135 | 60 | 73 | Inf | 1,0 | Inf |
| chr6:27857592-27858092:downstream_4960 | chr6 | 27858092 | 27858570 | - | HIST1H3J | NM | 23 | 4,8 | 1 | -4 | 7,11E-06 | NA | 20 | 11 | 6 | 6 | 7 | 6 | 47 | 84 | 1,5 | -3,3 | 4,8 |
| chr11:6469342-6469842:downstream_18366 | chr11 | 6469842 | 6495689 | - | TRIM3 | NM | 7 | 4,8 | 2 | -2 | 1,81E-02 | NA | 5 | 8 | 0 | 0 | 11 | 12 | 10 | 11 | Inf | 0,2 | Inf |
| chr12:121161443-121161943:downstream_10455 | chr12 | 121148237 | 121161443 | + | UNC119B | NM | 4 | 4,8 | 2 | -2 | 3,76E-02 | NA | 7 | 6 | 1 | 1 | 2 | 0 | 4 | 7 | 2,6 | -2,2 | 4,8 |
| chr1:113453969-113454469:downstream_17240 | chr1 | 113454469 | 113498975 | - | SLC16A1 | NM | 9 | 4,9 | 2 | -2 | 1,49E-02 | NA | 7 | 3 | 0 | 0 | 10 | 18 | 21 | 13 | Inf | -0,3 | Inf |
| chr6:27861669-27862169:downstream_4962 | chr6 | 27861202 | 27861669 | + | HIST1H2BO | NM | 58 | 4,9 | 1 | -4 | 1,13E-05 | 2,02E-03 | 7 | 14 | 1 | 2 | 40 | 40 | 157 | 205 | 2,7 | -2,2 | 4,9 |
| chr17:80015247-80015747:downstream_13445 | chr17 | 80015747 | 80023697 | - | DUS1L | NM | 13 | 4,9 | 2 | -3 | 6,86E-03 | NA | 11 | 8 | 0 | 0 | 29 | 24 | 18 | 16 | Inf | 0,6 | Inf |
| chr14:50363772-50364272:downstream_11132 | chr14 | 50359735 | 50363772 | + | ARF6 | NM | 21 | 4,9 | 1 | -4 | 5,63E-05 | NA | 23 | 25 | 1 | 2 | 23 | 17 | 49 | 32 | 4,0 | -1,0 | 5,0 |
| chr9:139005926-139006426:downstream_26240 | chr9 | 139006426 | 139010731 | - | C9orf69 | NM | 28 | 5,0 | 2 | -3 | 3,20E-03 | 7,50E-02 | 7 | 13 | 0 | 0 | 57 | 69 | 35 | 47 | Inf | 0,6 | Inf |
| chr2:219528166-219528666:downstream_21768 | chr2 | 219524378 | 219528166 | + | BCS1L | NM | 10 | 5,1 | 2 | -3 | 7,97E-03 | NA | 15 | 13 | 0 | 0 | 17 | 17 | 7 | 9 | Inf | 1,1 | Inf |
| chr8:110585904-110586404:downstream_26141 | chr8 | 110586404 | 110704020 | - | SYBU | NM | 8 | 5,1 | 2 | -3 | 8,26E-03 | NA | 1 | 13 | 4 | 5 | 2 | 0 | 18 | 22 | 0,6 | -4,6 | 5,2 |
| chr2:96939900-96940400:downstream_2121 | chr2 | 96931883 | 96939900 | + | CIAO1 | NM | 12 | 5,1 | 2 | -3 | 6,91E-03 | NA | 11 | 24 | 0 | 0 | 15 | 32 | 11 | 7 | Inf | 1,3 | Inf |
| chr3:183863099-183863599:downstream_3292 | chr3 | 183852809 | 183863099 | + | EIF2B5 | NM | 20 | 5,1 | 2 | -3 | 4,87E-03 | NA | 2 | 8 | 0 | 0 | 43 | 17 | 50 | 43 | Inf | -0,6 | Inf |
| chr2:220101391-220101891:downstream_21742 | chr2 | 220094478 | 220101391 | + | ANKZF1 | NM | 11 | 5,2 | 2 | -3 | 8,19E-03 | NA | 4 | 7 | 0 | 0 | 25 | 8 | 21 | 22 | Inf | -0,4 | Inf |
| chr8:144647862-144648362:downstream_6766 | chr8 | 144648362 | 144654928 | - | MROH6 | NM | 12 | 5,2 | 2 | -3 | 5,54E-03 | NA | 12 | 20 | 0 | 0 | 23 | 23 | 14 | 7 | Inf | 1,2 | Inf |
| chr12:53700739-53701239:downstream_18407 | chr12 | 53701239 | 53715412 | - | AAAS | NM | 16 | 5,2 | 2 | -3 | 4,83E-03 | NA | 2 | 7 | 0 | 0 | 29 | 15 | 33 | 38 | Inf | -0,7 | Inf |
| chr10:120794040-120794540:downstream_8744 | chr10 | 120794540 | 120840334 | - | EIF3A | NM | 22 | 5,2 | 2 | -3 | 2,58E-03 | NA | 42 | 29 | 43 | 44 | 0 | 0 | 6 | 9 | -0,3 | -Inf | Inf |
| chr15:32932150-32932650:downstream_19467 | chr15 | 32907344 | 32932150 | + | ARHGAP11A | NM | 4 | 5,4 | 2 | -2 | 2,36E-02 | NA | 2 | 8 | 4 | 7 | 0 | 0 | 8 | 7 | -0,2 | -Inf | Inf |
| chr6:26184458-26184958:downstream_4915 | chr6 | 26184023 | 26184458 | + | HIST1H2BE | NM | 55 | 5,4 | 1 | -4 | 2,56E-05 | 2,98E-03 | 5 | 15 | 0 | 2 | 42 | 36 | 160 | 179 | 3,6 | -2,1 | 5,8 |

| | | | | | | | | | | | | | | | | | | | | | | | |
|---|---|---|---|---|---|---|---|---|---|---|---|---|---|---|---|---|---|---|---|---|---|---|---|
| chr3:71003364-71003864:downstream_23155 | chr3 | 71003864 | 71633140 | - | FOXP1 | NM | 4 | 5,4 | 2 | -2 | 2,56E-02 | NA | 2 | 9 | 6 | 4 | 0 | 0 | 8 | 5 | 0,1 | -Inf | Inf |
| chr1:33401546-33402046:downstream_17189 | chr1 | 33402046 | 33430414 | - | RNF19B | NM | 13 | 5,6 | 2 | -3 | 2,11E-03 | NA | 15 | 21 | 0 | 0 | 24 | 23 | 14 | 11 | Inf | 0,9 | Inf |
| chr16:19510434-19510934:downstream_20243 | chr16 | 19421860 | 19510434 | + | TMC5 | NM | 6 | 5,6 | 2 | -2 | 1,25E-02 | NA | 2 | 11 | 10 | 5 | 0 | 0 | 11 | 7 | -0,2 | -Inf | Inf |
| chr1:204391257-204391757:downstream_1321 | chr1 | 204391757 | 204459474 | - | PIK3C2B | NM | 5 | 5,6 | 2 | -3 | 9,02E-03 | NA | 5 | 7 | 1 | 0 | 2 | 3 | 11 | 13 | 3,1 | -2,2 | 5,2 |
| chr9:140476030-140476530:downstream_7448 | chr9 | 140476530 | 140484937 | - | ZMYND19 | NM | 12 | 5,7 | 2 | -3 | 2,64E-03 | NA | 2 | 7 | 0 | 0 | 16 | 12 | 29 | 33 | Inf | -1,2 | Inf |
| chr11:65420566-65421066:downstream_18255 | chr11 | 65421066 | 65430443 | - | RELA | NM | 23 | 5,7 | 2 | -3 | 8,34E-04 | NA | 8 | 12 | 0 | 0 | 44 | 35 | 36 | 46 | Inf | -0,1 | Inf |
| chr6:26032819-26033319:downstream_4904 | chr6 | 26033319 | 26033796 | - | HIST1H2AB | NM | 178 | 5,7 | 1 | -6 | 2,82E-09 | 1,07E-06 | 6 | 7 | 1 | 3 | 43 | 18 | 647 | 698 | 1,4 | -4,5 | 5,9 |
| chr20:5100615-5101115:downstream_14873 | chr20 | 5100231 | 5100615 | + | PCNA-AS1 | NR | 15 | 5,8 | 2 | -3 | 9,28E-04 | NA | 42 | 26 | 0 | 1 | 14 | 15 | 18 | 7 | 6,4 | 0,2 | 6,2 |
| chr19:36245420-36245920:downstream_14230 | chr19 | 36239261 | 36245420 | + | LIN37 | NM | 12 | 5,8 | 2 | -3 | 2,05E-03 | NA | 19 | 10 | 0 | 0 | 17 | 21 | 8 | 21 | Inf | 0,4 | Inf |
| chr10:72188374-72188874:downstream_8554 | chr10 | 72163860 | 72188374 | + | EIF4EBP2 | NM | 15 | 5,8 | 2 | -3 | 1,69E-03 | NA | 7 | 2 | 0 | 0 | 15 | 15 | 53 | 29 | Inf | -1,4 | Inf |
| chr1:46744145-46744645:downstream_17155 | chr1 | 46713366 | 46744145 | + | RAD54L | NM | 5 | 6,0 | 3 | -2 | 1,83E-02 | NA | 5 | 8 | 0 | 0 | 0 | 10 | 4 | 14 | Inf | -0,9 | Inf |
| chr10:101156126-101156626:downstream_8330 | chr10 | 101156626 | 101190530 | - | GOT1 | NM | 11 | 6,0 | 2 | -3 | 4,23E-03 | NA | 1 | 10 | 0 | 0 | 19 | 4 | 28 | 26 | Inf | -1,3 | Inf |
| chr5:16472646-16473146:downstream_24113 | chr5 | 16473146 | 16617167 | - | FAM134B | NM | 5 | 6,2 | 2 | -3 | 7,85E-03 | NA | 5 | 15 | 3 | 2 | 1 | 0 | 8 | 6 | 2,1 | -4,1 | 6,2 |
| chr8:110252647-110253147:downstream_26035 | chr8 | 110253147 | 110346350 | - | NUDCD1 | NM | 10 | 6,2 | 2 | -3 | 1,77E-03 | NA | 21 | 6 | 15 | 15 | 0 | 0 | 7 | 19 | -0,2 | -Inf | Inf |
| chr17:39031640-39032140:downstream_13055 | chr17 | 39032140 | 39041495 | - | KRT20 | NM | 3 | 6,5 | 3 | -2 | 1,87E-02 | NA | 7 | 3 | 0 | 0 | 3 | 0 | 4 | 8 | Inf | -1,9 | Inf |
| chr19:33087406-33087906:downstream_13761 | chr19 | 33087906 | 33166102 | - | ANKRD27 | NM | 69 | 6,5 | 2 | -4 | 9,62E-05 | 9,72E-03 | 1 | 10 | 0 | 0 | 50 | 69 | 239 | 183 | Inf | -1,8 | Inf |
| chr6:26273640-26274140:downstream_4929 | chr6 | 26273203 | 26273640 | + | HIST1H2BI | NM | 116 | 7,7 | 1 | -10 | 3,82E-22 | 5,80E-19 | 193 | 194 | 6 | 2 | 67 | 40 | 185 | 243 | 5,6 | -2,0 | 7,6 |
| chr1:8939943-8940443:downstream_420 | chr1 | 8938893 | 8939943 | + | ENO1-AS1 | NR | 5 | 7,9 | 2 | -3 | 6,27E-04 | NA | 10 | 14 | 0 | 0 | 1 | 3 | 7 | 7 | Inf | -1,9 | Inf |
| chr6:27107457-27107957:downstream_4937 | chr6 | 27107087 | 27107457 | + | HIST1H4I | NM | 17 | 9,5 | 2 | -5 | 1,40E-07 | NA | 10 | 12 | 0 | 0 | 4 | 4 | 60 | 48 | Inf | -3,8 | Inf |
| chr4:159629865-159630365:downstream_23633 | chr4 | 159593252 | 159629865 | + | ETFDH | NM | 16 | 2,6 | 1 | -2 | 5,01E-02 | NA | 8 | 6 | 10 | 9 | 11 | 0 | 50 | 36 | -0,4 | -3,0 | 2,6 |
| chr1:243663393-243663893:downstream_320 | chr1 | 243419306 | 243663393 | + | SDCCAG8 | NM | 6 | -4,3 | 2 | 2 | 5,06E-02 | NA | 1 | 0 | 6 | 6 | 7 | 14 | 11 | 2 | -3,8 | 0,6 | -4,4 |
| chr16:23585710-23586210:downstream_12264 | chr16 | 23568861 | 23585710 | + | UBFD1 | NM | 127 | 2,2 | 1 | -2 | 5,09E-02 | 2,79E-01 | 7 | 15 | 0 | 2 | 181 | 426 | 186 | 197 | 3,2 | 0,7 | 2,5 |
| chr1:51440309-51440809:downstream_16584 | chr1 | 51434366 | 51440309 | + | CDKN2C | NM | 17 | -3,6 | 2 | 2 | 5,09E-02 | NA | 0 | 0 | 13 | 6 | 6 | 15 | 45 | 51 | -Inf | -2,2 | -Inf |
| chr3:87304698-87305198:downstream_23076 | chr3 | 87276412 | 87304698 | + | CHMP2B | NM | 69 | 1,7 | 1 | -2 | 5,09E-02 | 2,79E-01 | 7 | 11 | 4 | 5 | 91 | 103 | 189 | 143 | 0,9 | -0,8 | 1,7 |
| chr1:201432911-201433411:downstream_1290 | chr1 | 201433411 | 201438299 | - | PHLDA3 | NM | 23 | 2,4 | 1 | -2 | 5,12E-02 | NA | 24 | 29 | 3 | 1 | 50 | 45 | 18 | 12 | 3,9 | 1,6 | 2,2 |
| chr3:49720934-49721434:downstream_2963 | chr3 | 49711434 | 49720934 | + | APEH | NM | 19 | 2,8 | 1 | -2 | 5,15E-02 | NA | 3 | 10 | 1 | 1 | 41 | 23 | 42 | 34 | 2,5 | -0,2 | 2,8 |
| chr9:96208281-96208781:downstream_7192 | chr9 | 96208781 | 96215874 | - | FAM120AOS | NM | 70 | 1,2 | 1 | -2 | 5,19E-02 | 2,82E-01 | 89 | 109 | 43 | 43 | 75 | 64 | 77 | 62 | 1,2 | 0,0 | 1,2 |
| chr9:37587911-37588411:downstream_26545 | chr9 | 37588411 | 37592636 | - | TOMM5 | NM | 629 | 1,2 | 1 | -2 | 5,19E-02 | 2,82E-01 | 21 | 28 | 3 | 11 | 1439 | 1463 | 1105 | 962 | 1,9 | 0,5 | 1,4 |
| chr6:122763992-122764492:downstream_5328 | chr6 | 122764492 | 122792952 | - | SERINC1 | NM | 91 | 1,3 | 1 | -2 | 5,24E-02 | 2,84E-01 | 28 | 35 | 8 | 16 | 151 | 175 | 157 | 153 | 1,4 | 0,1 | 1,3 |
| chr8:146034529-146035029:downstream_6798 | chr8 | 146024260 | 146034529 | + | ZNF517 | NM | 4 | 5,2 | 3 | -2 | 5,25E-02 | NA | 1 | 10 | 7 | 2 | 0 | 0 | 3 | 7 | 0,3 | -Inf | Inf |
| chr4:141303710-141304210:downstream_23840 | chr4 | 141178439 | 141303710 | + | SCOC | NM | 40 | 1,6 | 1 | -2 | 5,31E-02 | 2,87E-01 | 24 | 38 | 8 | 5 | 74 | 77 | 54 | 42 | 2,2 | 0,7 | 1,6 |
| chr10:47000568-47001068:downstream_8503 | chr10 | 46993545 | 47000568 | + | GPRIN2 | NM | 15 | 2,3 | 1 | -2 | 5,40E-02 | NA | 11 | 17 | 24 | 17 | 4 | 2 | 21 | 22 | -0,5 | -2,8 | 2,3 |
| chr1:213072705-213073205:downstream_1360 | chr1 | 213031596 | 213072705 | + | FLVCR1 | NM | 12 | 2,3 | 1 | -2 | 5,42E-02 | NA | 11 | 10 | 7 | 7 | 7 | 6 | 18 | 27 | 0,6 | -1,8 | 2,3 |
| chr11:61731256-61731756:downstream_8889 | chr11 | 61731756 | 61735132 | - | FTH1 | NM | 153 | -1,1 | 1 | 2 | 5,42E-02 | 2,91E-01 | 27 | 16 | 36 | 47 | 241 | 348 | 256 | 257 | -0,9 | 0,2 | -1,1 |
| chr12:21787774-21788274:downstream_18631 | chr12 | 21788274 | 21810789 | - | LDHB | NM | 121 | -1,4 | 1 | 2 | 5,48E-02 | 2,92E-01 | 11 | 6 | 11 | 23 | 338 | 187 | 170 | 224 | -1,0 | 0,4 | -1,4 |
| chr4:113191211-113191711:downstream_23559 | chr4 | 113152894 | 113191211 | + | AP1AR | NM | 5 | 3,8 | 2 | -2 | 5,49E-02 | NA | 7 | 7 | 3 | 3 | 2 | 0 | 6 | 11 | 1,1 | -2,7 | 3,9 |
| chr12:102138774-102139274:downstream_10381 | chr12 | 102139274 | 102224645 | - | GNPTAB | NM | 26 | 2,0 | 1 | -2 | 5,51E-02 | NA | 7 | 4 | 4 | 5 | 28 | 17 | 72 | 74 | 0,3 | -1,7 | 2,0 |
| chr17:80446143-80446643:downstream_20603 | chr17 | 80416059 | 80446143 | + | NARF | NM | 15 | 2,2 | 1 | -2 | 5,51E-02 | NA | 6 | 9 | 4 | 6 | 15 | 8 | 40 | 35 | 0,5 | -1,8 | 2,3 |
| chr9:72323937-72324437:downstream_6814 | chr9 | 72324437 | 72374876 | - | PTAR1 | NM | 22 | 2,5 | 1 | -2 | 5,53E-02 | NA | 3 | 7 | 3 | 1 | 18 | 40 | 51 | 55 | 1,5 | -0,9 | 2,4 |
| chr10:96092482-96092982:downstream_8649 | chr10 | 96092982 | 96122733 | - | NOC3L | NM | 47 | 1,5 | 1 | -2 | 5,55E-02 | 2,92E-01 | 41 | 72 | 17 | 9 | 84 | 64 | 54 | 39 | 2,1 | 0,7 | 1,5 |
| chr17:7571219-7571719:downstream_20787 | chr17 | 7571719 | 7590868 | - | TP53 | NM | 37 | 3,2 | 2 | -2 | 5,56E-02 | 2,92E-01 | 9 | 0 | 1 | 0 | 70 | 70 | 61 | 81 | 2,7 | 0,0 | 2,7 |
| chr1:110889303-110889803:downstream_17163 | chr1 | 110881944 | 110889303 | + | RBM15 | NM | 49 | 1,3 | 1 | -2 | 5,56E-02 | 2,92E-01 | 41 | 41 | 29 | 25 | 56 | 39 | 86 | 72 | 0,6 | -0,7 | 1,3 |
| chr12:117581084-117581584:downstream_18564 | chr12 | 117581584 | 117628300 | - | FBXO21 | NM | 47 | 1,6 | 1 | -2 | 5,57E-02 | 2,92E-01 | 7 | 13 | 8 | 10 | 57 | 31 | 138 | 115 | 0,1 | -1,5 | 1,7 |
| chr1:231467976-231468476:downstream_309 | chr1 | 231468476 | 231473618 | - | EXOC8 | NM | 61 | 1,6 | 1 | -2 | 5,57E-02 | 2,92E-01 | 18 | 28 | 3 | 4 | 138 | 155 | 74 | 69 | 2,7 | 1,0 | 1,7 |
| chr11:10873750-10874250:downstream_18397 | chr11 | 10874250 | 10879620 | - | ZBED5 | NM | 114 | 1,2 | 1 | -2 | 5,58E-02 | 2,92E-01 | 41 | 67 | 24 | 33 | 203 | 133 | 184 | 224 | 0,9 | -0,3 | 1,2 |
| chr16:56718108-56718608:downstream_12370 | chr16 | 56716381 | 56718108 | + | MT1X | NM | 13 | -2,9 | 2 | 2 | 5,59E-02 | NA | 1 | 1 | 8 | 7 | 16 | 22 | 24 | 22 | -3,1 | -0,3 | -2,9 |
| chr7:127220181-127220681:downstream_6068 | chr7 | 127220681 | 127225654 | - | GCC1 | NM | 27 | 2,0 | 1 | -2 | 5,64E-02 | NA | 22 | 32 | 4 | 4 | 65 | 28 | 36 | 21 | 2,7 | 0,7 | 2,0 |
| chr1:231498996-231499496:downstream_1421 | chr1 | 231499496 | 231560790 | - | EGLN1 | NM | 109 | 1,1 | 1 | -2 | 5,64E-02 | 2,95E-01 | 32 | 29 | 58 | 46 | 76 | 78 | 287 | 268 | -0,8 | -1,8 | 1,1 |
| chr4:114371687-114372187:downstream_23582 | chr4 | 114372187 | 114683083 | - | CAMK2D | NM | 7 | -4,0 | 2 | 2 | 5,70E-02 | NA | 0 | 0 | 7 | 5 | 10 | 4 | 15 | 16 | -Inf | -1,2 | -Inf |
| chr11:75133345-75133845:downstream_18265 | chr11 | 75110534 | 75133345 | + | RPS3 | NM | 9 | -3,8 | 2 | 2 | 5,71E-02 | NA | 7 | 10 | 1 | 3 | 16 | 33 | 1 | 0 | 1,8 | 5,1 | -3,4 |
| chr13:113977741-113978241:downstream_10891 | chr13 | 113951468 | 113977741 | + | LAMP1 | NM | 103 | 2,0 | 1 | -2 | 5,71E-02 | 2,96E-01 | 11 | 8 | 4 | 0 | 219 | 254 | 150 | 176 | 2,1 | 0,5 | 1,6 |
| chr15:25334043-25334543:downstream_11573 | chr15 | 25333949 | 25334043 | + | SNORD116-21 | NR | 155 | -1,4 | 1 | 2 | 5,73E-02 | 2,96E-01 | 2 | 10 | 11 | 9 | 380 | 361 | 213 | 257 | -0,7 | 0,7 | -1,4 |
| chr14:62214977-62215477:downstream_19253 | chr14 | 62162118 | 62214977 | + | HIF1A | NM | 208 | 1,5 | 1 | -2 | 5,75E-02 | 2,96E-01 | 3 | 7 | 8 | 4 | 208 | 167 | 640 | 629 | -0,3 | -1,8 | 1,4 |
| chr1:247001901-247002401:downstream_324 | chr1 | 247002401 | 247094726 | - | AHCTF1 | NM | 7 | -4,0 | 2 | 2 | 5,75E-02 | NA | 0 | 0 | 11 | 3 | 7 | 4 | 13 | 16 | -Inf | -1,4 | -Inf |
| chr2:36581391-36581891:downstream_1935 | chr2 | 36581891 | 36582713 | - | LOC100288911 | NR | 26 | 2,0 | 1 | -2 | 5,77E-02 | NA | 31 | 37 | 11 | 1 | 44 | 35 | 19 | 30 | 2,5 | 0,7 | 1,8 |
| chr1:224567153-224567653:downstream_16610 | chr1 | 224544512 | 224567153 | + | CNIH4 | NM | 20 | 1,9 | 1 | -2 | 5,78E-02 | NA | 20 | 26 | 3 | 10 | 24 | 26 | 25 | 30 | 1,9 | -0,1 | 2,0 |
| chr14:103603776-103604276:downstream_11379 | chr14 | 103592663 | 103603776 | + | TNFAIP2 | NM | 98 | 2,4 | 1 | -2 | 5,78E-02 | 2,96E-01 | 4 | 5 | 1 | 1 | 157 | 193 | 191 | 234 | 2,0 | -0,3 | 2,3 |
| chr19:36128587-36129087:downstream_14223 | chr19 | 36119979 | 36128587 | + | RBM42 | NM | 45 | 2,5 | 1 | -2 | 5,78E-02 | 2,96E-01 | 4 | 11 | 0 | 2 | 99 | 92 | 57 | 94 | 3,2 | 0,3 | 2,8 |
| chr1:156109880-156110380:downstream_16890 | chr1 | 156052336 | 156109880 | + | LMNA | NM | 41 | 1,7 | 1 | -2 | 5,79E-02 | 2,96E-01 | 30 | 33 | 1 | 10 | 64 | 92 | 45 | 52 | 2,5 | 0,7 | 1,8 |
| chrX:106174091-106174591:downstream_26669 | chrX | 106143393 | 106174091 | + | CLDN2 | NM | 134 | -1,8 | 1 | 2 | 5,80E-02 | 2,96E-01 | 2 | 1 | 15 | 17 | 128 | 149 | 340 | 418 | -3,3 | -1,5 | -1,8 |
| chr6:126360420-126360920:downstream_4794 | chr6 | 126307575 | 126360420 | + | TRMT11 | NM | 26 | 1,7 | 1 | -2 | 5,82E-02 | NA | 17 | 23 | 11 | 7 | 26 | 37 | 45 | 43 | 1,2 | -0,5 | 1,7 |
| chr1:146651528-146652028:downstream_951 | chr1 | 146649429 | 146651528 | + | PDIA3P1 | NR | 517 | -1,9 | 1 | 2 | 5,84E-02 | 2,96E-01 | 3 | 0 | 3 | 7 | 1106 | 1184 | 832 | 998 | -1,5 | 0,3 | -1,8 |
| chr17:61822109-61822609:downstream_13299 | chr17 | 61822609 | 61851088 | - | CCDC47 | NM | 63 | 1,6 | 1 | -2 | 5,84E-02 | 2,96E-01 | 13 | 16 | 0 | 11 | 85 | 108 | 156 | 112 | 1,4 | -0,5 | 1,8 |
| chr15:45010357-45010857:downstream_11701 | chr15 | 45003684 | 45010357 | + | B2M | NM | 2699 | 1,1 | 1 | -2 | 5,88E-02 | 2,97E-01 | 29 | 39 | 17 | 5 | 5941 | 7219 | 4493 | 3847 | 1,7 | 0,7 | 1,0 |
| chr19:764318-764818:downstream_13829 | chr19 | 751145 | 764318 | + | MISP | NM | 25 | 2,6 | 1 | -2 | 5,91E-02 | NA | 4 | 6 | 1 | 1 | 41 | 42 | 47 | 58 | 2,2 | -0,4 | 2,5 |
| chr2:27846082-27846582:downstream_22336 | chr2 | 27805835 | 27846082 | + | ZNF512 | NM | 35 | 1,8 | 1 | -2 | 5,97E-02 | 3,01E-01 | 7 | 18 | 14 | 3 | 41 | 34 | 71 | 94 | 0,6 | -1,1 | 1,7 |
| chr11:14478548-14479048:downstream_17965 | chr11 | 14479048 | 14521441 | - | COPB1 | NM | 56 | 1,5 | 1 | -2 | 5,99E-02 | 3,01E-01 | 12 | 9 | 6 | 8 | 74 | 70 | 127 | 146 | 0,6 | -0,9 | 1,5 |
| chr9:78999932-79000432:downstream_7144 | chr9 | 79000432 | 79009444 | - | RFK | NM | 72 | 1,2 | 1 | -2 | 6,01E-02 | 3,01E-01 | 86 | 66 | 31 | 23 | 110 | 96 | 61 | 107 | 1,5 | 0,3 | 1,2 |
| chr11:126132879-126133379:downstream_8961 | chr11 | 126081618 | 126132879 | + | FAM118B | NM | 26 | 1,8 | 1 | -2 | 6,05E-02 | NA | 21 | 46 | 40 | 24 | 10 | 8 | 28 | 29 | 0,1 | -1,7 | 1,8 |
| chr12:99129211-99129711:downstream_18433 | chr12 | 99039077 | 99129211 | + | APAF1 | NM | 7 | 3,4 | 2 | -2 | 6,11E-02 | NA | 11 | 9 | 1 | 2 | 11 | 0 | 11 | 12 | 2,3 | -1,1 | 3,5 |
| chr5:175810439-175810939:downstream_24253 | chr5 | 175810939 | 175815763 | - | NOP16 | NM | 59 | 3,1 | 2 | -2 | 6,15E-02 | 3,07E-01 | 5 | 4 | 0 | 0 | 189 | 146 | 57 | 69 | Inf | 1,4 | Inf |
| chr2:170494254-170494754:downstream_2303 | chr2 | 170440849 | 170494254 | + | PPIG | NM | 27 | -1,7 | 1 | 2 | 6,18E-02 | NA | 9 | 6 | 35 | 23 | 41 | 25 | 43 | 35 | -2,0 | -0,3 | -1,7 |
| chr16:88875376-88875876:downstream_19874 | chr16 | 88875876 | 88878342 | - | APRT | NM | 57 | 2,9 | 2 | -2 | 6,18E-02 | 3,07E-01 | 7 | 8 | 0 | 1 | 146 | 139 | 71 | 82 | 4,1 | 0,9 | 3,2 |
| chr14:68143018-68143518:downstream_19362 | chr14 | 68143518 | 68162510 | - | RDH11 | NM | 199 | 2,0 | 1 | -2 | 6,20E-02 | 3,07E-01 | 9 | 4 | 3 | 1 | 368 | 448 | 426 | 334 | 1,8 | 0,1 | 1,7 |
| chr16:57461580-57462080:downstream_19932 | chr16 | 57462080 | 57481440 | - | CIAPIN1 | NM | 15 | 2,2 | 1 | -2 | 6,24E-02 | NA | 20 | 23 | 3 | 10 | 12 | 14 | 13 | 23 | 1,8 | -0,5 | 2,2 |
| chr2:133401836-133402336:downstream_22062 | chr2 | 133402336 | 133429070 | - | LYPD1 | NM | 12 | 3,4 | 2 | -2 | 6,25E-02 | NA | 3 | 11 | 1 | 0 | 24 | 19 | 13 | 23 | 3,3 | 0,3 | 3,0 |
| chr12:105765296-105765796:downstream_10396 | chr12 | 105724413 | 105765296 | + | C12orf75 | NM | 19 | -2,7 | 1 | 2 | 6,26E-02 | NA | 2 | 0 | 6 | 7 | 34 | 31 | 31 | 40 | -2,9 | -0,1 | -2,8 |
| chr8:64079783-64080283:downstream_6574 | chr8 | 64080283 | 64081001 | - | YTHDF3-AS1 | NR | 4 | -4,8 | 3 | 2 | 6,27E-02 | NA | 0 | 0 | 0 | 8 | 7 | 5 | 8 | 3 | -Inf | 0,1 | -Inf |
| chr19:44149746-44150246:downstream_21331 | chr19 | 44150246 | 44174498 | - | PLAUR | NM | 20 | 2,3 | 1 | -2 | 6,28E-02 | NA | 6 | 18 | 3 | 2 | 40 | 29 | 33 | 27 | 2,4 | 0,2 | 2,3 |
| chr8:67834283-67834783:downstream_26008 | chr8 | 67782983 | 67834283 | + | MCMDC2 | NM | 8 | -3,0 | 2 | 2 | 6,29E-02 | NA | 2 | 1 | 10 | 3 | 14 | 18 | 11 | 8 | -2,3 | 0,7 | -3,1 |
| chr14:81726493-81726993:downstream_19406 | chr14 | 81726993 | 81893748 | - | STON2 | NM | 7 | 3,9 | 2 | -2 | 6,31E-02 | NA | 5 | 6 | 0 | 0 | 7 | 22 | 10 | 7 | Inf | 0,8 | Inf |
| chr18:21110962-21111462:downstream_13593 | chr18 | 21111462 | 21166581 | - | NPC1 | NM | 46 | -2,1 | 1 | 2 | 6,31E-02 | 3,11E-01 | 2 | 1 | 14 | 11 | 51 | 55 | 113 | 121 | -3,2 | -1,1 | -2,1 |
| chr20:48570422-48570922:downstream_15074 | chr20 | 48552913 | 48570422 | + | RNF114 | NM | 29 | 2,0 | 1 | -2 | 6,32E-02 | 3,11E-01 | 15 | 15 | 3 | 2 | 68 | 55 | 39 | 31 | 2,8 | 0,8 | 2,0 |
| chr4:186285120-186285620:downstream_3550 | chr4 | 186131283 | 186285120 | + | SNX25 | NM | 11 | 3,1 | 2 | -2 | 6,33E-02 | NA | 2 | 2 | 13 | 7 | 1 | 1 | 40 | 25 | -2,1 | -5,2 | 3,1 |
| chr11:27675941-27676441:downstream_17891 | chr11 | 27676441 | 27743605 | - | BDNF | NM | 19 | 2,0 | 1 | -2 | 6,38E-02 | NA | 7 | 10 | 21 | 21 | 5 | 4 | 39 | 49 | -1,4 | -3,3 | 2,0 |
| chr17:4848444-4848944:downstream_12591 | chr17 | 4848944 | 4852381 | - | PFN1 | NM | 195 | -1,1 | 1 | 2 | 6,41E-02 | 3,13E-01 | 28 | 24 | 46 | 25 | 346 | 541 | 245 | 303 | -0,5 | 0,7 | -1,2 |
| chr18:56994555-56995055:downstream_13521 | chr18 | 56995055 | 57026508 | - | LMAN1 | NM | 83 | 1,4 | 1 | -2 | 6,44E-02 | 3,13E-01 | 11 | 17 | 6 | 7 | 113 | 158 | 141 | 209 | 1,1 | -0,4 | 1,5 |
| chr6:26021186-26021686:downstream_4901 | chr6 | 26020717 | 26021186 | + | HIST1H3A | NM | 29 | 1,8 | 1 | -2 | 6,44E-02 | 3,13E-01 | 4 | 7 | 10 | 7 | 20 | 12 | 82 | 90 | -0,7 | -2,5 | 1,8 |
| chr19:49611870-49612370:downstream_21525 | chr19 | 49588396 | 49611870 | + | SNRNP70 | NM | 44 | 1,8 | 1 | -2 | 6,45E-02 | 3,13E-01 | 10 | 15 | 3 | 2 | 94 | 93 | 64 | 71 | 2,3 | 0,5 | 1,8 |
| chr8:144805602-144806102:downstream_6771 | chr8 | 144806102 | 144815914 | - | FAM83H | NM | 19 | 3,1 | 2 | -2 | 6,46E-02 | NA | 7 | 9 | 0 | 1 | 50 | 38 | 28 | 24 | 4,2 | 0,8 | 3,4 |
| chr18:66340424-66340924:downstream_13663 | chr18 | 66340924 | 66382353 | - | TMX3 | NM | 40 | 2,5 | 1 | -2 | 6,46E-02 | 3,13E-01 | 2 | 7 | 1 | 1 | 59 | 75 | 97 | 75 | 2,1 | -0,4 | 2,4 |
| chr5:162946359-162946859:downstream_24219 | chr5 | 162930069 | 162946359 | + | MAT2B | NM | 37 | 1,6 | 1 | -2 | 6,48E-02 | 3,13E-01 | 20 | 24 | 4 | 13 | 44 | 60 | 54 | 76 | 1,3 | -0,3 | 1,7 |
| chr1:155867098-155867598:downstream_17188 | chr1 | 155867598 | 155881193 | - | RIT1 | NM | 14 | 2,6 | 1 | -2 | 6,49E-02 | NA | 20 | 4 | 0 | 3 | 21 | 22 | 24 | 20 | 2,8 | 0,0 | 2,8 |
| chr19:40533666-40534166:downstream_14296 | chr19 | 40534166 | 40562115 | - | ZNF780B | NM | 16 | 2,0 | 1 | -2 | 6,50E-02 | NA | 17 | 25 | 11 | 15 | 13 | 5 | 24 | 21 | 0,7 | -1,3 | 2,0 |
| chr12:121016347-121016847:downstream_18751 | chr12 | 121016847 | 121019201 | - | POP5 | NM | 27 | 2,0 | 1 | -2 | 6,50E-02 | NA | 19 | 14 | 7 | 5 | 57 | 15 | 39 | 62 | 1,4 | -0,5 | 1,9 |
| chr17:40931618-40932118:downstream_13131 | chr17 | 40925453 | 40931618 | + | VPS25 | NM | 37 | 1,5 | 1 | -2 | 6,53E-02 | 3,13E-01 | 25 | 35 | 13 | 7 | 63 | 50 | 39 | 66 | 1,6 | 0,1 | 1,5 |
| chr1:45976206-45976706:downstream_17120 | chr1 | 45976706 | 45988562 | - | PRDX1 | NM | 695 | -1,1 | 1 | 2 | 6,53E-02 | 3,13E-01 | 27 | 13 | 26 | 27 | 1793 | 1528 | 885 | 1258 | -0,4 | 0,6 | -1,1 |
| chr2:64119166-64119666:downstream_22317 | chr2 | 64119666 | 64246214 | - | VPS54 | NM | 12 | 3,4 | 2 | -2 | 6,55E-02 | NA | 4 | 5 | 18 | 21 | 0 | 1 | 28 | 16 | -2,1 | -5,5 | 3,3 |
| chr14:54940708-54941208:downstream_11148 | chr14 | 54941208 | 54955744 | - | GMFB | NM | 32 | 1,6 | 1 | -2 | 6,58E-02 | 3,15E-01 | 85 | 44 | 22 | 14 | 28 | 22 | 15 | 26 | 1,8 | 0,3 | 1,6 |
| chr5:73237818-73238318:downstream_24017 | chr5 | 72921982 | 73237818 | + | ARHGEF28 | NM | 10 | -3,6 | 2 | 2 | 6,59E-02 | NA | 0 | 0 | 6 | 11 | 5 | 8 | 24 | 30 | -Inf | -2,1 | -Inf |
| chr5:140999942-141000442:downstream_4488 | chr5 | 141000442 | 141016423 | - | HDAC3 | NM | 13 | 3,4 | 2 | -2 | 6,61E-02 | NA | 2 | 11 | 0 | 1 | 27 | 23 | 18 | 26 | 3,9 | 0,2 | 3,7 |
| chr11:32624419-32624919:downstream_18002 | chr11 | 32605312 | 32624419 | + | EIF3M | NM | 104 | 1,1 | 1 | -2 | 6,61E-02 | 3,15E-01 | 59 | 71 | 18 | 16 | 228 | 197 | 106 | 135 | 1,9 | 0,8 | 1,1 |
| chr11:61564714-61565214:downstream_9380 | chr11 | 61560108 | 61564714 | + | FEN1 | NM | 32 | 1,9 | 1 | -2 | 6,64E-02 | 3,16E-01 | 6 | 6 | 1 | 4 | 41 | 39 | 71 | 87 | 1,1 | -1,0 | 2,0 |
| chr5:180256453-180256953:downstream_4631 | chr5 | 180256953 | 180258618 | - | HEIH | NR | 20 | 2,1 | 1 | -2 | 6,71E-02 | NA | 9 | 6 | 6 | 2 | 25 | 24 | 35 | 57 | 1,0 | -0,9 | 1,9 |
| chr3:196079868-196080368:downstream_2571 | chr3 | 196080368 | 196159345 | - | UBXN7 | NM | 32 | -1,7 | 1 | 2 | 6,72E-02 | 3,18E-01 | 4 | 4 | 32 | 43 | 23 | 20 | 71 | 60 | -3,2 | -1,6 | -1,6 |
| chr12:120898970-120899470:downstream_10450 | chr12 | 120899470 | 120907558 | - | SRSF9 | NM | 50 | 1,3 | 1 | -2 | 6,72E-02 | 3,18E-01 | 33 | 35 | 24 | 20 | 75 | 37 | 90 | 89 | 0,7 | -0,7 | 1,3 |
| chr2:216902610-216903110:downstream_1791 | chr2 | 216903110 | 216946539 | - | PECR | NM | 14 | -3,3 | 2 | 2 | 6,82E-02 | NA | 1 | 0 | 6 | 5 | 22 | 23 | 24 | 30 | -3,7 | -0,3 | -3,4 |
| chrX:47089394-47089894:downstream_26659 | chrX | 47077527 | 47089394 | + | CDK16 | NM | 13 | 2,1 | 1 | -2 | 6,82E-02 | NA | 16 | 21 | 6 | 5 | 11 | 15 | 13 | 19 | 1,8 | -0,3 | 2,1 |
| chr1:93307481-93307981:downstream_831 | chr1 | 93297593 | 93307481 | + | RPL5 | NM | 53 | 1,5 | 1 | -2 | 6,84E-02 | 3,21E-01 | 31 | 36 | 4 | 12 | 104 | 90 | 43 | 102 | 2,0 | 0,4 | 1,6 |
| chr20:24943079-24943579:downstream_14930 | chr20 | 24943579 | 24973425 | - | APMAP | NM | 422 | 1,5 | 1 | -2 | 6,85E-02 | 3,21E-01 | 7 | 13 | 4 | 2 | 692 | 1062 | 811 | 784 | 1,6 | 0,1 | 1,4 |
| chr3:178734510-178735010:downstream_23522 | chr3 | 178735010 | 178789656 | - | ZMAT3 | NM | 11 | 2,8 | 2 | -2 | 6,85E-02 | NA | 12 | 12 | 4 | 0 | 20 | 11 | 22 | 9 | 2,5 | 0,0 | 2,5 |
| chr6:26200216-26200716:downstream_4918 | chr6 | 26199786 | 26200216 | + | HIST1H2BF | NM | 8 | 2,9 | 2 | -2 | 6,86E-02 | NA | 14 | 4 | 3 | 3 | 3 | 7 | 18 | 8 | 1,5 | -1,4 | 2,9 |
| chr7:56168765-56169265:downstream_5838 | chr7 | 56169265 | 56174187 | - | CHCHD2 | NM | 94 | 1,1 | 1 | -2 | 6,88E-02 | 3,22E-01 | 176 | 205 | 58 | 39 | 90 | 90 | 39 | 57 | 2,0 | 0,9 | 1,1 |
| chr10:75543406-75543906:downstream_8575 | chr10 | 75541807 | 75543406 | + | CHCHD1 | NM | 26 | 3,0 | 2 | -2 | 6,91E-02 | NA | 3 | 6 | 1 | 0 | 58 | 43 | 42 | 53 | 2,7 | 0,1 | 2,6 |
| chr1:59245962-59246462:downstream_760 | chr1 | 59246462 | 59249785 | - | JUN | NM | 10 | 2,9 | 2 | -2 | 6,91E-02 | NA | 14 | 11 | 0 | 2 | 18 | 17 | 13 | 9 | 3,9 | 0,7 | 3,2 |
| chr17:7485429-7485929:downstream_20400 | chr17 | 7482804 | 7485429 | + | CD68 | NM | 52 | 1,7 | 1 | -2 | 6,98E-02 | 3,25E-01 | 17 | 30 | 0 | 7 | 104 | 133 | 71 | 53 | 2,8 | 0,9 | 1,9 |
| chr15:56921873-56922373:downstream_19851 | chr15 | 56922373 | 57026284 | - | ZNF280D | NM | 8 | -3,8 | 2 | 2 | 6,99E-02 | NA | 0 | 0 | 10 | 3 | 10 | 3 | 19 | 16 | -Inf | -1,5 | -Inf |
| chr13:41386596-41387096:downstream_10715 | chr13 | 41363546 | 41386596 | + | SLC25A15 | NM | 34 | 2,3 | 1 | -2 | 7,00E-02 | 3,25E-01 | 14 | 19 | 3 | 0 | 77 | 104 | 25 | 34 | 3,6 | 1,6 | 2,0 |
| chr3:63989136-63989636:downstream_23016 | chr3 | 63850232 | 63989136 | + | ATXN7 | NM | 12 | 2,4 | 1 | -2 | 7,00E-02 | NA | 2 | 8 | 6 | 5 | 7 | 5 | 26 | 36 | 0,0 | -2,5 | 2,4 |
| chr6:34845054-34845554:downstream_25291 | chr6 | 34845554 | 34855848 | - | TAF11 | NM | 14 | 2,1 | 1 | -2 | 7,03E-02 | NA | 23 | 20 | 4 | 3 | 19 | 16 | 11 | 15 | 2,5 | 0,4 | 2,1 |
| chr5:145135406-145135906:downstream_24312 | chr5 | 145135906 | 145214932 | - | PRELID2 | NM | 4 | 4,5 | 2 | -2 | 7,04E-02 | NA | 1 | 5 | 6 | 6 | 0 | 0 | 6 | 9 | -1,0 | -Inf | Inf |
| chr11:85347583-85348083:downstream_18341 | chr11 | 85339616 | 85347583 | + | TMEM126B | NM | 73 | 1,3 | 1 | -2 | 7,06E-02 | 3,27E-01 | 14 | 29 | 11 | 13 | 115 | 96 | 178 | 126 | 0,8 | -0,5 | 1,4 |
| chr17:73163324-73163824:downstream_20754 | chr17 | 73163824 | 73179098 | - | SUMO2 | NM | 18 | 1,9 | 1 | -2 | 7,10E-02 | NA | 13 | 18 | 6 | 5 | 21 | 23 | 28 | 27 | 1,6 | -0,3 | 1,9 |
| chr1:16944250-16944750:downstream_494 | chr1 | 16944750 | 16957401 | - | CROCCP2 | NR | 15 | 2,0 | 1 | -2 | 7,10E-02 | NA | 15 | 16 | 8 | 8 | 10 | 14 | 19 | 27 | 0,9 | -1,0 | 1,9 |
| chr13:108886603-108887103:downstream_10871 | chr13 | 108870762 | 108886603 | + | ABHD13 | NM | 5 | 3,6 | 2 | -2 | 7,12E-02 | NA | 5 | 8 | 6 | 1 | 2 | 1 | 10 | 6 | 1,0 | -2,6 | 3,6 |
| chrX:47833749-47834249:downstream_27127 | chrX | 47834249 | 47863394 | - | ZNF182 | NM | 5 | -4,0 | 2 | 2 | 7,13E-02 | NA | 1 | 0 | 3 | 8 | 8 | 7 | 7 | 5 | -3,8 | 0,3 | -4,1 |
| chr16:69738569-69739069:downstream_20140 | chr16 | 69599868 | 69738569 | + | NFAT5 | NM | 37 | 1,9 | 1 | -2 | 7,14E-02 | 3,30E-01 | 2 | 6 | 1 | 9 | 19 | 21 | 129 | 108 | -0,5 | -2,6 | 2,1 |
| chr19:33078358-33078858:downstream_14191 | chr19 | 33072093 | 33078358 | + | PDCD5 | NM | 26 | 2,6 | 1 | -2 | 7,18E-02 | NA | 3 | 12 | 0 | 2 | 52 | 55 | 28 | 59 | 3,2 | 0,3 | 2,9 |
| chr1:227177065-227177565:downstream_16578 | chr1 | 227177565 | 227505826 | - | CDC42BPA | NM | 12 | -3,4 | 2 | 2 | 7,18E-02 | NA | 0 | 0 | 4 | 7 | 7 | 14 | 31 | 31 | -Inf | -1,6 | -Inf |
| chr11:33101000-33101500:downstream_9199 | chr11 | 33097695 | 33101000 | + | LINC00294 | NR | 26 | 1,9 | 1 | -2 | 7,20E-02 | NA | 11 | 4 | 3 | 7 | 32 | 20 | 70 | 61 | 0,6 | -1,3 | 2,0 |
| chr4:184369049-184369549:downstream_3921 | chr4 | 184365788 | 184369049 | + | CDKN2AIP | NM | 30 | 1,5 | 1 | -2 | 7,22E-02 | 3,33E-01 | 28 | 22 | 17 | 15 | 35 | 22 | 53 | 48 | 0,7 | -0,8 | 1,5 |
| chr2:224832431-224832931:downstream_2458 | chr2 | 224822120 | 224832431 | + | MRPL44 | NM | 18 | -2,6 | 1 | 2 | 7,25E-02 | NA | 2 | 0 | 4 | 10 | 29 | 25 | 39 | 36 | -3,1 | -0,5 | -2,6 |
| chr14:23494559-23495059:downstream_19351 | chr14 | 23495059 | 23504429 | - | PSMB5 | NM | 284 | 1,0 | 1 | -2 | 7,25E-02 | 3,33E-01 | 35 | 32 | 13 | 11 | 620 | 661 | 399 | 500 | 1,5 | 0,5 | 1,0 |
| chr12:21243040-21243540:downstream_9834 | chr12 | 21168629 | 21243040 | + | SLCO1B7 | NM | 19 | 2,5 | 1 | -2 | 7,25E-02 | NA | 9 | 40 | 46 | 21 | 4 | 0 | 10 | 22 | -0,5 | -3,0 | 2,5 |
| chr14:65009954-65010454:downstream_11180 | chr14 | 65007185 | 65009954 | + | HSPA2 | NM | 14 | 2,7 | 1 | -2 | 7,28E-02 | NA | 7 | 7 | 3 | 0 | 25 | 20 | 24 | 22 | 2,3 | 0,0 | 2,3 |
| chr17:12047148-12047648:downstream_20566 | chr17 | 11924134 | 12047148 | + | MAP2K4 | NM | 17 | 2,0 | 1 | -2 | 7,30E-02 | NA | 5 | 13 | 8 | 9 | 7 | 13 | 33 | 47 | 0,0 | -2,0 | 2,0 |
| chr9:104151748-104152248:downstream_7232 | chr9 | 104152248 | 104160919 | - | MRPL50 | NM | 70 | 1,1 | 1 | -2 | 7,31E-02 | 3,35E-01 | 69 | 62 | 25 | 36 | 96 | 85 | 106 | 82 | 1,1 | -0,1 | 1,2 |
| chr9:34251877-34252377:downstream_7035 | chr9 | 34252377 | 34329198 | - | KIF24 | NM | 7 | 3,2 | 2 | -2 | 7,35E-02 | NA | 3 | 9 | 10 | 3 | 2 | 0 | 13 | 13 | -0,1 | -3,4 | 3,3 |
| chr10:99217580-99218080:downstream_17658 | chr10 | 99218080 | 99258551 | - | MMS19 | NM | 23 | 2,6 | 1 | -2 | 7,37E-02 | NA | 7 | 3 | 0 | 2 | 22 | 53 | 42 | 55 | 2,5 | -0,4 | 2,9 |
| chr9:131458675-131459175:downstream_26494 | chr9 | 131445933 | 131458675 | + | SET | NM | 193 | 1,2 | 1 | -2 | 7,39E-02 | 3,35E-01 | 50 | 17 | 24 | 21 | 311 | 269 | 477 | 374 | 0,6 | -0,6 | 1,1 |
| chr3:127790526-127791026:downstream_3165 | chr3 | 127771211 | 127790526 | + | SEC61A1 | NM | 34 | 2,9 | 2 | -2 | 7,39E-02 | 3,35E-01 | 11 | 7 | 0 | 1 | 98 | 83 | 24 | 51 | 4,5 | 1,3 | 3,2 |
| chr10:15146256-15146756:downstream_17751 | chr10 | 15139178 | 15146256 | + | RPP38 | NM | 38 | 1,7 | 1 | -2 | 7,39E-02 | 3,35E-01 | 7 | 17 | 1 | 5 | 72 | 68 | 65 | 68 | 2,0 | 0,1 | 1,9 |
| chr7:128505903-128506403:downstream_25402 | chr7 | 128502856 | 128505903 | + | ATP6V1F | NM | 44 | 1,4 | 1 | -2 | 7,40E-02 | 3,35E-01 | 69 | 47 | 11 | 8 | 89 | 60 | 31 | 34 | 2,6 | 1,2 | 1,4 |
| chr11:64812300-64812800:downstream_9429 | chr11 | 64808375 | 64812300 | + | SAC3D1 | NM | 53 | 2,1 | 1 | -2 | 7,45E-02 | 3,35E-01 | 22 | 6 | 0 | 2 | 157 | 117 | 58 | 64 | 3,5 | 1,2 | 2,3 |
| chr16:87438380-87438880:downstream_12526 | chr16 | 87425800 | 87438380 | + | MAP1LC3B | NM | 55 | 2,3 | 1 | -2 | 7,47E-02 | 3,35E-01 | 4 | 7 | 1 | 1 | 99 | 123 | 99 | 109 | 2,3 | 0,1 | 2,2 |
| chr14:75548317-75548817:downstream_11232 | chr14 | 75548817 | 75593778 | - | NEK9 | NM | 36 | 1,6 | 1 | -2 | 7,47E-02 | 3,35E-01 | 10 | 13 | 10 | 11 | 37 | 26 | 121 | 62 | 0,1 | -1,5 | 1,6 |
| chr11:3695739-3696239:downstream_18179 | chr11 | 3696239 | 3819022 | - | NUP98 | NM | 14 | -3,3 | 2 | 2 | 7,48E-02 | NA | 0 | 0 | 4 | 11 | 9 | 8 | 33 | 46 | -Inf | -2,2 | -Inf |
| chr14:76116732-76117232:downstream_11237 | chr14 | 76117232 | 76127538 | - | C14orf1 | NM | 23 | 1,8 | 1 | -2 | 7,51E-02 | NA | 25 | 10 | 10 | 9 | 26 | 19 | 35 | 47 | 0,9 | -0,8 | 1,7 |
| chr2:232318958-232319458:downstream_2474 | chr2 | 232319458 | 232329205 | - | NCL | NM | 125 | 1,2 | 1 | -2 | 7,51E-02 | 3,36E-01 | 59 | 25 | 22 | 17 | 230 | 201 | 223 | 220 | 1,1 | 0,0 | 1,1 |
| chr1:52815764-52816264:downstream_133 | chr1 | 52816264 | 52831877 | - | CC2D1B | NM | 7 | 3,3 | 2 | -2 | 7,53E-02 | NA | 3 | 15 | 3 | 0 | 8 | 9 | 7 | 13 | 2,8 | -0,3 | 3,0 |
| chr9:86582497-86582997:downstream_26348 | chr9 | 86582997 | 86595569 | - | HNRNPK | NM | 49 | 1,2 | 1 | -2 | 7,56E-02 | 3,37E-01 | 80 | 58 | 43 | 41 | 35 | 36 | 50 | 49 | 0,7 | -0,5 | 1,2 |
| chrX:15841382-15841882:downstream_7611 | chrX | 15808573 | 15841382 | + | ZRSR2 | NM | 15 | 2,5 | 1 | -2 | 7,56E-02 | NA | 11 | 3 | 3 | 1 | 28 | 15 | 28 | 28 | 2,0 | -0,4 | 2,4 |
| chr14:24028271-24028771:downstream_19146 | chr14 | 24028771 | 24037045 | - | AP1G2 | NM | 23 | 2,2 | 1 | -2 | 7,62E-02 | NA | 2 | 9 | 1 | 3 | 39 | 15 | 63 | 52 | 1,3 | -1,1 | 2,3 |
| chr17:39847898-39848398:downstream_12565 | chr17 | 39845126 | 39847898 | + | EIF1 | NM | 119 | -1,1 | 1 | 2 | 7,65E-02 | 3,40E-01 | 43 | 26 | 49 | 29 | 279 | 254 | 121 | 153 | -0,2 | 1,0 | -1,1 |
| chr1:211548286-211548786:downstream_17351 | chr1 | 211499956 | 211548286 | + | TRAF5 | NM | 5 | 4,2 | 2 | -2 | 7,65E-02 | NA | 2 | 2 | 10 | 3 | 0 | 0 | 10 | 11 | -1,9 | -Inf | Inf |
| chr19:52795976-52796476:downstream_14556 | chr19 | 52772823 | 52795976 | + | ZNF766 | NM | 35 | 1,9 | 1 | -2 | 7,70E-02 | 3,41E-01 | 6 | 19 | 1 | 3 | 79 | 64 | 58 | 51 | 2,4 | 0,4 | 2,0 |

| | | | | | | | | | | | | | | | | | | | | | | | |
|---|---|---|---|---|---|---|---|---|---|---|---|---|---|---|---|---|---|---|---|---|---|---|---|
| chr3:158383702-158384202:downstream_3254 | chr3 | 158384202 | 158390482 | - | LXN | NM | 54 | -1,6 | 1 | 2 | 7,75E-02 | 3,43E-01 | 4 | 3 | 19 | 7 | 81 | 103 | 125 | 94 | -1,9 | -0,3 | -1,7 |
| chr12:4796720-4797220:downstream_9984 | chr12 | 4758263 | 4796720 | + | NDUFA9 | NM | 35 | -2,9 | 2 | 2 | 7,83E-02 | 3,45E-01 | 1 | 0 | 7 | 4 | 63 | 42 | 81 | 84 | -3,8 | -0,7 | -3,1 |
| chr16:11772435-11772935:downstream_20262 | chr16 | 11772935 | 11836734 | - | TXNDC11 | NM | 14 | 2,3 | 1 | -2 | 7,84E-02 | NA | 2 | 9 | 4 | 3 | 9 | 13 | 42 | 34 | 0,5 | -1,8 | 2,3 |
| chr10:99195165-99195665:downstream_8658 | chr10 | 99195665 | 99205768 | - | EXOSC1 | NM | 27 | 1,8 | 1 | -2 | 7,85E-02 | NA | 19 | 16 | 4 | 2 | 50 | 58 | 29 | 37 | 2,4 | 0,7 | 1,7 |
| chr12:98995778-98996278:downstream_18829 | chr12 | 98987402 | 98995778 | + | SLC25A3 | NM | 72 | 1,1 | 1 | -2 | 7,86E-02 | 3,45E-01 | 85 | 82 | 28 | 15 | 113 | 124 | 64 | 64 | 2,0 | 0,9 | 1,1 |
| chr15:69747884-69748384:downstream_19703 | chr15 | 69745158 | 69747884 | + | RPLP1 | NM | 1244 | -0,9 | 1 | 2 | 7,87E-02 | 3,45E-01 | 38 | 29 | 29 | 19 | 4175 | 3039 | 1314 | 1306 | 0,5 | 1,5 | -1,0 |
| chr1:61928460-61928960:downstream_17018 | chr1 | 61542945 | 61928460 | + | NFIA | NM | 4 | 4,2 | 2 | -2 | 7,89E-02 | NA | 0 | 11 | 1 | 2 | 2 | 0 | 6 | 12 | 1,5 | -2,9 | 4,3 |
| chr12:104744085-104744585:downstream_18900 | chr12 | 104609556 | 104744085 | + | TXNRD1 | NM | 545 | -1,0 | 1 | 2 | 7,93E-02 | 3,46E-01 | 10 | 11 | 28 | 57 | 671 | 705 | 1312 | 1564 | -2,1 | -1,1 | -1,0 |
| chr2:47386720-47387220:downstream_21788 | chr2 | 47387220 | 47404075 | - | CALM2 | NM | 174 | 1,0 | 1 | -2 | 7,93E-02 | 3,46E-01 | 127 | 118 | 61 | 57 | 320 | 206 | 195 | 307 | 1,0 | 0,1 | 1,0 |
| chr2:101886778-101887278:downstream_2146 | chr2 | 101869344 | 101886778 | + | CNOT11 | NM | 19 | 1,9 | 1 | -2 | 7,94E-02 | NA | 24 | 6 | 7 | 6 | 22 | 19 | 32 | 35 | 1,2 | -0,7 | 1,9 |
| chr17:40696466-40696966:downstream_13125 | chr17 | 40687950 | 40696466 | + | NAGLU | NM | 28 | -2,2 | 1 | 2 | 7,95E-02 | NA | 2 | 0 | 4 | 11 | 41 | 47 | 51 | 65 | -2,6 | -0,4 | -2,2 |
| chr7:30673648-30674148:downstream_5532 | chr7 | 30634180 | 30673648 | + | GARS | NM | 64 | 1,5 | 1 | -2 | 7,97E-02 | 3,46E-01 | 9 | 3 | 10 | 5 | 45 | 70 | 185 | 189 | -0,3 | -1,7 | 1,4 |
| chr12:57923332-57923832:downstream_18535 | chr12 | 57923832 | 57941114 | - | DCTN2 | NM | 27 | 1,8 | 1 | -2 | 7,98E-02 | NA | 5 | 13 | 6 | 4 | 28 | 39 | 65 | 59 | 0,9 | -0,9 | 1,8 |
| chr20:33515735-33516235:downstream_14984 | chr20 | 33516235 | 33543601 | - | GSS | NM | 14 | 2,8 | 2 | -2 | 8,00E-02 | NA | 5 | 4 | 3 | 0 | 27 | 11 | 22 | 42 | 1,7 | -0,8 | 2,4 |
| chr3:101579869-101580369:downstream_23307 | chr3 | 101546833 | 101579869 | + | NFKBIZ | NM | 24 | 2,1 | 1 | -2 | 8,00E-02 | NA | 17 | 6 | 3 | 2 | 41 | 54 | 39 | 33 | 2,4 | 0,4 | 2,0 |
| chr1:94334513-94335013:downstream_836 | chr1 | 94335013 | 94344762 | - | DNTTIP2 | NM | 26 | 1,6 | 1 | -2 | 8,02E-02 | NA | 24 | 29 | 8 | 4 | 37 | 49 | 28 | 32 | 2,1 | 0,5 | 1,6 |
| chr12:110889790-110890290:downstream_18584 | chr12 | 110890290 | 110906526 | - | GPN3 | NM | 9 | 3,3 | 2 | -2 | 8,03E-02 | NA | 8 | 6 | 3 | 0 | 20 | 1 | 14 | 20 | 2,3 | -0,7 | 3,0 |
| chr12:56214243-56214743:downstream_10247 | chr12 | 56214743 | 56223420 | - | DNAJC14 | NM | 45 | 1,4 | 1 | -2 | 8,04E-02 | 3,47E-01 | 21 | 38 | 11 | 20 | 42 | 70 | 63 | 97 | 0,9 | -0,5 | 1,4 |
| chr2:42577144-42577644:downstream_1949 | chr2 | 42577644 | 42588356 | - | COX7A2L | NM | 69 | 1,2 | 1 | -2 | 8,04E-02 | 3,47E-01 | 46 | 25 | 11 | 16 | 130 | 105 | 99 | 120 | 1,4 | 0,1 | 1,3 |
| chr7:55537800-55538300:downstream_25801 | chr7 | 55538300 | 55640200 | - | VOPP1 | NM | 20 | -1,8 | 1 | 2 | 8,06E-02 | NA | 6 | 4 | 54 | 36 | 7 | 10 | 18 | 25 | -3,2 | -1,4 | -1,9 |
| chr1:19541657-19542157:downstream_16691 | chr1 | 19542157 | 19578053 | - | EMC1 | NM | 8 | -3,0 | 2 | 2 | 8,07E-02 | NA | 2 | 0 | 7 | 9 | 11 | 11 | 13 | 14 | -3,3 | -0,3 | -3,0 |
| chr6:75962137-75962637:downstream_25312 | chr6 | 75962637 | 75994632 | - | TMEM30A | NM | 82 | -1,9 | 1 | 2 | 8,10E-02 | 3,49E-01 | 2 | 0 | 6 | 8 | 123 | 140 | 191 | 189 | -2,5 | -0,5 | -2,0 |
| chr12:27124005-27124505:downstream_9836 | chr12 | 27124505 | 27167339 | - | TM7SF3 | NM | 66 | 1,6 | 1 | -2 | 8,14E-02 | 3,49E-01 | 5 | 11 | 1 | 5 | 117 | 97 | 141 | 155 | 1,3 | -0,5 | 1,8 |
| chr20:25653330-25653830:downstream_22613 | chr20 | 25653830 | 25677540 | - | ZNF337 | NM | 8 | 3,3 | 2 | -2 | 8,15E-02 | NA | 12 | 9 | 1 | 0 | 15 | 14 | 8 | 7 | 3,9 | 1,0 | 2,9 |
| chr14:45376460-45376960:downstream_11122 | chr14 | 45366506 | 45376460 | + | C14orf28 | NM | 17 | 2,6 | 1 | -2 | 8,16E-02 | NA | 26 | 27 | 0 | 2 | 39 | 25 | 4 | 10 | 5,0 | 2,2 | 2,8 |
| chr15:43619473-43619973:downstream_11689 | chr15 | 43619973 | 43622820 | - | LCMT2 | NM | 13 | 2,7 | 2 | -2 | 8,21E-02 | NA | 7 | 20 | 1 | 1 | 22 | 30 | 7 | 20 | 3,6 | 1,0 | 2,6 |
| chr15:25339276-25339776:downstream_11569 | chr15 | 25339182 | 25339276 | + | SNORD116-24 | NR | 9 | -2,7 | 2 | 2 | 8,23E-02 | NA | 2 | 1 | 15 | 14 | 6 | 9 | 14 | 11 | -3,5 | -0,8 | -2,7 |
| chr10:101947623-101948123:downstream_8333 | chr10 | 101948123 | 101989344 | - | CHUK | NM | 34 | 1,7 | 1 | -2 | 8,24E-02 | 3,53E-01 | 9 | 44 | 10 | 13 | 25 | 53 | 64 | 52 | 1,2 | -0,6 | 1,8 |
| chr5:82348164-82348664:downstream_4332 | chr5 | 82348664 | 82373272 | - | TMEM167A | NM | 26 | 1,7 | 1 | -2 | 8,26E-02 | NA | 24 | 41 | 7 | 6 | 30 | 51 | 22 | 27 | 2,4 | 0,7 | 1,6 |
| chr11:114183238-114183738:downstream_9658 | chr11 | 114166534 | 114183238 | + | NNMT | NM | 26 | -2,9 | 2 | 2 | 8,29E-02 | NA | 0 | 1 | 31 | 18 | 12 | 7 | 71 | 66 | -5,7 | -2,9 | -2,8 |
| chrX:139005806-139006306:downstream_7962 | chrX | 139006306 | 139006390 | - | MIR505 | NR | 5 | -4,1 | 2 | 2 | 8,35E-02 | NA | 4 | 0 | 0 | 12 | 15 | 6 | 0 | 3 | -1,6 | 2,6 | -4,2 |
| chr2:73988824-73989324:downstream_2038 | chr2 | 73989324 | 74007284 | - | DUSP11 | NM | 40 | 1,5 | 1 | -2 | 8,36E-02 | 3,57E-01 | 24 | 36 | 4 | 9 | 70 | 79 | 56 | 41 | 2,2 | 0,6 | 1,5 |
| chr10:127524408-127524908:downstream_8364 | chr10 | 127524908 | 127569884 | - | DHX32 | NM | 50 | 1,7 | 1 | -2 | 8,37E-02 | 3,57E-01 | 6 | 7 | 10 | 3 | 67 | 26 | 138 | 147 | -0,1 | -1,6 | 1,6 |
| chr19:4795571-4796071:downstream_13906 | chr19 | 4791727 | 4795571 | + | FEM1A | NM | 22 | 2,1 | 1 | -2 | 8,38E-02 | NA | 9 | 14 | 3 | 1 | 51 | 41 | 28 | 30 | 2,6 | 0,7 | 2,0 |
| chrX:151121095-151121595:downstream_7569 | chrX | 151121595 | 151143151 | - | GABRE | NM | 18 | 2,1 | 1 | -2 | 8,43E-02 | NA | 7 | 9 | 1 | 2 | 33 | 24 | 28 | 39 | 2,0 | -0,2 | 2,2 |
| chr5:143537222-143537722:downstream_24443 | chr5 | 143537722 | 143550278 | - | YIPF5 | NM | 17 | 1,9 | 1 | -2 | 8,43E-02 | NA | 25 | 18 | 13 | 4 | 18 | 14 | 19 | 22 | 1,4 | -0,4 | 1,8 |
| chr12:109038384-109038884:downstream_18525 | chr12 | 109038884 | 109125326 | - | CORO1C | NM | 58 | 1,5 | 1 | -2 | 8,44E-02 | 3,58E-01 | 3 | 11 | 4 | 8 | 66 | 50 | 175 | 144 | 0,2 | -1,5 | 1,6 |
| chr10:105788991-105789491:downstream_17774 | chr10 | 105726942 | 105788991 | + | SLK | NM | 9 | 2,6 | 2 | -2 | 8,46E-02 | NA | 14 | 4 | 1 | 5 | 3 | 12 | 18 | 16 | 1,5 | -1,2 | 2,7 |
| chr2:139330805-139331305:downstream_1721 | chr2 | 139259349 | 139330805 | + | SPOPL | NM | 7 | 3,0 | 2 | -2 | 8,47E-02 | NA | 5 | 14 | 11 | 5 | 1 | 2 | 10 | 8 | 0,2 | -2,7 | 2,9 |
| chr5:147822399-147822899:downstream_24122 | chr5 | 147763497 | 147822399 | + | FBXO38 | NM | 6 | -3,7 | 2 | 2 | 8,48E-02 | NA | 0 | 0 | 3 | 8 | 6 | 5 | 11 | 16 | -Inf | -1,4 | -Inf |
| chr11:112104695-112105195:downstream_9648 | chr11 | 112097087 | 112104695 | + | PTS | NM | 23 | 2,1 | 1 | -2 | 8,49E-02 | NA | 28 | 3 | 10 | 2 | 31 | 24 | 43 | 42 | 1,4 | -0,6 | 2,0 |
| chr16:71891236-71891736:downstream_12460 | chr16 | 71879893 | 71891236 | + | ATXN1L | NM | 16 | 2,3 | 1 | -2 | 8,52E-02 | NA | 7 | 10 | 3 | 1 | 25 | 30 | 25 | 28 | 2,2 | 0,1 | 2,1 |
| chr6:28201264-28201764:downstream_25367 | chr6 | 28193028 | 28201264 | + | ZSCAN9 | NM | 9 | 3,2 | 2 | -2 | 8,52E-02 | NA | 11 | 6 | 0 | 1 | 15 | 20 | 11 | 10 | 4,3 | 0,8 | 3,5 |
| chr1:151954885-151955385:downstream_1008 | chr1 | 151955385 | 151966714 | - | S100A10 | NM | 398 | 1,0 | 1 | -2 | 8,54E-02 | 3,61E-01 | 46 | 27 | 10 | 18 | 1017 | 677 | 743 | 642 | 1,4 | 0,3 | 1,1 |
| chr8:118811101-118811601:downstream_6344 | chr8 | 118811601 | 119124058 | - | EXT1 | NM | 32 | -3,0 | 2 | 2 | 8,56E-02 | 3,61E-01 | 0 | 0 | 13 | 3 | 13 | 23 | 96 | 111 | -Inf | -2,5 | -Inf |
| chr4:41962824-41963324:downstream_3363 | chr4 | 41937136 | 41962824 | + | TMEM33 | NM | 24 | 1,7 | 1 | -2 | 8,57E-02 | NA | 31 | 20 | 10 | 5 | 35 | 32 | 39 | 21 | 1,8 | 0,2 | 1,6 |
| chr9:115427591-115428091:downstream_26361 | chr9 | 115249247 | 115427591 | + | KIAA1958 | NM | 4 | -4,2 | 2 | 2 | 8,57E-02 | NA | 0 | 0 | 7 | 7 | 1 | 5 | 6 | 7 | -Inf | -1,1 | -Inf |
| chr9:6010518-6011018:downstream_26473 | chr9 | 6011018 | 6015640 | - | RANBP6 | NM | 20 | 2,1 | 1 | -2 | 8,61E-02 | NA | 10 | 11 | 0 | 3 | 33 | 42 | 32 | 29 | 2,6 | 0,3 | 2,3 |
| chr8:124231695-124232195:downstream_6700 | chr8 | 124232195 | 124253638 | - | C8orf76 | NM | 9 | 2,5 | 1 | -2 | 8,62E-02 | NA | 7 | 7 | 3 | 2 | 9 | 10 | 13 | 20 | 1,7 | -0,8 | 2,5 |
| chr8:87480178-87480678:downstream_6327 | chr8 | 87354993 | 87480178 | + | WWP1 | NM | 25 | 1,7 | 1 | -2 | 8,63E-02 | NA | 7 | 11 | 15 | 21 | 7 | 13 | 75 | 50 | -1,0 | -2,7 | 1,7 |
| chr17:695741-696241:downstream_12720 | chr17 | 685512 | 695741 | + | RNMTL1 | NM | 34 | 2,1 | 1 | -2 | 8,66E-02 | 3,65E-01 | 5 | 17 | 3 | 1 | 98 | 50 | 51 | 43 | 2,6 | 0,7 | 2,0 |
| chr8:87573726-87574226:downstream_6326 | chr8 | 87526655 | 87573726 | + | CPNE3 | NM | 30 | 1,6 | 1 | -2 | 8,69E-02 | 3,65E-01 | 11 | 13 | 11 | 6 | 18 | 46 | 68 | 67 | 0,5 | -1,1 | 1,6 |
| chr2:85568577-85569077:downstream_2073 | chr2 | 85569077 | 85581821 | - | RETSAT | NM | 89 | -1,2 | 1 | 2 | 8,71E-02 | 3,65E-01 | 48 | 44 | 15 | 31 | 180 | 285 | 42 | 64 | 1,0 | 2,1 | -1,1 |
| chr8:146014653-146015153:downstream_26101 | chr8 | 146015153 | 146017805 | - | RPL8 | NM | 92 | 1,7 | 1 | -2 | 8,74E-02 | 3,65E-01 | 7 | 11 | 1 | 2 | 203 | 200 | 132 | 180 | 2,2 | 0,4 | 1,8 |
| chr19:50058224-50058724:downstream_21307 | chr19 | 50058724 | 50083829 | - | NOSIP | NM | 7 | 3,1 | 2 | -2 | 8,74E-02 | NA | 6 | 19 | 1 | 2 | 12 | 2 | 8 | 7 | 3,0 | -0,1 | 3,1 |
| chr9:32553023-32553523:downstream_26406 | chr9 | 32553523 | 32573182 | - | NDUFB6 | NM | 15 | -2,2 | 1 | 2 | 8,75E-02 | NA | 2 | 7 | 4 | 13 | 17 | 46 | 14 | 16 | -1,0 | 1,1 | -2,1 |
| chr5:177638184-177638684:downstream_24165 | chr5 | 177631507 | 177638184 | + | HNRNPAB | NM | 94 | 1,1 | 1 | -2 | 8,80E-02 | 3,65E-01 | 24 | 33 | 11 | 9 | 190 | 191 | 146 | 148 | 1,5 | 0,4 | 1,1 |
| chr17:42396492-42396992:downstream_20717 | chr17 | 42396992 | 42402217 | - | SLC25A39 | NM | 35 | 1,7 | 1 | -2 | 8,80E-02 | 3,65E-01 | 13 | 16 | 4 | 2 | 77 | 71 | 39 | 61 | 2,2 | 0,6 | 1,6 |
| chr11:46877767-46878267:downstream_8878 | chr11 | 46878267 | 46940173 | - | LRP4 | NM | 48 | 1,5 | 1 | -2 | 8,82E-02 | 3,65E-01 | 4 | 10 | 7 | 9 | 30 | 50 | 132 | 140 | -0,2 | -1,8 | 1,5 |
| chr22:37965410-37965910:downstream_15593 | chr22 | 37956470 | 37965410 | + | CDC42EP1 | NM | 7 | -2,9 | 2 | 2 | 8,88E-02 | NA | 3 | 0 | 7 | 7 | 11 | 12 | 10 | 3 | -2,1 | 0,8 | -3,0 |
| chr2:208626809-208627309:downstream_2392 | chr2 | 208627309 | 208634143 | - | FZD5 | NM | 31 | 1,4 | 1 | -2 | 8,91E-02 | 3,68E-01 | 16 | 18 | 14 | 28 | 18 | 21 | 70 | 62 | -0,3 | -1,7 | 1,5 |
| chr13:23902461-23902961:downstream_19082 | chr13 | 23902961 | 24007867 | - | SACS | NM | 15 | 2,3 | 1 | -2 | 9,00E-02 | NA | 5 | 5 | 1 | 2 | 21 | 19 | 32 | 32 | 1,7 | -0,7 | 2,3 |
| chr12:57031458-57031958:downstream_10261 | chr12 | 57031958 | 57039852 | - | ATP5B | NM | 266 | 1,1 | 1 | -2 | 9,03E-02 | 3,72E-01 | 20 | 33 | 11 | 10 | 490 | 635 | 426 | 508 | 1,3 | 0,3 | 1,1 |
| chr11:13409042-13409542:downstream_17899 | chr11 | 13409542 | 13484844 | - | BTBD10 | NM | 17 | 1,9 | 1 | -2 | 9,05E-02 | NA | 7 | 17 | 4 | 7 | 23 | 13 | 32 | 35 | 1,0 | -0,9 | 2,0 |
| chr8:121407582-121408082:downstream_6693 | chr8 | 121408082 | 121457647 | - | MRPL13 | NM | 17 | 2,4 | 1 | -2 | 9,09E-02 | NA | 17 | 1 | 4 | 1 | 21 | 29 | 39 | 26 | 1,9 | -0,4 | 2,2 |
| chr2:191557492-191557992:downstream_2357 | chr2 | 191513847 | 191557492 | + | NAB1 | NM | 13 | -3,1 | 2 | 2 | 9,12E-02 | NA | 1 | 0 | 7 | 5 | 17 | 17 | 25 | 29 | -3,9 | -0,6 | -3,2 |
| chr12:123773155-123773655:downstream_18812 | chr12 | 123773655 | 123834988 | - | SBNO1 | NM | 12 | 2,7 | 2 | -2 | 9,12E-02 | NA | 1 | 9 | 6 | 18 | 3 | 0 | 25 | 34 | -1,3 | -4,2 | 2,9 |
| chr19:40028946-40029446:downstream_14288 | chr19 | 40029446 | 40030838 | - | EID2 | NM | 54 | 1,3 | 1 | -2 | 9,14E-02 | 3,76E-01 | 17 | 34 | 6 | 10 | 89 | 116 | 83 | 80 | 1,7 | 0,3 | 1,4 |
| chr13:51544596-51545096:downstream_19078 | chr13 | 51483813 | 51544596 | + | RNASEH2B | NM | 6 | -3,4 | 2 | 2 | 9,15E-02 | NA | 2 | 0 | 8 | 11 | 2 | 11 | 3 | 11 | -3,6 | -0,1 | -3,5 |
| chr2:97513223-97513723:downstream_1526 | chr2 | 97513723 | 97523756 | - | ANKRD39 | NM | 12 | -2,3 | 1 | 2 | 9,16E-02 | NA | 2 | 2 | 10 | 7 | 22 | 16 | 22 | 13 | -2,2 | 0,1 | -2,3 |
| chr17:62534062-62534562:downstream_12675 | chr17 | 62503157 | 62534062 | + | CEP95 | NM | 34 | 1,7 | 1 | -2 | 9,17E-02 | 3,76E-01 | 39 | 10 | 6 | 7 | 59 | 60 | 38 | 55 | 2,0 | 0,4 | 1,6 |
| chr17:71204645-71205145:downstream_13334 | chr17 | 71189172 | 71204645 | + | COG1 | NM | 14 | 2,1 | 1 | -2 | 9,17E-02 | NA | 7 | 16 | 6 | 2 | 12 | 25 | 22 | 24 | 1,7 | -0,3 | 2,0 |
| chr11:82645699-82646199:downstream_9553 | chr11 | 82612736 | 82645699 | + | DDIAS | NM | 8 | 2,6 | 2 | -2 | 9,19E-02 | NA | 7 | 13 | 1 | 2 | 14 | 5 | 14 | 10 | 2,4 | -0,3 | 2,7 |
| chr12:107367813-107368313:downstream_10400 | chr12 | 107349543 | 107367813 | + | TMEM263 | NM | 60 | 1,3 | 1 | -2 | 9,20E-02 | 3,76E-01 | 41 | 22 | 36 | 21 | 77 | 36 | 139 | 105 | 0,2 | -1,1 | 1,3 |
| chr12:120119429-120119929:downstream_10443 | chr12 | 120105760 | 120119429 | + | PRKAB1 | NM | 18 | 2,8 | 2 | -2 | 9,21E-02 | NA | 15 | 10 | 1 | 0 | 41 | 47 | 13 | 14 | 4,2 | 1,7 | 2,4 |
| chr11:17353070-17353570:downstream_8854 | chr11 | 17298285 | 17353070 | + | NUCB2 | NM | 25 | -1,8 | 1 | 2 | 9,25E-02 | NA | 7 | 0 | 25 | 19 | 28 | 26 | 46 | 44 | -2,6 | -0,7 | -1,9 |
| chr2:68405488-68405988:downstream_1641 | chr2 | 68405988 | 68479651 | - | PPP3R1 | NM | 15 | 2,1 | 1 | -2 | 9,29E-02 | NA | 6 | 7 | 6 | 2 | 11 | 19 | 33 | 34 | 0,8 | -1,2 | 2,0 |
| chr12:13066600-13067100:downstream_10073 | chr12 | 13043955 | 13066600 | + | GPRC5A | NM | 204 | -0,9 | 1 | 2 | 9,29E-02 | 3,79E-01 | 43 | 77 | 121 | 112 | 270 | 359 | 301 | 349 | -1,0 | 0,0 | -0,9 |
| chr12:58122139-58122639:downstream_10284 | chr12 | 58120022 | 58122139 | + | AGAP2-AS1 | NR | 10 | 2,4 | 1 | -2 | 9,31E-02 | NA | 10 | 13 | 0 | 5 | 11 | 12 | 15 | 14 | 2,2 | -0,4 | 2,6 |
| chr5:177037346-177037846:downstream_4610 | chr5 | 177027118 | 177037346 | + | B4GALT7 | NM | 11 | 2,1 | 1 | -2 | 9,31E-02 | NA | 16 | 10 | 10 | 7 | 8 | 4 | 19 | 15 | 0,6 | -1,5 | 2,1 |
| chr12:50229325-50229825:downstream_10158 | chr12 | 50229825 | 50236912 | - | BCDIN3D | NM | 15 | 3,1 | 2 | -2 | 9,32E-02 | NA | 15 | 9 | 0 | 0 | 52 | 30 | 4 | 7 | Inf | 2,8 | Inf |
| chr9:4708398-4708898:downstream_6973 | chr9 | 4679552 | 4708398 | + | CDC37L1 | NM | 9 | 2,8 | 2 | -2 | 9,35E-02 | NA | 6 | 9 | 1 | 1 | 15 | 11 | 18 | 9 | 2,7 | -0,1 | 2,8 |
| chrX:123236505-123237005:downstream_27053 | chrX | 123094409 | 123236505 | + | STAG2 | NM | 26 | 1,9 | 1 | -2 | 9,39E-02 | NA | 7 | 3 | 25 | 9 | 10 | 3 | 67 | 82 | -1,7 | -3,6 | 1,8 |
| chr15:99675800-99676300:downstream_19799 | chr15 | 99645285 | 99675800 | + | SYNM | NM | 11 | 2,2 | 1 | -2 | 9,44E-02 | NA | 10 | 12 | 6 | 2 | 12 | 13 | 14 | 21 | 1,6 | -0,5 | 2,1 |
| chr17:34851098-34851598:downstream_20599 | chr17 | 34851598 | 34891305 | - | MYO19 | NM | 36 | 1,4 | 1 | -2 | 9,44E-02 | 3,81E-01 | 20 | 45 | 18 | 26 | 29 | 35 | 63 | 53 | 0,6 | -0,9 | 1,4 |
| chr3:101399433-101399933:downstream_3082 | chr3 | 101399933 | 101405563 | - | RPL24 | NM | 200 | 0,9 | 1 | -2 | 9,44E-02 | 3,81E-01 | 75 | 71 | 19 | 20 | 437 | 506 | 249 | 224 | 1,9 | 1,0 | 0,9 |
| chr1:228644564-228645064:downstream_1402 | chr1 | 228645064 | 228645560 | - | HIST3H2A | NM | 92 | 1,1 | 1 | -2 | 9,45E-02 | 3,81E-01 | 23 | 18 | 14 | 22 | 120 | 105 | 259 | 176 | 0,2 | -1,0 | 1,1 |
| chr1:200373574-200374074:downstream_17429 | chr1 | 200374074 | 200379186 | - | ZNF281 | NM | 44 | 1,4 | 1 | -2 | 9,46E-02 | 3,81E-01 | 13 | 20 | 7 | 8 | 74 | 62 | 103 | 63 | 1,1 | -0,3 | 1,4 |
| chr17:30189689-30190189:downstream_12965 | chr17 | 30190189 | 30228729 | - | UTP6 | NM | 59 | -1,4 | 1 | 2 | 9,48E-02 | 3,81E-01 | 7 | 3 | 10 | 18 | 100 | 110 | 117 | 108 | -1,4 | -0,1 | -1,3 |
| chr1:246931440-246931940:downstream_1459 | chr1 | 246887377 | 246931440 | + | SCCPDH | NM | 8 | -3,4 | 2 | 2 | 9,50E-02 | NA | 1 | 0 | 11 | 2 | 10 | 10 | 14 | 13 | -4,1 | -0,5 | -3,6 |
| chr22:50963209-50963709:downstream_22858 | chr22 | 50946644 | 50963209 | + | NCAPH2 | NM | 5 | 3,8 | 2 | -2 | 9,50E-02 | NA | 8 | 15 | 1 | 0 | 5 | 4 | 1 | 4 | 4,0 | 0,7 | 3,4 |
| chr5:111497814-111498314:downstream_4064 | chr5 | 111498314 | 111755010 | - | EPB41L4A | NM | 12 | 2,2 | 1 | -2 | 9,52E-02 | NA | 6 | 15 | 6 | 10 | 11 | 2 | 25 | 22 | 0,4 | -1,9 | 2,3 |
| chr21:35275256-35275756:downstream_15302 | chr21 | 35275756 | 35288158 | - | ATP5O | NM | 47 | 2,2 | 1 | -2 | 9,54E-02 | 3,83E-01 | 7 | 1 | 1 | 1 | 93 | 79 | 102 | 91 | 1,9 | -0,2 | 2,1 |
| chrX:48379202-48379702:downstream_26987 | chrX | 48367346 | 48379202 | + | PORCN | NM | 21 | 1,8 | 1 | -2 | 9,55E-02 | NA | 20 | 18 | 10 | 1 | 34 | 29 | 29 | 25 | 1,9 | 0,2 | 1,7 |
| chr4:74901811-74902311:downstream_3740 | chr4 | 74902311 | 74904490 | - | CXCL3 | NM | 5 | 3,9 | 2 | -2 | 9,57E-02 | NA | 1 | 4 | 8 | 9 | 0 | 0 | 10 | 8 | -1,9 | -Inf | Inf |
| chr18:10552766-10553266:downstream_13564 | chr18 | 10525872 | 10552766 | + | NAPG | NM | 5 | 3,4 | 2 | -2 | 9,60E-02 | NA | 5 | 8 | 6 | 2 | 2 | 0 | 11 | 4 | 0,8 | -2,6 | 3,5 |
| chr18:44680889-44681389:downstream_13507 | chr18 | 44681389 | 44702745 | - | IER3IP1 | NM | 24 | 1,7 | 1 | -2 | 9,60E-02 | NA | 11 | 8 | 7 | 3 | 35 | 25 | 60 | 40 | 0,9 | -0,7 | 1,6 |
| chr2:120974546-120975046:downstream_1708 | chr2 | 120975046 | 120980984 | - | TMEM185B | NM | 22 | 1,9 | 1 | -2 | 9,61E-02 | NA | 21 | 14 | 0 | 4 | 40 | 50 | 22 | 25 | 3,1 | 0,9 | 2,1 |
| chr6:85441715-85442215:downstream_4759 | chr6 | 85442215 | 85473954 | - | TBX18 | NM | 18 | 1,9 | 1 | -2 | 9,68E-02 | NA | 5 | 12 | 19 | 26 | 2 | 5 | 36 | 39 | -1,5 | -3,4 | 1,9 |
| chr7:42955961-42956461:downstream_5804 | chr7 | 42956461 | 42971805 | - | PSMA2 | NM | 22 | 1,9 | 1 | -2 | 9,71E-02 | NA | 21 | 9 | 8 | 3 | 44 | 15 | 32 | 47 | 1,4 | -0,4 | 1,8 |
| chr12:72074428-72074928:downstream_10317 | chr12 | 72057676 | 72074428 | + | THAP2 | NM | 8 | 3,6 | 2 | -2 | 9,73E-02 | NA | 15 | 1 | 0 | 0 | 15 | 25 | 3 | 9 | Inf | 1,8 | Inf |
| chr19:58317949-58318449:downstream_14708 | chr19 | 58318449 | 58326281 | - | ZNF552 | NM | 14 | 2,2 | 1 | -2 | 9,83E-02 | NA | 6 | 12 | 1 | 2 | 29 | 15 | 24 | 25 | 2,2 | -0,1 | 2,3 |
| chr9:130209452-130209952:downstream_7321 | chr9 | 130209952 | 130213711 | - | RPL12 | NM | 106 | 1,0 | 1 | -2 | 9,84E-02 | 3,92E-01 | 49 | 51 | 21 | 16 | 199 | 213 | 161 | 137 | 1,4 | 0,5 | 1,0 |
| chr22:39080766-39081266:downstream_15611 | chr22 | 39077953 | 39080766 | + | TOMM22 | NM | 35 | 1,7 | 1 | -2 | 9,85E-02 | 3,92E-01 | 13 | 8 | 6 | 1 | 67 | 70 | 63 | 54 | 1,7 | 0,2 | 1,5 |
| chr12:96260238-96260738:downstream_10368 | chr12 | 96252708 | 96260238 | + | SNRPF | NM | 19 | 2,2 | 1 | -2 | 9,87E-02 | NA | 6 | 15 | 3 | 1 | 40 | 33 | 36 | 16 | 2,6 | 0,5 | 2,1 |
| chr7:116559313-116559813:downstream_5613 | chr7 | 116502562 | 116559313 | + | CAPZA2 | NM | 42 | 1,6 | 1 | -2 | 9,87E-02 | 3,92E-01 | 11 | 2 | 10 | 5 | 39 | 39 | 125 | 107 | -0,1 | -1,6 | 1,4 |
| chr14:77583630-77584130:downstream_11245 | chr14 | 77564577 | 77583630 | + | CIPC | NM | 13 | 2,1 | 1 | -2 | 9,89E-02 | NA | 11 | 15 | 1 | 4 | 21 | 15 | 15 | 18 | 2,3 | 0,1 | 2,2 |
| chr3:101285290-101285790:downstream_3079 | chr3 | 101280679 | 101285290 | + | TRMT10C | NM | 44 | 1,4 | 1 | -2 | 9,90E-02 | 3,92E-01 | 41 | 28 | 6 | 7 | 89 | 93 | 25 | 61 | 2,5 | 1,1 | 1,4 |
| chrX:103440582-103441082:downstream_7534 | chrX | 103411155 | 103440582 | + | FAM199X | NM | 37 | 1,6 | 1 | -2 | 9,92E-02 | 3,92E-01 | 5 | 4 | 4 | 4 | 32 | 40 | 92 | 116 | 0,1 | -1,5 | 1,6 |
| chr7:43846941-43847441:downstream_25408 | chr7 | 43798271 | 43846941 | + | BLVRA | NM | 68 | 1,5 | 1 | -2 | 9,94E-02 | 3,92E-01 | 7 | 5 | 4 | 3 | 107 | 81 | 184 | 157 | 0,6 | -0,9 | 1,5 |
| chr15:35270041-35270541:downstream_11650 | chr15 | 35270541 | 35280497 | - | ZNF770 | NM | 47 | 1,5 | 1 | -2 | 9,95E-02 | 3,92E-01 | 24 | 15 | 3 | 6 | 74 | 127 | 77 | 51 | 2,2 | 0,7 | 1,6 |
| chr8:59495419-59495919:downstream_26111 | chr8 | 59465727 | 59495419 | + | SDCBP | NM | 36 | 1,4 | 1 | -2 | 1,00E-01 | 3,93E-01 | 14 | 19 | 7 | 6 | 59 | 63 | 68 | 55 | 1,4 | 0,0 | 1,4 |
| chrY:15032390-15032890:downstream_27152 | chrY | 15016018 | 15032390 | + | DDX3Y | NM | 8 | 2,9 | 2 | -2 | 1,00E-01 | NA | 6 | 5 | 0 | 2 | 11 | 9 | 19 | 9 | 2,7 | -0,5 | 3,2 |
| chr1:155111334-155111834:downstream_17253 | chr1 | 155107819 | 155111334 | + | SLC50A1 | NM | 32 | 2,2 | 1 | -2 | 1,00E-01 | 3,93E-01 | 4 | 14 | 3 | 0 | 76 | 72 | 49 | 37 | 2,7 | 0,8 | 1,9 |
| chr19:56088390-56088890:downstream_14664 | chr19 | 56088890 | 56092211 | - | ZNF579 | NM | 4 | 4,0 | 2 | -2 | 1,01E-01 | NA | 3 | 7 | 0 | 2 | 4 | 0 | 10 | 3 | 2,6 | -1,7 | 4,3 |
| chr1:180859415-180859915:downstream_17409 | chr1 | 180601145 | 180859415 | + | XPR1 | NM | 10 | 2,2 | 1 | -2 | 1,01E-01 | NA | 7 | 7 | 10 | 11 | 2 | 3 | 18 | 21 | -0,6 | -2,9 | 2,2 |
| chr10:96088148-96088648:downstream_17712 | chr10 | 95753745 | 96088148 | + | PLCE1 | NM | 9 | 2,3 | 1 | -2 | 1,01E-01 | NA | 6 | 4 | 4 | 7 | 5 | 3 | 19 | 25 | -0,2 | -2,5 | 2,4 |
| chr1:161006921-161007421:downstream_17368 | chr1 | 161007421 | 161008774 | - | TSTD1 | NM | 4 | -4,0 | 2 | 2 | 1,01E-01 | NA | 4 | 6 | 1 | 2 | 11 | 10 | 0 | 0 | 1,7 | Inf | -Inf |
| chr4:10441003-10441503:downstream_3619 | chr4 | 10441503 | 10459032 | - | ZNF518B | NM | 10 | 2,7 | 2 | -2 | 1,02E-01 | NA | 4 | 9 | 1 | 1 | 10 | 20 | 19 | 13 | 2,5 | -0,1 | 2,6 |
| chr17:19640797-19641297:downstream_20321 | chr17 | 19641297 | 19651746 | - | ALDH3A1 | NM | 790 | -1,4 | 1 | 2 | 1,02E-01 | 4,00E-01 | 5 | 0 | 6 | 7 | 1766 | 1576 | 1340 | 1623 | -1,3 | 0,2 | -1,5 |
| chr1:52838000-52838500:downstream_17050 | chr1 | 52838500 | 52870143 | - | ORC1 | NM | 6 | 2,9 | 2 | -2 | 1,03E-01 | NA | 4 | 8 | 4 | 4 | 0 | 5 | 13 | 12 | 0,5 | -2,4 | 2,9 |
| chr2:201688569-201689069:downstream_21775 | chr2 | 201676268 | 201688569 | + | BZW1 | NM | 27 | 1,5 | 1 | -2 | 1,03E-01 | NA | 21 | 17 | 8 | 7 | 32 | 49 | 36 | 49 | 1,4 | -0,1 | 1,4 |
| chr14:69253871-69254371:downstream_19444 | chr14 | 69254371 | 69262960 | - | ZFP36L1 | NM | 63 | -1,3 | 1 | 2 | 1,04E-01 | 4,04E-01 | 4 | 6 | 10 | 16 | 104 | 116 | 120 | 126 | -1,4 | -0,2 | -1,2 |
| chr9:95883270-95883770:downstream_7190 | chr9 | 95883770 | 95896570 | - | NINJ1 | NM | 27 | 2,6 | 2 | -2 | 1,04E-01 | NA | 7 | 10 | 1 | 0 | 86 | 53 | 28 | 30 | 3,5 | 1,3 | 2,3 |
| chrX:45605084-45605584:downstream_7686 | chrX | 45605584 | 45605694 | - | MIR221 | NR | 91 | -1,0 | 1 | 2 | 1,04E-01 | 4,04E-01 | 95 | 127 | 65 | 91 | 102 | 160 | 42 | 50 | 0,5 | 1,5 | -1,0 |
| chr18:71826204-71826704:downstream_13671 | chr18 | 71815745 | 71826204 | + | TIMM21 | NM | 16 | 2,3 | 1 | -2 | 1,04E-01 | NA | 2 | 15 | 0 | 3 | 19 | 34 | 24 | 35 | 2,4 | -0,2 | 2,5 |
| chr3:15687328-15687828:downstream_23026 | chr3 | 15642858 | 15687328 | + | BTD | NM | 10 | -2,3 | 1 | 2 | 1,04E-01 | NA | 3 | 2 | 13 | 8 | 20 | 10 | 10 | 14 | -2,0 | 0,3 | -2,3 |
| chr19:59095762-59096262:downstream_14732 | chr19 | 59086765 | 59095762 | + | CENPBD1P1 | NR | 22 | 1,8 | 1 | -2 | 1,05E-01 | NA | 10 | 5 | 1 | 5 | 33 | 27 | 56 | 43 | 1,2 | -0,7 | 1,9 |
| chr2:39009106-39009606:downstream_1942 | chr2 | 39005326 | 39009106 | + | GEMIN6 | NM | 95 | 1,0 | 1 | -2 | 1,05E-01 | 4,05E-01 | 35 | 31 | 21 | 16 | 187 | 122 | 180 | 172 | 0,8 | -0,2 | 1,0 |
| chr11:102249401-102249901:downstream_17895 | chr11 | 102217912 | 102249401 | + | BIRC2 | NM | 27 | 1,7 | 1 | -2 | 1,05E-01 | NA | 11 | 6 | 1 | 8 | 25 | 39 | 70 | 55 | 0,8 | -1,0 | 1,8 |
| chrX:76759855-76760355:downstream_26639 | chrX | 76760355 | 77041755 | - | ATRX | NM | 7 | -3,6 | 2 | 2 | 1,05E-01 | NA | 0 | 0 | 14 | 11 | 4 | 0 | 14 | 14 | -Inf | -2,8 | -Inf |
| chr1:28975246-28975746:downstream_593 | chr1 | 28975111 | 28975246 | + | RNU11 | NR | 65 | -1,3 | 1 | 2 | 1,05E-01 | 4,05E-01 | 6 | 6 | 11 | 8 | 119 | 176 | 96 | 98 | -0,8 | 0,6 | -1,4 |
| chr3:20026667-20027167:downstream_23375 | chr3 | 19988571 | 20026667 | + | RAB5A | NM | 64 | 1,2 | 1 | -2 | 1,05E-01 | 4,05E-01 | 16 | 10 | 13 | 10 | 65 | 93 | 139 | 168 | 0,2 | -1,0 | 1,2 |
| chr8:30642625-30643125:downstream_6517 | chr8 | 30643125 | 30670352 | - | PPP2CB | NM | 18 | 2,9 | 2 | -2 | 1,05E-01 | NA | 9 | 0 | 1 | 0 | 36 | 37 | 29 | 34 | 2,7 | 0,2 | 2,5 |
| chr22:32783061-32783561:downstream_15575 | chr22 | 32783561 | 32808274 | - | RTCB | NM | 29 | 1,4 | 1 | -2 | 1,05E-01 | 4,05E-01 | 19 | 16 | 8 | 8 | 35 | 46 | 45 | 57 | 1,1 | -0,3 | 1,4 |
| chr20:30194317-30194817:downstream_22441 | chr20 | 30193085 | 30194317 | + | ID1 | NM | 81 | -1,2 | 1 | 2 | 1,06E-01 | 4,05E-01 | 11 | 3 | 21 | 31 | 122 | 104 | 163 | 191 | -1,9 | -0,6 | -1,2 |
| chr5:132210582-132211082:downstream_4409 | chr5 | 132209357 | 132210582 | + | LEAP2 | NM | 25 | 1,7 | 1 | -2 | 1,06E-01 | NA | 77 | 45 | 8 | 7 | 28 | 14 | 11 | 6 | 3,0 | 1,3 | 1,7 |
| chr12:95413504-95414004:downstream_18712 | chr12 | 95414004 | 95467404 | - | NR2C1 | NM | 6 | 3,1 | 2 | -2 | 1,06E-01 | NA | 1 | 9 | 6 | 5 | 2 | 1 | 11 | 13 | -0,1 | -3,2 | 3,1 |
| chr16:2028751-2029251:downstream_12103 | chr16 | 2022063 | 2028751 | + | TBL3 | NM | 4 | 3,8 | 2 | -2 | 1,06E-01 | NA | 6 | 3 | 0 | 0 | 7 | 7 | 4 | 5 | Inf | 0,6 | Inf |
| chr6:170106402-170106902:downstream_5456 | chr6 | 170102256 | 170106402 | + | C6orf120 | NM | 36 | 1,5 | 1 | -2 | 1,07E-01 | 4,09E-01 | 51 | 22 | 7 | 3 | 65 | 85 | 26 | 30 | 2,8 | 1,4 | 1,4 |
| chr1:183594827-183595327:downstream_16490 | chr1 | 183595327 | 183605076 | - | ARPC5 | NM | 121 | 1,2 | 1 | -2 | 1,07E-01 | 4,09E-01 | 10 | 7 | 4 | 7 | 189 | 178 | 299 | 277 | 0,6 | -0,7 | 1,3 |
| chr12:6775142-6775642:downstream_18926 | chr12 | 6775642 | 6798738 | - | ZNF384 | NM | 13 | 2,2 | 1 | -2 | 1,07E-01 | NA | 3 | 14 | 3 | 3 | 22 | 9 | 26 | 25 | 1,5 | -0,8 | 2,2 |
| chr5:162872022-162872522:downstream_24048 | chr5 | 162864576 | 162872022 | + | CCNG1 | NM | 150 | 1,0 | 1 | -2 | 1,07E-01 | 4,09E-01 | 54 | 66 | 19 | 12 | 302 | 399 | 191 | 160 | 1,9 | 1,0 | 0,9 |
| chr22:31375380-31375880:downstream_22958 | chr22 | 31365196 | 31375380 | + | TUG1 | NR | 13 | -2,2 | 1 | 2 | 1,08E-01 | NA | 7 | 0 | 13 | 8 | 16 | 31 | 18 | 14 | -1,7 | 0,6 | -2,2 |
| chr11:65771617-65772117:downstream_17887 | chr11 | 65769549 | 65771617 | + | BANF1 | NM | 9 | 3,2 | 2 | -2 | 1,08E-01 | NA | 3 | 10 | 1 | 0 | 27 | 5 | 10 | 16 | 3,2 | 0,3 | 2,9 |
| chr13:44468068-44468568:downstream_19005 | chr13 | 44453419 | 44468068 | + | LACC1 | NM | 25 | 2,1 | 1 | -2 | 1,08E-01 | NA | 37 | 15 | 0 | 2 | 59 | 58 | 19 | 9 | 4,4 | 2,0 | 2,4 |
| chr1:204371991-204372491:downstream_1320 | chr1 | 204372491 | 204380945 | - | PPP1R15B | NM | 23 | 1,6 | 1 | -2 | 1,09E-01 | NA | 22 | 11 | 3 | 7 | 36 | 37 | 33 | 34 | 1,8 | 0,1 | 1,7 |
| chr3:196670884-196671384:downstream_3354 | chr3 | 196669493 | 196670884 | + | NCBP2-AS2 | NR | 5 | -3,0 | 2 | 2 | 1,09E-01 | NA | 5 | 3 | 4 | 7 | 7 | 12 | 3 | 1 | -0,5 | 2,4 | -2,9 |
| chr19:44681838-44682338:downstream_21618 | chr19 | 44669214 | 44681838 | + | ZNF226 | NM | 21 | 1,7 | 1 | -2 | 1,09E-01 | NA | 10 | 8 | 6 | 4 | 36 | 15 | 46 | 45 | 0,9 | -0,9 | 1,7 |
| chr9:132574720-132575220:downstream_7358 | chr9 | 132575220 | 132586441 | - | TOR1A | NM | 33 | 1,5 | 1 | -2 | 1,09E-01 | 4,15E-01 | 11 | 28 | 7 | 4 | 62 | 58 | 60 | 35 | 1,8 | 0,3 | 1,5 |
| chr14:21965774-21966274:downstream_11047 | chr14 | 21966274 | 21979517 | - | METTL3 | NM | 30 | 1,7 | 1 | -2 | 1,10E-01 | 4,15E-01 | 2 | 17 | 4 | 5 | 35 | 48 | 72 | 58 | 1,1 | -0,6 | 1,8 |
| chr17:57292611-57293111:downstream_13276 | chr17 | 57287370 | 57292611 | + | SMG8 | NM | 17 | 2,3 | 1 | -2 | 1,10E-01 | NA | 4 | 9 | 1 | 1 | 32 | 35 | 24 | 31 | 2,5 | 0,3 | 2,3 |
| chr12:111889517-111890017:downstream_9926 | chr12 | 111890017 | 112037480 | - | ATXN2 | NM | 22 | -2,3 | 1 | 2 | 1,10E-01 | NA | 2 | 0 | 21 | 15 | 23 | 5 | 64 | 48 | -4,5 | -2,0 | -2,4 |
| chr5:169031781-169032281:downstream_4569 | chr5 | 169010637 | 169031781 | + | SPDL1 | NM | 29 | 2,0 | 1 | -2 | 1,10E-01 | 4,16E-01 | 3 | 6 | 0 | 2 | 41 | 54 | 56 | 72 | 1,9 | -0,4 | 2,3 |
| chr17:43188507-43189007:downstream_13174 | chr17 | 43189007 | 43209900 | - | PLCD3 | NM | 47 | 1,3 | 1 | -2 | 1,10E-01 | 4,16E-01 | 19 | 23 | 15 | 6 | 70 | 74 | 102 | 66 | 1,0 | -0,2 | 1,2 |
| chr2:29025806-29026306:downstream_22158 | chr2 | 28974613 | 29025806 | + | PPP1CB | NM | 20 | 1,8 | 1 | -2 | 1,10E-01 | NA | 3 | 9 | 7 | 3 | 12 | 24 | 49 | 57 | 0,2 | -1,5 | 1,8 |

| | | | | | | | | | | | | | | | | | | | | | | | |
|---|---|---|---|---|---|---|---|---|---|---|---|---|---|---|---|---|---|---|---|---|---|---|---|
| chr5:1461041-1461541:downstream_4163 | chr5 | 1461541 | 1524076 | - | LPCAT1 | NM | 43 | -2,6 | 2 | 2 | 1,11E-01 | 4,16E-01 | 1 | 0 | 7 | 6 | 63 | 38 | 100 | 131 | -4,0 | -1,2 | -2,8 |
| chr4:186321390-186321890:downstream_3934 | chr4 | 186317839 | 186321390 | + | ANKRD37 | NM | 4 | -3,3 | 2 | 2 | 1,11E-01 | NA | 2 | 0 | 7 | 4 | 7 | 6 | 4 | 4 | -2,8 | 0,7 | -3,4 |
| chr14:20925511-20926011:downstream_19423 | chr14 | 20926011 | 20929771 | - | TMEM55B | NM | 14 | 1,8 | 1 | -2 | 1,11E-01 | NA | 15 | 20 | 11 | 6 | 13 | 10 | 18 | 19 | 1,1 | -0,7 | 1,8 |
| chr7:23349327-23349827:downstream_5525 | chr7 | 23349827 | 23509995 | - | IGF2BP3 | NM | 15 | 2,1 | 1 | -2 | 1,12E-01 | NA | 2 | 15 | 14 | 15 | 2 | 7 | 29 | 36 | -0,8 | -3,0 | 2,1 |
| chr2:178088685-178089185:downstream_2334 | chr2 | 178077421 | 178088685 | + | HNRNPA3 | NM | 60 | 1,1 | 1 | -2 | 1,12E-01 | 4,18E-01 | 113 | 131 | 46 | 57 | 45 | 27 | 36 | 29 | 1,2 | 0,1 | 1,1 |
| chr2:183789078-183789578:downstream_22105 | chr2 | 183789578 | 183903586 | - | NCKAP1 | NM | 73 | 1,2 | 1 | -2 | 1,12E-01 | 4,18E-01 | 5 | 13 | 25 | 19 | 41 | 38 | 227 | 220 | -1,3 | -2,5 | 1,2 |
| chr1:100616054-100616554:downstream_845 | chr1 | 100598705 | 100616054 | + | TRMT13 | NM | 11 | 2,3 | 1 | -2 | 1,12E-01 | NA | 6 | 7 | 1 | 2 | 13 | 20 | 14 | 27 | 2,0 | -0,3 | 2,3 |
| chr21:27096290-27096790:downstream_22623 | chr21 | 27096790 | 27107965 | - | ATP5J | NM | 13 | 2,5 | 2 | -2 | 1,12E-01 | NA | 6 | 18 | 1 | 1 | 15 | 38 | 13 | 14 | 3,4 | 1,0 | 2,4 |
| chr11:766722-767222:downstream_8983 | chr11 | 767222 | 777487 | - | PDDC1 | NM | 34 | 1,4 | 1 | -2 | 1,12E-01 | 4,18E-01 | 23 | 24 | 6 | 5 | 73 | 63 | 38 | 41 | 2,2 | 0,8 | 1,4 |
| chr2:71662191-71662691:downstream_22338 | chr2 | 71558884 | 71662191 | + | ZNF638 | NM | 61 | 1,3 | 1 | -2 | 1,12E-01 | 4,18E-01 | 25 | 13 | 29 | 16 | 76 | 31 | 145 | 156 | -0,2 | -1,5 | 1,2 |
| chr17:26674035-26674535:downstream_12922 | chr17 | 26662547 | 26674035 | + | TNFAIP1 | NM | 70 | 1,2 | 1 | -2 | 1,13E-01 | 4,18E-01 | 12 | 26 | 8 | 15 | 92 | 111 | 170 | 123 | 0,7 | -0,5 | 1,3 |
| chr6:26225844-26226344:downstream_4920 | chr6 | 26225382 | 26225844 | + | HIST1H3E | NM | 27 | 1,5 | 1 | -2 | 1,13E-01 | NA | 33 | 27 | 8 | 7 | 54 | 28 | 24 | 35 | 1,9 | 0,5 | 1,5 |
| chr15:68498829-68499329:downstream_11780 | chr15 | 68499329 | 68522080 | - | CLN6 | NM | 21 | -1,7 | 1 | 2 | 1,13E-01 | NA | 3 | 4 | 11 | 20 | 28 | 26 | 33 | 42 | -2,1 | -0,5 | -1,6 |
| chr10:72642764-72643264:downstream_17699 | chr10 | 72643264 | 72648543 | - | PCBD1 | NM | 54 | -1,3 | 1 | 2 | 1,13E-01 | 4,18E-01 | 9 | 5 | 18 | 14 | 97 | 104 | 63 | 121 | -1,2 | 0,1 | -1,4 |
| chr1:26608013-26608513:downstream_95 | chr1 | 26606212 | 26608013 | + | SH3BGRL3 | NM | 8 | 3,3 | 2 | -2 | 1,13E-01 | NA | 2 | 8 | 0 | 0 | 19 | 20 | 7 | 9 | Inf | 1,3 | Inf |
| chr21:26954586-26955086:downstream_15234 | chr21 | 26955086 | 26955536 | - | LINC00515 | NR | 10 | 3,3 | 2 | -2 | 1,13E-01 | NA | 7 | 19 | 0 | 0 | 14 | 32 | 1 | 5 | Inf | 2,9 | Inf |
| chr8:67541987-67542487:downstream_6585 | chr8 | 67542487 | 67579452 | - | VCPIP1 | NM | 17 | 1,7 | 1 | -2 | 1,14E-01 | NA | 19 | 13 | 17 | 6 | 12 | 14 | 29 | 28 | 0,5 | -1,1 | 1,6 |
| chr1:40025984-40026484:downstream_17054 | chr1 | 40026484 | 40042521 | - | PABPC4 | NM | 14 | 2,2 | 1 | -2 | 1,14E-01 | NA | 4 | 17 | 4 | 3 | 6 | 27 | 22 | 29 | 1,5 | -0,6 | 2,2 |
| chr20:43123244-43123744:downstream_22593 | chr20 | 43104525 | 43123244 | + | TTPAL | NM | 8 | 2,4 | 2 | -2 | 1,14E-01 | NA | 9 | 8 | 3 | 2 | 14 | 3 | 11 | 17 | 1,7 | -0,8 | 2,4 |
| chrX:38665783-38666283:downstream_26915 | chrX | 38660684 | 38665783 | + | MID1IP1 | NM | 32 | -1,6 | 1 | 2 | 1,14E-01 | 4,22E-01 | 4 | 3 | 17 | 4 | 56 | 65 | 50 | 58 | -1,6 | 0,2 | -1,7 |
| chr3:10047601-10048101:downstream_2818 | chr3 | 10048101 | 10052779 | - | LOC401052 | NM | 5 | 2,9 | 2 | -2 | 1,14E-01 | NA | 8 | 8 | 3 | 4 | 2 | 1 | 7 | 5 | 1,2 | -1,8 | 3,0 |
| chr2:203744822-203745322:downstream_2380 | chr2 | 203745322 | 203776949 | - | WDR12 | NM | 24 | 1,6 | 1 | -2 | 1,14E-01 | NA | 9 | 11 | 7 | 7 | 20 | 34 | 64 | 44 | 0,5 | -1,0 | 1,6 |
| chr8:82605390-82605890:downstream_6622 | chr8 | 82605890 | 82607207 | - | SLC10A5 | NM | 19 | -2,3 | 1 | 2 | 1,14E-01 | NA | 27 | 56 | 21 | 25 | 5 | 15 | 3 | 0 | 0,9 | 2,9 | -2,0 |
| chr8:37637286-37637786:downstream_6528 | chr8 | 37620100 | 37637286 | + | PROSC | NM | 14 | 2,1 | 1 | -2 | 1,14E-01 | NA | 21 | 8 | 3 | 2 | 23 | 28 | 15 | 16 | 2,7 | 0,7 | 2,0 |
| chr20:3911568-3912068:downstream_22537 | chr20 | 3912068 | 3996216 | - | RNF24 | NM | 5 | 3,6 | 2 | -2 | 1,14E-01 | NA | 1 | 7 | 6 | 8 | 1 | 0 | 10 | 10 | -0,9 | -4,6 | 3,7 |
| chr9:74476867-74477367:downstream_26203 | chr9 | 74477367 | 74526148 | - | ABHD17B | NM | 3 | 3,9 | 2 | -2 | 1,14E-01 | NA | 6 | 3 | 3 | 1 | 2 | 0 | 4 | 6 | 1,3 | -2,6 | 3,9 |
| chr3:9986725-9987225:downstream_2817 | chr3 | 9987225 | 9994078 | - | PRRT3 | NM | 7 | 3,2 | 2 | -2 | 1,15E-01 | NA | 6 | 12 | 14 | 11 | 1 | 0 | 7 | 5 | -0,5 | -3,9 | 3,4 |
| chr16:8888536-8889036:downstream_12178 | chr16 | 8889036 | 8891505 | - | TMEM186 | NM | 11 | 2,9 | 2 | -2 | 1,15E-01 | NA | 7 | 8 | 0 | 1 | 33 | 14 | 14 | 11 | 4,1 | 0,9 | 3,3 |
| chr2:30383399-30383899:downstream_22327 | chr2 | 30369749 | 30383399 | + | YPEL5 | NM | 17 | 2,0 | 1 | -2 | 1,15E-01 | NA | 10 | 8 | 6 | 1 | 35 | 13 | 36 | 26 | 1,5 | -0,4 | 1,8 |
| chr17:62119889-62120389:downstream_12672 | chr17 | 62120389 | 62207502 | - | ERN1 | NM | 5 | 3,6 | 2 | -2 | 1,15E-01 | NA | 4 | 3 | 13 | 8 | 0 | 0 | 6 | 6 | -1,6 | -Inf | Inf |
| chr6:56920037-56920537:downstream_24828 | chr6 | 56911329 | 56920037 | + | KIAA1586 | NM | 20 | 1,6 | 1 | -2 | 1,16E-01 | NA | 43 | 24 | 6 | 10 | 25 | 21 | 19 | 14 | 2,1 | 0,5 | 1,7 |
| chr6:153331531-153332031:downstream_4820 | chr6 | 153332031 | 153452389 | - | RGS17 | NM | 18 | -2,8 | 2 | 2 | 1,16E-01 | NA | 1 | 0 | 4 | 21 | 11 | 11 | 51 | 43 | -5,0 | -2,1 | -2,9 |
| chr1:220321383-220321883:downstream_1371 | chr1 | 220267454 | 220321383 | + | IARS2 | NM | 312 | -1,0 | 1 | 2 | 1,16E-01 | 4,28E-01 | 5 | 14 | 31 | 32 | 457 | 409 | 863 | 690 | -1,8 | -0,8 | -0,9 |
| chr3:179111008-179111508:downstream_3286 | chr3 | 179065479 | 179111008 | + | MFN1 | NM | 15 | 2,1 | 1 | -2 | 1,16E-01 | NA | 4 | 8 | 15 | 24 | 3 | 1 | 19 | 42 | -1,7 | -3,9 | 2,1 |
| chr14:104378124-104378624:downstream_19170 | chr14 | 104378624 | 104387903 | - | C14orf2 | NM | 143 | 1,4 | 1 | -2 | 1,16E-01 | 4,28E-01 | 12 | 9 | 1 | 4 | 259 | 384 | 214 | 260 | 1,9 | 0,4 | 1,5 |
| chr6:24786325-24786825:downstream_24781 | chr6 | 24775158 | 24786325 | + | GMNN | NM | 16 | 1,8 | 1 | -2 | 1,17E-01 | NA | 13 | 10 | 4 | 4 | 24 | 18 | 19 | 33 | 1,5 | -0,3 | 1,8 |
| chr1:94020218-94020718:downstream_16753 | chr1 | 93913687 | 94020218 | + | FNBP1L | NM | 12 | 2,2 | 1 | -2 | 1,17E-01 | NA | 2 | 7 | 7 | 13 | 3 | 2 | 33 | 27 | -1,3 | -3,5 | 2,3 |
| chr1:23685440-23685940:downstream_17431 | chr1 | 23685940 | 23696357 | - | ZNF436 | NM | 7 | 3,1 | 2 | -2 | 1,17E-01 | NA | 10 | 13 | 1 | 1 | 15 | 1 | 3 | 10 | 3,3 | 0,3 | 3,0 |
| chr17:79674556-79675056:downstream_13429 | chr17 | 79670399 | 79674556 | + | MRPL12 | NM | 5 | 3,1 | 2 | -2 | 1,17E-01 | NA | 8 | 8 | 0 | 2 | 11 | 2 | 7 | 7 | 3,3 | -0,1 | 3,4 |
| chr13:50103117-50103617:downstream_19045 | chr13 | 50069800 | 50103117 | + | PHF11 | NM | 17 | 2,1 | 1 | -2 | 1,17E-01 | NA | 4 | 8 | 0 | 2 | 28 | 31 | 39 | 26 | 2,3 | -0,2 | 2,4 |
| chrX:151995370-151995870:downstream_8024 | chrX | 151995870 | 151999301 | - | CETN2 | NM | 103 | -1,1 | 1 | 2 | 1,17E-01 | 4,30E-01 | 9 | 8 | 15 | 11 | 242 | 217 | 150 | 174 | -0,6 | 0,5 | -1,1 |
| chr11:35253949-35254449:downstream_17938 | chr11 | 35160416 | 35253949 | + | CD44 | NM | 279 | 1,2 | 1 | -2 | 1,17E-01 | 4,30E-01 | 2 | 14 | 4 | 7 | 390 | 354 | 813 | 652 | 0,4 | -1,0 | 1,4 |
| chr1:206764473-206764973:downstream_16685 | chr1 | 206764973 | 206785904 | - | EIF2D | NM | 5 | 3,4 | 2 | -2 | 1,17E-01 | NA | 6 | 4 | 0 | 1 | 11 | 4 | 4 | 13 | 3,5 | -0,2 | 3,7 |
| chr10:97820625-97821125:downstream_17512 | chr10 | 97803158 | 97820625 | + | CCNJ | NM | 7 | 2,5 | 2 | -2 | 1,18E-01 | NA | 2 | 10 | 4 | 3 | 4 | 4 | 14 | 14 | 0,7 | -1,8 | 2,5 |
| chr10:112052297-112052797:downstream_8718 | chr10 | 112052797 | 112064707 | - | SMNDC1 | NM | 40 | 1,2 | 1 | -2 | 1,18E-01 | 4,31E-01 | 60 | 75 | 36 | 20 | 23 | 43 | 29 | 33 | 1,3 | 0,1 | 1,2 |
| chr2:67637533-67638033:downstream_1638 | chr2 | 67624441 | 67637533 | + | ETAA1 | NM | 19 | 1,5 | 1 | -2 | 1,18E-01 | NA | 24 | 30 | 8 | 14 | 15 | 19 | 18 | 24 | 1,3 | -0,3 | 1,6 |
| chr2:101888897-101889397:downstream_2148 | chr2 | 101889397 | 101889511 | - | SNORD89 | NR | 10 | -2,3 | 1 | 2 | 1,18E-01 | NA | 3 | 2 | 4 | 15 | 7 | 25 | 11 | 13 | -1,9 | 0,4 | -2,3 |
| chr1:20808383-20808883:downstream_523 | chr1 | 20808883 | 20812728 | - | CAMK2N1 | NM | 25 | -2,7 | 2 | 2 | 1,19E-01 | NA | 1 | 0 | 3 | 7 | 28 | 38 | 51 | 73 | -3,6 | -0,9 | -2,7 |
| chr2:160796759-160797259:downstream_22148 | chr2 | 160797259 | 160919126 | - | PLA2R1 | NM | 9 | -3,1 | 2 | 2 | 1,19E-01 | NA | 1 | 0 | 10 | 11 | 3 | 9 | 19 | 15 | -4,7 | -1,5 | -3,2 |
| chr5:177575964-177576464:downstream_24247 | chr5 | 177576464 | 177580961 | - | NHP2 | NM | 19 | 1,7 | 1 | -2 | 1,19E-01 | NA | 25 | 33 | 6 | 4 | 21 | 32 | 15 | 12 | 2,6 | 0,9 | 1,6 |
| chr2:190610885-190611385:downstream_2353 | chr2 | 190611385 | 190627924 | - | OSGEPL1 | NM | 8 | 2,3 | 1 | -2 | 1,19E-01 | NA | 6 | 5 | 3 | 4 | 5 | 7 | 17 | 21 | 0,6 | -1,7 | 2,3 |
| chr5:137841281-137841781:downstream_24110 | chr5 | 137841781 | 137878989 | - | ETF1 | NM | 32 | 1,5 | 1 | -2 | 1,19E-01 | 4,32E-01 | 15 | 23 | 22 | 4 | 30 | 37 | 60 | 63 | 0,5 | -0,9 | 1,4 |
| chr4:147111213-147111713:downstream_3862 | chr4 | 147096834 | 147111213 | + | LSM6 | NM | 16 | 1,7 | 1 | -2 | 1,19E-01 | NA | 33 | 23 | 8 | 11 | 14 | 11 | 10 | 18 | 1,5 | -0,2 | 1,7 |
| chr5:108063025-108063525:downstream_4358 | chr5 | 108063525 | 108063962 | - | LINC01023 | NR | 5 | 3,4 | 2 | -2 | 1,19E-01 | NA | 9 | 8 | 0 | 1 | 10 | 4 | 7 | 2 | 4,3 | 0,5 | 3,8 |
| chr8:33358778-33359278:downstream_6519 | chr8 | 33342684 | 33358778 | + | MAK16 | NM | 47 | 1,6 | 1 | -2 | 1,19E-01 | 4,32E-01 | 15 | 11 | 3 | 2 | 115 | 118 | 56 | 55 | 2,6 | 1,1 | 1,5 |
| chr19:14625075-14625575:downstream_21102 | chr19 | 14625575 | 14629371 | - | DNAJB1 | NM | 68 | 1,4 | 1 | -2 | 1,19E-01 | 4,32E-01 | 8 | 5 | 6 | 3 | 95 | 104 | 160 | 165 | 0,6 | -0,7 | 1,3 |
| chr7:21941186-21941686:downstream_5522 | chr7 | 21582832 | 21941186 | + | DNAH11 | NM | 5 | -3,7 | 2 | 2 | 1,19E-01 | NA | 0 | 0 | 4 | 11 | 2 | 2 | 7 | 10 | -Inf | -1,9 | -Inf |
| chr8:61536203-61536703:downstream_26077 | chr8 | 61429468 | 61536203 | + | RAB2A | NM | 5 | -3,5 | 2 | 2 | 1,19E-01 | NA | 1 | 0 | 13 | 6 | 3 | 4 | 10 | 3 | -4,5 | -0,9 | -3,6 |
| chr5:41921738-41922238:downstream_4234 | chr5 | 41904469 | 41921738 | + | C5orf51 | NM | 14 | 1,9 | 1 | -2 | 1,20E-01 | NA | 19 | 10 | 4 | 2 | 20 | 23 | 18 | 17 | 2,1 | 0,3 | 1,8 |
| chr5:72801448-72801948:downstream_24028 | chr5 | 72794249 | 72801448 | + | BTF3 | NM | 67 | 1,1 | 1 | -2 | 1,20E-01 | 4,33E-01 | 29 | 44 | 13 | 10 | 156 | 109 | 68 | 108 | 1,7 | 0,6 | 1,1 |
| chr12:32146043-32146543:downstream_9842 | chr12 | 32112352 | 32146043 | + | KIAA1551 | NM | 28 | 1,7 | 1 | -2 | 1,20E-01 | NA | 3 | 15 | 1 | 8 | 31 | 36 | 67 | 62 | 0,9 | -0,9 | 1,8 |
| chr11:57468659-57469159:downstream_9322 | chr11 | 57435473 | 57468659 | + | ZDHHC5 | NM | 83 | 1,1 | 1 | -2 | 1,20E-01 | 4,33E-01 | 14 | 17 | 7 | 12 | 131 | 128 | 186 | 168 | 0,7 | -0,5 | 1,2 |
| chr7:38270272-38270772:downstream_5796 | chr7 | 38217807 | 38270272 | + | STARD3NL | NM | 61 | 1,2 | 1 | -2 | 1,21E-01 | 4,33E-01 | 10 | 9 | 13 | 11 | 57 | 58 | 182 | 149 | -0,4 | -1,5 | 1,1 |
| chr20:17921739-17922239:downstream_22569 | chr20 | 17922239 | 17949634 | - | SNX5 | NM | 55 | 1,3 | 1 | -2 | 1,21E-01 | 4,33E-01 | 11 | 10 | 6 | 5 | 83 | 103 | 116 | 107 | 1,0 | -0,3 | 1,3 |
| chr14:50884710-50885210:downstream_19296 | chr14 | 50885210 | 50999376 | - | MAP4K5 | NM | 17 | 1,7 | 1 | -2 | 1,21E-01 | NA | 7 | 7 | 11 | 11 | 12 | 5 | 50 | 37 | -0,6 | -2,4 | 1,7 |
| chr12:121407140-121407640:downstream_10458 | chr12 | 121407640 | 121410095 | - | HNF1A-AS1 | NR | 38 | -1,3 | 1 | 2 | 1,21E-01 | 4,33E-01 | 19 | 6 | 88 | 67 | 15 | 20 | 43 | 46 | -2,7 | -1,3 | -1,3 |
| chr9:110094475-110094975:downstream_26470 | chr9 | 110045516 | 110094475 | + | RAD23B | NM | 11 | 2,0 | 1 | -2 | 1,21E-01 | NA | 9 | 12 | 3 | 6 | 11 | 9 | 19 | 16 | 1,3 | -0,8 | 2,1 |
| chr4:175157309-175157809:downstream_3908 | chr4 | 175157809 | 175205402 | - | FBXO8 | NM | 9 | 2,4 | 2 | -2 | 1,21E-01 | NA | 9 | 12 | 14 | 9 | 4 | 0 | 15 | 9 | -0,2 | -2,6 | 2,4 |
| chr2:62094761-62095261:downstream_21803 | chr2 | 62095261 | 62115806 | - | CCT4 | NM | 60 | 1,2 | 1 | -2 | 1,21E-01 | 4,33E-01 | 20 | 39 | 4 | 11 | 141 | 108 | 64 | 94 | 2,0 | 0,7 | 1,3 |
| chr6:7586946-7587446:downstream_24732 | chr6 | 7541869 | 7586946 | + | DSP | NM | 17 | 2,3 | 1 | -2 | 1,22E-01 | NA | 8 | 0 | 1 | 2 | 19 | 27 | 32 | 48 | 1,4 | -0,8 | 2,2 |
| chr19:29697666-29698166:downstream_14181 | chr19 | 29698166 | 29704136 | - | UQCRFS1 | NM | 73 | 1,0 | 1 | -2 | 1,22E-01 | 4,36E-01 | 58 | 84 | 28 | 32 | 115 | 93 | 79 | 94 | 1,2 | 0,3 | 1,0 |
| chr17:46672598-46673098:downstream_13207 | chr17 | 46673098 | 46682334 | - | HOXB6 | NM | 22 | -1,8 | 1 | 2 | 1,22E-01 | NA | 2 | 3 | 6 | 10 | 24 | 60 | 43 | 28 | -1,5 | 0,2 | -1,8 |
| chr10:59950777-59951277:downstream_8529 | chr10 | 59951277 | 60027694 | - | IPMK | NM | 8 | 2,9 | 2 | -2 | 1,22E-01 | NA | 0 | 8 | 4 | 7 | 3 | 0 | 26 | 14 | -0,6 | -3,6 | 3,1 |
| chr9:123561009-123561509:downstream_7285 | chr9 | 123555774 | 123561009 | + | LOC100288842 | NR | 7 | -2,5 | 2 | 2 | 1,23E-01 | NA | 3 | 4 | 19 | 9 | 3 | 11 | 8 | 2 | -2,0 | 0,5 | -2,5 |
| chr2:9723595-9724095:downstream_1850 | chr2 | 9724095 | 9771184 | - | YWHAQ | NM | 65 | 1,3 | 1 | -2 | 1,23E-01 | 4,38E-01 | 12 | 17 | 13 | 2 | 115 | 115 | 114 | 135 | 1,0 | -0,1 | 1,1 |
| chr22:45999811-46000311:downstream_15676 | chr22 | 46000311 | 46001527 | - | LINC01589 | NR | 7 | -2,6 | 2 | 2 | 1,23E-01 | NA | 2 | 3 | 10 | 3 | 8 | 17 | 8 | 4 | -1,5 | 1,0 | -2,6 |
| chr6:79946517-79947017:downstream_5239 | chr6 | 79943390 | 79946517 | + | HMGN3-AS1 | NR | 10 | 2,5 | 2 | -2 | 1,24E-01 | NA | 34 | 15 | 4 | 11 | 7 | 0 | 1 | 10 | 1,7 | -0,8 | 2,5 |
| chr9:14010-14510:downstream_6957 | chr9 | 14510 | 29739 | - | WASH1 | NM | 10 | 2,4 | 2 | -2 | 1,24E-01 | NA | 5 | 6 | 1 | 2 | 22 | 5 | 19 | 22 | 1,8 | -0,6 | 2,5 |
| chr1:25548266-25548766:downstream_17297 | chr1 | 25548766 | 25559013 | - | SYF2 | NM | 12 | 2,7 | 2 | -2 | 1,25E-01 | NA | 11 | 9 | 1 | 0 | 20 | 38 | 10 | 11 | 3,9 | 1,5 | 2,3 |
| chr1:238643183-238643683:downstream_1443 | chr1 | 238643683 | 238649317 | - | LINC01139 | NR | 5 | 3,6 | 2 | -2 | 1,25E-01 | NA | 12 | 10 | 0 | 1 | 9 | 1 | 1 | 4 | 4,7 | 0,9 | 3,9 |
| chr2:178414380-178414880:downstream_2337 | chr2 | 178414880 | 178417524 | - | TTC30B | NM | 9 | 2,3 | 1 | -2 | 1,26E-01 | NA | 6 | 21 | 4 | 13 | 3 | 4 | 11 | 11 | 0,6 | -1,7 | 2,3 |
| chr7:95033673-95034173:downstream_25683 | chr7 | 95034173 | 95064384 | - | PON2 | NM | 1082 | -0,7 | 0 | 2 | 1,26E-01 | 4,44E-01 | 47 | 38 | 56 | 64 | 2083 | 2459 | 2057 | 1852 | -0,5 | 0,2 | -0,7 |
| chr2:230628052-230628552:downstream_22298 | chr2 | 230628552 | 230786725 | - | TRIP12 | NM | 15 | -2,8 | 2 | 2 | 1,26E-01 | NA | 1 | 0 | 14 | 10 | 5 | 15 | 33 | 41 | -4,9 | -1,9 | -2,9 |
| chr14:50583345-50583845:downstream_10940 | chr14 | 50583845 | 50698099 | - | SOS2 | NM | 9 | -3,1 | 2 | 2 | 1,26E-01 | NA | 0 | 0 | 3 | 11 | 5 | 4 | 25 | 21 | -Inf | -2,4 | -Inf |
| chr1:85622855-85623355:downstream_806 | chr1 | 85623355 | 85666728 | - | SYDE2 | NM | 6 | 2,9 | 2 | -2 | 1,26E-01 | NA | 2 | 12 | 6 | 6 | 3 | 0 | 13 | 9 | 0,2 | -2,7 | 3,0 |
| chr20:49550904-49551404:downstream_15084 | chr20 | 49551404 | 49575060 | - | DPM1 | NM | 31 | 1,4 | 1 | -2 | 1,26E-01 | 4,44E-01 | 10 | 17 | 4 | 7 | 45 | 52 | 56 | 58 | 1,2 | -0,2 | 1,5 |
| chr11:10532724-10533224:downstream_9144 | chr11 | 10533224 | 10562774 | - | RNF141 | NM | 4 | 3,5 | 2 | -2 | 1,26E-01 | NA | 5 | 7 | 1 | 0 | 2 | 7 | 4 | 6 | 3,1 | -0,1 | 3,2 |
| chr5:137772716-137773216:downstream_4093 | chr5 | 137688284 | 137772716 | + | KDM3B | NM | 32 | -2,6 | 2 | 2 | 1,26E-01 | 4,44E-01 | 0 | 0 | 8 | 4 | 12 | 23 | 102 | 105 | -Inf | -2,5 | -Inf |
| chr2:99234977-99235477:downstream_2133 | chr2 | 99225041 | 99234977 | + | UNC50 | NM | 46 | 1,3 | 1 | -2 | 1,26E-01 | 4,44E-01 | 7 | 11 | 7 | 6 | 72 | 46 | 109 | 112 | 0,4 | -0,9 | 1,3 |
| chr6:13621127-13621627:downstream_4874 | chr6 | 13615558 | 13621127 | + | NOL7 | NM | 44 | 1,2 | 1 | -2 | 1,26E-01 | 4,44E-01 | 52 | 26 | 33 | 21 | 41 | 44 | 89 | 47 | 0,5 | -0,7 | 1,2 |
| chr17:17920833-17921333:downstream_12872 | chr17 | 17921333 | 17942480 | - | ATPAF2 | NM | 6 | 3,7 | 2 | -2 | 1,26E-01 | NA | 0 | 11 | 0 | 0 | 6 | 20 | 7 | 6 | Inf | 1,0 | Inf |
| chr17:35873588-35874088:downstream_13007 | chr17 | 35849950 | 35873588 | + | DUSP14 | NM | 126 | 2,4 | 2 | -2 | 1,27E-01 | 4,44E-01 | 15 | 8 | 0 | 0 | 425 | 475 | 36 | 46 | Inf | 3,5 | Inf |
| chr12:112331228-112331728:downstream_18671 | chr12 | 112280031 | 112331228 | + | MAPKAPK5 | NM | 16 | 1,7 | 1 | -2 | 1,27E-01 | NA | 8 | 6 | 4 | 7 | 18 | 11 | 35 | 40 | 0,4 | -1,4 | 1,8 |
| chr12:83528067-83528567:downstream_9893 | chr12 | 83080933 | 83528067 | + | TMTC2 | NM | 18 | 2,7 | 2 | -2 | 1,27E-01 | NA | 2 | 7 | 32 | 51 | 1 | 0 | 28 | 25 | -3,2 | -6,0 | 2,9 |
| chr5:132531651-132532151:downstream_4086 | chr5 | 132532151 | 132948223 | - | FSTL4 | NM | 4 | -4,2 | 3 | 2 | 1,27E-01 | NA | 0 | 0 | 1 | 8 | 0 | 8 | 7 | 6 | -Inf | -0,7 | -Inf |
| chr17:3572963-3573463:downstream_20464 | chr17 | 3572016 | 3572963 | + | EMC6 | NM | 192 | 1,1 | 1 | -2 | 1,28E-01 | 4,44E-01 | 21 | 24 | 11 | 1 | 516 | 484 | 241 | 239 | 1,9 | 1,1 | 0,9 |
| chr3:149682190-149682690:downstream_23336 | chr3 | 149682690 | 149688741 | - | PFN2 | NM | 36 | 1,4 | 1 | -2 | 1,28E-01 | 4,44E-01 | 11 | 10 | 7 | 6 | 66 | 29 | 68 | 89 | 0,7 | -0,7 | 1,4 |
| chr17:42754304-42754804:downstream_20383 | chr17 | 42754804 | 42767165 | - | CCDC43 | NM | 33 | 1,3 | 1 | -2 | 1,28E-01 | 4,44E-01 | 24 | 25 | 7 | 8 | 51 | 64 | 35 | 53 | 1,7 | 0,4 | 1,3 |
| chr8:145538385-145538885:downstream_6784 | chr8 | 145515269 | 145538385 | + | HSF1 | NM | 31 | 1,6 | 1 | -2 | 1,28E-01 | 4,44E-01 | 10 | 6 | 1 | 3 | 61 | 55 | 49 | 66 | 1,7 | 0,0 | 1,7 |
| chr7:75955607-75956107:downstream_5915 | chr7 | 75956107 | 75988342 | - | YWHAG | NM | 127 | 1,1 | 1 | -2 | 1,29E-01 | 4,46E-01 | 19 | 12 | 19 | 5 | 185 | 193 | 316 | 272 | 0,3 | -0,6 | 1,0 |
| chr9:127704387-127704887:downstream_26490 | chr9 | 127704887 | 127905838 | - | SCAI | NM | 4 | 3,7 | 2 | -2 | 1,29E-01 | NA | 2 | 8 | 0 | 2 | 0 | 6 | 10 | 4 | 2,6 | -1,3 | 3,9 |
| chr1:165744685-165745185:downstream_1194 | chr1 | 165738165 | 165744685 | + | LOC100147773 | NR | 18 | -1,6 | 1 | 2 | 1,29E-01 | NA | 17 | 24 | 17 | 14 | 32 | 25 | 11 | 3 | 0,4 | 2,0 | -1,6 |
| chr11:33095109-33095609:downstream_18335 | chr11 | 33060962 | 33095109 | + | TCP11L1 | NM | 78 | -1,0 | 1 | 2 | 1,29E-01 | 4,46E-01 | 27 | 19 | 83 | 54 | 89 | 85 | 143 | 120 | -1,6 | -0,6 | -1,0 |
| chr3:184086383-184086883:downstream_23353 | chr3 | 184079501 | 184086383 | + | POLR2H | NM | 37 | 1,4 | 1 | -2 | 1,29E-01 | 4,46E-01 | 15 | 20 | 4 | 7 | 79 | 55 | 45 | 73 | 1,6 | 0,2 | 1,4 |
| chr9:108314714-108315214:downstream_26322 | chr9 | 108210314 | 108314714 | + | FSD1L | NM | 8 | -2,7 | 2 | 2 | 1,29E-01 | NA | 2 | 0 | 17 | 16 | 6 | 2 | 11 | 12 | -4,3 | -1,6 | -2,7 |
| chr17:7128160-7128660:downstream_12796 | chr17 | 7128660 | 7137863 | - | DVL2 | NM | 60 | 1,2 | 1 | -2 | 1,29E-01 | 4,46E-01 | 72 | 196 | 86 | 68 | 11 | 13 | 15 | 17 | 0,8 | -0,4 | 1,2 |
| chr9:102736818-102737318:downstream_7228 | chr9 | 102668914 | 102736818 | + | STX17 | NM | 10 | 2,3 | 2 | -2 | 1,30E-01 | NA | 15 | 17 | 10 | 17 | 3 | 0 | 6 | 8 | 0,3 | -2,1 | 2,4 |
| chr20:4682234-4682734:downstream_22515 | chr20 | 4666796 | 4682234 | + | PRNP | NM | 433 | 1,8 | 1 | -2 | 1,30E-01 | 4,47E-01 | 3 | 10 | 1 | 1 | 1247 | 929 | 710 | 560 | 2,5 | 0,8 | 1,8 |
| chr5:39104853-39105353:downstream_24133 | chr5 | 39105353 | 39270759 | - | FYB | NM | 27 | 1,6 | 1 | -2 | 1,30E-01 | NA | 9 | 3 | 3 | 4 | 38 | 33 | 65 | 62 | 0,8 | -0,8 | 1,6 |
| chr3:52027143-52027643:downstream_2986 | chr3 | 52027643 | 52029958 | - | RPL29 | NM | 101 | -1,1 | 1 | 2 | 1,30E-01 | 4,47E-01 | 11 | 11 | 15 | 7 | 280 | 255 | 93 | 138 | 0,0 | 1,2 | -1,2 |
| chr3:30735633-30736133:downstream_23465 | chr3 | 30647993 | 30735633 | + | TGFBR2 | NM | 115 | 1,0 | 1 | -2 | 1,31E-01 | 4,47E-01 | 13 | 7 | 67 | 59 | 35 | 22 | 369 | 349 | -2,7 | -3,6 | 1,0 |
| chr16:2821413-2821913:downstream_12125 | chr16 | 2802329 | 2821413 | + | SRRM2 | NM | 44 | 3,9 | 3 | -2 | 1,31E-01 | 4,47E-01 | 0 | 13 | 0 | 1 | 86 | 65 | 106 | 82 | 3,9 | -0,3 | 4,3 |
| chr2:150425646-150426146:downstream_2271 | chr2 | 150426146 | 150444330 | - | MMADHC | NM | 136 | 0,9 | 1 | -2 | 1,32E-01 | 4,47E-01 | 50 | 29 | 24 | 23 | 200 | 259 | 234 | 267 | 0,8 | -0,1 | 0,9 |
| chr8:145152430-145152930:downstream_6777 | chr8 | 145149937 | 145152430 | + | CYC1 | NM | 90 | 1,4 | 1 | -2 | 1,32E-01 | 4,47E-01 | 15 | 6 | 0 | 4 | 202 | 219 | 114 | 157 | 2,3 | 0,6 | 1,7 |
| chr9:130696873-130697373:downstream_7331 | chr9 | 130697373 | 130700763 | - | DPM2 | NM | 18 | 2,6 | 2 | -2 | 1,32E-01 | NA | 7 | 4 | 1 | 0 | 35 | 48 | 17 | 34 | 2,9 | 0,7 | 2,2 |
| chr4:41270446-41270946:downstream_3658 | chr4 | 41258897 | 41270446 | + | UCHL1 | NM | 322 | -0,9 | 1 | 2 | 1,32E-01 | 4,47E-01 | 21 | 10 | 18 | 21 | 764 | 744 | 438 | 564 | -0,3 | 0,6 | -0,9 |
| chr1:247463121-247463621:downstream_58 | chr1 | 247463621 | 247495045 | - | ZNF496 | NM | 12 | 2,1 | 1 | -2 | 1,32E-01 | NA | 4 | 10 | 17 | 16 | 4 | 0 | 26 | 19 | -1,3 | -3,5 | 2,2 |
| chr2:46613842-46614342:downstream_1617 | chr2 | 46524540 | 46613842 | + | EPAS1 | NM | 36 | -1,8 | 1 | 2 | 1,32E-01 | 4,47E-01 | 2 | 1 | 10 | 9 | 50 | 38 | 67 | 115 | -2,9 | -1,0 | -1,8 |
| chr11:76509198-76509698:downstream_18373 | chr11 | 76493356 | 76509198 | + | TSKU | NM | 145 | 0,8 | 1 | -2 | 1,32E-01 | 4,47E-01 | 158 | 161 | 116 | 110 | 122 | 160 | 154 | 184 | 0,5 | -0,3 | 0,8 |
| chr1:174981163-174981663:downstream_16546 | chr1 | 174968570 | 174981163 | + | CACYBP | NM | 14 | 1,7 | 1 | -2 | 1,32E-01 | NA | 20 | 22 | 11 | 14 | 9 | 7 | 15 | 15 | 0,7 | -0,9 | 1,7 |
| chr9:88878962-88879462:downstream_7161 | chr9 | 88879462 | 88897490 | - | ISCA1 | NM | 14 | 1,8 | 1 | -2 | 1,33E-01 | NA | 18 | 11 | 3 | 8 | 14 | 15 | 29 | 12 | 1,4 | -0,6 | 1,9 |
| chr17:80035713-80036213:downstream_13446 | chr17 | 80036213 | 80056106 | - | FASN | NM | 13 | 2,3 | 2 | -2 | 1,33E-01 | NA | 3 | 7 | 1 | 1 | 18 | 27 | 21 | 26 | 2,2 | -0,1 | 2,2 |
| chr11:102920912-102921412:downstream_9616 | chr11 | 102921412 | 102962944 | - | DCUN1D5 | NM | 4 | 3,8 | 3 | -2 | 1,33E-01 | NA | 1 | 10 | 0 | 1 | 7 | 1 | 4 | 7 | 3,7 | -0,5 | 4,2 |
| chr17:61909387-61909887:downstream_20654 | chr17 | 61904769 | 61909387 | + | PSMC5 | NM | 21 | 1,6 | 1 | -2 | 1,33E-01 | NA | 22 | 8 | 3 | 7 | 30 | 33 | 29 | 39 | 1,5 | -0,1 | 1,7 |
| chr7:150744869-150745369:downstream_25370 | chr7 | 150725508 | 150744869 | + | ABCB8 | NM | 6 | 3,0 | 2 | -2 | 1,33E-01 | NA | 2 | 9 | 0 | 7 | 5 | 0 | 15 | 11 | 0,8 | -2,4 | 3,2 |
| chr11:65273983-65274483:downstream_8899 | chr11 | 65265232 | 65273983 | + | MALAT1 | NR | 51 | -1,7 | 1 | 2 | 1,34E-01 | 4,50E-01 | 2 | 1 | 8 | 7 | 47 | 115 | 114 | 115 | -2,2 | -0,5 | -1,7 |
| chr16:85709812-85710312:downstream_20007 | chr16 | 85645028 | 85709812 | + | GSE1 | NM | 17 | -2,6 | 2 | 1 | 1,34E-01 | NA | 1 | 0 | 4 | 8 | 20 | 18 | 38 | 51 | -3,9 | -1,2 | -2,7 |
| chr11:73638781-73639281:downstream_18188 | chr11 | 73587743 | 73638781 | + | PAAF1 | NM | 9 | 2,5 | 2 | -1 | 1,34E-01 | NA | 3 | 7 | 4 | 0 | 9 | 10 | 11 | 25 | 1,3 | -1,0 | 2,2 |
| chr2:152972814-152973314:downstream_2277 | chr2 | 152973314 | 153032506 | - | STAM2 | NM | 13 | -2,7 | 2 | 1 | 1,34E-01 | NA | 1 | 0 | 10 | 6 | 9 | 16 | 33 | 32 | -4,3 | -1,4 | -2,9 |
| chr13:58303065-58303565:downstream_10774 | chr13 | 58205788 | 58303065 | + | PCDH17 | NM | 11 | -3,1 | 2 | 1 | 1,34E-01 | NA | 0 | 0 | 14 | 30 | 2 | 1 | 25 | 18 | -Inf | -4,1 | -Inf |
| chr10:102282985-102283485:downstream_17666 | chr10 | 102283485 | 102289680 | - | NDUFB8 | NM | 30 | -1,6 | 1 | 1 | 1,34E-01 | 4,51E-01 | 8 | 1 | 4 | 7 | 61 | 93 | 25 | 39 | -0,3 | 1,3 | -1,6 |
| chr16:31051488-31051988:downstream_20228 | chr16 | 31044415 | 31051488 | + | STX4 | NM | 12 | 1,9 | 1 | -1 | 1,35E-01 | NA | 16 | 15 | 15 | 11 | 8 | 2 | 15 | 16 | 0,2 | -1,6 | 1,9 |
| chr9:75486144-75486644:downstream_26367 | chr9 | 75486644 | 75489025 | - | LINC01474 | NR | 15 | -2,0 | 1 | 1 | 1,35E-01 | NA | 2 | 1 | 22 | 13 | 7 | 16 | 32 | 26 | -3,4 | -1,3 | -2,0 |
| chrX:119657945-119658445:downstream_26775 | chrX | 119658445 | 119709684 | - | CUL4B | NM | 12 | 2,0 | 1 | -1 | 1,35E-01 | NA | 7 | 3 | 7 | 3 | 10 | 5 | 26 | 34 | -0,1 | -2,0 | 1,9 |
| chr3:171317694-171318194:downstream_23346 | chr3 | 171318194 | 171528284 | - | PLD1 | NM | 12 | -3,0 | 2 | 1 | 1,35E-01 | NA | 0 | 0 | 11 | 6 | 2 | 9 | 42 | 25 | -Inf | -2,7 | -Inf |
| chr15:81047962-81048462:downstream_11512 | chr15 | 80987651 | 81047962 | + | ABHD17C | NM | 17 | 1,8 | 1 | -1 | 1,35E-01 | NA | 3 | 7 | 4 | 2 | 19 | 19 | 46 | 39 | 0,6 | -1,2 | 1,7 |
| chr13:77618291-77618791:downstream_10609 | chr13 | 77618791 | 77901177 | - | MYCBP2 | NM | 8 | -3,2 | 2 | 1 | 1,35E-01 | NA | 0 | 0 | 3 | 8 | 2 | 8 | 18 | 21 | -Inf | -2,0 | -Inf |
| chr12:88886069-88886569:downstream_18617 | chr12 | 88886569 | 88974250 | - | KITLG | NM | 23 | 1,5 | 1 | -1 | 1,36E-01 | NA | 8 | 7 | 4 | 9 | 22 | 21 | 51 | 61 | 0,2 | -1,4 | 1,6 |
| chr14:94516767-94517267:downstream_11281 | chr14 | 94517267 | 94547558 | - | DDX24 | NM | 144 | 1,6 | 1 | -1 | 1,36E-01 | 4,56E-01 | 2 | 10 | 1 | 2 | 299 | 331 | 213 | 297 | 2,0 | 0,3 | 1,7 |
| chr17:38496618-38497118:downstream_13046 | chr17 | 38497118 | 38499388 | - | RARA-AS1 | NR | 27 | 1,4 | 1 | -1 | 1,37E-01 | NA | 60 | 65 | 11 | 7 | 24 | 29 | 7 | 12 | 2,8 | 1,4 | 1,3 |
| chr17:41166476-41166976:downstream_13146 | chr17 | 41158741 | 41166476 | + | IFI35 | NM | 34 | 1,2 | 1 | -1 | 1,37E-01 | 4,58E-01 | 37 | 38 | 10 | 9 | 64 | 50 | 35 | 31 | 2,0 | 0,8 | 1,2 |
| chr16:30756517-30757017:downstream_12303 | chr16 | 30751511 | 30756517 | + | TMEM265 | NM | 3 | -3,5 | 2 | 1 | 1,37E-01 | NA | 3 | 0 | 6 | 5 | 9 | 2 | 1 | 2 | -1,7 | 1,8 | -3,5 |
| chr12:129277238-129277738:downstream_10513 | chr12 | 129277738 | 129308541 | - | SLC15A4 | NM | 18 | -2,2 | 1 | 1 | 1,37E-01 | NA | 0 | 2 | 3 | 7 | 34 | 25 | 38 | 34 | -2,3 | -0,3 | -2,0 |
| chr20:48809227-48809727:downstream_22381 | chr20 | 48807119 | 48809227 | + | CEBPB | NM | 25 | 1,7 | 1 | -1 | 1,37E-01 | NA | 3 | 8 | 0 | 6 | 33 | 27 | 71 | 50 | 0,9 | -1,0 | 2,0 |
| chr2:95812899-95813399:downstream_1670 | chr2 | 95813399 | 95825263 | - | ZNF514 | NM | 19 | 1,5 | 1 | -1 | 1,38E-01 | NA | 16 | 23 | 29 | 18 | 7 | 9 | 22 | 28 | -0,3 | -1,7 | 1,5 |
| chr2:8824583-8825083:downstream_1847 | chr2 | 8822112 | 8824583 | + | ID2 | NM | 55 | -1,4 | 1 | 1 | 1,38E-01 | 4,58E-01 | 13 | 3 | 7 | 4 | 124 | 206 | 47 | 34 | 0,5 | 2,0 | -1,5 |
| chr1:149855509-149856009:downstream_974 | chr1 | 149856009 | 149858232 | - | HIST2H2BE | NM | 21 | 1,7 | 1 | -1 | 1,38E-01 | NA | 20 | 23 | 0 | 4 | 39 | 48 | 11 | 21 | 3,4 | 1,4 | 2,0 |
| chr17:7215782-7216282:downstream_20461 | chr17 | 7210317 | 7215782 | + | EIF5A | NM | 123 | -0,9 | 1 | 1 | 1,38E-01 | 4,58E-01 | 21 | 25 | 33 | 17 | 331 | 238 | 157 | 159 | -0,1 | 0,8 | -1,0 |
| chr15:41591795-41592295:downstream_11675 | chr15 | 41576200 | 41591795 | + | OIP5-AS1 | NR | 49 | 1,1 | 1 | -1 | 1,38E-01 | 4,58E-01 | 55 | 60 | 100 | 86 | 9 | 12 | 31 | 39 | -0,7 | -1,8 | 1,1 |
| chr3:15450766-15451266:downstream_23270 | chr3 | 15451266 | 15469054 | - | METTL6 | NM | 9 | 2,1 | 1 | -1 | 1,38E-01 | NA | 7 | 14 | 4 | 2 | 11 | 9 | 11 | 15 | 1,7 | -0,4 | 2,0 |
| chr8:126034525-126035025:downstream_6710 | chr8 | 126010719 | 126034525 | + | SQLE | NM | 118 | 1,3 | 1 | -1 | 1,38E-01 | 4,58E-01 | 6 | 1 | 7 | 6 | 85 | 89 | 370 | 383 | -0,9 | -2,1 | 1,2 |
| chr11:85989852-85990352:downstream_17996 | chr11 | 85955425 | 85989852 | + | EED | NM | 30 | 1,5 | 1 | -1 | 1,39E-01 | 4,58E-01 | 38 | 53 | 14 | 10 | 16 | 58 | 36 | 17 | 1,9 | 0,5 | 1,5 |
| chrX:117928496-117928996:downstream_7892 | chrX | 117861558 | 117928496 | + | IL13RA1 | NM | 80 | -1,1 | 1 | 1 | 1,39E-01 | 4,58E-01 | 8 | 3 | 21 | 13 | 118 | 138 | 171 | 171 | -1,6 | -0,4 | -1,2 |
| chr10:5805703-5806203:downstream_8237 | chr10 | 5726800 | 5805703 | + | FAM208B | NM | 13 | 2,0 | 1 | -1 | 1,39E-01 | NA | 7 | 2 | 3 | 5 | 11 | 8 | 22 | 42 | 0,3 | -1,7 | 2,0 |
| chr5:179040678-179041178:downstream_24166 | chr5 | 179041178 | 179050722 | - | HNRNPH1 | NM | 163 | 0,9 | 1 | -1 | 1,39E-01 | 4,59E-01 | 41 | 24 | 24 | 21 | 318 | 214 | 299 | 367 | 0,6 | -0,3 | 0,9 |
| chr11:111597131-111597631:downstream_18224 | chr11 | 111597631 | 111637169 | - | PPP2R1B | NM | 6 | 2,5 | 2 | -1 | 1,39E-01 | NA | 5 | 7 | 0 | 3 | 7 | 6 | 13 | 11 | 1,8 | -0,9 | 2,7 |

| | | | | | | | | | | | | | | | | | | | | | | | |
|---|---|---|---|---|---|---|---|---|---|---|---|---|---|---|---|---|---|---|---|---|---|---|---|
| chr11:33724055-33724555:downstream_17939 | chr11 | 33724555 | 33758025 | - | CD59 | NM | 16 | 1,7 | 1 | -1 | 1,39E-01 | NA | 43 | 28 | 7 | 7 | 11 | 15 | 4 | 11 | 2,3 | 0,7 | 1,6 |
| chr17:41842988-41843488:downstream_13151 | chr17 | 41843488 | 41856368 | - | DUSP3 | NM | 50 | 1,1 | 1 | -1 | 1,40E-01 | 4,59E-01 | 13 | 22 | 10 | 10 | 85 | 71 | 100 | 92 | 0,8 | -0,3 | 1,1 |
| chr11:47185348-47185848:downstream_17873 | chr11 | 47185848 | 47198676 | - | ARFGAP2 | NM | 7 | 2,7 | 2 | -1 | 1,40E-01 | NA | 6 | 11 | 1 | 1 | 14 | 5 | 8 | 7 | 2,9 | 0,2 | 2,6 |
| chr19:13944829-13945329:downstream_14051 | chr19 | 13945329 | 13947103 | - | LOC284454 | NR | 20 | 1,8 | 1 | -1 | 1,40E-01 | NA | 5 | 15 | 1 | 3 | 36 | 37 | 39 | 24 | 2,1 | 0,2 | 1,9 |
| chr16:67878098-67878598:downstream_12424 | chr16 | 67876212 | 67878098 | + | THAP11 | NM | 12 | 2,4 | 2 | -1 | 1,40E-01 | NA | 5 | 5 | 0 | 2 | 24 | 17 | 13 | 31 | 2,6 | -0,1 | 2,6 |
| chr20:56287836-56288336:downstream_15105 | chr20 | 56285238 | 56287836 | + | NKILA | NR | 36 | 1,6 | 1 | -1 | 1,40E-01 | 4,60E-01 | 88 | 73 | 8 | 0 | 52 | 55 | 8 | 6 | 4,3 | 2,9 | 1,3 |
| chr6:160200144-160200644:downstream_24458 | chr6 | 160182988 | 160200144 | + | ACAT2 | NM | 53 | 1,1 | 1 | -1 | 1,41E-01 | 4,60E-01 | 14 | 23 | 8 | 14 | 79 | 74 | 83 | 126 | 0,7 | -0,5 | 1,2 |
| chr15:57580730-57581230:downstream_19802 | chr15 | 57210832 | 57580730 | + | TCF12 | NM | 34 | 1,7 | 1 | -1 | 1,41E-01 | 4,60E-01 | 2 | 16 | 18 | 21 | 22 | 4 | 107 | 79 | -1,1 | -2,8 | 1,7 |
| chr6:53131695-53132195:downstream_24745 | chr6 | 53132195 | 53213977 | - | ELOVL5 | NM | 21 | 1,5 | 1 | -1 | 1,41E-01 | NA | 5 | 14 | 10 | 7 | 19 | 19 | 43 | 52 | 0,2 | -1,3 | 1,5 |
| chr8:128199530-128200030:downstream_6718 | chr8 | 128200030 | 128209872 | - | CASC19 | NR | 5 | -3,4 | 2 | 1 | 1,41E-01 | NA | 0 | 0 | 4 | 7 | 3 | 3 | 10 | 10 | -Inf | -1,7 | -Inf |
| chr12:4669213-4669713:downstream_18782 | chr12 | 4647949 | 4669213 | + | RAD51AP1 | NM | 11 | 2,0 | 1 | -1 | 1,41E-01 | NA | 5 | 7 | 1 | 6 | 12 | 7 | 29 | 21 | 0,7 | -1,4 | 2,1 |
| chr1:21068670-21069170:downstream_89 | chr1 | 21069170 | 21113181 | - | HP1BP3 | NM | 15 | -2,0 | 1 | 1 | 1,42E-01 | NA | 0 | 3 | 13 | 8 | 20 | 15 | 29 | 34 | -2,8 | -0,9 | -1,9 |
| chr15:40765357-40765857:downstream_11447 | chr15 | 40763159 | 40765357 | + | CHST14 | NM | 15 | 2,0 | 1 | -1 | 1,42E-01 | NA | 7 | 4 | 1 | 2 | 22 | 27 | 25 | 29 | 1,9 | -0,1 | 2,0 |
| chr1:24088549-24089049:downstream_546 | chr1 | 24069855 | 24088549 | + | TCEB3 | NM | 8 | 2,5 | 2 | -1 | 1,42E-01 | NA | 7 | 5 | 3 | 2 | 11 | 3 | 6 | 25 | 1,4 | -1,1 | 2,4 |
| chr16:30580518-30581018:downstream_20294 | chr16 | 30581018 | 30583728 | - | ZNF688 | NM | 8 | 2,4 | 2 | -1 | 1,42E-01 | NA | 10 | 7 | 4 | 0 | 11 | 9 | 14 | 9 | 2,0 | -0,2 | 2,2 |
| chr5:6669675-6670175:downstream_4181 | chr5 | 6633499 | 6669675 | + | SRD5A1 | NM | 7 | -2,3 | 2 | 1 | 1,42E-01 | NA | 2 | 3 | 17 | 10 | 9 | 4 | 8 | 5 | -2,3 | -0,1 | -2,3 |
| chr16:416883-417383:downstream_12061 | chr16 | 417383 | 420569 | - | MRPL28 | NM | 14 | 2,0 | 1 | -1 | 1,42E-01 | NA | 3 | 8 | 3 | 1 | 18 | 27 | 26 | 29 | 1,6 | -0,3 | 1,9 |
| chr5:37291234-37291734:downstream_24264 | chr5 | 37291734 | 37371228 | - | NUP155 | NM | 44 | 1,2 | 1 | -1 | 1,43E-01 | 4,62E-01 | 12 | 7 | 24 | 30 | 24 | 15 | 120 | 122 | -1,5 | -2,7 | 1,2 |
| chr10:124924886-124925386:downstream_17495 | chr10 | 124913759 | 124924886 | + | BUB3 | NM | 57 | 1,1 | 1 | -1 | 1,43E-01 | 4,62E-01 | 29 | 22 | 10 | 13 | 82 | 113 | 70 | 118 | 1,2 | 0,1 | 1,1 |
| chr2:151324206-151324706:downstream_22193 | chr2 | 151324706 | 151344209 | - | RND3 | NM | 51 | -1,0 | 1 | 1 | 1,43E-01 | 4,62E-01 | 33 | 25 | 15 | 19 | 111 | 136 | 36 | 35 | 0,8 | 1,8 | -1,0 |
| chr15:25332901-25333401:downstream_11574 | chr15 | 25332807 | 25332901 | + | SNORD116-20 | NR | 172 | 0,8 | 1 | -1 | 1,43E-01 | 4,62E-01 | 442 | 378 | 188 | 212 | 36 | 47 | 28 | 42 | 1,0 | 0,3 | 0,8 |
| chr1:244571293-244571793:downstream_322 | chr1 | 244571793 | 244615436 | - | ADSS | NM | 28 | 1,3 | 1 | -1 | 1,43E-01 | 4,62E-01 | 43 | 29 | 26 | 11 | 20 | 32 | 29 | 34 | 1,0 | -0,3 | 1,3 |
| chrX:48969822-48970322:downstream_7716 | chrX | 48970322 | 48980151 | - | GPKOW | NM | 27 | 1,4 | 1 | -1 | 1,44E-01 | NA | 10 | 11 | 6 | 4 | 37 | 44 | 51 | 49 | 1,1 | -0,3 | 1,4 |
| chr9:101916473-101916973:downstream_26533 | chr9 | 101867370 | 101916473 | + | TGFBR1 | NM | 8 | 2,2 | 2 | -1 | 1,44E-01 | NA | 8 | 10 | 1 | 4 | 11 | 4 | 15 | 9 | 1,7 | -0,7 | 2,4 |
| chr17:76987297-76987797:downstream_20375 | chr17 | 76987797 | 77005899 | - | CANT1 | NM | 37 | 1,6 | 1 | -1 | 1,45E-01 | 4,65E-01 | 10 | 6 | 1 | 2 | 79 | 78 | 54 | 66 | 2,0 | 0,4 | 1,6 |
| chr15:23003603-23004103:downstream_19521 | chr15 | 22892648 | 23003603 | + | CYFIP1 | NM | 70 | -1,2 | 1 | 1 | 1,45E-01 | 4,65E-01 | 7 | 2 | 13 | 21 | 91 | 98 | 136 | 191 | -2,0 | -0,8 | -1,2 |
| chr8:30429778-30430278:downstream_26081 | chr8 | 30242016 | 30429778 | + | RBPMS | NM | 6 | 3,1 | 2 | -1 | 1,46E-01 | NA | 2 | 3 | 14 | 5 | 1 | 0 | 10 | 16 | -1,8 | -5,0 | 3,2 |
| chr5:1816167-1816667:downstream_4165 | chr5 | 1801495 | 1816167 | + | NDUFS6 | NM | 31 | 2,5 | 2 | -1 | 1,46E-01 | 4,65E-01 | 10 | 0 | 0 | 1 | 83 | 70 | 38 | 48 | 3,6 | 0,8 | 2,7 |
| chr4:184560287-184560787:downstream_23834 | chr4 | 184560787 | 184580372 | - | RWDD4 | NM | 5 | -3,5 | 2 | 1 | 1,46E-01 | NA | 16 | 4 | 4 | 2 | 3 | 13 | 0 | 0 | 1,8 | Inf | -Inf |
| chr3:176738041-176738541:downstream_2755 | chr3 | 176738541 | 176915048 | - | TBL1XR1 | NM | 64 | 1,1 | 1 | -1 | 1,46E-01 | 4,65E-01 | 10 | 19 | 40 | 43 | 31 | 25 | 225 | 121 | -1,5 | -2,6 | 1,1 |
| chrX:19377675-19378175:downstream_7479 | chrX | 19378175 | 19533379 | - | MAP3K15 | NM | 12 | 2,3 | 2 | -1 | 1,46E-01 | NA | 15 | 25 | 24 | 22 | 1 | 1 | 3 | 7 | -0,2 | -2,4 | 2,2 |
| chr1:151091007-151091507:downstream_991 | chr1 | 151043079 | 151091007 | + | GABPB2 | NM | 8 | -2,9 | 2 | 1 | 1,46E-01 | NA | 1 | 0 | 7 | 12 | 6 | 5 | 19 | 12 | -4,6 | -1,6 | -3,0 |
| chr8:101930303-101930803:downstream_26185 | chr8 | 101930803 | 101965623 | - | YWHAZ | NM | 92 | 1,0 | 1 | -1 | 1,46E-01 | 4,65E-01 | 9 | 19 | 15 | 10 | 133 | 114 | 220 | 219 | 0,2 | -0,8 | 1,0 |
| chr9:37436986-37437486:downstream_7077 | chr9 | 37422706 | 37436986 | + | GRHPR | NM | 78 | 1,1 | 1 | -1 | 1,46E-01 | 4,65E-01 | 14 | 29 | 8 | 7 | 156 | 169 | 134 | 107 | 1,5 | 0,4 | 1,1 |
| chr9:35736362-35736862:downstream_7056 | chr9 | 35736862 | 35749225 | - | GBA2 | NM | 20 | 1,8 | 1 | -1 | 1,47E-01 | NA | 3 | 13 | 3 | 2 | 20 | 45 | 35 | 41 | 1,6 | -0,2 | 1,8 |
| chr19:6467230-6467730:downstream_21084 | chr19 | 6464259 | 6467230 | + | CRB3 | NM | 11 | -2,2 | 2 | 1 | 1,47E-01 | NA | 3 | 0 | 7 | 4 | 19 | 24 | 22 | 7 | -1,8 | 0,5 | -2,3 |
| chr4:15071777-15072277:downstream_23601 | chr4 | 15004297 | 15071777 | + | CPEB2 | NM | 10 | -2,8 | 2 | 1 | 1,47E-01 | NA | 0 | 1 | 17 | 16 | 3 | 5 | 8 | 31 | -5,1 | -2,3 | -2,8 |
| chrX:153759104-153759604:downstream_26825 | chrX | 153759604 | 153775801 | - | G6PD | NM | 239 | -1,2 | 1 | 1 | 1,47E-01 | 4,65E-01 | 6 | 1 | 8 | 4 | 570 | 537 | 337 | 448 | -0,9 | 0,5 | -1,4 |
| chr15:80217196-80217696:downstream_11510 | chr15 | 80215112 | 80217196 | + | ST20-AS1 | NR | 5 | 3,3 | 2 | -1 | 1,47E-01 | NA | 3 | 9 | 0 | 0 | 9 | 14 | 1 | 6 | Inf | 1,7 | Inf |
| chr17:19580908-19581408:downstream_20322 | chr17 | 19552063 | 19580908 | + | ALDH3A2 | NM | 35 | -1,3 | 1 | 1 | 1,47E-01 | 4,65E-01 | 5 | 7 | 24 | 27 | 50 | 26 | 64 | 75 | -2,1 | -0,9 | -1,3 |
| chr5:36870962-36871462:downstream_4223 | chr5 | 36871462 | 36876796 | - | NIPBL-AS1 | NR | 142 | 0,9 | 1 | -1 | 1,47E-01 | 4,65E-01 | 123 | 153 | 8 | 13 | 332 | 400 | 47 | 62 | 3,7 | 2,7 | 0,9 |
| chr19:46190211-46190711:downstream_21526 | chr19 | 46190711 | 46195443 | - | SNRPD2 | NM | 51 | 2,3 | 2 | -1 | 1,48E-01 | 4,65E-01 | 4 | 5 | 0 | 1 | 155 | 96 | 68 | 77 | 3,4 | 0,8 | 2,7 |
| chr1:231413719-231414219:downstream_1420 | chr1 | 231376918 | 231413719 | + | GNPAT | NM | 42 | 1,5 | 1 | -1 | 1,48E-01 | 4,65E-01 | 2 | 12 | 3 | 3 | 85 | 54 | 93 | 86 | 1,2 | -0,4 | 1,6 |
| chr4:109551639-109552139:downstream_23830 | chr4 | 109541721 | 109551639 | + | RPL34 | NM | 53 | 1,0 | 1 | -1 | 1,48E-01 | 4,65E-01 | 58 | 62 | 17 | 16 | 104 | 71 | 42 | 53 | 1,9 | 0,9 | 1,0 |
| chr1:111489311-111489811:downstream_16917 | chr1 | 111489811 | 111506566 | - | LRIF1 | NM | 27 | 1,3 | 1 | -1 | 1,48E-01 | NA | 29 | 17 | 8 | 7 | 37 | 51 | 29 | 39 | 1,6 | 0,4 | 1,3 |
| chr15:80430735-80431235:downstream_19848 | chr15 | 80351909 | 80430735 | + | ZFAND6 | NM | 59 | 1,6 | 1 | -1 | 1,48E-01 | 4,65E-01 | 2 | 7 | 3 | 2 | 79 | 113 | 143 | 121 | 1,1 | -0,5 | 1,5 |
| chr9:115926422-115926922:downstream_7265 | chr9 | 115913237 | 115926422 | + | SLC31A2 | NM | 11 | 2,0 | 1 | -1 | 1,48E-01 | NA | 6 | 8 | 1 | 2 | 20 | 13 | 22 | 17 | 1,8 | -0,3 | 2,1 |
| chr22:33668561-33669061:downstream_22831 | chr22 | 33669061 | 34316416 | - | LARGE | NM | 22 | -2,6 | 2 | 1 | 1,48E-01 | NA | 0 | 0 | 4 | 12 | 8 | 8 | 70 | 74 | -Inf | -3,2 | -Inf |
| chr2:175294303-175294803:downstream_22208 | chr2 | 175260456 | 175294303 | + | SCRN3 | NM | 14 | 1,7 | 1 | -1 | 1,48E-01 | NA | 10 | 11 | 7 | 7 | 16 | 9 | 25 | 27 | 0,6 | -1,1 | 1,7 |
| chr19:46030240-46030740:downstream_14383 | chr19 | 46010687 | 46030240 | + | VASP | NM | 18 | 1,7 | 1 | -1 | 1,49E-01 | NA | 8 | 15 | 8 | 1 | 24 | 28 | 26 | 32 | 1,4 | -0,2 | 1,5 |
| chr8:103216228-103216728:downstream_26103 | chr8 | 103216728 | 103251346 | - | RRM2B | NM | 70 | 1,0 | 1 | -1 | 1,49E-01 | 4,68E-01 | 44 | 30 | 10 | 9 | 126 | 184 | 77 | 83 | 2,0 | 1,0 | 1,0 |
| chr17:37690818-37691318:downstream_20403 | chr17 | 37617763 | 37690818 | + | CDK12 | NM | 19 | 1,5 | 1 | -1 | 1,49E-01 | NA | 7 | 10 | 8 | 7 | 20 | 14 | 50 | 37 | 0,1 | -1,4 | 1,5 |
| chr1:169101960-169102460:downstream_238 | chr1 | 169075946 | 169101960 | + | ATP1B1 | NM | 627 | 0,8 | 1 | -1 | 1,50E-01 | 4,68E-01 | 13 | 15 | 33 | 21 | 495 | 664 | 1849 | 1923 | -0,9 | -1,7 | 0,8 |
| chr15:37102461-37102961:downstream_19483 | chr15 | 36871803 | 37102461 | + | C15orf41 | NM | 9 | -2,8 | 2 | 1 | 1,50E-01 | NA | 1 | 0 | 8 | 7 | 9 | 7 | 28 | 13 | -4,3 | -1,4 | -2,9 |
| chr17:30697468-30697968:downstream_20817 | chr17 | 30677156 | 30697468 | + | ZNF207 | NM | 108 | 0,8 | 1 | -1 | 1,50E-01 | 4,68E-01 | 115 | 98 | 88 | 100 | 112 | 66 | 122 | 159 | 0,2 | -0,7 | 0,8 |
| chr17:40177656-40178156:downstream_20609 | chr17 | 40169593 | 40177656 | + | NKIRAS2 | NM | 17 | 1,5 | 1 | -1 | 1,50E-01 | NA | 13 | 29 | 7 | 10 | 22 | 15 | 19 | 24 | 1,3 | -0,2 | 1,6 |
| chr10:123716102-123716602:downstream_17678 | chr10 | 123716602 | 123734743 | - | NSMCE4A | NM | 32 | 1,5 | 1 | -1 | 1,50E-01 | 4,69E-01 | 2 | 4 | 6 | 11 | 8 | 14 | 116 | 94 | -1,6 | -3,3 | 1,6 |
| chr13:51935200-51935700:downstream_18998 | chr13 | 51935700 | 52027275 | - | INTS6 | NM | 6 | 2,6 | 2 | -1 | 1,51E-01 | NA | 2 | 11 | 7 | 7 | 1 | 2 | 10 | 10 | -0,2 | -2,8 | 2,6 |
| chr12:52642709-52643209:downstream_10176 | chr12 | 52626953 | 52642709 | + | KRT7 | NM | 147 | 1,0 | 1 | -1 | 1,51E-01 | 4,70E-01 | 11 | 22 | 6 | 10 | 275 | 281 | 212 | 358 | 1,1 | 0,0 | 1,1 |
| chr14:97033453-97033953:downstream_19332 | chr14 | 96968712 | 97033453 | + | PAPOLA | NM | 57 | 1,1 | 1 | -1 | 1,51E-01 | 4,70E-01 | 10 | 29 | 14 | 17 | 73 | 68 | 125 | 124 | 0,3 | -0,8 | 1,1 |
| chr9:97330411-97330911:downstream_26436 | chr9 | 97317326 | 97330411 | + | PCAT7 | NR | 5 | 3,0 | 2 | -1 | 1,52E-01 | NA | 2 | 8 | 4 | 6 | 2 | 0 | 10 | 6 | -0,1 | -3,3 | 3,2 |
| chr11:130096073-130096573:downstream_18399 | chr11 | 130096573 | 130184607 | - | ZBTB44 | NM | 10 | -3,0 | 2 | 1 | 1,52E-01 | NA | 1 | 0 | 19 | 2 | 2 | 14 | 28 | 17 | -4,8 | -1,6 | -3,2 |
| chr13:48621358-48621858:downstream_19039 | chr13 | 48611702 | 48621358 | + | NUDT15 | NM | 11 | 2,7 | 2 | -1 | 1,52E-01 | NA | 7 | 1 | 1 | 0 | 22 | 23 | 15 | 21 | 2,6 | 0,3 | 2,3 |
| chr5:140042065-140042565:downstream_4462 | chr5 | 140027383 | 140042065 | + | IK | NM | 16 | 2,2 | 2 | -1 | 1,52E-01 | NA | 6 | 3 | 0 | 2 | 37 | 19 | 29 | 30 | 2,4 | -0,1 | 2,5 |
| chr4:110608872-110609372:downstream_3488 | chr4 | 110481354 | 110608872 | + | CCDC109B | NM | 6 | 2,8 | 2 | -1 | 1,52E-01 | NA | 1 | 9 | 3 | 7 | 0 | 4 | 18 | 9 | 0,0 | -2,8 | 2,8 |
| chr3:149104370-149104870:downstream_23472 | chr3 | 149095564 | 149104370 | + | TM4SF1-AS1 | NR | 4 | -3,4 | 2 | 1 | 1,52E-01 | NA | 1 | 0 | 3 | 7 | 4 | 6 | 7 | 3 | -3,5 | -0,1 | -3,5 |
| chr14:93648040-93648540:downstream_11279 | chr14 | 93648540 | 93651249 | - | MOAP1 | NM | 43 | -1,3 | 1 | 1 | 1,53E-01 | 4,73E-01 | 12 | 4 | 7 | 14 | 86 | 116 | 74 | 34 | -0,4 | 0,9 | -1,3 |
| chr13:73330336-73330836:downstream_18943 | chr13 | 73301886 | 73330336 | + | BORA | NM | 15 | 2,1 | 1 | -1 | 1,53E-01 | NA | 5 | 11 | 0 | 2 | 14 | 41 | 21 | 23 | 2,7 | 0,3 | 2,3 |
| chr19:49470136-49470636:downstream_14464 | chr19 | 49468565 | 49470136 | + | FTL | NM | 2208 | -0,7 | 0 | 1 | 1,53E-01 | 4,73E-01 | 36 | 38 | 21 | 27 | 6957 | 5528 | 2435 | 2623 | 0,6 | 1,3 | -0,7 |
| chr17:7126115-7126615:downstream_12794 | chr17 | 7126615 | 7126698 | - | MIR324 | NR | 16 | -1,5 | 1 | 1 | 1,53E-01 | NA | 15 | 12 | 11 | 19 | 21 | 27 | 10 | 9 | -0,1 | 1,4 | -1,5 |
| chr15:64979272-64979772:downstream_19656 | chr15 | 64979772 | 64995480 | - | OAZ2 | NM | 23 | 1,6 | 1 | -1 | 1,53E-01 | NA | 10 | 6 | 1 | 4 | 39 | 35 | 53 | 33 | 1,5 | -0,2 | 1,7 |
| chr13:52986236-52986736:downstream_19128 | chr13 | 52986736 | 53024813 | - | VPS36 | NM | 10 | 1,9 | 1 | -1 | 1,54E-01 | NA | 10 | 14 | 4 | 5 | 7 | 15 | 11 | 19 | 1,4 | -0,5 | 1,9 |
| chr11:62434923-62435423:downstream_9398 | chr11 | 62432778 | 62434923 | + | METTL12 | NM | 12 | 2,1 | 1 | -1 | 1,54E-01 | NA | 14 | 11 | 0 | 3 | 28 | 10 | 19 | 7 | 2,9 | 0,5 | 2,4 |
| chr5:89810944-89811444:downstream_24214 | chr5 | 89811444 | 89825401 | - | LYSMD3 | NM | 25 | 1,3 | 1 | -1 | 1,54E-01 | NA | 29 | 28 | 8 | 7 | 31 | 49 | 24 | 27 | 1,9 | 0,7 | 1,3 |
| chr11:67354124-67354624:downstream_9488 | chr11 | 67351065 | 67354124 | + | GSTP1 | NM | 420 | 1,0 | 1 | -1 | 1,54E-01 | 4,73E-01 | 20 | 14 | 10 | 2 | 1180 | 833 | 564 | 734 | 1,5 | 0,6 | 0,8 |
| chr3:149234521-149235021:downstream_23510 | chr3 | 149235021 | 149421060 | - | WWTR1 | NM | 114 | -2,2 | 2 | 1 | 1,54E-01 | 4,73E-01 | 1 | 0 | 10 | 5 | 105 | 104 | 344 | 348 | -4,2 | -1,7 | -2,4 |
| chr4:83273966-83274466:downstream_23694 | chr4 | 83274466 | 83295149 | - | HNRNPD | NM | 15 | 1,6 | 1 | -1 | 1,54E-01 | NA | 14 | 6 | 6 | 4 | 21 | 16 | 25 | 31 | 1,0 | -0,6 | 1,6 |
| chr6:166337035-166337535:downstream_24842 | chr6 | 166337535 | 166401527 | - | LINC00473 | NR | 99 | -1,0 | 1 | 1 | 1,54E-01 | 4,73E-01 | 12 | 5 | 29 | 43 | 127 | 99 | 262 | 217 | -2,1 | -1,1 | -1,0 |
| chr1:226032732-226033232:downstream_1390 | chr1 | 226033232 | 226070420 | - | TMEM63A | NM | 10 | 2,5 | 2 | -1 | 1,54E-01 | NA | 4 | 32 | 10 | 17 | 0 | 3 | 4 | 7 | 0,4 | -2,0 | 2,4 |
| chr7:83587158-83587658:downstream_5582 | chr7 | 83587658 | 83824217 | - | SEMA3A | NM | 42 | -1,3 | 1 | 1 | 1,54E-01 | 4,73E-01 | 2 | 4 | 36 | 42 | 22 | 15 | 116 | 103 | -3,8 | -2,5 | -1,3 |
| chr6:26249869-26250369:downstream_4923 | chr6 | 26250369 | 26250835 | - | HIST1H3F | NM | 6 | 2,8 | 2 | -1 | 1,54E-01 | NA | 10 | 2 | 4 | 2 | 6 | 0 | 7 | 15 | 0,8 | -1,9 | 2,7 |
| chr8:30624520-30625020:downstream_26173 | chr8 | 30601681 | 30624520 | + | UBXN8 | NM | 16 | 2,1 | 1 | -1 | 1,54E-01 | NA | 4 | 9 | 3 | 0 | 29 | 36 | 24 | 25 | 2,2 | 0,4 | 1,8 |
| chr20:37207504-37208004:downstream_22524 | chr20 | 37101448 | 37207504 | + | RALGAPB | NM | 9 | 2,2 | 2 | -1 | 1,55E-01 | NA | 3 | 9 | 1 | 10 | 7 | 2 | 17 | 23 | 0,1 | -2,2 | 2,3 |
| chr12:53840427-53840927:downstream_18764 | chr12 | 53835432 | 53840427 | + | PRR13 | NM | 85 | 1,1 | 1 | -1 | 1,55E-01 | 4,74E-01 | 18 | 20 | 7 | 4 | 174 | 224 | 134 | 102 | 1,8 | 0,8 | 1,0 |
| chr12:85638883-85639383:downstream_9894 | chr12 | 85430098 | 85638883 | + | LRRIQ1 | NM | 11 | -1,8 | 1 | 1 | 1,55E-01 | NA | 3 | 5 | 24 | 22 | 9 | 5 | 11 | 11 | -2,5 | -0,7 | -1,8 |
| chr5:60841999-60842499:downstream_3964 | chr5 | 60628099 | 60841999 | + | ZSWIM6 | NM | 6 | -3,3 | 2 | 1 | 1,55E-01 | NA | 0 | 0 | 7 | 11 | 4 | 0 | 14 | 9 | -Inf | -2,5 | -Inf |
| chr2:118589953-118590453:downstream_2210 | chr2 | 118572254 | 118589953 | + | DDX18 | NM | 55 | 1,2 | 1 | -1 | 1,55E-01 | 4,75E-01 | 25 | 28 | 8 | 8 | 65 | 153 | 58 | 94 | 1,7 | 0,5 | 1,2 |
| chr10:13358937-13359437:downstream_17762 | chr10 | 13359437 | 13390298 | - | SEPHS1 | NM | 20 | 1,6 | 1 | -1 | 1,56E-01 | NA | 4 | 5 | 6 | 2 | 15 | 24 | 57 | 46 | 0,1 | -1,4 | 1,6 |
| chr3:32566962-32567462:downstream_2870 | chr3 | 32567462 | 32612366 | - | DYNC1LI1 | NM | 42 | -1,6 | 1 | 1 | 1,56E-01 | 4,75E-01 | 1 | 3 | 7 | 11 | 76 | 39 | 111 | 88 | -2,2 | -0,8 | -1,5 |
| chr19:39326527-39327027:downstream_21174 | chr19 | 39327027 | 39342979 | - | HNRNPL | NM | 134 | 1,1 | 1 | -1 | 1,57E-01 | 4,77E-01 | 11 | 5 | 6 | 6 | 151 | 253 | 288 | 354 | 0,4 | -0,7 | 1,1 |
| chr22:30972111-30972611:downstream_22873 | chr22 | 30972611 | 31003000 | - | PES1 | NM | 9 | 2,7 | 2 | -1 | 1,57E-01 | NA | 6 | 6 | 0 | 1 | 17 | 21 | 6 | 16 | 3,8 | 0,8 | 3,0 |
| chr3:121488109-121488609:downstream_23210 | chr3 | 121488609 | 121553926 | - | IQCB1 | NM | 11 | 2,2 | 2 | -1 | 1,57E-01 | NA | 0 | 10 | 7 | 3 | 6 | 7 | 28 | 30 | -0,1 | -2,2 | 2,1 |
| chr16:2261102-2261602:downstream_12112 | chr16 | 2261602 | 2264822 | - | PGP | NM | 12 | 1,7 | 1 | -1 | 1,57E-01 | NA | 10 | 15 | 8 | 6 | 8 | 13 | 24 | 14 | 0,8 | -0,9 | 1,7 |
| chr19:16660147-16660647:downstream_14100 | chr19 | 16660647 | 16683193 | - | SLC35E1 | NM | 18 | 1,7 | 1 | -1 | 1,57E-01 | NA | 2 | 12 | 3 | 3 | 24 | 27 | 32 | 43 | 1,2 | -0,6 | 1,8 |
| chr4:38797375-38797875:downstream_3654 | chr4 | 38797875 | 38806412 | - | TLR1 | NM | 43 | -1,3 | 1 | 1 | 1,57E-01 | 4,78E-01 | 2 | 4 | 25 | 29 | 28 | 33 | 100 | 121 | -3,1 | -1,8 | -1,2 |
| chr12:41466213-41466713:downstream_18520 | chr12 | 41086243 | 41466213 | + | CNTN1 | NM | 297 | -0,8 | 1 | 1 | 1,58E-01 | 4,78E-01 | 7 | 10 | 29 | 30 | 404 | 366 | 793 | 733 | -1,8 | -1,0 | -0,8 |
| chr3:12624599-12625099:downstream_2828 | chr3 | 12625099 | 12705700 | - | RAF1 | NM | 9 | 2,4 | 2 | -1 | 1,58E-01 | NA | 3 | 6 | 0 | 5 | 12 | 1 | 18 | 25 | 0,9 | -1,7 | 2,6 |
| chr10:101515894-101516394:downstream_8672 | chr10 | 101491957 | 101515894 | + | CUTC | NM | 10 | 2,4 | 2 | -1 | 1,58E-01 | NA | 19 | 0 | 4 | 8 | 7 | 4 | 21 | 15 | 0,6 | -1,8 | 2,4 |
| chr10:114578504-114579004:downstream_8226 | chr10 | 114206755 | 114578504 | + | VTI1A | NM | 20 | 1,6 | 1 | -1 | 1,58E-01 | NA | 4 | 13 | 26 | 21 | 2 | 8 | 45 | 40 | -1,5 | -3,1 | 1,6 |
| chr9:140138159-140138659:downstream_7443 | chr9 | 140135710 | 140138159 | + | TUBB4B | NM | 107 | 1,7 | 1 | -1 | 1,59E-01 | 4,80E-01 | 6 | 6 | 1 | 1 | 267 | 261 | 164 | 150 | 2,4 | 0,7 | 1,6 |
| chr7:2474216-2474716:downstream_25450 | chr7 | 2443194 | 2474216 | + | CHST12 | NM | 4 | -3,7 | 3 | 1 | 1,59E-01 | NA | 0 | 0 | 15 | 8 | 0 | 2 | 1 | 8 | -Inf | -2,3 | -Inf |
| chr5:158583916-158584416:downstream_24338 | chr5 | 158584416 | 158637061 | - | RNF145 | NM | 91 | 1,3 | 1 | -1 | 1,59E-01 | 4,81E-01 | 1 | 5 | 11 | 4 | 41 | 51 | 345 | 266 | -1,4 | -2,7 | 1,3 |
| chrX:13729860-13730360:downstream_27084 | chrX | 13730360 | 13752754 | - | TRAPPC2 | NM | 15 | 1,7 | 1 | -1 | 1,60E-01 | NA | 9 | 26 | 8 | 6 | 9 | 22 | 24 | 16 | 1,3 | -0,4 | 1,7 |
| chr1:24799473-24799973:downstream_93 | chr1 | 24742244 | 24799473 | + | NIPAL3 | NM | 14 | -1,8 | 1 | 1 | 1,60E-01 | NA | 2 | 1 | 11 | 12 | 19 | 11 | 31 | 25 | -2,8 | -0,9 | -1,8 |
| chr14:50084905-50085405:downstream_11130 | chr14 | 50085405 | 50087403 | - | RPL36AL | NM | 626 | 0,8 | 1 | -1 | 1,60E-01 | 4,83E-01 | 52 | 61 | 18 | 17 | 1970 | 1170 | 1001 | 717 | 1,7 | 0,9 | 0,8 |
| chr10:14560055-14560555:downstream_17560 | chr10 | 14560555 | 14816896 | - | FAM107B | NM | 47 | 1,2 | 1 | -1 | 1,61E-01 | 4,83E-01 | 5 | 5 | 10 | 7 | 39 | 33 | 128 | 148 | -0,7 | -1,9 | 1,2 |
| chr6:159057006-159057506:downstream_24736 | chr6 | 159057506 | 159065818 | - | DYNLT1 | NM | 18 | 2,0 | 1 | -1 | 1,61E-01 | NA | 9 | 5 | 1 | 1 | 49 | 28 | 26 | 21 | 2,6 | 0,7 | 2,0 |
| chr1:155708800-155709300:downstream_16641 | chr1 | 155658881 | 155708800 | + | DAP3 | NM | 8 | -2,2 | 2 | 1 | 1,61E-01 | NA | 5 | 1 | 7 | 6 | 9 | 22 | 10 | 5 | -1,1 | 1,1 | -2,2 |
| chr8:21995032-21995532:downstream_6480 | chr8 | 21995532 | 21999448 | - | REEP4 | NM | 25 | 1,9 | 1 | -1 | 1,61E-01 | NA | 5 | 4 | 3 | 0 | 42 | 55 | 43 | 48 | 1,7 | 0,1 | 1,6 |
| chr12:88442289-88442789:downstream_9895 | chr12 | 88442789 | 88535993 | - | CEP290 | NM | 18 | 1,8 | 1 | -1 | 1,61E-01 | NA | 2 | 22 | 7 | 16 | 10 | 13 | 24 | 48 | 0,1 | -1,7 | 1,8 |
| chr3:120346514-120347014:downstream_3133 | chr3 | 120347014 | 120401418 | - | HGD | NM | 40 | -1,7 | 1 | 1 | 1,61E-01 | 4,85E-01 | 1 | 2 | 6 | 7 | 48 | 76 | 68 | 110 | -2,1 | -0,5 | -1,6 |
| chr1:173755840-173756340:downstream_1231 | chr1 | 173684079 | 173755840 | + | KLHL20 | NM | 4 | 3,4 | 2 | -1 | 1,62E-01 | NA | 3 | 1 | 13 | 4 | 0 | 0 | 4 | 9 | -2,0 | -Inf | Inf |
| chr6:170843703-170844203:downstream_5465 | chr6 | 170844203 | 170862417 | - | PSMB1 | NM | 90 | 0,8 | 1 | -1 | 1,62E-01 | 4,85E-01 | 50 | 36 | 25 | 28 | 124 | 151 | 142 | 162 | 0,7 | -0,1 | 0,8 |
| chr12:106456624-106457124:downstream_10398 | chr12 | 106457124 | 106533811 | - | NUAK1 | NM | 6 | 2,4 | 2 | -1 | 1,62E-01 | NA | 6 | 4 | 1 | 2 | 5 | 6 | 13 | 11 | 1,3 | -1,1 | 2,4 |
| chr10:91180759-91181259:downstream_8632 | chr10 | 91174324 | 91180759 | + | IFIT5 | NM | 10 | 2,0 | 1 | -1 | 1,62E-01 | NA | 11 | 11 | 1 | 2 | 18 | 15 | 13 | 11 | 2,5 | 0,4 | 2,0 |
| chr8:52729634-52730134:downstream_26043 | chr8 | 52730134 | 52811746 | - | PCMTD1 | NM | 41 | 1,2 | 1 | -1 | 1,62E-01 | 4,85E-01 | 9 | 22 | 15 | 30 | 20 | 38 | 109 | 87 | -0,5 | -1,8 | 1,2 |
| chr6:163999628-164000128:downstream_25158 | chr6 | 163835674 | 163999628 | + | QKI | NM | 8 | -3,0 | 2 | 1 | 1,62E-01 | NA | 0 | 0 | 21 | 10 | 1 | 2 | 13 | 18 | -Inf | -3,5 | -Inf |
| chr5:64813092-64813592:downstream_24062 | chr5 | 64813592 | 64858995 | - | CENPK | NM | 18 | 1,8 | 1 | -1 | 1,63E-01 | NA | 2 | 15 | 7 | 2 | 22 | 23 | 39 | 39 | 0,9 | -0,8 | 1,7 |
| chr7:29959218-29959718:downstream_25718 | chr7 | 29959718 | 30029905 | - | SCRN1 | NM | 38 | 1,4 | 1 | -1 | 1,63E-01 | 4,85E-01 | 3 | 6 | 7 | 2 | 36 | 45 | 95 | 112 | -0,1 | -1,3 | 1,3 |
| chr4:57896736-57897236:downstream_23701 | chr4 | 57897236 | 57976551 | - | IGFBP7 | NM | 67 | -1,2 | 1 | 1 | 1,63E-01 | 4,85E-01 | 5 | 5 | 7 | 4 | 195 | 159 | 72 | 91 | -0,2 | 1,1 | -1,3 |
| chr11:18299716-18300216:downstream_18050 | chr11 | 18300216 | 18343721 | - | HPS5 | NM | 22 | 1,6 | 1 | -1 | 1,63E-01 | NA | 5 | 10 | 3 | 2 | 38 | 36 | 39 | 41 | 1,5 | -0,1 | 1,6 |
| chr8:56980238-56980738:downstream_26102 | chr8 | 56980738 | 56987140 | - | RPS20 | NM | 36 | 1,2 | 1 | -1 | 1,63E-01 | 4,85E-01 | 72 | 75 | 8 | 7 | 57 | 44 | 8 | 16 | 3,2 | 2,1 | 1,1 |
| chr7:130561717-130562217:downstream_6092 | chr7 | 130562217 | 130562298 | - | MIR29B1 | NR | 29 | 1,7 | 1 | -1 | 1,63E-01 | 4,85E-01 | 37 | 33 | 71 | 71 | 2 | 0 | 7 | 10 | -1,0 | -2,8 | 1,7 |
| chr12:93970521-93971021:downstream_18845 | chr12 | 93963597 | 93970521 | + | SOCS2 | NM | 7 | -3,2 | 2 | 1 | 1,64E-01 | NA | 10 | 18 | 6 | 0 | 11 | 9 | 0 | 0 | 2,3 | Inf | -Inf |
| chr1:84964033-84964533:downstream_804 | chr1 | 84944919 | 84964033 | + | RPF1 | NM | 12 | 2,0 | 1 | -1 | 1,64E-01 | NA | 16 | 25 | 4 | 8 | 13 | 4 | 18 | 3 | 1,7 | -0,3 | 2,1 |
| chr1:15897693-15898193:downstream_479 | chr1 | 15898193 | 15911605 | - | AGMAT | NM | 15 | -1,5 | 1 | 1 | 1,64E-01 | NA | 13 | 11 | 28 | 22 | 11 | 15 | 8 | 10 | -1,1 | 0,5 | -1,5 |
| chr16:135850-136350:downstream_20123 | chr16 | 127017 | 135850 | + | MPG | NM | 6 | 2,5 | 2 | -1 | 1,64E-01 | NA | 5 | 9 | 3 | 1 | 7 | 5 | 13 | 4 | 1,9 | -0,6 | 2,5 |
| chr9:109773796-109774296:downstream_6909 | chr9 | 109625377 | 109773796 | + | ZNF462 | NM | 13 | -2,7 | 2 | 1 | 1,64E-01 | NA | 1 | 0 | 33 | 12 | 7 | 0 | 28 | 24 | -5,8 | -3,0 | -2,8 |
| chr8:104455680-104456180:downstream_6675 | chr8 | 104426941 | 104455680 | + | DCAF13 | NM | 28 | 1,3 | 1 | -1 | 1,64E-01 | NA | 35 | 17 | 13 | 17 | 19 | 40 | 43 | 38 | 0,8 | -0,5 | 1,3 |
| chr7:100276792-100277292:downstream_6002 | chr7 | 100271362 | 100276792 | + | GNB2 | NM | 5 | 3,7 | 3 | -1 | 1,64E-01 | NA | 3 | 6 | 0 | 0 | 20 | 0 | 1 | 10 | Inf | 0,9 | Inf |
| chr8:128808274-128808774:downstream_6722 | chr8 | 128808207 | 128808274 | + | MIR1204 | NR | 50 | 1,1 | 1 | -1 | 1,64E-01 | 4,86E-01 | 134 | 121 | 38 | 21 | 35 | 21 | 14 | 13 | 2,1 | 1,1 | 1,1 |
| chr10:70041916-70042416:downstream_17698 | chr10 | 70042416 | 70092684 | - | PBLD | NM | 14 | -1,9 | 1 | 1 | 1,64E-01 | NA | 3 | 0 | 11 | 7 | 24 | 15 | 25 | 30 | -2,5 | -0,5 | -2,0 |
| chr8:22877710-22878210:downstream_26095 | chr8 | 22844929 | 22877710 | + | RHOBTB2 | NM | 11 | 2,1 | 1 | -1 | 1,64E-01 | NA | 17 | 22 | 25 | 4 | 2 | 3 | 7 | 7 | 0,4 | -1,6 | 2,0 |
| chr1:150532870-150533370:downstream_985 | chr1 | 150533370 | 150547028 | - | ADAMTSL4-AS1 | NR | 14 | 1,8 | 1 | -1 | 1,65E-01 | NA | 28 | 18 | 1 | 3 | 20 | 25 | 14 | 4 | 3,3 | 1,3 | 2,0 |
| chr13:48836232-48836732:downstream_10750 | chr13 | 48807273 | 48836232 | + | ITM2B | NM | 90 | 1,3 | 1 | -1 | 1,65E-01 | 4,86E-01 | 20 | 3 | 3 | 5 | 200 | 175 | 150 | 165 | 1,5 | 0,3 | 1,3 |
| chr2:28002608-28003108:downstream_22085 | chr2 | 27994583 | 28002608 | + | MRPL33 | NM | 64 | 1,5 | 1 | -1 | 1,65E-01 | 4,86E-01 | 13 | 6 | 4 | 0 | 175 | 153 | 88 | 72 | 2,2 | 1,0 | 1,1 |
| chr3:195686291-195686791:downstream_2785 | chr3 | 195686791 | 195717150 | - | SDHAP1 | NR | 5 | 2,5 | 2 | -1 | 1,65E-01 | NA | 6 | 6 | 7 | 5 | 1 | 2 | 8 | 7 | 0,0 | -2,5 | 2,5 |
| chrX:23721276-23721776:downstream_26600 | chrX | 23721776 | 23761407 | - | ACOT9 | NM | 62 | 1,0 | 1 | -1 | 1,65E-01 | 4,86E-01 | 15 | 20 | 24 | 26 | 50 | 57 | 170 | 130 | -0,5 | -1,5 | 1,0 |
| chr19:8387280-8387780:downstream_13959 | chr19 | 8386383 | 8387280 | + | RPS28 | NM | 9 | -1,9 | 1 | 1 | 1,65E-01 | NA | 10 | 10 | 11 | 11 | 12 | 10 | 4 | 2 | -0,2 | 1,7 | -1,9 |
| chr20:47860614-47861114:downstream_14809 | chr20 | 47835831 | 47860614 | + | DDX27 | NM | 31 | 1,3 | 1 | -1 | 1,66E-01 | 4,86E-01 | 14 | 16 | 6 | 3 | 81 | 42 | 45 | 43 | 1,8 | 0,5 | 1,3 |
| chr5:108669909-108670409:downstream_4359 | chr5 | 108670409 | 108745675 | - | PJA2 | NM | 53 | -1,9 | 1 | 1 | 1,66E-01 | 4,86E-01 | 2 | 0 | 3 | 16 | 32 | 67 | 129 | 176 | -3,5 | -1,6 | -1,9 |
| chr2:105965271-105965771:downstream_21778 | chr2 | 105953815 | 105965271 | + | C2orf49 | NM | 15 | 1,8 | 1 | -1 | 1,66E-01 | NA | 12 | 15 | 1 | 3 | 15 | 39 | 24 | 11 | 2,6 | 0,6 | 1,9 |
| chr5:179204287-179204787:downstream_4140 | chr5 | 179159850 | 179204287 | + | MAML1 | NM | 15 | -1,9 | 1 | 1 | 1,66E-01 | NA | 1 | 2 | 7 | 7 | 14 | 32 | 31 | 30 | -2,3 | -0,4 | -1,9 |
| chr1:87213867-87214367:downstream_17232 | chr1 | 87170252 | 87213867 | + | SH3GLB1 | NM | 11 | 1,8 | 1 | -1 | 1,66E-01 | NA | 7 | 3 | 4 | 7 | 5 | 9 | 25 | 30 | -0,2 | -2,0 | 1,8 |
| chr11:60657519-60658019:downstream_9362 | chr11 | 60658019 | 60674061 | - | PRPF19 | NM | 27 | 1,4 | 1 | -1 | 1,67E-01 | NA | 11 | 9 | 0 | 7 | 48 | 43 | 56 | 43 | 1,6 | -0,1 | 1,7 |
| chr12:10740576-10741076:downstream_9819 | chr12 | 10741076 | 10752434 | - | KLRAP1 | NR | 13 | -1,7 | 1 | 1 | 1,67E-01 | NA | 9 | 8 | 7 | 7 | 15 | 42 | 8 | 7 | 0,2 | 1,9 | -1,7 |
| chr9:119159936-119160436:downstream_7281 | chr9 | 119160436 | 119162885 | - | PAPPA-AS1 | NR | 5 | -3,4 | 2 | 1 | 1,67E-01 | NA | 0 | 0 | 7 | 15 | 2 | 0 | 11 | 6 | -Inf | -2,8 | -Inf |
| chr6:37450196-37450696:downstream_5114 | chr6 | 37450696 | 37467700 | - | CCDC167 | NM | 4 | 2,7 | 2 | -1 | 1,67E-01 | NA | 5 | 7 | 4 | 2 | 2 | 2 | 6 | 6 | 1,0 | -1,7 | 2,7 |
| chrX:14861028-14861528:downstream_26814 | chrX | 14861528 | 14891184 | - | FANCB | NM | 7 | -3,2 | 2 | 1 | 1,67E-01 | NA | 0 | 0 | 13 | 9 | 0 | 4 | 19 | 8 | -Inf | -2,8 | -Inf |
| chr5:32124323-32124823:downstream_4211 | chr5 | 32124823 | 32174425 | - | GOLPH3 | NM | 24 | 1,5 | 1 | -1 | 1,67E-01 | NA | 4 | 6 | 4 | 3 | 23 | 34 | 70 | 51 | 0,4 | -1,1 | 1,5 |
| chr2:171823643-171824143:downstream_21943 | chr2 | 171784947 | 171823643 | + | GORASP2 | NM | 13 | -2,7 | 2 | 1 | 1,68E-01 | NA | 1 | 0 | 0 | 8 | 17 | 17 | 28 | 29 | -3,3 | -0,7 | -2,6 |
| chr19:58724928-58725428:downstream_21626 | chr19 | 58694355 | 58724928 | + | ZNF274 | NM | 17 | 1,7 | 1 | -1 | 1,68E-01 | NA | 4 | 7 | 1 | 4 | 29 | 14 | 42 | 36 | 1,0 | -0,9 | 1,8 |
| chr20:5917978-5918478:downstream_22589 | chr20 | 5918478 | 5931204 | - | TRMT6 | NM | 35 | 1,9 | 1 | -1 | 1,68E-01 | 4,92E-01 | 4 | 6 | 1 | 1 | 56 | 94 | 50 | 65 | 2,2 | 0,4 | 1,8 |
| chr17:48830072-48830572:downstream_20563 | chr17 | 48796925 | 48830072 | + | LUC7L3 | NM | 65 | 0,9 | 1 | -1 | 1,68E-01 | 4,92E-01 | 59 | 46 | 51 | 31 | 58 | 76 | 106 | 88 | 0,4 | -0,5 | 0,9 |
| chr14:55820329-55820829:downstream_19226 | chr14 | 55738020 | 55820329 | + | FBXO34 | NM | 20 | 1,7 | 1 | -1 | 1,68E-01 | NA | 6 | 0 | 13 | 10 | 5 | 6 | 67 | 58 | -2,0 | -3,5 | 1,6 |
| chr1:70717701-70718201:downstream_17279 | chr1 | 70671364 | 70717701 | + | SRSF11 | NM | 112 | -1,0 | 1 | 1 | 1,69E-01 | 4,92E-01 | 7 | 5 | 18 | 16 | 139 | 200 | 260 | 249 | -1,6 | -0,6 | -1,0 |

| | | | | | | | | | | | | | | | | | | | | | | | |
|---|---|---|---|---|---|---|---|---|---|---|---|---|---|---|---|---|---|---|---|---|---|---|---|
| chr4:110630644-110631144:downstream_3810 | chr4 | 110631144 | 110651242 | - | PLA2G12A | NM | 8 | 2,4 | 2 | -1 | 1,69E-01 | NA | 3 | 9 | 4 | 1 | 14 | 1 | 14 | 17 | 1,3 | -1,1 | 2,3 |
| chr1:181059979-181060479:downstream_1253 | chr1 | 181057637 | 181059979 | + | IER5 | NM | 76 | -0,9 | 1 | 1 | 1,69E-01 | 4,92E-01 | 76 | 65 | 43 | 26 | 154 | 159 | 38 | 44 | 1,0 | 1,9 | -0,9 |
| chr10:120439993-120440493:downstream_8353 | chr10 | 120440493 | 120514758 | - | CACUL1 | NM | 9 | 2,0 | 1 | -1 | 1,69E-01 | NA | 4 | 15 | 3 | 9 | 6 | 5 | 15 | 12 | 0,7 | -1,4 | 2,1 |
| chr17:58119885-58120385:downstream_13280 | chr17 | 58120385 | 58120466 | - | MIR4737 | NR | 3 | -3,5 | 3 | 1 | 1,70E-01 | NA | 4 | 0 | 6 | 5 | 1 | 9 | 1 | 1 | -1,4 | 2,1 | -3,5 |
| chr12:62991363-62991863:downstream_18687 | chr12 | 62860596 | 62991363 | + | MON2 | NM | 7 | -3,1 | 2 | 1 | 1,70E-01 | NA | 1 | 0 | 14 | 14 | 5 | 0 | 14 | 6 | -5,1 | -2,0 | -3,1 |
| chr2:54045956-54046456:downstream_21904 | chr2 | 54014067 | 54045956 | + | ERLEC1 | NM | 33 | 1,2 | 1 | -1 | 1,70E-01 | 4,93E-01 | 16 | 23 | 13 | 11 | 54 | 32 | 64 | 52 | 0,7 | -0,4 | 1,2 |
| chrX:117583923-117584423:downstream_27107 | chrX | 117480035 | 117583923 | + | WDR44 | NM | 10 | -2,8 | 2 | 1 | 1,70E-01 | NA | 0 | 0 | 4 | 9 | 9 | 1 | 24 | 36 | -Inf | -2,6 | -Inf |
| chr19:14163717-14164217:downstream_13735 | chr19 | 14142551 | 14163717 | + | IL27RA | NM | 16 | -1,6 | 1 | 1 | 1,70E-01 | NA | 10 | 1 | 10 | 9 | 32 | 34 | 18 | 18 | -0,8 | 0,9 | -1,7 |
| chr12:67708472-67708972:downstream_10300 | chr12 | 67663060 | 67708472 | + | CAND1 | NM | 25 | 1,3 | 1 | -1 | 1,70E-01 | NA | 15 | 10 | 28 | 34 | 9 | 8 | 56 | 43 | -1,3 | -2,6 | 1,3 |
| chr6:64293493-64293993:downstream_5212 | chr6 | 64281916 | 64293493 | + | PTP4A1 | NM | 76 | 0,8 | 1 | -1 | 1,70E-01 | 4,93E-01 | 50 | 59 | 40 | 49 | 86 | 79 | 122 | 120 | 0,3 | -0,5 | 0,8 |
| chr14:91923324-91923824:downstream_19397 | chr14 | 91923824 | 91976824 | - | SMEK1 | NM | 33 | 1,4 | 1 | -1 | 1,70E-01 | 4,93E-01 | 11 | 7 | 1 | 8 | 64 | 27 | 72 | 74 | 0,9 | -0,7 | 1,5 |
| chr17:7417935-7418435:downstream_12804 | chr17 | 7387697 | 7417935 | + | POLR2A | NM | 11 | 2,8 | 2 | -1 | 1,70E-01 | NA | 1 | 10 | 0 | 0 | 19 | 40 | 8 | 7 | Inf | 1,9 | Inf |
| chr10:73506813-73507313:downstream_8298 | chr10 | 73507313 | 73533337 | - | C10orf54 | NM | 4 | 3,6 | 3 | -1 | 1,71E-01 | NA | 2 | 8 | 0 | 1 | 8 | 0 | 4 | 6 | 3,6 | -0,3 | 3,9 |
| chr7:66134589-66135089:downstream_5870 | chr7 | 66119505 | 66134589 | + | LOC100996437 | NR | 9 | 2,2 | 2 | -1 | 1,71E-01 | NA | 13 | 28 | 8 | 1 | 3 | 6 | 6 | 3 | 2,2 | 0,0 | 2,1 |
| chr1:235612283-235612783:downstream_17312 | chr1 | 235530674 | 235612283 | + | TBCE | NM | 11 | 2,2 | 2 | -1 | 1,71E-01 | NA | 2 | 2 | 8 | 14 | 2 | 0 | 33 | 27 | -2,3 | -4,6 | 2,3 |
| chr18:33559250-33559750:downstream_20832 | chr18 | 33552587 | 33559250 | + | C18orf21 | NM | 39 | 1,3 | 1 | -1 | 1,72E-01 | 4,95E-01 | 10 | 16 | 3 | 3 | 96 | 82 | 57 | 48 | 2,1 | 0,8 | 1,4 |
| chr8:25365425-25365925:downstream_6501 | chr8 | 25316512 | 25365425 | + | CDCA2 | NM | 30 | -1,5 | 1 | 1 | 1,72E-01 | 4,95E-01 | 2 | 2 | 31 | 24 | 21 | 13 | 54 | 96 | -3,6 | -2,2 | -1,5 |
| chr12:102467472-102467972:downstream_9907 | chr12 | 102467972 | 102512361 | - | NUP37 | NM | 16 | 1,8 | 1 | -1 | 1,72E-01 | NA | 16 | 7 | 17 | 24 | 9 | 0 | 28 | 27 | -0,8 | -2,6 | 1,8 |
| chr18:2914090-2914590:downstream_13479 | chr18 | 2847027 | 2914090 | + | EMILIN2 | NM | 7 | 2,2 | 2 | -1 | 1,72E-01 | NA | 6 | 13 | 4 | 4 | 7 | 2 | 10 | 8 | 1,1 | -1,1 | 2,2 |
| chr17:16373962-16374462:downstream_20562 | chr17 | 16342300 | 16373962 | + | LRRC75A-AS1 | NR | 14 | 1,7 | 1 | -1 | 1,73E-01 | NA | 32 | 15 | 6 | 6 | 21 | 11 | 18 | 7 | 2,0 | 0,3 | 1,7 |
| chr7:130606705-130607205:downstream_6093 | chr7 | 130598222 | 130606705 | + | LOC100506860 | NR | 6 | -2,6 | 2 | 1 | 1,73E-01 | NA | 9 | 6 | 8 | 4 | 5 | 13 | 3 | 0 | 0,2 | 2,6 | -2,4 |
| chr1:43168020-43168520:downstream_677 | chr1 | 43148065 | 43168020 | + | YBX1 | NM | 17 | 2,1 | 2 | -1 | 1,73E-01 | NA | 4 | 44 | 0 | 7 | 34 | 12 | 8 | 25 | 2,7 | 0,5 | 2,2 |
| chrX:21903541-21904041:downstream_7482 | chrX | 21857655 | 21903541 | + | MBTPS2 | NM | 14 | 1,7 | 1 | -1 | 1,73E-01 | NA | 10 | 10 | 17 | 10 | 10 | 2 | 24 | 27 | -0,4 | -2,1 | 1,7 |
| chr6:38643201-38643701:downstream_4726 | chr6 | 38643701 | 38670952 | - | GLO1 | NM | 37 | 1,1 | 1 | -1 | 1,73E-01 | 4,99E-01 | 15 | 23 | 6 | 13 | 63 | 48 | 63 | 63 | 1,0 | -0,2 | 1,2 |
| chr19:12809507-12810007:downstream_21567 | chr19 | 12810007 | 12834810 | - | TNPO2 | NM | 11 | 2,0 | 1 | -1 | 1,74E-01 | NA | 11 | 9 | 0 | 2 | 27 | 15 | 10 | 16 | 3,0 | 0,7 | 2,3 |
| chr9:100666270-100666770:downstream_7221 | chr9 | 100666770 | 100684852 | - | C9orf156 | NM | 14 | -2,1 | 2 | 1 | 1,74E-01 | NA | 2 | 0 | 7 | 6 | 11 | 26 | 32 | 31 | -3,0 | -0,8 | -2,2 |
| chr5:150273453-150273953:downstream_24451 | chr5 | 150273953 | 150284545 | - | ZNF300 | NM | 20 | 1,3 | 1 | -1 | 1,74E-01 | NA | 15 | 12 | 11 | 10 | 17 | 22 | 38 | 37 | 0,4 | -0,9 | 1,3 |
| chr11:61556101-61556601:downstream_9379 | chr11 | 61556601 | 61560085 | - | TMEM258 | NM | 28 | 1,3 | 1 | -1 | 1,75E-01 | 5,01E-01 | 15 | 13 | 6 | 7 | 33 | 56 | 35 | 63 | 1,2 | -0,1 | 1,3 |
| chr20:30251760-30252260:downstream_22361 | chr20 | 30252260 | 30310656 | - | BCL2L1 | NM | 28 | 1,3 | 1 | -1 | 1,75E-01 | NA | 14 | 11 | 4 | 9 | 53 | 24 | 50 | 58 | 0,9 | -0,5 | 1,4 |
| chr15:68480404-68480904:downstream_11494 | chr15 | 68346571 | 68480404 | + | PIAS1 | NM | 62 | 1,1 | 1 | -1 | 1,75E-01 | 5,01E-01 | 7 | 11 | 65 | 39 | 19 | 8 | 192 | 157 | -2,6 | -3,7 | 1,1 |
| chr1:112310199-112310699:downstream_903 | chr1 | 112298189 | 112310199 | + | DDX20 | NM | 10 | 1,9 | 1 | -1 | 1,76E-01 | NA | 10 | 8 | 6 | 1 | 11 | 13 | 14 | 16 | 1,5 | -0,3 | 1,8 |
| chrX:128579977-128580477:downstream_27028 | chrX | 128580477 | 128657482 | - | SMARCA1 | NM | 47 | -1,1 | 1 | 1 | 1,76E-01 | 5,01E-01 | 6 | 5 | 17 | 32 | 56 | 44 | 99 | 119 | -2,2 | -1,1 | -1,1 |
| chr22:31023047-31023547:downstream_22936 | chr22 | 31003069 | 31023047 | + | TCN2 | NM | 6 | -2,4 | 2 | 1 | 1,76E-01 | NA | 1 | 5 | 4 | 11 | 8 | 8 | 3 | 6 | -1,5 | 0,9 | -2,4 |
| chr20:49493714-49494214:downstream_22360 | chr20 | 49411430 | 49493714 | + | BCAS4 | NM | 14 | -2,1 | 2 | 1 | 1,77E-01 | NA | 0 | 2 | 7 | 10 | 22 | 7 | 40 | 22 | -3,1 | -1,1 | -2,0 |
| chr22:18074402-18074902:downstream_22750 | chr22 | 18074902 | 18111588 | - | ATP6V1E1 | NM | 117 | 1,1 | 1 | -1 | 1,77E-01 | 5,01E-01 | 6 | 18 | 7 | 2 | 236 | 269 | 218 | 183 | 1,4 | 0,3 | 1,0 |
| chr5:180217040-180217540:downstream_24228 | chr5 | 180217540 | 180237137 | - | MGAT1 | NM | 49 | 1,3 | 1 | -1 | 1,77E-01 | 5,01E-01 | 4 | 8 | 4 | 3 | 76 | 72 | 117 | 107 | 0,7 | -0,6 | 1,3 |
| chr12:110788897-110789397:downstream_18447 | chr12 | 110719031 | 110788897 | + | ATP2A2 | NM | 56 | 1,1 | 1 | -1 | 1,77E-01 | 5,01E-01 | 6 | 5 | 15 | 13 | 26 | 36 | 166 | 185 | -1,4 | -2,5 | 1,1 |
| chr10:17682782-17683282:downstream_8433 | chr10 | 17683282 | 17685917 | - | STAM-AS1 | NR | 3 | 3,8 | 3 | -1 | 1,77E-01 | NA | 11 | 2 | 0 | 0 | 7 | 1 | 3 | 1 | Inf | 1,2 | Inf |
| chr7:45951343-45951843:downstream_25568 | chr7 | 45951843 | 45960871 | - | IGFBP3 | NM | 413 | -1,1 | 1 | 1 | 1,77E-01 | 5,01E-01 | 5 | 1 | 7 | 7 | 750 | 815 | 818 | 898 | -1,3 | -0,1 | -1,2 |
| chr1:212794119-212794619:downstream_16496 | chr1 | 212738675 | 212794119 | + | ATF3 | NM | 42 | -2,4 | 2 | 1 | 1,77E-01 | 5,01E-01 | 12 | 8 | 1 | 0 | 130 | 184 | 3 | 1 | 3,8 | 6,4 | -2,6 |
| chr12:121139667-121140167:downstream_18685 | chr12 | 121124926 | 121139667 | + | MLEC | NM | 131 | 1,0 | 1 | -1 | 1,78E-01 | 5,01E-01 | 20 | 11 | 4 | 11 | 292 | 210 | 221 | 281 | 1,1 | 0,0 | 1,1 |
| chr8:91633722-91634222:downstream_26153 | chr8 | 91634222 | 91658133 | - | TMEM64 | NM | 23 | 1,6 | 1 | -1 | 1,78E-01 | NA | 17 | 6 | 1 | 2 | 54 | 49 | 38 | 20 | 2,6 | 0,9 | 1,7 |
| chr7:26229055-26229555:downstream_25560 | chr7 | 26229555 | 26240413 | - | HNRNPA2B1 | NM | 175 | 0,7 | 0 | -1 | 1,78E-01 | 5,01E-01 | 206 | 163 | 88 | 97 | 255 | 215 | 186 | 189 | 1,0 | 0,3 | 0,7 |
| chr19:6535939-6536439:downstream_13928 | chr19 | 6531009 | 6535939 | + | TNFSF9 | NM | 39 | 1,8 | 1 | -1 | 1,78E-01 | 5,01E-01 | 10 | 9 | 0 | 2 | 78 | 133 | 36 | 42 | 3,5 | 1,4 | 2,1 |
| chr12:112277072-112277572:downstream_10424 | chr12 | 112277572 | 112280706 | - | MAPKAPK5-AS1 | NR | 40 | -1,3 | 1 | 1 | 1,79E-01 | 5,01E-01 | 5 | 2 | 4 | 7 | 90 | 96 | 46 | 71 | -0,7 | 0,7 | -1,3 |
| chr20:57570188-57570688:downstream_15116 | chr20 | 57556262 | 57570188 | + | NELFCD | NM | 12 | -2,0 | 1 | 1 | 1,79E-01 | NA | 1 | 3 | 1 | 8 | 27 | 20 | 14 | 21 | -1,4 | 0,5 | -1,8 |
| chr1:156564091-156564591:downstream_1113 | chr1 | 156561557 | 156564091 | + | APOA1BP | NM | 71 | -1,0 | 1 | 1 | 1,79E-01 | 5,01E-01 | 9 | 4 | 19 | 14 | 130 | 103 | 136 | 151 | -1,4 | -0,3 | -1,1 |
| chr19:38793699-38794199:downstream_21595 | chr19 | 38794199 | 38806606 | - | YIF1B | NM | 8 | 2,4 | 2 | -1 | 1,79E-01 | NA | 11 | 4 | 1 | 9 | 0 | 8 | 15 | 13 | 0,6 | -1,9 | 2,4 |
| chr4:189011925-189012425:downstream_23898 | chr4 | 189012425 | 189026741 | - | TRIML2 | NM | 38 | 1,3 | 1 | -1 | 1,79E-01 | 5,01E-01 | 9 | 7 | 3 | 9 | 59 | 37 | 75 | 109 | 0,4 | -0,9 | 1,4 |
| chr8:37997598-37998098:downstream_25855 | chr8 | 37963010 | 37997598 | + | ASH2L | NM | 5 | -3,0 | 2 | 1 | 1,79E-01 | NA | 2 | 0 | 6 | 6 | 1 | 15 | 3 | 11 | -2,8 | 0,3 | -3,1 |
| chr4:122588651-122589151:downstream_3833 | chr4 | 122589151 | 122618147 | - | ANXA5 | NM | 283 | -1,1 | 1 | 1 | 1,79E-01 | 5,01E-01 | 7 | 0 | 4 | 7 | 569 | 738 | 444 | 495 | -0,7 | 0,5 | -1,1 |
| chr6:82454946-82455446:downstream_5245 | chr6 | 82455446 | 82462428 | - | FAM46A | NM | 72 | -1,1 | 1 | 1 | 1,79E-01 | 5,01E-01 | 11 | 1 | 26 | 13 | 125 | 97 | 161 | 144 | -1,7 | -0,5 | -1,2 |
| chr2:75889334-75889834:downstream_2057 | chr2 | 75873908 | 75889334 | + | MRPL19 | NM | 35 | 1,2 | 1 | -1 | 1,80E-01 | 5,01E-01 | 15 | 15 | 13 | 7 | 35 | 60 | 72 | 59 | 0,6 | -0,5 | 1,1 |
| chr1:167509750-167510250:downstream_1202 | chr1 | 167510250 | 167523056 | - | CREG1 | NM | 333 | 1,4 | 1 | -1 | 1,80E-01 | 5,01E-01 | 8 | 7 | 1 | 2 | 803 | 924 | 458 | 459 | 2,3 | 0,9 | 1,4 |
| chr9:111704348-111704848:downstream_26271 | chr9 | 111704848 | 111775874 | - | CTNNAL1 | NM | 9 | 2,2 | 2 | -1 | 1,80E-01 | NA | 5 | 5 | 4 | 0 | 7 | 15 | 18 | 16 | 1,2 | -0,7 | 1,9 |
| chr19:52208443-52208943:downstream_14541 | chr19 | 52196592 | 52208443 | + | SPACA6P | NR | 12 | 1,6 | 1 | -1 | 1,81E-01 | NA | 19 | 27 | 8 | 8 | 9 | 7 | 8 | 9 | 1,5 | -0,1 | 1,6 |
| chr1:156706752-156707252:downstream_17200 | chr1 | 156698262 | 156706752 | + | RRNAD1 | NM | 6 | 2,3 | 2 | -1 | 1,81E-01 | NA | 3 | 6 | 4 | 6 | 4 | 1 | 8 | 19 | -0,1 | -2,4 | 2,3 |
| chr17:79845210-79845710:downstream_13440 | chr17 | 79845710 | 79849462 | - | ALYREF | NM | 44 | 1,1 | 1 | -1 | 1,81E-01 | 5,03E-01 | 19 | 34 | 10 | 16 | 52 | 79 | 60 | 81 | 1,0 | -0,1 | 1,1 |
| chr9:117361152-117361652:downstream_7277 | chr9 | 117349993 | 117361152 | + | ATP6V1G1 | NM | 16 | 1,6 | 1 | -1 | 1,81E-01 | NA | 15 | 22 | 1 | 5 | 22 | 30 | 19 | 10 | 2,6 | 0,8 | 1,8 |
| chr5:52249485-52249985:downstream_4015 | chr5 | 52084135 | 52249485 | + | ITGA1 | NM | 6 | -2,3 | 2 | 1 | 1,81E-01 | NA | 4 | 0 | 10 | 5 | 12 | 8 | 4 | 9 | -1,9 | 0,6 | -2,4 |
| chr22:25746884-25747384:downstream_22837 | chr22 | 25747384 | 25777544 | - | LRP5L | NM | 7 | -2,7 | 2 | 1 | 1,81E-01 | NA | 0 | 1 | 4 | 7 | 6 | 9 | 18 | 11 | -3,6 | -1,0 | -2,6 |
| chr10:91450556-91451056:downstream_8633 | chr10 | 91451056 | 91457685 | - | FLJ37201 | NR | 4 | 3,2 | 2 | -1 | 1,82E-01 | NA | 2 | 7 | 3 | 9 | 1 | 0 | 7 | 5 | -0,5 | -3,9 | 3,4 |
| chr17:18231370-18231870:downstream_12613 | chr17 | 18218593 | 18231370 | + | SMCR8 | NM | 18 | 1,7 | 1 | -1 | 1,82E-01 | NA | 3 | 12 | 3 | 2 | 24 | 41 | 38 | 25 | 1,7 | 0,1 | 1,7 |
| chrX:63404496-63404996:downstream_7764 | chrX | 63404996 | 63425624 | - | AMER1 | NM | 4 | 3,4 | 3 | -1 | 1,82E-01 | NA | 7 | 2 | 0 | 0 | 10 | 8 | 0 | 7 | Inf | 1,2 | Inf |
| chr6:74211179-74211679:downstream_25028 | chr6 | 74171453 | 74211179 | + | MTO1 | NM | 9 | 2,1 | 2 | -1 | 1,82E-01 | NA | 7 | 16 | 1 | 2 | 17 | 10 | 10 | 6 | 3,0 | 0,8 | 2,2 |
| chr12:56989980-56990480:downstream_10258 | chr12 | 56915608 | 56989980 | + | RBMS2 | NM | 7 | -2,8 | 2 | 1 | 1,82E-01 | NA | 0 | 1 | 13 | 11 | 1 | 6 | 11 | 12 | -4,6 | -1,8 | -2,8 |
| chr4:129440551-129441051:downstream_3505 | chr4 | 129349170 | 129440551 | + | LOC100507487 | NR | 8 | -2,8 | 2 | 1 | 1,83E-01 | NA | 0 | 0 | 3 | 7 | 7 | 3 | 25 | 21 | -Inf | -2,2 | -Inf |
| chr6:147891157-147891657:downstream_4675 | chr6 | 147829827 | 147891157 | + | SAMD5 | NM | 7 | -3,1 | 2 | 1 | 1,83E-01 | NA | 0 | 0 | 4 | 8 | 7 | 0 | 13 | 22 | -Inf | -2,2 | -Inf |
| chr2:37476303-37476803:downstream_22107 | chr2 | 37458773 | 37476303 | + | NDUFAF7 | NM | 23 | 1,3 | 1 | -1 | 1,83E-01 | NA | 47 | 19 | 24 | 12 | 23 | 12 | 22 | 24 | 0,9 | -0,4 | 1,3 |
| chr8:42691316-42691816:downstream_26150 | chr8 | 42691816 | 42698474 | - | THAP1 | NM | 21 | 1,8 | 1 | -1 | 1,83E-01 | NA | 6 | 8 | 1 | 2 | 60 | 22 | 42 | 25 | 2,1 | 0,3 | 1,8 |
| chr18:5288517-5289017:downstream_20973 | chr18 | 5289017 | 5297052 | - | ZBTB14 | NM | 5 | 3,2 | 2 | -1 | 1,83E-01 | NA | 3 | 7 | 7 | 10 | 1 | 0 | 0 | 13 | -0,7 | -4,0 | 3,3 |
| chr17:46973232-46973732:downstream_20346 | chr17 | 46970147 | 46973232 | + | ATP5G1 | NM | 22 | 1,4 | 1 | -1 | 1,84E-01 | NA | 15 | 27 | 8 | 2 | 47 | 28 | 24 | 25 | 1,9 | 0,6 | 1,3 |
| chr15:42191138-42191638:downstream_11457 | chr15 | 42191638 | 42264755 | - | EHD4 | NM | 11 | -1,7 | 1 | 1 | 1,84E-01 | NA | 8 | 3 | 19 | 18 | 9 | 11 | 13 | 8 | -1,8 | -0,1 | -1,7 |
| chrX:150577836-150578336:downstream_8015 | chrX | 150565656 | 150577836 | + | VMA21 | NM | 44 | 1,5 | 1 | -1 | 1,84E-01 | 5,10E-01 | 8 | 3 | 1 | 2 | 84 | 104 | 77 | 73 | 1,9 | 0,3 | 1,5 |
| chr8:97251125-97251625:downstream_26018 | chr8 | 97251625 | 97273841 | - | MTERF3 | NM | 12 | 2,1 | 2 | -1 | 1,84E-01 | NA | 5 | 4 | 1 | 1 | 19 | 24 | 19 | 23 | 2,0 | 0,0 | 2,0 |
| chr12:56507694-56508194:downstream_10251 | chr12 | 56498102 | 56507694 | + | PA2G4 | NM | 92 | 0,9 | 1 | -1 | 1,85E-01 | 5,10E-01 | 46 | 53 | 8 | 10 | 206 | 256 | 83 | 75 | 2,5 | 1,5 | 0,9 |
| chr2:99785225-99785725:downstream_2138 | chr2 | 99785725 | 99797492 | - | MITD1 | NM | 9 | 2,0 | 2 | -1 | 1,85E-01 | NA | 6 | 5 | 1 | 2 | 10 | 15 | 8 | 29 | 1,5 | -0,6 | 2,1 |
| chr10:46110448-46110948:downstream_17826 | chr10 | 46110948 | 46168261 | - | ZFAND4 | NM | 7 | 2,7 | 2 | -1 | 1,85E-01 | NA | 4 | 9 | 19 | 9 | 1 | 0 | 6 | 7 | -1,2 | -4,0 | 2,8 |
| chr1:186958113-186958613:downstream_263 | chr1 | 186798031 | 186958113 | + | PLA2G4A | NM | 125 | 1,0 | 1 | -1 | 1,85E-01 | 5,10E-01 | 2 | 10 | 8 | 9 | 132 | 106 | 404 | 326 | -0,5 | -1,6 | 1,1 |
| chr12:49667121-49667621:downstream_18896 | chr12 | 49621708 | 49667121 | + | TUBA1C | NM | 372 | 1,2 | 1 | -1 | 1,85E-01 | 5,10E-01 | 9 | 7 | 6 | 1 | 740 | 954 | 646 | 615 | 1,3 | 0,4 | 0,9 |
| chr2:136596695-136597195:downstream_2265 | chr2 | 136597195 | 136634047 | - | MCM6 | NM | 4 | 3,2 | 2 | -1 | 1,85E-01 | NA | 7 | 3 | 6 | 0 | 2 | 0 | 10 | 2 | 0,9 | -2,3 | 3,2 |
| chr4:139978370-139978870:downstream_23625 | chr4 | 139978870 | 140060630 | - | ELF2 | NM | 13 | 1,5 | 1 | -1 | 1,85E-01 | NA | 12 | 15 | 13 | 16 | 5 | 8 | 18 | 20 | -0,1 | -1,6 | 1,5 |
| chr2:202037411-202037911:downstream_21823 | chr2 | 201980826 | 202037411 | + | CFLAR | NM | 7 | -2,4 | 2 | 1 | 1,85E-01 | NA | 2 | 0 | 7 | 4 | 8 | 11 | 10 | 12 | -2,8 | -0,2 | -2,8 |
| chr6:72086162-72086662:downstream_5220 | chr6 | 72086662 | 72086734 | - | MIR30C2 | NR | 31 | 1,2 | 1 | -1 | 1,86E-01 | 5,10E-01 | 7 | 15 | 53 | 25 | 7 | 10 | 64 | 65 | -1,8 | -3,0 | 1,2 |
| chr1:231154203-231154703:downstream_1417 | chr1 | 231154703 | 231175995 | - | FAM89A | NM | 9 | 1,9 | 1 | -1 | 1,86E-01 | NA | 2 | 10 | 6 | 3 | 8 | 6 | 21 | 17 | 0,5 | -1,5 | 1,9 |
| chr10:62554610-62555110:downstream_17517 | chr10 | 62538088 | 62554610 | + | CDK1 | NM | 75 | 0,9 | 1 | -1 | 1,86E-01 | 5,10E-01 | 27 | 20 | 14 | 34 | 94 | 76 | 157 | 177 | 0,0 | -1,0 | 1,0 |
| chr19:42375484-42375984:downstream_14324 | chr19 | 42363987 | 42375484 | + | RPS19 | NM | 166 | 0,7 | 1 | -1 | 1,86E-01 | 5,10E-01 | 212 | 162 | 65 | 45 | 273 | 299 | 131 | 139 | 1,8 | 1,1 | 0,7 |
| chr3:39182841-39183341:downstream_2606 | chr3 | 39183341 | 39195102 | - | CSRNP1 | NM | 19 | 1,6 | 1 | -1 | 1,86E-01 | NA | 36 | 15 | 3 | 2 | 36 | 37 | 10 | 12 | 3,3 | 1,7 | 1,5 |
| chr13:48984681-48985181:downstream_19021 | chr13 | 48985181 | 49018840 | - | LPAR6 | NM | 15 | 1,6 | 1 | -1 | 1,86E-01 | NA | 15 | 20 | 4 | 2 | 29 | 20 | 8 | 18 | 2,4 | 0,9 | 1,5 |
| chr5:100142138-100142638:downstream_24397 | chr5 | 100142638 | 100238989 | - | ST8SIA4 | NM | 6 | 2,8 | 2 | -1 | 1,87E-01 | NA | 2 | 3 | 15 | 6 | 1 | 0 | 8 | 16 | -2,0 | -4,9 | 2,9 |
| chr1:201939789-201940289:downstream_1292 | chr1 | 201924618 | 201939789 | + | TIMM17A | NM | 17 | 1,6 | 1 | -1 | 1,87E-01 | NA | 8 | 14 | 4 | 1 | 33 | 32 | 21 | 21 | 2,1 | 0,7 | 1,5 |
| chr6:28057340-28057840:downstream_4705 | chr6 | 28048481 | 28057340 | + | ZNF165 | NM | 7 | 2,2 | 2 | -1 | 1,87E-01 | NA | 3 | 9 | 0 | 3 | 9 | 8 | 8 | 16 | 1,9 | -0,6 | 2,4 |
| chr3:136574734-136575234:downstream_23431 | chr3 | 136537860 | 136574734 | + | SLC35G2 | NM | 11 | 2,0 | 1 | -1 | 1,87E-01 | NA | 6 | 6 | 4 | 1 | 20 | 8 | 14 | 27 | 1,2 | -0,6 | 1,8 |
| chr10:124689918-124690418:downstream_8763 | chr10 | 124690418 | 124713919 | - | C10orf88 | NM | 8 | 2,1 | 2 | -1 | 1,87E-01 | NA | 7 | 6 | 6 | 0 | 9 | 8 | 17 | 12 | 1,1 | -0,8 | 1,9 |
| chr3:93845630-93846130:downstream_3057 | chr3 | 93781854 | 93845630 | + | NSUN3 | NM | 25 | 1,3 | 1 | -1 | 1,87E-01 | NA | 13 | 32 | 38 | 14 | 16 | 12 | 46 | 33 | -0,2 | -1,5 | 1,3 |
| chr6:109687216-109687716:downstream_24554 | chr6 | 109687716 | 109703762 | - | CD164 | NM | 177 | 1,2 | 1 | -1 | 1,87E-01 | 5,13E-01 | 11 | 4 | 3 | 2 | 323 | 468 | 263 | 343 | 1,5 | 0,4 | 1,2 |
| chr8:67954814-67955314:downstream_6590 | chr8 | 67955314 | 67974562 | - | COPS5 | NM | 18 | 1,6 | 1 | -1 | 1,87E-01 | NA | 7 | 16 | 1 | 3 | 29 | 40 | 24 | 21 | 2,3 | 0,6 | 1,7 |
| chr17:42269099-42269599:downstream_20782 | chr17 | 42264353 | 42269099 | + | TMUB2 | NM | 15 | 1,5 | 1 | -1 | 1,88E-01 | NA | 7 | 11 | 6 | 10 | 17 | 7 | 33 | 34 | 0,1 | -1,5 | 1,6 |
| chr16:75611779-75612279:downstream_12478 | chr16 | 75600248 | 75611779 | + | GABARAPL2 | NM | 68 | 1,1 | 1 | -1 | 1,88E-01 | 5,15E-01 | 18 | 13 | 8 | 2 | 141 | 162 | 93 | 108 | 1,5 | 0,6 | 0,9 |
| chr16:70407281-70407781:downstream_12021 | chr16 | 70380823 | 70407281 | + | DDX19A | NM | 17 | 1,6 | 1 | -1 | 1,89E-01 | NA | 11 | 7 | 4 | 2 | 23 | 33 | 21 | 38 | 1,4 | -0,1 | 1,5 |
| chr2:55463255-55463755:downstream_22091 | chr2 | 55463755 | 55496384 | - | MTIF2 | NM | 13 | 1,5 | 1 | -1 | 1,89E-01 | NA | 8 | 15 | 14 | 17 | 6 | 5 | 25 | 16 | -0,4 | -2,0 | 1,6 |
| chr6:99847340-99847840:downstream_25067 | chr6 | 99847840 | 99873207 | - | PNISR | NM | 22 | 1,3 | 1 | -1 | 1,89E-01 | NA | 20 | 16 | 38 | 25 | 8 | 7 | 17 | 46 | -0,8 | -2,1 | 1,3 |
| chr14:53502957-53503457:downstream_19200 | chr14 | 53503457 | 53620046 | - | DDHD1 | NM | 5 | -2,8 | 2 | 1 | 1,89E-01 | NA | 0 | 2 | 4 | 11 | 2 | 8 | 4 | 7 | -2,9 | -0,2 | -2,7 |
| chr2:70508317-70508817:downstream_2016 | chr2 | 70485230 | 70508317 | + | PCYOX1 | NM | 60 | 1,2 | 1 | -1 | 1,90E-01 | 5,15E-01 | 5 | 17 | 6 | 4 | 103 | 124 | 97 | 122 | 1,2 | 0,0 | 1,2 |
| chr6:10709970-10710470:downstream_4864 | chr6 | 10695187 | 10709970 | + | PAK1IP1 | NM | 43 | 1,3 | 1 | -1 | 1,90E-01 | 5,15E-01 | 12 | 16 | 1 | 5 | 131 | 66 | 56 | 60 | 2,2 | 0,8 | 1,4 |
| chr22:24237409-24237909:downstream_15517 | chr22 | 24236564 | 24237409 | + | MIF | NM | 107 | 1,1 | 1 | -1 | 1,90E-01 | 5,15E-01 | 11 | 13 | 8 | 0 | 250 | 257 | 145 | 176 | 1,5 | 0,7 | 0,8 |
| chr4:76735442-76735942:downstream_23912 | chr4 | 76649705 | 76735442 | + | USO1 | NM | 67 | -1,7 | 1 | 1 | 1,90E-01 | 5,15E-01 | 2 | 0 | 17 | 4 | 46 | 73 | 213 | 186 | -3,7 | -1,8 | -1,9 |
| chr2:32531658-32532158:downstream_1514 | chr2 | 32502957 | 32531658 | + | YIPF4 | NM | 154 | -0,6 | 0 | 1 | 1,90E-01 | 5,15E-01 | 62 | 62 | 141 | 131 | 177 | 172 | 257 | 232 | -1,1 | -0,5 | -0,6 |
| chr3:152555843-152556343:downstream_3242 | chr3 | 152552735 | 152555843 | + | P2RY1 | NM | 9 | 1,9 | 1 | -1 | 1,91E-01 | NA | 7 | 9 | 4 | 2 | 10 | 13 | 18 | 11 | 1,4 | -0,4 | 1,8 |
| chr14:39722575-39723075:downstream_10934 | chr14 | 39703124 | 39722575 | + | MIA2 | NM | 8 | 2,3 | 2 | -1 | 1,91E-01 | NA | 7 | 4 | 3 | 3 | 13 | 0 | 24 | 14 | 0,8 | -1,5 | 2,3 |
| chr17:61908940-61909440:downstream_13302 | chr17 | 61909440 | 61920351 | - | SMARCD2 | NM | 21 | -1,5 | 1 | 1 | 1,91E-01 | NA | 3 | 1 | 7 | 7 | 31 | 43 | 39 | 40 | -1,7 | -0,1 | -1,6 |
| chr2:170667766-170668266:downstream_22078 | chr2 | 170668266 | 170681441 | - | METTL5 | NM | 10 | 1,8 | 1 | -1 | 1,91E-01 | NA | 9 | 18 | 7 | 7 | 5 | 8 | 19 | 19 | 0,9 | -0,9 | 1,8 |
| chr1:152004481-152004981:downstream_1007 | chr1 | 152004981 | 152009511 | - | S100A11 | NM | 221 | 0,9 | 1 | -1 | 1,92E-01 | 5,18E-01 | 112 | 87 | 47 | 14 | 554 | 423 | 335 | 194 | 1,7 | 0,9 | 0,8 |
| chr2:85621370-85621870:downstream_21789 | chr2 | 85621870 | 85641197 | - | CAPG | NM | 14 | 1,9 | 1 | -1 | 1,92E-01 | NA | 1 | 12 | 1 | 2 | 19 | 23 | 18 | 34 | 1,7 | -0,3 | 2,0 |
| chr3:63997917-63998417:downstream_3028 | chr3 | 63989697 | 63997917 | + | PSMD6-AS2 | NR | 4 | -2,6 | 2 | 1 | 1,92E-01 | NA | 2 | 1 | 6 | 8 | 4 | 5 | 6 | 2 | -2,4 | 0,1 | -2,6 |
| chr20:3028896-3029396:downstream_14862 | chr20 | 3026674 | 3028896 | + | MRPS26 | NM | 25 | 1,6 | 1 | -1 | 1,92E-01 | NA | 2 | 13 | 1 | 2 | 48 | 53 | 36 | 48 | 2,0 | 0,3 | 1,7 |
| chr2:192290115-192290615:downstream_22097 | chr2 | 192110106 | 192290115 | + | MYO1B | NM | 24 | -2,2 | 2 | 1 | 1,92E-01 | NA | 0 | 1 | 8 | 16 | 12 | 11 | 56 | 84 | -4,7 | -2,6 | -2,1 |
| chr17:53854748-53855248:downstream_20624 | chr17 | 53828339 | 53854748 | + | PCTP | NM | 4 | 3,7 | 3 | -1 | 1,93E-01 | NA | 2 | 0 | 0 | 11 | 0 | 0 | 11 | 7 | -2,1 | -Inf | Inf |
| chr1:36883006-36883506:downstream_17052 | chr1 | 36883506 | 36916086 | - | OSCP1 | NM | 6 | 2,3 | 2 | -1 | 1,93E-01 | NA | 2 | 10 | 3 | 2 | 7 | 6 | 11 | 11 | 1,4 | -0,9 | 2,2 |
| chrX:24096927-24097427:downstream_7634 | chrX | 24073064 | 24096927 | + | EIF2S3 | NM | 107 | 0,9 | 1 | -1 | 1,93E-01 | 5,20E-01 | 15 | 8 | 6 | 11 | 184 | 156 | 239 | 234 | 0,5 | -0,5 | 1,0 |
| chr1:173833079-173833579:downstream_1234 | chr1 | 173832385 | 173833079 | + | GAS5-AS1 | NR | 13 | -1,7 | 1 | 1 | 1,93E-01 | NA | 7 | 0 | 13 | 11 | 15 | 23 | 17 | 19 | -1,7 | 0,1 | -1,8 |
| chr15:66184329-66184829:downstream_19689 | chr15 | 66161796 | 66184329 | + | RAB11A | NM | 61 | 1,2 | 1 | -1 | 1,93E-01 | 5,20E-01 | 33 | 8 | 4 | 19 | 74 | 107 | 128 | 116 | 0,8 | -0,4 | 1,2 |
| chr10:27475848-27476348:downstream_17645 | chr10 | 27443752 | 27475848 | + | MASTL | NM | 24 | 1,4 | 1 | -1 | 1,93E-01 | NA | 3 | 5 | 8 | 4 | 15 | 19 | 68 | 66 | -0,6 | -2,0 | 1,3 |
| chr18:67670542-67671042:downstream_13528 | chr18 | 67671042 | 67872962 | - | RTTN | NM | 5 | -2,9 | 2 | 1 | 1,94E-01 | NA | 1 | 0 | 7 | 16 | 2 | 3 | 6 | 10 | -4,8 | -1,8 | -3,0 |
| chr3:48954720-48955220:downstream_2946 | chr3 | 48955220 | 48956818 | - | ARIH2OS | NM | 10 | -1,8 | 1 | 1 | 1,94E-01 | NA | 9 | 4 | 6 | 5 | 16 | 28 | 4 | 6 | 0,3 | 2,2 | -1,9 |
| chr19:44637255-44637755:downstream_14361 | chr19 | 44617547 | 44637255 | + | ZNF225 | NM | 11 | -1,6 | 1 | 1 | 1,94E-01 | NA | 8 | 6 | 14 | 12 | 11 | 18 | 11 | 7 | -0,9 | 0,7 | -1,6 |
| chr2:128603339-128603839:downstream_2229 | chr2 | 128603839 | 128615729 | - | POLR2D | NM | 40 | -0,9 | 1 | 1 | 1,94E-01 | 5,20E-01 | 31 | 34 | 25 | 25 | 73 | 75 | 28 | 31 | 0,4 | 1,3 | -0,9 |
| chr6:116595521-116596021:downstream_5314 | chr6 | 116596021 | 116601280 | - | TSPYL1 | NM | 6 | -2,7 | 2 | 1 | 1,94E-01 | NA | 11 | 0 | 4 | 5 | 6 | 17 | 1 | 2 | 0,2 | 2,9 | -2,7 |
| chr12:56864767-56865267:downstream_10256 | chr12 | 56862300 | 56864767 | + | SPRYD4 | NM | 24 | 1,3 | 1 | -1 | 1,94E-01 | NA | 38 | 50 | 4 | 7 | 32 | 41 | 11 | 12 | 2,9 | 1,6 | 1,3 |
| chrX:48829630-48830130:downstream_7714 | chrX | 48830130 | 48858675 | - | GRIPAP1 | NM | 11 | 1,7 | 1 | -1 | 1,94E-01 | NA | 7 | 13 | 1 | 4 | 15 | 15 | 14 | 17 | 1,8 | -0,1 | 1,9 |
| chr18:673499-673999:downstream_13539 | chr18 | 657603 | 673499 | + | TYMS | NM | 31 | 1,4 | 1 | -1 | 1,94E-01 | 5,20E-01 | 7 | 19 | 0 | 5 | 74 | 62 | 40 | 42 | 2,4 | 0,7 | 1,7 |
| chr10:13180276-13180776:downstream_17688 | chr10 | 13142081 | 13180276 | + | OPTN | NM | 6 | 2,1 | 2 | -1 | 1,94E-01 | NA | 4 | 6 | 3 | 3 | 3 | 6 | 11 | 13 | 0,7 | -1,4 | 2,1 |
| chr17:20218072-20218572:downstream_20739 | chr17 | 19912648 | 20218072 | + | SPECC1 | NM | 7 | -2,7 | 2 | 1 | 1,95E-01 | NA | 1 | 0 | 7 | 4 | 4 | 11 | 14 | 14 | -3,8 | -0,9 | -2,8 |
| chr17:25640645-25641145:downstream_20811 | chr17 | 25621105 | 25640645 | + | WSB1 | NM | 70 | 0,9 | 1 | -1 | 1,95E-01 | 5,20E-01 | 21 | 20 | 26 | 23 | 51 | 96 | 164 | 158 | -0,3 | -1,1 | 0,9 |
| chr3:14714166-14714666:downstream_2836 | chr3 | 14693252 | 14714166 | + | CCDC174 | NM | 32 | 1,1 | 1 | -1 | 1,95E-01 | 5,20E-01 | 64 | 47 | 17 | 24 | 31 | 26 | 22 | 21 | 1,5 | 0,4 | 1,1 |
| chr20:50212813-50213313:downstream_14738 | chr20 | 50213313 | 50384950 | - | ATP9A | NM | 34 | -1,7 | 1 | 1 | 1,95E-01 | 5,20E-01 | 1 | 1 | 17 | 10 | 24 | 20 | 102 | 98 | -3,9 | -2,2 | -1,7 |
| chr12:74935232-74935732:downstream_10323 | chr12 | 74931550 | 74935232 | + | ATXN7L3B | NM | 38 | 1,1 | 1 | -1 | 1,95E-01 | 5,20E-01 | 49 | 41 | 28 | 11 | 54 | 37 | 46 | 35 | 1,2 | 0,2 | 1,0 |
| chr1:246703362-246703862:downstream_1457 | chr1 | 246703862 | 246729565 | - | TFB2M | NM | 11 | 2,1 | 2 | -1 | 1,95E-01 | NA | 2 | 12 | 3 | 0 | 21 | 20 | 14 | 16 | 2,3 | 0,5 | 1,8 |
| chr8:133861052-133861552:downstream_26049 | chr8 | 133787603 | 133861052 | + | PHF20L1 | NM | 10 | 1,9 | 1 | -1 | 1,95E-01 | NA | 8 | 7 | 1 | 7 | 15 | 3 | 19 | 21 | 0,8 | -1,2 | 2,0 |
| chr12:110478481-110478981:downstream_9924 | chr12 | 110478981 | 110505500 | - | C12orf76 | NM | 9 | 2,7 | 2 | -1 | 1,95E-01 | NA | 4 | 12 | 0 | 0 | 17 | 29 | 4 | 3 | Inf | 2,6 | Inf |
| chr7:111983989-111984489:downstream_5608 | chr7 | 111846642 | 111983989 | + | ZNF277 | NM | 17 | 1,6 | 1 | -1 | 1,95E-01 | NA | 7 | 15 | 46 | 16 | 4 | 2 | 22 | 26 | -1,4 | -3,0 | 1,6 |
| chr15:63445038-63445538:downstream_11749 | chr15 | 63445538 | 63449741 | - | RPS27L | NM | 171 | 0,7 | 1 | -1 | 1,96E-01 | 5,20E-01 | 472 | 425 | 74 | 65 | 116 | 150 | 33 | 33 | 2,7 | 2,0 | 0,7 |
| chr11:62573831-62574331:downstream_18322 | chr11 | 62574331 | 62599563 | - | STX5 | NM | 15 | 1,5 | 1 | -1 | 1,96E-01 | NA | 37 | 22 | 11 | 7 | 11 | 15 | 6 | 15 | 1,8 | 0,4 | 1,4 |
| chr1:222730743-222731243:downstream_17306 | chr1 | 222731243 | 222763275 | - | TAF1A | NM | 4 | -2,5 | 2 | 1 | 1,96E-01 | NA | 4 | 1 | 8 | 7 | 5 | 5 | 1 | 3 | -1,6 | 1,1 | -2,6 |
| chr12:65106721-65107221:downstream_9875 | chr12 | 65107221 | 65153226 | - | GNS | NM | 24 | 1,6 | 1 | -1 | 1,96E-01 | NA | 6 | 5 | 1 | 2 | 49 | 45 | 40 | 43 | 1,8 | 0,2 | 1,6 |
| chr3:179342288-179342788:downstream_23303 | chr3 | 179322574 | 179342288 | + | NDUFB5 | NM | 84 | 1,0 | 1 | -1 | 1,96E-01 | 5,21E-01 | 15 | 7 | 6 | 4 | 159 | 182 | 157 | 147 | 1,1 | 0,2 | 1,0 |
| chrX:51232362-51232862:downstream_7730 | chrX | 51232862 | 51239459 | - | NUDT11 | NM | 4 | 3,4 | 3 | -1 | 1,97E-01 | NA | 7 | 10 | 0 | 0 | 5 | 5 | 3 | 0 | Inf | 1,8 | Inf |
| chr17:1420782-1421282:downstream_12584 | chr17 | 1421282 | 1466110 | - | PITPNA | NM | 7 | 2,3 | 2 | -1 | 1,97E-01 | NA | 3 | 8 | 0 | 2 | 5 | 15 | 11 | 13 | 2,2 | -0,3 | 2,5 |
| chr3:12194067-12194567:downstream_2827 | chr3 | 12194567 | 12200851 | - | TIMP4 | NM | 38 | -1,2 | 1 | 1 | 1,97E-01 | 5,22E-01 | 20 | 8 | 8 | 16 | 67 | 110 | 29 | 42 | 0,2 | 1,3 | -1,1 |
| chr19:58052703-58053203:downstream_21666 | chr19 | 58053203 | 58071231 | - | ZNF550 | NM | 9 | 2,2 | 2 | -1 | 1,98E-01 | NA | 8 | 3 | 3 | 0 | 13 | 17 | 17 | 11 | 2,0 | 0,1 | 1,9 |
| chr5:126172712-126173212:downstream_24198 | chr5 | 126112314 | 126172712 | + | LMNB1 | NM | 42 | -2,1 | 2 | 1 | 1,98E-01 | 5,22E-01 | 1 | 0 | 11 | 6 | 18 | 44 | 136 | 122 | -4,4 | -2,1 | -2,3 |
| chr20:18542059-18542559:downstream_22545 | chr20 | 18488187 | 18542059 | + | SEC23B | NM | 15 | 1,5 | 1 | -1 | 1,98E-01 | NA | 7 | 13 | 6 | 3 | 21 | 18 | 18 | 30 | 1,2 | -0,3 | 1,5 |
| chr22:21981877-21982377:downstream_15498 | chr22 | 21982377 | 21984340 | - | YDJC | NM | 33 | -1,2 | 1 | 1 | 1,98E-01 | 5,22E-01 | 12 | 3 | 10 | 10 | 62 | 87 | 42 | 40 | -0,4 | 0,9 | -1,2 |
| chr16:67905219-67905719:downstream_12013 | chr16 | 67880818 | 67905219 | + | NUTF2 | NM | 99 | 0,9 | 1 | -1 | 1,98E-01 | 5,22E-01 | 18 | 35 | 6 | 11 | 231 | 216 | 113 | 164 | 1,7 | 0,7 | 1,0 |
| chr15:72879654-72880154:downstream_11793 | chr15 | 72879557 | 72879654 | + | MIR630 | NR | 40 | -1,1 | 1 | 1 | 1,98E-01 | 5,22E-01 | 32 | 17 | 88 | 92 | 24 | 11 | 22 | 37 | -1,9 | -0,8 | -1,1 |
| chr1:233119381-233119881:downstream_310 | chr1 | 233119881 | 233431459 | - | PCNXL2 | NM | 9 | 1,9 | 1 | -1 | 1,99E-01 | NA | 6 | 15 | 17 | 6 | 2 | 4 | 15 | 10 | -0,2 | -2,0 | 1,8 |

| | | | | | | | | | | | | | | | | | | | | | | | |
|---|---|---|---|---|---|---|---|---|---|---|---|---|---|---|---|---|---|---|---|---|---|---|---|
| chr11:8711419-8711919:downstream_9131 | chr11 | 8703994 | 8711419 | + | RPL27A | NM | 56 | -1,0 | 1 | 1 | 1,99E-01 | 5,22E-01 | 105 | 138 | 29 | 30 | 83 | 44 | 7 | 8 | 2,0 | 3,1 | -1,0 |
| chr1:28904549-28905049:downstream_592 | chr1 | 28905049 | 28908366 | - | SNHG12 | NR | 50 | 0,9 | 1 | -1 | 1,99E-01 | 5,22E-01 | 44 | 51 | 13 | 11 | 80 | 109 | 40 | 50 | 2,0 | 1,1 | 0,9 |
| chr5:148208197-148208697:downstream_4511 | chr5 | 148206155 | 148208197 | + | ADRB2 | NM | 21 | 2,8 | 2 | -1 | 2,00E-01 | NA | 67 | 51 | 0 | 0 | 25 | 21 | 1 | 0 | Inf | 5,1 | Inf |
| chr2:20231910-20232410:downstream_1879 | chr2 | 20232410 | 20251789 | - | LAPTM4A | NM | 356 | -1,2 | 1 | 1 | 2,00E-01 | 5,23E-01 | 1 | 4 | 7 | 4 | 652 | 735 | 686 | 756 | -1,2 | -0,1 | -1,2 |
| chr18:39661446-39661946:downstream_20912 | chr18 | 39535162 | 39661446 | + | PIK3C3 | NM | 10 | 2,4 | 2 | -1 | 2,01E-01 | NA | 1 | 4 | 11 | 13 | 2 | 0 | 42 | 7 | -2,4 | -4,9 | 2,5 |
| chr11:47260352-47260852:downstream_17846 | chr11 | 47260852 | 47270398 | - | ACP2 | NM | 21 | 1,3 | 1 | -1 | 2,01E-01 | NA | 14 | 9 | 3 | 6 | 31 | 38 | 38 | 31 | 1,4 | 0,0 | 1,4 |
| chr1:46743571-46744071:downstream_710 | chr1 | 46744071 | 46769038 | - | LRRC41 | NM | 9 | -1,9 | 1 | 1 | 2,01E-01 | NA | 2 | 1 | 7 | 8 | 16 | 9 | 14 | 16 | -2,2 | -0,2 | -1,9 |
| chr5:162880085-162880585:downstream_4558 | chr5 | 162880585 | 162887143 | - | NUDCD2 | NM | 40 | 1,1 | 1 | -1 | 2,01E-01 | 5,26E-01 | 5 | 13 | 7 | 7 | 53 | 49 | 96 | 91 | 0,3 | -0,9 | 1,2 |
| chr6:79610683-79611183:downstream_5238 | chr6 | 79577260 | 79610683 | + | IRAK1BP1 | NM | 5 | -2,6 | 2 | 1 | 2,02E-01 | NA | 2 | 0 | 8 | 7 | 2 | 8 | 8 | 6 | -3,3 | -0,6 | -2,7 |
| chr10:38412278-38412778:downstream_17836 | chr10 | 38383263 | 38412278 | + | ZNF37A | NM | 32 | 1,2 | 1 | -1 | 2,02E-01 | 5,27E-01 | 32 | 15 | 14 | 7 | 49 | 46 | 67 | 28 | 1,2 | 0,0 | 1,2 |
| chr10:54210136-54210636:downstream_17633 | chr10 | 54210636 | 54230293 | - | LINC01468 | NR | 30 | -1,2 | 1 | 1 | 2,02E-01 | 5,27E-01 | 4 | 4 | 18 | 25 | 31 | 24 | 64 | 71 | -2,5 | -1,3 | -1,2 |
| chr9:35088184-35088684:downstream_26444 | chr9 | 35088684 | 35096598 | - | PIGO | NM | 30 | 1,1 | 1 | -1 | 2,03E-01 | 5,27E-01 | 12 | 13 | 7 | 8 | 50 | 34 | 53 | 61 | 0,7 | -0,4 | 1,2 |
| chr4:90036052-90036552:downstream_3783 | chr4 | 90033967 | 90036052 | + | TIGD2 | NM | 24 | 1,3 | 1 | -1 | 2,03E-01 | NA | 41 | 35 | 36 | 23 | 17 | 4 | 18 | 21 | 0,4 | -0,9 | 1,3 |
| chr2:200828847-200829347:downstream_2371 | chr2 | 200820039 | 200828847 | + | C2orf47 | NM | 12 | 1,7 | 1 | -1 | 2,03E-01 | NA | 10 | 10 | 3 | 2 | 23 | 15 | 11 | 21 | 1,9 | 0,2 | 1,7 |
| chr12:6980110-6980610:downstream_18888 | chr12 | 6976583 | 6980110 | + | TPI1 | NM | 785 | 0,7 | 1 | -1 | 2,03E-01 | 5,27E-01 | 39 | 32 | 7 | 21 | 1631 | 1984 | 1162 | 1400 | 1,3 | 0,5 | 0,8 |
| chr6:26031316-26031816:downstream_4905 | chr6 | 26031816 | 26032288 | - | HIST1H3B | NM | 196 | 1,0 | 1 | -1 | 2,03E-01 | 5,27E-01 | 6 | 4 | 8 | 16 | 150 | 92 | 511 | 777 | -1,4 | -2,4 | 1,0 |
| chr5:110466200-110466700:downstream_4362 | chr5 | 110427869 | 110466200 | + | WDR36 | NM | 24 | 1,5 | 1 | -1 | 2,04E-01 | NA | 4 | 5 | 11 | 2 | 26 | 9 | 72 | 65 | -0,6 | -2,0 | 1,4 |
| chr15:101610317-101610817:downstream_11529 | chr15 | 101459419 | 101610317 | + | LRRK1 | NM | 7 | -2,1 | 2 | 1 | 2,04E-01 | NA | 2 | 2 | 4 | 6 | 7 | 20 | 6 | 12 | -1,5 | 0,6 | -2,1 |
| chr2:27353124-27353624:downstream_1900 | chr2 | 27353624 | 27357542 | - | PREB | NM | 54 | 1,0 | 1 | -1 | 2,05E-01 | 5,30E-01 | 24 | 36 | 33 | 10 | 78 | 61 | 97 | 97 | 0,5 | -0,5 | 0,9 |
| chr20:55214338-55214838:downstream_15100 | chr20 | 55204357 | 55214338 | + | TFAP2C | NM | 16 | 1,4 | 1 | -1 | 2,06E-01 | NA | 9 | 14 | 6 | 11 | 18 | 13 | 26 | 35 | 0,4 | -1,0 | 1,4 |
| chr7:150935913-150936413:downstream_25448 | chr7 | 150929574 | 150935913 | + | CHPF2 | NM | 11 | 1,8 | 1 | -1 | 2,07E-01 | NA | 8 | 5 | 3 | 2 | 8 | 20 | 22 | 16 | 1,3 | -0,4 | 1,7 |
| chr8:135812262-135812762:downstream_6735 | chr8 | 135812762 | 135812850 | - | MIR30B | NR | 43 | -1,0 | 1 | 1 | 2,07E-01 | 5,33E-01 | 58 | 53 | 113 | 84 | 10 | 12 | 13 | 7 | -0,8 | 0,2 | -1,0 |
| chr10:32297437-32297937:downstream_8470 | chr10 | 32297937 | 32345371 | - | KIF5B | NM | 130 | 1,0 | 1 | -1 | 2,07E-01 | 5,33E-01 | 3 | 15 | 10 | 5 | 209 | 175 | 352 | 269 | 0,3 | -0,7 | 1,0 |
| chr1:185265021-185265521:downstream_1267 | chr1 | 185265521 | 185286461 | - | IVNS1ABP | NM | 37 | 1,1 | 1 | -1 | 2,07E-01 | 5,33E-01 | 25 | 16 | 28 | 11 | 45 | 28 | 72 | 66 | 0,1 | -0,9 | 1,0 |
| chr14:24910547-24911047:downstream_19268 | chr14 | 24898491 | 24910547 | + | KHNYN | NM | 11 | 1,6 | 1 | -1 | 2,08E-01 | NA | 23 | 15 | 7 | 5 | 10 | 12 | 10 | 10 | 1,6 | 0,1 | 1,5 |
| chrX:73802310-73802810:downstream_27008 | chrX | 73802810 | 73834461 | - | RLIM | NM | 39 | 1,1 | 1 | -1 | 2,08E-01 | 5,34E-01 | 7 | 22 | 6 | 7 | 71 | 68 | 71 | 58 | 1,3 | 0,1 | 1,2 |
| chr6:76428001-76428501:downstream_25192 | chr6 | 76311224 | 76428001 | + | SENP6 | NM | 14 | -1,7 | 1 | 1 | 2,08E-01 | NA | 13 | 3 | 36 | 16 | 5 | 16 | 13 | 9 | -1,7 | 0,0 | -1,7 |
| chr5:82878122-82878622:downstream_24439 | chr5 | 82767492 | 82878122 | + | VCAN | NM | 27 | 1,2 | 1 | -1 | 2,08E-01 | NA | 5 | 5 | 15 | 8 | 15 | 13 | 85 | 71 | -1,3 | -2,5 | 1,2 |
| chr2:220237679-220238179:downstream_2446 | chr2 | 220238179 | 220252662 | - | DNPEP | NM | 6 | -2,3 | 2 | 1 | 2,08E-01 | NA | 2 | 0 | 4 | 7 | 10 | 8 | 7 | 8 | -2,1 | 0,2 | -2,3 |
| chr3:193415600-193416100:downstream_23319 | chr3 | 193310932 | 193415600 | + | OPA1 | NM | 7 | 2,6 | 2 | -1 | 2,09E-01 | NA | 2 | 0 | 3 | 7 | 2 | 0 | 22 | 17 | -1,9 | -4,6 | 2,7 |
| chr6:147711612-147712112:downstream_25282 | chr6 | 147525493 | 147711612 | + | STXBP5 | NM | 4 | -3,1 | 2 | 1 | 2,09E-01 | NA | 0 | 0 | 11 | 6 | 2 | 1 | 13 | 3 | -Inf | -2,6 | -Inf |
| chr1:210416440-210416940:downstream_1351 | chr1 | 210406194 | 210416440 | + | SERTAD4 | NM | 5 | 2,3 | 2 | -1 | 2,09E-01 | NA | 3 | 8 | 4 | 9 | 1 | 2 | 11 | 6 | -0,3 | -2,6 | 2,4 |
| chrX:57623910-57624410:downstream_7762 | chrX | 57618268 | 57623910 | + | ZXDB | NM | 19 | 1,4 | 1 | -1 | 2,09E-01 | NA | 12 | 10 | 3 | 2 | 40 | 38 | 26 | 23 | 2,1 | 0,7 | 1,4 |
| chr14:101416240-101416740:downstream_11333 | chr14 | 101416169 | 101416240 | + | SNORD114-1 | NR | 3 | 3,4 | 3 | -1 | 2,10E-01 | NA | 7 | 2 | 0 | 0 | 9 | 2 | 3 | 2 | Inf | 1,1 | Inf |
| chr1:36522063-36522563:downstream_16453 | chr1 | 36396682 | 36522063 | + | AGO3 | NM | 15 | 1,8 | 1 | -1 | 2,10E-01 | NA | 2 | 26 | 22 | 16 | 2 | 7 | 21 | 21 | -0,4 | -2,2 | 1,7 |
| chr17:41145707-41146207:downstream_13142 | chr17 | 41132581 | 41145707 | + | RUNDC1 | NM | 13 | 1,6 | 1 | -1 | 2,10E-01 | NA | 11 | 5 | 7 | 2 | 16 | 15 | 18 | 29 | 0,9 | -0,6 | 1,5 |
| chr1:27190947-27191447:downstream_575 | chr1 | 27189632 | 27190947 | + | SFN | NM | 21 | -1,7 | 1 | 1 | 2,10E-01 | NA | 7 | 7 | 4 | 0 | 65 | 76 | 4 | 7 | 1,7 | 3,7 | -2,1 |
| chr19:42585720-42586220:downstream_14327 | chr19 | 42580289 | 42585720 | + | ZNF574 | NM | 26 | -1,2 | 1 | 1 | 2,10E-01 | NA | 15 | 21 | 14 | 6 | 58 | 69 | 11 | 17 | 0,9 | 2,2 | -1,3 |
| chr19:38795646-38796146:downstream_14258 | chr19 | 38794803 | 38795646 | + | C19orf33 | NM | 347 | -0,8 | 1 | 1 | 2,10E-01 | 5,39E-01 | 10 | 12 | 10 | 14 | 809 | 838 | 643 | 441 | -0,1 | 0,6 | -0,8 |
| chr4:39499874-39500374:downstream_23904 | chr4 | 39500374 | 39529218 | - | UGDH | NM | 90 | 0,9 | 1 | -1 | 2,11E-01 | 5,39E-01 | 25 | 7 | 38 | 19 | 76 | 74 | 234 | 250 | -0,8 | -1,7 | 0,9 |
| chr2:174770686-174771186:downstream_22243 | chr2 | 174771186 | 174830430 | - | SP3 | NM | 14 | 1,4 | 1 | -1 | 2,11E-01 | NA | 16 | 16 | 13 | 20 | 9 | 4 | 13 | 22 | 0,0 | -1,4 | 1,5 |
| chr4:16259810-16260310:downstream_23879 | chr4 | 16228285 | 16259810 | + | TAPT1-AS1 | NR | 5 | 2,8 | 2 | -1 | 2,11E-01 | NA | 2 | 8 | 10 | 9 | 1 | 0 | 8 | 3 | -0,9 | -3,8 | 3,0 |
| chr8:95138893-95139393:downstream_25879 | chr8 | 95139393 | 95229531 | - | CDH17 | NM | 16 | -1,8 | 1 | 1 | 2,11E-01 | NA | 2 | 0 | 11 | 7 | 13 | 15 | 39 | 44 | -3,5 | -1,5 | -2,0 |
| chr11:64855874-64856374:downstream_9428 | chr11 | 64851693 | 64855874 | + | ZFPL1 | NM | 6 | 2,4 | 2 | -1 | 2,11E-01 | NA | 2 | 8 | 3 | 0 | 7 | 10 | 10 | 12 | 1,8 | -0,4 | 2,1 |
| chr21:43717354-43717854:downstream_22618 | chr21 | 43619798 | 43717354 | + | ABCG1 | NM | 6 | 2,1 | 2 | -1 | 2,12E-01 | NA | 5 | 5 | 7 | 8 | 3 | 0 | 14 | 10 | -0,6 | -2,9 | 2,2 |
| chr4:68480978-68481478:downstream_3452 | chr4 | 68481478 | 68566889 | - | UBA6 | NM | 5 | 3,0 | 2 | -1 | 2,12E-01 | NA | 6 | 0 | 11 | 13 | 0 | 0 | 3 | 8 | -2,1 | -Inf | Inf |
| chr17:1614297-1614797:downstream_20584 | chr17 | 1614797 | 1619566 | - | MIR22HG | NR | 59 | 2,0 | 2 | -1 | 2,12E-01 | 5,41E-01 | 7 | 15 | 0 | 1 | 176 | 196 | 51 | 23 | 4,7 | 2,3 | 2,4 |
| chr16:85710779-85711279:downstream_12035 | chr16 | 85711279 | 85722588 | - | GINS2 | NM | 8 | 2,0 | 2 | -1 | 2,12E-01 | NA | 3 | 11 | 1 | 3 | 8 | 10 | 7 | 18 | 1,6 | -0,5 | 2,1 |
| chr20:46285649-46286149:downstream_22578 | chr20 | 46286149 | 46415360 | - | SULF2 | NM | 13 | 1,6 | 1 | -1 | 2,13E-01 | NA | 5 | 6 | 1 | 3 | 15 | 21 | 33 | 20 | 1,2 | -0,6 | 1,8 |
| chr19:35446757-35447257:downstream_14209 | chr19 | 35447257 | 35454953 | - | ZNF792 | NM | 9 | 1,9 | 2 | -1 | 2,14E-01 | NA | 2 | 5 | 6 | 6 | 2 | 5 | 29 | 13 | -0,6 | -2,5 | 1,9 |
| chr6:160199029-160199529:downstream_25299 | chr6 | 160199529 | 160210735 | - | TCP1 | NM | 41 | 1,1 | 1 | -1 | 2,14E-01 | 5,46E-01 | 11 | 10 | 4 | 5 | 51 | 98 | 75 | 75 | 1,2 | 0,0 | 1,2 |
| chr5:137685418-137685918:downstream_24119 | chr5 | 137673223 | 137685418 | + | FAM53C | NM | 20 | 2,1 | 2 | -1 | 2,14E-01 | NA | 11 | 2 | 1 | 0 | 50 | 58 | 14 | 25 | 3,2 | 1,5 | 1,7 |
| chr3:28361263-28361763:downstream_2540 | chr3 | 28283123 | 28361263 | + | CMC1 | NM | 8 | 2,0 | 2 | -1 | 2,14E-01 | NA | 2 | 15 | 3 | 11 | 4 | 4 | 8 | 16 | 0,4 | -1,6 | 2,1 |
| chr22:42903840-42904340:downstream_15657 | chr22 | 42904340 | 42915829 | - | RRP7A | NM | 10 | -2,2 | 2 | 1 | 2,15E-01 | NA | 8 | 6 | 0 | 3 | 31 | 27 | 4 | 0 | 2,1 | 3,8 | -1,7 |
| chr3:123212862-123213362:downstream_3143 | chr3 | 123213362 | 123303924 | - | HACD2 | NM | 327 | -0,7 | 1 | 1 | 2,15E-01 | 5,48E-01 | 310 | 167 | 501 | 454 | 204 | 320 | 347 | 313 | -1,0 | -0,3 | -0,7 |
| chr8:6501140-6501640:downstream_26009 | chr8 | 6264112 | 6501140 | + | MCPH1 | NM | 6 | 2,9 | 2 | -1 | 2,16E-01 | NA | 0 | 9 | 11 | 11 | 1 | 0 | 6 | 11 | -1,3 | -4,3 | 3,0 |
| chr3:100467811-100468311:downstream_23463 | chr3 | 100428133 | 100467811 | + | TFG | NM | 50 | 1,1 | 1 | -1 | 2,16E-01 | 5,48E-01 | 11 | 8 | 15 | 5 | 51 | 67 | 100 | 139 | -0,1 | -1,0 | 0,9 |
| chr5:102884055-102884555:downstream_24263 | chr5 | 102884555 | 102898502 | - | NUDT12 | NM | 3 | -3,5 | 3 | 1 | 2,16E-01 | NA | 0 | 0 | 1 | 8 | 4 | 0 | 4 | 7 | -Inf | -1,4 | -Inf |
| chr9:3823627-3824127:downstream_26336 | chr9 | 3824127 | 4300035 | - | GLIS3 | NM | 18 | 2,0 | 2 | -1 | 2,16E-01 | NA | 1 | 1 | 10 | 3 | 3 | 1 | 68 | 53 | -2,9 | -4,8 | 2,0 |
| chr11:69479831-69480331:downstream_9501 | chr11 | 69480331 | 69490165 | - | ORAOV1 | NM | 9 | 2,4 | 2 | -1 | 2,16E-01 | NA | 18 | 9 | 0 | 1 | 11 | 26 | 6 | 2 | 5,0 | 2,2 | 2,8 |
| chr3:118930088-118930588:downstream_23019 | chr3 | 118930588 | 118959752 | - | B4GALT4 | NM | 292 | -0,7 | 1 | 1 | 2,16E-01 | 5,49E-01 | 15 | 13 | 35 | 25 | 473 | 497 | 636 | 644 | -1,1 | -0,4 | -0,7 |
| chr16:576346-576846:downstream_12065 | chr16 | 576846 | 577407 | - | LINC00235 | NR | 15 | 1,6 | 1 | -1 | 2,17E-01 | NA | 32 | 25 | 8 | 5 | 15 | 14 | 18 | 2 | 2,1 | 0,5 | 1,6 |
| chr11:66435858-66436358:downstream_18250 | chr11 | 66406087 | 66435858 | + | RBM4 | NM | 14 | 1,7 | 1 | -1 | 2,17E-01 | NA | 3 | 15 | 3 | 2 | 24 | 21 | 15 | 26 | 1,8 | 0,1 | 1,6 |
| chr17:63107212-63107712:downstream_13312 | chr17 | 63096929 | 63107212 | + | LOC100507002 | NR | 9 | -1,9 | 2 | 1 | 2,17E-01 | NA | 6 | 0 | 14 | 13 | 13 | 5 | 10 | 12 | -2,2 | -0,3 | -1,9 |
| chr12:112339706-112340206:downstream_10426 | chr12 | 112336866 | 112339706 | + | ADAM1A | NR | 14 | -1,7 | 1 | 1 | 2,18E-01 | NA | 3 | 0 | 8 | 10 | 15 | 17 | 39 | 15 | -2,5 | -0,7 | -1,8 |
| chr7:75518244-75518744:downstream_25709 | chr7 | 75508316 | 75518244 | + | RHBDD2 | NM | 20 | -1,4 | 1 | 1 | 2,18E-01 | NA | 6 | 3 | 10 | 2 | 55 | 41 | 18 | 28 | -0,5 | 1,1 | -1,6 |
| chr7:105730313-105730813:downstream_25764 | chr7 | 105730813 | 105753093 | - | SYPL1 | NM | 66 | 0,8 | 1 | -1 | 2,19E-01 | 5,54E-01 | 20 | 22 | 18 | 18 | 107 | 69 | 141 | 133 | 0,2 | -0,6 | 0,9 |
| chr4:125584703-125585203:downstream_23557 | chr4 | 125585203 | 125633887 | - | ANKRD50 | NM | 11 | 1,7 | 1 | -1 | 2,19E-01 | NA | 2 | 13 | 11 | 10 | 3 | 6 | 19 | 25 | -0,6 | -2,3 | 1,7 |
| chr1:149858518-149859018:downstream_979 | chr1 | 149859018 | 149859466 | - | HIST2H2AB | NM | 553 | 0,8 | 1 | -1 | 2,19E-01 | 5,54E-01 | 6 | 9 | 21 | 22 | 444 | 243 | 1952 | 1727 | -1,6 | -2,4 | 0,8 |
| chr19:9920442-9920942:downstream_13982 | chr19 | 9920942 | 9929731 | - | FBXL12 | NM | 18 | 1,9 | 2 | -1 | 2,19E-01 | NA | 10 | 5 | 1 | 1 | 65 | 18 | 28 | 16 | 2,7 | 0,9 | 1,8 |
| chr5:172658606-172659106:downstream_24251 | chr5 | 172659106 | 172662315 | - | NKX2-5 | NM | 5 | -2,7 | 2 | 1 | 2,19E-01 | NA | 7 | 3 | 0 | 4 | 18 | 8 | 0 | 2 | 1,2 | 4,0 | -2,8 |
| chr6:138409141-138409641:downstream_4673 | chr6 | 138409641 | 138428660 | - | PERP | NM | 10 | 1,7 | 1 | -1 | 2,20E-01 | NA | 10 | 13 | 1 | 3 | 15 | 13 | 14 | 7 | 2,3 | 0,4 | 1,9 |
| chr12:56438007-56438507:downstream_10249 | chr12 | 56435685 | 56438007 | + | RPS26 | NM | 24 | 1,3 | 1 | -1 | 2,20E-01 | NA | 67 | 69 | 14 | 16 | 11 | 10 | 0 | 10 | 2,2 | 1,0 | 1,2 |
| chr4:26757835-26758335:downstream_23882 | chr4 | 26585545 | 26757835 | + | TBC1D19 | NM | 8 | -2,1 | 2 | 1 | 2,20E-01 | NA | 5 | 0 | 17 | 18 | 2 | 8 | 10 | 6 | -2,8 | -0,7 | -2,1 |
| chr2:132241032-132241532:downstream_2255 | chr2 | 132241532 | 132250064 | - | MZT2A | NM | 16 | 1,4 | 1 | -1 | 2,20E-01 | NA | 7 | 15 | 3 | 7 | 20 | 19 | 33 | 20 | 1,1 | -0,4 | 1,5 |
| chr11:61169619-61170119:downstream_17966 | chr11 | 61170119 | 61197464 | - | CPSF7 | NM | 28 | 1,1 | 1 | -1 | 2,20E-01 | 5,56E-01 | 15 | 12 | 4 | 12 | 42 | 32 | 64 | 45 | 0,7 | -0,6 | 1,2 |
| chr9:2654485-2654985:downstream_26569 | chr9 | 2621792 | 2654485 | + | VLDLR | NM | 13 | 1,9 | 2 | -1 | 2,21E-01 | NA | 7 | 0 | 15 | 13 | 4 | 0 | 36 | 30 | -2,1 | -4,0 | 1,9 |
| chr13:52950802-52951302:downstream_19110 | chr13 | 52951302 | 52980629 | - | THSD1 | NM | 4 | 3,0 | 2 | -1 | 2,21E-01 | NA | 2 | 8 | 0 | 0 | 10 | 9 | 3 | 3 | Inf | 1,6 | Inf |
| chr12:118470042-118470542:downstream_18797 | chr12 | 118454505 | 118470042 | + | RFC5 | NM | 27 | 1,1 | 1 | -1 | 2,21E-01 | NA | 7 | 10 | 10 | 13 | 21 | 22 | 67 | 69 | -0,5 | -1,6 | 1,1 |
| chr3:185206888-185207388:downstream_3303 | chr3 | 185207388 | 185216845 | - | TMEM41A | NM | 18 | 1,3 | 1 | -1 | 2,21E-01 | NA | 18 | 7 | 7 | 7 | 21 | 23 | 24 | 36 | 0,9 | -0,4 | 1,3 |
| chr8:30041155-30041655:downstream_6281 | chr8 | 30013812 | 30041155 | + | DCTN6 | NM | 31 | -1,2 | 1 | 1 | 2,21E-01 | 5,57E-01 | 8 | 2 | 13 | 9 | 55 | 57 | 54 | 48 | -1,1 | 0,1 | -1,2 |
| chr2:9545314-9545814:downstream_21994 | chr2 | 9545814 | 9563643 | - | ITGB1BP1 | NM | 5 | 2,3 | 2 | -1 | 2,21E-01 | NA | 3 | 9 | 3 | 2 | 6 | 1 | 8 | 7 | 1,2 | -1,2 | 2,4 |
| chr2:219314248-219314748:downstream_2422 | chr2 | 219283837 | 219314248 | + | VIL1 | NM | 11 | 1,8 | 1 | -1 | 2,21E-01 | NA | 16 | 15 | 25 | 19 | 2 | 0 | 8 | 5 | -0,5 | -2,4 | 1,9 |
| chrX:16862642-16863142:downstream_27091 | chrX | 16804554 | 16862642 | + | TXLNG | NM | 14 | -1,5 | 1 | 1 | 2,21E-01 | NA | 7 | 9 | 17 | 11 | 12 | 32 | 6 | 20 | -0,8 | 0,8 | -1,6 |
| chr16:1469244-1469744:downstream_12088 | chr16 | 1469744 | 1470801 | - | C16orf91 | NM | 45 | 2,9 | 2 | -1 | 2,22E-01 | 5,59E-01 | 1 | 14 | 0 | 0 | 149 | 139 | 18 | 42 | Inf | 2,3 | Inf |
| chr19:51300449-51300949:downstream_21032 | chr19 | 51300949 | 51308110 | - | C19orf48 | NM | 18 | 1,5 | 1 | -1 | 2,22E-01 | NA | 8 | 14 | 0 | 7 | 32 | 24 | 19 | 40 | 1,6 | -0,1 | 1,6 |
| chr4:129796379-129796879:downstream_23707 | chr4 | 129730777 | 129796379 | + | JADE1 | NM | 17 | 1,8 | 2 | -1 | 2,22E-01 | NA | 3 | 0 | 14 | 3 | 1 | 6 | 58 | 51 | -2,4 | -4,0 | 1,6 |
| chr8:135816618-135817118:downstream_6736 | chr8 | 135817118 | 135817188 | - | MIR30D | NR | 31 | 1,1 | 1 | -1 | 2,22E-01 | 5,59E-01 | 49 | 36 | 61 | 65 | 6 | 3 | 17 | 12 | -0,6 | -1,8 | 1,2 |
| chr6:27776445-27776945:downstream_4948 | chr6 | 27775976 | 27776445 | + | HIST1H2AI | NM | 65 | 0,8 | 1 | -1 | 2,23E-01 | 5,59E-01 | 20 | 11 | 42 | 37 | 32 | 42 | 159 | 178 | -1,4 | -2,2 | 0,8 |
| chr19:36031139-36031639:downstream_14219 | chr19 | 36031639 | 36036931 | - | TMEM147-AS1 | NR | 14 | 1,4 | 1 | -1 | 2,23E-01 | NA | 15 | 33 | 8 | 6 | 12 | 17 | 13 | 11 | 1,8 | 0,4 | 1,4 |
| chr12:53874946-53875446:downstream_18729 | chr12 | 53845885 | 53874946 | + | PCBP2 | NM | 11 | 1,7 | 1 | -1 | 2,23E-01 | NA | 4 | 6 | 1 | 2 | 16 | 15 | 22 | 21 | 1,4 | -0,4 | 1,8 |
| chr19:39999121-39999621:downstream_21099 | chr19 | 39989556 | 39999121 | + | DLL3 | NM | 8 | 2,0 | 2 | -1 | 2,23E-01 | NA | 14 | 5 | 4 | 1 | 7 | 15 | 13 | 7 | 1,9 | 0,1 | 1,8 |
| chr8:146072893-146073393:downstream_26198 | chr8 | 146052902 | 146072893 | + | ZNF7 | NM | 10 | 2,0 | 2 | -1 | 2,23E-01 | NA | 20 | 7 | 1 | 1 | 15 | 22 | 8 | 4 | 3,6 | 1,6 | 2,0 |
| chr12:58351052-58351552:downstream_10287 | chr12 | 58335444 | 58351052 | + | XRCC6BP1 | NM | 6 | 2,3 | 2 | -1 | 2,24E-01 | NA | 3 | 14 | 7 | 2 | 5 | 1 | 8 | 6 | 1,0 | -1,3 | 2,2 |
| chr3:118924000-118924500:downstream_3121 | chr3 | 118892424 | 118924000 | + | UPK1B | NM | 60 | 1,1 | 1 | -1 | 2,24E-01 | 5,62E-01 | 7 | 14 | 7 | 2 | 138 | 97 | 97 | 119 | 1,1 | 0,1 | 1,0 |
| chr6:151679694-151680194:downstream_24472 | chr6 | 151561133 | 151679694 | + | AKAP12 | NM | 18 | 1,4 | 1 | -1 | 2,24E-01 | NA | 7 | 3 | 3 | 7 | 16 | 20 | 45 | 46 | 0,1 | -1,3 | 1,4 |
| chr12:56118726-56119226:downstream_18492 | chr12 | 56119226 | 56123457 | - | CD63 | NM | 346 | -0,8 | 1 | 1 | 2,25E-01 | 5,62E-01 | 11 | 9 | 8 | 8 | 881 | 955 | 399 | 495 | 0,2 | 1,0 | -0,8 |
| chr1:45976739-45977239:downstream_706 | chr1 | 45965855 | 45976739 | + | MMACHC | NM | 34 | 1,4 | 1 | -1 | 2,25E-01 | 5,62E-01 | 42 | 117 | 49 | 43 | 2 | 7 | 1 | 10 | 0,8 | -0,4 | 1,2 |
| chr12:66516348-66516848:downstream_10297 | chr12 | 66516848 | 66524533 | - | LLPH | NM | 24 | 1,2 | 1 | -1 | 2,25E-01 | NA | 19 | 22 | 7 | 2 | 50 | 43 | 25 | 21 | 2,1 | 1,0 | 1,1 |
| chr9:35812456-35812956:downstream_7064 | chr9 | 35812956 | 35815042 | - | HINT2 | NM | 18 | 1,8 | 1 | -1 | 2,25E-01 | NA | 4 | 5 | 0 | 2 | 41 | 35 | 19 | 39 | 2,4 | 0,4 | 2,1 |
| chr14:59015549-59016049:downstream_19270 | chr14 | 58894102 | 59015549 | + | KIAA0586 | NM | 5 | 2,5 | 2 | -1 | 2,25E-01 | NA | 1 | 3 | 4 | 6 | 2 | 0 | 10 | 17 | -1,4 | -4,1 | 2,6 |
| chrX:106183463-106183963:downstream_26929 | chrX | 106183963 | 106243474 | - | MORC4 | NM | 114 | 0,8 | 1 | -1 | 2,26E-01 | 5,62E-01 | 19 | 30 | 11 | 24 | 200 | 162 | 224 | 245 | 0,5 | -0,4 | 0,9 |
| chr2:37428274-37428774:downstream_1937 | chr2 | 37428774 | 37458740 | - | CEBPZ | NM | 58 | 1,0 | 1 | -1 | 2,26E-01 | 5,62E-01 | 8 | 12 | 3 | 6 | 123 | 100 | 113 | 99 | 1,2 | 0,1 | 1,1 |
| chr1:1336775-1337275:downstream_373 | chr1 | 1337275 | 1342693 | - | MRPL20 | NM | 126 | 1,0 | 1 | -1 | 2,27E-01 | 5,62E-01 | 13 | 3 | 0 | 8 | 222 | 222 | 245 | 294 | 1,0 | -0,3 | 1,2 |
| chr17:26654850-26655350:downstream_20520 | chr17 | 26655350 | 26662515 | - | IFT20 | NM | 72 | 0,8 | 1 | -1 | 2,27E-01 | 5,62E-01 | 35 | 54 | 24 | 22 | 112 | 123 | 109 | 98 | 1,0 | 0,2 | 0,8 |
| chr17:68176183-68176683:downstream_13324 | chr17 | 68165675 | 68176183 | + | KCNJ2 | NM | 14 | -2,2 | 2 | 1 | 2,28E-01 | NA | 1 | 0 | 19 | 11 | 7 | 3 | 40 | 29 | -5,2 | -2,9 | -2,3 |
| chr4:175410827-175411327:downstream_23697 | chr4 | 175411327 | 175444049 | - | HPGD | NM | 21 | 1,3 | 1 | -1 | 2,28E-01 | NA | 2 | 10 | 4 | 6 | 28 | 19 | 46 | 56 | 0,3 | -1,1 | 1,4 |
| chr1:149872348-149872848:downstream_977 | chr1 | 149871154 | 149872348 | + | BOLA1 | NM | 29 | 1,1 | 1 | -1 | 2,28E-01 | 5,65E-01 | 22 | 11 | 14 | 10 | 24 | 47 | 56 | 52 | 0,5 | -0,6 | 1,1 |
| chr2:238007489-238007989:downstream_21846 | chr2 | 237994083 | 238007489 | + | COPS8 | NM | 35 | 1,0 | 1 | -1 | 2,29E-01 | 5,65E-01 | 65 | 63 | 25 | 11 | 48 | 23 | 17 | 24 | 1,8 | 0,8 | 1,0 |
| chr12:120532398-120532898:downstream_18777 | chr12 | 120532898 | 120554643 | - | RAB35 | NM | 8 | 2,1 | 2 | -1 | 2,29E-01 | NA | 5 | 4 | 0 | 2 | 9 | 18 | 10 | 16 | 2,4 | 0,1 | 2,4 |
| chr6:26104565-26105065:downstream_4914 | chr6 | 26104175 | 26104565 | + | HIST1H4C | NM | 114 | 1,1 | 1 | -1 | 2,29E-01 | 5,66E-01 | 14 | 6 | 10 | 2 | 286 | 140 | 170 | 285 | 0,8 | -0,1 | 0,9 |
| chr15:89002208-89002708:downstream_11876 | chr15 | 89002708 | 89010633 | - | MRPL46 | NM | 27 | 1,1 | 1 | -1 | 2,29E-01 | NA | 24 | 15 | 6 | 11 | 41 | 42 | 36 | 39 | 1,3 | 0,1 | 1,1 |
| chr12:19675173-19675673:downstream_18416 | chr12 | 19592607 | 19675173 | + | AEBP2 | NM | 7 | -2,1 | 2 | 1 | 2,29E-01 | NA | 2 | 0 | 10 | 8 | 7 | 5 | 8 | 11 | -2,9 | -0,7 | -2,2 |
| chr22:30884397-30884897:downstream_22905 | chr22 | 30884897 | 30901698 | - | SEC14L4 | NM | 5 | 2,8 | 2 | -1 | 2,30E-01 | NA | 0 | 14 | 6 | 7 | 2 | 0 | 6 | 6 | 0,1 | -2,8 | 2,9 |
| chr19:36525386-36525886:downstream_14238 | chr19 | 36525886 | 36545664 | - | THAP8 | NM | 8 | 2,1 | 2 | -1 | 2,30E-01 | NA | 5 | 0 | 3 | 7 | 3 | 2 | 25 | 17 | -0,9 | -3,0 | 2,1 |
| chr2:25961752-25962252:downstream_1587 | chr2 | 25962252 | 26101312 | - | ASXL2 | NM | 40 | 1,0 | 1 | -1 | 2,30E-01 | 5,67E-01 | 14 | 12 | 82 | 72 | 7 | 4 | 71 | 58 | -2,6 | -3,6 | 1,0 |
| chr2:55462989-55463489:downstream_22197 | chr2 | 55459038 | 55462989 | + | RPS27A | NM | 101 | 0,8 | 1 | -1 | 2,30E-01 | 5,67E-01 | 44 | 31 | 6 | 12 | 298 | 196 | 106 | 116 | 2,1 | 1,2 | 0,9 |
| chr9:80032399-80032899:downstream_26570 | chr9 | 79792360 | 80032399 | + | VPS13A | NM | 17 | -1,3 | 1 | 1 | 2,31E-01 | NA | 7 | 3 | 21 | 21 | 13 | 18 | 26 | 30 | -2,2 | -0,9 | -1,3 |
| chr3:113161064-113161564:downstream_3106 | chr3 | 113161564 | 113234034 | - | SPICE1 | NM | 4 | -3,1 | 3 | 1 | 2,31E-01 | NA | 1 | 0 | 6 | 7 | 0 | 7 | 4 | 7 | -3,9 | -0,7 | -3,2 |
| chr15:65737497-65737997:downstream_19531 | chr15 | 65737997 | 65810035 | - | DPP8 | NM | 47 | -0,9 | 1 | 1 | 2,31E-01 | 5,67E-01 | 37 | 22 | 111 | 98 | 13 | 23 | 36 | 31 | -1,8 | -0,9 | -0,9 |
| chr6:80752244-80752744:downstream_25330 | chr6 | 80714321 | 80752244 | + | TTK | NM | 17 | 1,4 | 1 | -1 | 2,31E-01 | NA | 2 | 5 | 8 | 6 | 10 | 9 | 47 | 47 | -1,0 | -2,3 | 1,4 |
| chr20:39753126-39753626:downstream_14796 | chr20 | 39657461 | 39753126 | + | TOP1 | NM | 12 | 1,5 | 1 | -1 | 2,31E-01 | NA | 5 | 11 | 10 | 7 | 9 | 7 | 18 | 28 | -0,1 | -1,5 | 1,5 |
| chr11:64894250-64894750:downstream_18330 | chr11 | 64894750 | 64902003 | - | SYVN1 | NM | 22 | 1,3 | 1 | -1 | 2,31E-01 | NA | 11 | 48 | 10 | 8 | 33 | 23 | 15 | 26 | 1,7 | 0,4 | 1,3 |
| chr8:6260576-6261076:downstream_6382 | chr8 | 6261076 | 6264069 | - | LOC100287015 | NR | 8 | 1,8 | 2 | -1 | 2,31E-01 | NA | 19 | 9 | 4 | 3 | 9 | 5 | 8 | 5 | 1,9 | 0,1 | 1,8 |
| chr16:75476641-75477141:downstream_20247 | chr16 | 75477141 | 75498635 | - | TMEM170A | NM | 15 | 1,5 | 1 | -1 | 2,32E-01 | NA | 9 | 2 | 3 | 3 | 19 | 21 | 35 | 29 | 0,8 | -0,7 | 1,5 |
| chr7:34967992-34968492:downstream_5539 | chr7 | 34968492 | 35077653 | - | DPY19L1 | NM | 114 | -0,8 | 1 | 1 | 2,32E-01 | 5,67E-01 | 5 | 11 | 25 | 27 | 176 | 112 | 287 | 271 | -1,7 | -1,0 | -0,8 |
| chr1:150594098-150594598:downstream_16694 | chr1 | 150594598 | 150602098 | - | ENSA | NM | 132 | -0,8 | 1 | 1 | 2,33E-01 | 5,67E-01 | 10 | 6 | 22 | 17 | 235 | 180 | 333 | 250 | -1,3 | -0,5 | -0,9 |
| chr4:24521314-24521814:downstream_3640 | chr4 | 24521814 | 24521913 | - | MIR573 | NR | 13 | 1,7 | 1 | -1 | 2,33E-01 | NA | 16 | 1 | 26 | 13 | 3 | 3 | 22 | 22 | -1,2 | -2,9 | 1,7 |
| chr17:40961649-40962149:downstream_13134 | chr17 | 40962149 | 40976310 | - | BECN1 | NM | 112 | 1,0 | 1 | -1 | 2,33E-01 | 5,67E-01 | 16 | 10 | 0 | 9 | 214 | 266 | 200 | 178 | 1,5 | 0,3 | 1,2 |
| chr4:96079601-96080101:downstream_23572 | chr4 | 95679127 | 96079601 | + | BMPR1B | NM | 44 | 0,9 | 1 | -1 | 2,34E-01 | 5,67E-01 | 19 | 13 | 72 | 89 | 9 | 6 | 74 | 74 | -2,4 | -3,3 | 1,0 |
| chr1:94984219-94984719:downstream_16436 | chr1 | 94883932 | 94984219 | + | ABCD3 | NM | 41 | -1,4 | 1 | 1 | 2,34E-01 | 5,67E-01 | 2 | 0 | 7 | 11 | 30 | 51 | 102 | 121 | -2,9 | -1,5 | -1,5 |
| chr14:56768031-56768531:downstream_10908 | chr14 | 56585092 | 56768031 | + | PELI2 | NM | 7 | 2,6 | 2 | -1 | 2,34E-01 | NA | 0 | 3 | 0 | 10 | 1 | 1 | 19 | 21 | -1,8 | -4,5 | 2,7 |
| chr20:42824080-42824580:downstream_15020 | chr20 | 42824580 | 42839546 | - | OSER1 | NM | 7 | 2,0 | 2 | -1 | 2,34E-01 | NA | 9 | 19 | 6 | 1 | 5 | 7 | 3 | 7 | 2,1 | 0,3 | 1,8 |
| chr13:53050766-53051266:downstream_18954 | chr13 | 53029494 | 53050766 | + | CKAP2 | NM | 7 | 2,2 | 2 | -1 | 2,34E-01 | NA | 0 | 12 | 3 | 2 | 5 | 9 | 7 | 21 | 1,1 | -1,1 | 2,2 |
| chr16:58740534-58741034:downstream_20002 | chr16 | 58741034 | 58768261 | - | GOT2 | NM | 78 | 1,4 | 1 | -1 | 2,34E-01 | 5,67E-01 | 8 | 0 | 1 | 2 | 153 | 166 | 136 | 161 | 1,4 | 0,1 | 1,3 |
| chr4:24528587-24529087:downstream_3641 | chr4 | 24529087 | 24586184 | - | DHX15 | NM | 43 | 0,9 | 1 | -1 | 2,34E-01 | 5,67E-01 | 33 | 53 | 49 | 52 | 22 | 28 | 46 | 61 | -0,2 | -1,1 | 0,9 |
| chr6:41034030-41034530:downstream_5124 | chr6 | 41034530 | 41040188 | - | OARD1 | NM | 37 | -1,1 | 1 | 1 | 2,35E-01 | 5,67E-01 | 6 | 4 | 17 | 12 | 62 | 44 | 86 | 64 | -1,6 | -0,5 | -1,1 |
| chr3:49453909-49454409:downstream_2957 | chr3 | 49449638 | 49453909 | + | TCTA | NM | 26 | 1,2 | 1 | -1 | 2,35E-01 | NA | 25 | 19 | 4 | 7 | 37 | 59 | 39 | 16 | 2,1 | 0,8 | 1,3 |
| chr2:11321277-11321777:downstream_1573 | chr2 | 11321777 | 11484711 | - | ROCK2 | NM | 85 | 0,7 | 1 | -1 | 2,35E-01 | 5,67E-01 | 38 | 44 | 149 | 135 | 25 | 21 | 150 | 117 | -1,8 | -2,5 | 0,7 |
| chr6:129897739-129898239:downstream_4670 | chr6 | 129898239 | 130031370 | - | ARHGAP18 | NM | 77 | 0,8 | 1 | -1 | 2,35E-01 | 5,67E-01 | 20 | 9 | 72 | 64 | 22 | 26 | 188 | 212 | -2,3 | -3,1 | 0,8 |
| chr12:125395690-125396190:downstream_10496 | chr12 | 125396190 | 125399587 | - | UBC | NM | 231 | 0,6 | 1 | -1 | 2,35E-01 | 5,67E-01 | 103 | 71 | 67 | 45 | 424 | 366 | 363 | 408 | 0,6 | 0,0 | 0,6 |
| chr2:233448355-233448855:downstream_21896 | chr2 | 233415296 | 233448355 | + | EIF4E2 | NM | 8 | 1,8 | 2 | -1 | 2,35E-01 | NA | 7 | 2 | 1 | 6 | 7 | 5 | 22 | 16 | 0,2 | -1,7 | 1,9 |
| chr9:22452472-22452972:downstream_7016 | chr9 | 22446839 | 22452472 | + | DMRTA1 | NM | 14 | 1,8 | 2 | -1 | 2,35E-01 | NA | 6 | 45 | 17 | 19 | 7 | 0 | 11 | 6 | 0,5 | -1,4 | 1,9 |
| chr15:23813166-23813666:downstream_11552 | chr15 | 23810453 | 23813166 | + | MKRN3 | NM | 29 | -1,6 | 1 | 1 | 2,35E-01 | 5,67E-01 | 94 | 64 | 33 | 21 | 7 | 13 | 1 | 1 | 1,6 | 3,1 | -1,6 |
| chr7:128301052-128301552:downstream_6076 | chr7 | 128281294 | 128301052 | + | LINC01000 | NR | 15 | -1,6 | 1 | 1 | 2,36E-01 | NA | 2 | 0 | 19 | 17 | 9 | 6 | 36 | 32 | -3,9 | -2,2 | -1,7 |
| chr7:91501690-91502190:downstream_25640 | chr7 | 91502190 | 91510034 | - | MTERF1 | NM | 24 | 1,2 | 1 | -1 | 2,36E-01 | NA | 12 | 33 | 4 | 13 | 37 | 31 | 29 | 34 | 1,4 | 0,1 | 1,3 |
| chr12:117468953-117469453:downstream_18565 | chr12 | 117348760 | 117468953 | + | FBXW8 | NM | 10 | -1,9 | 2 | 1 | 2,36E-01 | NA | 0 | 2 | 6 | 7 | 11 | 11 | 26 | 16 | -2,7 | -0,9 | -1,8 |
| chr1:145585992-145586492:downstream_947 | chr1 | 145586492 | 145589435 | - | NUDT17 | NM | 8 | -2,0 | 2 | 1 | 2,37E-01 | NA | 11 | 2 | 8 | 9 | 22 | 3 | 1 | 7 | -0,5 | 1,6 | -2,1 |
| chr4:10075462-10075962:downstream_23993 | chr4 | 10075962 | 10118573 | - | WDR1 | NM | 80 | 0,9 | 1 | -1 | 2,37E-01 | 5,71E-01 | 8 | 11 | 6 | 7 | 120 | 150 | 189 | 149 | 0,6 | -0,3 | 1,0 |
| chr9:37779210-37779710:downstream_26301 | chr9 | 37779710 | 37785089 | - | EXOSC3 | NM | 29 | -1,0 | 1 | 1 | 2,38E-01 | 5,71E-01 | 42 | 37 | 40 | 26 | 29 | 34 | 15 | 11 | 0,2 | 1,2 | -1,0 |
| chr15:59912210-59912710:downstream_11740 | chr15 | 59903981 | 59912210 | + | GCNT3 | NM | 517 | 0,5 | 0 | -1 | 2,38E-01 | 5,71E-01 | 68 | 81 | 200 | 147 | 437 | 395 | 1461 | 1343 | -1,2 | -1,8 | 0,5 |
| chr6:160180790-160181290:downstream_5425 | chr6 | 160181290 | 160183364 | - | LOC100129518 | NR | 4 | 2,6 | 2 | -1 | 2,38E-01 | NA | 4 | 7 | 1 | 4 | 3 | 0 | 6 | 5 | 1,0 | -1,7 | 2,7 |
| chr4:128838459-128838959:downstream_3503 | chr4 | 128838959 | 128887139 | - | MFSD8 | NM | 10 | -2,0 | 2 | 1 | 2,39E-01 | NA | 2 | 0 | 10 | 2 | 15 | 10 | 24 | 19 | -2,9 | -0,8 | -2,1 |
| chr8:99203886-99204386:downstream_6336 | chr8 | 99204386 | 99306621 | - | NIPAL2 | NM | 16 | -1,3 | 1 | 1 | 2,39E-01 | NA | 9 | 4 | 15 | 16 | 22 | 19 | 25 | 17 | -1,3 | 0,0 | -1,3 |
| chr15:101843632-101844132:downstream_19661 | chr15 | 101844132 | 102030187 | - | PCSK6 | NM | 27 | -1,1 | 1 | 1 | 2,40E-01 | NA | 5 | 5 | 17 | 16 | 39 | 29 | 65 | 39 | -1,7 | -0,6 | -1,1 |
| chr1:247262763-247263263:downstream_17434 | chr1 | 247263263 | 247267674 | - | ZNF669 | NM | 14 | -1,3 | 1 | 1 | 2,40E-01 | NA | 11 | 18 | 11 | 14 | 20 | 23 | 4 | 10 | 0,2 | 1,6 | -1,4 |

| | | | | | | | | | | | | | | | | | | | | | | | |
|---|---|---|---|---|---|---|---|---|---|---|---|---|---|---|---|---|---|---|---|---|---|---|---|
| chr7:19734584-19735084:downstream_5725 | chr7 | 19735084 | 19748660 | - | TWISTNB | NM | 17 | 1,5 | 1 | -1 | 2,40E-01 | NA | 12 | 10 | 1 | 2 | 44 | 33 | 14 | 17 | 2,9 | 1,3 | 1,5 |
| chr3:160969795-160970295:downstream_2735 | chr3 | 160939098 | 160969795 | + | NMD3 | NM | 31 | 1,3 | 1 | -1 | 2,40E-01 | 5,75E-01 | 18 | 5 | 4 | 3 | 78 | 44 | 56 | 38 | 1,6 | 0,4 | 1,2 |
| chr1:95362004-95362504:downstream_16612 | chr1 | 95362504 | 95392779 | - | CNN3 | NM | 16 | -1,6 | 1 | 1 | 2,40E-01 | NA | 2 | 1 | 7 | 8 | 20 | 18 | 22 | 51 | -2,5 | -0,9 | -1,6 |
| chr15:101108927-101109427:downstream_11918 | chr15 | 101109427 | 101142445 | - | LINS | NM | 14 | -1,6 | 1 | 1 | 2,40E-01 | NA | 2 | 1 | 8 | 4 | 17 | 27 | 29 | 21 | -1,9 | -0,2 | -1,7 |
| chr1:226331879-226332379:downstream_300 | chr1 | 226332379 | 226374423 | - | ACBD3 | NM | 32 | 0,9 | 1 | -1 | 2,41E-01 | 5,76E-01 | 40 | 27 | 39 | 25 | 20 | 25 | 46 | 36 | 0,1 | -0,9 | 0,9 |
| chr19:49315773-49316273:downstream_14456 | chr19 | 49316273 | 49339934 | - | HSD17B14 | NM | 12 | -1,6 | 1 | 1 | 2,41E-01 | NA | 8 | 1 | 10 | 12 | 25 | 12 | 19 | 11 | -1,3 | 0,3 | -1,6 |
| chr8:101166230-101166730:downstream_6663 | chr8 | 101162838 | 101166230 | + | POLR2K | NM | 42 | 1,0 | 1 | -1 | 2,41E-01 | 5,76E-01 | 24 | 16 | 8 | 3 | 99 | 83 | 46 | 52 | 1,8 | 0,9 | 0,9 |
| chr12:54788544-54789044:downstream_10223 | chr12 | 54789044 | 54813050 | - | ITGA5 | NM | 9 | 2,1 | 2 | -1 | 2,41E-01 | NA | 2 | 7 | 0 | 2 | 15 | 15 | 8 | 19 | 2,5 | 0,2 | 2,3 |
| chr17:18834599-18835099:downstream_20648 | chr17 | 18759611 | 18834599 | + | PRPSAP2 | NM | 13 | 1,5 | 1 | -1 | 2,42E-01 | NA | 4 | 9 | 3 | 3 | 14 | 23 | 28 | 23 | 1,1 | -0,5 | 1,5 |
| chr17:79647723-79648223:downstream_13433 | chr17 | 79648223 | 79650954 | - | ARL16 | NM | 32 | 1,4 | 1 | -1 | 2,42E-01 | 5,76E-01 | 6 | 3 | 0 | 2 | 61 | 67 | 68 | 52 | 1,8 | 0,1 | 1,7 |
| chr11:73583212-73583712:downstream_9527 | chr11 | 73583712 | 73587890 | - | COA4 | NM | 77 | 0,9 | 1 | -1 | 2,42E-01 | 5,76E-01 | 22 | 21 | 6 | 7 | 214 | 145 | 99 | 103 | 1,7 | 0,8 | 0,9 |
| chr3:81538349-81538849:downstream_2551 | chr3 | 81538849 | 81810950 | - | GBE1 | NM | 22 | -1,7 | 1 | 1 | 2,43E-01 | NA | 2 | 0 | 6 | 7 | 16 | 34 | 64 | 45 | -2,9 | -1,1 | -1,8 |
| chr10:64568239-64568739:downstream_8536 | chr10 | 64564515 | 64568239 | + | ADO | NM | 78 | -0,9 | 1 | 1 | 2,43E-01 | 5,78E-01 | 24 | 20 | 6 | 18 | 171 | 251 | 70 | 64 | 0,9 | 1,7 | -0,8 |
| chr17:38459413-38459913:downstream_13044 | chr17 | 38444145 | 38459413 | + | CDC6 | NM | 16 | 1,3 | 1 | -1 | 2,43E-01 | NA | 12 | 13 | 24 | 7 | 13 | 6 | 28 | 30 | -0,3 | -1,6 | 1,3 |
| chr3:14153076-14153576:downstream_23074 | chr3 | 14153576 | 14166371 | - | CHCHD4 | NM | 16 | 1,7 | 1 | -1 | 2,44E-01 | NA | 3 | 7 | 3 | 0 | 29 | 36 | 21 | 27 | 1,8 | 0,4 | 1,4 |
| chr14:54893146-54893646:downstream_11147 | chr14 | 54893646 | 54908148 | - | CNIH1 | NM | 107 | -0,8 | 1 | 1 | 2,44E-01 | 5,79E-01 | 70 | 21 | 54 | 70 | 194 | 168 | 124 | 157 | -0,4 | 0,4 | -0,8 |
| chr20:62375403-62375903:downstream_15159 | chr20 | 62371210 | 62375403 | + | SLC2A4RG | NM | 10 | 1,9 | 2 | -1 | 2,44E-01 | NA | 6 | 4 | 0 | 2 | 23 | 9 | 14 | 21 | 2,0 | -0,1 | 2,1 |
| chr6:4994779-4995279:downstream_25179 | chr6 | 4995279 | 5004297 | - | RPP40 | NM | 7 | 2,1 | 2 | -1 | 2,44E-01 | NA | 4 | 5 | 3 | 0 | 9 | 11 | 14 | 8 | 1,7 | -0,2 | 1,9 |
| chr17:62223831-62224331:downstream_13306 | chr17 | 62223698 | 62223831 | + | SNORA76C | NR | 41 | -0,9 | 1 | 1 | 2,45E-01 | 5,79E-01 | 65 | 87 | 63 | 69 | 16 | 17 | 6 | 9 | 0,2 | 1,2 | -1,0 |
| chr1:161726954-161727454:downstream_1182 | chr1 | 161719557 | 161726954 | + | DUSP12 | NM | 27 | 1,3 | 1 | -1 | 2,45E-01 | NA | 7 | 4 | 3 | 4 | 28 | 46 | 64 | 57 | 0,6 | -0,7 | 1,3 |
| chr11:95565857-95566357:downstream_17949 | chr11 | 95523624 | 95565857 | + | CEP57 | NM | 7 | -2,5 | 2 | 1 | 2,45E-01 | NA | 2 | 0 | 11 | 2 | 19 | 0 | 11 | 15 | -3,1 | -0,5 | -2,6 |
| chr1:67872992-67873492:downstream_17225 | chr1 | 67873492 | 67896123 | - | SERBP1 | NM | 21 | 1,3 | 1 | -1 | 2,45E-01 | NA | 15 | 13 | 0 | 5 | 52 | 37 | 22 | 25 | 2,5 | 0,9 | 1,5 |
| chr3:8616354-8616854:downstream_2802 | chr3 | 8613467 | 8616354 | + | LINC00312 | NR | 6 | -2,7 | 2 | 1 | 2,45E-01 | NA | 0 | 0 | 10 | 7 | 3 | 0 | 14 | 14 | -Inf | -3,1 | -Inf |
| chr5:150069851-150070351:downstream_4109 | chr5 | 150070351 | 150080669 | - | RBM22 | NM | 54 | 1,3 | 1 | -1 | 2,45E-01 | 5,80E-01 | 10 | 6 | 4 | 0 | 108 | 167 | 63 | 78 | 1,9 | 1,0 | 0,9 |
| chr20:33765165-33765665:downstream_14986 | chr20 | 33759739 | 33765165 | + | PROCR | NM | 262 | -0,7 | 1 | 1 | 2,46E-01 | 5,80E-01 | 24 | 13 | 15 | 16 | 740 | 593 | 359 | 336 | 0,2 | 0,9 | -0,7 |
| chr14:74432680-74433180:downstream_10911 | chr14 | 74433180 | 74486026 | - | ENTPD5 | NM | 29 | 1,1 | 1 | -1 | 2,46E-01 | 5,80E-01 | 7 | 26 | 17 | 24 | 20 | 23 | 54 | 62 | -0,3 | -1,4 | 1,1 |
| chr10:112364392-112364892:downstream_8345 | chr10 | 112327448 | 112364392 | + | SMC3 | NM | 19 | 1,3 | 1 | -1 | 2,46E-01 | NA | 7 | 7 | 7 | 10 | 11 | 18 | 31 | 60 | -0,3 | -1,6 | 1,3 |
| chr8:124827690-124828190:downstream_6702 | chr8 | 124780881 | 124827690 | + | FAM91A1 | NM | 17 | 1,3 | 1 | -1 | 2,46E-01 | NA | 10 | 4 | 6 | 6 | 12 | 24 | 38 | 37 | 0,3 | -1,0 | 1,3 |
| chr11:812876-813376:downstream_8987 | chr11 | 809935 | 812876 | + | RPLP2 | NM | 885 | 1,2 | 1 | -1 | 2,46E-01 | 5,80E-01 | 7 | 7 | 1 | 2 | 2511 | 2110 | 1283 | 1156 | 2,1 | 0,9 | 1,2 |
| chr12:104382656-104383156:downstream_10391 | chr12 | 104359592 | 104382656 | + | TDG | NM | 23 | 1,1 | 1 | -1 | 2,47E-01 | NA | 25 | 13 | 8 | 13 | 29 | 28 | 28 | 43 | 0,8 | -0,3 | 1,1 |
| chr6:44221625-44222125:downstream_24810 | chr6 | 44213902 | 44221625 | + | HSP90AB1 | NM | 143 | -0,8 | 1 | 1 | 2,47E-01 | 5,80E-01 | 12 | 39 | 32 | 33 | 277 | 308 | 202 | 244 | -0,3 | 0,4 | -0,7 |
| chr19:53961515-53962015:downstream_21700 | chr19 | 53935226 | 53961515 | + | ZNF761 | NM | 9 | 2,1 | 2 | -1 | 2,47E-01 | NA | 18 | 3 | 4 | 3 | 16 | 0 | 7 | 18 | 1,5 | -0,6 | 2,1 |
| chr11:118617972-118618472:downstream_17981 | chr11 | 118618472 | 118661972 | - | DDX6 | NM | 19 | -1,5 | 1 | 1 | 2,47E-01 | NA | 2 | 0 | 8 | 4 | 25 | 27 | 45 | 38 | -2,4 | -0,7 | -1,7 |
| chr4:56771244-56771744:downstream_23638 | chr4 | 56719815 | 56771244 | + | EXOC1 | NM | 29 | 1,2 | 1 | -1 | 2,47E-01 | 5,81E-01 | 6 | 2 | 4 | 6 | 36 | 17 | 92 | 73 | -0,4 | -1,6 | 1,3 |
| chr11:85667713-85668213:downstream_18207 | chr11 | 85668213 | 85780923 | - | PICALM | NM | 5 | -2,0 | 2 | 1 | 2,48E-01 | NA | 2 | 2 | 6 | 5 | 7 | 11 | 7 | 3 | -1,3 | 0,7 | -2,0 |
| chr8:48648975-48649475:downstream_6549 | chr8 | 48649475 | 48650726 | - | CEBPD | NM | 51 | 1,0 | 1 | -1 | 2,49E-01 | 5,84E-01 | 2 | 9 | 24 | 16 | 25 | 19 | 173 | 143 | -1,8 | -2,8 | 1,0 |
| chr5:94799098-94799598:downstream_4343 | chr5 | 94799598 | 94890709 | - | TTC37 | NM | 21 | -1,5 | 1 | 1 | 2,49E-01 | NA | 2 | 0 | 8 | 7 | 16 | 34 | 54 | 44 | -2,6 | -1,0 | -1,6 |
| chr17:18710026-18710526:downstream_12887 | chr17 | 18684581 | 18710026 | + | TVP23B | NM | 19 | 1,5 | 1 | -1 | 2,49E-01 | NA | 23 | 3 | 3 | 5 | 46 | 20 | 25 | 30 | 1,7 | 0,2 | 1,5 |
| chr20:43570270-43570770:downstream_15032 | chr20 | 43570770 | 43589114 | - | TOMM34 | NM | 8 | 2,1 | 2 | -1 | 2,50E-01 | NA | 19 | 17 | 1 | 1 | 6 | 18 | 1 | 4 | 4,0 | 2,1 | 1,9 |
| chr12:56366573-56367073:downstream_18495 | chr12 | 56360552 | 56366573 | + | CDK2 | NM | 18 | 1,4 | 1 | -1 | 2,50E-01 | NA | 9 | 6 | 0 | 5 | 24 | 38 | 29 | 34 | 1,6 | 0,0 | 1,6 |
| chr10:101909346-101909846:downstream_17558 | chr10 | 101909846 | 101945814 | - | ERLIN1 | NM | 18 | 1,4 | 1 | -1 | 2,50E-01 | NA | 4 | 8 | 0 | 6 | 24 | 28 | 32 | 46 | 1,0 | -0,6 | 1,6 |
| chr10:120068071-120068571:downstream_17564 | chr10 | 120068571 | 120101839 | - | FAM204A | NM | 10 | -1,6 | 1 | 1 | 2,50E-01 | NA | 11 | 15 | 15 | 19 | 6 | 8 | 6 | 1 | -0,4 | 1,1 | -1,5 |
| chr11:65633914-65634414:downstream_9455 | chr11 | 65627871 | 65633914 | + | MUS81 | NM | 14 | 1,5 | 1 | -1 | 2,50E-01 | NA | 3 | 7 | 4 | 2 | 18 | 15 | 28 | 31 | 0,6 | -0,8 | 1,4 |
| chr2:210885644-210886144:downstream_21999 | chr2 | 210886144 | 211036094 | - | KANSL1L | NM | 14 | 1,6 | 1 | -1 | 2,50E-01 | NA | 2 | 22 | 8 | 3 | 10 | 21 | 18 | 27 | 1,0 | -0,5 | 1,6 |
| chr10:133780686-133781186:downstream_8787 | chr10 | 133781186 | 133795517 | - | BNIP3 | NM | 41 | 1,1 | 1 | -1 | 2,51E-01 | 5,86E-01 | 11 | 7 | 3 | 5 | 81 | 77 | 65 | 80 | 1,2 | 0,1 | 1,1 |
| chr22:19160345-19160845:downstream_15461 | chr22 | 19159218 | 19160345 | + | LINC01311 | NR | 11 | -1,7 | 1 | 1 | 2,52E-01 | NA | 15 | 13 | 6 | 2 | 20 | 27 | 1 | 2 | 1,8 | 3,6 | -1,8 |
| chr3:128338312-128338812:downstream_3171 | chr3 | 128338812 | 128369719 | - | RPN1 | NM | 191 | 1,1 | 1 | -1 | 2,52E-01 | 5,89E-01 | 2 | 8 | 4 | 1 | 412 | 370 | 342 | 391 | 1,0 | 0,1 | 0,9 |
| chr1:67454302-67454802:downstream_16954 | chr1 | 67390577 | 67454302 | + | MIER1 | NM | 8 | -1,8 | 2 | 1 | 2,53E-01 | NA | 1 | 6 | 7 | 8 | 12 | 11 | 11 | 5 | -1,2 | 0,5 | -1,7 |
| chr19:52188115-52188615:downstream_14542 | chr19 | 52188615 | 52196709 | - | SPACA6P-AS | NR | 6 | 2,2 | 2 | -1 | 2,53E-01 | NA | 3 | 7 | 10 | 8 | 2 | 0 | 10 | 5 | -0,8 | -3,2 | 2,3 |
| chr15:83684680-83685180:downstream_19480 | chr15 | 83685180 | 83736106 | - | BTBD1 | NM | 52 | -1,0 | 1 | 1 | 2,54E-01 | 5,91E-01 | 2 | 5 | 10 | 8 | 92 | 85 | 129 | 87 | -1,3 | -0,3 | -1,0 |
| chr1:87575681-87576181:downstream_16807 | chr1 | 87380334 | 87575681 | + | HS2ST1 | NM | 4 | -2,8 | 2 | 1 | 2,54E-01 | NA | 0 | 0 | 8 | 7 | 2 | 1 | 6 | 11 | -Inf | -2,7 | -Inf |
| chr12:388722-389222:downstream_9801 | chr12 | 389222 | 498621 | - | KDM5A | NM | 36 | 0,9 | 1 | -1 | 2,54E-01 | 5,92E-01 | 13 | 14 | 51 | 38 | 11 | 13 | 77 | 74 | -1,7 | -2,6 | 0,9 |
| chr8:86126753-86127253:downstream_25913 | chr8 | 86089618 | 86126753 | + | E2F5 | NM | 9 | 1,6 | 1 | -1 | 2,54E-01 | NA | 6 | 8 | 13 | 6 | 6 | 2 | 15 | 16 | -0,4 | -2,1 | 1,6 |
| chr1:116934986-116935486:downstream_16500 | chr1 | 116935486 | 116961244 | - | ATP1A1-AS1 | NR | 12 | 1,4 | 1 | -1 | 2,54E-01 | NA | 16 | 19 | 6 | 10 | 11 | 9 | 10 | 13 | 1,2 | -0,2 | 1,4 |
| chr7:79848725-79849225:downstream_25536 | chr7 | 79764139 | 79848725 | + | GNAI1 | NM | 13 | 1,4 | 1 | -1 | 2,55E-01 | NA | 7 | 12 | 15 | 7 | 5 | 11 | 26 | 21 | -0,3 | -1,6 | 1,3 |
| chr3:100074478-100074978:downstream_3076 | chr3 | 100053561 | 100074478 | + | NIT2 | NM | 37 | -0,9 | 1 | 1 | 2,55E-01 | 5,92E-01 | 16 | 22 | 28 | 20 | 43 | 83 | 42 | 39 | -0,3 | 0,6 | -0,9 |
| chr2:196602426-196602926:downstream_22225 | chr2 | 196521531 | 196602426 | + | SLC39A10 | NM | 33 | 1,3 | 1 | -1 | 2,55E-01 | 5,92E-01 | 2 | 8 | 4 | 2 | 54 | 48 | 74 | 72 | 0,7 | -0,5 | 1,2 |
| chr3:130735555-130736055:downstream_23015 | chr3 | 130569368 | 130735555 | + | ATP2C1 | NM | 17 | -1,4 | 1 | 1 | 2,55E-01 | NA | 2 | 2 | 7 | 8 | 35 | 16 | 29 | 39 | -1,8 | -0,4 | -1,4 |
| chr7:77409120-77409620:downstream_5578 | chr7 | 77325742 | 77409120 | + | RSBN1L | NM | 10 | -1,9 | 2 | 1 | 2,55E-01 | NA | 2 | 0 | 6 | 21 | 2 | 12 | 21 | 18 | -3,4 | -1,5 | -1,9 |
| chr17:62493873-62494373:downstream_13307 | chr17 | 62494373 | 62502484 | - | DDX5 | NM | 40 | 1,0 | 1 | -1 | 2,56E-01 | 5,92E-01 | 33 | 38 | 11 | 11 | 68 | 69 | 64 | 24 | 1,6 | 0,6 | 1,0 |
| chr11:117156404-117156904:downstream_18262 | chr11 | 117103340 | 117156404 | + | RNF214 | NM | 9 | 1,9 | 2 | -1 | 2,56E-01 | NA | 2 | 7 | 8 | 7 | 7 | 0 | 32 | 9 | -0,7 | -2,7 | 2,0 |
| chr13:37523407-37523907:downstream_18933 | chr13 | 37523907 | 37573504 | - | ALG5 | NM | 10 | 1,6 | 1 | -1 | 2,56E-01 | NA | 4 | 7 | 1 | 4 | 15 | 10 | 18 | 23 | 1,0 | -0,7 | 1,7 |
| chr19:44104587-44105087:downstream_21681 | chr19 | 44100543 | 44104587 | + | ZNF576 | NM | 9 | -1,6 | 1 | 1 | 2,56E-01 | NA | 10 | 7 | 4 | 14 | 13 | 16 | 6 | 6 | -0,1 | 1,4 | -1,5 |
| chr1:180084015-180084515:downstream_252 | chr1 | 179923907 | 180084015 | + | CEP350 | NM | 8 | -2,3 | 2 | 1 | 2,56E-01 | NA | 0 | 0 | 3 | 7 | 4 | 4 | 24 | 23 | -Inf | -2,6 | -Inf |
| chr5:137522836-137523336:downstream_4441 | chr5 | 137523336 | 137549032 | - | CDC23 | NM | 15 | 1,3 | 1 | -1 | 2,57E-01 | NA | 10 | 13 | 7 | 7 | 19 | 15 | 28 | 21 | 0,7 | -0,5 | 1,3 |
| chr8:80830594-80831094:downstream_6322 | chr8 | 80831094 | 80942506 | - | MRPS28 | NM | 27 | 1,3 | 1 | -1 | 2,57E-01 | NA | 7 | 11 | 17 | 4 | 37 | 9 | 50 | 80 | -0,3 | -1,5 | 1,2 |
| chr12:13068852-13069352:downstream_10070 | chr12 | 13068762 | 13068852 | + | MIR614 | NR | 42 | -0,8 | 1 | 1 | 2,57E-01 | 5,94E-01 | 35 | 25 | 47 | 57 | 41 | 44 | 36 | 48 | -0,8 | 0,0 | -0,8 |
| chr1:249143716-249144216:downstream_1502 | chr1 | 249132376 | 249143716 | + | ZNF672 | NM | 14 | 1,4 | 1 | -1 | 2,57E-01 | NA | 11 | 7 | 1 | 7 | 20 | 18 | 29 | 19 | 1,2 | -0,3 | 1,5 |
| chr17:7215477-7215977:downstream_12799 | chr17 | 7215977 | 7218658 | - | GPS2 | NM | 47 | 0,9 | 1 | -1 | 2,57E-01 | 5,94E-01 | 20 | 44 | 10 | 11 | 78 | 103 | 54 | 59 | 1,6 | 0,7 | 0,9 |
| chr6:109629423-109629923:downstream_5294 | chr6 | 109615505 | 109629423 | + | CCDC162P | NR | 4 | -2,7 | 2 | 1 | 2,57E-01 | NA | 0 | 5 | 6 | 6 | 1 | 9 | 3 | 1 | -1,2 | 1,4 | -2,6 |
| chr5:36102913-36103413:downstream_4222 | chr5 | 36103413 | 36152015 | - | LMBRD2 | NM | 21 | -1,1 | 1 | 1 | 2,58E-01 | NA | 11 | 6 | 29 | 25 | 15 | 22 | 31 | 25 | -1,7 | -0,6 | -1,1 |
| chr15:52838741-52839241:downstream_19469 | chr15 | 52839241 | 52861643 | - | ARPP19 | NM | 52 | 0,9 | 1 | -1 | 2,58E-01 | 5,94E-01 | 21 | 24 | 21 | 12 | 51 | 97 | 85 | 107 | 0,5 | -0,4 | 0,8 |
| chr1:168171351-168171851:downstream_17323 | chr1 | 168148082 | 168171351 | + | TIPRL | NM | 20 | 1,2 | 1 | -1 | 2,58E-01 | NA | 9 | 3 | 4 | 9 | 15 | 22 | 47 | 52 | -0,2 | -1,4 | 1,3 |
| chr2:207179150-207179650:downstream_22330 | chr2 | 207139364 | 207179150 | + | ZDBF2 | NM | 10 | 1,6 | 1 | -1 | 2,58E-01 | NA | 6 | 4 | 7 | 7 | 9 | 1 | 26 | 21 | -0,6 | -2,2 | 1,7 |
| chr20:55093942-55094442:downstream_22541 | chr20 | 55043640 | 55093942 | + | RTFDC1 | NM | 43 | -1,0 | 1 | 1 | 2,58E-01 | 5,95E-01 | 4 | 4 | 8 | 10 | 74 | 75 | 70 | 100 | -1,2 | -0,2 | -1,0 |
| chr4:173961558-173962058:downstream_3392 | chr4 | 172734574 | 173961558 | + | GALNTL6 | NM | 10 | -1,9 | 2 | 1 | 2,59E-01 | NA | 3 | 0 | 10 | 13 | 2 | 17 | 14 | 21 | -2,8 | -0,9 | -2,0 |
| chr13:76434006-76434506:downstream_19019 | chr13 | 76194569 | 76434006 | + | LMO7 | NM | 11 | 1,5 | 1 | -1 | 2,59E-01 | NA | 21 | 5 | 7 | 11 | 9 | 6 | 13 | 17 | 0,5 | -1,0 | 1,5 |
| chr8:38120287-38120787:downstream_25896 | chr8 | 38089008 | 38120287 | + | DDHD2 | NM | 30 | 1,1 | 1 | -1 | 2,59E-01 | 5,97E-01 | 7 | 15 | 10 | 10 | 24 | 44 | 60 | 71 | 0,1 | -1,0 | 1,1 |
| chr10:126676005-126676505:downstream_8768 | chr10 | 126630691 | 126676005 | + | ZRANB1 | NM | 9 | 1,6 | 1 | -1 | 2,60E-01 | NA | 7 | 17 | 13 | 11 | 6 | 1 | 14 | 7 | 0,1 | -1,6 | 1,7 |
| chr17:49375292-49375792:downstream_13243 | chr17 | 49337896 | 49375292 | + | UTP18 | NM | 11 | 1,6 | 1 | -1 | 2,60E-01 | NA | 6 | 6 | 7 | 2 | 16 | 3 | 22 | 24 | 0,3 | -1,3 | 1,6 |
| chr9:124905837-124906337:downstream_6933 | chr9 | 124906337 | 124922098 | - | NDUFA8 | NM | 21 | -1,4 | 1 | 1 | 2,60E-01 | NA | 4 | 0 | 4 | 7 | 28 | 50 | 32 | 44 | -1,4 | 0,0 | -1,4 |
| chr20:3805954-3806454:downstream_22350 | chr20 | 3801170 | 3805954 | + | AP5S1 | NM | 10 | -1,6 | 1 | 1 | 2,60E-01 | NA | 2 | 1 | 6 | 5 | 15 | 17 | 17 | 14 | -1,6 | 0,1 | -1,7 |
| chr17:70398962-70399462:downstream_12685 | chr17 | 70399462 | 70588943 | - | LINC00673 | NR | 18 | 1,5 | 1 | -1 | 2,61E-01 | NA | 0 | 4 | 7 | 13 | 3 | 4 | 49 | 66 | -2,4 | -4,0 | 1,6 |
| chr6:28089072-28089572:downstream_25364 | chr6 | 28089572 | 28105094 | - | ZSCAN16-AS1 | NR | 4 | -2,4 | 2 | 1 | 2,61E-01 | NA | 3 | 6 | 1 | 3 | 7 | 14 | 1 | 1 | 1,0 | 3,2 | -2,2 |
| chr16:24584183-24584683:downstream_20189 | chr16 | 24550907 | 24584183 | + | RBBP6 | NM | 29 | -1,1 | 1 | 1 | 2,61E-01 | 6,00E-01 | 7 | 3 | 6 | 4 | 76 | 72 | 29 | 31 | 0,1 | 1,3 | -1,2 |
| chr5:115628987-115629487:downstream_24070 | chr5 | 115420672 | 115628987 | + | COMMD10 | NM | 30 | -1,2 | 1 | 1 | 2,63E-01 | 6,02E-01 | 4 | 0 | 10 | 9 | 39 | 37 | 71 | 71 | -2,2 | -0,9 | -1,3 |
| chr11:74718743-74719243:downstream_8921 | chr11 | 74699949 | 74718743 | + | NEU3 | NM | 61 | -0,8 | 1 | 1 | 2,63E-01 | 6,02E-01 | 51 | 35 | 26 | 15 | 160 | 127 | 35 | 42 | 1,1 | 1,9 | -0,8 |
| chr11:30359770-30360270:downstream_9195 | chr11 | 30344645 | 30359770 | + | ARL14EP | NM | 24 | 1,1 | 1 | -1 | 2,63E-01 | NA | 27 | 53 | 18 | 5 | 31 | 24 | 21 | 13 | 1,8 | 0,7 | 1,1 |
| chr9:119164600-119165100:downstream_6927 | chr9 | 118916070 | 119164600 | + | PAPPA | NM | 31 | -1,9 | 2 | 1 | 2,63E-01 | 6,02E-01 | 0 | 0 | 3 | 7 | 12 | 15 | 111 | 103 | -Inf | -3,0 | -Inf |
| chr10:103827795-103828295:downstream_8688 | chr10 | 103825123 | 103827795 | + | HPS6 | NM | 21 | 1,1 | 1 | -1 | 2,64E-01 | NA | 17 | 28 | 3 | 10 | 32 | 37 | 19 | 25 | 1,8 | 0,6 | 1,2 |
| chr11:16779901-16780401:downstream_8807 | chr11 | 16760147 | 16779901 | + | C11orf58 | NM | 17 | 1,3 | 1 | -1 | 2,64E-01 | NA | 7 | 13 | 4 | 2 | 25 | 34 | 29 | 21 | 1,5 | 0,2 | 1,3 |
| chr19:45910090-45910590:downstream_21120 | chr19 | 45910590 | 45927177 | - | ERCC1 | NM | 6 | 2,1 | 2 | -1 | 2,64E-01 | NA | 11 | 3 | 3 | 1 | 14 | 3 | 6 | 11 | 1,9 | 0,0 | 1,9 |
| chr7:98992404-98992904:downstream_5968 | chr7 | 98972297 | 98992404 | + | ARPC1B | NM | 38 | -1,2 | 1 | 1 | 2,64E-01 | 6,03E-01 | 6 | 1 | 4 | 11 | 75 | 61 | 50 | 95 | -1,2 | -0,1 | -1,1 |
| chr14:59972124-59972624:downstream_19265 | chr14 | 59951160 | 59972124 | + | JKAMP | NM | 19 | -1,2 | 1 | 1 | 2,64E-01 | NA | 7 | 3 | 15 | 6 | 24 | 39 | 26 | 30 | -1,2 | 0,2 | -1,3 |
| chr5:52410956-52411456:downstream_24208 | chr5 | 52405671 | 52410956 | + | LOC257396 | NR | 4 | 2,7 | 2 | -1 | 2,64E-01 | NA | 4 | 5 | 0 | 1 | 4 | 7 | 3 | 5 | 3,4 | 0,5 | 2,9 |
| chr14:31914742-31915242:downstream_11098 | chr14 | 31915242 | 31926680 | - | DTD2 | NM | 21 | 1,1 | 1 | -1 | 2,64E-01 | NA | 20 | 20 | 11 | 12 | 15 | 33 | 28 | 31 | 0,8 | -0,3 | 1,1 |
| chr1:16160209-16160709:downstream_485 | chr1 | 16160709 | 16174642 | - | FLJ37453 | NR | 22 | -1,1 | 1 | 1 | 2,64E-01 | NA | 22 | 8 | 19 | 11 | 38 | 42 | 21 | 16 | 0,0 | 1,1 | -1,2 |
| chr14:31568820-31569320:downstream_10926 | chr14 | 31569320 | 31676729 | - | HECTD1 | NM | 47 | -1,4 | 1 | 1 | 2,65E-01 | 6,03E-01 | 2 | 0 | 8 | 9 | 49 | 28 | 131 | 152 | -3,4 | -1,9 | -1,5 |
| chr4:119200978-119201478:downstream_23864 | chr4 | 119199916 | 119200978 | + | SNHG8 | NR | 56 | 0,8 | 1 | -1 | 2,65E-01 | 6,03E-01 | 56 | 37 | 35 | 25 | 82 | 60 | 75 | 77 | 0,6 | -0,1 | 0,7 |
| chr20:49578399-49578899:downstream_15083 | chr20 | 49575350 | 49578399 | + | MOCS3 | NM | 26 | 1,0 | 1 | -1 | 2,65E-01 | NA | 40 | 66 | 14 | 9 | 22 | 34 | 11 | 13 | 2,2 | 1,2 | 1,0 |
| chr3:50324662-50325162:downstream_2975 | chr3 | 50325162 | 50330026 | - | IFRD2 | NM | 16 | 1,3 | 1 | -1 | 2,65E-01 | NA | 8 | 8 | 6 | 2 | 28 | 25 | 28 | 27 | 1,1 | 0,0 | 1,2 |
| chr20:60883615-60884115:downstream_15130 | chr20 | 60884115 | 60942368 | - | LAMA5 | NM | 21 | 1,2 | 1 | -1 | 2,66E-01 | NA | 11 | 21 | 3 | 10 | 37 | 28 | 21 | 41 | 1,3 | 0,1 | 1,3 |
| chr11:5683924-5684424:downstream_18368 | chr11 | 5684424 | 5706339 | - | TRIM5 | NM | 4 | -2,4 | 2 | 1 | 2,66E-01 | NA | 0 | 4 | 3 | 8 | 6 | 6 | 1 | 5 | -1,5 | 0,9 | -2,4 |
| chr17:72765499-72765999:downstream_12688 | chr17 | 72744750 | 72765499 | + | SLC9A3R1 | NM | 9 | 2,0 | 2 | -1 | 2,66E-01 | NA | 7 | 2 | 1 | 1 | 21 | 11 | 21 | 7 | 2,1 | 0,2 | 1,9 |
| chr7:139879440-139879940:downstream_6113 | chr7 | 139877060 | 139879440 | + | JHDM1D-AS1 | NR | 11 | 2,2 | 2 | -1 | 2,66E-01 | NA | 11 | 9 | 0 | 0 | 22 | 38 | 6 | 1 | Inf | 3,2 | Inf |
| chr1:38268113-38268613:downstream_114 | chr1 | 38268613 | 38273865 | - | YRDC | NM | 16 | 1,9 | 2 | -1 | 2,67E-01 | NA | 7 | 4 | 0 | 1 | 42 | 44 | 18 | 12 | 3,8 | 1,5 | 2,3 |
| chr2:74731669-74732169:downstream_2053 | chr2 | 74732169 | 74734821 | - | PCGF1 | NM | 7 | -1,8 | 2 | 1 | 2,67E-01 | NA | 3 | 2 | 6 | 6 | 15 | 7 | 6 | 8 | -1,1 | 0,7 | -1,8 |
| chr1:207968861-207969361:downstream_16570 | chr1 | 207925382 | 207968861 | + | CD46 | NM | 115 | -0,8 | 1 | 1 | 2,67E-01 | 6,07E-01 | 10 | 6 | 14 | 17 | 185 | 216 | 227 | 246 | -1,0 | -0,2 | -0,8 |
| chr5:95220301-95220801:downstream_4052 | chr5 | 95220801 | 95297775 | - | ELL2 | NM | 17 | 1,3 | 1 | -1 | 2,67E-01 | NA | 12 | 5 | 13 | 6 | 10 | 19 | 31 | 40 | -0,1 | -1,3 | 1,2 |
| chr14:94593617-94594117:downstream_11280 | chr14 | 94594117 | 94595957 | - | IFI27L2 | NM | 15 | 1,4 | 1 | -1 | 2,68E-01 | NA | 4 | 9 | 4 | 1 | 23 | 31 | 26 | 25 | 1,4 | 0,1 | 1,3 |
| chr2:27586718-27587218:downstream_21895 | chr2 | 27587218 | 27593324 | - | EIF2B4 | NM | 27 | 1,2 | 1 | -1 | 2,68E-01 | NA | 4 | 8 | 3 | 3 | 28 | 59 | 58 | 50 | 1,0 | -0,3 | 1,3 |
| chr11:85367597-85368097:downstream_18340 | chr11 | 85358962 | 85367597 | + | TMEM126A | NM | 15 | 1,4 | 1 | -1 | 2,68E-01 | NA | 12 | 15 | 1 | 4 | 33 | 22 | 10 | 21 | 2,3 | 0,9 | 1,5 |
| chr3:156256841-156257341:downstream_23441 | chr3 | 156257341 | 156272989 | - | SSR3 | NM | 29 | 1,0 | 1 | -1 | 2,68E-01 | 6,07E-01 | 46 | 27 | 33 | 25 | 26 | 15 | 42 | 22 | 0,3 | -0,7 | 1,0 |
| chr1:204337057-204337557:downstream_1319 | chr1 | 204337557 | 204338847 | - | LINC00628 | NR | 5 | 2,1 | 2 | -1 | 2,68E-01 | NA | 7 | 9 | 1 | 2 | 7 | 5 | 1 | 8 | 2,3 | 0,2 | 2,1 |
| chr3:158410360-158410860:downstream_23161 | chr3 | 158362316 | 158410360 | + | GFM1 | NM | 47 | 0,8 | 1 | -1 | 2,69E-01 | 6,07E-01 | 33 | 16 | 24 | 31 | 53 | 39 | 100 | 83 | -0,2 | -1,0 | 0,8 |
| chr18:48700419-48700919:downstream_13629 | chr18 | 48700919 | 48724051 | - | MEX3C | NM | 39 | 1,1 | 1 | -1 | 2,69E-01 | 6,07E-01 | 7 | 10 | 6 | 3 | 48 | 85 | 96 | 54 | 0,9 | -0,2 | 1,0 |
| chr8:145135551-145136051:downstream_6780 | chr8 | 145133521 | 145135551 | + | EXOSC4 | NM | 21 | 1,4 | 1 | -1 | 2,69E-01 | NA | 9 | 21 | 1 | 2 | 59 | 39 | 14 | 20 | 3,0 | 1,5 | 1,4 |
| chr5:96496070-96496570:downstream_24334 | chr5 | 96496570 | 96519005 | - | RIOK2 | NM | 11 | 1,5 | 1 | -1 | 2,69E-01 | NA | 20 | 9 | 7 | 2 | 11 | 17 | 10 | 14 | 1,6 | 0,3 | 1,3 |
| chr19:50886285-50886785:downstream_21308 | chr19 | 50879679 | 50886285 | + | NR1H2 | NM | 31 | 1,0 | 1 | -1 | 2,69E-01 | 6,07E-01 | 20 | 6 | 8 | 7 | 50 | 42 | 51 | 63 | 0,7 | -0,3 | 1,0 |
| chr16:2011561-2012061:downstream_12097 | chr16 | 2012061 | 2014827 | - | RPS2 | NM | 32 | -1,0 | 1 | 1 | 2,69E-01 | 6,07E-01 | 11 | 5 | 17 | 9 | 49 | 72 | 57 | 33 | -0,7 | 0,4 | -1,1 |
| chr1:24022915-24023415:downstream_17195 | chr1 | 24018268 | 24022915 | + | RPL11 | NM | 55 | 0,8 | 1 | -1 | 2,69E-01 | 6,07E-01 | 29 | 31 | 11 | 7 | 115 | 126 | 56 | 64 | 1,8 | 1,0 | 0,8 |
| chr15:76193388-76193888:downstream_19825 | chr15 | 76135626 | 76193388 | + | UBE2Q2 | NM | 9 | 1,6 | 1 | -1 | 2,69E-01 | NA | 2 | 7 | 6 | 11 | 4 | 3 | 17 | 22 | -0,8 | -2,5 | 1,7 |
| chr9:136218159-136218659:downstream_26522 | chr9 | 136218659 | 136223361 | - | SURF1 | NM | 8 | 1,8 | 2 | -1 | 2,70E-01 | NA | 23 | 14 | 4 | 2 | 4 | 7 | 4 | 2 | 2,5 | 0,7 | 1,7 |
| chr5:41941672-41942172:downstream_24123 | chr5 | 41925353 | 41941672 | + | FBXO4 | NM | 9 | 1,7 | 2 | -1 | 2,70E-01 | NA | 2 | 16 | 4 | 2 | 11 | 12 | 8 | 14 | 1,7 | 0,0 | 1,7 |
| chr16:89807332-89807832:downstream_20289 | chr16 | 89787392 | 89807332 | + | ZNF276 | NM | 9 | 1,7 | 2 | -1 | 2,70E-01 | NA | 7 | 4 | 4 | 5 | 11 | 3 | 8 | 30 | 0,2 | -1,5 | 1,7 |
| chr11:100861656-100862156:downstream_8811 | chr11 | 100558406 | 100861656 | + | ARHGAP42 | NM | 16 | -1,5 | 1 | 1 | 2,70E-01 | NA | 2 | 0 | 22 | 21 | 7 | 6 | 40 | 32 | -4,1 | -2,6 | -1,6 |
| chr19:37620651-37621151:downstream_13765 | chr19 | 37569381 | 37620651 | + | ZNF420 | NM | 6 | -2,1 | 2 | 1 | 2,70E-01 | NA | 2 | 0 | 6 | 6 | 10 | 5 | 15 | 8 | -2,8 | -0,7 | -2,1 |
| chr18:9538106-9538606:downstream_13562 | chr18 | 9475529 | 9538106 | + | RALBP1 | NM | 6 | -2,2 | 2 | 1 | 2,71E-01 | NA | 2 | 0 | 6 | 5 | 3 | 12 | 14 | 7 | -2,7 | -0,5 | -2,2 |
| chrUn_gl000220:126696-127196:downstream_16416 | chrUn | gl000220 | 97128 | NR | 126696 | LOC100507412 | 4 | -3,0 | 3 | 1 | 2,71E-01 | NA | 5 | 4 | 1 | 0 | 15 | 10 | 0 | 0 | 2,7 | Inf | -Inf |
| chr5:27496508-27497008:downstream_4204 | chr5 | 27472398 | 27496508 | + | LINC01021 | NR | 37 | 1,0 | 1 | -1 | 2,72E-01 | 6,12E-01 | 37 | 34 | 6 | 2 | 68 | 110 | 18 | 21 | 3,1 | 2,2 | 0,9 |
| chr3:149455756-149456256:downstream_3229 | chr3 | 149456256 | 149470286 | - | COMMD2 | NM | 9 | 1,8 | 2 | -1 | 2,72E-01 | NA | 5 | 7 | 1 | 2 | 6 | 25 | 10 | 18 | 1,9 | 0,2 | 1,8 |
| chr4:74962253-74962753:downstream_3739 | chr4 | 74962753 | 74964997 | - | CXCL2 | NM | 12 | 1,6 | 1 | -1 | 2,73E-01 | NA | 10 | 0 | 11 | 7 | 7 | 4 | 17 | 42 | -0,9 | -2,5 | 1,6 |
| chr18:60994470-60994970:downstream_13654 | chr18 | 60994970 | 61034506 | - | KDSR | NM | 21 | 1,3 | 1 | -1 | 2,73E-01 | NA | 7 | 3 | 4 | 2 | 37 | 27 | 50 | 39 | 0,7 | -0,5 | 1,2 |
| chr7:17829884-17830384:downstream_5517 | chr7 | 17830384 | 17980131 | - | SNX13 | NM | 9 | -2,3 | 2 | 1 | 2,73E-01 | NA | 1 | 0 | 15 | 7 | 8 | 0 | 22 | 19 | -4,7 | -2,3 | -2,4 |
| chr21:44262689-44263189:downstream_22727 | chr21 | 44263189 | 44299699 | - | WDR4 | NM | 5 | 2,0 | 2 | -1 | 2,73E-01 | NA | 7 | 5 | 1 | 2 | 8 | 3 | 7 | 7 | 1,7 | -0,4 | 2,0 |
| chr7:93620499-93620999:downstream_5947 | chr7 | 93620999 | 93633690 | - | BET1 | NM | 19 | -1,3 | 1 | 1 | 2,73E-01 | NA | 37 | 35 | 14 | 14 | 20 | 26 | 8 | 0 | 1,4 | 2,5 | -1,1 |
| chr12:123742506-123743006:downstream_18464 | chr12 | 123717843 | 123742506 | + | C12orf65 | NM | 13 | -1,9 | 2 | 1 | 2,73E-01 | NA | 2 | 0 | 10 | 3 | 25 | 4 | 29 | 29 | -3,0 | -1,0 | -2,0 |
| chr9:33384447-33384947:downstream_7027 | chr9 | 33384947 | 33402517 | - | AQP7 | NM | 8 | -1,7 | 2 | 1 | 2,74E-01 | NA | 1 | 5 | 6 | 5 | 11 | 19 | 13 | 5 | -0,9 | 0,8 | -1,7 |
| chr5:179265077-179265577:downstream_24391 | chr5 | 179233387 | 179265077 | + | SQSTM1 | NM | 9 | 1,6 | 1 | -1 | 2,74E-01 | NA | 11 | 6 | 4 | 2 | 19 | 8 | 13 | 14 | 1,5 | 0,0 | 1,5 |
| chr9:91620069-91620569:downstream_6886 | chr9 | 91606323 | 91620069 | + | S1PR3 | NM | 41 | 1,0 | 1 | -1 | 2,74E-01 | 6,14E-01 | 6 | 9 | 4 | 5 | 65 | 69 | 74 | 99 | 0,7 | -0,4 | 1,0 |
| chr11:6631547-6632047:downstream_9103 | chr11 | 6632047 | 6633475 | - | TAF10 | NM | 51 | -0,8 | 1 | 1 | 2,74E-01 | 6,14E-01 | 28 | 17 | 29 | 17 | 118 | 84 | 50 | 65 | 0,0 | 0,8 | -0,9 |
| chr7:112756272-112756772:downstream_25598 | chr7 | 112756772 | 112758637 | - | LINC00998 | NR | 63 | 1,0 | 1 | -1 | 2,74E-01 | 6,14E-01 | 11 | 6 | 4 | 2 | 149 | 148 | 100 | 85 | 1,6 | 0,7 | 0,9 |
| chr1:8071278-8071778:downstream_417 | chr1 | 8071778 | 8086393 | - | ERRFI1 | NM | 27 | -1,2 | 1 | 1 | 2,74E-01 | NA | 5 | 0 | 22 | 21 | 17 | 17 | 60 | 70 | -3,2 | -1,9 | -1,2 |
| chr1:116947396-116947896:downstream_16499 | chr1 | 116915794 | 116947396 | + | ATP1A1 | NM | 157 | 1,0 | 1 | -1 | 2,75E-01 | 6,14E-01 | 10 | 0 | 4 | 3 | 270 | 248 | 335 | 386 | 0,4 | -0,5 | 0,9 |
| chr2:203070402-203070902:downstream_22267 | chr2 | 203070902 | 203103322 | - | SUMO1 | NM | 9 | -1,6 | 1 | 1 | 2,75E-01 | NA | 2 | 6 | 4 | 7 | 11 | 26 | 10 | 9 | -0,5 | 1,0 | -1,5 |
| chr19:45412652-45413152:downstream_21004 | chr19 | 45409005 | 45412652 | + | APOE | NM | 11 | 2,1 | 2 | -1 | 2,76E-01 | NA | 2 | 8 | 0 | 0 | 33 | 36 | 4 | 7 | Inf | 2,6 | Inf |
| chr2:68403094-68403594:downstream_2005 | chr2 | 68385004 | 68403094 | + | PNO1 | NM | 36 | 0,9 | 1 | -1 | 2,76E-01 | 6,16E-01 | 29 | 25 | 4 | 8 | 81 | 74 | 25 | 45 | 2,1 | 1,1 | 1,0 |
| chr15:91541145-91541645:downstream_19838 | chr15 | 91541645 | 91565851 | - | VPS33B | NM | 9 | 1,5 | 1 | -1 | 2,76E-01 | NA | 18 | 14 | 4 | 7 | 9 | 5 | 8 | 7 | 1,4 | -0,1 | 1,6 |
| chr8:103838029-103838529:downstream_25858 | chr8 | 103838529 | 103876428 | - | AZIN1 | NM | 11 | 1,4 | 1 | -1 | 2,76E-01 | NA | 8 | 12 | 7 | 6 | 10 | 11 | 11 | 22 | 0,6 | -0,7 | 1,3 |
| chr16:11022247-11022747:downstream_12184 | chr16 | 11022747 | 11036257 | - | DEXI | NM | 9 | 1,7 | 2 | -1 | 2,76E-01 | NA | 16 | 6 | 1 | 2 | 12 | 19 | 7 | 7 | 2,9 | 1,1 | 1,7 |
| chr8:59364060-59364560:downstream_6567 | chr8 | 59323822 | 59364060 | + | UBXN2B | NM | 4 | -2,4 | 2 | 1 | 2,76E-01 | NA | 2 | 0 | 4 | 6 | 3 | 7 | 1 | 9 | -2,6 | -0,1 | -2,6 |
| chr8:105500958-105501458:downstream_25996 | chr8 | 105501458 | 105601252 | - | LRP12 | NM | 16 | -1,6 | 1 | 1 | 2,77E-01 | NA | 0 | 2 | 15 | 11 | 6 | 12 | 35 | 44 | -3,7 | -2,2 | -1,6 |
| chr3:122599986-122600486:downstream_2682 | chr3 | 122513900 | 122599986 | + | DIRC2 | NM | 37 | -0,9 | 1 | 1 | 2,77E-01 | 6,17E-01 | 25 | 25 | 78 | 57 | 16 | 28 | 40 | 25 | -1,4 | -0,6 | -0,9 |
| chr7:86954163-86954663:downstream_5929 | chr7 | 86954663 | 86974808 | - | TP53TG1 | NR | 10 | -1,8 | 2 | 1 | 2,77E-01 | NA | 15 | 10 | 3 | 2 | 24 | 24 | 0 | 2 | 2,3 | 4,3 | -2,0 |
| chr12:105196774-105197274:downstream_9916 | chr12 | 105197274 | 105322472 | - | SLC41A2 | NM | 66 | -0,8 | 1 | 1 | 2,77E-01 | 6,17E-01 | 12 | 4 | 78 | 77 | 30 | 25 | 154 | 146 | -3,3 | -2,4 | -0,8 |
| chr11:118873170-118873670:downstream_9682 | chr11 | 118873670 | 118874303 | - | RPL23AP64 | NR | 7 | -1,6 | 2 | 1 | 2,78E-01 | NA | 5 | 6 | 7 | 6 | 12 | 11 | 6 | 3 | -0,2 | 1,4 | -1,6 |

| | | | | | | | | | | | | | | | | | | | | | | | |
|---|---|---|---|---|---|---|---|---|---|---|---|---|---|---|---|---|---|---|---|---|---|---|---|
| chr3:33911199-33911699:downstream_23332 | chr3 | 33840062 | 33911199 | + | PDCD6IP | NM | 35 | 1,1 | 1 | -1 | 2,78E-01 | 6,17E-01 | 7 | 6 | 3 | 12 | 50 | 17 | 104 | 77 | -0,2 | -1,4 | 1,2 |
| chr12:95702289-95702789:downstream_10363 | chr12 | 95702195 | 95702289 | + | MIR331 | NR | 40 | -1,0 | 1 | 1 | 2,78E-01 | 6,17E-01 | 9 | 15 | 141 | 94 | 4 | 7 | 29 | 24 | -3,3 | -2,3 | -1,0 |
| chr2:24337178-24337678:downstream_1888 | chr2 | 24337678 | 24346347 | - | PFN4 | NM | 12 | 1,6 | 1 | -1 | 2,78E-01 | NA | 27 | 36 | 4 | 2 | 11 | 11 | 4 | 2 | 3,4 | 1,9 | 1,6 |
| chr17:10583148-10583648:downstream_12840 | chr17 | 10583648 | 10600885 | - | SCO1 | NM | 52 | 0,8 | 1 | -1 | 2,78E-01 | 6,17E-01 | 20 | 15 | 10 | 8 | 93 | 100 | 95 | 77 | 1,0 | 0,2 | 0,8 |
| chr1:62146218-62146718:downstream_768 | chr1 | 62146718 | 62191095 | - | TM2D1 | NM | 7 | 2,1 | 2 | -1 | 2,79E-01 | NA | 8 | 5 | 4 | 2 | 12 | 0 | 17 | 6 | 1,2 | -0,9 | 2,0 |
| chr2:227595532-227596032:downstream_2461 | chr2 | 227596032 | 227663506 | - | IRS1 | NM | 5 | 2,3 | 2 | -1 | 2,79E-01 | NA | 1 | 10 | 3 | 7 | 0 | 4 | 11 | 6 | 0,2 | -2,1 | 2,3 |
| chr10:81587358-81587858:downstream_8593 | chr10 | 81585657 | 81587358 | + | LOC642361 | NR | 9 | -1,8 | 2 | 1 | 2,79E-01 | NA | 20 | 13 | 14 | 10 | 7 | 4 | 3 | 0 | 0,5 | 2,0 | -1,5 |
| chr9:21330517-21331017:downstream_7005 | chr9 | 21331017 | 21335429 | - | KLHL9 | NM | 33 | 0,9 | 1 | -1 | 2,79E-01 | 6,18E-01 | 44 | 21 | 24 | 16 | 28 | 47 | 38 | 48 | 0,7 | -0,2 | 0,9 |
| chr12:120564513-120565013:downstream_9795 | chr12 | 120565013 | 120632513 | - | GCN1L1 | NM | 69 | 0,8 | 1 | -1 | 2,80E-01 | 6,18E-01 | 12 | 18 | 17 | 11 | 85 | 112 | 121 | 177 | 0,2 | -0,6 | 0,8 |
| chr18:29128814-29129314:downstream_13496 | chr18 | 29078026 | 29128814 | + | DSG2 | NM | 107 | 1,8 | 2 | -1 | 2,80E-01 | 6,18E-01 | 2 | 22 | 13 | 3 | 124 | 130 | 310 | 251 | 0,6 | -1,1 | 1,7 |
| chr16:69376648-69377148:downstream_12437 | chr16 | 69377148 | 69385712 | - | TMED6 | NM | 4 | 2,6 | 2 | -1 | 2,80E-01 | NA | 5 | 5 | 1 | 0 | 2 | 10 | 6 | 2 | 2,8 | 0,5 | 2,3 |
| chr8:37615319-37615819:downstream_25926 | chr8 | 37594096 | 37615319 | + | ERLIN2 | NM | 14 | 1,6 | 1 | -1 | 2,81E-01 | NA | 2 | 8 | 0 | 2 | 24 | 29 | 17 | 30 | 2,0 | 0,2 | 1,8 |
| chr15:26788193-26788693:downstream_19557 | chr15 | 26788693 | 27018935 | - | GABRB3 | NM | 50 | -1,1 | 1 | 1 | 2,81E-01 | 6,19E-01 | 4 | 0 | 10 | 13 | 65 | 41 | 131 | 137 | -2,5 | -1,3 | -1,2 |
| chr1:169365780-169366280:downstream_239 | chr1 | 169337193 | 169365780 | + | BLZF1 | NM | 8 | -1,6 | 1 | 1 | 2,81E-01 | NA | 3 | 8 | 4 | 6 | 24 | 12 | 7 | 4 | 0,1 | 1,7 | -1,5 |
| chr6:160527583-160528083:downstream_4678 | chr6 | 160390130 | 160527583 | + | IGF2R | NM | 10 | 1,6 | 1 | -1 | 2,82E-01 | NA | 3 | 6 | 8 | 18 | 1 | 4 | 17 | 24 | -1,5 | -3,1 | 1,6 |
| chr5:133737255-133737755:downstream_4418 | chr5 | 133737755 | 133747598 | - | CDKN2AIPNL | NM | 25 | -1,1 | 1 | 1 | 2,82E-01 | NA | 5 | 8 | 4 | 15 | 42 | 54 | 40 | 33 | -0,6 | 0,4 | -1,0 |
| chr20:42214273-42214773:downstream_22548 | chr20 | 42187634 | 42214273 | + | SGK2 | NM | 13 | -1,4 | 1 | 1 | 2,82E-01 | NA | 5 | 1 | 22 | 11 | 7 | 14 | 18 | 23 | -2,5 | -1,0 | -1,5 |
| chr14:55516206-55516706:downstream_19401 | chr14 | 55493843 | 55516206 | + | SOCS4 | NM | 32 | 1,0 | 1 | -1 | 2,82E-01 | 6,20E-01 | 49 | 83 | 13 | 9 | 49 | 28 | 8 | 16 | 2,6 | 1,7 | 0,9 |
| chr2:113334729-113335229:downstream_22154 | chr2 | 113299491 | 113334729 | + | POLR1B | NM | 12 | -1,4 | 1 | 1 | 2,83E-01 | NA | 18 | 17 | 14 | 12 | 14 | 10 | 6 | 2 | 0,4 | 1,7 | -1,3 |
| chr1:118503049-118503549:downstream_36 | chr1 | 118472371 | 118503049 | + | WDR3 | NM | 15 | 1,5 | 1 | -1 | 2,83E-01 | NA | 8 | 4 | 0 | 2 | 33 | 29 | 24 | 19 | 2,3 | 0,5 | 1,8 |
| chr22:50882930-50883430:downstream_15707 | chr22 | 50883430 | 50913500 | - | SBF1 | NM | 6 | -1,9 | 2 | 1 | 2,83E-01 | NA | 5 | 7 | 6 | 4 | 7 | 13 | 3 | 2 | 0,3 | 2,1 | -1,8 |
| chr2:46811827-46812327:downstream_1619 | chr2 | 46769866 | 46811827 | + | RHOQ | NM | 10 | -1,5 | 1 | 1 | 2,83E-01 | NA | 4 | 14 | 14 | 14 | 15 | 6 | 7 | 5 | -0,7 | 0,8 | -1,4 |
| chr20:22561141-22561641:downstream_22429 | chr20 | 22561641 | 22566101 | - | FOXA2 | NM | 21 | -1,2 | 1 | 1 | 2,83E-01 | NA | 2 | 6 | 13 | 13 | 14 | 41 | 46 | 37 | -1,8 | -0,6 | -1,2 |
| chr7:56023033-56023533:downstream_5836 | chr7 | 56019610 | 56023033 | + | MRPS17 | NM | 131 | -0,9 | 1 | 1 | 2,83E-01 | 6,20E-01 | 9 | 2 | 3 | 7 | 365 | 329 | 185 | 148 | 0,2 | 1,1 | -0,8 |
| chr5:96110385-96110885:downstream_24044 | chr5 | 95997740 | 96110385 | + | CAST | NM | 129 | 0,7 | 1 | -1 | 2,83E-01 | 6,20E-01 | 17 | 6 | 32 | 30 | 107 | 68 | 376 | 396 | -1,4 | -2,1 | 0,7 |
| chr1:149858961-149859461:downstream_975 | chr1 | 149858524 | 149858961 | + | HIST2H2AC | NM | 305 | 0,7 | 1 | -1 | 2,83E-01 | 6,20E-01 | 11 | 10 | 25 | 16 | 346 | 222 | 950 | 860 | -1,0 | -1,7 | 0,6 |
| chr7:37945034-37945534:downstream_5794 | chr7 | 37945534 | 37956525 | - | SFRP4 | NM | 10 | -2,3 | 2 | 1 | 2,83E-01 | NA | 38 | 17 | 8 | 0 | 4 | 11 | 0 | 0 | 2,7 | Inf | -Inf |
| chr2:233208678-233209178:downstream_21870 | chr2 | 232826292 | 233208678 | + | DIS3L2 | NM | 10 | -1,7 | 2 | 1 | 2,83E-01 | NA | 0 | 3 | 13 | 11 | 8 | 6 | 14 | 23 | -3,0 | -1,4 | -1,6 |
| chr2:202345574-202346074:downstream_22265 | chr2 | 202316391 | 202345574 | + | STRADB | NM | 8 | 1,5 | 1 | -1 | 2,83E-01 | NA | 11 | 10 | 10 | 7 | 5 | 5 | 10 | 11 | 0,4 | -1,1 | 1,4 |
| chr5:141379733-141380233:downstream_4492 | chr5 | 141380233 | 141392620 | - | GNPDA1 | NM | 18 | 1,2 | 1 | -1 | 2,83E-01 | NA | 8 | 10 | 4 | 5 | 26 | 27 | 29 | 32 | 1,0 | -0,2 | 1,2 |
| chr12:118583390-118583890:downstream_10439 | chr12 | 118573869 | 118583390 | + | PEBP1 | NM | 400 | -0,8 | 1 | 1 | 2,84E-01 | 6,22E-01 | 12 | 1 | 10 | 5 | 889 | 1139 | 575 | 573 | -0,2 | 0,8 | -1,0 |
| chr15:40866893-40867393:downstream_19704 | chr15 | 40861491 | 40866893 | + | RPUSD2 | NM | 6 | 2,1 | 2 | -1 | 2,84E-01 | NA | 11 | 1 | 0 | 2 | 9 | 7 | 7 | 8 | 2,3 | 0,1 | 2,3 |
| chr4:101944086-101944586:downstream_23804 | chr4 | 101944586 | 102268628 | - | PPP3CA | NM | 19 | -1,5 | 1 | 1 | 2,84E-01 | NA | 2 | 0 | 10 | 6 | 19 | 15 | 47 | 51 | -3,3 | -1,6 | -1,7 |
| chrX:135043730-135044230:downstream_7952 | chrX | 135044230 | 135056134 | - | MMGT1 | NM | 35 | -0,9 | 1 | 1 | 2,85E-01 | 6,22E-01 | 6 | 7 | 10 | 8 | 84 | 61 | 56 | 52 | -0,5 | 0,4 | -1,0 |
| chr14:74523867-74524367:downstream_19145 | chr14 | 74524367 | 74551196 | - | ALDH6A1 | NM | 6 | -2,0 | 2 | 1 | 2,86E-01 | NA | 1 | 2 | 6 | 6 | 2 | 14 | 10 | 7 | -2,0 | -0,1 | -1,9 |
| chr14:75127454-75127954:downstream_11228 | chr14 | 75127954 | 75179807 | - | AREL1 | NM | 32 | 1,1 | 1 | -1 | 2,86E-01 | 6,23E-01 | 7 | 5 | 8 | 8 | 41 | 15 | 72 | 98 | -0,5 | -1,6 | 1,1 |
| chr19:14198728-14199228:downstream_14063 | chr19 | 14199228 | 14201232 | - | SAMD1 | NM | 19 | 1,1 | 1 | -1 | 2,86E-01 | NA | 28 | 32 | 14 | 27 | 7 | 13 | 11 | 18 | 0,5 | -0,6 | 1,1 |
| chr13:31233686-31234186:downstream_10573 | chr13 | 31191829 | 31233686 | + | USPL1 | NM | 14 | -1,3 | 1 | 1 | 2,86E-01 | NA | 2 | 12 | 8 | 16 | 18 | 23 | 14 | 16 | -0,8 | 0,4 | -1,3 |
| chr11:18388590-18389090:downstream_18043 | chr11 | 18343815 | 18388590 | + | GTF2H1 | NM | 9 | 1,4 | 1 | -1 | 2,87E-01 | NA | 13 | 16 | 6 | 4 | 8 | 12 | 10 | 7 | 1,6 | 0,2 | 1,4 |
| chr21:38887679-38888179:downstream_22652 | chr21 | 38739858 | 38887679 | + | DYRK1A | NM | 13 | -2,0 | 2 | 1 | 2,87E-01 | NA | 0 | 0 | 4 | 7 | 4 | 6 | 28 | 52 | -Inf | -3,0 | -Inf |
| chr15:50999236-50999736:downstream_11722 | chr15 | 50999736 | 51057910 | - | SPPL2A | NM | 73 | -0,6 | 1 | 1 | 2,88E-01 | 6,27E-01 | 47 | 55 | 103 | 115 | 62 | 48 | 78 | 74 | -1,1 | -0,5 | -0,6 |
| chr2:219314473-219314973:downstream_1796 | chr2 | 219314973 | 219433084 | - | USP37 | NM | 7 | 2,2 | 2 | -1 | 2,88E-01 | NA | 0 | 15 | 7 | 9 | 0 | 4 | 10 | 8 | 0,0 | -2,2 | 2,2 |
| chr19:28297688-28298188:downstream_21252 | chr19 | 28284374 | 28297688 | + | LOC101927151 | NR | 19 | -1,1 | 1 | 1 | 2,88E-01 | NA | 35 | 30 | 22 | 25 | 21 | 11 | 7 | 4 | 0,5 | 1,5 | -1,1 |
| chr17:46709351-46709851:downstream_13205 | chr17 | 46709851 | 46709921 | - | MIR196A1 | NR | 7 | 1,7 | 2 | -1 | 2,89E-01 | NA | 5 | 10 | 6 | 2 | 3 | 12 | 17 | 7 | 1,0 | -0,6 | 1,7 |
| chr5:34959069-34959569:downstream_24093 | chr5 | 34929697 | 34959069 | + | DNAJC21 | NM | 17 | 1,2 | 1 | -1 | 2,89E-01 | NA | 37 | 38 | 6 | 11 | 18 | 10 | 6 | 8 | 2,2 | 1,0 | 1,2 |
| chr17:5288845-5289345:downstream_12775 | chr17 | 5289345 | 5323059 | - | NUP88 | NM | 17 | 1,1 | 1 | -1 | 2,89E-01 | NA | 12 | 13 | 19 | 10 | 10 | 13 | 33 | 23 | -0,2 | -1,3 | 1,1 |
| chr2:207027653-207028153:downstream_21890 | chr2 | 207024317 | 207027653 | + | EEF1B2 | NM | 163 | 0,9 | 1 | -1 | 2,90E-01 | 6,29E-01 | 11 | 16 | 4 | 3 | 491 | 366 | 231 | 177 | 1,9 | 1,1 | 0,8 |
| chr12:124082688-124083188:downstream_10493 | chr12 | 124069075 | 124082688 | + | TMED2 | NM | 511 | 0,5 | 1 | -1 | 2,90E-01 | 6,29E-01 | 28 | 31 | 21 | 23 | 986 | 925 | 1066 | 1006 | 0,4 | -0,1 | 0,6 |
| chr12:56399309-56399809:downstream_18858 | chr12 | 56391042 | 56399309 | + | SUOX | NM | 17 | 1,4 | 1 | -1 | 2,90E-01 | NA | 7 | 0 | 3 | 7 | 9 | 17 | 40 | 56 | -0,5 | -1,9 | 1,4 |
| chr4:57259028-57259528:downstream_3445 | chr4 | 57259528 | 57301802 | - | PPAT | NM | 12 | 1,4 | 1 | -1 | 2,90E-01 | NA | 10 | 16 | 15 | 3 | 13 | 7 | 18 | 16 | 0,5 | -0,8 | 1,3 |
| chr2:209100450-209100950:downstream_21975 | chr2 | 209100950 | 209119867 | - | IDH1 | NM | 48 | -0,9 | 1 | 1 | 2,90E-01 | 6,29E-01 | 5 | 5 | 6 | 15 | 94 | 69 | 92 | 97 | -1,1 | -0,2 | -0,8 |
| chr9:95606812-95607312:downstream_26584 | chr9 | 95607312 | 95640320 | - | ZNF484 | NM | 7 | 1,8 | 2 | -1 | 2,90E-01 | NA | 9 | 10 | 7 | 12 | 4 | 0 | 4 | 11 | 0,0 | -1,9 | 1,8 |
| chr17:28853832-28854332:downstream_20499 | chr17 | 28804425 | 28853832 | + | GOSR1 | NM | 10 | -1,8 | 2 | 1 | 2,91E-01 | NA | 1 | 1 | 15 | 11 | 9 | 1 | 21 | 21 | -3,9 | -2,1 | -1,8 |
| chr18:34359487-34359987:downstream_20962 | chr18 | 34359987 | 34409179 | - | TPGS2 | NM | 9 | -1,6 | 2 | 1 | 2,91E-01 | NA | 1 | 3 | 14 | 8 | 5 | 10 | 19 | 9 | -2,6 | -1,0 | -1,6 |
| chr5:122358577-122359077:downstream_4388 | chr5 | 122359077 | 122372425 | - | PPIC | NM | 40 | -0,9 | 1 | 1 | 2,91E-01 | 6,31E-01 | 3 | 9 | 14 | 15 | 67 | 54 | 93 | 66 | -1,3 | -0,4 | -0,9 |
| chr12:22600979-22601479:downstream_18471 | chr12 | 22601479 | 22697480 | - | C2CD5 | NM | 9 | -2,3 | 2 | 1 | 2,92E-01 | NA | 0 | 0 | 4 | 8 | 7 | 0 | 15 | 37 | -Inf | -3,0 | -Inf |
| chr2:74760683-74761183:downstream_21972 | chr2 | 74756531 | 74760683 | + | HTRA2 | NM | 6 | 1,8 | 2 | -1 | 2,93E-01 | NA | 13 | 6 | 3 | 2 | 10 | 3 | 6 | 7 | 1,8 | 0,1 | 1,8 |
| chr16:83846607-83847107:downstream_12497 | chr16 | 83841507 | 83846607 | + | HSBP1 | NM | 27 | 1,3 | 1 | -1 | 2,93E-01 | NA | 10 | 7 | 1 | 2 | 64 | 73 | 22 | 39 | 2,4 | 1,2 | 1,3 |
| chr1:32663886-32664386:downstream_608 | chr1 | 32645344 | 32663886 | + | TXLNA | NM | 10 | 1,5 | 1 | -1 | 2,93E-01 | NA | 14 | 9 | 0 | 6 | 14 | 16 | 17 | 8 | 2,0 | 0,3 | 1,7 |
| chr15:44819429-44819929:downstream_11699 | chr15 | 44719578 | 44819429 | + | CTDSPL2 | NM | 8 | 1,7 | 2 | -1 | 2,94E-01 | NA | 3 | 11 | 11 | 14 | 0 | 4 | 8 | 14 | -0,9 | -2,5 | 1,7 |
| chr7:141464997-141465497:downstream_6119 | chr7 | 141463896 | 141464997 | + | TAS2R3 | NM | 16 | 1,2 | 1 | -1 | 2,94E-01 | NA | 38 | 28 | 13 | 6 | 9 | 19 | 7 | 10 | 1,9 | 0,8 | 1,1 |
| chr1:236372209-236372709:downstream_313 | chr1 | 236305831 | 236372209 | + | GPR137B | NM | 10 | -1,5 | 1 | 1 | 2,95E-01 | NA | 3 | 1 | 6 | 12 | 19 | 6 | 18 | 18 | -2,1 | -0,6 | -1,5 |
| chr1:27273362-27273862:downstream_12 | chr1 | 27248212 | 27273362 | + | NUDC | NM | 5 | 2,0 | 2 | -1 | 2,95E-01 | NA | 4 | 8 | 1 | 3 | 4 | 4 | 3 | 10 | 1,3 | -0,7 | 2,0 |
| chr2:74732192-74732692:downstream_2052 | chr2 | 74729743 | 74732192 | + | LBX2-AS1 | NR | 9 | -1,5 | 1 | 1 | 2,95E-01 | NA | 6 | 4 | 14 | 8 | 5 | 16 | 10 | 7 | -1,2 | 0,3 | -1,5 |
| chr3:172539264-172539764:downstream_23125 | chr3 | 172468474 | 172539264 | + | ECT2 | NM | 39 | 1,1 | 1 | -1 | 2,96E-01 | 6,40E-01 | 7 | 0 | 14 | 9 | 25 | 15 | 107 | 135 | -1,6 | -2,6 | 1,0 |
| chr18:9134343-9134843:downstream_13560 | chr18 | 9102627 | 9134343 | + | NDUFV2 | NM | 13 | 1,4 | 1 | -1 | 2,96E-01 | NA | 3 | 21 | 8 | 6 | 15 | 14 | 14 | 27 | 0,8 | -0,5 | 1,3 |
| chr1:245008359-245008859:downstream_1455 | chr1 | 244998638 | 245008359 | + | COX20 | NM | 13 | 1,4 | 1 | -1 | 2,96E-01 | NA | 10 | 20 | 6 | 2 | 33 | 8 | 10 | 18 | 1,9 | 0,6 | 1,3 |
| chr1:202846909-202847409:downstream_1301 | chr1 | 202847409 | 202858385 | - | RABIF | NM | 7 | -1,8 | 2 | 1 | 2,96E-01 | NA | 4 | 0 | 10 | 7 | 7 | 9 | 7 | 10 | -2,0 | -0,1 | -1,9 |
| chr1:186640443-186640943:downstream_48 | chr1 | 186640943 | 186649559 | - | PTGS2 | NM | 432 | -0,5 | 1 | 1 | 2,97E-01 | 6,40E-01 | 45 | 25 | 60 | 52 | 805 | 768 | 962 | 738 | -0,7 | -0,1 | -0,6 |
| chr7:24733322-24733822:downstream_25638 | chr7 | 24612964 | 24733322 | + | MPP6 | NM | 13 | 1,3 | 1 | -1 | 2,97E-01 | NA | 3 | 8 | 14 | 15 | 2 | 7 | 32 | 27 | -1,4 | -2,7 | 1,3 |
| chr3:189615068-189615568:downstream_23484 | chr3 | 189349215 | 189615068 | + | TP63 | NM | 7 | -2,4 | 2 | 1 | 2,97E-01 | NA | 0 | 0 | 8 | 3 | 0 | 6 | 21 | 16 | -Inf | -2,7 | -Inf |
| chr11:89933096-89933596:downstream_17953 | chr11 | 89933596 | 89956532 | - | CHORDC1 | NM | 58 | 0,7 | 1 | -1 | 2,97E-01 | 6,40E-01 | 59 | 42 | 19 | 15 | 100 | 110 | 70 | 48 | 1,6 | 0,8 | 0,7 |
| chr4:129189891-129190391:downstream_3842 | chr4 | 129190391 | 129209984 | - | PGRMC2 | NM | 15 | 1,2 | 1 | -1 | 2,97E-01 | NA | 10 | 7 | 6 | 11 | 13 | 12 | 39 | 21 | 0,0 | -1,3 | 1,2 |
| chr3:27413711-27414211:downstream_23433 | chr3 | 27414211 | 27525911 | - | SLC4A7 | NM | 18 | -1,4 | 1 | 1 | 2,97E-01 | NA | 2 | 0 | 11 | 21 | 7 | 13 | 38 | 55 | -3,7 | -2,2 | -1,5 |
| chr18:3278282-3278782:downstream_20901 | chr18 | 3262110 | 3278282 | + | MYL12B | NM | 474 | 0,6 | 1 | -1 | 2,98E-01 | 6,40E-01 | 29 | 26 | 4 | 18 | 1096 | 1119 | 768 | 733 | 1,3 | 0,6 | 0,8 |
| chr17:73001892-73002392:downstream_13351 | chr17 | 72983726 | 73001892 | + | CDR2L | NM | 27 | -1,5 | 1 | 1 | 2,98E-01 | NA | 1 | 1 | 8 | 4 | 48 | 13 | 77 | 67 | -2,8 | -1,2 | -1,6 |
| chr9:132589063-132589563:downstream_7359 | chr9 | 132589563 | 132597572 | - | C9orf78 | NM | 52 | 1,1 | 1 | -1 | 2,98E-01 | 6,40E-01 | 7 | 11 | 0 | 3 | 117 | 151 | 58 | 69 | 2,5 | 1,1 | 1,4 |
| chr2:196997806-196998306:downstream_1764 | chr2 | 196998306 | 197036336 | - | STK17B | NM | 5 | 2,3 | 2 | -1 | 2,98E-01 | NA | 7 | 2 | 1 | 0 | 7 | 14 | 6 | 6 | 2,7 | 0,8 | 1,9 |
| chr20:60883918-60884418:downstream_22347 | chr20 | 60877951 | 60883918 | + | ADRM1 | NM | 56 | -1,0 | 1 | 1 | 2,99E-01 | 6,41E-01 | 6 | 4 | 6 | 2 | 143 | 167 | 64 | 57 | 0,3 | 1,4 | -1,1 |
| chr1:207191365-207191865:downstream_16525 | chr1 | 207191865 | 207206101 | - | C1orf116 | NM | 13 | 1,3 | 1 | -1 | 2,99E-01 | NA | 7 | 5 | 3 | 3 | 13 | 22 | 22 | 25 | 0,9 | -0,4 | 1,3 |
| chr7:12370008-12370508:downstream_5713 | chr7 | 12370508 | 12443852 | - | VWDE | NM | 5 | 1,9 | 2 | -1 | 2,99E-01 | NA | 2 | 6 | 7 | 7 | 2 | 1 | 7 | 11 | -0,9 | -2,8 | 2,0 |
| chr6:35438558-35439058:downstream_5097 | chr6 | 35436177 | 35438558 | + | RPL10A | NM | 61 | 0,7 | 1 | -1 | 3,00E-01 | 6,41E-01 | 64 | 69 | 26 | 21 | 89 | 106 | 56 | 53 | 1,5 | 0,8 | 0,7 |
| chr1:110138454-110138954:downstream_34 | chr1 | 110091185 | 110138454 | + | GNAI3 | NM | 7 | 1,6 | 2 | -1 | 3,00E-01 | NA | 5 | 10 | 10 | 8 | 2 | 2 | 7 | 10 | -0,3 | -1,9 | 1,6 |
| chr20:30157370-30157870:downstream_22438 | chr20 | 30102212 | 30157370 | + | HM13 | NM | 110 | -0,9 | 1 | 1 | 3,01E-01 | 6,41E-01 | 6 | 2 | 11 | 5 | 189 | 230 | 192 | 243 | -1,1 | -0,1 | -1,0 |
| chrX:118919966-118920466:downstream_7902 | chrX | 118920466 | 118925622 | - | RPL39 | NM | 29 | 1,1 | 1 | -1 | 3,01E-01 | 6,41E-01 | 14 | 9 | 3 | 3 | 61 | 67 | 29 | 43 | 1,9 | 0,8 | 1,1 |
| chr1:151022946-151023446:downstream_16579 | chr1 | 151023446 | 151032125 | - | CDC42SE1 | NM | 39 | -0,8 | 1 | 1 | 3,01E-01 | 6,41E-01 | 23 | 24 | 22 | 21 | 61 | 83 | 46 | 30 | 0,1 | 0,9 | -0,8 |
| chrX:118671611-118672111:downstream_26778 | chrX | 118672111 | 118699397 | - | CXorf56 | NM | 32 | 1,0 | 1 | -1 | 3,01E-01 | 6,41E-01 | 6 | 10 | 3 | 4 | 61 | 58 | 63 | 52 | 1,2 | 0,1 | 1,1 |
| chr8:67833664-67834164:downstream_6588 | chr8 | 67834164 | 67837777 | - | SNHG6 | NR | 78 | -0,7 | 1 | 1 | 3,01E-01 | 6,41E-01 | 23 | 9 | 17 | 17 | 193 | 147 | 106 | 111 | -0,1 | 0,7 | -0,8 |
| chr16:57850831-57851331:downstream_12390 | chr16 | 57844548 | 57850831 | + | LOC388282 | NM | 13 | -1,4 | 1 | 1 | 3,01E-01 | NA | 7 | 0 | 10 | 23 | 15 | 9 | 25 | 19 | -2,3 | -0,9 | -1,4 |
| chr4:53609183-53609683:downstream_23632 | chr4 | 53609683 | 53617807 | - | ERVMER34-1 | NM | 3 | 3,1 | 3 | -1 | 3,02E-01 | NA | 3 | 6 | 0 | 0 | 9 | 0 | 3 | 1 | Inf | 1,3 | Inf |
| chr8:94753047-94753547:downstream_6639 | chr8 | 94752338 | 94753047 | + | RBM12B-AS1 | NR | 9 | -2,4 | 2 | 1 | 3,02E-01 | NA | 9 | 3 | 1 | 0 | 23 | 39 | 0 | 1 | 3,1 | 6,2 | -3,1 |
| chr12:44121909-44122409:downstream_18775 | chr12 | 44122409 | 44152620 | - | PUS7L | NM | 42 | -0,8 | 1 | 1 | 3,02E-01 | 6,43E-01 | 11 | 12 | 24 | 34 | 56 | 45 | 82 | 73 | -1,4 | -0,6 | -0,8 |
| chr22:36906396-36906896:downstream_15584 | chr22 | 36906896 | 36925277 | - | EIF3D | NM | 9 | -1,5 | 2 | 1 | 3,03E-01 | NA | 4 | 6 | 1 | 6 | 23 | 20 | 3 | 8 | 0,5 | 2,0 | -1,5 |
| chr7:115898837-115899337:downstream_25774 | chr7 | 115850546 | 115898837 | + | TES | NM | 28 | 1,1 | 1 | -1 | 3,03E-01 | NA | 7 | 3 | 1 | 5 | 33 | 46 | 68 | 57 | 0,6 | -0,7 | 1,2 |
| chr1:46653852-46654352:downstream_17098 | chr1 | 46654352 | 46685977 | - | POMGNT1 | NM | 12 | 1,5 | 1 | -1 | 3,03E-01 | NA | 9 | 2 | 6 | 2 | 20 | 7 | 29 | 18 | 0,6 | -0,8 | 1,4 |
| chr11:2949002-2949502:downstream_9021 | chr11 | 2949502 | 2950650 | - | PHLDA2 | NM | 5 | -2,2 | 2 | 1 | 3,04E-01 | NA | 7 | 13 | 6 | 2 | 8 | 5 | 1 | 0 | 1,3 | 3,2 | -1,9 |
| chr11:8997830-8998330:downstream_9134 | chr11 | 8986221 | 8997830 | + | TMEM9B-AS1 | NR | 6 | 1,6 | 2 | -1 | 3,04E-01 | NA | 9 | 9 | 4 | 6 | 4 | 5 | 10 | 6 | 0,8 | -0,8 | 1,6 |
| chr2:136542633-136543133:downstream_2264 | chr2 | 136499188 | 136542633 | + | UBXN4 | NM | 47 | 1,0 | 1 | -1 | 3,05E-01 | 6,47E-01 | 5 | 8 | 10 | 3 | 52 | 72 | 88 | 141 | 0,0 | -0,9 | 0,8 |
| chr19:52536176-52536676:downstream_14549 | chr19 | 52536676 | 52552073 | - | ZNF432 | NM | 17 | 1,3 | 1 | -1 | 3,05E-01 | NA | 19 | 16 | 45 | 19 | 7 | 0 | 22 | 11 | -0,9 | -2,2 | 1,4 |
| chr15:57177867-57178367:downstream_11733 | chr15 | 57178367 | 57210697 | - | LOC145783 | NR | 6 | 1,9 | 2 | -1 | 3,05E-01 | NA | 7 | 9 | 4 | 2 | 7 | 3 | 13 | 1 | 1,5 | -0,5 | 2,0 |
| chr20:62567384-62567884:downstream_15164 | chr20 | 62526454 | 62567384 | + | DNAJC5 | NM | 22 | 1,2 | 1 | -1 | 3,05E-01 | NA | 8 | 5 | 1 | 4 | 29 | 47 | 47 | 32 | 1,2 | 0,0 | 1,3 |
| chr1:156261977-156262477:downstream_16771 | chr1 | 156262477 | 156265480 | - | GLMP | NM | 30 | 0,9 | 1 | -1 | 3,06E-01 | 6,48E-01 | 28 | 22 | 15 | 11 | 46 | 38 | 49 | 32 | 0,9 | 0,0 | 0,9 |
| chr13:51102779-51103279:downstream_18971 | chr13 | 50656304 | 51102779 | + | DLEU1 | NR | 8 | -1,7 | 2 | 1 | 3,06E-01 | NA | 3 | 0 | 17 | 16 | 2 | 5 | 13 | 9 | -3,3 | -1,6 | -1,7 |
| chr5:68426899-68427399:downstream_24367 | chr5 | 68389775 | 68426899 | + | SLC30A5 | NM | 11 | 1,5 | 1 | -1 | 3,06E-01 | NA | 9 | 4 | 0 | 11 | 12 | 5 | 21 | 26 | 0,2 | -1,5 | 1,6 |
| chr20:37048738-37049238:downstream_22565 | chr20 | 37049238 | 37064018 | - | SNHG17 | NR | 36 | 0,8 | 1 | -1 | 3,07E-01 | 6,49E-01 | 52 | 40 | 11 | 20 | 52 | 52 | 33 | 30 | 1,6 | 0,7 | 0,8 |
| chr2:20096013-20096513:downstream_1875 | chr2 | 20096513 | 20101744 | - | TTC32 | NM | 28 | 1,0 | 1 | -1 | 3,07E-01 | NA | 54 | 35 | 11 | 10 | 37 | 40 | 8 | 25 | 2,1 | 1,2 | 0,9 |
| chr4:120217073-120217573:downstream_23578 | chr4 | 120217573 | 120225600 | - | C4orf3 | NM | 5 | -2,2 | 2 | 1 | 3,07E-01 | NA | 5 | 6 | 4 | 0 | 11 | 15 | 0 | 2 | 1,4 | 4,0 | -2,6 |
| chr11:94149968-94150468:downstream_18134 | chr11 | 94150468 | 94227040 | - | MRE11A | NM | 4 | 2,2 | 2 | -1 | 3,08E-01 | NA | 7 | 1 | 4 | 4 | 3 | 0 | 3 | 12 | 0,0 | -2,2 | 2,2 |
| chr9:34343711-34344211:downstream_26421 | chr9 | 34329503 | 34343711 | + | NUDT2 | NM | 19 | -1,3 | 1 | 1 | 3,08E-01 | NA | 2 | 0 | 10 | 8 | 17 | 18 | 50 | 47 | -2,9 | -1,4 | -1,4 |
| chr21:45182188-45182688:downstream_15349 | chr21 | 45138977 | 45182188 | + | PDXK | NM | 11 | -1,6 | 2 | 1 | 3,08E-01 | NA | 2 | 0 | 7 | 7 | 9 | 11 | 26 | 26 | -3,1 | -1,4 | -1,7 |
| chr22:46726707-46727207:downstream_15686 | chr22 | 46692637 | 46726707 | + | GTSE1 | NM | 5 | -2,0 | 2 | 1 | 3,08E-01 | NA | 2 | 0 | 7 | 9 | 2 | 5 | 6 | 11 | -3,3 | -1,2 | -2,1 |
| chr2:61704568-61705068:downstream_1628 | chr2 | 61705068 | 61765418 | - | XPO1 | NM | 30 | 1,1 | 1 | -1 | 3,08E-01 | 6,51E-01 | 2 | 6 | 1 | 9 | 27 | 24 | 81 | 94 | -0,5 | -1,8 | 1,3 |
| chr17:26934481-26934981:downstream_12930 | chr17 | 26934981 | 26941211 | - | SGK494 | NM | 13 | 1,4 | 1 | -1 | 3,08E-01 | NA | 5 | 15 | 10 | 12 | 14 | 2 | 19 | 25 | -0,1 | -1,5 | 1,4 |
| chr7:44604515-44605015:downstream_25481 | chr7 | 44605015 | 44614137 | - | DDX56 | NM | 29 | 0,9 | 1 | -1 | 3,09E-01 | 6,52E-01 | 54 | 23 | 17 | 12 | 34 | 43 | 24 | 29 | 1,4 | 0,6 | 0,9 |
| chr3:40503863-40504363:downstream_23392 | chr3 | 40498782 | 40503863 | + | RPL14 | NM | 118 | 0,7 | 1 | -1 | 3,09E-01 | 6,52E-01 | 13 | 20 | 7 | 7 | 277 | 261 | 180 | 180 | 1,3 | 0,6 | 0,7 |
| chr10:85913311-85913811:downstream_8602 | chr10 | 85899184 | 85913311 | + | GHITM | NM | 16 | 1,1 | 1 | -1 | 3,10E-01 | NA | 32 | 18 | 7 | 9 | 22 | 15 | 7 | 19 | 1,6 | 0,5 | 1,1 |
| chr9:7795990-7796490:downstream_6982 | chr9 | 7796490 | 7799799 | - | TMEM261 | NM | 35 | 0,8 | 1 | -1 | 3,10E-01 | 6,53E-01 | 60 | 54 | 8 | 11 | 69 | 44 | 15 | 21 | 2,5 | 1,7 | 0,9 |
| chr2:198573114-198573614:downstream_2367 | chr2 | 198570027 | 198573114 | + | mars-02 | NM | 15 | 1,2 | 1 | -1 | 3,10E-01 | NA | 15 | 33 | 7 | 11 | 18 | 10 | 7 | 17 | 1,4 | 0,2 | 1,2 |
| chr2:96939573-96940073:downstream_2119 | chr2 | 96940073 | 96971307 | - | SNRNP200 | NM | 11 | 1,6 | 2 | -1 | 3,10E-01 | NA | 7 | 4 | 3 | 0 | 22 | 22 | 14 | 13 | 2,0 | 0,7 | 1,3 |
| chr1:27205372-27205872:downstream_578 | chr1 | 27205872 | 27216869 | - | GPN2 | NM | 23 | 1,0 | 1 | -1 | 3,10E-01 | NA | 15 | 44 | 11 | 9 | 32 | 29 | 22 | 20 | 1,6 | 0,5 | 1,0 |
| chr6:170881958-170882458:downstream_25295 | chr6 | 170863420 | 170881958 | + | TBP | NM | 8 | 1,6 | 2 | -1 | 3,11E-01 | NA | 6 | 8 | 1 | 2 | 15 | 14 | 11 | 9 | 2,1 | 0,5 | 1,6 |
| chr6:37143204-37143704:downstream_25061 | chr6 | 37137921 | 37143204 | + | PIM1 | NM | 28 | -1,4 | 1 | 1 | 3,11E-01 | NA | 0 | 2 | 7 | 7 | 20 | 30 | 75 | 80 | -2,9 | -1,6 | -1,3 |
| chr4:6677774-6678274:downstream_3593 | chr4 | 6675820 | 6677774 | + | LOC93622 | NR | 19 | 1,2 | 1 | -1 | 3,11E-01 | NA | 21 | 11 | 0 | 11 | 28 | 30 | 24 | 25 | 1,6 | 0,3 | 1,3 |
| chr11:65484643-65485143:downstream_9446 | chr11 | 65485143 | 65488409 | - | RNASEH2C | NM | 5 | 1,9 | 2 | -1 | 3,11E-01 | NA | 8 | 2 | 1 | 2 | 10 | 2 | 8 | 9 | 1,4 | -0,6 | 2,0 |
| chr3:186838240-186838740:downstream_3315 | chr3 | 186838740 | 186857263 | - | RPL39L | NM | 12 | -1,5 | 1 | 1 | 3,11E-01 | NA | 4 | 0 | 11 | 3 | 19 | 17 | 26 | 16 | -1,8 | -0,2 | -1,6 |
| chr6:160873611-160874111:downstream_4826 | chr6 | 160769404 | 160873611 | + | SLC22A3 | NM | 92 | -0,6 | 1 | 1 | 3,12E-01 | 6,55E-01 | 15 | 11 | 39 | 40 | 110 | 105 | 237 | 180 | -1,6 | -1,0 | -0,6 |
| chr8:1734736-1735236:downstream_6377 | chr8 | 1711869 | 1734736 | + | CLN8 | NM | 11 | 1,5 | 1 | -1 | 3,12E-01 | NA | 23 | 12 | 1 | 2 | 20 | 23 | 7 | 4 | 3,5 | 2,0 | 1,6 |
| chr1:109941152-109941652:downstream_17130 | chr1 | 109941652 | 109969108 | - | PSMA5 | NM | 5 | 2,4 | 2 | -1 | 3,12E-01 | NA | 8 | 0 | 1 | 0 | 9 | 11 | 4 | 9 | 2,5 | 0,6 | 2,0 |
| chr1:162569633-162570133:downstream_1186 | chr1 | 162531295 | 162569633 | + | UAP1 | NM | 18 | 1,2 | 1 | -1 | 3,12E-01 | NA | 8 | 2 | 6 | 3 | 28 | 14 | 38 | 43 | 0,2 | -1,0 | 1,1 |
| chr6:26095469-26095969:downstream_24795 | chr6 | 26087508 | 26095469 | + | HFE | NM | 57 | -0,7 | 1 | 1 | 3,13E-01 | 6,55E-01 | 19 | 14 | 22 | 35 | 90 | 85 | 89 | 105 | -0,8 | -0,1 | -0,7 |
| chr5:172194592-172195092:downstream_4580 | chr5 | 172195092 | 172198203 | - | DUSP1 | NM | 46 | 0,9 | 1 | -1 | 3,13E-01 | 6,55E-01 | 11 | 36 | 11 | 17 | 62 | 75 | 70 | 86 | 0,7 | -0,2 | 0,9 |
| chr1:9211226-9211726:downstream_422 | chr1 | 9211726 | 9211836 | - | MIR34A | NR | 24 | -1,1 | 1 | 1 | 3,13E-01 | NA | 54 | 37 | 49 | 36 | 11 | 3 | 3 | 3 | 0,1 | 1,2 | -1,1 |
| chr19:44741421-44741921:downstream_21619 | chr19 | 44716680 | 44741421 | + | ZNF227 | NM | 26 | 0,9 | 1 | -1 | 3,13E-01 | NA | 9 | 15 | 14 | 6 | 32 | 33 | 46 | 54 | 0,3 | -0,6 | 0,9 |
| chr12:69973571-69974071:downstream_18571 | chr12 | 69864128 | 69973571 | + | FRS2 | NM | 9 | 1,6 | 2 | -1 | 3,13E-01 | NA | 11 | 10 | 24 | 11 | 2 | 0 | 7 | 7 | -0,8 | -2,5 | 1,7 |
| chr19:11040916-11041416:downstream_14003 | chr19 | 11039423 | 11040916 | + | C19orf52 | NM | 19 | 1,2 | 1 | -1 | 3,13E-01 | NA | 43 | 12 | 4 | 6 | 33 | 28 | 11 | 14 | 2,5 | 1,3 | 1,2 |
| chr6:26986644-26987144:downstream_4935 | chr6 | 26987144 | 26988085 | - | LOC100270746 | NR | 17 | 1,2 | 1 | -1 | 3,13E-01 | NA | 28 | 41 | 13 | 3 | 15 | 21 | 13 | 6 | 2,1 | 1,0 | 1,1 |
| chr10:102032534-102033034:downstream_17492 | chr10 | 102033034 | 102046469 | - | BLOC1S2 | NM | 7 | 2,0 | 2 | -1 | 3,14E-01 | NA | 9 | 20 | 3 | 0 | 7 | 14 | 0 | 6 | 3,4 | 1,8 | 1,6 |
| chr15:75246942-75247442:downstream_11811 | chr15 | 75247442 | 75249775 | - | RPP25 | NM | 10 | -1,3 | 1 | 1 | 3,15E-01 | NA | 5 | 9 | 10 | 7 | 19 | 17 | 4 | 12 | -0,3 | 1,1 | -1,4 |
| chr17:47006422-47006922:downstream_13212 | chr17 | 46985730 | 47006422 | + | UBE2Z | NM | 61 | 0,7 | 1 | -1 | 3,15E-01 | 6,57E-01 | 41 | 28 | 17 | 49 | 63 | 69 | 95 | 123 | 0,1 | -0,7 | 0,8 |
| chr19:39221171-39221671:downstream_13771 | chr19 | 39138266 | 39221171 | + | ACTN4 | NM | 204 | -0,9 | 1 | 1 | 3,15E-01 | 6,57E-01 | 0 | 6 | 4 | 6 | 382 | 427 | 347 | 463 | -0,8 | 0,0 | -0,8 |
| chr15:42834219-42834719:downstream_11686 | chr15 | 42834719 | 42841002 | - | LRRC57 | NM | 9 | 1,5 | 1 | -1 | 3,15E-01 | NA | 8 | 5 | 3 | 2 | 16 | 12 | 10 | 16 | 1,6 | 0,1 | 1,4 |
| chr11:66431964-66432464:downstream_18251 | chr11 | 66432464 | 66445392 | - | RBM4B | NM | 11 | 1,7 | 2 | -1 | 3,15E-01 | NA | 0 | 11 | 1 | 2 | 11 | 25 | 19 | 16 | 1,8 | 0,0 | 1,8 |
| chr15:25233379-25233879:downstream_11561 | chr15 | 25230006 | 25233379 | + | PWAR5 | NR | 42 | -0,8 | 1 | 1 | 3,15E-01 | 6,57E-01 | 24 | 29 | 60 | 48 | 25 | 54 | 56 | 37 | -1,0 | -0,2 | -0,8 |
| chr9:97223202-97223702:downstream_7200 | chr9 | 97136832 | 97223202 | + | HIATL1 | NM | 30 | 1,0 | 1 | -1 | 3,15E-01 | 6,57E-01 | 7 | 4 | 6 | 7 | 34 | 33 | 61 | 91 | -0,2 | -1,2 | 1,0 |
| chr6:42907008-42907508:downstream_5148 | chr6 | 42896859 | 42907008 | + | CNPY3 | NM | 27 | -1,0 | 1 | 1 | 3,15E-01 | NA | 4 | 5 | 3 | 13 | 41 | 57 | 45 | 47 | -0,8 | 0,1 | -0,9 |
| chr1:180256851-180257351:downstream_44 | chr1 | 180257351 | 180472022 | - | ACBD6 | NM | 9 | -1,5 | 2 | 1 | 3,15E-01 | NA | 8 | 0 | 17 | 21 | 7 | 5 | 11 | 7 | -2,2 | -0,6 | -1,6 |
| chr19:57805436-57805936:downstream_13807 | chr19 | 57791852 | 57805436 | + | ZNF460 | NM | 106 | -0,6 | 1 | 1 | 3,15E-01 | 6,57E-01 | 77 | 70 | 79 | 69 | 139 | 191 | 117 | 108 | 0,0 | 0,5 | -0,6 |
| chr3:151546276-151546776:downstream_3240 | chr3 | 151531860 | 151546276 | + | AADAC | NM | 22 | 1,1 | 1 | -1 | 3,16E-01 | NA | 9 | 12 | 10 | 7 | 16 | 37 | 60 | 27 | 0,3 | -0,7 | 1,0 |
| chr13:37582950-37583450:downstream_19102 | chr13 | 37583450 | 37633850 | - | SUPT20H | NM | 13 | 1,2 | 1 | -1 | 3,16E-01 | NA | 7 | 6 | 6 | 7 | 16 | 9 | 17 | 38 | 0,1 | -1,1 | 1,2 |
| chr2:120023227-120023727:downstream_22261 | chr2 | 119981383 | 120023227 | + | STEAP3 | NM | 28 | -1,2 | 1 | 1 | 3,16E-01 | 6,58E-01 | 3 | 0 | 3 | 10 | 45 | 33 | 56 | 75 | -2,0 | -0,8 | -1,2 |
| chr14:62213256-62213756:downstream_11173 | chr14 | 62213756 | 62215807 | - | HIF1A-AS2 | NR | 60 | -0,7 | 1 | 1 | 3,17E-01 | 6,58E-01 | 36 | 38 | 45 | 17 | 120 | 109 | 64 | 50 | 0,3 | 1,0 | -0,8 |
| chr8:100889722-100890222:downstream_6660 | chr8 | 100890222 | 100906242 | - | COX6C | NM | 18 | 1,5 | 2 | -1 | 3,17E-01 | NA | 8 | 4 | 3 | 0 | 63 | 24 | 13 | 33 | 2,1 | 0,9 | 1,2 |
| chr3:185864489-185864989:downstream_23111 | chr3 | 185864989 | 186080023 | - | DGKG | NM | 7 | 1,8 | 2 | -1 | 3,17E-01 | NA | 7 | 4 | 19 | 11 | 2 | 0 | 10 | 7 | -1,5 | -3,4 | 1,9 |
| chr17:55915786-55916286:downstream_13258 | chr17 | 55916286 | 55927433 | - | MRPS23 | NM | 38 | 0,8 | 1 | -1 | 3,17E-01 | 6,58E-01 | 55 | 69 | 25 | 18 | 42 | 46 | 22 | 29 | 1,5 | 0,8 | 0,7 |
| chr14:92433742-92434242:downstream_11271 | chr14 | 92434242 | 92506403 | - | TRIP11 | NM | 7 | -1,9 | 2 | 1 | 3,17E-01 | NA | 1 | 1 | 8 | 16 | 3 | 2 | 13 | 8 | -3,8 | -2,0 | -1,8 |
| chr8:76479061-76479561:downstream_6609 | chr8 | 76452202 | 76479061 | + | HNF4G | NM | 5 | -2,5 | 3 | 1 | 3,18E-01 | NA | 0 | 0 | 7 | 7 | 2 | 0 | 6 | 14 | -Inf | -3,0 | -Inf |

| | | | | | | | | | | | | | | | | | | | | | | | |
|---|---|---|---|---|---|---|---|---|---|---|---|---|---|---|---|---|---|---|---|---|---|---|---|
| chr9:35681489-35681989:downstream_26550 | chr9 | 35681989 | 35690053 | - | TPM2 | NM | 11 | 1,4 | 1 | -1 | 3,18E-01 | NA | 5 | 8 | 3 | 2 | 15 | 18 | 11 | 25 | 1,3 | -0,1 | 1,3 |
| chr5:132210570-132211070:downstream_4087 | chr5 | 132211070 | 132299354 | - | AFF4 | NM | 57 | 0,9 | 1 | -1 | 3,19E-01 | 6,60E-01 | 2 | 4 | 8 | 10 | 37 | 28 | 210 | 153 | -1,5 | -2,5 | 1,0 |
| chr19:49519182-49519682:downstream_14462 | chr19 | 49497155 | 49519182 | + | RUVBL2 | NM | 18 | 1,3 | 1 | -1 | 3,19E-01 | NA | 4 | 9 | 3 | 1 | 50 | 29 | 22 | 28 | 1,8 | 0,6 | 1,2 |
| chr15:78575077-78575577:downstream_19842 | chr15 | 78575577 | 78592068 | - | WDR61 | NM | 15 | 1,2 | 1 | -1 | 3,20E-01 | NA | 24 | 21 | 3 | 9 | 14 | 25 | 7 | 16 | 2,0 | 0,7 | 1,2 |
| chr11:120960354-120960854:downstream_18333 | chr11 | 120894802 | 120960354 | + | TBCEL | NM | 16 | 1,2 | 1 | -1 | 3,20E-01 | NA | 11 | 3 | 13 | 16 | 8 | 6 | 43 | 25 | -1,1 | -2,3 | 1,2 |
| chr13:31708610-31709110:downstream_18993 | chr13 | 31709110 | 31736525 | - | HSPH1 | NM | 18 | 1,1 | 1 | -1 | 3,20E-01 | NA | 43 | 39 | 15 | 18 | 10 | 8 | 3 | 11 | 1,3 | 0,3 | 1,0 |
| chr3:148747403-148747903:downstream_23193 | chr3 | 148747903 | 148804341 | - | HLTF | NM | 22 | 1,1 | 1 | -1 | 3,21E-01 | NA | 5 | 7 | 13 | 5 | 13 | 23 | 40 | 68 | -0,6 | -1,6 | 1,0 |
| chr3:45054158-45054658:downstream_2911 | chr3 | 45017740 | 45054158 | + | EXOSC7 | NR | 7 | -1,5 | 2 | 1 | 3,21E-01 | NA | 7 | 8 | 7 | 6 | 17 | 8 | 1 | 6 | 0,2 | 1,8 | -1,6 |
| chr19:50161906-50162406:downstream_14494 | chr19 | 50145381 | 50161906 | + | SCAF1 | NM | 5 | 2,3 | 2 | -1 | 3,21E-01 | NA | 10 | 1 | 0 | 1 | 11 | 9 | 1 | 7 | 3,7 | 1,1 | 2,6 |
| chr11:94695286-94695786:downstream_9593 | chr11 | 94695786 | 94706776 | - | CWC15 | NM | 14 | 1,6 | 2 | -1 | 3,21E-01 | NA | 8 | 20 | 3 | 0 | 28 | 36 | 13 | 3 | 3,4 | 2,0 | 1,3 |
| chr14:35743284-35743784:downstream_19269 | chr14 | 35591526 | 35743284 | + | KIAA0391 | NM | 32 | -1,1 | 1 | 1 | 3,21E-01 | 6,63E-01 | 0 | 4 | 7 | 16 | 28 | 27 | 103 | 67 | -2,6 | -1,6 | -1,0 |
| chr11:94865815-94866315:downstream_9599 | chr11 | 94822973 | 94865815 | + | ENDOD1 | NM | 63 | -0,9 | 1 | 1 | 3,21E-01 | 6,63E-01 | 5 | 1 | 7 | 7 | 103 | 112 | 114 | 154 | -1,3 | -0,3 | -1,0 |
| chr14:52115735-52116235:downstream_11137 | chr14 | 52116235 | 52118462 | - | FRMD6-AS1 | NR | 12 | 1,3 | 1 | -1 | 3,21E-01 | NA | 14 | 16 | 6 | 4 | 24 | 8 | 10 | 14 | 1,6 | 0,4 | 1,2 |
| chr5:150649953-150650453:downstream_24147 | chr5 | 150632612 | 150649953 | + | GM2A | NM | 25 | -0,9 | 1 | 1 | 3,22E-01 | NA | 20 | 20 | 38 | 24 | 25 | 29 | 31 | 15 | -0,6 | 0,3 | -0,9 |
| chr11:77032559-77033059:downstream_18190 | chr11 | 77033059 | 77185108 | - | PAK1 | NM | 17 | -1,5 | 2 | 1 | 3,22E-01 | NA | 2 | 0 | 7 | 4 | 14 | 25 | 50 | 35 | -2,8 | -1,1 | -1,6 |
| chr10:30598229-30598729:downstream_8465 | chr10 | 30598729 | 30638267 | - | MTPAP | NM | 14 | 1,2 | 1 | -1 | 3,22E-01 | NA | 25 | 13 | 21 | 7 | 13 | 5 | 18 | 12 | 0,4 | -0,8 | 1,2 |
| chr12:69995357-69995857:downstream_18488 | chr12 | 69979207 | 69995357 | + | CCT2 | NM | 36 | 1,0 | 1 | -1 | 3,22E-01 | 6,64E-01 | 3 | 10 | 1 | 6 | 62 | 56 | 63 | 84 | 0,9 | -0,3 | 1,2 |
| chr21:34875737-34876237:downstream_22659 | chr21 | 34876237 | 34915198 | - | GART | NM | 43 | 0,7 | 1 | -1 | 3,23E-01 | 6,64E-01 | 16 | 13 | 19 | 21 | 46 | 34 | 97 | 94 | -0,5 | -1,3 | 0,8 |
| chr2:239360891-239361391:downstream_1818 | chr2 | 239335625 | 239360891 | + | ASB1 | NM | 7 | 1,6 | 2 | -1 | 3,23E-01 | NA | 6 | 14 | 0 | 4 | 12 | 9 | 6 | 10 | 2,2 | 0,4 | 1,8 |
| chr9:77597372-77597872:downstream_7142 | chr9 | 77597872 | 77643310 | - | C9orf41 | NM | 72 | -0,6 | 1 | 1 | 3,23E-01 | 6,64E-01 | 35 | 58 | 77 | 99 | 60 | 77 | 77 | 92 | -0,9 | -0,3 | -0,6 |
| chrX:77384744-77385244:downstream_7819 | chrX | 77385244 | 77395179 | - | TAF9B | NM | 43 | -0,8 | 1 | 1 | 3,23E-01 | 6,64E-01 | 20 | 48 | 46 | 34 | 47 | 68 | 40 | 38 | -0,2 | 0,6 | -0,8 |
| chr16:22095972-22096472:downstream_11935 | chr16 | 22019455 | 22095972 | + | C16orf52 | NM | 4 | 2,2 | 2 | -1 | 3,23E-01 | NA | 3 | 3 | 3 | 7 | 2 | 0 | 8 | 4 | -0,6 | -2,9 | 2,3 |
| chr3:37408370-37408870:downstream_23173 | chr3 | 37284681 | 37408370 | + | GOLGA4 | NM | 7 | -1,7 | 2 | 1 | 3,24E-01 | NA | 2 | 0 | 10 | 8 | 2 | 10 | 14 | 11 | -2,9 | -1,1 | -1,8 |
| chr10:71909460-71909960:downstream_17758 | chr10 | 71909960 | 71930285 | - | SAR1A | NM | 34 | 0,9 | 1 | -1 | 3,24E-01 | 6,65E-01 | 28 | 11 | 8 | 13 | 60 | 46 | 45 | 64 | 0,9 | 0,0 | 0,9 |
| chr13:79892502-79893002:downstream_19075 | chr13 | 79893002 | 79980356 | - | RBM26 | NM | 12 | -1,5 | 2 | 1 | 3,25E-01 | NA | 0 | 4 | 19 | 11 | 3 | 13 | 18 | 27 | -3,0 | -1,5 | -1,4 |
| chr19:18390003-18390503:downstream_21200 | chr19 | 18390503 | 18392466 | - | JUND | NM | 23 | 1,0 | 1 | -1 | 3,25E-01 | NA | 5 | 7 | 4 | 7 | 20 | 35 | 43 | 63 | 0,1 | -1,0 | 1,1 |
| chr17:647160-647660:downstream_12581 | chr17 | 647660 | 655501 | - | GEMIN4 | NM | 13 | 1,2 | 1 | -1 | 3,25E-01 | NA | 17 | 14 | 3 | 5 | 24 | 16 | 11 | 13 | 2,0 | 0,8 | 1,2 |
| chr2:37331783-37332283:downstream_21894 | chr2 | 37332283 | 37384190 | - | EIF2AK2 | NM | 62 | -0,7 | 1 | 1 | 3,25E-01 | 6,66E-01 | 21 | 23 | 127 | 62 | 35 | 38 | 93 | 96 | -2,1 | -1,4 | -0,7 |
| chr17:73232186-73232686:downstream_20490 | chr17 | 73232686 | 73258474 | - | GGA3 | NM | 7 | 1,6 | 2 | -1 | 3,26E-01 | NA | 5 | 6 | 1 | 2 | 14 | 6 | 8 | 14 | 1,5 | -0,2 | 1,7 |
| chr8:86125787-86126287:downstream_25868 | chr8 | 86126287 | 86132651 | - | C8orf59 | NM | 29 | -0,9 | 1 | 1 | 3,26E-01 | 6,67E-01 | 24 | 40 | 18 | 15 | 61 | 42 | 21 | 8 | 1,0 | 1,8 | -0,9 |
| chr2:231914427-231914927:downstream_2472 | chr2 | 231902280 | 231914427 | + | C2orf72 | NM | 16 | -1,2 | 1 | 1 | 3,26E-01 | NA | 2 | 2 | 25 | 16 | 8 | 8 | 36 | 29 | -3,2 | -2,0 | -1,2 |
| chr5:32604185-32604685:downstream_4212 | chr5 | 32585604 | 32604185 | + | SUB1 | NM | 12 | 1,2 | 1 | -1 | 3,26E-01 | NA | 18 | 17 | 11 | 4 | 18 | 7 | 13 | 11 | 1,2 | 0,0 | 1,2 |
| chr10:18833635-18834135:downstream_8203 | chr10 | 18834135 | 18940566 | - | NSUN6 | NM | 5 | -1,7 | 2 | 1 | 3,27E-01 | NA | 2 | 4 | 8 | 9 | 3 | 7 | 3 | 7 | -1,7 | 0,1 | -1,8 |
| chr12:53346685-53347185:downstream_18623 | chr12 | 53342654 | 53346685 | + | KRT18 | NM | 254 | 0,6 | 1 | -1 | 3,27E-01 | 6,68E-01 | 17 | 15 | 25 | 35 | 210 | 290 | 637 | 799 | -0,9 | -1,5 | 0,6 |
| chr20:34633039-34633539:downstream_14998 | chr20 | 34633539 | 34638882 | - | LINC00657 | NR | 90 | 0,8 | 1 | -1 | 3,27E-01 | 6,68E-01 | 31 | 12 | 4 | 5 | 197 | 289 | 72 | 112 | 2,2 | 1,4 | 0,8 |
| chr7:72307978-72308478:downstream_25716 | chr7 | 72299951 | 72307978 | + | SBDSP1 | NR | 26 | 0,9 | 1 | -1 | 3,27E-01 | NA | 30 | 35 | 7 | 7 | 36 | 60 | 13 | 25 | 2,3 | 1,4 | 0,9 |
| chrX:70521018-70521518:downstream_26944 | chrX | 70503041 | 70521018 | + | NONO | NM | 107 | 0,7 | 1 | -1 | 3,28E-01 | 6,68E-01 | 21 | 14 | 11 | 9 | 173 | 247 | 164 | 217 | 0,8 | 0,1 | 0,6 |
| chr19:44353050-44353550:downstream_21628 | chr19 | 44331443 | 44353050 | + | ZNF283 | NM | 16 | 1,1 | 1 | -1 | 3,28E-01 | NA | 8 | 22 | 6 | 7 | 15 | 32 | 18 | 23 | 1,3 | 0,2 | 1,1 |
| chr7:37991542-37992042:downstream_25502 | chr7 | 37960162 | 37991542 | + | EPDR1 | NM | 136 | -0,7 | 1 | 1 | 3,28E-01 | 6,68E-01 | 7 | 19 | 13 | 33 | 214 | 267 | 289 | 247 | -0,8 | -0,2 | -0,6 |
| chr2:189896140-189896640:downstream_1759 | chr2 | 189896640 | 190044605 | - | COL5A2 | NM | 9 | 1,7 | 2 | -1 | 3,29E-01 | NA | 9 | 5 | 19 | 19 | 2 | 0 | 10 | 6 | -1,5 | -3,3 | 1,8 |
| chr3:14983785-14984285:downstream_23150 | chr3 | 14984285 | 14989948 | - | FGD5-AS1 | NR | 31 | 0,9 | 1 | -1 | 3,29E-01 | 6,68E-01 | 4 | 9 | 6 | 5 | 45 | 41 | 65 | 71 | 0,3 | -0,7 | 1,0 |
| chr15:58886902-58887402:downstream_11480 | chr15 | 58887402 | 59042177 | - | ADAM10 | NM | 78 | -1,0 | 1 | 1 | 3,30E-01 | 6,69E-01 | 3 | 0 | 10 | 16 | 68 | 57 | 188 | 285 | -3,0 | -1,9 | -1,0 |
| chr18:48611411-48611911:downstream_13628 | chr18 | 48556582 | 48611411 | + | SMAD4 | NM | 11 | -1,4 | 1 | 1 | 3,30E-01 | NA | 4 | 1 | 10 | 4 | 13 | 23 | 25 | 11 | -1,5 | 0,0 | -1,4 |
| chr1:16786584-16787084:downstream_17014 | chr1 | 16767166 | 16786584 | + | NECAP2 | NM | 20 | 1,1 | 1 | -1 | 3,30E-01 | NA | 5 | 4 | 3 | 4 | 33 | 19 | 51 | 42 | 0,3 | -0,8 | 1,2 |
| chr7:7288280-7288780:downstream_5708 | chr7 | 7222245 | 7288280 | + | C1GALT1 | NM | 6 | 1,8 | 2 | -1 | 3,30E-01 | NA | 4 | 9 | 8 | 2 | 0 | 6 | 8 | 8 | 0,2 | -1,5 | 1,8 |
| chr11:34124157-34124657:downstream_17922 | chr11 | 34073229 | 34124157 | + | CAPRIN1 | NM | 17 | 1,1 | 1 | -1 | 3,30E-01 | NA | 33 | 19 | 33 | 21 | 6 | 2 | 10 | 8 | -0,1 | -1,2 | 1,2 |
| chr16:2204523-2205023:downstream_12108 | chr16 | 2205023 | 2205106 | - | SNORD60 | NR | 90 | -0,6 | 1 | 1 | 3,31E-01 | 6,70E-01 | 55 | 72 | 45 | 51 | 154 | 180 | 68 | 99 | 0,4 | 1,0 | -0,6 |
| chr3:179053323-179053823:downstream_23528 | chr3 | 179040778 | 179053323 | + | ZNF639 | NM | 10 | -1,3 | 1 | 1 | 3,31E-01 | NA | 3 | 6 | 7 | 7 | 19 | 14 | 14 | 7 | -0,7 | 0,6 | -1,3 |
| chr7:45139198-45139698:downstream_25769 | chr7 | 45139698 | 45151346 | - | TBRG4 | NM | 9 | 1,6 | 2 | -1 | 3,31E-01 | NA | 10 | 3 | 0 | 3 | 21 | 7 | 11 | 14 | 1,9 | 0,2 | 1,8 |
| chr1:43849078-43849578:downstream_16946 | chr1 | 43849578 | 43855483 | - | MED8 | NM | 13 | 1,2 | 1 | -1 | 3,31E-01 | NA | 6 | 4 | 4 | 3 | 16 | 16 | 24 | 34 | 0,4 | -0,8 | 1,2 |
| chr3:88207115-88207615:downstream_3053 | chr3 | 88198892 | 88207115 | + | C3orf38 | NM | 42 | 0,7 | 1 | -1 | 3,31E-01 | 6,70E-01 | 58 | 40 | 29 | 16 | 57 | 51 | 38 | 44 | 1,1 | 0,4 | 0,7 |
| chr11:111935002-111935502:downstream_9642 | chr11 | 111895537 | 111935002 | + | DLAT | NM | 8 | 1,5 | 1 | -1 | 3,32E-01 | NA | 3 | 8 | 7 | 4 | 3 | 8 | 14 | 16 | 0,0 | -1,4 | 1,4 |
| chr2:160625094-160625594:downstream_22071 | chr2 | 160568967 | 160625094 | + | MARCH7 | NM | 45 | 0,7 | 1 | -1 | 3,32E-01 | 6,70E-01 | 20 | 16 | 33 | 19 | 36 | 46 | 95 | 98 | -0,5 | -1,2 | 0,7 |
| chr3:155543800-155544300:downstream_23430 | chr3 | 155544300 | 155572248 | - | SLC33A1 | NM | 22 | -1,3 | 1 | 1 | 3,32E-01 | NA | 2 | 0 | 1 | 10 | 31 | 25 | 49 | 58 | -2,2 | -0,9 | -1,3 |
| chr11:2425108-2425608:downstream_18375 | chr11 | 2421717 | 2425108 | + | TSSC4 | NM | 15 | -1,1 | 1 | 1 | 3,32E-01 | NA | 9 | 5 | 6 | 5 | 41 | 32 | 14 | 11 | 0,4 | 1,5 | -1,1 |
| chr17:6943440-6943940:downstream_12786 | chr17 | 6939393 | 6943440 | + | SLC16A13 | NM | 8 | 1,5 | 2 | -1 | 3,33E-01 | NA | 6 | 4 | 10 | 8 | 5 | 1 | 13 | 18 | -0,9 | -2,4 | 1,5 |
| chr4:52886360-52886860:downstream_3672 | chr4 | 52886860 | 52904485 | - | SGCB | NM | 46 | 2,0 | 2 | -1 | 3,33E-01 | 6,72E-01 | 3 | 24 | 7 | 0 | 92 | 79 | 95 | 64 | 2,0 | 0,1 | 1,9 |
| chr3:40559712-40560212:downstream_23526 | chr3 | 40547529 | 40559712 | + | ZNF620 | NM | 5 | 1,7 | 2 | -1 | 3,33E-01 | NA | 7 | 6 | 4 | 1 | 7 | 5 | 6 | 8 | 1,3 | -0,3 | 1,6 |
| chr8:38710546-38711046:downstream_26142 | chr8 | 38585703 | 38710546 | + | TACC1 | NM | 23 | 1,2 | 1 | -1 | 3,33E-01 | NA | 4 | 1 | 8 | 5 | 17 | 9 | 50 | 92 | -1,4 | -2,5 | 1,1 |
| chr4:17844338-17844838:downstream_23718 | chr4 | 17844838 | 18023483 | - | LCORL | NM | 11 | 1,6 | 2 | -1 | 3,34E-01 | NA | 15 | 1 | 17 | 24 | 0 | 4 | 18 | 11 | -1,4 | -2,9 | 1,5 |
| chr11:3847707-3848207:downstream_9031 | chr11 | 3848207 | 3862213 | - | RHOG | NM | 12 | 1,2 | 1 | -1 | 3,34E-01 | NA | 10 | 17 | 4 | 5 | 20 | 14 | 10 | 16 | 1,6 | 0,4 | 1,2 |
| chr7:89794141-89794641:downstream_5587 | chr7 | 89783688 | 89794141 | + | STEAP1 | NM | 87 | 0,8 | 1 | -1 | 3,34E-01 | 6,73E-01 | 7 | 4 | 7 | 6 | 114 | 119 | 191 | 249 | -0,2 | -0,9 | 0,7 |
| chr17:33290205-33290705:downstream_12980 | chr17 | 33288548 | 33290205 | + | ZNF830 | NM | 30 | 0,9 | 1 | -1 | 3,34E-01 | 6,73E-01 | 25 | 56 | 8 | 7 | 67 | 38 | 21 | 16 | 2,4 | 1,5 | 0,9 |
| chr6:170103501-170104001:downstream_25058 | chr6 | 170104001 | 170124106 | - | PHF10 | NM | 9 | 1,5 | 2 | -1 | 3,35E-01 | NA | 2 | 16 | 4 | 4 | 14 | 7 | 11 | 16 | 1,1 | -0,4 | 1,5 |
| chr9:74867140-74867640:downstream_26329 | chr9 | 74729510 | 74867140 | + | GDA | NM | 11 | -1,5 | 2 | 1 | 3,35E-01 | NA | 2 | 0 | 4 | 10 | 12 | 11 | 21 | 25 | -2,5 | -1,0 | -1,5 |
| chr20:36033835-36034335:downstream_22573 | chr20 | 35973087 | 36033835 | + | SRC | NM | 29 | -1,0 | 1 | 1 | 3,36E-01 | 6,74E-01 | 7 | 0 | 7 | 13 | 41 | 41 | 64 | 57 | -1,6 | -0,6 | -1,0 |
| chr1:202300284-202300784:downstream_1299 | chr1 | 202300784 | 202311094 | - | UBE2T | NM | 29 | 1,1 | 1 | -1 | 3,36E-01 | 6,74E-01 | 11 | 7 | 7 | 3 | 43 | 53 | 22 | 88 | 0,8 | -0,2 | 1,0 |
| chr2:25012635-25013135:downstream_1890 | chr2 | 25013135 | 25016251 | - | PTRHD1 | NM | 26 | 0,9 | 1 | -1 | 3,36E-01 | NA | 7 | 18 | 6 | 6 | 47 | 45 | 42 | 38 | 1,2 | 0,2 | 1,0 |
| chr14:95998748-95999248:downstream_19399 | chr14 | 95999248 | 96001209 | - | SNHG10 | NR | 58 | 0,7 | 1 | -1 | 3,36E-01 | 6,75E-01 | 177 | 123 | 31 | 27 | 46 | 32 | 17 | 9 | 2,4 | 1,6 | 0,8 |
| chr3:40581285-40581785:downstream_23527 | chr3 | 40566368 | 40581285 | + | ZNF621 | NM | 5 | 2,0 | 2 | -1 | 3,36E-01 | NA | 3 | 12 | 3 | 4 | 2 | 4 | 0 | 10 | 1,1 | -0,8 | 2,0 |
| chr8:90945063-90945563:downstream_26025 | chr8 | 90945563 | 90996952 | - | NBN | NM | 26 | 1,2 | 1 | -1 | 3,37E-01 | NA | 0 | 19 | 15 | 14 | 17 | 18 | 68 | 54 | -0,6 | -1,8 | 1,2 |
| chr10:124191871-124192371:downstream_17714 | chr10 | 124134093 | 124191871 | + | PLEKHA1 | NM | 17 | -1,6 | 2 | 1 | 3,37E-01 | NA | 2 | 0 | 18 | 15 | 14 | 0 | 47 | 42 | -4,3 | -2,7 | -1,6 |
| chr5:8463209-8463709:downstream_4187 | chr5 | 8457799 | 8463209 | + | MIR4458HG | NR | 23 | -1,0 | 1 | 1 | 3,37E-01 | NA | 50 | 38 | 8 | 8 | 33 | 43 | 1 | 5 | 2,4 | 3,6 | -1,2 |
| chr1:24122029-24122529:downstream_551 | chr1 | 24117645 | 24122029 | + | LYPLA2 | NM | 10 | 1,5 | 2 | -1 | 3,37E-01 | NA | 3 | 11 | 0 | 4 | 18 | 13 | 10 | 19 | 1,8 | 0,1 | 1,7 |
| chrX:20142135-20142635:downstream_7624 | chrX | 20142635 | 20159966 | - | EIF1AX | NM | 22 | 1,0 | 1 | -1 | 3,38E-01 | NA | 13 | 18 | 8 | 3 | 41 | 36 | 38 | 17 | 1,4 | 0,5 | 1,0 |
| chr19:7747748-7748248:downstream_21570 | chr19 | 7745706 | 7747748 | + | TRAPPC5 | NM | 51 | 1,0 | 1 | -1 | 3,38E-01 | 6,77E-01 | 4 | 10 | 6 | 1 | 114 | 101 | 70 | 100 | 1,1 | 0,3 | 0,8 |
| chr11:12966284-12966784:downstream_8850 | chr11 | 12695968 | 12966284 | + | TEAD1 | NM | 24 | -1,4 | 1 | 1 | 3,38E-01 | NA | 2 | 0 | 3 | 9 | 15 | 33 | 71 | 62 | -2,9 | -1,5 | -1,4 |
| chr19:45978437-45978937:downstream_21138 | chr19 | 45971252 | 45978437 | + | FOSB | NM | 9 | -2,4 | 3 | 1 | 3,39E-01 | NA | 10 | 2 | 0 | 0 | 20 | 43 | 0 | 0 | Inf | Inf | NaN |
| chr19:44664462-44664962:downstream_21622 | chr19 | 44645709 | 44664462 | + | ZNF234 | NM | 5 | -1,8 | 2 | 1 | 3,39E-01 | NA | 3 | 1 | 13 | 6 | 3 | 6 | 10 | 2 | -2,1 | -0,3 | -1,8 |
| chr11:6632105-6632605:downstream_18063 | chr11 | 6624937 | 6632105 | + | ILK | NM | 21 | 1,0 | 1 | -1 | 3,39E-01 | NA | 17 | 34 | 6 | 21 | 24 | 17 | 13 | 33 | 1,0 | -0,1 | 1,1 |
| chr2:191371118-191371618:downstream_2356 | chr2 | 191371618 | 191399468 | - | NEMP2 | NM | 8 | -1,5 | 2 | 1 | 3,40E-01 | NA | 3 | 0 | 11 | 6 | 8 | 9 | 10 | 18 | -2,4 | -0,7 | -1,7 |
| chr17:33594761-33595261:downstream_12985 | chr17 | 33570085 | 33594761 | + | SLFN5 | NM | 57 | -0,7 | 1 | 1 | 3,41E-01 | 6,82E-01 | 98 | 47 | 90 | 81 | 46 | 36 | 36 | 24 | -0,2 | 0,4 | -0,7 |
| chr2:223423617-223424117:downstream_1557 | chr2 | 223289321 | 223423617 | + | SGPP2 | NM | 6 | -2,1 | 2 | 1 | 3,41E-01 | NA | 1 | 0 | 4 | 8 | 2 | 6 | 10 | 15 | -3,9 | -1,7 | -2,2 |
| chr8:38132060-38132560:downstream_26180 | chr8 | 38132560 | 38239790 | - | WHSC1L1 | NM | 34 | 1,0 | 1 | -1 | 3,41E-01 | 6,82E-01 | 14 | 14 | 95 | 91 | 4 | 0 | 25 | 33 | -2,8 | -3,8 | 1,1 |
| chr16:46693088-46693588:downstream_12338 | chr16 | 46693588 | 46723144 | - | VPS35 | NM | 60 | -0,7 | 1 | 1 | 3,42E-01 | 6,82E-01 | 12 | 4 | 18 | 26 | 85 | 73 | 118 | 142 | -1,5 | -0,7 | -0,7 |
| chr11:66306234-66306734:downstream_9470 | chr11 | 66306734 | 66313671 | - | ZDHHC24 | NM | 29 | -0,8 | 1 | 1 | 3,42E-01 | 6,82E-01 | 21 | 43 | 17 | 20 | 42 | 56 | 15 | 16 | 0,8 | 1,6 | -0,8 |
| chr14:24612073-24612573:downstream_11075 | chr14 | 24612573 | 24615855 | - | PSME2 | NM | 18 | 1,1 | 1 | -1 | 3,43E-01 | NA | 9 | 12 | 6 | 1 | 31 | 40 | 24 | 19 | 1,7 | 0,7 | 1,0 |
| chr5:89753519-89754019:downstream_4341 | chr5 | 89754019 | 89770585 | - | MBLAC2 | NM | 9 | 1,4 | 2 | -1 | 3,43E-01 | NA | 5 | 6 | 4 | 1 | 14 | 12 | 17 | 12 | 1,1 | -0,2 | 1,3 |
| chr7:73611429-73611929:downstream_25498 | chr7 | 73588705 | 73611429 | + | EIF4H | NM | 11 | 1,2 | 1 | -1 | 3,43E-01 | NA | 14 | 15 | 8 | 10 | 12 | 4 | 7 | 17 | 0,6 | -0,6 | 1,2 |
| chr18:43663609-43664109:downstream_20829 | chr18 | 43664109 | 43684199 | - | ATP5A1 | NM | 154 | -0,7 | 1 | 1 | 3,43E-01 | 6,82E-01 | 7 | 7 | 13 | 4 | 350 | 378 | 210 | 266 | -0,2 | 0,6 | -0,9 |
| chr10:95288849-95289349:downstream_17519 | chr10 | 95256368 | 95288849 | + | CEP55 | NM | 20 | -1,4 | 1 | 1 | 3,43E-01 | NA | 2 | 0 | 10 | 4 | 24 | 15 | 42 | 68 | -3,1 | -1,5 | -1,6 |
| chr2:74652882-74653382:downstream_2045 | chr2 | 74648884 | 74652882 | + | WDR54 | NM | 82 | -1,0 | 1 | 1 | 3,43E-01 | 6,82E-01 | 4 | 0 | 1 | 7 | 154 | 142 | 177 | 169 | -1,1 | -0,2 | -0,9 |
| chr15:59149734-59150234:downstream_19545 | chr15 | 59063392 | 59149734 | + | FAM63B | NM | 9 | 1,4 | 2 | -1 | 3,44E-01 | NA | 5 | 21 | 11 | 19 | 2 | 2 | 6 | 8 | -0,2 | -1,7 | 1,5 |
| chr10:88809458-88809958:downstream_8613 | chr10 | 88809958 | 88854776 | - | GLUD1 | NM | 18 | -1,2 | 1 | 1 | 3,44E-01 | NA | 2 | 2 | 4 | 8 | 21 | 28 | 38 | 41 | -1,8 | -0,7 | -1,1 |
| chr2:3609340-3609840:downstream_22192 | chr2 | 3605975 | 3609340 | + | RNASEH1-AS1 | NR | 8 | 1,4 | 1 | -1 | 3,44E-01 | NA | 7 | 12 | 4 | 3 | 7 | 13 | 15 | 6 | 1,3 | -0,1 | 1,4 |
| chr19:48260323-48260823:downstream_14419 | chr19 | 48248792 | 48260323 | + | GLTSCR2 | NM | 7 | 1,5 | 2 | -1 | 3,45E-01 | NA | 16 | 8 | 7 | 2 | 7 | 6 | 4 | 8 | 1,3 | 0,0 | 1,4 |
| chr10:95662491-95662991:downstream_17772 | chr10 | 95653729 | 95662491 | + | SLC35G1 | NM | 15 | 1,1 | 1 | -1 | 3,45E-01 | NA | 20 | 27 | 13 | 13 | 18 | 4 | 13 | 13 | 0,9 | -0,2 | 1,1 |
| chr8:53534517-53535017:downstream_26080 | chr8 | 53535017 | 53627026 | - | RB1CC1 | NM | 12 | -1,5 | 2 | 1 | 3,45E-01 | NA | 2 | 0 | 8 | 8 | 10 | 8 | 31 | 30 | -3,3 | -1,8 | -1,6 |
| chr15:89175512-89176012:downstream_11880 | chr15 | 89164526 | 89175512 | + | AEN | NM | 61 | 0,8 | 1 | -1 | 3,45E-01 | 6,85E-01 | 32 | 28 | 3 | 5 | 154 | 188 | 43 | 37 | 3,0 | 2,1 | 0,9 |
| chr19:8919751-8920251:downstream_21671 | chr19 | 8920251 | 8942980 | - | ZNF558 | NM | 7 | -1,7 | 2 | 1 | 3,45E-01 | NA | 2 | 0 | 11 | 6 | 7 | 3 | 14 | 16 | -3,4 | -1,6 | -1,8 |
| chr7:73094747-73095247:downstream_5893 | chr7 | 73095247 | 73097781 | - | DNAJC30 | NM | 11 | -1,3 | 1 | 1 | 3,46E-01 | NA | 9 | 10 | 4 | 7 | 14 | 34 | 3 | 8 | 0,8 | 2,1 | -1,3 |
| chr16:68335726-68336226:downstream_20213 | chr16 | 68298418 | 68335726 | + | SLC7A6 | NM | 10 | 1,3 | 1 | -1 | 3,46E-01 | NA | 6 | 15 | 4 | 3 | 15 | 13 | 8 | 15 | 1,5 | 0,2 | 1,3 |
| chr12:48541071-48541571:downstream_10137 | chr12 | 48541571 | 48551377 | - | ASB8 | NM | 8 | 1,6 | 2 | -1 | 3,46E-01 | NA | 4 | 14 | 0 | 2 | 11 | 16 | 8 | 6 | 2,8 | 1,0 | 1,9 |
| chr7:4834026-4834526:downstream_5688 | chr7 | 4815261 | 4834026 | + | AP5Z1 | NM | 5 | -2,0 | 2 | 1 | 3,47E-01 | NA | 9 | 2 | 4 | 1 | 16 | 7 | 3 | 0 | 1,1 | 3,0 | -1,9 |
| chr3:38296979-38297479:downstream_2604 | chr3 | 38207025 | 38296979 | + | OXSR1 | NM | 31 | -1,2 | 1 | 1 | 3,47E-01 | 6,85E-01 | 0 | 3 | 8 | 10 | 17 | 40 | 97 | 75 | -2,6 | -1,6 | -1,0 |
| chr9:127619657-127620157:downstream_7315 | chr9 | 127620157 | 127624240 | - | RPL35 | NM | 248 | 0,6 | 1 | -1 | 3,47E-01 | 6,85E-01 | 24 | 14 | 4 | 8 | 629 | 620 | 288 | 396 | 1,6 | 0,9 | 0,7 |
| chr14:39616514-39617014:downstream_19426 | chr14 | 39617014 | 39639634 | - | TRAPPC6B | NM | 28 | 0,9 | 1 | -1 | 3,47E-01 | 6,85E-01 | 27 | 11 | 8 | 6 | 33 | 69 | 36 | 34 | 1,4 | 0,5 | 0,9 |
| chr15:41673248-41673748:downstream_19654 | chr15 | 41624891 | 41673248 | + | NUSAP1 | NM | 60 | -1,2 | 1 | 1 | 3,48E-01 | 6,85E-01 | 2 | 0 | 11 | 9 | 62 | 17 | 203 | 179 | -3,6 | -2,3 | -1,4 |
| chr8:125465266-125465766:downstream_6706 | chr8 | 125463047 | 125465266 | + | TRMT12 | NM | 56 | 0,9 | 1 | -1 | 3,48E-01 | 6,85E-01 | 18 | 6 | 3 | 3 | 144 | 138 | 53 | 81 | 2,0 | 1,1 | 0,9 |
| chr5:151184915-151185415:downstream_24134 | chr5 | 151151475 | 151184915 | + | G3BP1 | NM | 294 | -0,5 | 0 | 1 | 3,48E-01 | 6,85E-01 | 91 | 59 | 223 | 173 | 332 | 292 | 628 | 557 | -1,4 | -0,9 | -0,5 |
| chr7:99917762-99918262:downstream_5993 | chr7 | 99918262 | 99933930 | - | PMS2P1 | NR | 8 | -1,4 | 1 | 1 | 3,48E-01 | NA | 11 | 3 | 11 | 8 | 10 | 11 | 6 | 5 | -0,4 | 1,0 | -1,4 |
| chr4:7061279-7061779:downstream_3599 | chr4 | 7061779 | 7069800 | - | GRPEL1 | NM | 60 | 0,7 | 1 | -1 | 3,49E-01 | 6,85E-01 | 71 | 74 | 22 | 10 | 127 | 103 | 36 | 41 | 2,2 | 1,6 | 0,6 |
| chr2:71377232-71377732:downstream_2024 | chr2 | 71357443 | 71377232 | + | MPHOSPH10 | NM | 5 | 2,0 | 2 | -1 | 3,49E-01 | NA | 5 | 5 | 3 | 0 | 11 | 2 | 8 | 4 | 1,8 | 0,0 | 1,8 |
| chr12:64203338-64203838:downstream_18882 | chr12 | 64173582 | 64203338 | + | TMEM5 | NM | 39 | 0,9 | 1 | -1 | 3,49E-01 | 6,85E-01 | 11 | 13 | 0 | 11 | 53 | 81 | 79 | 62 | 1,0 | -0,1 | 1,1 |
| chr3:52937082-52937582:downstream_2630 | chr3 | 52937582 | 53080089 | - | SFMBT1 | NM | 9 | -1,9 | 2 | 1 | 3,49E-01 | NA | 1 | 0 | 13 | 10 | 4 | 2 | 17 | 25 | -4,8 | -2,8 | -2,0 |
| chr6:119230335-119230835:downstream_5326 | chr6 | 119215240 | 119230335 | + | ASF1A | NM | 12 | 1,2 | 1 | -1 | 3,49E-01 | NA | 5 | 16 | 11 | 8 | 5 | 13 | 18 | 18 | 0,1 | -1,0 | 1,2 |
| chr5:140024993-140025493:downstream_24414 | chr5 | 140019011 | 140024993 | + | TMCO6 | NM | 30 | 0,8 | 1 | -1 | 3,49E-01 | 6,85E-01 | 36 | 73 | 28 | 25 | 27 | 15 | 18 | 19 | 1,0 | 0,2 | 0,8 |
| chr19:58866549-58867049:downstream_14726 | chr19 | 58863335 | 58866549 | + | A1BG-AS1 | NR | 7 | -1,6 | 2 | 1 | 3,49E-01 | NA | 4 | 1 | 10 | 13 | 7 | 5 | 3 | 14 | -2,2 | -0,6 | -1,6 |
| chr9:21453766-21454266:downstream_6846 | chr9 | 21454266 | 21559697 | - | MIR31HG | NR | 18 | -1,0 | 1 | 1 | 3,49E-01 | NA | 6 | 10 | 24 | 24 | 13 | 19 | 14 | 35 | -1,6 | -0,6 | -1,0 |
| chr1:112004525-112005025:downstream_899 | chr1 | 111991742 | 112004525 | + | ATP5F1 | NM | 26 | -0,9 | 1 | 1 | 3,50E-01 | NA | 60 | 47 | 24 | 28 | 24 | 13 | 4 | 5 | 1,1 | 2,0 | -0,9 |
| chr14:55005334-55005834:downstream_11149 | chr14 | 54976586 | 55005334 | + | CGRRF1 | NM | 45 | -0,8 | 1 | 1 | 3,51E-01 | 6,86E-01 | 24 | 9 | 36 | 24 | 50 | 80 | 70 | 67 | -0,9 | -0,1 | -0,8 |
| chr9:74965840-74966340:downstream_26579 | chr9 | 74966340 | 74980163 | - | ZFAND5 | NM | 56 | 0,7 | 1 | -1 | 3,51E-01 | 6,86E-01 | 38 | 44 | 21 | 11 | 119 | 90 | 51 | 74 | 1,4 | 0,7 | 0,6 |
| chr7:116438440-116438940:downstream_25631 | chr7 | 116312458 | 116438440 | + | MET | NM | 185 | 0,8 | 1 | -1 | 3,51E-01 | 6,86E-01 | 2 | 4 | 3 | 7 | 154 | 158 | 632 | 524 | -0,9 | -1,9 | 1,0 |
| chr15:89859035-89859535:downstream_19672 | chr15 | 89859535 | 89878026 | - | POLG | NM | 29 | 0,9 | 1 | -1 | 3,52E-01 | 6,86E-01 | 38 | 28 | 11 | 7 | 25 | 71 | 26 | 22 | 1,8 | 1,0 | 0,9 |
| chr17:4839925-4840425:downstream_20715 | chr17 | 4840425 | 4843462 | - | SLC25A11 | NM | 49 | 0,6 | 1 | -1 | 3,53E-01 | 6,86E-01 | 46 | 39 | 21 | 21 | 69 | 79 | 57 | 57 | 1,0 | 0,4 | 0,6 |
| chr2:74698584-74699084:downstream_2046 | chr2 | 74699084 | 74699942 | - | MRPL53 | NM | 74 | 0,9 | 1 | -1 | 3,53E-01 | 6,86E-01 | 8 | 9 | 4 | 2 | 182 | 180 | 113 | 94 | 1,5 | 0,8 | 0,7 |
| chr6:109307139-109307639:downstream_25195 | chr6 | 109307639 | 109415708 | - | SESN1 | NM | 43 | 1,1 | 1 | -1 | 3,53E-01 | 6,86E-01 | 37 | 35 | 4 | 0 | 90 | 150 | 17 | 7 | 4,1 | 3,3 | 0,8 |
| chr8:128753680-128754180:downstream_6721 | chr8 | 128748314 | 128753680 | + | MYC | NM | 33 | -0,8 | 1 | 1 | 3,53E-01 | 6,86E-01 | 34 | 13 | 40 | 48 | 37 | 26 | 28 | 39 | -0,9 | -0,1 | -0,8 |
| chr6:64425418-64425918:downstream_25059 | chr6 | 64345706 | 64425418 | + | PHF3 | NM | 21 | 0,9 | 1 | -1 | 3,54E-01 | NA | 21 | 13 | 32 | 43 | 6 | 5 | 26 | 19 | -1,1 | -2,1 | 1,0 |
| chr3:160396235-160396735:downstream_3262 | chr3 | 160394947 | 160396235 | + | ARL14 | NM | 12 | 1,8 | 2 | -1 | 3,54E-01 | NA | 23 | 18 | 1 | 0 | 34 | 17 | 6 | 0 | 4,9 | 3,2 | 1,7 |
| chr17:47906458-47906958:downstream_20529 | chr17 | 47865980 | 47906458 | + | KAT7 | NM | 10 | 1,4 | 2 | -1 | 3,55E-01 | NA | 7 | 2 | 19 | 16 | 3 | 0 | 14 | 23 | -2,1 | -3,5 | 1,4 |
| chr11:111597641-111598141:downstream_8945 | chr11 | 111473114 | 111597641 | + | SIK2 | NM | 6 | 1,9 | 2 | -1 | 3,55E-01 | NA | 3 | 1 | 4 | 11 | 2 | 0 | 11 | 12 | -1,9 | -3,8 | 2,0 |
| chr13:52028851-52029351:downstream_10762 | chr13 | 52027480 | 52028851 | + | INTS6-AS1 | NR | 18 | 1,0 | 1 | -1 | 3,55E-01 | NA | 27 | 16 | 13 | 7 | 21 | 24 | 14 | 25 | 1,1 | 0,2 | 0,9 |
| chr4:140397069-140397569:downstream_3510 | chr4 | 140374960 | 140397069 | + | RAB33B | NM | 8 | 1,5 | 2 | -1 | 3,56E-01 | NA | 5 | 14 | 3 | 1 | 12 | 14 | 10 | 4 | 2,4 | 0,9 | 1,5 |
| chr5:154197167-154197667:downstream_3980 | chr5 | 154092461 | 154197167 | + | LARP1 | NM | 66 | -0,9 | 1 | 1 | 3,56E-01 | 6,91E-01 | 2 | 2 | 7 | 7 | 104 | 89 | 161 | 156 | -1,6 | -0,7 | -0,9 |
| chr5:86687743-86688243:downstream_24327 | chr5 | 86564069 | 86687743 | + | RASA1 | NM | 14 | 1,1 | 1 | -1 | 3,58E-01 | NA | 3 | 6 | 13 | 11 | 6 | 7 | 39 | 30 | -1,4 | -2,5 | 1,1 |
| chr3:53918725-53919225:downstream_3011 | chr3 | 53919225 | 53925989 | - | SELK | NM | 23 | 0,9 | 1 | -1 | 3,58E-01 | NA | 13 | 15 | 7 | 5 | 31 | 52 | 39 | 25 | 1,3 | 0,4 | 0,9 |
| chrX:128936763-128937263:downstream_27124 | chrX | 128937263 | 128977910 | - | ZDHHC9 | NM | 11 | 1,3 | 1 | -1 | 3,58E-01 | NA | 7 | 8 | 1 | 3 | 17 | 18 | 8 | 21 | 1,6 | 0,3 | 1,4 |
| chr3:14530118-14530618:downstream_2583 | chr3 | 14530618 | 14581850 | - | GRIP2 | NM | 5 | 2,2 | 2 | -1 | 3,58E-01 | NA | 1 | 6 | 8 | 11 | 0 | 1 | 10 | 2 | -1,5 | -3,7 | 2,1 |
| chr14:39652422-39652922:downstream_11118 | chr14 | 39644386 | 39652422 | + | PNN | NM | 43 | 0,7 | 1 | -1 | 3,59E-01 | 6,94E-01 | 46 | 84 | 19 | 37 | 53 | 39 | 35 | 32 | 1,2 | 0,5 | 0,8 |
| chrX:3735075-3735575:downstream_7588 | chrX | 3735575 | 3761935 | - | LOC389906 | NR | 31 | 0,9 | 1 | -1 | 3,59E-01 | 6,94E-01 | 31 | 46 | 4 | 6 | 58 | 74 | 21 | 11 | 3,0 | 2,0 | 0,9 |
| chr14:24607673-24608173:downstream_11072 | chr14 | 24608173 | 24610797 | - | EMC9 | NM | 17 | 1,2 | 1 | -1 | 3,60E-01 | NA | 7 | 23 | 3 | 19 | 8 | 22 | 26 | 25 | 0,5 | -0,8 | 1,2 |
| chr3:185361026-185361526:downstream_23202 | chr3 | 185361526 | 185542827 | - | IGF2BP2 | NM | 6 | 1,6 | 2 | -1 | 3,60E-01 | NA | 2 | 4 | 3 | 7 | 2 | 2 | 14 | 12 | -0,9 | -2,6 | 1,7 |
| chr2:208620896-208621396:downstream_21802 | chr2 | 208576263 | 208620896 | + | CCNYL1 | NM | 12 | 1,1 | 1 | -1 | 3,60E-01 | NA | 5 | 12 | 7 | 12 | 7 | 9 | 26 | 16 | -0,2 | -1,4 | 1,2 |
| chr14:100745371-100745871:downstream_11310 | chr14 | 100705101 | 100745371 | + | YY1 | NM | 26 | 0,8 | 1 | -1 | 3,60E-01 | NA | 40 | 30 | 28 | 40 | 16 | 8 | 21 | 21 | 0,0 | -0,8 | 0,9 |
| chr12:34181236-34181736:downstream_10125 | chr12 | 34175215 | 34181236 | + | ALG10 | NM | 7 | 1,5 | 2 | -1 | 3,61E-01 | NA | 7 | 10 | 10 | 16 | 1 | 2 | 6 | 7 | -0,6 | -2,1 | 1,5 |
| chr10:53454745-53455245:downstream_8524 | chr10 | 53455245 | 53459355 | - | CSTF2T | NM | 24 | 0,9 | 1 | -1 | 3,61E-01 | NA | 47 | 45 | 18 | 10 | 20 | 25 | 10 | 14 | 1,7 | 0,9 | 0,8 |
| chr1:241755951-241756451:downstream_1450 | chr1 | 241756451 | 241803701 | - | OPN3 | NM | 16 | -1,1 | 1 | 1 | 3,61E-01 | NA | 2 | 4 | 14 | 8 | 15 | 19 | 22 | 42 | -2,0 | -0,9 | -1,1 |
| chr14:23081265-23081765:downstream_10904 | chr14 | 23067146 | 23081265 | - | ABHD4 | NM | 35 | 0,8 | 1 | -1 | 3,61E-01 | 6,97E-01 | 33 | 45 | 3 | 9 | 75 | 73 | 18 | 24 | 2,7 | 1,8 | 0,9 |
| chr20:47905795-47906295:downstream_22608 | chr20 | 47894714 | 47905795 | + | ZFAS1 | NR | 75 | -0,6 | 1 | 1 | 3,62E-01 | 6,97E-01 | 30 | 17 | 24 | 20 | 156 | 167 | 78 | 111 | 0,1 | 0,8 | -0,6 |
| chr17:38293045-38293545:downstream_13043 | chr17 | 38278789 | 38293045 | + | MSL1 | NM | 13 | 1,2 | 1 | -1 | 3,62E-01 | NA | 4 | 16 | 3 | 3 | 20 | 28 | 14 | 19 | 1,8 | 0,5 | 1,2 |
| chr5:172396774-172397274:downstream_4581 | chr5 | 172386438 | 172396774 | + | RPL26L1 | NM | 12 | 1,7 | 2 | -1 | 3,62E-01 | NA | 2 | 11 | 1 | 0 | 30 | 35 | 6 | 13 | 3,1 | 1,8 | 1,3 |
| chr3:38456467-38456967:downstream_2605 | chr3 | 38388250 | 38456467 | + | XYLB | NM | 6 | -1,6 | 2 | 1 | 3,62E-01 | NA | 2 | 0 | 6 | 7 | 6 | 9 | 11 | 11 | -2,3 | -0,7 | -1,7 |
| chr12:2939770-2940270:downstream_9978 | chr12 | 2921787 | 2939770 | + | ITFG2 | NR | 13 | 1,2 | 1 | -1 | 3,63E-01 | NA | 2 | 6 | 11 | 4 | 7 | 9 | 46 | 21 | -0,9 | -2,1 | 1,2 |
| chr5:139623374-139623874:downstream_4096 | chr5 | 139554652 | 139623374 | + | CYSTM1 | NM | 70 | -0,9 | 1 | 1 | 3,63E-01 | 6,98E-01 | 7 | 1 | 6 | 3 | 181 | 176 | 74 | 116 | -0,1 | 0,9 | -1,0 |
| chr1:93828148-93828648:downstream_833 | chr1 | 93811477 | 93828148 | + | DR1 | NM | 54 | 0,6 | 1 | -1 | 3,63E-01 | 6,98E-01 | 83 | 79 | 25 | 30 | 84 | 55 | 40 | 33 | 1,6 | 0,9 | 0,6 |

| | | | | | | | | | | | | | | | | | | | | | | | |
|---|---|---|---|---|---|---|---|---|---|---|---|---|---|---|---|---|---|---|---|---|---|---|---|
| chr8:125562227-125562727:downstream_26028 | chr8 | 125551342 | 125562227 | + | NDUFB9 | NM | 118 | -0,7 | 1 | 1 | 3,63E-01 | 6,98E-01 | 8 | 5 | 7 | 7 | 255 | 311 | 153 | 200 | -0,1 | 0,7 | -0,7 |
| chr16:2820914-2821414:downstream_20237 | chr16 | 2821414 | 2827297 | - | TCEB2 | NM | 12 | 1,2 | 1 | -1 | 3,64E-01 | NA | 7 | 10 | 1 | 2 | 28 | 20 | 14 | 15 | 2,1 | 0,8 | 1,3 |
| chr10:44051292-44051792:downstream_17830 | chr10 | 44051792 | 44070066 | - | ZNF239 | NM | 13 | 1,4 | 2 | -1 | 3,64E-01 | NA | 20 | 16 | 3 | 0 | 18 | 39 | 4 | 5 | 3,7 | 2,6 | 1,1 |
| chr8:117786921-117787421:downstream_6684 | chr8 | 117778741 | 117786921 | + | UTP23 | NM | 21 | 1,0 | 1 | -1 | 3,64E-01 | NA | 33 | 14 | 6 | 4 | 40 | 43 | 14 | 18 | 2,3 | 1,4 | 0,9 |
| chr19:58027791-58028291:downstream_14699 | chr19 | 58011221 | 58027791 | + | ZNF773 | NR | 14 | 1,3 | 1 | -1 | 3,64E-01 | NA | 16 | 32 | 4 | 0 | 27 | 22 | 3 | 6 | 3,5 | 2,5 | 1,0 |
| chr5:53180113-53180613:downstream_3962 | chr5 | 53180613 | 53606403 | - | ARL15 | NM | 6 | -2,2 | 2 | 1 | 3,64E-01 | NA | 0 | 0 | 8 | 11 | 2 | 0 | 13 | 11 | -Inf | -3,8 | -Inf |
| chr2:86369280-86369780:downstream_2088 | chr2 | 86333304 | 86369280 | + | PTCD3 | NM | 30 | 0,8 | 1 | -1 | 3,64E-01 | 6,99E-01 | 24 | 23 | 8 | 10 | 54 | 51 | 35 | 34 | 1,4 | 0,6 | 0,8 |
| chr14:58731582-58732082:downstream_19349 | chr14 | 58732082 | 58764855 | - | PSMA3-AS1 | NR | 18 | 1,1 | 1 | -1 | 3,64E-01 | NA | 9 | 1 | 6 | 3 | 19 | 26 | 49 | 33 | 0,2 | -0,9 | 1,0 |
| chr5:102421203-102421703:downstream_4354 | chr5 | 102421703 | 102455842 | - | GIN1 | NM | 8 | -1,5 | 2 | 1 | 3,65E-01 | NA | 0 | 6 | 10 | 4 | 8 | 17 | 17 | 5 | -1,3 | 0,2 | -1,5 |
| chr2:157442915-157443415:downstream_21946 | chr2 | 157291964 | 157442915 | + | GPD2 | NM | 33 | 1,1 | 1 | -1 | 3,66E-01 | 7,01E-01 | 1 | 3 | 6 | 5 | 25 | 9 | 89 | 123 | -1,5 | -2,6 | 1,1 |
| chr20:2638540-2639040:downstream_22442 | chr20 | 2639040 | 2644865 | - | IDH3B | NM | 25 | 0,9 | 1 | -1 | 3,67E-01 | NA | 19 | 48 | 6 | 20 | 35 | 25 | 24 | 21 | 1,4 | 0,4 | 1,0 |
| chr3:158414396-158414896:downstream_23378 | chr3 | 158414896 | 158450275 | - | RARRES1 | NM | 38 | -0,8 | 1 | 1 | 3,67E-01 | 7,02E-01 | 6 | 10 | 11 | 7 | 67 | 95 | 51 | 53 | -0,2 | 0,6 | -0,8 |
| chr17:71244087-71244587:downstream_13333 | chr17 | 71244587 | 71258019 | - | CPSF4L | NM | 6 | 1,9 | 2 | -1 | 3,67E-01 | NA | 1 | 17 | 1 | 2 | 3 | 11 | 10 | 2 | 2,2 | 0,3 | 2,0 |
| chr9:104122198-104122698:downstream_26226 | chr9 | 104122698 | 104147287 | - | BAAT | NM | 15 | -1,1 | 1 | 1 | 3,67E-01 | NA | 3 | 3 | 13 | 18 | 11 | 13 | 29 | 29 | -2,3 | -1,3 | -1,0 |
| chr5:6371538-6372038:downstream_4178 | chr5 | 6372038 | 6378639 | - | MED10 | NM | 12 | 1,4 | 2 | -1 | 3,67E-01 | NA | 8 | 6 | 0 | 2 | 25 | 29 | 4 | 25 | 2,5 | 0,9 | 1,6 |
| chr1:179808601-179809101:downstream_17346 | chr1 | 179809101 | 179846941 | - | TOR1AIP2 | NM | 4 | 2,1 | 2 | -1 | 3,67E-01 | NA | 4 | 10 | 0 | 1 | 9 | 4 | 3 | 2 | 4,1 | 1,5 | 2,5 |
| chr16:52471417-52471917:downstream_20250 | chr16 | 52471917 | 52581714 | - | TOX3 | NM | 7 | -1,9 | 2 | 1 | 3,68E-01 | NA | 0 | 0 | 4 | 6 | 3 | 2 | 25 | 18 | -Inf | -3,1 | -Inf |
| chr11:95501605-95502105:downstream_9601 | chr11 | 95502105 | 95522954 | - | FAM76B | NM | 4 | 1,7 | 2 | -1 | 3,68E-01 | NA | 3 | 7 | 4 | 1 | 5 | 3 | 6 | 7 | 1,0 | -0,6 | 1,6 |
| chr17:35837226-35837726:downstream_20757 | chr17 | 35766976 | 35837226 | + | TADA2A | NM | 25 | 0,8 | 1 | -1 | 3,68E-01 | NA | 10 | 16 | 25 | 29 | 12 | 14 | 47 | 46 | -1,0 | -1,9 | 0,8 |
| chr9:12823059-12823559:downstream_6984 | chr9 | 12775011 | 12823059 | + | LURAP1L | NM | 8 | 1,7 | 2 | -1 | 3,68E-01 | NA | 4 | 0 | 15 | 8 | 2 | 0 | 17 | 15 | -2,5 | -4,3 | 1,7 |
| chr21:18965467-18965967:downstream_22628 | chr21 | 18965967 | 18985268 | - | BTG3 | NM | 15 | -1,2 | 1 | 1 | 3,70E-01 | NA | 5 | 0 | 6 | 7 | 19 | 33 | 25 | 30 | -1,3 | -0,1 | -1,2 |
| chr2:111394774-111395274:downstream_21774 | chr2 | 111395274 | 111435684 | - | BUB1 | NM | 9 | -1,4 | 2 | 1 | 3,70E-01 | NA | 1 | 3 | 6 | 11 | 9 | 10 | 8 | 25 | -2,2 | -0,9 | -1,3 |
| chr13:73329039-73329539:downstream_18970 | chr13 | 73329539 | 73356266 | - | DIS3 | NM | 13 | 1,1 | 1 | -1 | 3,70E-01 | NA | 18 | 15 | 8 | 4 | 19 | 12 | 17 | 7 | 1,4 | 0,3 | 1,1 |
| chr5:140476964-140477464:downstream_4468 | chr5 | 140474190 | 140476964 | + | PCDHB2 | NM | 5 | 1,6 | 2 | -1 | 3,70E-01 | NA | 6 | 6 | 3 | 2 | 8 | 4 | 7 | 7 | 1,4 | -0,2 | 1,5 |
| chr11:119005765-119006265:downstream_18047 | chr11 | 118992232 | 119005765 | + | HINFP | NM | 5 | -1,9 | 2 | 1 | 3,70E-01 | NA | 1 | 1 | 7 | 4 | 1 | 10 | 11 | 7 | -2,6 | -0,8 | -1,9 |
| chr12:102591298-102591798:downstream_10384 | chr12 | 102513955 | 102591298 | + | PARPBP | NM | 18 | 1,1 | 1 | -1 | 3,71E-01 | NA | 2 | 4 | 8 | 6 | 14 | 8 | 49 | 51 | -1,2 | -2,2 | 1,0 |
| chr2:75889331-75889831:downstream_21934 | chr2 | 75889831 | 75938111 | - | GCFC2 | NM | 11 | 1,1 | 1 | -1 | 3,71E-01 | NA | 8 | 14 | 8 | 7 | 11 | 9 | 21 | 10 | 0,5 | -0,6 | 1,2 |
| chr19:42460332-42460832:downstream_14326 | chr19 | 42460832 | 42463528 | - | RABAC1 | NM | 53 | -0,8 | 1 | 1 | 3,71E-01 | 7,09E-01 | 3 | 7 | 4 | 7 | 147 | 91 | 86 | 80 | -0,2 | 0,5 | -0,7 |
| chr2:27531859-27532359:downstream_1912 | chr2 | 27532359 | 27545969 | - | MPV17 | NM | 70 | 0,8 | 1 | -1 | 3,71E-01 | 7,09E-01 | 11 | 22 | 8 | 2 | 146 | 196 | 88 | 81 | 1,6 | 1,0 | 0,6 |
| chr3:122128961-122129461:downstream_3138 | chr3 | 122103022 | 122128961 | + | FAM162A | NM | 16 | -1,1 | 1 | 1 | 3,72E-01 | NA | 5 | 0 | 4 | 7 | 27 | 30 | 24 | 31 | -1,1 | 0,1 | -1,2 |
| chr7:116201239-116201739:downstream_25429 | chr7 | 116164838 | 116201239 | + | CAV1 | NM | 72 | 1,9 | 2 | -1 | 3,72E-01 | 7,09E-01 | 19 | 0 | 3 | 2 | 142 | 133 | 104 | 176 | 1,8 | 0,0 | 1,9 |
| chr2:152104227-152104727:downstream_2273 | chr2 | 152104727 | 152118389 | - | RBM43 | NM | 7 | -1,6 | 2 | 1 | 3,72E-01 | NA | 3 | 0 | 3 | 9 | 13 | 6 | 17 | 7 | -1,9 | -0,3 | -1,6 |
| chr6:53512198-53512698:downstream_5202 | chr6 | 53512698 | 53530506 | - | KLHL31 | NM | 2 | 2,7 | 3 | -1 | 3,73E-01 | NA | 7 | 1 | 0 | 0 | 6 | 3 | 0 | 2 | Inf | 1,8 | Inf |
| chr15:85185694-85186194:downstream_19707 | chr15 | 85174690 | 85185694 | + | SCAND2P | NR | 10 | -1,2 | 1 | 1 | 3,73E-01 | NA | 7 | 10 | 8 | 12 | 6 | 22 | 4 | 10 | -0,3 | 1,0 | -1,3 |
| chr10:103543699-103544199:downstream_17650 | chr10 | 103544199 | 103578222 | - | MGEA5 | NM | 36 | 0,9 | 1 | -1 | 3,74E-01 | 7,12E-01 | 5 | 26 | 17 | 5 | 59 | 46 | 58 | 74 | 0,5 | -0,3 | 0,8 |
| chr1:45959097-45959597:downstream_16558 | chr1 | 45959597 | 45965751 | - | CCDC163P | NR | 5 | 2,0 | 2 | -1 | 3,74E-01 | NA | 4 | 10 | 1 | 0 | 7 | 12 | 6 | 1 | 3,3 | 1,5 | 1,8 |
| chrX:47344626-47345126:downstream_7697 | chrX | 47342114 | 47344626 | + | LINC01560 | NR | 14 | 1,1 | 1 | -1 | 3,74E-01 | NA | 15 | 34 | 14 | 9 | 8 | 11 | 11 | 7 | 1,1 | 0,0 | 1,1 |
| chr9:128127290-128127790:downstream_26325 | chr9 | 128024072 | 128127290 | + | GAPVD1 | NM | 23 | -1,0 | 1 | 1 | 3,74E-01 | NA | 2 | 3 | 18 | 12 | 11 | 27 | 61 | 48 | -2,5 | -1,5 | -1,0 |
| chr11:121959310-121959810:downstream_8956 | chr11 | 121959810 | 122073770 | - | MIR100HG | NR | 9 | 1,6 | 2 | -1 | 3,74E-01 | NA | 1 | 12 | 6 | 12 | 7 | 0 | 18 | 15 | -0,5 | -2,2 | 1,6 |
| chr9:15463564-15464064:downstream_26458 | chr9 | 15464064 | 15511003 | - | PSIP1 | NM | 19 | -1,2 | 1 | 1 | 3,74E-01 | NA | 2 | 0 | 6 | 7 | 21 | 20 | 47 | 46 | -2,4 | -1,2 | -1,2 |
| chr22:35820495-35820995:downstream_15581 | chr22 | 35796115 | 35820495 | + | MCM5 | NM | 7 | -1,6 | 2 | 1 | 3,74E-01 | NA | 3 | 1 | 13 | 1 | 11 | 9 | 8 | 11 | -1,7 | 0,0 | -1,7 |
| chrX:30322038-30322538:downstream_7652 | chrX | 30322538 | 30327495 | - | NR0B1 | NM | 53 | 0,9 | 1 | -1 | 3,75E-01 | 7,13E-01 | 10 | 9 | 1 | 3 | 138 | 127 | 72 | 60 | 2,0 | 1,0 | 1,0 |
| chrX:77382324-77382824:downstream_7818 | chrX | 77359665 | 77382324 | + | PGK1 | NM | 107 | 0,5 | 1 | -1 | 3,76E-01 | 7,13E-01 | 29 | 39 | 45 | 27 | 133 | 155 | 224 | 206 | -0,1 | -0,6 | 0,5 |
| chr17:61896292-61896792:downstream_13303 | chr17 | 61896792 | 61905031 | - | FTSJ3 | NM | 54 | 0,7 | 1 | -1 | 3,76E-01 | 7,13E-01 | 14 | 13 | 10 | 9 | 85 | 99 | 102 | 104 | 0,5 | -0,2 | 0,7 |
| chr11:126132313-126132813:downstream_18314 | chr11 | 126132813 | 126138877 | - | SRPR | NM | 54 | 0,7 | 1 | -1 | 3,76E-01 | 7,13E-01 | 20 | 38 | 7 | 15 | 122 | 92 | 57 | 80 | 1,4 | 0,6 | 0,8 |
| chr12:10564413-10564913:downstream_18619 | chr12 | 10564913 | 10573194 | - | KLRC3 | NM | 11 | -1,2 | 1 | 1 | 3,77E-01 | NA | 2 | 5 | 15 | 18 | 7 | 8 | 24 | 10 | -2,4 | -1,2 | -1,1 |
| chr16:67209640-67210140:downstream_20146 | chr16 | 67204399 | 67209640 | + | NOL3 | NM | 21 | 1,0 | 1 | -1 | 3,77E-01 | NA | 5 | 6 | 4 | 2 | 30 | 39 | 36 | 45 | 0,7 | -0,2 | 0,9 |
| chr4:142155850-142156350:downstream_24000 | chr4 | 142142040 | 142155850 | + | ZNF330 | NM | 17 | 1,1 | 1 | -1 | 3,77E-01 | NA | 16 | 2 | 6 | 4 | 22 | 28 | 25 | 30 | 0,9 | -0,1 | 1,0 |
| chr12:32898977-32899477:downstream_10124 | chr12 | 32899477 | 32908887 | - | YARS2 | NM | 24 | 0,8 | 1 | -1 | 3,77E-01 | NA | 42 | 23 | 19 | 23 | 20 | 21 | 24 | 22 | 0,6 | -0,2 | 0,8 |
| chr6:146118771-146119271:downstream_5386 | chr6 | 146119271 | 146135921 | - | FBXO30 | NM | 13 | 1,2 | 1 | -1 | 3,78E-01 | NA | 21 | 3 | 10 | 22 | 7 | 5 | 15 | 21 | -0,4 | -1,6 | 1,2 |
| chr20:30532257-30532757:downstream_14781 | chr20 | 30532757 | 30539883 | - | PDRG1 | NM | 9 | 1,3 | 1 | -1 | 3,78E-01 | NA | 11 | 22 | 4 | 5 | 4 | 15 | 6 | 7 | 1,9 | 0,6 | 1,3 |
| chr1:6474793-6475293:downstream_409 | chr1 | 6475293 | 6479979 | - | HES2 | NM | 13 | -2,0 | 2 | 1 | 3,78E-01 | NA | 18 | 2 | 1 | 0 | 40 | 43 | 1 | 0 | 3,8 | 5,9 | -2,1 |
| chr3:37026856-37027356:downstream_2874 | chr3 | 37027356 | 37034795 | - | EPM2AIP1 | NM | 19 | 1,0 | 1 | -1 | 3,78E-01 | NA | 38 | 44 | 6 | 7 | 26 | 15 | 10 | 4 | 2,7 | 1,6 | 1,1 |
| chr1:151264381-151264881:downstream_17437 | chr1 | 151254030 | 151264381 | + | ZNF687 | NM | 4 | 1,7 | 2 | -1 | 3,78E-01 | NA | 4 | 8 | 1 | 3 | 5 | 2 | 3 | 7 | 1,3 | -0,5 | 1,8 |
| chr16:3348307-3348807:downstream_12147 | chr16 | 3348807 | 3355439 | - | TIGD7 | NM | 5 | 2,1 | 2 | -1 | 3,78E-01 | NA | 6 | 7 | 0 | 1 | 2 | 18 | 6 | 2 | 3,9 | 1,5 | 2,5 |
| chr2:191833261-191833761:downstream_22259 | chr2 | 191833761 | 191878976 | - | STAT1 | NM | 40 | 0,7 | 1 | -1 | 3,79E-01 | 7,16E-01 | 10 | 8 | 8 | 11 | 50 | 52 | 79 | 106 | -0,1 | -0,9 | 0,7 |
| chr17:65336118-65336618:downstream_20656 | chr17 | 65336618 | 65362721 | - | PSMD12 | NM | 116 | -0,5 | 1 | 1 | 3,79E-01 | 7,16E-01 | 78 | 126 | 109 | 136 | 106 | 153 | 97 | 120 | -0,3 | 0,3 | -0,5 |
| chr2:198256197-198256697:downstream_22215 | chr2 | 198256697 | 198299817 | - | SF3B1 | NM | 236 | 0,5 | 1 | -1 | 3,79E-01 | 7,16E-01 | 20 | 17 | 25 | 34 | 277 | 272 | 678 | 564 | -0,7 | -1,2 | 0,5 |
| chr1:151312615-151313115:downstream_17176 | chr1 | 151313115 | 151319769 | - | RFX5 | NM | 26 | -1,2 | 1 | 1 | 3,79E-01 | NA | 0 | 2 | 6 | 8 | 33 | 12 | 79 | 71 | -2,8 | -1,7 | -1,1 |
| chr19:33369403-33369903:downstream_13760 | chr19 | 33369903 | 33462935 | - | CEP89 | NM | 5 | 1,8 | 2 | -1 | 3,79E-01 | NA | 2 | 7 | 6 | 11 | 2 | 0 | 7 | 4 | -0,8 | -2,8 | 1,9 |
| chr11:61920635-61921135:downstream_18065 | chr11 | 61891444 | 61920635 | + | INCENP | NM | 32 | -0,9 | 1 | 1 | 3,80E-01 | 7,16E-01 | 8 | 1 | 28 | 37 | 26 | 12 | 75 | 68 | -2,8 | -1,9 | -0,9 |
| chr19:33795963-33796463:downstream_14199 | chr19 | 33793762 | 33795963 | + | CEBPA-AS1 | NR | 5 | 1,6 | 2 | -1 | 3,80E-01 | NA | 3 | 5 | 3 | 7 | 5 | 0 | 10 | 11 | -0,3 | -2,1 | 1,7 |
| chr17:74561430-74561930:downstream_20731 | chr17 | 74553845 | 74561430 | + | SNHG16 | NR | 49 | -0,7 | 1 | 1 | 3,80E-01 | 7,16E-01 | 37 | 19 | 17 | 16 | 102 | 119 | 39 | 43 | 0,8 | 1,4 | -0,7 |
| chr7:98963885-98964385:downstream_25392 | chr7 | 98923495 | 98963885 | + | ARPC1A | NM | 26 | 1,0 | 1 | -1 | 3,80E-01 | NA | 7 | 1 | 7 | 3 | 34 | 24 | 82 | 50 | -0,3 | -1,2 | 0,9 |
| chr6:33539822-33540322:downstream_5088 | chr6 | 33540322 | 33548070 | - | BAK1 | NM | 6 | 1,7 | 2 | -1 | 3,81E-01 | NA | 19 | 9 | 1 | 1 | 11 | 7 | 3 | 2 | 3,6 | 2,0 | 1,7 |
| chr5:98266713-98267213:downstream_4350 | chr5 | 98264837 | 98266713 | + | LOC100289230 | NR | 18 | 0,9 | 1 | -1 | 3,81E-01 | NA | 22 | 26 | 10 | 12 | 20 | 19 | 19 | 14 | 1,1 | 0,2 | 0,9 |
| chr6:116914764-116915264:downstream_25185 | chr6 | 116892529 | 116914764 | + | RWDD1 | NM | 16 | 1,0 | 1 | -1 | 3,81E-01 | NA | 15 | 13 | 3 | 8 | 33 | 17 | 24 | 19 | 1,3 | 0,2 | 1,1 |
| chr8:22477446-22477946:downstream_6487 | chr8 | 22477946 | 22526661 | - | BIN3 | NM | 10 | 1,3 | 2 | -1 | 3,81E-01 | NA | 4 | 13 | 6 | 1 | 21 | 8 | 15 | 11 | 1,4 | 0,2 | 1,2 |
| chr7:94186328-94186828:downstream_5949 | chr7 | 94139169 | 94186328 | + | CASD1 | NM | 24 | 1,0 | 1 | -1 | 3,81E-01 | NA | 6 | 1 | 10 | 21 | 7 | 8 | 63 | 74 | -2,2 | -3,2 | 1,0 |
| chr6:149915720-149916220:downstream_5392 | chr6 | 149914751 | 149915720 | + | RPS18P9 | NR | 16 | -0,9 | 1 | 1 | 3,82E-01 | NA | 8 | 14 | 17 | 17 | 21 | 16 | 14 | 17 | -0,6 | 0,3 | -0,9 |
| chr21:40769815-40770315:downstream_22728 | chr21 | 40752212 | 40769815 | + | WRB | NM | 14 | 1,1 | 1 | -1 | 3,83E-01 | NA | 5 | 8 | 3 | 6 | 24 | 11 | 18 | 35 | 0,6 | -0,6 | 1,2 |
| chr12:93797024-93797524:downstream_18715 | chr12 | 93771698 | 93797024 | + | NUDT4 | NM | 16 | 1,0 | 1 | -1 | 3,83E-01 | NA | 19 | 17 | 14 | 12 | 15 | 11 | 29 | 9 | 0,5 | -0,6 | 1,0 |
| chr12:19529333-19529833:downstream_18745 | chr12 | 19282625 | 19529333 | + | PLEKHA5 | NM | 6 | -2,1 | 2 | 1 | 3,83E-01 | NA | 0 | 0 | 4 | 11 | 2 | 0 | 10 | 20 | -Inf | -3,6 | -Inf |
| chr19:41930910-41931410:downstream_21024 | chr19 | 41903693 | 41930910 | + | BCKDHA | NM | 10 | 1,4 | 2 | -1 | 3,83E-01 | NA | 4 | 6 | 1 | 1 | 28 | 15 | 15 | 10 | 2,2 | 0,8 | 1,4 |
| chr10:88684945-88685445:downstream_8310 | chr10 | 88516395 | 88684945 | + | BMPR1A | NM | 20 | 1,3 | 1 | -1 | 3,83E-01 | NA | 2 | 1 | 11 | 25 | 3 | 0 | 65 | 55 | -3,8 | -5,2 | 1,4 |
| chr12:42475147-42475647:downstream_18590 | chr12 | 42475647 | 42538673 | - | GXYLT1 | NM | 9 | 1,3 | 1 | -1 | 3,83E-01 | NA | 12 | 5 | 0 | 6 | 14 | 14 | 11 | 14 | 1,6 | 0,1 | 1,4 |
| chr12:94701555-94702055:downstream_18498 | chr12 | 94702055 | 94853764 | - | CEP83 | NM | 4 | 1,8 | 2 | -1 | 3,83E-01 | NA | 4 | 2 | 4 | 6 | 0 | 3 | 13 | 4 | -0,7 | -2,5 | 1,8 |
| chr4:183810743-183811243:downstream_23615 | chr4 | 183811243 | 183838630 | - | DCTD | NM | 15 | 1,1 | 1 | -1 | 3,83E-01 | NA | 9 | 5 | 1 | 4 | 24 | 28 | 17 | 30 | 1,3 | 0,2 | 1,2 |
| chr1:159752333-159752833:downstream_1149 | chr1 | 159750758 | 159752333 | + | DUSP23 | NM | 45 | -0,8 | 1 | 1 | 3,83E-01 | 7,21E-01 | 8 | 3 | 3 | 13 | 86 | 94 | 95 | 62 | -0,5 | 0,2 | -0,7 |
| chr20:32442173-32442673:downstream_14978 | chr20 | 32399109 | 32442173 | + | CHMP4B | NM | 14 | 1,3 | 2 | -1 | 3,84E-01 | NA | 3 | 9 | 3 | 1 | 11 | 45 | 15 | 23 | 1,7 | 0,5 | 1,2 |
| chr5:132440709-132441209:downstream_4412 | chr5 | 132387661 | 132440709 | + | HSPA4 | NM | 54 | -0,6 | 1 | 1 | 3,85E-01 | 7,23E-01 | 34 | 35 | 68 | 71 | 53 | 42 | 64 | 65 | -1,0 | -0,4 | -0,6 |
| chr22:31721289-31721789:downstream_22871 | chr22 | 31721789 | 31742249 | - | PATZ1 | NM | 6 | 1,5 | 2 | -1 | 3,85E-01 | NA | 2 | 6 | 6 | 9 | 3 | 1 | 11 | 11 | -0,8 | -2,4 | 1,5 |
| chrX:154351349-154351849:downstream_26650 | chrX | 154299694 | 154351349 | + | BRCC3 | NM | 19 | 0,9 | 1 | -1 | 3,86E-01 | NA | 7 | 22 | 18 | 21 | 13 | 12 | 36 | 27 | -0,4 | -1,4 | 0,9 |
| chr5:96365115-96365615:downstream_24199 | chr5 | 96271345 | 96365115 | + | LNPEP | NM | 166 | -0,5 | 1 | 1 | 3,86E-01 | 7,23E-01 | 71 | 32 | 182 | 213 | 86 | 140 | 301 | 303 | -1,9 | -1,4 | -0,5 |
| chr12:20837041-20837541:downstream_18732 | chr12 | 20522178 | 20837041 | + | PDE3A | NM | 34 | 0,9 | 1 | -1 | 3,86E-01 | 7,23E-01 | 11 | 2 | 46 | 39 | 10 | 4 | 68 | 88 | -2,7 | -3,5 | 0,8 |
| chr1:109473044-109473544:downstream_190 | chr1 | 109419602 | 109473044 | + | GPSM2 | NM | 14 | 1,0 | 1 | -1 | 3,86E-01 | NA | 7 | 11 | 14 | 17 | 2 | 12 | 28 | 22 | -0,8 | -1,8 | 1,0 |
| chr1:113243368-113243868:downstream_16977 | chr1 | 113216933 | 113243368 | + | MOV10 | NM | 15 | 1,0 | 1 | -1 | 3,86E-01 | NA | 8 | 4 | 3 | 7 | 16 | 21 | 39 | 24 | 0,4 | -0,7 | 1,1 |
| chr1:180238297-180238797:downstream_1250 | chr1 | 180238797 | 180243816 | - | LHX4-AS1 | NR | 4 | -2,2 | 3 | 1 | 3,87E-01 | NA | 1 | 0 | 17 | 2 | 2 | 0 | 8 | 3 | -4,6 | -2,3 | -2,3 |
| chr1:43828873-43829373:downstream_687 | chr1 | 43824625 | 43828873 | + | CDC20 | NM | 6 | 1,6 | 2 | -1 | 3,87E-01 | NA | 1 | 16 | 4 | 6 | 3 | 4 | 7 | 6 | 0,8 | -0,8 | 1,6 |
| chr3:3190706-3191206:downstream_23489 | chr3 | 3168599 | 3190706 | + | TRNT1 | NM | 39 | 0,7 | 1 | -1 | 3,88E-01 | 7,24E-01 | 26 | 38 | 15 | 16 | 51 | 68 | 50 | 44 | 1,0 | 0,3 | 0,7 |
| chr9:33371155-33371655:downstream_26410 | chr9 | 33290417 | 33371155 | + | NFX1 | NM | 9 | -1,2 | 1 | 1 | 3,88E-01 | NA | 3 | 3 | 10 | 10 | 8 | 13 | 6 | 21 | -1,7 | -0,4 | -1,3 |
| chr14:20810729-20811229:downstream_11024 | chr14 | 20811229 | 20811570 | - | RPPH1 | NR | 11567 | 1,0 | 1 | -1 | 3,88E-01 | 7,24E-01 | 9 | 2 | 0 | 2 | 39377 | 28157 | 12182 | 12805 | 2,7 | 1,4 | 1,3 |
| chr19:3543696-3544196:downstream_21280 | chr19 | 3544196 | 3557582 | - | MFSD12 | NM | 6 | 1,8 | 2 | -1 | 3,88E-01 | NA | 0 | 16 | 4 | 1 | 13 | 2 | 7 | 8 | 1,7 | 0,0 | 1,7 |
| chr2:162268228-162268728:downstream_1732 | chr2 | 162164785 | 162268228 | + | PSMD14 | NM | 59 | -0,9 | 1 | 1 | 3,88E-01 | 7,24E-01 | 3 | 2 | 7 | 6 | 128 | 69 | 135 | 125 | -1,3 | -0,4 | -0,9 |
| chr16:15131209-15131709:downstream_20154 | chr16 | 15131709 | 15149936 | - | NTAN1 | NM | 6 | -1,4 | 2 | 1 | 3,88E-01 | NA | 5 | 6 | 6 | 8 | 6 | 9 | 3 | 4 | -0,4 | 1,1 | -1,4 |
| chr12:56555366-56555866:downstream_18695 | chr12 | 56552044 | 56555366 | + | MYL6 | NM | 119 | 0,6 | 1 | -1 | 3,88E-01 | 7,24E-01 | 17 | 27 | 10 | 6 | 305 | 282 | 118 | 185 | 1,5 | 1,0 | 0,6 |
| chr19:47730278-47730778:downstream_14407 | chr19 | 47730198 | 47730278 | + | MIR3190 | NR | 5 | -2,2 | 3 | 1 | 3,89E-01 | NA | 6 | 9 | 1 | 0 | 15 | 10 | 0 | 0 | 3,4 | Inf | -Inf |
| chr11:93496268-93496768:downstream_17905 | chr11 | 93474759 | 93496268 | + | C11orf54 | NM | 22 | -0,9 | 1 | 1 | 3,89E-01 | NA | 19 | 7 | 29 | 40 | 19 | 13 | 28 | 20 | -1,4 | -0,6 | -0,8 |
| chr10:21782920-21783420:downstream_8438 | chr10 | 21783420 | 21786213 | - | CASC10 | NM | 10 | -1,2 | 1 | 1 | 3,89E-01 | NA | 10 | 12 | 11 | 19 | 7 | 8 | 8 | 2 | -0,5 | 0,6 | -1,1 |
| chr10:35296978-35297478:downstream_17531 | chr10 | 35297478 | 35379570 | - | CUL2 | NM | 4 | 2,0 | 2 | -1 | 3,89E-01 | NA | 4 | 5 | 0 | 8 | 2 | 0 | 4 | 6 | 0,1 | -2,0 | 2,1 |
| chr12:27850566-27851066:downstream_10113 | chr12 | 27849427 | 27850566 | + | REP15 | NM | 5 | 2,1 | 2 | -1 | 3,90E-01 | NA | 2 | 0 | 8 | 7 | 1 | 0 | 18 | 5 | -2,7 | -4,8 | 2,1 |
| chr19:58929692-58930192:downstream_14724 | chr19 | 58920062 | 58929692 | + | ZNF584 | NM | 10 | -1,2 | 1 | 1 | 3,90E-01 | NA | 11 | 7 | 10 | 3 | 20 | 15 | 6 | 5 | 0,5 | 1,7 | -1,3 |
| chr15:83659809-83660309:downstream_11858 | chr15 | 83654954 | 83659809 | + | FAM103A1 | NM | 51 | 0,8 | 1 | -1 | 3,91E-01 | 7,27E-01 | 20 | 19 | 4 | 2 | 113 | 167 | 49 | 31 | 2,6 | 1,8 | 0,7 |
| chr7:99208456-99208956:downstream_25544 | chr7 | 99195901 | 99208456 | + | GS1-259H13.2 | NM | 5 | 1,8 | 2 | -1 | 3,91E-01 | NA | 4 | 12 | 1 | 1 | 11 | 2 | 3 | 3 | 2,8 | 1,0 | 1,8 |
| chr8:35652181-35652681:downstream_6233 | chr8 | 35092974 | 35652181 | + | UNC5D | NM | 4 | 2,3 | 3 | -1 | 3,92E-01 | NA | 2 | 0 | 3 | 14 | 0 | 0 | 11 | 5 | -3,4 | -Inf | Inf |
| chr20:19983103-19983603:downstream_22536 | chr20 | 19867164 | 19983103 | + | RIN2 | NM | 23 | -1,0 | 1 | 1 | 3,92E-01 | NA | 4 | 2 | 18 | 25 | 26 | 4 | 45 | 60 | -2,8 | -1,8 | -1,0 |
| chr12:113630173-113630673:downstream_18802 | chr12 | 113623330 | 113630173 | + | RITA1 | NM | 17 | 0,9 | 1 | -1 | 3,92E-01 | NA | 15 | 27 | 10 | 9 | 17 | 22 | 17 | 16 | 1,2 | 0,3 | 0,9 |
| chr11:124506443-124506943:downstream_9726 | chr11 | 124506393 | 124506443 | + | MIR7641-2 | NR | 6 | 1,6 | 2 | -1 | 3,92E-01 | NA | 5 | 9 | 3 | 1 | 2 | 15 | 4 | 8 | 1,9 | 0,5 | 1,5 |
| chr15:25324297-25324797:downstream_11589 | chr15 | 25324203 | 25324297 | + | SNORD116-13 | NR | 130 | 0,4 | 1 | -1 | 3,92E-01 | 7,30E-01 | 228 | 205 | 138 | 137 | 96 | 84 | 85 | 71 | 0,7 | 0,2 | 0,5 |
| chr14:51724372-51724872:downstream_11135 | chr14 | 51706885 | 51724372 | + | TMX1 | NM | 10 | 1,3 | 2 | -1 | 3,93E-01 | NA | 3 | 8 | 6 | 0 | 15 | 15 | 18 | 15 | 1,0 | -0,1 | 1,1 |
| chr19:5691678-5692178:downstream_21383 | chr19 | 5690271 | 5691678 | + | RPL36 | NM | 126 | 1,0 | 1 | -1 | 3,93E-01 | 7,30E-01 | 6 | 8 | 1 | 1 | 445 | 304 | 113 | 127 | 2,6 | 1,6 | 1,0 |
| chr7:98991797-98992297:downstream_5970 | chr7 | 98992297 | 99006305 | - | PDAP1 | NM | 11 | 1,1 | 1 | -1 | 3,93E-01 | NA | 7 | 10 | 4 | 7 | 9 | 15 | 19 | 15 | 0,6 | -0,5 | 1,1 |
| chr2:15771235-15771735:downstream_1867 | chr2 | 15731744 | 15771235 | + | DDX1 | NM | 24 | 0,8 | 1 | -1 | 3,93E-01 | NA | 17 | 17 | 14 | 11 | 22 | 37 | 32 | 39 | 0,5 | -0,3 | 0,8 |
| chr5:74895646-74896146:downstream_4035 | chr5 | 74807656 | 74895646 | + | POLK | NM | 12 | -1,1 | 1 | 1 | 3,94E-01 | NA | 3 | 3 | 17 | 21 | 3 | 11 | 13 | 25 | -2,6 | -1,4 | -1,2 |
| chr12:21069843-21070343:downstream_9777 | chr12 | 20963637 | 21069843 | + | SLCO1B3 | NM | 59 | -0,6 | 1 | 1 | 3,94E-01 | 7,31E-01 | 13 | 8 | 25 | 27 | 56 | 99 | 106 | 140 | -1,3 | -0,7 | -0,7 |
| chr12:75895716-75896216:downstream_9888 | chr12 | 75874512 | 75895716 | + | GLIPR1 | NM | 20 | 0,9 | 1 | -1 | 3,94E-01 | NA | 32 | 37 | 25 | 20 | 13 | 7 | 14 | 10 | 0,6 | -0,3 | 0,9 |
| chr1:150937148-150937648:downstream_16590 | chr1 | 150937648 | 150947479 | - | CERS2 | NM | 164 | 0,5 | 1 | -1 | 3,95E-01 | 7,32E-01 | 20 | 35 | 21 | 11 | 294 | 385 | 253 | 291 | 0,8 | 0,3 | 0,5 |
| chr17:38783475-38783975:downstream_12566 | chr17 | 38783975 | 38804103 | - | SMARCE1 | NM | 55 | 0,6 | 1 | -1 | 3,95E-01 | 7,32E-01 | 12 | 20 | 17 | 10 | 89 | 80 | 99 | 110 | 0,3 | -0,3 | 0,6 |
| chr15:102172679-102173179:downstream_19811 | chr15 | 102173179 | 102192594 | - | TM2D3 | NM | 3 | 2,5 | 3 | -1 | 3,95E-01 | NA | 6 | 5 | 0 | 0 | 3 | 6 | 0 | 2 | Inf | 2,5 | Inf |
| chr5:139062680-139063180:downstream_4452 | chr5 | 139028300 | 139062680 | + | CXXC5 | NM | 32 | -0,7 | 1 | 1 | 3,96E-01 | 7,32E-01 | 17 | 28 | 43 | 57 | 15 | 33 | 32 | 34 | -1,2 | -0,4 | -0,7 |
| chr11:74551454-74551954:downstream_18394 | chr11 | 74551954 | 74660232 | - | XRRA1 | NM | 11 | 1,2 | 1 | -1 | 3,96E-01 | NA | 2 | 11 | 13 | 9 | 6 | 5 | 22 | 20 | -0,8 | -2,0 | 1,2 |
| chr6:153291157-153291657:downstream_24764 | chr6 | 153291657 | 153304740 | - | FBXO5 | NM | 5 | -1,7 | 2 | 1 | 3,97E-01 | NA | 3 | 0 | 4 | 6 | 6 | 7 | 4 | 7 | -1,6 | 0,1 | -1,7 |
| chr11:34168458-34168958:downstream_18163 | chr11 | 34127110 | 34168458 | + | NAT10 | NM | 25 | 0,9 | 1 | -1 | 3,97E-01 | NA | 8 | 10 | 3 | 6 | 33 | 59 | 42 | 43 | 1,1 | 0,1 | 0,9 |
| chr3:56653659-56654159:downstream_23142 | chr3 | 56654159 | 56717135 | - | FAM208A | NM | 32 | 1,0 | 1 | -1 | 3,97E-01 | 7,32E-01 | 1 | 4 | 3 | 7 | 15 | 26 | 114 | 82 | -1,1 | -2,2 | 1,1 |
| chr4:110350618-110351118:downstream_3809 | chr4 | 110351118 | 110354973 | - | SEC24B-AS1 | NR | 14 | 1,2 | 1 | -1 | 3,97E-01 | NA | 25 | 32 | 19 | 11 | 10 | 0 | 3 | 9 | 0,9 | -0,3 | 1,2 |
| chr16:16236930-16237430:downstream_11972 | chr16 | 16043433 | 16236930 | + | ABCC1 | NM | 75 | -1,1 | 1 | 1 | 3,98E-01 | 7,32E-01 | 0 | 2 | 8 | 4 | 61 | 81 | 203 | 240 | -2,7 | -1,6 | -1,0 |
| chr17:38544272-38544772:downstream_13048 | chr17 | 38544772 | 38574202 | - | TOP2A | NM | 45 | 0,9 | 1 | -1 | 3,98E-01 | 7,32E-01 | 5 | 0 | 7 | 12 | 27 | 15 | 145 | 153 | -2,0 | -2,8 | 0,8 |
| chr19:38122888-38123388:downstream_13770 | chr19 | 38123388 | 38146313 | - | ZFP30 | NM | 14 | 1,2 | 1 | -1 | 3,98E-01 | NA | 2 | 4 | 14 | 8 | 7 | 1 | 43 | 34 | -2,0 | -3,2 | 1,2 |
| chr7:141479188-141479688:downstream_6118 | chr7 | 141478288 | 141479188 | + | TAS2R4 | NM | 11 | 1,2 | 1 | -1 | 3,98E-01 | NA | 18 | 10 | 15 | 4 | 11 | 4 | 15 | 7 | 0,5 | -0,7 | 1,2 |
| chr1:6281359-6281859:downstream_403 | chr1 | 6266188 | 6281359 | + | RNF207 | NM | 9 | 1,2 | 1 | -1 | 3,99E-01 | NA | 7 | 16 | 3 | 11 | 8 | 8 | 13 | 9 | 0,8 | -0,4 | 1,3 |
| chr4:90164928-90165428:downstream_3382 | chr4 | 90165428 | 90229161 | - | GPRIN3 | NM | 32 | -0,8 | 1 | 1 | 3,99E-01 | 7,34E-01 | 11 | 3 | 49 | 70 | 11 | 11 | 39 | 60 | -3,1 | -2,2 | -0,8 |
| chrX:96855597-96856097:downstream_26786 | chrX | 95939661 | 96855597 | + | DIAPH2 | NM | 18 | -1,0 | 1 | 1 | 4,00E-01 | NA | 3 | 1 | 28 | 31 | 6 | 5 | 29 | 41 | -3,8 | -2,7 | -1,1 |
| chr1:205681446-205681946:downstream_1330 | chr1 | 205681946 | 205719372 | - | NUCKS1 | NM | 16 | -1,1 | 1 | 1 | 4,00E-01 | NA | 2 | 4 | 4 | 14 | 12 | 26 | 26 | 37 | -1,7 | -0,7 | -1,0 |
| chr7:6448246-6448746:downstream_25478 | chr7 | 6448746 | 6487643 | - | DAGLB | NM | 12 | 1,1 | 1 | -1 | 4,00E-01 | NA | 6 | 13 | 4 | 11 | 15 | 6 | 18 | 21 | 0,3 | -0,9 | 1,2 |
| chr2:36778278-36778778:downstream_1516 | chr2 | 36583369 | 36778278 | + | CRIM1 | NM | 71 | -0,9 | 1 | 1 | 4,01E-01 | 7,36E-01 | 2 | 1 | 8 | 9 | 67 | 79 | 152 | 246 | -2,4 | -1,4 | -0,9 |
| chr20:826922-827422:downstream_22424 | chr20 | 814339 | 826922 | + | FAM110A | NM | 3 | -2,2 | 3 | 1 | 4,01E-01 | NA | 7 | 3 | 0 | 4 | 4 | 7 | 0 | 1 | 1,3 | 3,7 | -2,4 |
| chr6:63985355-63985855:downstream_24840 | chr6 | 63985855 | 64029882 | - | LGSN | NM | 27 | -0,9 | 1 | 1 | 4,01E-01 | NA | 9 | 0 | 21 | 27 | 15 | 26 | 54 | 61 | -2,4 | -1,5 | -1,0 |
| chr12:53584654-53585154:downstream_10201 | chr12 | 53574534 | 53584654 | + | ZNF740 | NM | 14 | -1,0 | 1 | 1 | 4,01E-01 | NA | 4 | 7 | 13 | 16 | 19 | 12 | 22 | 21 | -1,4 | -0,5 | -0,9 |
| chr16:58554431-58554931:downstream_20204 | chr16 | 58549382 | 58554431 | + | SETD6 | NM | 21 | -0,9 | 1 | 1 | 4,02E-01 | NA | 15 | 14 | 24 | 18 | 15 | 40 | 31 | 15 | -0,6 | 0,3 | -0,8 |
| chr11:118473747-118474247:downstream_17872 | chr11 | 118443101 | 118473747 | + | ARCN1 | NM | 23 | 0,9 | 1 | -1 | 4,02E-01 | NA | 4 | 11 | 4 | 10 | 33 | 20 | 36 | 63 | 0,1 | -0,9 | 1,0 |
| chr9:123714113-123714613:downstream_6825 | chr9 | 123714613 | 123812554 | - | C5 | NM | 78 | -1,0 | 1 | 1 | 4,02E-01 | 7,38E-01 | 2 | 0 | 10 | 5 | 110 | 52 | 213 | 230 | -2,6 | -1,4 | -1,1 |
| chr6:111349466-111349966:downstream_25177 | chr6 | 111303219 | 111349466 | + | RPF2 | NM | 19 | 1,0 | 1 | -1 | 4,02E-01 | NA | 5 | 9 | 0 | 5 | 33 | 38 | 19 | 41 | 1,5 | 0,2 | 1,2 |
| chr19:23406106-23406606:downstream_14174 | chr19 | 23406606 | 23433172 | - | ZNF724P | NR | 7 | 1,9 | 2 | -1 | 4,02E-01 | NA | 6 | 0 | 19 | 11 | 0 | 1 | 11 | 6 | -2,4 | -4,1 | 1,7 |
| chr9:104234939-104235439:downstream_26539 | chr9 | 104235439 | 104249503 | - | TMEM246 | NM | 9 | 1,3 | 2 | -1 | 4,02E-01 | NA | 7 | 3 | 15 | 8 | 5 | 0 | 18 | 13 | -1,3 | -2,7 | 1,4 |
| chr5:180274110-180274610:downstream_24447 | chr5 | 180274610 | 180288286 | - | ZFP62 | NM | 26 | 0,8 | 1 | -1 | 4,03E-01 | NA | 15 | 29 | 18 | 23 | 31 | 16 | 35 | 42 | 0,1 | -0,7 | 0,8 |
| chr2:120439694-120440194:downstream_22287 | chr2 | 120436742 | 120439694 | + | TMEM177 | NM | 28 | 0,8 | 1 | -1 | 4,03E-01 | 7,38E-01 | 13 | 10 | 8 | 5 | 52 | 44 | 38 | 55 | 0,8 | 0,1 | 0,7 |
| chrX:13787480-13787980:downstream_7474 | chrX | 13752831 | 13787480 | + | OFD1 | NM | 10 | 1,2 | 1 | -1 | 4,03E-01 | NA | 5 | 21 | 11 | 13 | 7 | 2 | 13 | 7 | 0,1 | -1,2 | 1,3 |
| chr12:66737423-66737923:downstream_9876 | chr12 | 66696334 | 66737423 | + | HELB | NM | 8 | 1,3 | 2 | -1 | 4,04E-01 | NA | 9 | 7 | 10 | 17 | 4 | 0 | 11 | 7 | -0,8 | -2,2 | 1,4 |
| chr16:107669-108169:downstream_12049 | chr16 | 103828 | 107669 | + | SNRNP25 | NM | 5 | 1,7 | 2 | -1 | 4,05E-01 | NA | 7 | 4 | 3 | 2 | 8 | 2 | 11 | 2 | 1,3 | -0,3 | 1,7 |
| chr6:134308638-134309138:downstream_25296 | chr6 | 134273307 | 134308638 | + | TBPL1 | NM | 9 | 1,2 | 1 | -1 | 4,05E-01 | NA | 3 | 8 | 7 | 4 | 11 | 5 | 17 | 20 | 0,0 | -1,2 | 1,2 |
| chr9:140341522-140342022:downstream_26418 | chr9 | 140342022 | 140353786 | - | NSMF | NM | 10 | 1,3 | 2 | -1 | 4,05E-01 | NA | 29 | 23 | 3 | 2 | 12 | 7 | 6 | 0 | 3,3 | 1,8 | 1,6 |
| chr7:44842722-44843222:downstream_25688 | chr7 | 44836234 | 44842722 | + | PPIA | NM | 24 | 0,9 | 1 | -1 | 4,07E-01 | NA | 7 | 11 | 3 | 6 | 37 | 50 | 28 | 52 | 1,1 | 0,1 | 0,9 |
| chr11:118894385-118894885:downstream_9687 | chr11 | 118889240 | 118894385 | + | TRAPPC4 | NM | 25 | 0,8 | 1 | -1 | 4,07E-01 | NA | 8 | 9 | 7 | 4 | 43 | 38 | 40 | 49 | 0,6 | -0,1 | 0,8 |
| chr1:24999772-25000272:downstream_17277 | chr1 | 24969593 | 24999772 | + | SRRM1 | NM | 21 | -1,0 | 1 | 1 | 4,08E-01 | NA | 5 | 0 | 8 | 4 | 36 | 34 | 40 | 41 | -1,4 | -0,2 | -1,1 |
| chr3:39454032-39454532:downstream_23397 | chr3 | 39448179 | 39454032 | + | RPSA | NM | 48 | -0,6 | 1 | 1 | 4,08E-01 | 7,47E-01 | 39 | 58 | 32 | 25 | 67 | 101 | 18 | 43 | 0,8 | 1,5 | -0,7 |
| chr19:58376491-58376991:downstream_21686 | chr19 | 58361180 | 58376491 | + | ZNF587 | NM | 6 | 1,4 | 2 | -1 | 4,09E-01 | NA | 2 | 8 | 4 | 8 | 3 | 3 | 11 | 9 | -0,3 | -1,7 | 1,4 |

| | | | | | | | | | | | | | | | | | | | | | | | |
|---|---|---|---|---|---|---|---|---|---|---|---|---|---|---|---|---|---|---|---|---|---|---|---|
| chr19:37883966-37884466:downstream_13768 | chr19 | 37862058 | 37883966 | + | ZNF527 | NM | 5 | 1,7 | 2 | -1 | 4,09E-01 | NA | 4 | 0 | 7 | 9 | 2 | 0 | 14 | 8 | -2,0 | -3,8 | 1,8 |
| chr10:75879914-75880414:downstream_17814 | chr10 | 75757871 | 75879914 | + | VCL | NM | 94 | 2,0 | 2 | -1 | 4,09E-01 | 7,48E-01 | 0 | 10 | 0 | 4 | 113 | 143 | 230 | 250 | 1,2 | -0,9 | 2,1 |
| chr14:45584301-45584801:downstream_11124 | chr14 | 45584801 | 45603732 | - | FKBP3 | NM | 21 | 0,8 | 1 | -1 | 4,09E-01 | NA | 11 | 15 | 7 | 9 | 33 | 27 | 24 | 40 | 0,7 | -0,1 | 0,8 |
| chr22:39129592-39130092:downstream_15610 | chr22 | 39101806 | 39129592 | + | GTPBP1 | NM | 7 | 1,3 | 2 | -1 | 4,10E-01 | NA | 15 | 11 | 1 | 4 | 7 | 12 | 1 | 8 | 2,2 | 0,9 | 1,3 |
| chr8:76134851-76135351:downstream_25875 | chr8 | 76135351 | 76191124 | - | CASC9 | NR | 17 | -1,3 | 2 | 1 | 4,10E-01 | NA | 1 | 2 | 4 | 7 | 41 | 5 | 33 | 45 | -2,0 | -0,8 | -1,2 |
| chr11:67131138-67131638:downstream_17957 | chr11 | 67131638 | 67141648 | - | CLCF1 | NM | 29 | -0,7 | 1 | 1 | 4,10E-01 | 7,49E-01 | 11 | 7 | 15 | 13 | 53 | 43 | 46 | 43 | -0,6 | 0,1 | -0,7 |
| chr6:151791234-151791734:downstream_24515 | chr6 | 151773392 | 151791234 | + | ARMT1 | NM | 21 | 0,9 | 1 | -1 | 4,11E-01 | NA | 15 | 4 | 13 | 6 | 27 | 22 | 42 | 39 | 0,1 | -0,7 | 0,8 |
| chr4:177253396-177253896:downstream_3913 | chr4 | 177241089 | 177253396 | + | SPCS3 | NM | 27 | -0,8 | 1 | 1 | 4,11E-01 | NA | 5 | 6 | 24 | 14 | 28 | 27 | 45 | 67 | -1,8 | -1,0 | -0,8 |
| chr12:59265436-59265936:downstream_18662 | chr12 | 59265936 | 59314319 | - | LRIG3 | NM | 15 | 1,1 | 1 | -1 | 4,12E-01 | NA | 1 | 5 | 15 | 16 | 2 | 5 | 40 | 36 | -2,5 | -3,6 | 1,1 |
| chr20:34541038-34541538:downstream_22542 | chr20 | 34541538 | 34543281 | - | SCAND1 | NM | 20 | 1,1 | 1 | -1 | 4,12E-01 | NA | 13 | 6 | 3 | 1 | 46 | 56 | 11 | 25 | 2,4 | 1,5 | 0,9 |
| chr1:230829731-230830231:downstream_16615 | chr1 | 230778201 | 230829731 | + | COG2 | NM | 26 | 0,9 | 1 | -1 | 4,12E-01 | NA | 9 | 2 | 8 | 2 | 38 | 36 | 78 | 39 | 0,1 | -0,7 | 0,8 |
| chr15:25344742-25345242:downstream_11567 | chr15 | 25344644 | 25344742 | + | SNORD116-26 | NR | 27 | 0,7 | 1 | -1 | 4,12E-01 | NA | 30 | 35 | 28 | 35 | 18 | 15 | 26 | 26 | 0,0 | -0,7 | 0,7 |
| chr9:38570860-38571360:downstream_7083 | chr9 | 38571360 | 38620360 | - | ANKRD18A | NM | 11 | 1,1 | 1 | -1 | 4,12E-01 | NA | 6 | 13 | 10 | 11 | 3 | 12 | 13 | 22 | -0,2 | -1,2 | 1,1 |
| chr17:42981047-42981547:downstream_20380 | chr17 | 42977079 | 42981047 | + | CCDC103 | NM | 6 | -1,6 | 2 | 1 | 4,13E-01 | NA | 2 | 0 | 3 | 10 | 6 | 3 | 10 | 11 | -3,0 | -1,3 | -1,7 |
| chr2:120196096-120196596:downstream_1706 | chr2 | 120189445 | 120196096 | + | TMEM37 | NM | 19 | -1,0 | 1 | 1 | 4,13E-01 | NA | 1 | 2 | 14 | 11 | 14 | 9 | 54 | 49 | -3,2 | -2,2 | -1,0 |
| chr13:107195161-107195661:downstream_10628 | chr13 | 107195661 | 107220514 | - | ARGLU1 | NM | 24 | 0,8 | 1 | -1 | 4,13E-01 | NA | 9 | 15 | 13 | 10 | 24 | 33 | 47 | 44 | 0,1 | -0,7 | 0,7 |
| chr5:77780537-77781037:downstream_4039 | chr5 | 77781037 | 77944648 | - | LHFPL2 | NM | 22 | -1,5 | 2 | 1 | 4,14E-01 | NA | 0 | 0 | 6 | 7 | 7 | 3 | 79 | 73 | -Inf | -3,9 | -Inf |
| chr12:122079946-122080446:downstream_10460 | chr12 | 122064454 | 122079946 | + | ORAI1 | NM | 6 | -1,3 | 2 | 1 | 4,14E-01 | NA | 8 | 5 | 13 | 8 | 5 | 5 | 3 | 3 | -0,7 | 0,7 | -1,4 |
| chr2:20649204-20649704:downstream_1880 | chr2 | 20646831 | 20649204 | + | RHOB | NM | 62 | -0,7 | 1 | 1 | 4,15E-01 | 7,57E-01 | 22 | 9 | 10 | 12 | 109 | 193 | 64 | 74 | 0,5 | 1,1 | -0,7 |
| chr14:57879466-57879966:downstream_11159 | chr14 | 57857270 | 57879466 | + | NAA30 | NM | 24 | 0,8 | 1 | -1 | 4,15E-01 | NA | 6 | 17 | 10 | 10 | 26 | 33 | 47 | 41 | 0,2 | -0,6 | 0,8 |
| chrX:106304617-106305117:downstream_27002 | chrX | 106305117 | 106362057 | - | RBM41 | NM | 11 | 1,0 | 1 | -1 | 4,15E-01 | NA | 7 | 8 | 8 | 8 | 9 | 8 | 24 | 14 | -0,1 | -1,2 | 1,0 |
| chr19:39923346-39923846:downstream_14282 | chr19 | 39923846 | 39926618 | - | RPS16 | NM | 51 | 1,0 | 1 | -1 | 4,16E-01 | 7,58E-01 | 7 | 5 | 4 | 0 | 168 | 85 | 63 | 77 | 1,4 | 0,9 | 0,6 |
| chr7:134364567-134365067:downstream_25409 | chr7 | 134331530 | 134364567 | + | BPGM | NM | 10 | 1,3 | 2 | -1 | 4,16E-01 | NA | 1 | 9 | 4 | 2 | 17 | 11 | 26 | 13 | 0,7 | -0,5 | 1,2 |
| chr19:55912149-55912649:downstream_14658 | chr19 | 55912649 | 55919325 | - | UBE2S | NM | 10 | 1,1 | 1 | -1 | 4,16E-01 | NA | 10 | 15 | 3 | 7 | 15 | 9 | 11 | 9 | 1,4 | 0,3 | 1,2 |
| chr5:137805004-137805504:downstream_4444 | chr5 | 137801180 | 137805004 | + | EGR1 | NM | 13 | -1,9 | 2 | 1 | 4,17E-01 | NA | 8 | 14 | 0 | 0 | 32 | 49 | 0 | 0 | Inf | Inf | NaN |
| chr11:77376773-77377273:downstream_8927 | chr11 | 77377273 | 77531880 | - | RSF1 | NM | 37 | -0,8 | 1 | 1 | 4,18E-01 | 7,60E-01 | 3 | 5 | 42 | 34 | 10 | 22 | 107 | 71 | -3,2 | -2,5 | -0,8 |
| chr3:129024135-129024635:downstream_23194 | chr3 | 128997683 | 129024135 | + | HMCES | NM | 10 | 1,2 | 1 | -1 | 4,18E-01 | NA | 2 | 9 | 8 | 7 | 10 | 3 | 18 | 24 | -0,5 | -1,7 | 1,2 |
| chr1:150670034-150670534:downstream_16805 | chr1 | 150670534 | 150693364 | - | HORMAD1 | NM | 31 | -0,7 | 1 | 1 | 4,18E-01 | 7,60E-01 | 7 | 9 | 75 | 60 | 8 | 8 | 43 | 39 | -3,1 | -2,4 | -0,7 |
| chr19:38899728-38900228:downstream_14261 | chr19 | 38893774 | 38899728 | + | FAM98C | NM | 9 | 1,2 | 1 | -1 | 4,19E-01 | NA | 9 | 6 | 1 | 5 | 15 | 11 | 10 | 16 | 1,2 | 0,0 | 1,2 |
| chr16:3271824-3272324:downstream_20283 | chr16 | 3272324 | 3285457 | - | ZNF200 | NM | 6 | 1,5 | 2 | -1 | 4,19E-01 | NA | 4 | 10 | 0 | 3 | 9 | 7 | 3 | 9 | 2,1 | 0,4 | 1,7 |
| chr17:19807264-19807764:downstream_12897 | chr17 | 19807764 | 19881169 | - | AKAP10 | NM | 8 | 1,4 | 2 | -1 | 4,19E-01 | NA | 7 | 1 | 11 | 7 | 6 | 0 | 11 | 24 | -1,2 | -2,6 | 1,4 |
| chr15:48431124-48431624:downstream_19639 | chr15 | 48431624 | 48470558 | - | MYEF2 | NM | 14 | -1,1 | 1 | 1 | 4,19E-01 | NA | 4 | 0 | 11 | 23 | 7 | 9 | 25 | 36 | -3,1 | -1,9 | -1,1 |
| chr9:135565470-135565970:downstream_7371 | chr9 | 135545727 | 135565470 | + | GTF3C4 | NM | 25 | 0,7 | 1 | -1 | 4,19E-01 | NA | 26 | 30 | 14 | 22 | 34 | 18 | 29 | 28 | 0,6 | -0,1 | 0,8 |
| chr4:148556672-148557172:downstream_3866 | chr4 | 148538538 | 148556672 | + | TMEM184C | NM | 41 | 0,7 | 1 | -1 | 4,20E-01 | 7,62E-01 | 77 | 44 | 70 | 35 | 32 | 11 | 28 | 31 | 0,2 | -0,5 | 0,7 |
| chr6:155155378-155155878:downstream_25189 | chr6 | 155054511 | 155155378 | + | SCAF8 | NM | 20 | -1,0 | 1 | 1 | 4,20E-01 | NA | 1 | 3 | 8 | 6 | 14 | 37 | 38 | 57 | -1,9 | -0,9 | -1,0 |
| chr8:96168913-96169413:downstream_6648 | chr8 | 96145948 | 96168913 | + | PLEKHF2 | NM | 10 | 1,1 | 1 | -1 | 4,20E-01 | NA | 7 | 17 | 1 | 7 | 15 | 11 | 8 | 13 | 1,5 | 0,3 | 1,2 |
| chrX:100882831-100883331:downstream_26623 | chrX | 100878119 | 100882831 | + | ARMCX3 | NM | 27 | 0,8 | 1 | -1 | 4,20E-01 | NA | 13 | 15 | 14 | 7 | 24 | 48 | 47 | 44 | 0,4 | -0,3 | 0,7 |
| chr11:62506108-62506608:downstream_9397 | chr11 | 62495951 | 62506108 | + | TTC9C | NM | 28 | 0,7 | 1 | -1 | 4,20E-01 | 7,62E-01 | 26 | 27 | 31 | 46 | 17 | 12 | 29 | 38 | -0,5 | -1,2 | 0,7 |
| chr12:10977444-10977944:downstream_10052 | chr12 | 10977944 | 10978868 | - | TAS2R10 | NM | 6 | 1,6 | 2 | -1 | 4,20E-01 | NA | 2 | 5 | 10 | 11 | 2 | 0 | 10 | 7 | -1,7 | -3,4 | 1,7 |
| chr3:9519838-9520338:downstream_23415 | chr3 | 9439383 | 9519838 | + | SETD5 | NM | 16 | 1,0 | 1 | -1 | 4,21E-01 | NA | 2 | 8 | 3 | 7 | 7 | 25 | 42 | 36 | -0,1 | -1,3 | 1,1 |
| chr3:105376608-105377108:downstream_2660 | chr3 | 105377108 | 105587887 | - | CBLB | NM | 5 | 2,1 | 3 | -1 | 4,21E-01 | NA | 1 | 0 | 8 | 7 | 0 | 0 | 8 | 16 | -4,2 | -Inf | Inf |
| chr15:86292589-86293089:downstream_19458 | chr15 | 85923846 | 86292589 | + | AKAP13 | NM | 12 | -1,6 | 2 | 1 | 4,22E-01 | NA | 0 | 0 | 4 | 10 | 2 | 3 | 39 | 41 | -Inf | -3,9 | -Inf |
| chr3:10195354-10195854:downstream_23504 | chr3 | 10183318 | 10195354 | + | VHL | NM | 32 | 0,8 | 1 | -1 | 4,22E-01 | 7,62E-01 | 9 | 9 | 6 | 3 | 51 | 75 | 46 | 57 | 1,0 | 0,3 | 0,7 |
| chr12:62803501-62804001:downstream_18904 | chr12 | 62654120 | 62803501 | + | USP15 | NM | 5 | -1,6 | 2 | 1 | 4,22E-01 | NA | 2 | 0 | 10 | 3 | 7 | 2 | 7 | 13 | -3,0 | -1,3 | -1,8 |
| chr12:14923153-14923653:downstream_10081 | chr12 | 14923653 | 14924065 | - | HIST4H4 | NM | 111 | 0,5 | 1 | -1 | 4,22E-01 | 7,62E-01 | 98 | 130 | 57 | 59 | 188 | 132 | 99 | 126 | 1,0 | 0,5 | 0,5 |
| chr9:100618997-100619497:downstream_7219 | chr9 | 100615536 | 100618997 | + | FOXE1 | NM | 6 | 1,5 | 2 | -1 | 4,23E-01 | NA | 3 | 9 | 1 | 6 | 11 | 0 | 10 | 10 | 0,7 | -0,9 | 1,6 |
| chr18:56417371-56417871:downstream_20888 | chr18 | 56338617 | 56417371 | + | MALT1 | NM | 24 | 0,8 | 1 | -1 | 4,23E-01 | NA | 2 | 10 | 33 | 30 | 7 | 5 | 49 | 59 | -2,4 | -3,2 | 0,9 |
| chr4:152025804-152026304:downstream_23832 | chr4 | 152020724 | 152025804 | + | RPS3A | NM | 59 | 0,5 | 1 | -1 | 4,23E-01 | 7,62E-01 | 105 | 113 | 25 | 25 | 82 | 69 | 29 | 22 | 2,1 | 1,6 | 0,6 |
| chr17:16344818-16345318:downstream_20561 | chr17 | 16345318 | 16395505 | - | LRRC75A | NM | 33 | 0,8 | 1 | -1 | 4,23E-01 | 7,62E-01 | 44 | 84 | 17 | 19 | 15 | 51 | 11 | 20 | 1,8 | 1,1 | 0,8 |
| chr15:79677061-79677561:downstream_19813 | chr15 | 79603403 | 79677061 | + | TMED3 | NM | 6 | -1,5 | 2 | 1 | 4,23E-01 | NA | 2 | 1 | 7 | 4 | 13 | 2 | 14 | 9 | -2,1 | -0,6 | -1,5 |
| chr11:82534908-82535408:downstream_18226 | chr11 | 82535408 | 82611557 | - | PRCP | NM | 109 | -0,6 | 1 | 1 | 4,23E-01 | 7,62E-01 | 15 | 6 | 22 | 15 | 208 | 174 | 174 | 256 | -0,8 | -0,2 | -0,6 |
| chr18:3597377-3597877:downstream_13543 | chr18 | 3594111 | 3597377 | + | DLGAP1-AS1 | NR | 36 | -0,6 | 1 | 1 | 4,23E-01 | 7,62E-01 | 18 | 11 | 39 | 33 | 34 | 40 | 49 | 69 | -1,3 | -0,7 | -0,7 |
| chr15:89021861-89022361:downstream_19637 | chr15 | 89010683 | 89021861 | + | MRPS11 | NM | 19 | -0,8 | 1 | 1 | 4,24E-01 | NA | 18 | 9 | 11 | 15 | 26 | 40 | 17 | 20 | 0,0 | 0,9 | -0,8 |
| chr19:17390162-17390662:downstream_21019 | chr19 | 17378184 | 17390162 | + | BABAM1 | NM | 10 | 1,4 | 2 | -1 | 4,24E-01 | NA | 1 | 9 | 1 | 1 | 26 | 18 | 11 | 15 | 2,1 | 0,8 | 1,3 |
| chr19:40574558-40575058:downstream_21702 | chr19 | 40575058 | 40596845 | - | ZNF780A | NM | 40 | -0,7 | 1 | 1 | 4,24E-01 | 7,63E-01 | 8 | 7 | 14 | 23 | 47 | 56 | 77 | 89 | -1,3 | -0,7 | -0,6 |
| chrX:57145614-57146114:downstream_27038 | chrX | 57146114 | 57147989 | - | SPIN2B | NM | 10 | -1,1 | 1 | 1 | 4,24E-01 | NA | 13 | 12 | 15 | 19 | 3 | 11 | 7 | 2 | -0,5 | 0,6 | -1,0 |
| chr6:43489246-43489746:downstream_5152 | chr6 | 43484776 | 43489246 | + | POLR1C | NM | 19 | 1,1 | 1 | -1 | 4,25E-01 | NA | 24 | 3 | 1 | 2 | 59 | 42 | 10 | 15 | 3,2 | 2,0 | 1,1 |
| chr2:27004099-27004599:downstream_1589 | chr2 | 26987141 | 27004099 | + | SLC35F6 | NM | 102 | 0,4 | 1 | -1 | 4,25E-01 | 7,64E-01 | 82 | 85 | 125 | 89 | 84 | 76 | 128 | 148 | -0,4 | -0,8 | 0,4 |
| chr1:85730959-85731459:downstream_808 | chr1 | 85731459 | 85742587 | - | BCL10 | NM | 18 | 0,9 | 1 | -1 | 4,25E-01 | NA | 31 | 22 | 3 | 3 | 38 | 30 | 10 | 6 | 3,1 | 2,1 | 1,0 |
| chr2:27599597-27600097:downstream_22337 | chr2 | 27600097 | 27603611 | - | ZNF513 | NM | 12 | -1,0 | 1 | 1 | 4,27E-01 | NA | 8 | 5 | 8 | 7 | 15 | 30 | 7 | 19 | -0,3 | 0,8 | -1,1 |
| chr1:110750572-110751072:downstream_888 | chr1 | 110751072 | 110752609 | - | KCNC4-AS1 | NR | 3 | -2,6 | 3 | 1 | 4,27E-01 | NA | 8 | 0 | 0 | 2 | 6 | 5 | 0 | 0 | 2,3 | Inf | -Inf |
| chr3:197682721-197683221:downstream_3360 | chr3 | 197677051 | 197682721 | + | RPL35A | NM | 32 | 0,7 | 1 | -1 | 4,28E-01 | 7,67E-01 | 50 | 26 | 7 | 14 | 55 | 52 | 18 | 31 | 1,9 | 1,1 | 0,7 |
| chr2:217366188-217366688:downstream_2408 | chr2 | 217363519 | 217366188 | + | RPL37A | NM | 229 | 0,4 | 1 | -1 | 4,28E-01 | 7,67E-01 | 122 | 144 | 56 | 55 | 406 | 533 | 224 | 290 | 1,3 | 0,9 | 0,4 |
| chr19:41830788-41831288:downstream_14314 | chr19 | 41816093 | 41830788 | + | CCDC97 | NM | 16 | 1,1 | 1 | -1 | 4,28E-01 | NA | 2 | 23 | 4 | 2 | 39 | 28 | 11 | 21 | 2,1 | 1,0 | 1,0 |
| chr11:47260769-47261269:downstream_17980 | chr11 | 47236492 | 47260769 | + | DDB2 | NM | 36 | 0,7 | 1 | -1 | 4,29E-01 | 7,67E-01 | 11 | 14 | 3 | 5 | 75 | 90 | 53 | 40 | 1,6 | 0,8 | 0,8 |
| chr19:4044715-4045215:downstream_13715 | chr19 | 4045215 | 4066816 | - | ZBTB7A | NM | 8 | 1,2 | 1 | -1 | 4,29E-01 | NA | 9 | 14 | 6 | 7 | 3 | 8 | 10 | 5 | 0,8 | -0,4 | 1,2 |
| chr3:45722755-45723255:downstream_2914 | chr3 | 45636322 | 45722755 | + | LIMD1 | NM | 6 | 1,4 | 2 | -1 | 4,29E-01 | NA | 2 | 5 | 10 | 5 | 4 | 0 | 7 | 16 | -1,0 | -2,5 | 1,5 |
| chr5:102539224-102539724:downstream_24307 | chr5 | 102455957 | 102539224 | + | PPIP5K2 | NM | 13 | 1,1 | 1 | -1 | 4,29E-01 | NA | 8 | 1 | 22 | 11 | 6 | 1 | 29 | 23 | -1,9 | -3,0 | 1,1 |
| chr14:93653431-93653931:downstream_19422 | chr14 | 93651295 | 93653431 | + | TMEM251 | NM | 34 | 0,7 | 1 | -1 | 4,29E-01 | 7,68E-01 | 33 | 26 | 24 | 10 | 34 | 61 | 33 | 50 | 0,8 | 0,2 | 0,6 |
| chr8:49829738-49830238:downstream_6551 | chr8 | 49830238 | 49833999 | - | SNAI2 | NM | 3 | 2,2 | 3 | -1 | 4,30E-01 | NA | 7 | 6 | 0 | 0 | 11 | 2 | 0 | 2 | Inf | 2,9 | Inf |
| chr15:51739420-51739920:downstream_19528 | chr15 | 51739920 | 51914967 | - | DMXL2 | NM | 11 | 1,2 | 2 | -1 | 4,30E-01 | NA | 3 | 1 | 13 | 7 | 5 | 1 | 25 | 34 | -2,2 | -3,3 | 1,2 |
| chr3:15708243-15708743:downstream_23001 | chr3 | 15708743 | 15901053 | - | ANKRD28 | NM | 11 | -1,3 | 2 | 1 | 4,30E-01 | NA | 1 | 1 | 17 | 12 | 2 | 5 | 28 | 21 | -4,0 | -2,7 | -1,3 |
| chr6:28127250-28127750:downstream_25358 | chr6 | 28109687 | 28127250 | + | ZKSCAN8 | NM | 7 | 1,4 | 2 | -1 | 4,31E-01 | NA | 7 | 2 | 3 | 0 | 17 | 10 | 13 | 7 | 1,6 | 0,4 | 1,2 |
| chrX:77160881-77161381:downstream_7816 | chrX | 77154960 | 77160881 | + | COX7B | NM | 99 | 0,7 | 1 | -1 | 4,31E-01 | 7,68E-01 | 5 | 12 | 3 | 5 | 246 | 180 | 145 | 197 | 1,1 | 0,3 | 0,8 |
| chr14:24787242-24787742:downstream_19294 | chr14 | 24780704 | 24787242 | + | LTB4R | NM | 11 | 1,1 | 1 | -1 | 4,32E-01 | NA | 2 | 9 | 3 | 2 | 22 | 17 | 11 | 22 | 1,3 | 0,2 | 1,1 |
| chr11:65621784-65622284:downstream_9447 | chr11 | 65622284 | 65625804 | - | CFL1 | NM | 308 | 0,5 | 1 | -1 | 4,32E-01 | 7,68E-01 | 15 | 25 | 8 | 4 | 899 | 787 | 323 | 401 | 1,7 | 1,2 | 0,5 |
| chr3:107530176-107530676:downstream_23020 | chr3 | 107241782 | 107530176 | + | BBX | NM | 5 | -2,0 | 3 | 1 | 4,32E-01 | NA | 0 | 0 | 6 | 10 | 0 | 2 | 4 | 17 | -Inf | -3,5 | -Inf |
| chr1:62917475-62917975:downstream_17385 | chr1 | 62901974 | 62917475 | + | USP1 | NM | 35 | 0,8 | 1 | -1 | 4,32E-01 | 7,68E-01 | 7 | 2 | 6 | 3 | 48 | 48 | 103 | 63 | -0,1 | -0,8 | 0,7 |
| chr11:130745265-130745765:downstream_18310 | chr11 | 130745765 | 130786382 | - | SNX19 | NM | 28 | 0,8 | 1 | -1 | 4,32E-01 | 7,68E-01 | 2 | 14 | 4 | 10 | 43 | 28 | 61 | 66 | 0,1 | -0,8 | 0,9 |
| chr1:32697205-32697705:downstream_610 | chr1 | 32687970 | 32697205 | + | EIF3I | NM | 77 | -0,7 | 1 | 1 | 4,33E-01 | 7,68E-01 | 7 | 2 | 3 | 5 | 210 | 173 | 95 | 121 | 0,1 | 0,8 | -0,7 |
| chr7:6443598-6444098:downstream_25702 | chr7 | 6414125 | 6443598 | + | RAC1 | NM | 56 | -0,8 | 1 | 1 | 4,33E-01 | 7,68E-01 | 5 | 0 | 10 | 4 | 88 | 87 | 131 | 123 | -1,5 | -0,5 | -1,0 |
| chr11:64011450-64011950:downstream_9419 | chr11 | 64011950 | 64014413 | - | PPP1R14B | NM | 11 | 1,2 | 2 | -1 | 4,34E-01 | NA | 6 | 3 | 4 | 0 | 25 | 14 | 15 | 21 | 1,0 | 0,1 | 0,9 |
| chr17:6546132-6546632:downstream_12594 | chr17 | 6546632 | 6554954 | - | MED31 | NM | 20 | -0,8 | 1 | 1 | 4,34E-01 | NA | 37 | 24 | 17 | 21 | 17 | 27 | 3 | 11 | 0,7 | 1,6 | -0,9 |
| chr22:38879452-38879952:downstream_22828 | chr22 | 38864066 | 38879452 | + | KDELR3 | NM | 23 | 0,9 | 1 | -1 | 4,34E-01 | NA | 11 | 43 | 21 | 18 | 33 | 5 | 31 | 22 | 0,4 | -0,5 | 0,9 |
| chr13:52607736-52608236:downstream_10766 | chr13 | 52598826 | 52607736 | + | UTP14C | NM | 10 | 1,2 | 1 | -1 | 4,34E-01 | NA | 7 | 1 | 3 | 7 | 11 | 5 | 26 | 18 | -0,3 | -1,5 | 1,2 |
| chr17:43229468-43229968:downstream_13173 | chr17 | 43224683 | 43229468 | + | HEXIM1 | NM | 69 | -0,8 | 1 | 1 | 4,34E-01 | 7,70E-01 | 4 | 5 | 1 | 4 | 150 | 236 | 81 | 74 | 0,7 | 1,3 | -0,6 |
| chr9:136207250-136207750:downstream_26392 | chr9 | 136207750 | 136215011 | - | MED22 | NM | 7 | 1,2 | 2 | -1 | 4,34E-01 | NA | 2 | 7 | 3 | 5 | 7 | 6 | 15 | 11 | 0,3 | -1,0 | 1,3 |
| chr1:180941349-180941849:downstream_17291 | chr1 | 180941849 | 180992257 | - | STX6 | NM | 9 | 1,2 | 2 | -1 | 4,36E-01 | NA | 7 | 7 | 3 | 2 | 25 | 8 | 18 | 6 | 1,7 | 0,5 | 1,2 |
| chr11:123594134-123594634:downstream_18404 | chr11 | 123594634 | 123612391 | - | ZNF202 | NM | 10 | 1,1 | 1 | -1 | 4,36E-01 | NA | 27 | 18 | 4 | 2 | 10 | 14 | 3 | 4 | 2,8 | 1,8 | 1,0 |
| chr11:95565543-95566043:downstream_18155 | chr11 | 95566043 | 95657371 | - | MTMR2 | NM | 7 | -1,3 | 2 | 1 | 4,36E-01 | NA | 0 | 7 | 8 | 11 | 5 | 10 | 10 | 8 | -1,6 | -0,3 | -1,2 |
| chrX:64731961-64732461:downstream_26879 | chrX | 64732461 | 64754686 | - | LAS1L | NM | 18 | -1,0 | 1 | 1 | 4,36E-01 | NA | 6 | 0 | 7 | 4 | 36 | 30 | 26 | 34 | -1,0 | 0,1 | -1,1 |
| chr17:7759417-7759917:downstream_12812 | chr17 | 7758383 | 7759417 | + | TMEM88 | NM | 4 | 2,0 | 3 | -1 | 4,37E-01 | NA | 3 | 9 | 0 | 0 | 8 | 10 | 3 | 0 | Inf | 2,7 | Inf |
| chr10:129894424-129894924:downstream_17655 | chr10 | 129894924 | 129924468 | - | MKI67 | NM | 19 | -0,9 | 1 | 1 | 4,37E-01 | NA | 3 | 1 | 13 | 9 | 16 | 18 | 36 | 52 | -2,4 | -1,4 | -1,0 |
| chr3:136055498-136055998:downstream_2564 | chr3 | 136055998 | 136471245 | - | STAG1 | NM | 22 | -0,9 | 1 | 1 | 4,37E-01 | NA | 2 | 6 | 18 | 32 | 7 | 20 | 46 | 46 | -2,6 | -1,8 | -0,8 |
| chr1:9989275-9989775:downstream_427 | chr1 | 9989775 | 10002826 | - | LZIC | NM | 23 | -0,7 | 1 | 1 | 4,38E-01 | NA | 11 | 14 | 36 | 28 | 25 | 13 | 38 | 24 | -1,4 | -0,7 | -0,7 |
| chr14:50109769-50110269:downstream_19338 | chr14 | 50110269 | 50155098 | - | POLE2 | NM | 12 | 1,4 | 2 | -1 | 4,38E-01 | NA | 4 | 40 | 3 | 13 | 0 | 19 | 6 | 13 | 1,5 | 0,0 | 1,4 |
| chr2:38294245-38294745:downstream_1940 | chr2 | 38294745 | 38303323 | - | CYP1B1 | NM | 16 | -0,9 | 1 | 1 | 4,38E-01 | NA | 4 | 9 | 4 | 5 | 32 | 43 | 17 | 12 | 0,5 | 1,4 | -0,9 |
| chr19:17924385-17924885:downstream_14117 | chr19 | 17905918 | 17924385 | + | B3GNT3 | NM | 17 | -0,8 | 1 | 1 | 4,38E-01 | NA | 6 | 5 | 10 | 15 | 19 | 25 | 28 | 30 | -1,2 | -0,4 | -0,8 |
| chr16:3431580-3432080:downstream_20304 | chr16 | 3432080 | 3451065 | - | ZSCAN32 | NM | 17 | 0,9 | 1 | -1 | 4,40E-01 | NA | 7 | 3 | 6 | 2 | 24 | 24 | 38 | 29 | 0,3 | -0,5 | 0,8 |
| chr7:6727563-6728063:downstream_25821 | chr7 | 6728063 | 6746566 | - | ZNF12 | NM | 18 | 0,8 | 1 | -1 | 4,40E-01 | NA | 40 | 23 | 14 | 18 | 20 | 7 | 10 | 15 | 1,0 | 0,1 | 0,8 |
| chr15:34157927-34158427:downstream_11440 | chr15 | 34158427 | 34331303 | - | AVEN | NM | 9 | -1,4 | 2 | 1 | 4,40E-01 | NA | 1 | 1 | 14 | 10 | 2 | 6 | 29 | 9 | -3,7 | -2,4 | -1,4 |
| chr5:72210215-72210715:downstream_24422 | chr5 | 72112417 | 72210215 | + | TNPO1 | NM | 62 | -0,6 | 1 | 1 | 4,41E-01 | 7,82E-01 | 24 | 7 | 53 | 44 | 59 | 61 | 121 | 125 | -1,6 | -1,0 | -0,6 |
| chr12:69668138-69668638:downstream_18528 | chr12 | 69633316 | 69668138 | + | CPSF6 | NM | 28 | -0,8 | 1 | 1 | 4,42E-01 | NA | 41 | 6 | 47 | 42 | 24 | 18 | 15 | 30 | -0,9 | -0,1 | -0,8 |
| chr22:39918691-39919191:downstream_22749 | chr22 | 39916563 | 39918691 | + | ATF4 | NM | 172 | 0,5 | 1 | -1 | 4,43E-01 | 7,82E-01 | 47 | 32 | 24 | 15 | 429 | 336 | 281 | 215 | 1,0 | 0,6 | 0,4 |
| chr7:97922275-97922775:downstream_25410 | chr7 | 97910978 | 97922275 | + | BRI3 | NM | 9 | -1,1 | 1 | 1 | 4,43E-01 | NA | 13 | 9 | 19 | 16 | 6 | 4 | 4 | 3 | -0,7 | 0,4 | -1,1 |
| chr3:156424557-156425057:downstream_23470 | chr3 | 156392204 | 156424557 | + | TIPARP | NM | 18 | 0,8 | 1 | -1 | 4,43E-01 | NA | 25 | 25 | 13 | 13 | 21 | 15 | 15 | 16 | 1,0 | 0,2 | 0,8 |
| chr14:50042889-50043389:downstream_19368 | chr14 | 50043389 | 50053134 | - | RPS29 | NM | 50 | 0,6 | 1 | -1 | 4,43E-01 | 7,82E-01 | 41 | 56 | 7 | 9 | 120 | 109 | 32 | 26 | 2,6 | 2,0 | 0,6 |
| chr9:36212059-36212559:downstream_26264 | chr9 | 36190852 | 36212059 | + | CLTA | NM | 205 | 0,7 | 1 | -1 | 4,43E-01 | 7,82E-01 | 7 | 5 | 4 | 2 | 460 | 475 | 341 | 349 | 1,0 | 0,4 | 0,5 |
| chr17:6547861-6548361:downstream_12780 | chr17 | 6544221 | 6547861 | + | TXNDC17 | NM | 26 | 0,8 | 1 | -1 | 4,43E-01 | NA | 24 | 43 | 18 | 6 | 55 | 19 | 24 | 21 | 1,5 | 0,7 | 0,7 |
| chr15:75889923-75890423:downstream_19784 | chr15 | 75890423 | 75918719 | - | SNUPN | NM | 13 | 0,9 | 1 | -1 | 4,44E-01 | NA | 3 | 6 | 11 | 7 | 10 | 7 | 29 | 32 | -1,0 | -1,9 | 0,9 |
| chr5:159819654-159820154:downstream_24444 | chr5 | 159820154 | 159827104 | - | ZBED8 | NM | 14 | 0,9 | 1 | -1 | 4,44E-01 | NA | 15 | 15 | 7 | 7 | 19 | 15 | 21 | 11 | 1,0 | 0,1 | 0,9 |
| chr7:26226756-26227256:downstream_5473 | chr7 | 26191846 | 26226756 | + | NFE2L3 | NM | 9 | 1,2 | 2 | -1 | 4,44E-01 | NA | 7 | 21 | 4 | 8 | 2 | 13 | 6 | 9 | 1,2 | 0,0 | 1,2 |
| chr6:100955570-100956070:downstream_24516 | chr6 | 100956070 | 101329248 | - | ASCC3 | NM | 4 | -2,1 | 3 | 1 | 4,44E-01 | NA | 0 | 0 | 7 | 5 | 2 | 0 | 8 | 7 | -Inf | -3,2 | -Inf |
| chr16:25044098-25044598:downstream_12271 | chr16 | 25043061 | 25044098 | + | LOC554206 | NR | 5 | 1,4 | 2 | -1 | 4,45E-01 | NA | 7 | 4 | 1 | 4 | 2 | 8 | 7 | 7 | 1,0 | -0,4 | 1,4 |
| chr1:192980995-192981495:downstream_17381 | chr1 | 192981495 | 193028523 | - | UCHL5 | NM | 16 | -0,8 | 1 | 1 | 4,45E-01 | NA | 8 | 6 | 17 | 10 | 20 | 21 | 17 | 26 | -0,9 | 0,0 | -0,9 |
| chr4:42089551-42090051:downstream_3428 | chr4 | 41992522 | 42089551 | + | SLC30A9 | NM | 20 | -0,9 | 1 | 1 | 4,46E-01 | NA | 2 | 4 | 14 | 6 | 22 | 28 | 42 | 42 | -1,6 | -0,7 | -0,9 |
| chr15:67819641-67820141:downstream_11776 | chr15 | 67813521 | 67819641 | + | C15orf61 | NM | 12 | 0,9 | 1 | -1 | 4,46E-01 | NA | 18 | 19 | 11 | 17 | 7 | 7 | 11 | 8 | 0,4 | -0,5 | 0,9 |
| chr17:43582748-43583248:downstream_13178 | chr17 | 43583248 | 43597889 | - | LRRC37A4P | NR | 8 | -1,5 | 2 | 1 | 4,46E-01 | NA | 2 | 0 | 8 | 11 | 11 | 0 | 25 | 5 | -3,0 | -1,5 | -1,5 |
| chr17:12894428-12894928:downstream_20462 | chr17 | 12894928 | 12921381 | - | ELAC2 | NM | 12 | 1,0 | 1 | -1 | 4,47E-01 | NA | 5 | 15 | 13 | 7 | 11 | 10 | 26 | 12 | 0,0 | -0,9 | 1,0 |
| chr19:42234341-42234841:downstream_21061 | chr19 | 42212503 | 42234341 | + | CEACAM5 | NM | 7 | 1,3 | 2 | -1 | 4,47E-01 | NA | 1 | 10 | 8 | 3 | 5 | 4 | 11 | 12 | -0,1 | -1,4 | 1,3 |
| chr8:125500859-125501359:downstream_6705 | chr8 | 125487007 | 125500859 | + | RNF139 | NM | 33 | 0,6 | 1 | -1 | 4,47E-01 | 7,88E-01 | 35 | 38 | 18 | 21 | 41 | 45 | 40 | 30 | 0,9 | 0,3 | 0,6 |
| chr2:178094530-178095030:downstream_22113 | chr2 | 178095030 | 178129859 | - | NFE2L2 | NM | 20 | 0,8 | 1 | -1 | 4,47E-01 | NA | 6 | 6 | 21 | 15 | 8 | 10 | 53 | 41 | -1,6 | -2,4 | 0,8 |
| chr12:53873775-53874275:downstream_18670 | chr12 | 53874275 | 53893444 | - | MAP3K12 | NM | 14 | -1,0 | 1 | 1 | 4,48E-01 | NA | 2 | 2 | 6 | 7 | 20 | 18 | 26 | 29 | -1,5 | -0,5 | -1,0 |
| chr12:6547666-6548166:downstream_9811 | chr12 | 6548166 | 6560683 | - | CD27-AS1 | NR | 3 | 2,1 | 3 | -1 | 4,48E-01 | NA | 5 | 4 | 0 | 0 | 3 | 8 | 1 | 1 | Inf | 2,3 | Inf |
| chr3:50391679-50392179:downstream_2976 | chr3 | 50392179 | 50396939 | - | TMEM115 | NM | 17 | 0,8 | 1 | -1 | 4,48E-01 | NA | 28 | 16 | 7 | 7 | 25 | 22 | 11 | 16 | 1,6 | 0,8 | 0,8 |
| chr17:46942607-46943107:downstream_20373 | chr17 | 46908349 | 46942607 | + | CALCOCO2 | NM | 14 | 0,9 | 1 | -1 | 4,48E-01 | NA | 11 | 8 | 28 | 23 | 1 | 7 | 18 | 19 | -1,4 | -2,3 | 0,9 |
| chr14:39820397-39820897:downstream_19192 | chr14 | 39734475 | 39820397 | + | CTAGE5 | NM | 6 | 1,5 | 2 | -1 | 4,49E-01 | NA | 0 | 15 | 3 | 9 | 3 | 3 | 11 | 4 | 0,3 | -1,3 | 1,6 |
| chr18:47309373-47309873:downstream_13511 | chr18 | 47309873 | 47340251 | - | ACAA2 | NM | 28 | 0,7 | 1 | -1 | 4,49E-01 | 7,91E-01 | 12 | 13 | 13 | 5 | 50 | 39 | 46 | 49 | 0,5 | -0,1 | 0,6 |
| chr6:135281016-135281516:downstream_24793 | chr6 | 135281516 | 135376036 | - | HBS1L | NM | 9 | 1,2 | 2 | -1 | 4,50E-01 | NA | 15 | 5 | 8 | 7 | 3 | 9 | 17 | 4 | 0,4 | -0,8 | 1,2 |
| chr9:26902867-26903367:downstream_7020 | chr9 | 26903367 | 26947468 | - | PLAA | NM | 13 | -1,1 | 1 | 1 | 4,50E-01 | NA | 6 | 0 | 18 | 4 | 15 | 12 | 14 | 31 | -2,0 | -0,7 | -1,2 |
| chr2:48034092-48034592:downstream_22088 | chr2 | 48010220 | 48034092 | + | MSH6 | NM | 43 | 0,7 | 1 | -1 | 4,50E-01 | 7,91E-01 | 13 | 6 | 29 | 23 | 38 | 13 | 83 | 139 | -1,5 | -2,1 | 0,7 |
| chrX:16862274-16862774:downstream_27000 | chrX | 16862774 | 16888534 | - | RBBP7 | NM | 182 | -0,6 | 1 | 1 | 4,50E-01 | 7,91E-01 | 4 | 6 | 7 | 7 | 359 | 363 | 301 | 409 | -0,5 | 0,0 | -0,6 |
| chr1:212898994-212899494:downstream_17034 | chr1 | 212899494 | 212965139 | - | NSL1 | NM | 4 | -1,8 | 2 | 1 | 4,50E-01 | NA | 2 | 0 | 1 | 9 | 3 | 4 | 13 | 1 | -2,7 | -0,9 | -1,8 |
| chr6:43588260-43588760:downstream_25070 | chr6 | 43543877 | 43588260 | + | POLH | NM | 14 | -1,0 | 1 | 1 | 4,51E-01 | NA | 7 | 26 | 3 | 4 | 33 | 34 | 3 | 4 | 2,2 | 3,3 | -1,0 |
| chr5:14509458-14509958:downstream_3990 | chr5 | 14143828 | 14509458 | + | TRIO | NM | 15 | 1,1 | 1 | -1 | 4,51E-01 | NA | 3 | 7 | 32 | 39 | 2 | 0 | 17 | 22 | -2,8 | -4,0 | 1,2 |
| chr15:67487533-67488033:downstream_19727 | chr15 | 67358194 | 67487533 | + | SMAD3 | NM | 60 | 0,6 | 1 | -1 | 4,51E-01 | 7,91E-01 | 4 | 5 | 6 | 13 | 50 | 50 | 181 | 170 | -1,1 | -1,8 | 0,7 |
| chr1:109606497-109606997:downstream_868 | chr1 | 109606997 | 109618624 | - | TAF13 | NM | 56 | 0,6 | 1 | -1 | 4,51E-01 | 7,91E-01 | 13 | 38 | 10 | 6 | 134 | 127 | 75 | 47 | 1,7 | 1,1 | 0,6 |
| chr12:106903976-106904476:downstream_18750 | chr12 | 106751435 | 106903976 | + | POLR3B | NM | 4 | 2,1 | 3 | -1 | 4,52E-01 | NA | 1 | 0 | 4 | 9 | 0 | 0 | 8 | 12 | -4,0 | -Inf | Inf |
| chr2:46857315-46857815:downstream_1963 | chr2 | 46844310 | 46857315 | + | CRIPT | NM | 12 | 0,9 | 1 | -1 | 4,52E-01 | NA | 16 | 7 | 13 | 8 | 12 | 8 | 17 | 16 | 0,2 | -0,7 | 0,9 |
| chr1:40706593-40707093:downstream_118 | chr1 | 40627040 | 40706593 | + | RLF | NM | 6 | -1,5 | 2 | 1 | 4,52E-01 | NA | 2 | 0 | 10 | 3 | 7 | 2 | 10 | 15 | -3,0 | -1,4 | -1,6 |
| chr15:22873891-22874391:downstream_19823 | chr15 | 22833394 | 22873891 | + | TUBGCP5 | NM | 10 | -1,1 | 1 | 1 | 4,52E-01 | NA | 2 | 2 | 6 | 7 | 8 | 19 | 18 | 18 | -1,5 | -0,4 | -1,1 |
| chr4:48039080-48039580:downstream_3669 | chr4 | 48018790 | 48039080 | + | NIPAL1 | NM | 5 | -1,8 | 2 | 1 | 4,52E-01 | NA | 13 | 14 | 0 | 3 | 2 | 8 | 0 | 0 | 3,0 | Inf | -Inf |
| chr8:23430063-23430563:downstream_6495 | chr8 | 23386362 | 23430063 | + | SLC25A37 | NM | 24 | -0,8 | 1 | 1 | 4,52E-01 | NA | 14 | 4 | 60 | 35 | 15 | 5 | 36 | 24 | -2,4 | -1,6 | -0,8 |
| chr2:39109850-39110350:downstream_1944 | chr2 | 39103102 | 39109850 | + | MORN2 | NM | 7 | 1,2 | 2 | -1 | 4,52E-01 | NA | 14 | 10 | 1 | 2 | 14 | 10 | 4 | 5 | 2,6 | 1,4 | 1,2 |
| chr16:57495187-57495687:downstream_12385 | chr16 | 57481336 | 57495187 | + | COQ9 | NM | 18 | 0,8 | 1 | -1 | 4,53E-01 | NA | 7 | 15 | 14 | 7 | 29 | 11 | 29 | 36 | 0,1 | -0,7 | 0,8 |
| chr9:96872138-96872638:downstream_26464 | chr9 | 96793075 | 96872138 | + | PTPDC1 | NM | 5 | -1,5 | 2 | 1 | 4,53E-01 | NA | 2 | 2 | 6 | 6 | 2 | 10 | 1 | 11 | -1,7 | -0,1 | -1,6 |
| chr17:8091150-8091650:downstream_12821 | chr17 | 8091650 | 8093564 | - | C17orf59 | NM | 8 | 1,1 | 1 | -1 | 4,53E-01 | NA | 16 | 5 | 7 | 5 | 11 | 5 | 8 | 10 | 0,8 | -0,2 | 1,1 |
| chr14:53241410-53241910:downstream_11144 | chr14 | 53241910 | 53258386 | - | GNPNAT1 | NM | 41 | 0,7 | 1 | -1 | 4,53E-01 | 7,93E-01 | 18 | 4 | 8 | 11 | 55 | 65 | 82 | 84 | 0,2 | -0,5 | 0,7 |
| chr1:205270690-205271190:downstream_1325 | chr1 | 205271190 | 205290919 | - | NUAK2 | NM | 23 | 0,8 | 1 | -1 | 4,53E-01 | NA | 17 | 25 | 13 | 7 | 46 | 27 | 15 | 38 | 1,1 | 0,5 | 0,7 |
| chr3:125247201-125247701:downstream_2693 | chr3 | 125247701 | 125314381 | - | OSBPL11 | NM | 9 | 1,3 | 2 | -1 | 4,54E-01 | NA | 3 | 0 | 14 | 7 | 1 | 2 | 25 | 16 | -2,7 | -3,9 | 1,2 |
| chr2:61414089-61414589:downstream_1627 | chr2 | 61414589 | 61697849 | - | USP34 | NM | 22 | 0,7 | 1 | -1 | 4,54E-01 | NA | 6 | 10 | 22 | 17 | 14 | 10 | 36 | 65 | -1,4 | -2,1 | 0,7 |
| chr12:7085165-7085665:downstream_10006 | chr12 | 7079943 | 7085165 | + | EMG1 | NM | 75 | 0,5 | 1 | -1 | 4,55E-01 | 7,94E-01 | 84 | 113 | 74 | 56 | 70 | 73 | 63 | 65 | 0,6 | 0,2 | 0,4 |
| chr7:135046046-135046546:downstream_25455 | chr7 | 135046546 | 135194875 | - | CNOT4 | NM | 8 | -1,5 | 2 | 1 | 4,55E-01 | NA | 1 | 0 | 8 | 7 | 4 | 3 | 19 | 22 | -4,2 | -2,6 | -1,6 |
| chr1:65298914-65299414:downstream_149 | chr1 | 65210777 | 65298914 | + | RAVER2 | NM | 11 | -1,0 | 1 | 1 | 4,55E-01 | NA | 5 | 8 | 22 | 11 | 4 | 15 | 17 | 8 | -1,4 | -0,4 | -1,0 |
| chr2:61279125-61279625:downstream_1983 | chr2 | 61244811 | 61279125 | + | PEX13 | NM | 13 | -1,4 | 2 | 1 | 4,55E-01 | NA | 1 | 0 | 3 | 7 | 2 | 15 | 45 | 34 | -3,6 | -2,1 | -1,5 |
| chr18:9960018-9960518:downstream_20970 | chr18 | 9913954 | 9960018 | + | VAPA | NM | 13 | -1,1 | 1 | 1 | 4,55E-01 | NA | 16 | 0 | 11 | 11 | 20 | 19 | 15 | 10 | -0,4 | 0,7 | -1,1 |

| | | | | | | | | | | | | | | | | | | | | | | | |
|---|---|---|---|---|---|---|---|---|---|---|---|---|---|---|---|---|---|---|---|---|---|---|---|
| chr2:36778903-36779403:downstream_21921 | chr2 | 36779403 | 36825332 | - | FEZ2 | NM | 22 | -0,7 | 1 | 1 | 4,56E-01 | NA | 4 | 8 | 14 | 9 | 39 | 28 | 36 | 41 | -1,0 | -0,2 | -0,8 |
| chr4:155503779-155504279:downstream_23650 | chr4 | 155504279 | 155511897 | - | FGA | NM | 101 | -0,5 | 1 | 1 | 4,56E-01 | 7,94E-01 | 14 | 7 | 19 | 18 | 118 | 219 | 184 | 229 | -0,9 | -0,3 | -0,6 |
| chr1:1337426-1337926:downstream_374 | chr1 | 1334909 | 1337426 | + | LOC148413 | NR | 55 | 0,6 | 1 | -1 | 4,56E-01 | 7,94E-01 | 22 | 20 | 14 | 13 | 60 | 132 | 90 | 91 | 0,6 | 0,1 | 0,6 |
| chr9:15461627-15462127:downstream_6989 | chr9 | 15422781 | 15461627 | + | SNAPC3 | NM | 29 | -0,6 | 1 | 1 | 4,56E-01 | 7,94E-01 | 26 | 22 | 46 | 42 | 21 | 24 | 15 | 36 | -0,9 | -0,2 | -0,7 |
| chr19:16302857-16303357:downstream_14096 | chr19 | 16296211 | 16302857 | + | FAM32A | NM | 150 | 1,4 | 2 | -1 | 4,57E-01 | 7,95E-01 | 8 | 0 | 0 | 2 | 340 | 395 | 217 | 235 | 2,3 | 0,7 | 1,6 |
| chr22:42222303-42222803:downstream_22767 | chr22 | 42196625 | 42222303 | + | CCDC134 | NM | 5 | 1,5 | 2 | -1 | 4,57E-01 | NA | 2 | 4 | 6 | 7 | 2 | 0 | 8 | 7 | -1,0 | -2,6 | 1,6 |
| chr2:242293441-242293941:downstream_22213 | chr2 | 242254601 | 242293441 | + | sept-02 | NM | 113 | 0,6 | 1 | -1 | 4,57E-01 | 7,95E-01 | 7 | 3 | 3 | 5 | 205 | 195 | 239 | 244 | 0,4 | -0,3 | 0,7 |
| chr6:160099648-160100148:downstream_25219 | chr6 | 160100148 | 160114353 | - | SOD2 | NM | 13 | 1,0 | 1 | -1 | 4,57E-01 | NA | 7 | 3 | 3 | 2 | 12 | 28 | 31 | 15 | 0,8 | -0,2 | 1,0 |
| chr5:156564950-156565450:downstream_24223 | chr5 | 156565450 | 156569921 | - | MED7 | NM | 44 | 0,6 | 1 | -1 | 4,58E-01 | 7,95E-01 | 24 | 14 | 13 | 11 | 79 | 72 | 82 | 55 | 0,7 | 0,1 | 0,6 |
| chr12:12982915-12983415:downstream_18538 | chr12 | 12966279 | 12982915 | + | DDX47 | NM | 24 | 0,8 | 1 | -1 | 4,58E-01 | NA | 6 | 9 | 4 | 3 | 59 | 29 | 33 | 46 | 0,9 | 0,2 | 0,8 |
| chr9:110246632-110247132:downstream_7251 | chr9 | 110247132 | 110252047 | - | KLF4 | NM | 5 | 1,8 | 2 | -1 | 4,59E-01 | NA | 6 | 3 | 0 | 0 | 22 | 6 | 3 | 2 | Inf | 2,6 | Inf |
| chr9:5784071-5784571:downstream_6979 | chr9 | 5784571 | 5833081 | - | ERMP1 | NM | 58 | -0,6 | 1 | 1 | 4,59E-01 | 7,97E-01 | 15 | 4 | 33 | 31 | 69 | 47 | 118 | 148 | -1,8 | -1,2 | -0,6 |
| chr6:138734582-138735082:downstream_5372 | chr6 | 138725335 | 138734582 | + | HEBP2 | NM | 22 | 0,8 | 1 | -1 | 4,60E-01 | NA | 3 | 15 | 10 | 2 | 31 | 39 | 42 | 39 | 0,5 | -0,2 | 0,7 |
| chr1:153964631-153965131:downstream_1075 | chr1 | 153963238 | 153964631 | + | RPS27 | NM | 104 | 0,4 | 1 | -1 | 4,61E-01 | 7,98E-01 | 64 | 75 | 33 | 21 | 241 | 183 | 97 | 120 | 1,4 | 1,0 | 0,4 |
| chr2:32248471-32248971:downstream_1930 | chr2 | 32248971 | 32264844 | - | DPY30 | NM | 24 | -0,9 | 1 | 1 | 4,61E-01 | NA | 5 | 0 | 4 | 8 | 41 | 31 | 56 | 43 | -1,3 | -0,5 | -0,9 |
| chr16:81418698-81419198:downstream_12493 | chr16 | 81418622 | 81418698 | + | MIR4720 | NR | 15 | 0,9 | 1 | -1 | 4,62E-01 | NA | 15 | 22 | 18 | 26 | 1 | 12 | 17 | 11 | -0,3 | -1,2 | 0,9 |
| chr7:100882392-100882892:downstream_6014 | chr7 | 100882892 | 100888371 | - | FIS1 | NM | 16 | -0,9 | 1 | 1 | 4,62E-01 | NA | 8 | 1 | 10 | 6 | 41 | 15 | 29 | 19 | -0,8 | 0,2 | -1,0 |
| chr17:42638630-42639130:downstream_13167 | chr17 | 42634811 | 42638630 | + | FZD2 | NM | 6 | -1,7 | 2 | 1 | 4,62E-01 | NA | 0 | 0 | 4 | 8 | 1 | 2 | 15 | 17 | -Inf | -3,6 | -Inf |
| chr11:65120451-65120951:downstream_9439 | chr11 | 65101224 | 65120451 | + | DPF2 | NM | 11 | -1,0 | 1 | 1 | 4,62E-01 | NA | 6 | 10 | 7 | 2 | 22 | 30 | 7 | 7 | 0,8 | 1,9 | -1,1 |
| chr8:68109383-68109883:downstream_6240 | chr8 | 68109883 | 68255912 | - | ARFGEF1 | NM | 17 | -1,4 | 2 | 1 | 4,63E-01 | NA | 0 | 0 | 7 | 6 | 7 | 1 | 45 | 68 | -Inf | -3,9 | -Inf |
| chr6:41900784-41901284:downstream_5133 | chr6 | 41888964 | 41900784 | + | BYSL | NM | 29 | 0,7 | 1 | -1 | 4,63E-01 | 8,02E-01 | 17 | 39 | 6 | 10 | 59 | 52 | 25 | 26 | 1,9 | 1,1 | 0,7 |
| chr5:138667366-138667866:downstream_24220 | chr5 | 138609440 | 138667366 | + | MATR3 | NM | 15 | 0,8 | 1 | -1 | 4,64E-01 | NA | 18 | 18 | 15 | 11 | 15 | 11 | 13 | 18 | 0,5 | -0,3 | 0,8 |
| chr17:73313656-73314156:downstream_20504 | chr17 | 73314156 | 73401790 | - | GRB2 | NM | 15 | 1,0 | 1 | -1 | 4,64E-01 | NA | 0 | 7 | 3 | 9 | 10 | 11 | 31 | 48 | -0,8 | -1,9 | 1,1 |
| chr3:120043075-120043575:downstream_2677 | chr3 | 120043575 | 120068186 | - | LRRC58 | NM | 24 | 0,8 | 1 | -1 | 4,64E-01 | NA | 15 | 12 | 8 | 17 | 43 | 11 | 45 | 44 | 0,1 | -0,7 | 0,8 |
| chr3:152888413-152888913:downstream_3243 | chr3 | 152880000 | 152888413 | + | RAP2B | NM | 60 | 0,8 | 1 | -1 | 4,64E-01 | 8,02E-01 | 54 | 63 | 4 | 1 | 177 | 160 | 6 | 14 | 4,6 | 4,1 | 0,4 |
| chr11:44105569-44106069:downstream_17844 | chr11 | 44087728 | 44105569 | + | ACCS | NM | 6 | 1,2 | 2 | -1 | 4,64E-01 | NA | 7 | 7 | 4 | 4 | 5 | 5 | 8 | 5 | 0,8 | -0,4 | 1,2 |
| chr5:139008018-139008518:downstream_24434 | chr5 | 138940750 | 139008018 | + | UBE2D2 | NM | 7 | 1,3 | 2 | -1 | 4,65E-01 | NA | 2 | 8 | 1 | 4 | 9 | 6 | 18 | 5 | 0,8 | -0,6 | 1,4 |
| chr7:97920461-97920961:downstream_5962 | chr7 | 97920961 | 98030427 | - | BAIAP2L1 | NM | 9 | 1,0 | 1 | -1 | 4,65E-01 | NA | 6 | 5 | 7 | 5 | 7 | 9 | 15 | 18 | -0,2 | -1,1 | 1,0 |
| chr10:127542264-127542764:downstream_17488 | chr10 | 127512103 | 127542264 | + | BCCIP | NM | 18 | 0,8 | 1 | -1 | 4,65E-01 | NA | 24 | 22 | 6 | 2 | 36 | 32 | 7 | 12 | 2,5 | 1,8 | 0,7 |
| chr10:7849762-7850262:downstream_17485 | chr10 | 7830092 | 7849762 | + | ATP5C1 | NM | 23 | 0,7 | 1 | -1 | 4,65E-01 | NA | 6 | 9 | 8 | 8 | 33 | 20 | 53 | 48 | -0,2 | -0,9 | 0,7 |
| chr4:57897328-57897828:downstream_23801 | chr4 | 57844805 | 57897328 | + | POLR2B | NM | 32 | 0,7 | 1 | -1 | 4,66E-01 | 8,04E-01 | 33 | 70 | 26 | 21 | 24 | 39 | 13 | 33 | 1,1 | 0,5 | 0,6 |
| chr19:58416641-58417141:downstream_21640 | chr19 | 58417141 | 58427984 | - | ZNF417 | NM | 13 | 0,9 | 1 | -1 | 4,67E-01 | NA | 9 | 18 | 4 | 15 | 9 | 16 | 17 | 17 | 0,5 | -0,4 | 0,9 |
| chr2:204082717-204083217:downstream_1779 | chr2 | 203879601 | 204082717 | + | NBEAL1 | NM | 27 | -0,7 | 1 | 1 | 4,67E-01 | NA | 10 | 11 | 54 | 32 | 5 | 26 | 43 | 34 | -2,1 | -1,3 | -0,8 |
| chr4:83814104-83814604:downstream_23885 | chr4 | 83814604 | 83822069 | - | THAP9-AS1 | NR | 17 | -0,9 | 1 | 1 | 4,67E-01 | NA | 14 | 30 | 13 | 7 | 15 | 41 | 4 | 9 | 1,1 | 2,1 | -0,9 |
| chr8:91064227-91064727:downstream_6632 | chr8 | 91013579 | 91064227 | + | DECR1 | NM | 38 | 0,8 | 1 | -1 | 4,67E-01 | 8,05E-01 | 7 | 9 | 10 | 0 | 62 | 85 | 71 | 63 | 0,6 | 0,1 | 0,5 |
| chr12:121015397-121015897:downstream_9943 | chr12 | 120972131 | 121015397 | + | RNF10 | NM | 9 | 1,0 | 1 | -1 | 4,67E-01 | NA | 3 | 10 | 6 | 2 | 11 | 12 | 15 | 12 | 0,7 | -0,3 | 1,0 |
| chr11:44641315-44641815:downstream_17942 | chr11 | 44587140 | 44641315 | + | CD82 | NM | 6 | 1,3 | 2 | -1 | 4,67E-01 | NA | 6 | 8 | 6 | 1 | 11 | 3 | 7 | 8 | 1,1 | -0,1 | 1,2 |
| chr10:71871522-71872022:downstream_17459 | chr10 | 71872022 | 71892690 | - | AIFM2 | NM | 8 | 1,1 | 1 | -1 | 4,67E-01 | NA | 11 | 14 | 7 | 4 | 11 | 4 | 8 | 6 | 1,2 | 0,1 | 1,1 |
| chr1:227918389-227918889:downstream_16833 | chr1 | 227918889 | 227923112 | - | JMJD4 | NM | 4 | 1,6 | 2 | -1 | 4,68E-01 | NA | 8 | 4 | 0 | 3 | 8 | 1 | 4 | 4 | 1,9 | 0,1 | 1,7 |
| chr3:100081802-100082302:downstream_3075 | chr3 | 100082302 | 100120242 | - | TOMM70A | NM | 18 | -0,9 | 1 | 1 | 4,68E-01 | NA | 3 | 1 | 8 | 7 | 20 | 23 | 38 | 43 | -1,8 | -0,9 | -0,9 |
| chr5:79775688-79776188:downstream_24448 | chr5 | 79703831 | 79775688 | + | ZFYVE16 | NM | 12 | 0,9 | 1 | -1 | 4,68E-01 | NA | 18 | 12 | 15 | 17 | 8 | 3 | 15 | 8 | -0,1 | -1,1 | 1,0 |
| chr1:20825440-20825940:downstream_524 | chr1 | 20825940 | 20834674 | - | MUL1 | NM | 15 | 0,9 | 1 | -1 | 4,69E-01 | NA | 14 | 8 | 3 | 2 | 31 | 34 | 17 | 12 | 2,0 | 1,2 | 0,9 |
| chrX:119759028-119759528:downstream_26651 | chrX | 119759528 | 119764005 | - | C1GALT1C1 | NM | 54 | -0,6 | 1 | 1 | 4,69E-01 | 8,06E-01 | 5 | 7 | 8 | 6 | 107 | 121 | 83 | 92 | -0,3 | 0,4 | -0,6 |
| chr17:27077976-27078476:downstream_12938 | chr17 | 27071022 | 27077976 | + | TRAF4 | NM | 18 | 1,3 | 2 | -1 | 4,69E-01 | NA | 18 | 2 | 0 | 1 | 52 | 57 | 6 | 8 | 4,6 | 3,0 | 1,6 |
| chr2:175195370-175195870:downstream_2316 | chr2 | 175190754 | 175195370 | + | LINC01305 | NR | 13 | -0,9 | 1 | 1 | 4,69E-01 | NA | 8 | 3 | 22 | 7 | 13 | 13 | 22 | 13 | -1,4 | -0,5 | -1,0 |
| chr9:33573001-33573501:downstream_6804 | chr9 | 33524410 | 33573001 | + | ANKRD18B | NM | 4 | -1,9 | 3 | 1 | 4,69E-01 | NA | 0 | 0 | 6 | 6 | 2 | 0 | 10 | 6 | -Inf | -3,3 | -Inf |
| chr8:144103827-144104327:downstream_25999 | chr8 | 144099901 | 144103827 | + | LY6E | NM | 91 | -0,6 | 1 | 1 | 4,69E-01 | 8,06E-01 | 7 | 7 | 1 | 7 | 275 | 219 | 97 | 120 | 0,7 | 1,2 | -0,4 |
| chr19:42733837-42734337:downstream_14331 | chr19 | 42734337 | 42746736 | - | GSK3A | NM | 30 | 0,7 | 1 | -1 | 4,70E-01 | 8,06E-01 | 7 | 15 | 7 | 8 | 50 | 44 | 58 | 52 | 0,5 | -0,2 | 0,7 |
| chr12:79985244-79985744:downstream_10330 | chr12 | 79985744 | 80084790 | - | PAWR | NM | 35 | 0,6 | 1 | -1 | 4,70E-01 | 8,06E-01 | 34 | 46 | 74 | 59 | 13 | 6 | 19 | 28 | -0,7 | -1,3 | 0,6 |
| chr1:28526289-28526789:downstream_588 | chr1 | 28526789 | 28559542 | - | DNAJC8 | NM | 19 | 0,9 | 1 | -1 | 4,70E-01 | NA | 7 | 2 | 6 | 2 | 30 | 28 | 29 | 45 | 0,4 | -0,4 | 0,7 |
| chr17:33900772-33901272:downstream_12989 | chr17 | 33900675 | 33900772 | + | SNORD7 | NR | 15 | -0,9 | 1 | 1 | 4,70E-01 | NA | 2 | 7 | 6 | 7 | 40 | 15 | 29 | 13 | -0,5 | 0,4 | -0,9 |
| chr9:127114719-127115219:downstream_26407 | chr9 | 127019884 | 127114719 | + | NEK6 | NM | 9 | -1,1 | 1 | 1 | 4,70E-01 | NA | 9 | 5 | 19 | 17 | 2 | 8 | 7 | 5 | -1,4 | -0,3 | -1,1 |
| chr7:102096166-102096666:downstream_5598 | chr7 | 102096666 | 102105321 | - | ALKBH4 | NM | 4 | 1,5 | 2 | -1 | 4,70E-01 | NA | 2 | 9 | 1 | 5 | 4 | 1 | 3 | 7 | 0,7 | -0,9 | 1,6 |
| chr7:56078243-56078743:downstream_5560 | chr7 | 56078743 | 56119268 | - | PSPH | NM | 20 | 0,8 | 1 | -1 | 4,70E-01 | NA | 17 | 16 | 14 | 11 | 14 | 32 | 40 | 16 | 0,4 | -0,3 | 0,8 |
| chr7:99689850-99690350:downstream_25626 | chr7 | 99690350 | 99699563 | - | MCM7 | NM | 18 | -0,9 | 1 | 1 | 4,71E-01 | NA | 6 | 1 | 3 | 15 | 19 | 30 | 28 | 42 | -1,4 | -0,5 | -0,9 |
| chr3:197340397-197340897:downstream_3358 | chr3 | 197340897 | 197354752 | - | LOC220729 | NR | 39 | -0,6 | 1 | 1 | 4,71E-01 | 8,07E-01 | 47 | 37 | 53 | 66 | 20 | 37 | 22 | 31 | -0,5 | 0,1 | -0,6 |
| chr11:65540868-65541368:downstream_9456 | chr11 | 65541368 | 65548062 | - | AP5B1 | NM | 15 | 1,0 | 1 | -1 | 4,71E-01 | NA | 2 | 8 | 3 | 2 | 31 | 25 | 32 | 20 | 1,1 | 0,1 | 1,0 |
| chr12:51640501-51641001:downstream_18532 | chr12 | 51632507 | 51640501 | + | DAZAP2 | NM | 9 | 1,1 | 1 | -1 | 4,71E-01 | NA | 25 | 19 | 4 | 7 | 6 | 5 | 1 | 4 | 1,9 | 0,9 | 1,0 |
| chr4:164441691-164442191:downstream_3894 | chr4 | 164415672 | 164441691 | + | TMA16 | NM | 32 | 0,6 | 1 | -1 | 4,71E-01 | 8,07E-01 | 39 | 31 | 24 | 18 | 34 | 41 | 40 | 26 | 0,7 | 0,2 | 0,6 |
| chr12:50514240-50514740:downstream_18526 | chr12 | 50505763 | 50514240 | + | COX14 | NM | 62 | 0,5 | 1 | -1 | 4,71E-01 | 8,07E-01 | 17 | 15 | 13 | 12 | 115 | 96 | 132 | 98 | 0,4 | -0,1 | 0,5 |
| chr4:68588222-68588722:downstream_3699 | chr4 | 68566995 | 68588222 | + | UBA6-AS1 | NR | 8 | 1,1 | 2 | -1 | 4,72E-01 | NA | 11 | 13 | 15 | 7 | 6 | 0 | 7 | 6 | 0,0 | -1,2 | 1,2 |
| chr4:17801777-17802277:downstream_3638 | chr4 | 17802277 | 17812381 | - | DCAF16 | NM | 11 | 1,0 | 1 | -1 | 4,72E-01 | NA | 5 | 19 | 4 | 7 | 8 | 21 | 10 | 16 | 1,2 | 0,2 | 1,0 |
| chr1:65523616-65524116:downstream_778 | chr1 | 65524116 | 65524191 | - | MIR101-1 | NR | 8 | -1,0 | 1 | 1 | 4,72E-01 | NA | 5 | 7 | 10 | 16 | 4 | 10 | 8 | 7 | -1,1 | -0,1 | -1,0 |
| chr8:77878503-77879003:downstream_6610 | chr8 | 77879003 | 77879086 | - | MIR3149 | NR | 10 | -1,1 | 2 | 1 | 4,72E-01 | NA | 11 | 11 | 14 | 8 | 7 | 16 | 0 | 10 | 0,0 | 1,2 | -1,2 |
| chr12:22843608-22844108:downstream_18559 | chr12 | 22778075 | 22843608 | + | ETNK1 | NM | 19 | 0,8 | 1 | -1 | 4,72E-01 | NA | 7 | 6 | 7 | 15 | 16 | 15 | 56 | 34 | -0,7 | -1,5 | 0,8 |
| chr19:36378642-36379142:downstream_14236 | chr19 | 36379142 | 36391552 | - | NFKBID | NM | 4 | 2,0 | 3 | -1 | 4,72E-01 | NA | 1 | 0 | 4 | 9 | 0 | 0 | 10 | 10 | -4,0 | -Inf | Inf |
| chr14:60765805-60766305:downstream_19339 | chr14 | 60712469 | 60765805 | + | PPM1A | NM | 10 | 1,1 | 1 | -1 | 4,74E-01 | NA | 3 | 7 | 1 | 2 | 13 | 23 | 19 | 11 | 1,4 | 0,2 | 1,2 |
| chr15:42062141-42062641:downstream_19619 | chr15 | 41952609 | 42062141 | + | MGA | NM | 30 | -0,6 | 1 | 1 | 4,74E-01 | 8,09E-01 | 7 | 6 | 40 | 30 | 16 | 20 | 56 | 68 | -2,4 | -1,8 | -0,7 |
| chr14:75203390-75203890:downstream_11229 | chr14 | 75179849 | 75203390 | + | FCF1 | NM | 14 | 0,9 | 1 | -1 | 4,75E-01 | NA | 21 | 5 | 11 | 7 | 17 | 13 | 15 | 21 | 0,6 | -0,3 | 0,8 |
| chr17:62540234-62540734:downstream_12673 | chr17 | 62540734 | 62658386 | - | SMURF2 | NM | 14 | -0,9 | 1 | 1 | 4,75E-01 | NA | 6 | 2 | 17 | 26 | 7 | 10 | 19 | 30 | -2,5 | -1,6 | -0,9 |
| chr13:95671582-95672082:downstream_18931 | chr13 | 95672082 | 95953700 | - | ABCC4 | NM | 47 | 0,6 | 1 | -1 | 4,75E-01 | 8,10E-01 | 5 | 8 | 6 | 25 | 37 | 31 | 110 | 159 | -1,3 | -2,0 | 0,7 |
| chr11:118885921-118886421:downstream_9685 | chr11 | 118886421 | 118889057 | - | RPS25 | NM | 21 | -0,9 | 1 | 1 | 4,75E-01 | NA | 31 | 9 | 8 | 14 | 20 | 63 | 6 | 20 | 0,8 | 1,7 | -0,9 |
| chr12:68056444-68056944:downstream_18548 | chr12 | 68042511 | 68056444 | + | DYRK2 | NM | 42 | 0,6 | 1 | -1 | 4,76E-01 | 8,10E-01 | 28 | 23 | 50 | 23 | 50 | 15 | 67 | 77 | -0,5 | -1,1 | 0,6 |
| chr7:93555826-93556326:downstream_5946 | chr7 | 93551015 | 93555826 | + | GNG11 | NM | 61 | -0,6 | 1 | 1 | 4,76E-01 | 8,10E-01 | 12 | 3 | 14 | 12 | 119 | 93 | 109 | 130 | -0,8 | -0,2 | -0,6 |
| chr1:23630680-23631180:downstream_16803 | chr1 | 23631180 | 23670857 | - | HNRNPR | NM | 32 | 0,7 | 1 | -1 | 4,76E-01 | 8,10E-01 | 59 | 34 | 63 | 63 | 11 | 1 | 15 | 13 | -0,4 | -1,2 | 0,8 |
| chr11:32623125-32623625:downstream_8867 | chr11 | 32623625 | 32816204 | - | CCDC73 | NM | 14 | 1,0 | 1 | -1 | 4,76E-01 | NA | 9 | 15 | 0 | 6 | 11 | 41 | 8 | 20 | 2,0 | 0,9 | 1,2 |
| chr5:179022404-179022904:downstream_4624 | chr5 | 179022904 | 179030742 | - | LOC101928445 | NR | 6 | 1,7 | 2 | -1 | 4,77E-01 | NA | 12 | 4 | 0 | 0 | 20 | 10 | 3 | 0 | Inf | 3,4 | Inf |
| chr8:29919757-29920257:downstream_6282 | chr8 | 29920257 | 30108213 | - | MIR548O2 | NR | 6 | 1,5 | 2 | -1 | 4,77E-01 | NA | 2 | 21 | 1 | 1 | 2 | 18 | 4 | 2 | 3,4 | 1,8 | 1,5 |
| chr15:55830584-55831084:downstream_11476 | chr15 | 55831084 | 55881050 | - | PYGO1 | NM | 7 | 1,1 | 2 | -1 | 4,78E-01 | NA | 4 | 12 | 3 | 2 | 13 | 8 | 7 | 8 | 1,6 | 0,5 | 1,1 |
| chr12:7246645-7247145:downstream_18469 | chr12 | 7247145 | 7261874 | - | C1RL | NM | 38 | -0,7 | 1 | 1 | 4,78E-01 | 8,12E-01 | 7 | 6 | 11 | 18 | 32 | 76 | 64 | 90 | -1,2 | -0,5 | -0,6 |
| chr2:33808227-33808727:downstream_21912 | chr2 | 33808727 | 33824429 | - | FAM98A | NM | 19 | 0,8 | 1 | -1 | 4,78E-01 | NA | 12 | 18 | 15 | 4 | 33 | 16 | 32 | 20 | 0,7 | -0,1 | 0,7 |
| chr2:187628512-187629012:downstream_1756 | chr2 | 187558788 | 187628512 | + | FAM171B | NM | 16 | -0,9 | 1 | 1 | 4,78E-01 | NA | 5 | 0 | 18 | 24 | 6 | 9 | 32 | 31 | -3,1 | -2,1 | -1,0 |
| chr17:38631579-38632079:downstream_13049 | chr17 | 38632079 | 38657854 | - | TNS4 | NM | 9 | -1,0 | 1 | 1 | 4,79E-01 | NA | 4 | 4 | 6 | 7 | 10 | 17 | 11 | 10 | -0,6 | 0,4 | -1,0 |
| chr3:48333306-48333806:downstream_23310 | chr3 | 48333806 | 48343175 | - | NME6 | NM | 5 | -1,3 | 2 | 1 | 4,79E-01 | NA | 5 | 1 | 7 | 6 | 8 | 4 | 3 | 7 | -1,1 | 0,2 | -1,4 |
| chr1:202696031-202696531:downstream_277 | chr1 | 202696531 | 202777549 | - | KDM5B | NM | 14 | 0,9 | 1 | -1 | 4,80E-01 | NA | 2 | 5 | 6 | 6 | 10 | 12 | 31 | 40 | -0,8 | -1,7 | 0,9 |
| chr16:3072120-3072620:downstream_20014 | chr16 | 3072620 | 3074287 | - | HCFC1R1 | NM | 38 | 0,8 | 1 | -1 | 4,80E-01 | 8,15E-01 | 4 | 14 | 3 | 2 | 112 | 74 | 49 | 46 | 1,7 | 1,0 | 0,8 |
| chr7:15239442-15239942:downstream_5513 | chr7 | 15239942 | 15601640 | - | AGMO | NM | 7 | 1,1 | 2 | -1 | 4,80E-01 | NA | 9 | 4 | 11 | 6 | 2 | 5 | 10 | 10 | -0,4 | -1,4 | 1,0 |
| chr2:24289953-24290453:downstream_1887 | chr2 | 24290453 | 24299314 | - | SF3B6 | NM | 18 | 0,9 | 1 | -1 | 4,80E-01 | NA | 6 | 4 | 1 | 2 | 45 | 29 | 22 | 35 | 1,3 | 0,4 | 1,0 |
| chr8:56650802-56651302:downstream_26154 | chr8 | 56651302 | 56685966 | - | TMEM68 | NM | 19 | 0,9 | 1 | -1 | 4,81E-01 | NA | 10 | 5 | 4 | 0 | 45 | 45 | 19 | 20 | 1,8 | 1,2 | 0,6 |
| chr11:57429337-57429837:downstream_17958 | chr11 | 57425215 | 57429337 | + | CLP1 | NM | 45 | -0,9 | 1 | 1 | 4,81E-01 | 8,15E-01 | 14 | 4 | 1 | 2 | 154 | 154 | 8 | 20 | 2,5 | 3,5 | -0,9 |
| chr16:70283633-70284133:downstream_12450 | chr16 | 70284133 | 70285833 | - | EXOSC6 | NM | 89 | 0,4 | 1 | -1 | 4,81E-01 | 8,15E-01 | 101 | 133 | 83 | 68 | 80 | 94 | 85 | 65 | 0,6 | 0,2 | 0,4 |
| chr1:202993197-202993697:downstream_1305 | chr1 | 202976533 | 202993197 | + | TMEM183A | NM | 26 | -0,6 | 1 | 1 | 4,82E-01 | NA | 16 | 9 | 15 | 13 | 45 | 46 | 24 | 41 | -0,2 | 0,5 | -0,7 |
| chr5:441450-441950:downstream_4147 | chr5 | 441950 | 443275 | - | EXOC3-AS1 | NR | 14 | -0,8 | 1 | 1 | 4,82E-01 | NA | 21 | 15 | 14 | 7 | 23 | 16 | 8 | 5 | 0,7 | 1,6 | -0,8 |
| chr3:3190816-3191316:downstream_23099 | chr3 | 3191316 | 3221401 | - | CRBN | NM | 5 | 1,4 | 2 | -1 | 4,83E-01 | NA | 6 | 5 | 13 | 3 | 3 | 0 | 4 | 8 | -0,6 | -1,9 | 1,3 |
| chr5:148874956-148875456:downstream_24075 | chr5 | 148875456 | 148931115 | - | CSNK1A1 | NM | 193 | 0,3 | 0 | -1 | 4,83E-01 | 8,17E-01 | 181 | 190 | 326 | 254 | 102 | 97 | 209 | 186 | -0,6 | -1,0 | 0,3 |
| chr7:55275031-55275531:downstream_25495 | chr7 | 55086724 | 55275031 | + | EGFR | NM | 250 | -0,4 | 1 | 1 | 4,84E-01 | 8,17E-01 | 66 | 74 | 481 | 363 | 102 | 75 | 468 | 372 | -2,6 | -2,2 | -0,4 |
| chr10:104576021-104576521:downstream_17820 | chr10 | 104503726 | 104576021 | + | WBP1L | NM | 12 | -1,3 | 2 | 1 | 4,84E-01 | NA | 1 | 0 | 3 | 7 | 3 | 12 | 38 | 32 | -3,6 | -2,2 | -1,4 |
| chr3:186507181-186507681:downstream_23385 | chr3 | 186507681 | 186524484 | - | RFC4 | NM | 19 | 0,8 | 1 | -1 | 4,84E-01 | NA | 2 | 7 | 1 | 9 | 22 | 17 | 42 | 56 | -0,3 | -1,3 | 1,0 |
| chr7:32620052-32620552:downstream_5536 | chr7 | 32620552 | 32758780 | - | DPY19L1P1 | NR | 4 | 1,7 | 2 | -1 | 4,84E-01 | NA | 2 | 1 | 10 | 5 | 1 | 0 | 6 | 10 | -2,5 | -4,2 | 1,7 |
| chr17:47006958-47007458:downstream_13211 | chr17 | 47007458 | 47022154 | - | SNF8 | NM | 17 | 0,9 | 1 | -1 | 4,84E-01 | NA | 5 | 4 | 4 | 2 | 34 | 22 | 32 | 31 | 0,6 | -0,2 | 0,8 |
| chr1:109639554-109640054:downstream_871 | chr1 | 109633402 | 109639554 | + | TMEM167B | NM | 18 | 0,9 | 1 | -1 | 4,84E-01 | NA | 6 | 4 | 4 | 0 | 44 | 35 | 32 | 21 | 1,2 | 0,6 | 0,6 |
| chr5:72879202-72879702:downstream_24438 | chr5 | 72861565 | 72879202 | + | UTP15 | NM | 22 | -0,7 | 1 | 1 | 4,85E-01 | NA | 21 | 31 | 7 | 11 | 46 | 39 | 8 | 11 | 1,5 | 2,2 | -0,7 |
| chr1:197007820-197008320:downstream_1280 | chr1 | 197008320 | 197036397 | - | F13B | NM | 4 | 1,5 | 2 | -1 | 4,86E-01 | NA | 7 | 4 | 11 | 2 | 2 | 0 | 1 | 7 | -0,4 | -1,8 | 1,5 |
| chrX:47305060-47305560:downstream_27129 | chrX | 47305560 | 47342345 | - | ZNF41 | NM | 11 | -0,9 | 1 | 1 | 4,86E-01 | NA | 1 | 6 | 17 | 13 | 7 | 10 | 19 | 20 | -2,2 | -1,3 | -0,9 |
| chr3:48724935-48725435:downstream_23209 | chr3 | 48725435 | 48754711 | - | IP6K2 | NM | 19 | 1,0 | 1 | -1 | 4,88E-01 | NA | 2 | 13 | 1 | 2 | 44 | 48 | 25 | 13 | 2,3 | 1,3 | 1,0 |
| chr9:130207651-130208151:downstream_26587 | chr9 | 130186652 | 130207651 | + | ZNF79 | NM | 61 | -0,5 | 1 | 1 | 4,88E-01 | 8,24E-01 | 95 | 58 | 15 | 16 | 154 | 109 | 26 | 11 | 2,3 | 2,8 | -0,5 |
| chr15:83425958-83426458:downstream_19729 | chr15 | 83419315 | 83425958 | + | SNHG21 | NR | 25 | 0,7 | 1 | -1 | 4,88E-01 | NA | 21 | 28 | 4 | 5 | 53 | 56 | 15 | 17 | 2,4 | 1,7 | 0,7 |
| chr8:90729126-90729626:downstream_6631 | chr8 | 90729626 | 90769939 | - | LOC101929709 | NR | 19 | 0,7 | 1 | -1 | 4,88E-01 | NA | 11 | 7 | 17 | 19 | 7 | 15 | 39 | 34 | -1,0 | -1,7 | 0,7 |
| chr15:70339629-70340129:downstream_19810 | chr15 | 70340129 | 70390256 | - | TLE3 | NM | 10 | -0,9 | 1 | 1 | 4,88E-01 | NA | 2 | 7 | 11 | 5 | 11 | 18 | 15 | 11 | -0,8 | 0,2 | -1,0 |
| chr11:33719153-33719653:downstream_9201 | chr11 | 33719653 | 33722286 | - | C11orf91 | NM | 10 | -1,2 | 2 | 1 | 4,88E-01 | NA | 11 | 24 | 0 | 4 | 19 | 18 | 1 | 1 | 3,1 | 4,1 | -1,0 |
| chr10:94210940-94211440:downstream_17604 | chr10 | 94211440 | 94333852 | - | IDE | NM | 14 | 1,0 | 1 | -1 | 4,89E-01 | NA | 0 | 6 | 1 | 11 | 8 | 9 | 38 | 40 | -1,1 | -2,2 | 1,2 |
| chr22:30166402-30166902:downstream_22964 | chr22 | 30163357 | 30166402 | + | UQCR10 | NM | 8 | 1,2 | 2 | -1 | 4,89E-01 | NA | 7 | 3 | 1 | 1 | 24 | 14 | 6 | 11 | 2,2 | 1,1 | 1,1 |
| chr9:130702360-130702860:downstream_26302 | chr9 | 130702860 | 130742812 | - | FAM102A | NM | 10 | -1,1 | 2 | 1 | 4,90E-01 | NA | 2 | 0 | 4 | 6 | 7 | 17 | 22 | 23 | -2,0 | -0,9 | -1,2 |
| chr2:170668571-170669071:downstream_22254 | chr2 | 170655321 | 170668571 | + | SSB | NM | 53 | -0,6 | 1 | 1 | 4,90E-01 | 8,26E-01 | 15 | 4 | 6 | 14 | 110 | 112 | 89 | 71 | -0,1 | 0,5 | -0,5 |
| chr16:30564584-30565084:downstream_20298 | chr16 | 30565084 | 30569642 | - | ZNF764 | NM | 15 | -0,8 | 1 | 1 | 4,90E-01 | NA | 11 | 3 | 8 | 7 | 20 | 33 | 18 | 16 | -0,2 | 0,6 | -0,8 |
| chr3:134073686-134074186:downstream_22998 | chr3 | 134074186 | 134094259 | - | AMOTL2 | NM | 12 | -0,9 | 1 | 1 | 4,90E-01 | NA | 5 | 14 | 1 | 5 | 25 | 38 | 6 | 7 | 1,5 | 2,4 | -0,8 |
| chr7:40171841-40172341:downstream_5800 | chr7 | 40172341 | 40174251 | - | MPLKIP | NM | 52 | 0,5 | 1 | -1 | 4,90E-01 | 8,26E-01 | 27 | 24 | 8 | 19 | 99 | 90 | 85 | 66 | 0,9 | 0,3 | 0,6 |
| chr10:114587491-114587991:downstream_8726 | chr10 | 114583254 | 114587491 | + | LOC103344931 | NR | 5 | -1,7 | 2 | 1 | 4,91E-01 | NA | 0 | 0 | 3 | 7 | 1 | 2 | 7 | 17 | -Inf | -3,1 | -Inf |
| chr1:28609002-28609502:downstream_589 | chr1 | 28585962 | 28609002 | + | SESN2 | NM | 20 | 1,3 | 2 | -1 | 4,91E-01 | NA | 9 | 5 | 0 | 0 | 60 | 81 | 1 | 7 | Inf | 4,0 | Inf |
| chr20:30779806-30780306:downstream_14961 | chr20 | 30780306 | 30795546 | - | PLAGL2 | NM | 11 | 1,0 | 1 | -1 | 4,92E-01 | NA | 2 | 10 | 3 | 2 | 18 | 21 | 13 | 21 | 1,2 | 0,2 | 1,0 |
| chr19:36729675-36730175:downstream_21606 | chr19 | 36705503 | 36729675 | + | ZNF146 | NM | 147 | 0,6 | 1 | -1 | 4,92E-01 | 8,26E-01 | 2 | 9 | 6 | 2 | 311 | 256 | 335 | 258 | 0,5 | -0,1 | 0,5 |
| chr12:10755863-10756363:downstream_18669 | chr12 | 10756363 | 10766208 | - | MAGOHB | NM | 31 | 0,6 | 1 | -1 | 4,92E-01 | 8,26E-01 | 54 | 34 | 11 | 21 | 37 | 48 | 21 | 25 | 1,5 | 0,9 | 0,6 |
| chr11:61166335-61166835:downstream_18347 | chr11 | 61159831 | 61166335 | + | TMEM216 | NM | 17 | 0,8 | 1 | -1 | 4,92E-01 | NA | 8 | 9 | 3 | 3 | 28 | 41 | 28 | 16 | 1,5 | 0,6 | 0,8 |
| chr12:2914587-2915087:downstream_9975 | chr12 | 2904107 | 2914587 | + | FKBP4 | NM | 4 | -1,4 | 2 | 1 | 4,92E-01 | NA | 5 | 4 | 6 | 2 | 6 | 5 | 1 | 2 | 0,3 | 1,8 | -1,5 |
| chr1:203823256-203823756:downstream_278 | chr1 | 203764750 | 203823256 | + | ZC3H11A | NM | 48 | 0,5 | 1 | -1 | 4,92E-01 | 8,27E-01 | 8 | 19 | 24 | 18 | 41 | 58 | 90 | 129 | -0,6 | -1,1 | 0,5 |
| chr12:105562912-105563412:downstream_18615 | chr12 | 105501491 | 105562912 | + | KIAA1033 | NM | 25 | 0,7 | 1 | -1 | 4,93E-01 | NA | 6 | 7 | 11 | 18 | 20 | 13 | 51 | 74 | -1,2 | -1,9 | 0,7 |
| chr2:55198826-55199326:downstream_22201 | chr2 | 55199326 | 55277734 | - | RTN4 | NM | 210 | -0,5 | 1 | 1 | 4,93E-01 | 8,27E-01 | 10 | 2 | 11 | 11 | 320 | 387 | 452 | 483 | -1,0 | -0,4 | -0,5 |
| chr18:33569286-33569786:downstream_20927 | chr18 | 33569786 | 33647557 | - | RPRD1A | NM | 5 | 1,2 | 2 | -1 | 4,93E-01 | NA | 4 | 5 | 4 | 2 | 7 | 4 | 7 | 9 | 0,6 | -0,5 | 1,1 |
| chr11:65089900-65090400:downstream_9440 | chr11 | 65082288 | 65089900 | + | CDC42EP2 | NM | 22 | -0,8 | 1 | 1 | 4,94E-01 | NA | 14 | 23 | 71 | 33 | 7 | 7 | 18 | 5 | -1,5 | -0,8 | -0,7 |
| chr12:110928192-110928692:downstream_10420 | chr12 | 110906231 | 110928192 | + | FAM216A | NM | 6 | 1,2 | 2 | -1 | 4,94E-01 | NA | 2 | 11 | 4 | 3 | 7 | 6 | 7 | 11 | 0,7 | -0,4 | 1,1 |
| chr17:67538470-67538970:downstream_12684 | chr17 | 67410837 | 67538470 | + | MAP2K6 | NM | 39 | -0,6 | 1 | 1 | 4,94E-01 | 8,27E-01 | 8 | 7 | 42 | 33 | 18 | 33 | 83 | 85 | -2,3 | -1,7 | -0,6 |
| chr8:54958426-54958926:downstream_26005 | chr8 | 54958926 | 55014577 | - | LYPLA1 | NM | 13 | 0,9 | 1 | -1 | 4,94E-01 | NA | 6 | 4 | 3 | 6 | 18 | 15 | 33 | 21 | 0,2 | -0,8 | 0,9 |
| chr2:63270599-63271099:downstream_22036 | chr2 | 63271099 | 63275656 | - | LOC100132215 | NR | 7 | 1,3 | 2 | -1 | 4,94E-01 | NA | 7 | 14 | 3 | 0 | 8 | 15 | 3 | 3 | 2,9 | 1,9 | 1,0 |
| chr12:49329491-49329991:downstream_10148 | chr12 | 49329991 | 49351252 | - | ARF3 | NM | 59 | -0,5 | 1 | 1 | 4,94E-01 | 8,27E-01 | 7 | 18 | 29 | 49 | 49 | 66 | 128 | 124 | -1,6 | -1,1 | -0,5 |
| chr10:126675917-126676417:downstream_17529 | chr10 | 126676417 | 126849624 | - | CTBP2 | NM | 7 | 1,1 | 2 | -1 | 4,94E-01 | NA | 8 | 1 | 10 | 8 | 4 | 2 | 11 | 14 | -1,0 | -2,1 | 1,1 |
| chr17:66042970-66043470:downstream_13317 | chr17 | 66031847 | 66042970 | + | KPNA2 | NM | 24 | 0,7 | 1 | -1 | 4,95E-01 | NA | 9 | 18 | 7 | 12 | 32 | 35 | 25 | 52 | 0,5 | -0,2 | 0,7 |
| chr13:20247599-20248099:downstream_10645 | chr13 | 20207787 | 20247599 | + | MPHOSPH8 | NM | 8 | -1,3 | 2 | 1 | 4,95E-01 | NA | 7 | 0 | 10 | 10 | 19 | 0 | 8 | 14 | -1,6 | -0,3 | -1,3 |
| chr14:53241705-53242205:downstream_19409 | chr14 | 53196882 | 53241705 | + | STYX | NM | 17 | 0,7 | 1 | -1 | 4,95E-01 | NA | 14 | 10 | 11 | 5 | 19 | 27 | 22 | 27 | 0,5 | -0,1 | 0,7 |
| chr6:102517958-102518458:downstream_24788 | chr6 | 101846860 | 102517958 | + | GRIK2 | NM | 10 | -1,4 | 2 | 1 | 4,96E-01 | NA | 0 | 0 | 8 | 4 | 2 | 3 | 28 | 35 | -Inf | -3,8 | -Inf |
| chr7:44865987-44866487:downstream_25551 | chr7 | 44866487 | 44887725 | - | H2AFV | NM | 37 | 0,6 | 1 | -1 | 4,96E-01 | 8,29E-01 | 10 | 6 | 6 | 12 | 50 | 44 | 82 | 89 | -0,2 | -0,8 | 0,6 |
| chr9:86274377-86274877:downstream_26565 | chr9 | 86274877 | 86323168 | - | UBQLN1 | NM | 21 | -0,8 | 1 | 1 | 4,96E-01 | NA | 7 | 0 | 11 | 7 | 41 | 20 | 38 | 47 | -1,3 | -0,4 | -0,9 |
| chr3:194409766-194410266:downstream_3342 | chr3 | 194406621 | 194409766 | + | FAM43A | NM | 9 | 1,3 | 2 | -1 | 4,96E-01 | NA | 0 | 22 | 0 | 3 | 8 | 25 | 8 | 6 | 2,8 | 1,2 | 1,5 |
| chr9:125579875-125580375:downstream_7306 | chr9 | 125580375 | 125590935 | - | PDCL | NM | 7 | 1,1 | 2 | -1 | 4,97E-01 | NA | 5 | 3 | 13 | 10 | 1 | 3 | 15 | 7 | -1,5 | -2,6 | 1,1 |
| chr10:82192753-82193253:downstream_17565 | chr10 | 82167584 | 82192753 | + | FAM213A | NM | 33 | -0,5 | 1 | 1 | 4,97E-01 | 8,30E-01 | 24 | 18 | 35 | 21 | 38 | 48 | 35 | 43 | -0,4 | 0,2 | -0,6 |
| chr3:150348234-150348734:downstream_2729 | chr3 | 150321065 | 150348234 | + | SELT | NM | 21 | 0,7 | 1 | -1 | 4,98E-01 | NA | 33 | 40 | 18 | 6 | 27 | 22 | 15 | 10 | 1,6 | 1,0 | 0,7 |
| chr9:117408703-117409203:downstream_7276 | chr9 | 117373705 | 117408703 | + | C9orf91 | NM | 7 | -1,0 | 1 | 1 | 4,98E-01 | NA | 3 | 6 | 13 | 7 | 6 | 9 | 8 | 7 | -1,1 | -0,1 | -1,0 |
| chr1:149784279-149784779:downstream_964 | chr1 | 149784779 | 149785236 | - | HIST2H3D | NM | 7 | -1,1 | 2 | 1 | 4,98E-01 | NA | 5 | 2 | 7 | 10 | 2 | 13 | 7 | 10 | -1,3 | -0,2 | -1,1 |
| chr19:55903451-55903951:downstream_21382 | chr19 | 55897299 | 55903451 | + | RPL28 | NM | 18 | 0,8 | 1 | -1 | 4,98E-01 | NA | 5 | 6 | 3 | 6 | 27 | 24 | 43 | 30 | 0,3 | -0,5 | 0,9 |
| chr10:105886143-105886643:downstream_17764 | chr10 | 105881815 | 105886143 | + | SFR1 | NM | 21 | -0,8 | 1 | 1 | 4,98E-01 | NA | 2 | 7 | 1 | 7 | 31 | 59 | 29 | 30 | -0,1 | 0,6 | -0,7 |
| chr15:50652607-50653107:downstream_11720 | chr15 | 50652525 | 50652607 | + | MIR4712 | NR | 39 | 0,5 | 1 | -1 | 4,98E-01 | 8,30E-01 | 67 | 37 | 35 | 43 | 27 | 36 | 35 | 33 | 0,4 | -0,1 | 0,5 |
| chr9:125611231-125611731:downstream_26476 | chr9 | 125611731 | 125667562 | - | RC3H2 | NM | 31 | -0,6 | 1 | 1 | 4,98E-01 | 8,30E-01 | 13 | 31 | 38 | 41 | 24 | 34 | 31 | 39 | -0,8 | -0,3 | -0,6 |
| chr21:26957467-26957967:downstream_22685 | chr21 | 26957967 | 26979801 | - | MRPL39 | NM | 13 | -0,9 | 1 | 1 | 4,99E-01 | NA | 15 | 19 | 4 | 6 | 35 | 15 | 3 | 5 | 1,8 | 2,7 | -0,9 |
| chr8:90940095-90940595:downstream_26037 | chr8 | 90914095 | 90940095 | + | OSGIN2 | NM | 15 | -0,9 | 1 | 1 | 4,99E-01 | NA | 15 | 0 | 18 | 16 | 15 | 15 | 17 | 21 | -1,2 | -0,3 | -0,9 |
| chr11:45834567-45835067:downstream_18300 | chr11 | 45825622 | 45834567 | + | SLC35C1 | NM | 8 | -1,2 | 2 | 1 | 4,99E-01 | NA | 4 | 0 | 3 | 10 | 14 | 5 | 8 | 17 | -1,6 | -0,5 | -1,2 |
| chr1:118405606-118406106:downstream_16765 | chr1 | 118406106 | 118472302 | - | GDAP2 | NM | 10 | -1,0 | 1 | 1 | 4,99E-01 | NA | 2 | 1 | 10 | 14 | 8 | 4 | 18 | 24 | -2,8 | -1,8 | -1,0 |
| chr5:130730382-130730882:downstream_24058 | chr5 | 130599701 | 130730382 | + | CDC42SE2 | NM | 32 | -1,2 | 2 | 1 | 5,00E-01 | 8,32E-01 | 0 | 0 | 7 | 4 | 7 | 9 | 139 | 93 | -Inf | -3,9 | -Inf |

| | | | | | | | | | | | | | | | | | | | | | | | |
|---|---|---|---|---|---|---|---|---|---|---|---|---|---|---|---|---|---|---|---|---|---|---|---|
| chr2:228425938-228426438:downstream_21732 | chr2 | 228336887 | 228425938 | + | AGFG1 | NM | 26 | 0,6 | 1 | -1 | 5,00E-01 | NA | 5 | 8 | 11 | 13 | 23 | 19 | 64 | 64 | -0,9 | -1,6 | 0,7 |
| chr12:122715593-122716093:downstream_10468 | chr12 | 122716093 | 122751068 | - | VPS33A | NM | 20 | 0,7 | 1 | -1 | 5,00E-01 | NA | 6 | 11 | 8 | 11 | 25 | 16 | 42 | 39 | -0,2 | -0,9 | 0,7 |
| chr1:15988217-15988717:downstream_480 | chr1 | 15986363 | 15988217 | + | RSC1A1 | NM | 22 | -0,7 | 1 | 1 | 5,01E-01 | NA | 12 | 4 | 18 | 39 | 24 | 10 | 40 | 33 | -1,8 | -1,1 | -0,7 |
| chr14:51099797-51100297:downstream_10939 | chr14 | 51100297 | 51135071 | - | SAV1 | NM | 14 | 0,9 | 1 | -1 | 5,01E-01 | NA | 17 | 6 | 3 | 2 | 26 | 38 | 11 | 11 | 2,4 | 1,5 | 0,8 |
| chr7:43666978-43667478:downstream_5548 | chr7 | 43622691 | 43666978 | + | STK17A | NM | 26 | 0,7 | 1 | -1 | 5,01E-01 | NA | 32 | 54 | 10 | 8 | 34 | 42 | 19 | 7 | 2,3 | 1,5 | 0,7 |
| chr9:135972606-135973106:downstream_26471 | chr9 | 135973106 | 136024607 | - | RALGDS | NM | 6 | 1,5 | 2 | -1 | 5,01E-01 | NA | 2 | 15 | 0 | 1 | 15 | 11 | 3 | 2 | 4,4 | 2,6 | 1,9 |
| chr15:75950968-75951468:downstream_11818 | chr15 | 75941347 | 75950968 | + | SNX33 | NM | 6 | -1,3 | 2 | 1 | 5,01E-01 | NA | 5 | 0 | 6 | 10 | 2 | 11 | 6 | 10 | -1,7 | -0,3 | -1,3 |
| chr12:53700965-53701465:downstream_10203 | chr12 | 53693469 | 53700965 | + | C12orf10 | NM | 10 | 1,0 | 1 | -1 | 5,02E-01 | NA | 10 | 23 | 4 | 9 | 18 | 3 | 7 | 10 | 1,3 | 0,3 | 1,0 |
| chr2:228226373-228226873:downstream_2463 | chr2 | 228226873 | 228244022 | - | TM4SF20 | NM | 348 | -0,4 | 1 | 1 | 5,02E-01 | 8,34E-01 | 50 | 24 | 78 | 91 | 417 | 504 | 881 | 735 | -1,2 | -0,8 | -0,4 |
| chr21:34636818-34637318:downstream_22663 | chr21 | 34602199 | 34636818 | + | IFNAR2 | NM | 16 | -0,9 | 1 | 1 | 5,03E-01 | NA | 2 | 0 | 13 | 21 | 6 | 6 | 42 | 39 | -3,8 | -2,8 | -1,0 |
| chr20:17588652-17589152:downstream_22411 | chr20 | 17550598 | 17588652 | + | DSTN | NM | 740 | -0,3 | 0 | 1 | 5,03E-01 | 8,34E-01 | 711 | 551 | 679 | 563 | 995 | 896 | 790 | 732 | 0,0 | 0,3 | -0,3 |
| chr17:46200105-46200605:downstream_20732 | chr17 | 46184919 | 46200105 | + | SNX11 | NM | 10 | 0,9 | 1 | -1 | 5,03E-01 | NA | 7 | 22 | 7 | 9 | 15 | 4 | 7 | 12 | 0,9 | -0,1 | 0,9 |
| chr5:140697556-140698056:downstream_4486 | chr5 | 140698056 | 140700351 | - | TAF7 | NM | 63 | 0,5 | 1 | -1 | 5,03E-01 | 8,34E-01 | 31 | 22 | 6 | 13 | 160 | 122 | 71 | 76 | 1,5 | 0,9 | 0,6 |
| chr22:40762575-40763075:downstream_22735 | chr22 | 40742503 | 40762575 | + | ADSL | NM | 17 | 0,9 | 1 | -1 | 5,04E-01 | NA | 5 | 0 | 4 | 9 | 11 | 10 | 53 | 45 | -1,4 | -2,3 | 0,8 |
| chr3:50686728-50687228:downstream_23262 | chr3 | 50649292 | 50686728 | + | MAPKAPK3 | NM | 10 | 1,0 | 1 | -1 | 5,04E-01 | NA | 7 | 14 | 4 | 2 | 13 | 18 | 17 | 3 | 1,7 | 0,7 | 1,0 |
| chr12:49521066-49521566:downstream_10150 | chr12 | 49521566 | 49525304 | - | TUBA1B | NM | 168 | -0,4 | 1 | 1 | 5,04E-01 | 8,34E-01 | 11 | 18 | 25 | 15 | 315 | 322 | 299 | 343 | -0,5 | 0,0 | -0,4 |
| chr15:72490869-72491369:downstream_19667 | chr15 | 72491369 | 72523727 | - | PKM | NM | 1735 | -0,5 | 1 | 1 | 5,05E-01 | 8,34E-01 | 7 | 3 | 3 | 5 | 3928 | 4836 | 2387 | 2714 | 0,3 | 0,8 | -0,5 |
| chr10:75195685-75196185:downstream_17720 | chr10 | 75196185 | 75255782 | - | PPP3CB | NM | 6 | -1,3 | 2 | 1 | 5,05E-01 | NA | 2 | 0 | 11 | 3 | 3 | 5 | 14 | 13 | -3,1 | -1,7 | -1,4 |
| chr8:22108680-22109180:downstream_6484 | chr8 | 22102618 | 22108680 | + | POLR3D | NM | 30 | -0,6 | 1 | 1 | 5,05E-01 | 8,34E-01 | 32 | 26 | 19 | 10 | 43 | 72 | 19 | 18 | 1,0 | 1,6 | -0,6 |
| chr8:109499136-109499636:downstream_6680 | chr8 | 109455852 | 109499136 | + | EMC2 | NM | 38 | 0,6 | 1 | -1 | 5,05E-01 | 8,34E-01 | 8 | 11 | 13 | 7 | 52 | 51 | 100 | 60 | -0,1 | -0,6 | 0,5 |
| chr18:52568239-52568739:downstream_20842 | chr18 | 52568739 | 52626739 | - | CCDC68 | NM | 12 | -1,3 | 2 | 1 | 5,06E-01 | NA | 1 | 0 | 11 | 2 | 10 | 3 | 35 | 35 | -4,0 | -2,5 | -1,5 |
| chr1:161069845-161070345:downstream_228 | chr1 | 161070345 | 161087866 | - | PFDN2 | NM | 29 | 0,7 | 1 | -1 | 5,06E-01 | 8,35E-01 | 4 | 9 | 4 | 2 | 65 | 60 | 39 | 48 | 1,1 | 0,5 | 0,6 |
| chr3:16347594-16348094:downstream_2844 | chr3 | 16306666 | 16347594 | + | OXNAD1 | NM | 11 | 0,8 | 1 | -1 | 5,06E-01 | NA | 11 | 8 | 8 | 20 | 7 | 5 | 19 | 11 | -0,6 | -1,4 | 0,9 |
| chr21:45193045-45193545:downstream_15350 | chr21 | 45193545 | 45196256 | - | CSTB | NM | 16 | 0,9 | 1 | -1 | 5,06E-01 | NA | 2 | 8 | 1 | 3 | 37 | 22 | 18 | 36 | 1,1 | 0,1 | 1,0 |
| chr20:2815460-2815960:downstream_22498 | chr20 | 2815960 | 2821889 | - | PCED1A | NM | 27 | -0,6 | 1 | 1 | 5,06E-01 | NA | 14 | 15 | 14 | 15 | 33 | 64 | 33 | 32 | 0,0 | 0,6 | -0,6 |
| chr11:36504816-36505316:downstream_18363 | chr11 | 36505316 | 36531863 | - | TRAF6 | NM | 15 | 0,8 | 1 | -1 | 5,07E-01 | NA | 11 | 15 | 10 | 16 | 19 | 6 | 24 | 20 | 0,0 | -0,8 | 0,8 |
| chr14:77807613-77808113:downstream_11251 | chr14 | 77808113 | 77843396 | - | TMED8 | NM | 17 | -0,8 | 1 | 1 | 5,07E-01 | NA | 25 | 11 | 51 | 25 | 5 | 6 | 8 | 5 | -1,1 | -0,3 | -0,8 |
| chr1:43390545-43391045:downstream_683 | chr1 | 43391045 | 43424847 | - | SLC2A1 | NM | 5 | 1,2 | 2 | -1 | 5,07E-01 | NA | 10 | 4 | 1 | 2 | 9 | 8 | 3 | 6 | 2,2 | 1,0 | 1,2 |
| chr13:92006829-92007329:downstream_19027 | chr13 | 92000073 | 92006829 | + | MIR17HG | NR | 50 | 0,5 | 1 | -1 | 5,07E-01 | 8,35E-01 | 56 | 58 | 19 | 20 | 93 | 77 | 39 | 41 | 1,5 | 1,1 | 0,5 |
| chr6:74224972-74225472:downstream_5232 | chr6 | 74225472 | 74230755 | - | EEF1A1 | NM | 52 | 0,5 | 1 | -1 | 5,07E-01 | 8,35E-01 | 54 | 38 | 25 | 25 | 96 | 61 | 45 | 75 | 0,9 | 0,4 | 0,5 |
| chr19:40735723-40736223:downstream_20993 | chr19 | 40736223 | 40791302 | - | AKT2 | NM | 6 | 1,2 | 2 | -1 | 5,08E-01 | NA | 11 | 10 | 3 | 2 | 6 | 9 | 6 | 2 | 2,2 | 1,0 | 1,2 |
| chr12:64541613-64542113:downstream_9873 | chr12 | 64238540 | 64541613 | + | SRGAP1 | NM | 6 | -1,5 | 2 | 1 | 5,08E-01 | NA | 0 | 0 | 3 | 7 | 2 | 2 | 15 | 22 | -Inf | -3,4 | -Inf |
| chr15:34432374-34432874:downstream_11439 | chr15 | 34432874 | 34502297 | - | KATNBL1 | NM | 13 | -0,8 | 1 | 1 | 5,08E-01 | NA | 10 | 4 | 24 | 16 | 8 | 12 | 17 | 16 | -1,6 | -0,7 | -0,8 |
| chr2:128461307-128461807:downstream_22320 | chr2 | 128461807 | 128568761 | - | WDR33 | NM | 4 | -1,6 | 2 | 1 | 5,08E-01 | NA | 0 | 1 | 7 | 5 | 1 | 3 | 11 | 5 | -3,6 | -2,1 | -1,5 |
| chr7:2273425-2273925:downstream_5681 | chr7 | 2273925 | 2281833 | - | FTSJ2 | NM | 19 | -0,7 | 1 | 1 | 5,09E-01 | NA | 5 | 10 | 6 | 7 | 46 | 35 | 15 | 29 | 0,2 | 0,9 | -0,7 |
| chr7:54819439-54819939:downstream_25720 | chr7 | 54819939 | 54826939 | - | SEC61G | NM | 30 | 0,6 | 1 | -1 | 5,10E-01 | 8,38E-01 | 18 | 8 | 8 | 6 | 54 | 57 | 39 | 51 | 0,9 | 0,3 | 0,6 |
| chr3:196280558-196281058:downstream_3349 | chr3 | 196281058 | 196295413 | - | WDR53 | NM | 17 | 0,9 | 1 | -1 | 5,10E-01 | NA | 1 | 10 | 1 | 5 | 30 | 23 | 32 | 33 | 0,7 | -0,3 | 1,0 |
| chr8:125474237-125474737:downstream_26098 | chr8 | 125474737 | 125486804 | - | RNF139-AS1 | NR | 5 | -1,8 | 3 | 1 | 5,10E-01 | NA | 2 | 18 | 0 | 2 | 13 | 2 | 0 | 0 | 3,7 | Inf | -Inf |
| chr17:68162601-68163101:downstream_13325 | chr17 | 68163101 | 68165543 | - | KCNJ2-AS1 | NR | 4 | -1,4 | 2 | 1 | 5,10E-01 | NA | 2 | 3 | 1 | 11 | 1 | 9 | 1 | 7 | -1,2 | 0,3 | -1,4 |
| chr3:49758408-49758908:downstream_23169 | chr3 | 49758908 | 49761407 | - | GMPPB | NM | 19 | -0,7 | 1 | 1 | 5,10E-01 | NA | 25 | 26 | 33 | 33 | 12 | 8 | 11 | 6 | -0,4 | 0,2 | -0,6 |
| chr20:33864804-33865304:downstream_14787 | chr20 | 33814538 | 33864804 | + | MMP24 | NM | 6 | 1,1 | 2 | -1 | 5,11E-01 | NA | 5 | 8 | 11 | 7 | 4 | 1 | 3 | 12 | -0,5 | -1,6 | 1,1 |
| chr1:207975287-207975787:downstream_1344 | chr1 | 207975787 | 207975868 | - | MIR29B2 | NR | 25 | -0,8 | 1 | 1 | 5,11E-01 | NA | 28 | 14 | 79 | 61 | 7 | 0 | 6 | 9 | -1,7 | -1,0 | -0,7 |
| chr16:50269219-50269719:downstream_20163 | chr16 | 50186828 | 50269219 | + | PAPD5 | NM | 4 | -1,4 | 2 | 1 | 5,11E-01 | NA | 2 | 0 | 4 | 8 | 5 | 1 | 4 | 7 | -2,3 | -0,9 | -1,5 |
| chr12:27955973-27956473:downstream_10114 | chr12 | 27933186 | 27955973 | + | KLHL42 | NM | 9 | 0,9 | 1 | -1 | 5,11E-01 | NA | 7 | 4 | 8 | 5 | 8 | 8 | 14 | 22 | -0,4 | -1,2 | 0,8 |
| chr16:3707537-3708037:downstream_20258 | chr16 | 3708037 | 3767598 | - | TRAP1 | NM | 11 | 1,0 | 1 | -1 | 5,12E-01 | NA | 2 | 10 | 4 | 1 | 22 | 20 | 13 | 18 | 1,3 | 0,5 | 0,8 |
| chr4:48863834-48864334:downstream_23781 | chr4 | 48833059 | 48863834 | + | OCIAD1 | NM | 6 | -1,1 | 2 | 1 | 5,12E-01 | NA | 4 | 1 | 7 | 5 | 4 | 11 | 10 | 6 | -1,2 | -0,1 | -1,2 |
| chr15:59929760-59930260:downstream_11739 | chr15 | 59930260 | 59949737 | - | GTF2A2 | NM | 7 | -1,0 | 2 | 1 | 5,12E-01 | NA | 13 | 4 | 7 | 7 | 10 | 9 | 1 | 6 | 0,2 | 1,4 | -1,1 |
| chr1:151263772-151264272:downstream_17079 | chr1 | 151264272 | 151300191 | - | PI4KB | NM | 5 | 1,4 | 2 | -1 | 5,12E-01 | NA | 0 | 12 | 0 | 3 | 6 | 9 | 6 | 7 | 1,8 | 0,2 | 1,6 |
| chr7:23349180-23349680:downstream_5737 | chr7 | 23338939 | 23349180 | + | MALSU1 | NM | 31 | 0,6 | 1 | -1 | 5,13E-01 | 8,41E-01 | 26 | 31 | 21 | 29 | 47 | 13 | 25 | 56 | 0,2 | -0,4 | 0,6 |
| chr6:28137373-28137873:downstream_4967 | chr6 | 28129538 | 28137373 | + | ZNF192P1 | NR | 11 | 1,2 | 2 | -1 | 5,13E-01 | NA | 26 | 29 | 0 | 2 | 12 | 15 | 3 | 0 | 5,1 | 3,3 | 1,8 |
| chr12:104343480-104343980:downstream_10389 | chr12 | 104343980 | 104350993 | - | C12orf73 | NM | 19 | 0,7 | 1 | -1 | 5,13E-01 | NA | 37 | 33 | 21 | 11 | 17 | 14 | 13 | 9 | 1,2 | 0,5 | 0,6 |
| chr9:14610568-14611068:downstream_6810 | chr9 | 14611068 | 14693480 | - | ZDHHC21 | NM | 14 | -0,8 | 1 | 1 | 5,13E-01 | NA | 3 | 1 | 11 | 16 | 7 | 12 | 36 | 29 | -2,7 | -1,8 | -0,9 |
| chr5:150087808-150088308:downstream_24086 | chr5 | 150088308 | 150138657 | - | DCTN4 | NM | 27 | 0,7 | 1 | -1 | 5,13E-01 | NA | 4 | 11 | 13 | 11 | 13 | 37 | 60 | 65 | -0,7 | -1,3 | 0,7 |
| chr14:101455537-101456037:downstream_11356 | chr14 | 101455466 | 101455537 | + | SNORD114-28 | NR | 7 | 1,2 | 2 | -1 | 5,14E-01 | NA | 7 | 2 | 0 | 2 | 15 | 15 | 7 | 9 | 2,4 | 1,0 | 1,4 |
| chr4:103715039-103715539:downstream_23902 | chr4 | 103715539 | 103790050 | - | UBE2D3 | NM | 11 | 1,0 | 2 | -1 | 5,14E-01 | NA | 12 | 1 | 0 | 6 | 24 | 10 | 18 | 16 | 1,2 | 0,0 | 1,2 |
| chr16:31500295-31500795:downstream_12319 | chr16 | 31500795 | 31519740 | - | C16orf58 | NM | 84 | -0,5 | 1 | 1 | 5,14E-01 | 8,41E-01 | 4 | 7 | 4 | 8 | 145 | 208 | 154 | 142 | -0,2 | 0,3 | -0,4 |
| chr7:77269388-77269888:downstream_25698 | chr7 | 77166772 | 77269388 | + | PTPN12 | NM | 46 | -1,1 | 2 | 1 | 5,14E-01 | 8,41E-01 | 1 | 0 | 6 | 7 | 17 | 26 | 178 | 130 | -4,0 | -2,8 | -1,2 |
| chr12:112463992-112464492:downstream_10427 | chr12 | 112464492 | 112546635 | - | NAA25 | NM | 16 | 0,7 | 1 | -1 | 5,14E-01 | NA | 3 | 9 | 18 | 10 | 8 | 11 | 39 | 34 | -1,2 | -2,0 | 0,7 |
| chr20:280963-281463:downstream_14837 | chr20 | 278203 | 280963 | + | ZCCHC3 | NM | 19 | 1,0 | 1 | -1 | 5,15E-01 | NA | 9 | 3 | 0 | 2 | 50 | 57 | 21 | 14 | 2,9 | 1,6 | 1,2 |
| chr20:49202416-49202916:downstream_15081 | chr20 | 49202322 | 49202416 | + | MIR645 | NR | 6 | -1,2 | 2 | 1 | 5,15E-01 | NA | 5 | 0 | 6 | 6 | 14 | 2 | 7 | 8 | -1,2 | 0,1 | -1,3 |
| chr20:25278648-25279148:downstream_14932 | chr20 | 25228705 | 25278648 | + | PYGB | NM | 43 | 0,7 | 1 | -1 | 5,15E-01 | 8,41E-01 | 6 | 5 | 10 | 0 | 67 | 77 | 72 | 105 | 0,1 | -0,3 | 0,4 |
| chr3:194788507-194789007:downstream_23514 | chr3 | 194789007 | 194991895 | - | XXYLT1 | NM | 21 | 0,8 | 1 | -1 | 5,15E-01 | NA | 7 | 0 | 15 | 9 | 11 | 9 | 65 | 50 | -1,9 | -2,6 | 0,7 |
| chr14:70232499-70232999:downstream_11201 | chr14 | 70232999 | 70234430 | - | LOC100289511 | NR | 7 | 1,1 | 2 | -1 | 5,15E-01 | NA | 6 | 18 | 6 | 1 | 9 | 9 | 8 | 2 | 1,9 | 0,8 | 1,1 |
| chr15:65442283-65442783:downstream_11765 | chr15 | 65442783 | 65477563 | - | CLPX | NM | 27 | -0,6 | 1 | 1 | 5,15E-01 | NA | 14 | 12 | 15 | 38 | 23 | 34 | 40 | 39 | -1,1 | -0,5 | -0,6 |
| chr1:196716634-196717134:downstream_16592 | chr1 | 196621007 | 196716634 | + | CFH | NM | 145 | -0,6 | 1 | 1 | 5,16E-01 | 8,41E-01 | 5 | 0 | 10 | 6 | 238 | 157 | 405 | 339 | -1,7 | -0,9 | -0,8 |
| chr15:44064804-44065304:downstream_11413 | chr15 | 44038589 | 44064804 | + | PDIA3 | NM | 28 | 0,6 | 1 | -1 | 5,16E-01 | 8,41E-01 | 7 | 17 | 13 | 10 | 41 | 34 | 61 | 45 | 0,1 | -0,5 | 0,6 |
| chr1:43318146-43318646:downstream_17438 | chr1 | 43312243 | 43318146 | + | ZNF691 | NM | 7 | 1,0 | 2 | -1 | 5,16E-01 | NA | 5 | 8 | 1 | 2 | 11 | 13 | 7 | 8 | 1,7 | 0,6 | 1,1 |
| chr1:202859723-202860223:downstream_16860 | chr1 | 202860223 | 202897772 | - | KLHL12 | NM | 10 | 0,8 | 1 | -1 | 5,16E-01 | NA | 10 | 7 | 7 | 7 | 10 | 12 | 17 | 14 | 0,3 | -0,5 | 0,8 |
| chr8:102208765-102209265:downstream_26199 | chr8 | 102209265 | 102218292 | - | ZNF706 | NM | 32 | 0,6 | 1 | -1 | 5,16E-01 | 8,41E-01 | 29 | 29 | 7 | 5 | 54 | 87 | 24 | 19 | 2,3 | 1,7 | 0,6 |
| chr14:74962271-74962771:downstream_19260 | chr14 | 74960422 | 74962271 | + | ISCA2 | NM | 18 | 0,7 | 1 | -1 | 5,17E-01 | NA | 15 | 12 | 17 | 14 | 8 | 21 | 31 | 24 | -0,2 | -0,9 | 0,7 |
| chrX:10412849-10413349:downstream_26914 | chrX | 10413349 | 10851809 | - | MID1 | NM | 34 | -0,8 | 1 | 1 | 5,17E-01 | 8,41E-01 | 2 | 0 | 17 | 11 | 15 | 19 | 127 | 82 | -3,5 | -2,6 | -0,9 |
| chr20:5975852-5976352:downstream_22470 | chr20 | 5931297 | 5975852 | + | MCM8 | NM | 41 | 0,5 | 1 | -1 | 5,17E-01 | 8,41E-01 | 36 | 15 | 61 | 87 | 17 | 8 | 49 | 57 | -1,5 | -2,1 | 0,6 |
| chr16:66444803-66445303:downstream_12398 | chr16 | 66442426 | 66444803 | + | LINC00920 | NR | 6 | -1,2 | 2 | 1 | 5,17E-01 | NA | 1 | 2 | 4 | 12 | 2 | 6 | 10 | 13 | -2,6 | -1,5 | -1,1 |
| chr12:124420455-124420955:downstream_18486 | chr12 | 124420955 | 124457532 | - | CCDC92 | NM | 23 | -0,7 | 1 | 1 | 5,17E-01 | NA | 6 | 3 | 7 | 9 | 49 | 25 | 51 | 34 | -0,9 | -0,2 | -0,7 |
| chr1:40306205-40306705:downstream_659 | chr1 | 40306705 | 40349177 | - | TRIT1 | NM | 29 | 0,6 | 1 | -1 | 5,17E-01 | 8,41E-01 | 17 | 13 | 18 | 9 | 41 | 33 | 45 | 52 | 0,1 | -0,4 | 0,5 |
| chr4:84184476-84184976:downstream_3763 | chr4 | 84184976 | 84206067 | - | COQ2 | NM | 72 | 0,6 | 1 | -1 | 5,18E-01 | 8,41E-01 | 6 | 5 | 3 | 7 | 127 | 104 | 143 | 180 | 0,2 | -0,5 | 0,7 |
| chr5:125962944-125963444:downstream_4081 | chr5 | 125936606 | 125962944 | + | PHAX | NM | 14 | 0,9 | 1 | -1 | 5,19E-01 | NA | 9 | 5 | 1 | 2 | 30 | 32 | 13 | 20 | 1,8 | 0,9 | 0,9 |
| chr2:55511607-55512107:downstream_1977 | chr2 | 55509454 | 55511607 | + | PRORSD1P | NR | 18 | 0,7 | 1 | -1 | 5,19E-01 | NA | 8 | 17 | 4 | 3 | 28 | 46 | 17 | 19 | 1,8 | 1,1 | 0,7 |
| chr11:47595013-47595513:downstream_18236 | chr11 | 47586981 | 47595013 | + | PTPMT1 | NM | 19 | 0,7 | 1 | -1 | 5,19E-01 | NA | 11 | 39 | 13 | 9 | 31 | 15 | 18 | 15 | 1,2 | 0,5 | 0,7 |
| chr9:100845365-100845865:downstream_7222 | chr9 | 100818958 | 100845365 | + | NANS | NM | 24 | 0,7 | 1 | -1 | 5,19E-01 | NA | 4 | 9 | 7 | 7 | 24 | 35 | 45 | 60 | -0,2 | -0,8 | 0,7 |
| chr3:194307901-194308401:downstream_23479 | chr3 | 194308401 | 194354150 | - | TMEM44 | NM | 12 | 0,8 | 1 | -1 | 5,19E-01 | NA | 7 | 8 | 10 | 9 | 10 | 10 | 22 | 19 | -0,3 | -1,1 | 0,8 |
| chr20:35519726-35520226:downstream_14790 | chr20 | 35520226 | 35580246 | - | SAMHD1 | NM | 8 | -1,0 | 2 | 1 | 5,19E-01 | NA | 4 | 2 | 10 | 11 | 4 | 9 | 8 | 13 | -1,8 | -0,7 | -1,0 |
| chr14:103811361-103811861:downstream_19218 | chr14 | 103800338 | 103811361 | + | EIF5 | NM | 94 | 0,5 | 1 | -1 | 5,19E-01 | 8,42E-01 | 25 | 36 | 6 | 9 | 327 | 176 | 103 | 74 | 2,1 | 1,5 | 0,6 |
| chr1:50906434-50906934:downstream_20 | chr1 | 50906934 | 51425936 | - | FAF1 | NM | 5 | 1,8 | 3 | -1 | 5,19E-01 | NA | 2 | 0 | 1 | 21 | 0 | 0 | 3 | 11 | -3,8 | -Inf | Inf |
| chr3:51435336-51435836:downstream_2982 | chr3 | 51428698 | 51435336 | + | RBM15B | NM | 19 | 0,8 | 1 | -1 | 5,19E-01 | NA | 11 | 5 | 0 | 4 | 47 | 39 | 29 | 16 | 2,0 | 0,9 | 1,0 |
| chr3:160152749-160153249:downstream_23437 | chr3 | 160117091 | 160152749 | + | SMC4 | NM | 33 | 0,6 | 1 | -1 | 5,20E-01 | 8,42E-01 | 7 | 6 | 11 | 7 | 51 | 27 | 65 | 89 | -0,4 | -1,0 | 0,6 |
| chr19:54711515-54712015:downstream_14636 | chr19 | 54704725 | 54711515 | + | RPS9 | NM | 55 | -0,6 | 1 | 1 | 5,21E-01 | 8,42E-01 | 28 | 11 | 6 | 28 | 149 | 82 | 57 | 83 | 0,2 | 0,7 | -0,5 |
| chr9:127620170-127620670:downstream_26572 | chr9 | 127615695 | 127620170 | + | WDR38 | NM | 27 | -0,7 | 1 | 1 | 5,21E-01 | NA | 31 | 94 | 22 | 26 | 11 | 24 | 6 | 3 | 1,4 | 2,0 | -0,6 |
| chr12:32898584-32899084:downstream_18545 | chr12 | 32832133 | 32898584 | + | DNM1L | NM | 18 | 0,7 | 1 | -1 | 5,21E-01 | NA | 11 | 18 | 17 | 23 | 11 | 12 | 19 | 30 | -0,5 | -1,1 | 0,7 |
| chr1:761085-761585:downstream_349 | chr1 | 761585 | 762902 | - | LINC00115 | NR | 15 | 0,9 | 1 | -1 | 5,21E-01 | NA | 28 | 20 | 1 | 2 | 23 | 33 | 6 | 3 | 3,6 | 2,7 | 1,0 |
| chr6:28246000-28246500:downstream_25365 | chr6 | 28234787 | 28246000 | + | ZSCAN26 | NM | 9 | -1,0 | 2 | 1 | 5,21E-01 | NA | 5 | 0 | 7 | 5 | 11 | 14 | 17 | 11 | -1,3 | -0,2 | -1,1 |
| chr11:93915137-93915637:downstream_9586 | chr11 | 93862093 | 93915137 | + | PANX1 | NM | 29 | -0,6 | 1 | 1 | 5,21E-01 | 8,42E-01 | 9 | 2 | 11 | 21 | 35 | 29 | 68 | 56 | -1,6 | -1,0 | -0,6 |
| chr14:35870215-35870715:downstream_11107 | chr14 | 35870715 | 35873960 | - | NFKBIA | NM | 170 | 0,3 | 1 | -1 | 5,21E-01 | 8,42E-01 | 48 | 49 | 24 | 35 | 318 | 355 | 278 | 251 | 0,7 | 0,3 | 0,4 |
| chr14:60610999-60611499:downstream_11164 | chr14 | 60611499 | 60632211 | - | DHRS7 | NM | 125 | 0,5 | 1 | -1 | 5,22E-01 | 8,42E-01 | 13 | 8 | 6 | 3 | 295 | 325 | 189 | 163 | 1,2 | 0,8 | 0,4 |
| chr12:48362661-48363161:downstream_18877 | chr12 | 48357329 | 48362661 | + | TMEM106C | NM | 46 | 0,7 | 1 | -1 | 5,22E-01 | 8,42E-01 | 7 | 4 | 3 | 2 | 88 | 109 | 64 | 89 | 1,0 | 0,4 | 0,6 |
| chr11:71212581-71213081:downstream_8913 | chr11 | 71164216 | 71212581 | + | NADSYN1 | NM | 10 | 1,0 | 2 | -1 | 5,23E-01 | NA | 3 | 8 | 0 | 5 | 28 | 5 | 18 | 16 | 1,2 | -0,1 | 1,2 |
| chr7:50511326-50511826:downstream_25520 | chr7 | 50511826 | 50518088 | - | FIGNL1 | NM | 33 | -0,7 | 1 | 1 | 5,24E-01 | 8,42E-01 | 0 | 7 | 4 | 15 | 51 | 32 | 90 | 68 | -1,5 | -0,9 | -0,6 |
| chr3:186507685-186508185:downstream_3312 | chr3 | 186501360 | 186507685 | + | EIF4A2 | NM | 77 | 0,5 | 1 | -1 | 5,24E-01 | 8,42E-01 | 28 | 15 | 14 | 33 | 107 | 99 | 175 | 144 | -0,1 | -0,6 | 0,5 |
| chr3:153992956-153993456:downstream_23114 | chr3 | 153993456 | 154042286 | - | DHX36 | NM | 40 | 0,5 | 1 | -1 | 5,24E-01 | 8,42E-01 | 13 | 7 | 18 | 22 | 31 | 36 | 92 | 100 | -1,0 | -1,5 | 0,5 |
| chr17:28513493-28513993:downstream_20615 | chr17 | 28443824 | 28513493 | + | NSRP1 | NM | 8 | 1,0 | 2 | -1 | 5,24E-01 | NA | 9 | 2 | 6 | 11 | 2 | 7 | 19 | 8 | -0,6 | -1,6 | 1,0 |
| chr17:8123447-8123947:downstream_12597 | chr17 | 8123947 | 8127361 | - | LINC00324 | NR | 167 | -1,0 | 2 | 1 | 5,24E-01 | 8,42E-01 | 10 | 3 | 0 | 0 | 697 | 607 | 7 | 11 | Inf | 6,1 | Inf |
| chr15:44160617-44161117:downstream_19846 | chr15 | 44119111 | 44160617 | + | WDR76 | NM | 6 | -1,1 | 2 | 1 | 5,24E-01 | NA | 3 | 6 | 4 | 10 | 2 | 15 | 4 | 7 | -0,6 | 0,5 | -1,1 |
| chr8:145750559-145751059:downstream_25997 | chr8 | 145743348 | 145750559 | + | LRRC14 | NM | 8 | 0,9 | 1 | -1 | 5,26E-01 | NA | 14 | 9 | 4 | 2 | 15 | 10 | 7 | 7 | 1,8 | 0,9 | 0,9 |
| chr2:86252950-86253450:downstream_2086 | chr2 | 86253450 | 86333278 | - | POLR1A | NM | 26 | -0,6 | 1 | 1 | 5,26E-01 | NA | 25 | 10 | 38 | 39 | 20 | 19 | 33 | 25 | -1,1 | -0,6 | -0,6 |
| chrX:57002302-57002802:downstream_7507 | chrX | 57002802 | 57021988 | - | SPIN3 | NR | 7 | 1,1 | 2 | -1 | 5,27E-01 | NA | 3 | 7 | 7 | 11 | 3 | 2 | 3 | 16 | -0,8 | -1,9 | 1,1 |
| chr7:148923339-148923839:downstream_25823 | chr7 | 148892553 | 148923339 | + | ZNF282 | NM | 12 | 0,8 | 1 | -1 | 5,27E-01 | NA | 13 | 5 | 7 | 8 | 14 | 11 | 17 | 19 | 0,2 | -0,5 | 0,8 |
| chr16:3459364-3459864:downstream_20282 | chr16 | 3451189 | 3459364 | + | ZNF174 | NM | 9 | -1,0 | 2 | 1 | 5,27E-01 | NA | 0 | 8 | 8 | 6 | 17 | 11 | 8 | 17 | -0,9 | 0,1 | -1,0 |
| chr3:87274838-87275338:downstream_3051 | chr3 | 87275338 | 87275427 | - | MIR4795 | NR | 10 | -1,1 | 2 | 1 | 5,27E-01 | NA | 13 | 29 | 3 | 3 | 11 | 17 | 0 | 2 | 2,8 | 4,1 | -1,3 |
| chr12:56703712-56704212:downstream_18519 | chr12 | 56704212 | 56710128 | - | CNPY2 | NM | 61 | -0,4 | 1 | 1 | 5,28E-01 | 8,45E-01 | 44 | 37 | 32 | 24 | 110 | 124 | 54 | 65 | 0,5 | 1,0 | -0,4 |
| chr14:71049437-71049937:downstream_19301 | chr14 | 71049937 | 71067407 | - | MED6 | NM | 9 | -0,9 | 1 | 1 | 5,28E-01 | NA | 8 | 14 | 7 | 10 | 19 | 5 | 7 | 3 | 0,4 | 1,2 | -0,8 |
| chr20:44471914-44472414:downstream_22568 | chr20 | 44462469 | 44471914 | + | SNX21 | NM | 9 | 1,0 | 2 | -1 | 5,28E-01 | NA | 7 | 15 | 14 | 11 | 7 | 0 | 10 | 7 | -0,3 | -1,3 | 1,1 |
| chr4:110745893-110746393:downstream_23665 | chr4 | 110736665 | 110745893 | + | GAR1 | NM | 9 | 1,0 | 2 | -1 | 5,28E-01 | NA | 7 | 17 | 3 | 2 | 17 | 13 | 4 | 6 | 2,5 | 1,6 | 0,9 |
| chr1:156563599-156564099:downstream_16781 | chr1 | 156564099 | 156571279 | - | GPATCH4 | NM | 35 | -0,5 | 1 | 1 | 5,28E-01 | 8,45E-01 | 36 | 41 | 29 | 15 | 68 | 46 | 25 | 21 | 0,8 | 1,3 | -0,5 |
| chr11:108018891-108019391:downstream_8824 | chr11 | 107992257 | 108018891 | + | ACAT1 | NM | 10 | 0,9 | 1 | -1 | 5,28E-01 | NA | 8 | 11 | 1 | 4 | 23 | 14 | 8 | 12 | 1,8 | 0,8 | 1,0 |
| chr20:3904502-3905002:downstream_22496 | chr20 | 3869485 | 3904502 | + | PANK2 | NM | 42 | 0,5 | 1 | -1 | 5,29E-01 | 8,45E-01 | 41 | 43 | 22 | 17 | 72 | 59 | 50 | 34 | 1,1 | 0,6 | 0,5 |
| chr13:77578888-77579388:downstream_10546 | chr13 | 77579388 | 77601331 | - | FBXL3 | NM | 11 | 0,8 | 1 | -1 | 5,29E-01 | NA | 15 | 17 | 8 | 9 | 7 | 15 | 7 | 12 | 0,9 | 0,2 | 0,7 |
| chr17:27582353-27582853:downstream_12943 | chr17 | 27582853 | 27621166 | - | NUFIP2 | NM | 41 | 0,5 | 1 | -1 | 5,29E-01 | 8,45E-01 | 11 | 9 | 14 | 6 | 49 | 74 | 92 | 74 | 0,0 | -0,4 | 0,5 |
| chr3:178952497-178952997:downstream_2757 | chr3 | 178866310 | 178952497 | + | PIK3CA | NM | 25 | -0,6 | 1 | 1 | 5,29E-01 | NA | 5 | 2 | 21 | 23 | 20 | 10 | 65 | 57 | -2,7 | -2,0 | -0,7 |
| chr2:131104197-131104697:downstream_2244 | chr2 | 131100488 | 131104197 | + | IMP4 | NM | 36 | -0,5 | 1 | 1 | 5,30E-01 | 8,45E-01 | 18 | 35 | 36 | 25 | 48 | 46 | 39 | 38 | -0,2 | 0,3 | -0,5 |
| chr13:73651680-73652180:downstream_19003 | chr13 | 73629113 | 73651680 | + | KLF5 | NM | 46 | -0,6 | 1 | 1 | 5,30E-01 | 8,45E-01 | 3 | 2 | 7 | 10 | 51 | 60 | 138 | 101 | -1,7 | -1,1 | -0,6 |
| chr3:12574820-12575320:downstream_23492 | chr3 | 12525930 | 12574820 | + | TSEN2 | NM | 12 | -1,1 | 2 | 1 | 5,30E-01 | NA | 2 | 0 | 6 | 5 | 3 | 18 | 28 | 34 | -2,7 | -1,5 | -1,2 |
| chr11:18552449-18552949:downstream_18382 | chr11 | 18552949 | 18610293 | - | UEVLD | NM | 67 | 0,4 | 1 | -1 | 5,30E-01 | 8,45E-01 | 59 | 54 | 71 | 78 | 56 | 44 | 107 | 67 | -0,4 | -0,8 | 0,4 |
| chr7:95212308-95212808:downstream_5952 | chr7 | 95212808 | 95225925 | - | PDK4 | NM | 98 | -0,7 | 1 | 1 | 5,31E-01 | 8,45E-01 | 2 | 0 | 14 | 24 | 43 | 29 | 358 | 317 | -4,0 | -3,2 | -0,7 |
| chr10:75535976-75536476:downstream_17580 | chr10 | 75532048 | 75535976 | + | FUT11 | NM | 28 | -0,6 | 1 | 1 | 5,31E-01 | NA | 6 | 1 | 11 | 11 | 39 | 25 | 64 | 66 | -1,7 | -1,0 | -0,7 |
| chr17:53039328-53039828:downstream_13247 | chr17 | 52978051 | 53039328 | + | TOM1L1 | NM | 26 | -0,6 | 1 | 1 | 5,31E-01 | NA | 26 | 17 | 43 | 25 | 15 | 33 | 25 | 25 | -0,6 | 0,0 | -0,6 |
| chr3:57307498-57307998:downstream_2633 | chr3 | 57261764 | 57307498 | + | APPL1 | NM | 7 | -1,3 | 2 | 1 | 5,32E-01 | NA | 1 | 0 | 6 | 8 | 5 | 1 | 19 | 19 | -4,1 | -2,7 | -1,4 |
| chr7:99131445-99131945:downstream_25818 | chr7 | 99102272 | 99131445 | + | ZKSCAN5 | NM | 14 | 0,7 | 1 | -1 | 5,32E-01 | NA | 19 | 13 | 4 | 7 | 24 | 16 | 14 | 11 | 1,4 | 0,7 | 0,8 |
| chr10:102754158-102754658:downstream_17499 | chr10 | 102747292 | 102754158 | + | C10orf2 | NM | 58 | 0,8 | 1 | -1 | 5,32E-01 | 8,45E-01 | 8 | 6 | 0 | 2 | 176 | 182 | 53 | 38 | 3,1 | 2,0 | 1,1 |
| chr3:28363343-28363843:downstream_23017 | chr3 | 28363843 | 28390618 | - | AZI2 | NM | 14 | -0,9 | 1 | 1 | 5,32E-01 | NA | 7 | 0 | 7 | 4 | 18 | 35 | 15 | 25 | -0,6 | 0,4 | -1,0 |
| chr16:11961484-11961984:downstream_20009 | chr16 | 11961984 | 12010519 | - | GSPT1 | NM | 55 | 0,6 | 1 | -1 | 5,32E-01 | 8,45E-01 | 7 | 6 | 7 | 6 | 53 | 112 | 116 | 130 | 0,0 | -0,6 | 0,5 |
| chr4:164047359-164047859:downstream_23766 | chr4 | 164047859 | 164088073 | - | NAF1 | NM | 7 | -1,1 | 2 | 1 | 5,33E-01 | NA | 9 | 4 | 8 | 8 | 16 | 0 | 6 | 4 | -0,4 | 0,8 | -1,1 |
| chr5:43045370-43045870:downstream_4237 | chr5 | 43042235 | 43045370 | + | LOC153684 | NR | 24 | 0,6 | 1 | -1 | 5,33E-01 | NA | 23 | 32 | 18 | 23 | 20 | 22 | 40 | 11 | 0,4 | -0,3 | 0,7 |
| chr5:137086572-137087072:downstream_4437 | chr5 | 137087072 | 137090039 | - | HNRNPA0 | NM | 33 | 0,5 | 1 | -1 | 5,33E-01 | 8,45E-01 | 40 | 24 | 35 | 42 | 19 | 28 | 33 | 45 | -0,3 | -0,7 | 0,5 |
| chr16:67263182-67263682:downstream_12411 | chr16 | 67261015 | 67263182 | + | TMEM208 | NM | 38 | 0,5 | 1 | -1 | 5,33E-01 | 8,45E-01 | 28 | 15 | 6 | 7 | 99 | 77 | 36 | 37 | 1,8 | 1,3 | 0,6 |
| chr7:47314251-47314751:downstream_5554 | chr7 | 47314751 | 47621742 | - | TNS3 | NM | 34 | -0,9 | 1 | 1 | 5,33E-01 | 8,45E-01 | 2 | 0 | 10 | 7 | 11 | 33 | 89 | 126 | -3,3 | -2,3 | -1,0 |
| chr17:58743640-58744140:downstream_12555 | chr17 | 58677543 | 58743640 | + | PPM1D | NM | 9 | 1,0 | 2 | -1 | 5,33E-01 | NA | 6 | 6 | 0 | 2 | 19 | 27 | 7 | 9 | 2,8 | 1,5 | 1,3 |
| chr19:49563896-49564396:downstream_14480 | chr19 | 49564396 | 49567124 | - | NTF4 | NM | 4 | 1,8 | 3 | -1 | 5,34E-01 | NA | 2 | 14 | 0 | 0 | 2 | 11 | 1 | 0 | Inf | 3,2 | Inf |
| chr1:53151513-53152013:downstream_735 | chr1 | 53152013 | 53164038 | - | COA7 | NM | 13 | 0,8 | 1 | -1 | 5,34E-01 | NA | 11 | 17 | 13 | 2 | 17 | 13 | 21 | 7 | 0,9 | 0,1 | 0,8 |
| chr5:140086239-140086739:downstream_4459 | chr5 | 140080031 | 140086239 | + | ZMAT2 | NM | 68 | -0,5 | 1 | 1 | 5,34E-01 | 8,45E-01 | 7 | 10 | 6 | 10 | 140 | 166 | 106 | 103 | 0,1 | 0,6 | -0,4 |
| chr7:21940016-21940516:downstream_25438 | chr7 | 21940516 | 21985542 | - | CDCA7L | NM | 10 | 0,9 | 1 | -1 | 5,35E-01 | NA | 1 | 12 | 7 | 11 | 5 | 9 | 21 | 18 | -0,6 | -1,5 | 0,9 |
| chr17:15932907-15933407:downstream_20606 | chr17 | 15933407 | 16118874 | - | NCOR1 | NM | 13 | 0,8 | 1 | -1 | 5,35E-01 | NA | 7 | 6 | 33 | 20 | 4 | 1 | 15 | 21 | -2,0 | -2,8 | 0,8 |
| chr15:81289257-81289757:downstream_11848 | chr15 | 81289757 | 81289814 | - | MIR4514 | NR | 8 | 1,3 | 2 | -1 | 5,36E-01 | NA | 2 | 9 | 0 | 0 | 25 | 25 | 4 | 1 | Inf | 3,3 | Inf |
| chr4:53527665-53528165:downstream_3675 | chr4 | 53525572 | 53527665 | + | USP46-AS1 | NR | 17 | -0,7 | 1 | 1 | 5,36E-01 | NA | 5 | 12 | 11 | 7 | 24 | 35 | 21 | 21 | -0,2 | 0,5 | -0,7 |
| chr3:113807268-113807768:downstream_23373 | chr3 | 113775581 | 113807268 | + | QTRTD1 | NM | 16 | 0,8 | 1 | -1 | 5,37E-01 | NA | 7 | 11 | 0 | 9 | 27 | 25 | 26 | 25 | 0,9 | 0,0 | 0,9 |
| chr3:120321258-120321758:downstream_23302 | chr3 | 120315127 | 120321258 | + | NDUFB4 | NM | 19 | 0,6 | 1 | -1 | 5,37E-01 | NA | 23 | 44 | 21 | 16 | 11 | 18 | 8 | 16 | 0,9 | 0,3 | 0,6 |
| chr11:60618561-60619061:downstream_9363 | chr11 | 60609428 | 60618561 | + | CCDC86 | NM | 30 | -0,8 | 1 | 1 | 5,37E-01 | 8,48E-01 | 7 | 7 | 0 | 2 | 76 | 122 | 7 | 16 | 2,5 | 3,1 | -0,6 |
| chr3:112720221-112720721:downstream_23185 | chr3 | 112709799 | 112720221 | + | GTPBP8 | NM | 38 | 0,5 | 1 | -1 | 5,37E-01 | 8,48E-01 | 24 | 32 | 35 | 33 | 33 | 35 | 60 | 52 | -0,3 | -0,7 | 0,5 |
| chr4:110969728-110970228:downstream_23626 | chr4 | 110970228 | 111119820 | - | ELOVL6 | NM | 5 | 1,5 | 2 | -1 | 5,38E-01 | NA | 2 | 0 | 11 | 7 | 1 | 0 | 11 | 7 | -2,9 | -4,4 | 1,5 |
| chr6:3020110-3020610:downstream_25039 | chr6 | 3000049 | 3020110 | + | NQO2 | NM | 10 | -0,9 | 1 | 1 | 5,38E-01 | NA | 2 | 2 | 4 | 6 | 16 | 15 | 17 | 22 | -1,2 | -0,3 | -0,8 |
| chr17:34946276-34946776:downstream_13003 | chr17 | 34900736 | 34946276 | + | GGNBP2 | NM | 58 | -0,6 | 1 | 1 | 5,38E-01 | 8,48E-01 | 4 | 0 | 6 | 9 | 57 | 79 | 174 | 134 | -1,8 | -1,2 | -0,7 |
| chr2:112972938-112973438:downstream_1701 | chr2 | 112973438 | 113012664 | - | ZC3H8 | NM | 30 | 0,6 | 1 | -1 | 5,38E-01 | 8,48E-01 | 8 | 5 | 6 | 5 | 66 | 32 | 61 | 58 | 0,3 | -0,3 | 0,6 |
| chr3:112322732-112323232:downstream_23052 | chr3 | 112323232 | 112359990 | - | CCDC80 | NM | 11 | -0,9 | 1 | 1 | 5,39E-01 | NA | 2 | 1 | 14 | 16 | 3 | 5 | 25 | 25 | -3,5 | -2,6 | -0,9 |
| chr1:56959918-56960418:downstream_139 | chr1 | 56960418 | 57045257 | - | PPAP2B | NM | 7 | 1,0 | 2 | -1 | 5,40E-01 | NA | 2 | 2 | 6 | 9 | 4 | 1 | 11 | 24 | -1,7 | -2,8 | 1,1 |
| chr4:129132298-129132798:downstream_23716 | chr4 | 128982420 | 129132298 | + | LARP1B | NM | 8 | -1,3 | 2 | 1 | 5,40E-01 | NA | 1 | 0 | 10 | 7 | 5 | 0 | 26 | 13 | -4,4 | -3,0 | -1,4 |
| chr18:669823-670323:downstream_20864 | chr18 | 670323 | 712662 | - | ENOSF1 | NM | 21 | 0,7 | 1 | -1 | 5,40E-01 | NA | 3 | 12 | 11 | 1 | 35 | 29 | 45 | 34 | 0,3 | -0,3 | 0,6 |
| chr5:133727799-133728299:downstream_4420 | chr5 | 133706866 | 133727799 | + | UBE2B | NM | 11 | 0,8 | 1 | -1 | 5,40E-01 | NA | 20 | 20 | 13 | 5 | 14 | 4 | 6 | 7 | 1,2 | 0,5 | 0,8 |

| | | | | | | | | | | | | | | | | | | | | | | | |
|---|---|---|---|---|---|---|---|---|---|---|---|---|---|---|---|---|---|---|---|---|---|---|---|
| chr12:120927640-120928140:downstream_10454 | chr12 | 120928140 | 120933749 | - | NRAV | NR | 57 | -0,4 | 1 | 1 | 5,40E-01 | 8,50E-01 | 62 | 42 | 13 | 20 | 135 | 126 | 29 | 32 | 1,7 | 2,1 | -0,4 |
| chr16:15503543-15504043:downstream_20124 | chr16 | 15489610 | 15503543 | + | MPV17L | NM | 10 | 0,9 | 1 | -1 | 5,41E-01 | NA | 11 | 11 | 15 | 1 | 11 | 7 | 14 | 10 | 0,4 | -0,4 | 0,8 |
| chr9:39072263-39072763:downstream_6862 | chr9 | 39072763 | 39288300 | - | CNTNAP3 | NM | 26 | 0,6 | 1 | -1 | 5,41E-01 | NA | 22 | 15 | 53 | 70 | 4 | 4 | 28 | 14 | -1,7 | -2,4 | 0,6 |
| chr11:64083463-64083963:downstream_18371 | chr11 | 64083963 | 64085556 | - | TRMT112 | NM | 32 | -0,6 | 1 | 1 | 5,41E-01 | 8,50E-01 | 3 | 10 | 3 | 6 | 62 | 101 | 25 | 48 | 0,6 | 1,2 | -0,6 |
| chr7:22896305-22896805:downstream_5732 | chr7 | 22896231 | 22896305 | + | SNORD93 | NR | 33 | 0,5 | 1 | -1 | 5,41E-01 | 8,50E-01 | 16 | 18 | 8 | 8 | 76 | 51 | 45 | 43 | 1,1 | 0,5 | 0,5 |
| chr15:45774179-45774679:downstream_11711 | chr15 | 45774679 | 45815002 | - | SLC30A4 | NM | 3 | 1,8 | 3 | -1 | 5,42E-01 | NA | 1 | 0 | 4 | 7 | 0 | 0 | 6 | 8 | -3,7 | -Inf | Inf |
| chr6:10974541-10975041:downstream_4866 | chr6 | 10887063 | 10974541 | + | SYCP2L | NM | 4 | -1,7 | 3 | 1 | 5,42E-01 | NA | 0 | 0 | 11 | 5 | 1 | 0 | 10 | 3 | -Inf | -4,0 | -Inf |
| chr12:109548798-109549298:downstream_18903 | chr12 | 109535398 | 109548798 | + | UNG | NM | 35 | -0,7 | 1 | 1 | 5,42E-01 | 8,50E-01 | 2 | 5 | 1 | 10 | 71 | 44 | 60 | 84 | -0,8 | -0,3 | -0,5 |
| chr15:34521696-34522196:downstream_19721 | chr15 | 34522196 | 34630265 | - | SLC12A6 | NM | 31 | 0,5 | 1 | -1 | 5,42E-01 | 8,50E-01 | 24 | 36 | 22 | 19 | 24 | 50 | 40 | 32 | 0,5 | 0,0 | 0,5 |
| chr2:27292841-27293341:downstream_1902 | chr2 | 27293341 | 27294567 | - | OST4 | NM | 133 | 0,3 | 1 | -1 | 5,43E-01 | 8,50E-01 | 154 | 122 | 53 | 46 | 237 | 238 | 97 | 114 | 1,5 | 1,2 | 0,3 |
| chr18:12661454-12661954:downstream_13574 | chr18 | 12661954 | 12702776 | - | CEP76 | NR | 3 | -1,8 | 3 | 1 | 5,43E-01 | NA | 6 | 5 | 0 | 1 | 2 | 12 | 0 | 0 | 3,7 | Inf | -Inf |
| chr1:41154251-41154751:downstream_119 | chr1 | 41154751 | 41157933 | - | NFYC-AS1 | NR | 15 | 0,7 | 1 | -1 | 5,43E-01 | NA | 32 | 16 | 10 | 8 | 19 | 16 | 14 | 7 | 1,4 | 0,7 | 0,7 |
| chr17:48358846-48359346:downstream_20779 | chr17 | 48348766 | 48358846 | + | TMEM92 | NM | 4 | 1,3 | 2 | -1 | 5,44E-01 | NA | 7 | 0 | 3 | 7 | 3 | 1 | 10 | 4 | -0,4 | -1,7 | 1,4 |
| chr13:45602404-45602904:downstream_10735 | chr13 | 45563686 | 45602404 | + | GPALPP1 | NM | 12 | -0,8 | 1 | 1 | 5,44E-01 | NA | 8 | 1 | 7 | 11 | 14 | 21 | 13 | 25 | -1,0 | -0,1 | -0,8 |
| chr12:100733914-100734414:downstream_9772 | chr12 | 100661548 | 100733914 | + | SCYL2 | NM | 42 | 0,5 | 1 | -1 | 5,44E-01 | 8,51E-01 | 26 | 29 | 117 | 89 | 10 | 2 | 33 | 31 | -1,9 | -2,5 | 0,6 |
| chr2:9628910-9629410:downstream_1848 | chr2 | 9629410 | 9695917 | - | ADAM17 | NM | 120 | 0,3 | 1 | -1 | 5,44E-01 | 8,51E-01 | 71 | 90 | 103 | 144 | 111 | 77 | 185 | 183 | -0,6 | -1,0 | 0,4 |
| chr9:114697654-114698154:downstream_6915 | chr9 | 114659045 | 114697654 | + | UGCG | NM | 32 | 0,7 | 1 | -1 | 5,45E-01 | 8,51E-01 | 7 | 0 | 3 | 8 | 41 | 30 | 88 | 83 | -0,6 | -1,3 | 0,7 |
| chr16:72118255-72118755:downstream_20263 | chr16 | 72118755 | 72128215 | - | TXNL4B | NM | 11 | 0,9 | 2 | -1 | 5,45E-01 | NA | 1 | 13 | 1 | 2 | 22 | 24 | 14 | 13 | 1,8 | 0,8 | 1,0 |
| chr20:30311212-30311712:downstream_14953 | chr20 | 30309309 | 30311212 | + | ABALON | NR | 5 | 1,2 | 2 | -1 | 5,45E-01 | NA | 12 | 2 | 0 | 3 | 11 | 4 | 4 | 5 | 2,1 | 0,7 | 1,4 |
| chr2:219138416-219138916:downstream_1556 | chr2 | 219138916 | 219157280 | - | TMBIM1 | NM | 26 | 0,6 | 1 | -1 | 5,45E-01 | NA | 9 | 6 | 6 | 2 | 57 | 54 | 31 | 45 | 1,0 | 0,6 | 0,5 |
| chr7:25157769-25158269:downstream_5742 | chr7 | 25158269 | 25164980 | - | CYCS | NM | 43 | 0,5 | 1 | -1 | 5,46E-01 | 8,51E-01 | 54 | 65 | 6 | 9 | 102 | 77 | 17 | 16 | 3,0 | 2,5 | 0,6 |
| chr18:43427073-43427573:downstream_13505 | chr18 | 43427573 | 43547305 | - | EPG5 | NM | 4 | 1,3 | 2 | -1 | 5,46E-01 | NA | 1 | 3 | 4 | 6 | 2 | 0 | 6 | 11 | -1,4 | -2,8 | 1,4 |
| chr1:90494094-90494594:downstream_17430 | chr1 | 90460677 | 90494094 | + | ZNF326 | NM | 6 | 1,0 | 2 | -1 | 5,47E-01 | NA | 6 | 8 | 4 | 2 | 7 | 7 | 8 | 3 | 1,2 | 0,2 | 1,0 |
| chr5:71505397-71505897:downstream_4031 | chr5 | 71403117 | 71505397 | + | MAP1B | NM | 43 | 0,7 | 1 | -1 | 5,47E-01 | 8,51E-01 | 3 | 0 | 6 | 10 | 20 | 20 | 146 | 143 | -2,2 | -2,9 | 0,6 |
| chr1:183114727-183115227:downstream_256 | chr1 | 182992594 | 183114727 | + | LAMC1 | NM | 43 | -0,6 | 1 | 1 | 5,47E-01 | 8,51E-01 | 4 | 0 | 8 | 8 | 37 | 57 | 96 | 133 | -2,0 | -1,3 | -0,7 |
| chr10:47088320-47088820:downstream_17673 | chr10 | 47083533 | 47088320 | + | NPY4R | NM | 9 | -0,9 | 1 | 1 | 5,47E-01 | NA | 1 | 7 | 13 | 11 | 8 | 7 | 15 | 11 | -1,7 | -0,8 | -0,8 |
| chr12:105388505-105389005:downstream_9915 | chr12 | 105380097 | 105388505 | + | C12orf45 | NM | 81 | 0,4 | 1 | -1 | 5,47E-01 | 8,51E-01 | 68 | 64 | 21 | 16 | 155 | 194 | 65 | 61 | 1,8 | 1,5 | 0,4 |
| chr5:38937522-38938022:downstream_24333 | chr5 | 38938022 | 39074510 | - | RICTOR | NM | 17 | 0,9 | 2 | -1 | 5,48E-01 | NA | 2 | 0 | 4 | 10 | 3 | 4 | 61 | 52 | -3,1 | -4,0 | 0,9 |
| chr13:21727233-21727733:downstream_19086 | chr13 | 21727733 | 21750741 | - | SKA3 | NM | 14 | -0,7 | 1 | 1 | 5,48E-01 | NA | 2 | 6 | 18 | 11 | 11 | 14 | 22 | 29 | -1,8 | -1,1 | -0,7 |
| chr1:111659453-111659953:downstream_192 | chr1 | 111659953 | 111682838 | - | DRAM2 | NM | 58 | -0,5 | 1 | 1 | 5,48E-01 | 8,52E-01 | 4 | 3 | 4 | 7 | 107 | 96 | 129 | 110 | -0,7 | -0,2 | -0,5 |
| chr12:7084846-7085346:downstream_10005 | chr12 | 7085346 | 7125842 | - | LPCAT3 | NM | 25 | -0,7 | 1 | 1 | 5,49E-01 | NA | 5 | 2 | 1 | 7 | 50 | 48 | 38 | 48 | -0,4 | 0,2 | -0,6 |
| chr12:10375724-10376224:downstream_10046 | chr12 | 10365488 | 10375724 | + | GABARAPL1 | NM | 23 | 0,7 | 1 | -1 | 5,49E-01 | NA | 11 | 3 | 4 | 7 | 27 | 45 | 35 | 51 | 0,4 | -0,2 | 0,7 |
| chr15:50793280-50793780:downstream_19835 | chr15 | 50716573 | 50793280 | + | USP8 | NM | 29 | -0,6 | 1 | 1 | 5,49E-01 | 8,53E-01 | 3 | 1 | 14 | 8 | 29 | 20 | 83 | 73 | -2,4 | -1,7 | -0,7 |
| chr17:45527092-45527592:downstream_13187 | chr17 | 45527592 | 45569986 | - | MRPL45P2 | NR | 6 | 1,0 | 2 | -1 | 5,51E-01 | NA | 10 | 9 | 6 | 3 | 4 | 5 | 4 | 4 | 1,1 | 0,1 | 1,0 |
| chr14:65000408-65000908:downstream_19441 | chr14 | 64971291 | 65000408 | + | ZBTB1 | NM | 7 | -1,0 | 2 | 1 | 5,51E-01 | NA | 0 | 5 | 3 | 7 | 7 | 15 | 13 | 10 | -1,0 | 0,0 | -0,9 |
| chr2:219528086-219528586:downstream_2426 | chr2 | 219528586 | 219536781 | - | RNF25 | NM | 13 | 0,7 | 1 | -1 | 5,51E-01 | NA | 10 | 15 | 4 | 2 | 26 | 23 | 11 | 11 | 1,9 | 1,2 | 0,7 |
| chr7:90020769-90021269:downstream_25549 | chr7 | 89975978 | 90020769 | + | GTPBP10 | NM | 12 | 0,7 | 1 | -1 | 5,51E-01 | NA | 7 | 15 | 11 | 3 | 18 | 13 | 19 | 13 | 0,6 | -0,1 | 0,7 |
| chr5:112228776-112229276:downstream_24395 | chr5 | 112196884 | 112228776 | + | SRP19 | NM | 27 | 0,6 | 1 | -1 | 5,51E-01 | NA | 7 | 15 | 4 | 2 | 78 | 54 | 28 | 30 | 1,8 | 1,2 | 0,6 |
| chr5:177575571-177576071:downstream_24335 | chr5 | 177557961 | 177575571 | + | RMND5B | NM | 33 | 0,5 | 1 | -1 | 5,51E-01 | 8,54E-01 | 14 | 22 | 18 | 16 | 37 | 45 | 53 | 55 | 0,1 | -0,4 | 0,5 |
| chr7:29606911-29607411:downstream_5767 | chr7 | 29603426 | 29606911 | + | PRR15 | NM | 22 | -0,8 | 1 | 1 | 5,51E-01 | NA | 1 | 2 | 10 | 10 | 20 | 9 | 75 | 52 | -2,8 | -2,1 | -0,7 |
| chr19:11033448-11033948:downstream_13727 | chr19 | 10982252 | 11033448 | + | CARM1 | NM | 11 | 0,9 | 1 | -1 | 5,51E-01 | NA | 2 | 19 | 8 | 5 | 8 | 16 | 8 | 20 | 0,7 | -0,2 | 0,9 |
| chr14:23347291-23347791:downstream_11057 | chr14 | 23340959 | 23347291 | + | LRP10 | NM | 166 | 0,3 | 0 | -1 | 5,52E-01 | 8,54E-01 | 157 | 170 | 116 | 101 | 212 | 224 | 167 | 185 | 0,6 | 0,3 | 0,3 |
| chr10:50662025-50662525:downstream_8281 | chr10 | 50662525 | 50747169 | - | ERCC6 | NM | 8 | -0,9 | 1 | 1 | 5,52E-01 | NA | 7 | 7 | 19 | 10 | 5 | 4 | 6 | 5 | -1,1 | -0,3 | -0,9 |
| chr9:104325626-104326126:downstream_7234 | chr9 | 104296130 | 104325626 | + | RNF20 | NM | 28 | -0,6 | 1 | 1 | 5,52E-01 | NA | 3 | 13 | 22 | 14 | 42 | 24 | 47 | 55 | -1,2 | -0,6 | -0,6 |
| chr5:142656995-142657495:downstream_24257 | chr5 | 142657495 | 142815077 | - | NR3C1 | NM | 17 | 0,8 | 1 | -1 | 5,52E-01 | NA | 7 | 0 | 10 | 13 | 5 | 11 | 43 | 45 | -1,8 | -2,5 | 0,7 |
| chrX:118721799-118722299:downstream_26941 | chrX | 118722299 | 118739846 | - | NKRF | NM | 18 | 0,7 | 1 | -1 | 5,52E-01 | NA | 6 | 4 | 4 | 2 | 39 | 29 | 22 | 40 | 0,7 | 0,1 | 0,6 |
| chr12:48055214-48055714:downstream_18806 | chr12 | 48055714 | 48099844 | - | RPAP3 | NM | 26 | 0,6 | 1 | -1 | 5,53E-01 | NA | 39 | 16 | 43 | 42 | 17 | 4 | 21 | 28 | -0,6 | -1,2 | 0,6 |
| chr14:73588076-73588576:downstream_10961 | chr14 | 73525220 | 73588076 | + | RBM25 | NM | 37 | -0,6 | 1 | 1 | 5,53E-01 | 8,54E-01 | 15 | 1 | 15 | 23 | 56 | 36 | 56 | 91 | -1,3 | -0,7 | -0,6 |
| chr1:173768187-173768687:downstream_16586 | chr1 | 173768687 | 173793777 | - | CENPL | NM | 13 | 0,7 | 1 | -1 | 5,53E-01 | NA | 8 | 10 | 21 | 19 | 7 | 2 | 17 | 18 | -1,2 | -1,9 | 0,8 |
| chr4:155455648-155456148:downstream_23800 | chr4 | 155456148 | 155471585 | - | PLRG1 | NM | 32 | 0,6 | 1 | -1 | 5,53E-01 | 8,54E-01 | 15 | 4 | 10 | 2 | 62 | 56 | 53 | 53 | 0,6 | 0,2 | 0,5 |
| chrX:65814981-65815481:downstream_26795 | chrX | 65815481 | 65859140 | - | EDA2R | NM | 30 | -0,5 | 1 | 1 | 5,53E-01 | 8,54E-01 | 40 | 30 | 26 | 27 | 32 | 42 | 19 | 21 | 0,4 | 0,9 | -0,5 |
| chr17:5289659-5290159:downstream_20665 | chr17 | 5185557 | 5289659 | + | RABEP1 | NM | 5 | 1,3 | 2 | -1 | 5,54E-01 | NA | 0 | 6 | 7 | 7 | 2 | 0 | 4 | 11 | -1,2 | -2,6 | 1,4 |
| chr12:102122846-102123346:downstream_9908 | chr12 | 102091416 | 102122846 | + | CHPT1 | NM | 18 | -0,6 | 1 | 1 | 5,54E-01 | NA | 5 | 14 | 13 | 20 | 15 | 29 | 25 | 26 | -0,8 | -0,2 | -0,6 |
| chr6:22194616-22195116:downstream_4654 | chr6 | 21666674 | 22194616 | + | CASC15 | NR | 11 | 0,8 | 1 | -1 | 5,54E-01 | NA | 4 | 4 | 13 | 7 | 9 | 3 | 24 | 27 | -1,3 | -2,1 | 0,8 |
| chrX:100096509-100097009:downstream_26679 | chrX | 100075347 | 100096509 | + | CSTF2 | NM | 10 | -0,9 | 2 | 1 | 5,55E-01 | NA | 3 | 8 | 3 | 13 | 21 | 6 | 15 | 7 | -0,5 | 0,2 | -0,8 |
| chr16:69358946-69359446:downstream_12436 | chr16 | 69345286 | 69358946 | + | VPS4A | NM | 14 | 0,8 | 1 | -1 | 5,55E-01 | NA | 11 | 5 | 3 | 2 | 34 | 30 | 13 | 16 | 1,9 | 1,2 | 0,7 |
| chr5:110100861-110101361:downstream_24365 | chr5 | 110073836 | 110100861 | + | SLC25A46 | NM | 11 | 0,8 | 1 | -1 | 5,55E-01 | NA | 2 | 8 | 3 | 2 | 12 | 27 | 21 | 16 | 1,0 | 0,1 | 0,8 |
| chr14:93668736-93669236:downstream_11277 | chr14 | 93669236 | 93673459 | - | C14orf142 | NM | 14 | 0,8 | 1 | -1 | 5,55E-01 | NA | 2 | 9 | 4 | 2 | 35 | 12 | 26 | 25 | 0,6 | -0,2 | 0,8 |
| chr1:6280752-6281252:downstream_8 | chr1 | 6281252 | 6296044 | - | ICMT | NM | 40 | 0,6 | 1 | -1 | 5,56E-01 | 8,56E-01 | 2 | 13 | 6 | 5 | 50 | 94 | 68 | 85 | 0,5 | -0,1 | 0,6 |
| chr19:52449011-52449511:downstream_21690 | chr19 | 52430687 | 52449011 | + | ZNF613 | NM | 8 | -1,0 | 2 | 1 | 5,56E-01 | NA | 3 | 0 | 6 | 5 | 6 | 13 | 13 | 16 | -1,7 | -0,6 | -1,1 |
| chr19:12840953-12841453:downstream_13733 | chr19 | 12841453 | 12845529 | - | C19orf43 | NM | 68 | 0,5 | 1 | -1 | 5,56E-01 | 8,56E-01 | 13 | 9 | 6 | 2 | 189 | 156 | 65 | 101 | 1,4 | 1,1 | 0,4 |
| chr10:1090141-1090641:downstream_17605 | chr10 | 1068576 | 1090141 | + | IDI2-AS1 | NR | 8 | 1,1 | 2 | -1 | 5,56E-01 | NA | 11 | 21 | 1 | 1 | 17 | 8 | 4 | 0 | 3,9 | 2,6 | 1,3 |
| chr9:125669834-125670334:downstream_7307 | chr9 | 125670334 | 125675607 | - | ZBTB6 | NM | 29 | 0,5 | 1 | -1 | 5,57E-01 | 8,56E-01 | 16 | 18 | 6 | 6 | 66 | 63 | 33 | 28 | 1,6 | 1,1 | 0,5 |
| chr5:111498198-111498698:downstream_4363 | chr5 | 111496222 | 111498198 | + | EPB41L4A-AS1 | NR | 110 | 0,3 | 1 | -1 | 5,57E-01 | 8,56E-01 | 130 | 110 | 40 | 40 | 203 | 189 | 90 | 75 | 1,6 | 1,2 | 0,3 |
| chr16:48393945-48394445:downstream_20208 | chr16 | 48394445 | 48419229 | - | SIAH1 | NM | 15 | 0,7 | 1 | -1 | 5,57E-01 | NA | 6 | 15 | 7 | 5 | 26 | 20 | 26 | 16 | 0,8 | 0,1 | 0,7 |
| chr9:131483199-131483699:downstream_6940 | chr9 | 131464801 | 131483199 | + | PKN3 | NM | 10 | 0,8 | 1 | -1 | 5,57E-01 | NA | 7 | 15 | 3 | 7 | 13 | 12 | 8 | 11 | 1,2 | 0,3 | 0,9 |
| chr1:233114219-233114719:downstream_1427 | chr1 | 233086369 | 233114219 | + | NTPCR | NM | 48 | 0,4 | 1 | -1 | 5,58E-01 | 8,57E-01 | 36 | 76 | 33 | 38 | 59 | 47 | 40 | 51 | 0,7 | 0,2 | 0,4 |
| chr3:128886157-128886657:downstream_23089 | chr3 | 128886657 | 128902810 | - | CNBP | NM | 16 | -0,7 | 1 | 1 | 5,59E-01 | NA | 21 | 17 | 8 | 7 | 26 | 35 | 14 | 2 | 1,3 | 1,9 | -0,6 |
| chr7:100956147-100956647:downstream_25567 | chr7 | 100956647 | 100965104 | - | IFT22 | NM | 6 | 1,1 | 2 | -1 | 5,59E-01 | NA | 4 | 5 | 4 | 0 | 2 | 15 | 7 | 9 | 1,1 | 0,2 | 0,9 |
| chr1:117116519-117117019:downstream_16817 | chr1 | 117117019 | 117210377 | - | IGSF3 | NM | 29 | -0,7 | 1 | 1 | 5,61E-01 | 8,60E-01 | 3 | 0 | 4 | 11 | 27 | 30 | 81 | 80 | -2,2 | -1,5 | -0,7 |
| chr7:100485843-100486343:downstream_6006 | chr7 | 100486343 | 100487339 | - | UFSP1 | NM | 34 | 0,5 | 1 | -1 | 5,61E-01 | 8,60E-01 | 45 | 75 | 17 | 20 | 44 | 38 | 29 | 8 | 1,7 | 1,1 | 0,6 |
| chr10:105206019-105206519:downstream_8706 | chr10 | 105156411 | 105206019 | + | PDCD11 | NM | 20 | 0,6 | 1 | -1 | 5,61E-01 | NA | 20 | 10 | 11 | 4 | 33 | 35 | 18 | 32 | 0,9 | 0,4 | 0,5 |
| chr6:43004854-43005354:downstream_24648 | chr6 | 43005354 | 43021683 | - | CUL7 | NM | 8 | 1,0 | 2 | -1 | 5,61E-01 | NA | 7 | 5 | 6 | 0 | 19 | 6 | 10 | 10 | 1,1 | 0,3 | 0,8 |
| chr5:68737890-68738390:downstream_24217 | chr5 | 68710938 | 68737890 | + | MARVELD2 | NM | 16 | -0,6 | 1 | 1 | 5,62E-01 | NA | 8 | 10 | 15 | 23 | 14 | 15 | 21 | 20 | -1,1 | -0,5 | -0,6 |
| chr4:71556268-71556768:downstream_3724 | chr4 | 71554195 | 71556268 | + | UTP3 | NM | 62 | -0,5 | 1 | 1 | 5,63E-01 | 8,62E-01 | 24 | 6 | 7 | 5 | 197 | 163 | 57 | 39 | 1,3 | 1,9 | -0,6 |
| chr2:70508005-70508505:downstream_1646 | chr2 | 70508505 | 70520869 | - | SNRPG | NM | 24 | -0,7 | 1 | 1 | 5,63E-01 | NA | 54 | 55 | 29 | 41 | 7 | 3 | 3 | 2 | 0,6 | 1,2 | -0,6 |
| chr1:236711804-236712304:downstream_314 | chr1 | 236712304 | 236767841 | - | HEATR1 | NM | 45 | -0,4 | 1 | 1 | 5,64E-01 | 8,62E-01 | 24 | 12 | 56 | 36 | 36 | 46 | 72 | 79 | -1,3 | -0,9 | -0,5 |
| chr15:43036041-43036541:downstream_11460 | chr15 | 43036541 | 43213007 | - | TTBK2 | NM | 5 | -1,3 | 2 | 1 | 5,64E-01 | NA | 0 | 1 | 7 | 6 | 2 | 2 | 8 | 11 | -3,7 | -2,5 | -1,2 |
| chr15:25304781-25305281:downstream_11583 | chr15 | 25304683 | 25304781 | + | SNORD116-4 | NR | 4 | -1,5 | 3 | 1 | 5,64E-01 | NA | 0 | 0 | 3 | 7 | 2 | 0 | 7 | 11 | -Inf | -3,5 | -Inf |
| chr3:167401194-167401694:downstream_23331 | chr3 | 167401694 | 167452651 | - | PDCD10 | NM | 14 | -0,7 | 1 | 1 | 5,64E-01 | NA | 4 | 2 | 17 | 10 | 7 | 16 | 35 | 25 | -2,1 | -1,4 | -0,8 |
| chr1:8920558-8921058:downstream_16693 | chr1 | 8921058 | 8939151 | - | ENO1 | NM | 477 | 0,3 | 1 | -1 | 5,64E-01 | 8,62E-01 | 29 | 42 | 22 | 25 | 1049 | 974 | 715 | 958 | 0,6 | 0,3 | 0,3 |
| chr12:88443937-88444437:downstream_10343 | chr12 | 88429267 | 88443937 | + | C12orf29 | NM | 9 | 1,0 | 2 | -1 | 5,64E-01 | NA | 6 | 3 | 3 | 0 | 19 | 24 | 8 | 13 | 1,6 | 1,0 | 0,6 |
| chr2:72405943-72406443:downstream_1650 | chr2 | 72406443 | 73053177 | - | EXOC6B | NM | 42 | -0,5 | 1 | 1 | 5,65E-01 | 8,62E-01 | 30 | 32 | 136 | 80 | 6 | 10 | 28 | 11 | -1,8 | -1,3 | -0,5 |
| chr14:32628934-32629434:downstream_19150 | chr14 | 32546494 | 32628934 | + | ARHGAP5 | NM | 10 | 0,9 | 2 | -1 | 5,66E-01 | NA | 8 | 3 | 13 | 11 | 8 | 0 | 22 | 11 | -1,1 | -2,1 | 0,9 |
| chr20:31027122-31027622:downstream_22356 | chr20 | 30946146 | 31027122 | + | ASXL1 | NM | 11 | 0,8 | 1 | -1 | 5,66E-01 | NA | 1 | 8 | 18 | 6 | 5 | 5 | 17 | 31 | -1,5 | -2,3 | 0,8 |
| chr19:52829180-52829680:downstream_21653 | chr19 | 52800421 | 52829180 | + | ZNF480 | NM | 17 | 0,6 | 1 | -1 | 5,66E-01 | NA | 4 | 10 | 7 | 4 | 24 | 27 | 35 | 29 | 0,3 | -0,3 | 0,6 |
| chr3:170582164-170582664:downstream_3273 | chr3 | 170582664 | 170588045 | - | RPL22L1 | NM | 13 | 0,7 | 1 | -1 | 5,67E-01 | NA | 12 | 27 | 4 | 7 | 20 | 17 | 11 | 7 | 1,8 | 1,0 | 0,8 |
| chr8:101268786-101269286:downstream_26100 | chr8 | 101269286 | 101348446 | - | RNF19A | NM | 49 | 1,1 | 2 | -1 | 5,67E-01 | 8,64E-01 | 14 | 0 | 8 | 23 | 36 | 23 | 132 | 154 | -1,2 | -2,3 | 1,1 |
| chr1:219386207-219386707:downstream_16927 | chr1 | 219347172 | 219386207 | + | LYPLAL1 | NM | 7 | 0,8 | 1 | -1 | 5,67E-01 | NA | 5 | 9 | 8 | 5 | 8 | 4 | 11 | 10 | 0,0 | -0,8 | 0,8 |
| chr9:119463579-119464079:downstream_26553 | chr9 | 119449580 | 119463579 | + | TRIM32 | NM | 68 | -0,4 | 1 | 1 | 5,67E-01 | 8,64E-01 | 110 | 101 | 39 | 28 | 117 | 96 | 29 | 23 | 1,7 | 2,0 | -0,4 |
| chr9:80881983-80882483:downstream_26261 | chr9 | 80850990 | 80881983 | + | CEP78 | NM | 9 | 1,9 | 3 | -1 | 5,67E-01 | NA | 11 | 0 | 1 | 5 | 16 | 0 | 18 | 17 | 0,9 | -1,1 | 2,0 |
| chr10:24872037-24872537:downstream_8256 | chr10 | 24872537 | 25012597 | - | ARHGAP21 | NM | 13 | -1,1 | 2 | 1 | 5,67E-01 | NA | 0 | 1 | 7 | 6 | 7 | 5 | 43 | 34 | -3,7 | -2,8 | -0,9 |
| chr4:169849608-169850108:downstream_23787 | chr4 | 169418216 | 169849608 | + | PALLD | NM | 17 | 0,7 | 1 | -1 | 5,68E-01 | NA | 7 | 1 | 11 | 2 | 21 | 14 | 35 | 46 | -0,6 | -1,2 | 0,6 |
| chr8:23088439-23088939:downstream_6492 | chr8 | 23082733 | 23088439 | + | LOC389641 | NR | 7 | -1,1 | 2 | 1 | 5,68E-01 | NA | 21 | 0 | 6 | 3 | 10 | 12 | 1 | 2 | 1,3 | 2,5 | -1,2 |
| chr3:98297789-98298289:downstream_2654 | chr3 | 98298289 | 98312455 | - | CPOX | NM | 37 | -0,6 | 1 | 1 | 5,68E-01 | 8,64E-01 | 5 | 0 | 3 | 11 | 53 | 46 | 110 | 71 | -1,5 | -0,9 | -0,6 |
| chr15:49622002-49622502:downstream_19559 | chr15 | 49447955 | 49622002 | + | GALK2 | NM | 10 | 0,8 | 1 | -1 | 5,68E-01 | NA | 4 | 6 | 6 | 20 | 7 | 1 | 14 | 22 | -1,4 | -2,3 | 0,9 |
| chr21:33041243-33041743:downstream_15283 | chr21 | 33031934 | 33041243 | + | SOD1 | NM | 184 | -1,1 | 2 | 1 | 5,68E-01 | 8,64E-01 | 14 | 0 | 15 | 5 | 363 | 505 | 220 | 348 | -0,5 | 0,6 | -1,2 |
| chr1:36389899-36390399:downstream_636 | chr1 | 36348809 | 36389899 | + | AGO1 | NM | 10 | 0,8 | 1 | -1 | 5,69E-01 | NA | 4 | 5 | 17 | 17 | 5 | 0 | 14 | 21 | -1,9 | -2,8 | 0,9 |
| chr7:135361160-135361660:downstream_6102 | chr7 | 135347220 | 135361160 | + | C7orf73 | NM | 10 | 0,8 | 1 | -1 | 5,69E-01 | NA | 13 | 8 | 10 | 3 | 9 | 15 | 14 | 11 | 0,7 | 0,0 | 0,7 |
| chr9:125679807-125680307:downstream_26575 | chr9 | 125680307 | 125693830 | - | ZBTB26 | NM | 5 | 1,0 | 2 | -1 | 5,69E-01 | NA | 6 | 5 | 3 | 2 | 7 | 6 | 3 | 10 | 1,0 | 0,1 | 1,0 |
| chr12:101786397-101786897:downstream_18439 | chr12 | 101786897 | 101801598 | - | ARL1 | NM | 46 | 0,5 | 1 | -1 | 5,69E-01 | 8,64E-01 | 5 | 17 | 6 | 13 | 67 | 75 | 77 | 104 | 0,3 | -0,3 | 0,6 |
| chr19:34720420-34720920:downstream_21259 | chr19 | 34663351 | 34720420 | + | LSM14A | NM | 14 | 0,6 | 1 | -1 | 5,69E-01 | NA | 7 | 5 | 10 | 8 | 15 | 11 | 26 | 31 | -0,6 | -1,2 | 0,6 |
| chr17:46839092-46839592:downstream_20796 | chr17 | 46839592 | 46894469 | - | TTLL6 | NM | 11 | -0,9 | 2 | 1 | 5,69E-01 | NA | 0 | 2 | 6 | 7 | 10 | 7 | 25 | 34 | -2,6 | -1,8 | -0,8 |
| chr3:98233816-98234316:downstream_23084 | chr3 | 98234316 | 98241910 | - | CLDND1 | NM | 52 | 0,5 | 1 | -1 | 5,69E-01 | 8,64E-01 | 7 | 3 | 4 | 5 | 76 | 100 | 114 | 109 | 0,2 | -0,3 | 0,5 |
| chr3:43764217-43764717:downstream_2902 | chr3 | 43732374 | 43764217 | + | ABHD5 | NM | 15 | 0,7 | 1 | -1 | 5,70E-01 | NA | 21 | 35 | 7 | 2 | 24 | 22 | 7 | 5 | 2,6 | 1,9 | 0,6 |
| chr16:1890203-1890703:downstream_19984 | chr16 | 1877224 | 1890203 | + | FAHD1 | NM | 11 | 1,0 | 2 | -1 | 5,71E-01 | NA | 15 | 46 | 0 | 4 | 18 | 4 | 3 | 1 | 3,9 | 2,6 | 1,3 |
| chr3:185631857-185632357:downstream_23486 | chr3 | 185632357 | 185655924 | - | TRA2B | NM | 22 | 0,6 | 1 | -1 | 5,71E-01 | NA | 28 | 21 | 6 | 9 | 46 | 30 | 18 | 16 | 1,8 | 1,2 | 0,6 |
| chr8:33227843-33228343:downstream_6285 | chr8 | 33228343 | 33330664 | - | FUT10 | NM | 12 | 0,8 | 1 | -1 | 5,71E-01 | NA | 4 | 0 | 8 | 7 | 3 | 7 | 43 | 22 | -2,0 | -2,7 | 0,8 |
| chr12:21621343-21621843:downstream_18794 | chr12 | 21621843 | 21654603 | - | RECQL | NM | 70 | 0,5 | 1 | -1 | 5,71E-01 | 8,65E-01 | 7 | 0 | 4 | 6 | 106 | 81 | 189 | 167 | -0,4 | -0,9 | 0,5 |
| chr16:70367735-70368235:downstream_19963 | chr16 | 70323669 | 70367735 | + | DDX19B | NM | 24 | -0,6 | 1 | 1 | 5,72E-01 | NA | 7 | 2 | 13 | 12 | 32 | 26 | 49 | 53 | -1,4 | -0,8 | -0,6 |
| chr1:25568239-25568739:downstream_561 | chr1 | 25568739 | 25573985 | - | RSRP1 | NM | 56 | 0,5 | 1 | -1 | 5,72E-01 | 8,65E-01 | 12 | 16 | 8 | 2 | 132 | 152 | 49 | 74 | 1,5 | 1,2 | 0,3 |
| chr12:16056410-16056910:downstream_10090 | chr12 | 16035287 | 16056410 | + | STRAP | NM | 83 | 0,4 | 1 | -1 | 5,72E-01 | 8,65E-01 | 14 | 23 | 10 | 11 | 146 | 196 | 110 | 153 | 0,8 | 0,4 | 0,4 |
| chr9:34613041-34613541:downstream_26277 | chr9 | 34613541 | 34620520 | - | DCTN3 | NM | 37 | 0,6 | 1 | -1 | 5,72E-01 | 8,65E-01 | 7 | 2 | 4 | 2 | 59 | 86 | 74 | 63 | 0,5 | 0,1 | 0,5 |
| chr3:50361839-50362339:downstream_2978 | chr3 | 50362339 | 50365669 | - | TUSC2 | NM | 25 | 0,7 | 1 | -1 | 5,73E-01 | NA | 9 | 4 | 0 | 2 | 75 | 59 | 22 | 28 | 2,4 | 1,4 | 1,0 |
| chr2:8996200-8996700:downstream_1570 | chr2 | 8996700 | 9143876 | - | MBOAT2 | NM | 13 | -0,7 | 1 | 1 | 5,73E-01 | NA | 5 | 9 | 8 | 11 | 11 | 27 | 19 | 14 | -0,5 | 0,2 | -0,7 |
| chr17:55015060-55015560:downstream_13251 | chr17 | 55015560 | 55038411 | - | COIL | NM | 23 | 0,5 | 1 | -1 | 5,73E-01 | NA | 10 | 23 | 8 | 14 | 37 | 27 | 28 | 37 | 0,6 | 0,0 | 0,6 |
| chr2:37326387-37326887:downstream_21944 | chr2 | 37311593 | 37326387 | + | GPATCH11 | NM | 9 | -0,8 | 1 | 1 | 5,73E-01 | NA | 2 | 4 | 11 | 19 | 3 | 6 | 14 | 15 | -2,4 | -1,7 | -0,8 |
| chr2:44547962-44548462:downstream_1957 | chr2 | 44502596 | 44547962 | + | SLC3A1 | NM | 14 | 0,7 | 1 | -1 | 5,73E-01 | NA | 6 | 12 | 10 | 19 | 8 | 10 | 15 | 32 | -0,7 | -1,4 | 0,7 |
| chr10:64914791-64915291:downstream_17675 | chr10 | 64893006 | 64914791 | + | NRBF2 | NM | 16 | 0,6 | 1 | -1 | 5,73E-01 | NA | 20 | 21 | 7 | 10 | 25 | 17 | 22 | 7 | 1,3 | 0,6 | 0,7 |
| chr1:153633763-153634263:downstream_16826 | chr1 | 153634263 | 153643504 | - | ILF2 | NM | 64 | 0,4 | 1 | -1 | 5,74E-01 | 8,65E-01 | 13 | 24 | 10 | 7 | 136 | 157 | 71 | 98 | 1,2 | 0,8 | 0,4 |
| chr15:75212116-75212616:downstream_11810 | chr15 | 75212616 | 75230495 | - | COX5A | NM | 19 | 0,6 | 1 | -1 | 5,74E-01 | NA | 22 | 19 | 11 | 13 | 16 | 31 | 25 | 16 | 0,8 | 0,2 | 0,6 |
| chr2:27272050-27272550:downstream_1905 | chr2 | 27272550 | 27273132 | - | AGBL5-AS1 | NR | 4 | -1,4 | 3 | 1 | 5,74E-01 | NA | 9 | 15 | 0 | 2 | 5 | 4 | 0 | 0 | 3,3 | Inf | -Inf |
| chr17:16286059-16286559:downstream_20799 | chr17 | 16284106 | 16286059 | + | UBB | NM | 297 | 0,3 | 1 | -1 | 5,74E-01 | 8,65E-01 | 37 | 15 | 24 | 28 | 572 | 443 | 597 | 663 | 0,0 | -0,3 | 0,3 |
| chr8:141540763-141541263:downstream_25844 | chr8 | 141541263 | 141645646 | - | AGO2 | NM | 27 | -0,5 | 1 | 1 | 5,74E-01 | NA | 7 | 6 | 36 | 43 | 15 | 7 | 56 | 44 | -2,7 | -2,2 | -0,5 |
| chr1:229576543-229577043:downstream_55 | chr1 | 229577043 | 229644088 | - | NUP133 | NM | 28 | 0,6 | 1 | -1 | 5,74E-01 | 8,65E-01 | 3 | 4 | 19 | 15 | 11 | 12 | 90 | 71 | -2,3 | -2,8 | 0,5 |
| chr12:124105482-124105982:downstream_10494 | chr12 | 124086671 | 124105482 | + | DDX55 | NM | 27 | 0,5 | 1 | -1 | 5,75E-01 | NA | 26 | 22 | 17 | 14 | 47 | 26 | 32 | 33 | 0,7 | 0,2 | 0,5 |
| chr14:21168761-21169261:downstream_19366 | chr14 | 21152371 | 21168761 | + | RNASE4 | NM | 281 | 0,3 | 1 | -1 | 5,75E-01 | 8,65E-01 | 16 | 24 | 13 | 14 | 529 | 660 | 518 | 478 | 0,6 | 0,3 | 0,4 |
| chr16:2320773-2321273:downstream_12116 | chr16 | 2320713 | 2320773 | + | MIR3677 | NR | 5 | -1,1 | 2 | 1 | 5,76E-01 | NA | 8 | 7 | 1 | 2 | 8 | 15 | 1 | 1 | 2,3 | 3,4 | -1,1 |
| chr17:5335598-5336098:downstream_12776 | chr17 | 5336098 | 5342471 | - | C1QBP | NM | 523 | 0,3 | 1 | -1 | 5,76E-01 | 8,66E-01 | 32 | 61 | 38 | 49 | 738 | 1099 | 1087 | 1080 | 0,1 | -0,2 | 0,3 |
| chr1:53679271-53679771:downstream_16527 | chr1 | 53679771 | 53686311 | - | C1orf123 | NM | 25 | 0,6 | 1 | -1 | 5,77E-01 | NA | 5 | 12 | 6 | 4 | 37 | 53 | 47 | 32 | 0,8 | 0,2 | 0,6 |
| chr3:131180544-131181044:downstream_2708 | chr3 | 131181044 | 131221860 | - | MRPL3 | NM | 72 | 0,4 | 1 | -1 | 5,77E-01 | 8,67E-01 | 11 | 14 | 13 | 18 | 98 | 97 | 185 | 138 | -0,3 | -0,7 | 0,4 |
| chr6:149912067-149912567:downstream_5394 | chr6 | 149887527 | 149912067 | + | GINM1 | NM | 94 | 0,4 | 1 | -1 | 5,77E-01 | 8,67E-01 | 203 | 97 | 145 | 92 | 66 | 43 | 56 | 55 | 0,3 | 0,0 | 0,4 |
| chr11:86151649-86152149:downstream_18124 | chr11 | 86152149 | 86383678 | - | ME3 | NM | 8 | -1,0 | 2 | 1 | 5,78E-01 | NA | 2 | 0 | 13 | 7 | 6 | 2 | 14 | 16 | -3,0 | -2,0 | -1,0 |
| chr11:93468595-93469095:downstream_9584 | chr11 | 93469095 | 93474703 | - | TAF1D | NM | 235 | 0,3 | 0 | -1 | 5,78E-01 | 8,67E-01 | 268 | 285 | 262 | 243 | 219 | 175 | 241 | 191 | 0,1 | -0,1 | 0,3 |
| chrX:107174867-107175367:downstream_26916 | chrX | 107069083 | 107174867 | + | MID2 | NM | 3 | -1,8 | 3 | 1 | 5,78E-01 | NA | 0 | 0 | 1 | 10 | 0 | 1 | 3 | 7 | -Inf | -3,3 | -Inf |
| chr17:18440614-18441114:downstream_12616 | chr17 | 18441114 | 18528930 | - | CCDC144B | NR | 6 | 1,1 | 2 | -1 | 5,80E-01 | NA | 3 | 0 | 11 | 6 | 2 | 0 | 11 | 16 | -2,4 | -3,5 | 1,1 |
| chr12:31478879-31479379:downstream_10120 | chr12 | 31477249 | 31478879 | + | FLJ13224 | NR | 11 | -1,3 | 2 | 1 | 5,81E-01 | NA | 13 | 13 | 0 | 0 | 42 | 21 | 0 | 0 | Inf | Inf | NaN |
| chr17:45761004-45761504:downstream_20534 | chr17 | 45727203 | 45761004 | + | KPNB1 | NM | 155 | -0,3 | 1 | 1 | 5,81E-01 | 8,69E-01 | 18 | 19 | 42 | 61 | 196 | 144 | 429 | 330 | -1,5 | -1,2 | -0,3 |
| chr2:28637516-28638016:downstream_1921 | chr2 | 28615778 | 28637516 | + | FOSL2 | NM | 78 | 0,3 | 1 | -1 | 5,81E-01 | 8,69E-01 | 90 | 87 | 78 | 80 | 59 | 77 | 81 | 71 | 0,2 | -0,2 | 0,3 |
| chr6:72011973-72012473:downstream_5221 | chr6 | 71998476 | 72011973 | + | OGFRL1 | NM | 31 | -0,5 | 1 | 1 | 5,81E-01 | 8,69E-01 | 11 | 8 | 4 | 8 | 84 | 61 | 43 | 25 | 0,6 | 1,1 | -0,4 |
| chr7:127009596-127010096:downstream_6066 | chr7 | 127010096 | 127032778 | - | ZNF800 | NM | 38 | 0,4 | 1 | -1 | 5,82E-01 | 8,69E-01 | 69 | 30 | 29 | 27 | 46 | 40 | 31 | 34 | 0,8 | 0,4 | 0,4 |
| chr1:145375705-145376205:downstream_16905 | chr1 | 145376205 | 145382404 | - | LOC101928979 | NR | 5 | 1,3 | 2 | -1 | 5,83E-01 | NA | 2 | 0 | 3 | 10 | 2 | 0 | 8 | 12 | -2,4 | -3,7 | 1,3 |
| chr7:140705460-140705960:downstream_25639 | chr7 | 140705960 | 140714781 | - | MRPS33 | NM | 75 | 0,4 | 1 | -1 | 5,83E-01 | 8,69E-01 | 30 | 13 | 14 | 11 | 133 | 168 | 131 | 102 | 0,7 | 0,4 | 0,4 |
| chr3:57547768-57548268:downstream_3016 | chr3 | 57541980 | 57547768 | + | PDE12 | NM | 26 | 0,5 | 1 | -1 | 5,83E-01 | NA | 33 | 34 | 7 | 12 | 54 | 34 | 19 | 18 | 1,8 | 1,2 | 0,5 |
| chr6:49519808-49520308:downstream_5174 | chr6 | 49518112 | 49519808 | + | C6orf141 | NM | 55 | 0,4 | 1 | -1 | 5,83E-01 | 8,70E-01 | 10 | 22 | 13 | 13 | 96 | 86 | 113 | 86 | 0,3 | -0,1 | 0,4 |
| chr2:98271901-98272401:downstream_2132 | chr2 | 98272401 | 98280561 | - | ACTR1B | NM | 17 | 0,6 | 1 | -1 | 5,84E-01 | NA | 8 | 5 | 8 | 7 | 20 | 18 | 28 | 38 | -0,2 | -0,8 | 0,6 |
| chr14:20756800-20757300:downstream_11026 | chr14 | 20757300 | 20774153 | - | TTC5 | NM | 17 | 0,6 | 1 | -1 | 5,84E-01 | NA | 16 | 28 | 22 | 16 | 12 | 10 | 15 | 13 | 0,2 | -0,4 | 0,6 |
| chr18:21715574-21716074:downstream_20964 | chr18 | 21572736 | 21715574 | + | TTC39C | NM | 3 | -1,7 | 3 | 1 | 5,84E-01 | NA | 0 | 0 | 3 | 11 | 1 | 0 | 1 | 11 | -Inf | -3,9 | -Inf |
| chr17:38438439-38438939:downstream_12635 | chr17 | 38375573 | 38438439 | + | WIPF2 | NM | 24 | -0,5 | 1 | 1 | 5,84E-01 | NA | 9 | 12 | 31 | 34 | 13 | 20 | 47 | 28 | -1,7 | -1,2 | -0,5 |
| chr6:154725932-154726432:downstream_4821 | chr6 | 154726432 | 154831753 | - | CNKSR3 | NM | 13 | 0,8 | 1 | -1 | 5,84E-01 | NA | 6 | 0 | 6 | 8 | 11 | 6 | 33 | 35 | -1,3 | -2,0 | 0,7 |
| chr5:54721409-54721909:downstream_4017 | chr5 | 54603575 | 54721409 | + | SKIV2L2 | NM | 21 | 0,5 | 1 | -1 | 5,85E-01 | NA | 11 | 14 | 18 | 20 | 10 | 22 | 26 | 46 | -0,6 | -1,2 | 0,5 |
| chr20:462837-463337:downstream_22392 | chr20 | 463337 | 524482 | - | CSNK2A1 | NM | 95 | -0,3 | 1 | 1 | 5,85E-01 | 8,70E-01 | 46 | 34 | 141 | 103 | 50 | 80 | 148 | 162 | -1,6 | -1,3 | -0,3 |
| chr9:102648099-102648599:downstream_7229 | chr9 | 102648599 | 102668883 | - | STX17-AS1 | NR | 6 | 1,0 | 2 | -1 | 5,85E-01 | NA | 7 | 16 | 0 | 7 | 7 | 6 | 7 | 2 | 1,8 | 0,6 | 1,2 |
| chr9:33920401-33920901:downstream_7033 | chr9 | 33817181 | 33920401 | + | UBE2R2 | NM | 12 | -0,9 | 2 | 1 | 5,85E-01 | NA | 2 | 0 | 3 | 10 | 6 | 15 | 26 | 30 | -2,4 | -1,5 | -0,9 |
| chr11:13753893-13754393:downstream_9153 | chr11 | 13690205 | 13753893 | + | FAR1 | NM | 9 | -0,9 | 2 | 1 | 5,85E-01 | NA | 1 | 3 | 1 | 9 | 11 | 12 | 15 | 22 | -1,5 | -0,8 | -0,7 |
| chr6:86386224-86386724:downstream_5252 | chr6 | 86386724 | 86388451 | - | SNHG5 | NR | 173 | -0,4 | 1 | 1 | 5,85E-01 | 8,70E-01 | 507 | 285 | 152 | 110 | 168 | 91 | 42 | 26 | 1,6 | 1,9 | -0,3 |

| | | | | | | | | | | | | | | | | | | | | | | | |
|---|---|---|---|---|---|---|---|---|---|---|---|---|---|---|---|---|---|---|---|---|---|---|---|
| chr10:81914379-81914879:downstream_17471 | chr10 | 81914879 | 81965433 | - | ANXA11 | NM | 13 | -0,6 | 1 | 1 | 5,85E-01 | NA | 7 | 7 | 13 | 18 | 14 | 9 | 17 | 17 | -1,2 | -0,6 | -0,6 |
| chr20:23025769-23026269:downstream_14914 | chr20 | 23026269 | 23030301 | - | THBD | NM | 5 | -1,2 | 2 | 1 | 5,86E-01 | NA | 2 | 0 | 10 | 7 | 3 | 0 | 10 | 5 | -3,4 | -2,2 | -1,2 |
| chrX:46184858-46185358:downstream_7688 | chrX | 46185358 | 46187109 | - | LINC01186 | NR | 9 | -0,9 | 2 | 1 | 5,86E-01 | NA | 10 | 0 | 19 | 14 | 8 | 3 | 10 | 11 | -1,8 | -0,9 | -0,9 |
| chr12:69239324-69239824:downstream_18674 | chr12 | 69201951 | 69239324 | + | MDM2 | NM | 88 | 0,4 | 1 | -1 | 5,86E-01 | 8,70E-01 | 36 | 32 | 1 | 7 | 275 | 259 | 45 | 50 | 3,1 | 2,5 | 0,6 |
| chr8:126036002-126036502:downstream_6355 | chr8 | 126036502 | 126104061 | - | KIAA0196 | NM | 9 | 0,9 | 2 | -1 | 5,86E-01 | NA | 7 | 0 | 6 | 14 | 2 | 7 | 18 | 22 | -1,4 | -2,3 | 0,8 |
| chr3:52574586-52575086:downstream_3002 | chr3 | 52570620 | 52574586 | + | SMIM4 | NM | 16 | 0,6 | 1 | -1 | 5,86E-01 | NA | 29 | 15 | 6 | 6 | 19 | 35 | 14 | 7 | 2,0 | 1,3 | 0,7 |
| chr17:36889649-36890149:downstream_13016 | chr17 | 36890149 | 36904558 | - | PCGF2 | NM | 5 | 1,1 | 2 | -1 | 5,87E-01 | NA | 2 | 6 | 8 | 3 | 0 | 6 | 8 | 10 | -0,6 | -1,6 | 1,0 |
| chr15:72878896-72879396:downstream_11500 | chr15 | 72766666 | 72878896 | + | ARIH1 | NM | 94 | 0,4 | 1 | -1 | 5,87E-01 | 8,70E-01 | 21 | 6 | 24 | 27 | 87 | 109 | 281 | 198 | -0,9 | -1,3 | 0,4 |
| chr5:40830929-40831429:downstream_4231 | chr5 | 40831429 | 40835387 | - | RPL37 | NM | 62 | 0,4 | 1 | -1 | 5,87E-01 | 8,70E-01 | 63 | 29 | 13 | 10 | 140 | 148 | 50 | 41 | 2,0 | 1,7 | 0,4 |
| chr7:106841688-106842188:downstream_25457 | chr7 | 106842188 | 107204959 | - | COG5 | NM | 13 | -0,8 | 1 | 1 | 5,87E-01 | NA | 2 | 2 | 3 | 27 | 5 | 9 | 22 | 32 | -2,8 | -2,0 | -0,8 |
| chr2:220100819-220101319:downstream_21939 | chr2 | 220101319 | 220110151 | - | GLB1L | NM | 9 | -0,8 | 1 | 1 | 5,87E-01 | NA | 4 | 5 | 6 | 3 | 14 | 21 | 11 | 9 | 0,0 | 0,8 | -0,8 |
| chr17:38613982-38614482:downstream_13047 | chr17 | 38599675 | 38613982 | + | IGFBP4 | NM | 440 | -0,3 | 1 | 1 | 5,87E-01 | 8,70E-01 | 15 | 23 | 32 | 31 | 643 | 799 | 910 | 1066 | -0,7 | -0,5 | -0,3 |
| chr9:34085880-34086380:downstream_7034 | chr9 | 34086380 | 34126771 | - | DCAF12 | NM | 18 | -0,7 | 1 | 1 | 5,87E-01 | NA | 3 | 0 | 6 | 6 | 22 | 20 | 47 | 39 | -1,8 | -1,0 | -0,8 |
| chr19:58906171-58906671:downstream_14727 | chr19 | 58898635 | 58906171 | + | RPS5 | NM | 173 | -0,4 | 1 | 1 | 5,88E-01 | 8,70E-01 | 45 | 11 | 17 | 13 | 482 | 448 | 157 | 213 | 0,9 | 1,3 | -0,4 |
| chr12:62995030-62995530:downstream_10289 | chr12 | 62995530 | 62997214 | - | LINC01465 | NR | 10 | 0,8 | 2 | -1 | 5,88E-01 | NA | 15 | 14 | 4 | 0 | 20 | 18 | 6 | 3 | 2,8 | 2,1 | 0,6 |
| chr4:119633425-119633925:downstream_3823 | chr4 | 119606524 | 119633425 | + | METTL14 | NM | 14 | -0,7 | 1 | 1 | 5,88E-01 | NA | 7 | 2 | 8 | 7 | 20 | 21 | 26 | 18 | -0,8 | -0,1 | -0,7 |
| chr2:99215285-99215785:downstream_1678 | chr2 | 99215785 | 99224981 | - | COA5 | NM | 11 | -0,7 | 1 | 1 | 5,88E-01 | NA | 15 | 16 | 13 | 6 | 18 | 13 | 10 | 2 | 0,8 | 1,4 | -0,7 |
| chr12:21684210-21684710:downstream_10098 | chr12 | 21679255 | 21684210 | + | SPX | NM | 27 | -0,5 | 1 | 1 | 5,88E-01 | NA | 51 | 57 | 39 | 36 | 15 | 8 | 6 | 6 | 0,5 | 1,0 | -0,5 |
| chr7:102955133-102955633:downstream_6030 | chr7 | 102937872 | 102955133 | + | PMPCB | NM | 22 | 0,5 | 1 | -1 | 5,88E-01 | NA | 25 | 22 | 26 | 12 | 24 | 16 | 26 | 20 | 0,3 | -0,2 | 0,5 |
| chr8:37603573-37604073:downstream_6522 | chr8 | 37604073 | 37605564 | - | LOC728024 | NR | 8 | -1,0 | 2 | 1 | 5,88E-01 | NA | 2 | 15 | 1 | 9 | 11 | 16 | 8 | 1 | 0,8 | 1,6 | -0,8 |
| chr7:44616993-44617493:downstream_25779 | chr7 | 44617493 | 44621894 | - | TMED4 | NM | 45 | -0,5 | 1 | 1 | 5,89E-01 | 8,71E-01 | 7 | 15 | 4 | 9 | 104 | 117 | 50 | 49 | 0,8 | 1,2 | -0,4 |
| chr3:44702283-44702783:downstream_2612 | chr3 | 44690232 | 44702283 | + | ZNF35 | NM | 14 | 0,6 | 1 | -1 | 5,89E-01 | NA | 13 | 15 | 7 | 3 | 22 | 27 | 8 | 17 | 1,5 | 0,9 | 0,5 |
| chrX:67261685-67262185:downstream_7512 | chrX | 67262185 | 67653299 | - | OPHN1 | NM | 3 | 1,4 | 3 | -1 | 5,89E-01 | NA | 1 | 2 | 6 | 6 | 1 | 0 | 1 | 8 | -2,0 | -3,6 | 1,5 |
| chr17:17396534-17397034:downstream_12868 | chr17 | 17380299 | 17396534 | + | MED9 | NM | 20 | -0,6 | 1 | 1 | 5,89E-01 | NA | 6 | 10 | 7 | 5 | 61 | 28 | 19 | 25 | 0,4 | 1,0 | -0,6 |
| chr11:9469674-9470174:downstream_8816 | chr11 | 9406168 | 9469674 | + | IPO7 | NM | 43 | 0,4 | 1 | -1 | 5,90E-01 | 8,71E-01 | 8 | 10 | 8 | 12 | 56 | 61 | 113 | 78 | -0,2 | -0,7 | 0,5 |
| chr17:27050865-27051365:downstream_20770 | chr17 | 27051365 | 27053949 | - | TLCD1 | NM | 24 | 0,5 | 1 | -1 | 5,90E-01 | NA | 12 | 25 | 6 | 7 | 55 | 40 | 22 | 26 | 1,5 | 1,0 | 0,6 |
| chr19:45169428-45169928:downstream_21365 | chr19 | 45147097 | 45169428 | + | PVR | NM | 23 | -0,6 | 1 | 1 | 5,90E-01 | NA | 9 | 12 | 26 | 10 | 36 | 23 | 21 | 48 | -0,8 | -0,2 | -0,6 |
| chr4:54326103-54326603:downstream_23656 | chr4 | 54243819 | 54326103 | + | FIP1L1 | NM | 8 | 0,8 | 2 | -1 | 5,90E-01 | NA | 6 | 3 | 8 | 8 | 2 | 6 | 10 | 17 | -0,9 | -1,7 | 0,8 |
| chr19:6379079-6379579:downstream_13922 | chr19 | 6379579 | 6393291 | - | GTF2F1 | NM | 16 | -0,6 | 1 | 1 | 5,91E-01 | NA | 15 | 10 | 6 | 11 | 21 | 38 | 13 | 14 | 0,6 | 1,2 | -0,6 |
| chr16:70412837-70413337:downstream_12452 | chr16 | 70413337 | 70472991 | - | ST3GAL2 | NM | 21 | 0,6 | 1 | -1 | 5,91E-01 | NA | 6 | 18 | 15 | 7 | 33 | 16 | 42 | 27 | 0,1 | -0,5 | 0,6 |
| chr6:99879190-99879690:downstream_4769 | chr6 | 99872746 | 99879190 | + | LOC101927365 | NR | 13 | -0,7 | 1 | 1 | 5,91E-01 | NA | 18 | 29 | 6 | 12 | 9 | 24 | 3 | 5 | 1,4 | 2,1 | -0,7 |
| chr17:56167618-56168118:downstream_13261 | chr17 | 56160779 | 56167618 | + | DYNLL2 | NM | 79 | -0,4 | 1 | 1 | 5,91E-01 | 8,72E-01 | 127 | 131 | 113 | 98 | 26 | 75 | 35 | 29 | 0,3 | 0,7 | -0,4 |
| chr6:86317001-86317501:downstream_25284 | chr6 | 86317501 | 86353043 | - | SYNCRIP | NM | 31 | -0,4 | 1 | 1 | 5,91E-01 | 8,72E-01 | 29 | 37 | 53 | 26 | 24 | 33 | 25 | 24 | -0,3 | 0,2 | -0,5 |
| chr10:98745585-98746085:downstream_8654 | chr10 | 98741040 | 98745585 | + | C10orf12 | NM | 22 | -0,6 | 1 | 1 | 5,91E-01 | NA | 16 | 6 | 53 | 30 | 9 | 11 | 32 | 17 | -1,9 | -1,3 | -0,6 |
| chr2:101636155-101636655:downstream_22196 | chr2 | 101618690 | 101636155 | + | RPL31 | NM | 27 | 0,7 | 1 | -1 | 5,92E-01 | NA | 20 | 2 | 3 | 2 | 52 | 89 | 26 | 25 | 2,1 | 1,5 | 0,6 |
| chr11:64084212-64084712:downstream_18007 | chr11 | 64072999 | 64084212 | + | ESRRA | NM | 38 | -0,5 | 1 | 1 | 5,92E-01 | 8,73E-01 | 6 | 3 | 8 | 10 | 65 | 47 | 106 | 62 | -1,1 | -0,6 | -0,5 |
| chr8:66514190-66514690:downstream_25852 | chr8 | 66514690 | 66546452 | - | ARMC1 | NM | 31 | 0,6 | 1 | -1 | 5,93E-01 | 8,73E-01 | 5 | 1 | 10 | 3 | 30 | 28 | 96 | 77 | -1,2 | -1,6 | 0,4 |
| chr17:10614875-10615375:downstream_12839 | chr17 | 10600926 | 10614875 | + | ADPRM | NM | 14 | 0,6 | 1 | -1 | 5,94E-01 | NA | 19 | 24 | 8 | 10 | 13 | 20 | 10 | 11 | 1,2 | 0,7 | 0,6 |
| chr19:46146092-46146592:downstream_21114 | chr19 | 46144751 | 46146092 | + | EML2-AS1 | NR | 9 | 1,2 | 2 | -1 | 5,94E-01 | NA | 23 | 17 | 0 | 1 | 29 | 3 | 0 | 2 | 5,6 | 4,3 | 1,3 |
| chr14:21819130-21819630:downstream_11046 | chr14 | 21819630 | 21852425 | - | SUPT16H | NM | 73 | 0,4 | 1 | -1 | 5,95E-01 | 8,74E-01 | 58 | 26 | 47 | 47 | 74 | 95 | 111 | 129 | -0,2 | -0,5 | 0,3 |
| chr1:32681297-32681797:downstream_612 | chr1 | 32681797 | 32687926 | - | TMEM234 | NM | 14 | -0,7 | 1 | 1 | 5,95E-01 | NA | 16 | 3 | 29 | 8 | 19 | 9 | 14 | 18 | -1,0 | -0,2 | -0,7 |
| chr9:26992634-26993134:downstream_6850 | chr9 | 26993134 | 27005691 | - | LRRC19 | NM | 5 | 1,5 | 3 | -1 | 5,95E-01 | NA | 2 | 27 | 0 | 1 | 2 | 9 | 0 | 1 | 5,1 | 3,7 | 1,5 |
| chr8:71549000-71549500:downstream_6313 | chr8 | 71549500 | 71581447 | - | LACTB2 | NM | 23 | 0,5 | 1 | -1 | 5,95E-01 | NA | 13 | 17 | 6 | 14 | 34 | 34 | 32 | 33 | 0,6 | 0,1 | 0,6 |
| chr16:1819820-1820320:downstream_12092 | chr16 | 1820320 | 1821710 | - | NME3 | NM | 7 | 0,8 | 2 | -1 | 5,96E-01 | NA | 6 | 11 | 4 | 7 | 10 | 4 | 11 | 7 | 0,5 | -0,4 | 0,9 |
| chr5:58264365-58264865:downstream_24295 | chr5 | 58264865 | 59783925 | - | PDE4D | NM | 54 | 0,7 | 1 | -1 | 5,96E-01 | 8,75E-01 | 2 | 0 | 19 | 11 | 11 | 3 | 198 | 189 | -4,3 | -4,8 | 0,6 |
| chr14:24708350-24708850:downstream_19420 | chr14 | 24708850 | 24711880 | - | TINF2 | NM | 14 | -0,6 | 1 | 1 | 5,97E-01 | NA | 8 | 5 | 11 | 7 | 23 | 20 | 19 | 18 | -0,4 | 0,2 | -0,6 |
| chr13:32973809-32974309:downstream_10579 | chr13 | 32889616 | 32973809 | + | BRCA2 | NM | 9 | -0,8 | 1 | 1 | 5,97E-01 | NA | 2 | 4 | 22 | 19 | 3 | 2 | 15 | 8 | -2,9 | -2,2 | -0,7 |
| chr10:3190821-3191321:downstream_8385 | chr10 | 3183792 | 3190821 | + | PITRM1-AS1 | NR | 5 | 1,2 | 2 | -1 | 5,98E-01 | NA | 0 | 12 | 1 | 1 | 5 | 15 | 7 | 2 | 2,4 | 1,2 | 1,2 |
| chr4:99983960-99984460:downstream_3790 | chr4 | 99916787 | 99983960 | + | METAP1 | NM | 45 | -0,5 | 1 | 1 | 5,98E-01 | 8,77E-01 | 2 | 3 | 10 | 7 | 36 | 58 | 141 | 103 | -1,9 | -1,4 | -0,5 |
| chr3:8609811-8610311:downstream_23239 | chr3 | 8543492 | 8609811 | + | LMCD1 | NM | 11 | 0,8 | 2 | -1 | 5,98E-01 | NA | 1 | 3 | 15 | 10 | 4 | 0 | 24 | 29 | -2,8 | -3,7 | 0,9 |
| chr9:19452500-19453000:downstream_6994 | chr9 | 19408924 | 19452500 | + | ACER2 | NM | 75 | 0,5 | 1 | -1 | 5,99E-01 | 8,78E-01 | 75 | 75 | 3 | 3 | 185 | 239 | 6 | 17 | 4,6 | 4,2 | 0,4 |
| chr8:144682485-144682985:downstream_6767 | chr8 | 144680073 | 144682485 | + | TIGD5 | NM | 17 | 0,6 | 1 | -1 | 6,00E-01 | NA | 10 | 5 | 7 | 7 | 17 | 28 | 40 | 22 | 0,1 | -0,5 | 0,6 |
| chr15:40327797-40328297:downstream_11449 | chr15 | 40226324 | 40327797 | + | EIF2AK4 | NM | 10 | 0,7 | 1 | -1 | 6,01E-01 | NA | 7 | 14 | 13 | 6 | 6 | 11 | 11 | 12 | 0,1 | -0,5 | 0,7 |
| chr2:190340288-190340788:downstream_22322 | chr2 | 190306158 | 190340288 | + | WDR75 | NM | 53 | 0,4 | 1 | -1 | 6,01E-01 | 8,78E-01 | 30 | 30 | 24 | 20 | 111 | 51 | 88 | 66 | 0,5 | 0,1 | 0,4 |
| chr3:42823899-42824399:downstream_23192 | chr3 | 42824399 | 42846027 | - | HIGD1A | NM | 12 | -0,8 | 1 | 1 | 6,01E-01 | NA | 26 | 25 | 17 | 10 | 3 | 8 | 3 | 1 | 0,9 | 1,6 | -0,7 |
| chr16:2289372-2289872:downstream_19978 | chr16 | 2289872 | 2301602 | - | ECI1 | NM | 9 | 0,7 | 1 | -1 | 6,02E-01 | NA | 11 | 6 | 4 | 6 | 15 | 9 | 13 | 11 | 0,8 | 0,1 | 0,7 |
| chr16:20744485-20744985:downstream_20241 | chr16 | 20744985 | 20753286 | - | THUMPD1 | NM | 70 | 0,4 | 1 | -1 | 6,03E-01 | 8,78E-01 | 8 | 11 | 8 | 7 | 132 | 116 | 163 | 112 | 0,3 | -0,2 | 0,4 |
| chr16:3341459-3341959:downstream_12149 | chr16 | 3333486 | 3341459 | + | ZNF263 | NM | 48 | 0,4 | 1 | -1 | 6,03E-01 | 8,78E-01 | 29 | 27 | 15 | 14 | 90 | 87 | 70 | 50 | 0,9 | 0,6 | 0,4 |
| chr16:9213555-9214055:downstream_12179 | chr16 | 9185536 | 9213555 | + | C16orf72 | NM | 20 | -0,6 | 1 | 1 | 6,03E-01 | NA | 18 | 7 | 8 | 11 | 54 | 23 | 26 | 15 | 0,4 | 0,9 | -0,5 |
| chr14:50250031-50250531:downstream_19318 | chr14 | 50250531 | 50319791 | - | NEMF | NM | 14 | 0,7 | 1 | -1 | 6,03E-01 | NA | 3 | 7 | 17 | 4 | 14 | 6 | 29 | 34 | -1,1 | -1,7 | 0,6 |
| chr12:66530215-66530715:downstream_18874 | chr12 | 66530715 | 66563852 | - | TMBIM4 | NM | 42 | 0,5 | 1 | -1 | 6,03E-01 | 8,78E-01 | 10 | 4 | 4 | 5 | 81 | 81 | 67 | 83 | 0,6 | 0,1 | 0,5 |
| chr16:19717173-19717673:downstream_12236 | chr16 | 19717673 | 19729492 | - | KNOP1 | NM | 49 | 0,4 | 1 | -1 | 6,04E-01 | 8,78E-01 | 18 | 40 | 14 | 29 | 68 | 78 | 61 | 87 | 0,4 | 0,0 | 0,4 |
| chr13:41765210-41765710:downstream_10719 | chr13 | 41765710 | 41768702 | - | KBTBD7 | NM | 12 | 0,6 | 1 | -1 | 6,04E-01 | NA | 7 | 14 | 13 | 9 | 13 | 8 | 21 | 14 | -0,1 | -0,7 | 0,6 |
| chr9:33041349-33041849:downstream_7025 | chr9 | 33041849 | 33076714 | - | SMU1 | NM | 6 | 0,9 | 2 | -1 | 6,04E-01 | NA | 8 | 5 | 3 | 2 | 5 | 15 | 4 | 7 | 1,6 | 0,7 | 0,8 |
| chr6:136577500-136578000:downstream_24524 | chr6 | 136578000 | 136610989 | - | BCLAF1 | NM | 31 | -0,4 | 1 | 1 | 6,04E-01 | 8,78E-01 | 8 | 19 | 15 | 20 | 45 | 52 | 40 | 52 | -0,3 | 0,1 | -0,4 |
| chr5:149823291-149823791:downstream_24341 | chr5 | 149823791 | 149829319 | - | RPS14 | NM | 45 | 0,4 | 1 | -1 | 6,04E-01 | 8,78E-01 | 61 | 47 | 17 | 31 | 68 | 62 | 29 | 47 | 1,2 | 0,8 | 0,4 |
| chr12:100430362-100430862:downstream_18902 | chr12 | 100430862 | 100536642 | - | UHRF1BP1L | NM | 34 | -0,5 | 1 | 1 | 6,04E-01 | 8,78E-01 | 3 | 5 | 24 | 45 | 20 | 8 | 96 | 73 | -3,1 | -2,6 | -0,5 |
| chr11:35832603-35833103:downstream_8871 | chr11 | 35684299 | 35832603 | + | TRIM44 | NM | 31 | -0,5 | 1 | 1 | 6,04E-01 | 8,78E-01 | 5 | 0 | 18 | 21 | 17 | 16 | 82 | 91 | -3,0 | -2,4 | -0,6 |
| chr17:29704695-29705195:downstream_20608 | chr17 | 29421944 | 29704695 | + | NF1 | NM | 15 | 0,8 | 2 | -1 | 6,04E-01 | NA | 3 | 0 | 3 | 16 | 2 | 8 | 45 | 44 | -2,5 | -3,2 | 0,7 |
| chr1:40223402-40223902:downstream_116 | chr1 | 40223902 | 40254533 | - | BMP8B | NM | 15 | 0,7 | 1 | -1 | 6,05E-01 | NA | 13 | 15 | 36 | 28 | 7 | 0 | 15 | 9 | -1,2 | -1,9 | 0,7 |
| chr4:114820939-114821439:downstream_3818 | chr4 | 114821439 | 114900878 | - | ARSJ | NM | 5 | -1,2 | 2 | 1 | 6,05E-01 | NA | 2 | 0 | 6 | 7 | 6 | 0 | 4 | 14 | -2,9 | -1,7 | -1,2 |
| chr17:73269976-73270476:downstream_12691 | chr17 | 73267379 | 73269976 | + | LOC100287042 | NR | 14 | 0,6 | 1 | -1 | 6,05E-01 | NA | 17 | 36 | 14 | 11 | 10 | 9 | 7 | 7 | 1,1 | 0,4 | 0,6 |
| chr16:19869792-19870292:downstream_20004 | chr16 | 19870292 | 19896956 | - | GPRC5B | NM | 9 | -0,7 | 1 | 1 | 6,06E-01 | NA | 3 | 10 | 17 | 16 | 3 | 7 | 4 | 11 | -1,4 | -0,6 | -0,8 |
| chr5:71655180-71655680:downstream_24319 | chr5 | 71616193 | 71655180 | + | PTCD2 | NM | 10 | 0,7 | 1 | -1 | 6,06E-01 | NA | 10 | 8 | 17 | 12 | 5 | 5 | 17 | 9 | -0,7 | -1,4 | 0,7 |
| chr19:51869851-51870351:downstream_14532 | chr19 | 51870351 | 51872257 | - | CLDND2 | NM | 5 | -0,9 | 2 | 1 | 6,06E-01 | NA | 6 | 1 | 6 | 7 | 5 | 7 | 10 | 2 | -1,0 | -0,1 | -0,9 |
| chr2:152333860-152334360:downstream_22189 | chr2 | 152266396 | 152333860 | + | RIF1 | NM | 36 | -0,5 | 1 | 1 | 6,06E-01 | 8,78E-01 | 2 | 4 | 14 | 17 | 26 | 28 | 114 | 81 | -2,3 | -1,9 | -0,5 |
| chr6:127520626-127521126:downstream_5339 | chr6 | 127440047 | 127520626 | + | RSPO3 | NM | 3 | -1,2 | 2 | 1 | 6,07E-01 | NA | 0 | 2 | 4 | 6 | 3 | 1 | 4 | 6 | -2,4 | -1,2 | -1,1 |
| chr17:8261230-8261730:downstream_12829 | chr17 | 8261730 | 8263859 | - | LOC100128288 | NR | 15 | 0,8 | 2 | -1 | 6,07E-01 | NA | 20 | 35 | 0 | 2 | 21 | 35 | 3 | 2 | 4,5 | 3,4 | 1,0 |
| chr1:225978168-225978668:downstream_17276 | chr1 | 225965514 | 225978168 | + | SRP9 | NM | 169 | 0,3 | 1 | -1 | 6,07E-01 | 8,78E-01 | 14 | 21 | 7 | 20 | 325 | 311 | 359 | 295 | 0,4 | 0,0 | 0,4 |
| chr21:44833897-44834397:downstream_15346 | chr21 | 44834397 | 44847002 | - | SIK1 | NM | 132 | 1,0 | 2 | -1 | 6,07E-01 | 8,78E-01 | 14 | 0 | 4 | 13 | 141 | 152 | 360 | 372 | -0,3 | -1,3 | 1,0 |
| chr10:93387696-93388196:downstream_8640 | chr10 | 93388196 | 93392858 | - | PPP1R3C | NM | 21 | 0,6 | 1 | -1 | 6,07E-01 | NA | 39 | 22 | 6 | 3 | 47 | 32 | 11 | 6 | 2,8 | 2,2 | 0,6 |
| chr1:192685520-192686020:downstream_1274 | chr1 | 192685457 | 192685520 | + | MIR4426 | NR | 257 | 0,3 | 1 | -1 | 6,07E-01 | 8,78E-01 | 82 | 33 | 15 | 20 | 811 | 561 | 212 | 320 | 1,7 | 1,4 | 0,3 |
| chr9:99517646-99518146:downstream_7209 | chr9 | 99518146 | 99540328 | - | ZNF510 | NM | 16 | -0,6 | 1 | 1 | 6,08E-01 | NA | 9 | 15 | 42 | 18 | 7 | 11 | 14 | 16 | -1,4 | -0,8 | -0,6 |
| chr14:69929107-69929607:downstream_19392 | chr14 | 69865095 | 69929107 | + | SLC39A9 | NM | 20 | -0,6 | 1 | 1 | 6,08E-01 | NA | 3 | 3 | 3 | 9 | 36 | 25 | 31 | 51 | -0,9 | -0,4 | -0,5 |
| chr7:138876732-138877232:downstream_25794 | chr7 | 138818489 | 138876732 | + | TTC26 | NM | 7 | -0,9 | 2 | 1 | 6,08E-01 | NA | 4 | 0 | 6 | 6 | 12 | 4 | 8 | 15 | -1,5 | -0,5 | -1,0 |
| chr5:175937075-175937575:downstream_4130 | chr5 | 175875355 | 175937075 | + | FAF2 | NM | 81 | -0,5 | 1 | 1 | 6,08E-01 | 8,78E-01 | 4 | 1 | 11 | 11 | 90 | 65 | 231 | 236 | -2,1 | -1,6 | -0,5 |
| chr19:44790000-44790500:downstream_14363 | chr19 | 44790500 | 44809178 | - | ZNF235 | NM | 7 | 0,8 | 2 | -1 | 6,08E-01 | NA | 8 | 5 | 3 | 7 | 9 | 5 | 10 | 8 | 0,5 | -0,4 | 0,9 |
| chr7:72982776-72983276:downstream_5890 | chr7 | 72983276 | 72993013 | - | TBL2 | NM | 28 | 0,5 | 1 | -1 | 6,08E-01 | 8,78E-01 | 24 | 17 | 7 | 14 | 51 | 44 | 31 | 36 | 1,0 | 0,5 | 0,5 |
| chr14:101373305-101373805:downstream_11327 | chr14 | 101361106 | 101373305 | + | MEG8 | NR | 5 | 1,1 | 2 | -1 | 6,08E-01 | NA | 12 | 1 | 4 | 11 | 2 | 2 | 0 | 8 | -0,2 | -1,2 | 1,0 |
| chr12:80166842-80167342:downstream_18756 | chr12 | 80167342 | 80329235 | - | PPP1R12A | NM | 12 | 0,9 | 2 | -1 | 6,09E-01 | NA | 2 | 0 | 11 | 5 | 4 | 0 | 45 | 28 | -3,3 | -4,2 | 0,9 |
| chr8:127564182-127564682:downstream_6713 | chr8 | 127564682 | 127570711 | - | FAM84B | NM | 35 | 0,5 | 1 | -1 | 6,09E-01 | 8,78E-01 | 22 | 19 | 0 | 4 | 88 | 112 | 13 | 21 | 3,3 | 2,6 | 0,8 |
| chr3:148890983-148891483:downstream_23195 | chr3 | 148847370 | 148890983 | + | HPS3 | NM | 46 | 0,4 | 1 | -1 | 6,09E-01 | 8,78E-01 | 27 | 70 | 93 | 65 | 18 | 18 | 53 | 28 | -0,7 | -1,2 | 0,4 |
| chr12:58196639-58197139:downstream_18893 | chr12 | 58176527 | 58196639 | + | TSFM | NM | 6 | 0,9 | 2 | -1 | 6,09E-01 | NA | 8 | 6 | 7 | 7 | 7 | 0 | 7 | 6 | 0,0 | -0,9 | 0,9 |
| chr12:57923931-57924431:downstream_10275 | chr12 | 57916658 | 57923931 | + | MBD6 | NM | 12 | 0,7 | 1 | -1 | 6,09E-01 | NA | 8 | 1 | 3 | 2 | 24 | 22 | 19 | 16 | 1,0 | 0,4 | 0,7 |
| chr11:46764583-46765083:downstream_17956 | chr11 | 46765083 | 46867859 | - | CKAP5 | NM | 39 | -0,6 | 1 | 1 | 6,10E-01 | 8,78E-01 | 3 | 0 | 7 | 7 | 28 | 53 | 107 | 104 | -2,1 | -1,4 | -0,7 |
| chr5:40824864-40825364:downstream_4232 | chr5 | 40825364 | 40829244 | - | LOC100506548 | NR | 38 | -0,4 | 1 | 1 | 6,10E-01 | 8,78E-01 | 39 | 24 | 10 | 11 | 94 | 83 | 24 | 20 | 1,6 | 2,0 | -0,4 |
| chr11:27359560-27360060:downstream_17931 | chr11 | 27360060 | 27384795 | - | CCDC34 | NM | 23 | 0,5 | 1 | -1 | 6,10E-01 | NA | 13 | 4 | 7 | 4 | 37 | 44 | 33 | 39 | 0,6 | 0,2 | 0,5 |
| chr17:8173809-8174309:downstream_12826 | chr17 | 8152595 | 8173809 | + | PFAS | NM | 17 | 0,6 | 1 | -1 | 6,11E-01 | NA | 7 | 7 | 3 | 7 | 28 | 25 | 28 | 31 | 0,5 | -0,1 | 0,6 |
| chr2:149545136-149545636:downstream_1726 | chr2 | 149402559 | 149545136 | + | EPC2 | NM | 8 | -0,8 | 2 | 1 | 6,11E-01 | NA | 0 | 3 | 13 | 12 | 3 | 3 | 18 | 13 | -3,1 | -2,3 | -0,8 |
| chr1:173832538-173833038:downstream_1233 | chr1 | 173833038 | 173837125 | - | GAS5 | NR | 64 | 0,3 | 1 | -1 | 6,11E-01 | 8,79E-01 | 128 | 97 | 63 | 54 | 38 | 65 | 31 | 36 | 0,9 | 0,6 | 0,3 |
| chr9:117546414-117546914:downstream_26543 | chr9 | 117546914 | 117568408 | - | TNFSF15 | NM | 70 | -0,6 | 1 | 1 | 6,12E-01 | 8,79E-01 | 2 | 0 | 8 | 18 | 24 | 42 | 262 | 204 | -3,4 | -2,8 | -0,6 |
| chr3:25705863-25706363:downstream_2860 | chr3 | 25706363 | 25706430 | - | MIR4442 | NR | 15 | -0,6 | 1 | 1 | 6,12E-01 | NA | 28 | 8 | 24 | 22 | 12 | 9 | 8 | 9 | -0,3 | 0,3 | -0,6 |
| chr15:75930925-75931425:downstream_11821 | chr15 | 75931425 | 75932664 | - | IMP3 | NM | 173 | 0,3 | 1 | -1 | 6,13E-01 | 8,79E-01 | 30 | 39 | 14 | 7 | 452 | 510 | 156 | 180 | 1,7 | 1,5 | 0,2 |
| chr1:10520102-10520602:downstream_16653 | chr1 | 10520602 | 10532613 | - | DFFA | NM | 8 | 0,8 | 2 | -1 | 6,13E-01 | NA | 2 | 9 | 8 | 11 | 7 | 2 | 11 | 16 | -0,9 | -1,7 | 0,8 |
| chr9:20341162-20341662:downstream_26396 | chr9 | 20341662 | 20622542 | - | MLLT3 | NM | 8 | 0,8 | 2 | -1 | 6,13E-01 | NA | 2 | 3 | 11 | 11 | 2 | 3 | 15 | 17 | -2,1 | -2,8 | 0,8 |
| chr7:43602938-43603438:downstream_25554 | chr7 | 43152197 | 43602938 | + | HECW1 | NM | 3 | -1,5 | 3 | 1 | 6,13E-01 | NA | 0 | 0 | 7 | 6 | 1 | 0 | 8 | 4 | -Inf | -3,9 | -Inf |
| chr17:48945732-48946232:downstream_13239 | chr17 | 48944039 | 48945732 | + | TOB1-AS1 | NR | 5 | 0,9 | 2 | -1 | 6,13E-01 | NA | 3 | 2 | 8 | 5 | 1 | 3 | 11 | 7 | -1,4 | -2,3 | 0,9 |
| chr7:105888231-105888731:downstream_5477 | chr7 | 105888731 | 105925638 | - | NAMPT | NM | 13 | 0,6 | 1 | -1 | 6,14E-01 | NA | 15 | 10 | 17 | 11 | 15 | 4 | 18 | 14 | -0,2 | -0,8 | 0,6 |
| chr19:40927908-40928408:downstream_14302 | chr19 | 40928408 | 40931932 | - | SERTAD1 | NM | 51 | -0,5 | 1 | 1 | 6,14E-01 | 8,79E-01 | 67 | 47 | 10 | 2 | 160 | 102 | 13 | 5 | 3,3 | 3,9 | -0,6 |
| chr19:48969367-48969867:downstream_14445 | chr19 | 48958963 | 48969367 | + | KCNJ14 | NM | 7 | -0,8 | 2 | 1 | 6,14E-01 | NA | 7 | 5 | 3 | 3 | 12 | 16 | 3 | 6 | 0,9 | 1,7 | -0,8 |
| chr12:116395880-116396380:downstream_9934 | chr12 | 116396380 | 116714991 | - | MED13L | NM | 64 | 0,4 | 1 | -1 | 6,14E-01 | 8,79E-01 | 2 | 14 | 71 | 72 | 7 | 19 | 192 | 137 | -3,2 | -3,6 | 0,5 |
| chr12:57910438-57910938:downstream_10273 | chr12 | 57881735 | 57910438 | + | MARS | NM | 88 | 0,4 | 1 | -1 | 6,15E-01 | 8,79E-01 | 4 | 5 | 3 | 6 | 137 | 148 | 171 | 227 | 0,1 | -0,5 | 0,5 |
| chr6:57050012-57050512:downstream_5209 | chr6 | 57037103 | 57050012 | + | BAG2 | NM | 16 | -0,7 | 1 | 1 | 6,15E-01 | NA | 10 | 0 | 35 | 15 | 12 | 5 | 38 | 15 | -2,3 | -1,6 | -0,7 |
| chr7:141356027-141356527:downstream_6117 | chr7 | 141356527 | 141401953 | - | KIAA1147 | NM | 32 | -0,5 | 1 | 1 | 6,15E-01 | 8,79E-01 | 9 | 1 | 38 | 58 | 11 | 9 | 78 | 51 | -3,3 | -2,7 | -0,5 |
| chr3:15089518-15090018:downstream_2837 | chr3 | 15090018 | 15106816 | - | MRPS25 | NM | 29 | 0,4 | 1 | -1 | 6,15E-01 | 8,79E-01 | 41 | 47 | 36 | 34 | 20 | 15 | 19 | 18 | 0,3 | -0,1 | 0,4 |
| chr1:38146741-38147241:downstream_16524 | chr1 | 38147241 | 38156267 | - | C1orf109 | NM | 10 | -0,7 | 1 | 1 | 6,16E-01 | NA | 4 | 9 | 3 | 3 | 18 | 29 | 8 | 6 | 1,1 | 1,7 | -0,7 |
| chr10:5260910-5261410:downstream_8200 | chr10 | 5238797 | 5260910 | + | AKR1C4 | NM | 4 | -1,1 | 2 | 1 | 6,16E-01 | NA | 2 | 0 | 4 | 7 | 2 | 4 | 7 | 4 | -2,1 | -1,0 | -1,1 |
| chr16:22430366-22430866:downstream_12256 | chr16 | 22430866 | 22449036 | - | RRN3P3 | NR | 14 | -0,5 | 1 | 1 | 6,16E-01 | NA | 8 | 10 | 18 | 12 | 15 | 15 | 22 | 14 | -0,8 | -0,2 | -0,5 |
| chr12:13029070-13029570:downstream_10072 | chr12 | 13028410 | 13029070 | + | RPL13AP20 | NR | 13 | -0,6 | 1 | 1 | 6,16E-01 | NA | 7 | 9 | 7 | 5 | 32 | 21 | 13 | 13 | 0,4 | 1,0 | -0,6 |
| chr14:24678286-24678786:downstream_11080 | chr14 | 24678786 | 24683036 | - | CHMP4A | NM | 37 | -0,4 | 1 | 1 | 6,17E-01 | 8,80E-01 | 16 | 28 | 32 | 24 | 55 | 44 | 60 | 36 | -0,3 | 0,1 | -0,4 |
| chr19:18499986-18500486:downstream_14128 | chr19 | 18496967 | 18499986 | + | GDF15 | NM | 36 | -0,5 | 1 | 0 | 6,17E-01 | 8,80E-01 | 12 | 1 | 6 | 5 | 82 | 89 | 50 | 42 | 0,3 | 0,9 | -0,6 |
| chr22:38338465-38338965:downstream_15604 | chr22 | 38302154 | 38338465 | + | MICALL1 | NM | 5 | 1,1 | 2 | 0 | 6,17E-01 | NA | 3 | 14 | 3 | 0 | 6 | 9 | 3 | 2 | 2,6 | 1,7 | 0,9 |
| chr8:146075050-146075550:downstream_25886 | chr8 | 146075550 | 146078963 | - | COMMD5 | NM | 11 | 0,7 | 1 | 0 | 6,17E-01 | NA | 9 | 15 | 1 | 5 | 14 | 24 | 10 | 7 | 2,0 | 1,2 | 0,8 |
| chr7:16831763-16832263:downstream_5719 | chr7 | 16832263 | 16844738 | - | AGR2 | NM | 2664 | -0,2 | 0 | 0 | 6,18E-01 | 8,80E-01 | 1454 | 1323 | 1239 | 1307 | 4659 | 4223 | 3355 | 3756 | 0,1 | 0,3 | -0,2 |
| chr1:156711398-156711898:downstream_16792 | chr1 | 156711898 | 156722240 | - | HDGF | NM | 218 | 0,3 | 1 | 0 | 6,19E-01 | 8,80E-01 | 11 | 20 | 17 | 9 | 381 | 459 | 406 | 438 | 0,3 | 0,0 | 0,3 |
| chr17:18873880-18874380:downstream_12889 | chr17 | 18874380 | 18908060 | - | FAM83G | NM | 22 | 0,5 | 1 | 0 | 6,19E-01 | NA | 19 | 43 | 28 | 20 | 15 | 19 | 14 | 22 | 0,4 | -0,1 | 0,5 |
| chr4:44702697-44703197:downstream_3430 | chr4 | 44680432 | 44702697 | + | GUF1 | NM | 49 | 0,4 | 1 | 0 | 6,19E-01 | 8,80E-01 | 23 | 52 | 13 | 15 | 98 | 95 | 53 | 41 | 1,5 | 1,0 | 0,4 |
| chr19:57957191-57957691:downstream_14697 | chr19 | 57946692 | 57957191 | + | ZNF749 | NM | 6 | -0,8 | 2 | 0 | 6,19E-01 | NA | 6 | 3 | 14 | 9 | 5 | 3 | 4 | 7 | -1,4 | -0,6 | -0,8 |
| chr1:169822229-169822729:downstream_1218 | chr1 | 169764549 | 169822229 | + | C1orf112 | NM | 15 | -0,6 | 1 | 0 | 6,19E-01 | NA | 3 | 7 | 21 | 21 | 9 | 10 | 32 | 21 | -2,0 | -1,5 | -0,5 |
| chr12:72181150-72181650:downstream_10319 | chr12 | 72148657 | 72181150 | + | RAB21 | NM | 74 | 0,3 | 1 | 0 | 6,19E-01 | 8,80E-01 | 35 | 51 | 24 | 38 | 107 | 126 | 110 | 101 | 0,5 | 0,1 | 0,4 |
| chr12:123255611-123256111:downstream_10478 | chr12 | 123237367 | 123255611 | + | DENR | NM | 18 | 0,6 | 1 | 0 | 6,19E-01 | NA | 7 | 10 | 3 | 4 | 43 | 28 | 26 | 21 | 1,2 | 0,6 | 0,6 |
| chr7:87463313-87463813:downstream_5930 | chr7 | 87463813 | 87505692 | - | SLC25A40 | NM | 3 | -1,3 | 3 | 0 | 6,20E-01 | NA | 1 | 0 | 6 | 5 | 2 | 0 | 6 | 7 | -3,7 | -2,3 | -1,4 |
| chr10:79816571-79817071:downstream_17752 | chr10 | 79793517 | 79816571 | + | RPS24 | NM | 17 | 1,1 | 2 | 0 | 6,20E-01 | NA | 28 | 30 | 0 | 0 | 31 | 44 | 1 | 0 | Inf | 5,7 | Inf |
| chr20:1148426-1148926:downstream_22517 | chr20 | 1093905 | 1148426 | + | PSMF1 | NM | 12 | 0,6 | 1 | 0 | 6,20E-01 | NA | 12 | 11 | 13 | 9 | 14 | 7 | 13 | 16 | 0,1 | -0,5 | 0,6 |
| chr10:112659764-112660264:downstream_17703 | chr10 | 112631552 | 112659764 | + | PDCD4 | NM | 21 | 0,5 | 1 | 0 | 6,21E-01 | NA | 5 | 10 | 13 | 14 | 24 | 11 | 61 | 31 | -0,9 | -1,4 | 0,6 |
| chr2:24251705-24252205:downstream_21777 | chr2 | 24252205 | 24270296 | - | C2orf44 | NM | 24 | 0,5 | 1 | 0 | 6,21E-01 | NA | 9 | 15 | 4 | 7 | 47 | 46 | 19 | 43 | 1,1 | 0,6 | 0,5 |
| chr7:120988404-120988904:downstream_25512 | chr7 | 120988904 | 121036422 | - | FAM3C | NM | 8 | -0,7 | 1 | 0 | 6,21E-01 | NA | 6 | 4 | 10 | 19 | 7 | 2 | 7 | 10 | -1,6 | -0,9 | -0,7 |
| chr10:5020158-5020658:downstream_8392 | chr10 | 5005453 | 5020158 | + | AKR1C1 | NM | 276 | -0,3 | 1 | 0 | 6,21E-01 | 8,82E-01 | 48 | 34 | 67 | 72 | 285 | 545 | 536 | 619 | -0,8 | -0,5 | -0,3 |
| chr8:117857672-117858172:downstream_6685 | chr8 | 117858172 | 117887105 | - | RAD21 | NM | 21 | 0,5 | 1 | 0 | 6,22E-01 | NA | 5 | 6 | 14 | 7 | 18 | 20 | 49 | 52 | -0,9 | -1,4 | 0,5 |
| chr1:169821714-169822214:downstream_17217 | chr1 | 169822214 | 169863100 | - | SCYL3 | NM | 18 | 0,6 | 1 | 0 | 6,22E-01 | NA | 7 | 5 | 6 | 16 | 13 | 15 | 58 | 21 | -0,9 | -1,5 | 0,6 |
| chr22:42968765-42969265:downstream_15658 | chr22 | 42969265 | 42978017 | - | RRP7BP | NR | 12 | 0,6 | 1 | 0 | 6,23E-01 | NA | 7 | 23 | 8 | 3 | 14 | 19 | 10 | 10 | 1,4 | 0,8 | 0,6 |
| chr22:38612517-38613017:downstream_22839 | chr22 | 38597938 | 38612517 | + | MAFF | NM | 22 | 0,9 | 2 | 0 | 6,23E-01 | NA | 8 | 4 | 0 | 1 | 57 | 88 | 4 | 17 | 3,9 | 2,8 | 1,1 |
| chr12:58142026-58142526:downstream_10282 | chr12 | 58138783 | 58142026 | + | TSPAN31 | NM | 30 | -0,4 | 1 | 0 | 6,23E-01 | 8,82E-01 | 16 | 26 | 13 | 17 | 64 | 43 | 32 | 25 | 0,5 | 0,9 | -0,4 |
| chr16:71678328-71678828:downstream_20169 | chr16 | 71678828 | 71758604 | - | PHLPP2 | NM | 11 | 0,9 | 2 | 0 | 6,23E-01 | NA | 0 | 3 | 1 | 11 | 7 | 1 | 38 | 26 | -2,1 | -3,1 | 1,0 |
| chr3:172347934-172348434:downstream_23297 | chr3 | 172348434 | 172429008 | - | NCEH1 | NM | 126 | -0,4 | 1 | 0 | 6,24E-01 | 8,82E-01 | 6 | 2 | 3 | 11 | 151 | 247 | 288 | 303 | -0,9 | -0,6 | -0,3 |
| chr18:77898228-77898728:downstream_13687 | chr18 | 77866914 | 77898228 | + | ADNP2 | NM | 29 | 0,4 | 1 | 0 | 6,24E-01 | 8,82E-01 | 37 | 24 | 11 | 17 | 34 | 51 | 29 | 25 | 1,1 | 0,7 | 0,5 |
| chr8:99053437-99053937:downstream_6656 | chr8 | 99053937 | 99057818 | - | RPL30 | NM | 41 | 0,5 | 1 | 0 | 6,24E-01 | 8,82E-01 | 11 | 15 | 3 | 4 | 120 | 97 | 47 | 33 | 2,0 | 1,4 | 0,5 |
| chr22:32014537-32015037:downstream_22911 | chr22 | 31892124 | 32014537 | + | SFI1 | NM | 3 | 1,1 | 2 | 0 | 6,26E-01 | NA | 2 | 8 | 1 | 1 | 2 | 9 | 3 | 2 | 2,1 | 1,0 | 1,1 |
| chr3:158262624-158263124:downstream_23399 | chr3 | 157827840 | 158262624 | + | RSRC1 | NM | 19 | 0,6 | 1 | 0 | 6,27E-01 | NA | 9 | 8 | 26 | 53 | 7 | 0 | 24 | 25 | -2,3 | -2,9 | 0,7 |
| chr1:11085549-11086049:downstream_432 | chr1 | 11072678 | 11085549 | + | TARDBP | NM | 21 | 0,5 | 1 | 0 | 6,27E-01 | NA | 20 | 21 | 4 | 11 | 39 | 32 | 19 | 21 | 1,4 | 0,8 | 0,6 |
| chr10:37635956-37636456:downstream_8267 | chr10 | 37598113 | 37635956 | + | LINC00993 | NR | 45 | -0,4 | 1 | 0 | 6,27E-01 | 8,86E-01 | 21 | 39 | 82 | 112 | 15 | 15 | 38 | 39 | -1,7 | -1,3 | -0,4 |
| chr2:169627960-169628460:downstream_21819 | chr2 | 169628460 | 169642939 | - | CERS6-AS1 | NR | 6 | 0,9 | 2 | 0 | 6,28E-01 | NA | 3 | 7 | 8 | 6 | 0 | 6 | 11 | 4 | -0,5 | -1,4 | 0,9 |
| chr1:150781680-150782180:downstream_16489 | chr1 | 150782180 | 150849244 | - | ARNT | NM | 10 | -0,8 | 2 | 0 | 6,28E-01 | NA | 0 | 7 | 13 | 4 | 5 | 17 | 19 | 12 | -1,3 | -0,5 | -0,8 |
| chr20:10604027-10604527:downstream_14743 | chr20 | 10415950 | 10604027 | + | SLX4IP | NM | 21 | 0,5 | 1 | 0 | 6,28E-01 | NA | 10 | 6 | 21 | 16 | 11 | 16 | 45 | 43 | -1,3 | -1,7 | 0,4 |
| chr8:57869987-57870487:downstream_6561 | chr8 | 57870487 | 57906430 | - | IMPAD1 | NM | 96 | -0,4 | 1 | 0 | 6,29E-01 | 8,87E-01 | 6 | 1 | 7 | 10 | 93 | 170 | 257 | 226 | -1,3 | -0,9 | -0,5 |
| chr13:80130212-80130712:downstream_19036 | chr13 | 80055258 | 80130212 | + | NDFIP2 | NM | 10 | 0,7 | 2 | 0 | 6,29E-01 | NA | 5 | 0 | 3 | 8 | 8 | 5 | 26 | 23 | -1,2 | -1,9 | 0,8 |

| | | | | | | | | | | | | | | | | | | | | | | | |
|---|---|---|---|---|---|---|---|---|---|---|---|---|---|---|---|---|---|---|---|---|---|---|---|
| chr16:2011976-2012476:downstream_12104 | chr16 | 2009516 | 2011976 | + | NDUFB10 | NM | 76 | 0,3 | 1 | 0 | 6,29E-01 | 8,87E-01 | 14 | 18 | 11 | 10 | 167 | 140 | 111 | 140 | 0,6 | 0,3 | 0,3 |
| chr17:40553966-40554466:downstream_13122 | chr17 | 40554466 | 40575338 | - | PTRF | NM | 14 | -0,6 | 1 | 0 | 6,29E-01 | NA | 6 | 3 | 0 | 6 | 33 | 34 | 17 | 16 | 0,6 | 1,0 | -0,5 |
| chr1:220957596-220958096:downstream_1373 | chr1 | 220921675 | 220957596 | + | MARC2 | NM | 25 | -0,4 | 1 | 0 | 6,30E-01 | NA | 15 | 10 | 36 | 39 | 14 | 17 | 38 | 34 | -1,6 | -1,2 | -0,4 |
| chr18:44633280-44633780:downstream_13623 | chr18 | 44633780 | 44676871 | - | HDHD2 | NM | 21 | -0,5 | 1 | 0 | 6,30E-01 | NA | 2 | 10 | 8 | 5 | 39 | 40 | 31 | 33 | -0,2 | 0,3 | -0,5 |
| chr1:54684056-54684556:downstream_746 | chr1 | 54665839 | 54684056 | + | MRPL37 | NM | 12 | -0,7 | 1 | 0 | 6,30E-01 | NA | 11 | 2 | 6 | 2 | 18 | 38 | 8 | 12 | 0,6 | 1,4 | -0,8 |
| chr12:30781414-30781914:downstream_18607 | chr12 | 30781914 | 30848929 | - | IPO8 | NM | 37 | -0,5 | 1 | 0 | 6,31E-01 | 8,88E-01 | 1 | 5 | 15 | 21 | 28 | 15 | 110 | 98 | -2,7 | -2,3 | -0,4 |
| chrX:100600143-100600643:downstream_27077 | chrX | 100600643 | 100603957 | - | TIMM8A | NM | 29 | 0,4 | 1 | 0 | 6,31E-01 | 8,88E-01 | 28 | 52 | 15 | 9 | 54 | 39 | 17 | 21 | 1,7 | 1,3 | 0,4 |
| chr11:67165883-67166383:downstream_18245 | chr11 | 67159422 | 67165883 | + | RAD9A | NM | 11 | 0,6 | 1 | 0 | 6,31E-01 | NA | 12 | 22 | 10 | 13 | 8 | 7 | 4 | 11 | 0,6 | 0,0 | 0,6 |
| chr19:33204702-33205202:downstream_14192 | chr19 | 33182866 | 33204702 | + | NUDT19 | NM | 22 | -0,5 | 1 | 0 | 6,32E-01 | NA | 5 | 16 | 19 | 20 | 28 | 21 | 33 | 34 | -0,9 | -0,4 | -0,4 |
| chr7:104621693-104622193:downstream_6033 | chr7 | 104622193 | 104631612 | - | LINC01004 | NR | 18 | 0,6 | 1 | 0 | 6,32E-01 | NA | 10 | 18 | 10 | 1 | 33 | 36 | 14 | 21 | 1,4 | 1,0 | 0,4 |
| chr12:48733292-48733792:downstream_18929 | chr12 | 48733792 | 48745029 | - | ZNF641 | NM | 11 | -0,6 | 1 | 0 | 6,32E-01 | NA | 3 | 4 | 10 | 7 | 11 | 15 | 11 | 30 | -1,3 | -0,6 | -0,7 |
| chr5:74657926-74658426:downstream_24162 | chr5 | 74632992 | 74657926 | + | HMGCR | NM | 26 | -0,5 | 1 | 0 | 6,32E-01 | NA | 7 | 2 | 15 | 12 | 30 | 23 | 63 | 59 | -1,7 | -1,2 | -0,5 |
| chr1:52812358-52812858:downstream_17425 | chr1 | 52607765 | 52812358 | + | ZFYVE9 | NM | 6 | 1,0 | 2 | 0 | 6,32E-01 | NA | 0 | 9 | 8 | 3 | 6 | 1 | 11 | 7 | -0,4 | -1,4 | 1,0 |
| chr12:56664750-56665250:downstream_18524 | chr12 | 56660641 | 56664750 | + | COQ10A | NM | 40 | 0,5 | 1 | 0 | 6,32E-01 | 8,89E-01 | 15 | 2 | 4 | 4 | 88 | 88 | 58 | 61 | 1,0 | 0,6 | 0,4 |
| chr10:99520664-99521164:downstream_17827 | chr10 | 99496877 | 99520664 | + | ZFYVE27 | NM | 17 | 0,6 | 1 | 0 | 6,33E-01 | NA | 15 | 8 | 0 | 5 | 29 | 50 | 19 | 11 | 2,2 | 1,4 | 0,8 |
| chr10:112657987-112658487:downstream_17487 | chr10 | 112658487 | 112679124 | - | BBIP1 | NM | 11 | -0,7 | 1 | 0 | 6,33E-01 | NA | 5 | 0 | 3 | 8 | 11 | 20 | 24 | 21 | -1,2 | -0,5 | -0,7 |
| chr10:90699731-90700231:downstream_8626 | chr10 | 90692440 | 90699731 | + | ACTA2-AS1 | NR | 5 | -0,9 | 2 | 0 | 6,34E-01 | NA | 7 | 2 | 10 | 3 | 10 | 2 | 6 | 4 | -0,6 | 0,3 | -0,9 |
| chr3:25638895-25639395:downstream_2592 | chr3 | 25639395 | 25705863 | - | TOP2B | NM | 36 | 0,4 | 1 | 0 | 6,34E-01 | 8,90E-01 | 3 | 4 | 11 | 12 | 22 | 25 | 95 | 118 | -1,7 | -2,2 | 0,5 |
| chr5:156345792-156346292:downstream_24413 | chr5 | 156346292 | 156390266 | - | TIMD4 | NM | 10 | 0,7 | 1 | 0 | 6,34E-01 | NA | 7 | 10 | 22 | 16 | 2 | 3 | 7 | 9 | -1,2 | -1,8 | 0,6 |
| chr7:150904422-150904922:downstream_25371 | chr7 | 150904922 | 150924460 | - | ABCF2 | NM | 4 | -1,3 | 3 | 0 | 6,34E-01 | NA | 7 | 3 | 1 | 0 | 14 | 9 | 1 | 0 | 2,8 | 4,0 | -1,3 |
| chr16:30708524-30709024:downstream_12305 | chr16 | 30709024 | 30709810 | - | LOC730183 | NM | 8 | -0,7 | 1 | 0 | 6,34E-01 | NA | 6 | 5 | 7 | 5 | 15 | 11 | 8 | 10 | -0,2 | 0,5 | -0,7 |
| chr10:101471002-101471502:downstream_8329 | chr10 | 101419262 | 101471002 | + | ENTPD7 | NM | 7 | 0,8 | 2 | 0 | 6,34E-01 | NA | 7 | 4 | 14 | 13 | 4 | 0 | 11 | 7 | -1,3 | -2,1 | 0,8 |
| chr5:138731955-138732455:downstream_4450 | chr5 | 138732455 | 138739776 | - | SPATA24 | NM | 4 | -0,9 | 2 | 0 | 6,34E-01 | NA | 4 | 2 | 3 | 7 | 4 | 6 | 7 | 2 | -0,8 | 0,1 | -0,8 |
| chr4:17627251-17627751:downstream_3637 | chr4 | 17616250 | 17627251 | + | MED28 | NM | 20 | -0,5 | 1 | 0 | 6,34E-01 | NA | 20 | 29 | 24 | 15 | 20 | 28 | 8 | 17 | 0,4 | 0,9 | -0,5 |
| chr2:225334366-225334866:downstream_21854 | chr2 | 225334866 | 225450114 | - | CUL3 | NM | 4 | -1,2 | 2 | 0 | 6,34E-01 | NA | 1 | 0 | 7 | 6 | 2 | 0 | 7 | 9 | -4,0 | -2,7 | -1,3 |
| chr1:173827682-173828182:downstream_245 | chr1 | 173793796 | 173827682 | + | DARS2 | NM | 63 | -0,4 | 1 | 0 | 6,35E-01 | 8,90E-01 | 9 | 8 | 8 | 17 | 68 | 140 | 124 | 129 | -0,6 | -0,3 | -0,3 |
| chr7:102952420-102952920:downstream_25488 | chr7 | 102952920 | 102985320 | - | DNAJC2 | NM | 27 | 0,4 | 1 | 0 | 6,36E-01 | NA | 31 | 44 | 42 | 19 | 28 | 12 | 19 | 24 | 0,3 | -0,1 | 0,4 |
| chr14:57935517-57936017:downstream_19165 | chr14 | 57936017 | 57960581 | - | C14orf105 | NM | 7 | 0,8 | 2 | 0 | 6,36E-01 | NA | 1 | 5 | 6 | 11 | 4 | 1 | 10 | 17 | -1,5 | -2,4 | 0,9 |
| chr6:42857502-42858002:downstream_5144 | chr6 | 42858002 | 42858554 | - | C6orf226 | NM | 8 | -0,8 | 2 | 0 | 6,36E-01 | NA | 15 | 18 | 4 | 2 | 13 | 13 | 0 | 2 | 2,3 | 3,4 | -1,1 |
| chr8:87484077-87484577:downstream_26097 | chr8 | 87484577 | 87521009 | - | RMDN1 | NM | 24 | 0,5 | 1 | 0 | 6,36E-01 | NA | 14 | 5 | 3 | 7 | 33 | 60 | 33 | 41 | 0,9 | 0,3 | 0,6 |
| chr9:32450832-32451332:downstream_26205 | chr9 | 32384600 | 32450832 | + | ACO1 | NM | 35 | 0,4 | 1 | 0 | 6,36E-01 | 8,90E-01 | 10 | 13 | 10 | 10 | 46 | 63 | 54 | 72 | 0,2 | -0,2 | 0,4 |
| chr19:44978144-44978644:downstream_21611 | chr19 | 44978644 | 45004575 | - | ZNF180 | NM | 11 | 0,6 | 1 | 0 | 6,36E-01 | NA | 12 | 19 | 8 | 11 | 8 | 12 | 7 | 11 | 0,7 | 0,1 | 0,6 |
| chr17:42268672-42269172:downstream_20349 | chr17 | 42269172 | 42275529 | - | ATXN7L3 | NM | 9 | -0,7 | 1 | 0 | 6,36E-01 | NA | 7 | 2 | 7 | 8 | 13 | 10 | 15 | 8 | -0,7 | -0,1 | -0,7 |
| chr3:9808353-9808853:downstream_23318 | chr3 | 9791627 | 9808353 | + | OGG1 | NM | 9 | 0,7 | 1 | 0 | 6,36E-01 | NA | 16 | 4 | 8 | 5 | 8 | 12 | 8 | 11 | 0,6 | 0,0 | 0,6 |
| chr15:34522366-34522866:downstream_19540 | chr15 | 34517197 | 34522366 | + | EMC4 | NM | 1194 | -0,3 | 1 | 0 | 6,36E-01 | 8,90E-01 | 35 | 19 | 26 | 12 | 3039 | 3034 | 1794 | 1593 | 0,5 | 0,8 | -0,4 |
| chr19:36238056-36238556:downstream_21351 | chr19 | 36236477 | 36238056 | + | PSENEN | NM | 97 | 0,3 | 1 | 0 | 6,36E-01 | 8,90E-01 | 43 | 75 | 31 | 35 | 190 | 162 | 116 | 126 | 0,8 | 0,5 | 0,3 |
| chr12:27119581-27120081:downstream_18567 | chr12 | 27091304 | 27119581 | + | FGFR1OP2 | NM | 20 | 0,6 | 1 | 0 | 6,36E-01 | NA | 7 | 7 | 4 | 2 | 48 | 42 | 40 | 15 | 1,2 | 0,7 | 0,5 |
| chr19:53858122-53858622:downstream_14577 | chr19 | 53837001 | 53858122 | + | ZNF845 | NM | 6 | -0,8 | 2 | 0 | 6,36E-01 | NA | 9 | 5 | 3 | 9 | 12 | 4 | 4 | 4 | 0,2 | 1,0 | -0,7 |
| chr4:144478642-144479142:downstream_3515 | chr4 | 144434615 | 144478642 | + | SMARCA5 | NM | 7 | 0,7 | 2 | 0 | 6,37E-01 | NA | 5 | 5 | 13 | 4 | 2 | 7 | 10 | 13 | -0,8 | -1,4 | 0,7 |
| chr8:128219126-128219626:downstream_6717 | chr8 | 128219626 | 128231513 | - | CCAT1 | NR | 11 | 0,6 | 1 | 0 | 6,37E-01 | NA | 7 | 8 | 18 | 22 | 6 | 1 | 8 | 20 | -1,4 | -2,1 | 0,7 |
| chr2:191068859-191069359:downstream_21965 | chr2 | 191069359 | 191184771 | - | HIBCH | NM | 19 | -0,5 | 1 | 0 | 6,37E-01 | NA | 4 | 3 | 14 | 13 | 23 | 10 | 32 | 54 | -2,0 | -1,4 | -0,5 |
| chr1:214510477-214510977:downstream_1364 | chr1 | 214454564 | 214510477 | + | SMYD2 | NM | 16 | -1,4 | 3 | 0 | 6,37E-01 | NA | 1 | 0 | 3 | 7 | 24 | 0 | 49 | 47 | -3,5 | -2,0 | -1,5 |
| chr2:190424815-190425315:downstream_2351 | chr2 | 190425315 | 190445537 | - | SLC40A1 | NM | 49 | 0,5 | 1 | 0 | 6,38E-01 | 8,92E-01 | 6 | 0 | 8 | 11 | 33 | 32 | 185 | 113 | -1,8 | -2,2 | 0,4 |
| chr11:64887598-64888098:downstream_9432 | chr11 | 64888098 | 64889672 | - | FAU | NM | 100 | 0,3 | 1 | 0 | 6,39E-01 | 8,92E-01 | 39 | 48 | 13 | 11 | 215 | 308 | 79 | 90 | 1,9 | 1,6 | 0,3 |
| chr2:89050452-89050952:downstream_1668 | chr2 | 88991175 | 89050452 | + | RPIA | NM | 13 | 0,6 | 1 | 0 | 6,39E-01 | NA | 1 | 10 | 6 | 3 | 15 | 23 | 29 | 21 | 0,2 | -0,4 | 0,6 |
| chr2:98264657-98265157:downstream_2131 | chr2 | 98262520 | 98264657 | + | COX5B | NM | 43 | 0,4 | 1 | 0 | 6,39E-01 | 8,92E-01 | 38 | 26 | 25 | 9 | 84 | 65 | 38 | 60 | 0,9 | 0,6 | 0,3 |
| chr5:131816800-131817300:downstream_4404 | chr5 | 131817300 | 131826465 | - | IRF1 | NM | 13 | 0,6 | 1 | 0 | 6,39E-01 | NA | 36 | 20 | 4 | 4 | 19 | 13 | 3 | 4 | 2,8 | 2,2 | 0,6 |
| chr3:23965187-23965687:downstream_23393 | chr3 | 23958294 | 23965187 | + | RPL15 | NM | 14 | 1,0 | 2 | 0 | 6,39E-01 | NA | 40 | 13 | 1 | 0 | 24 | 33 | 3 | 0 | 5,2 | 4,3 | 0,9 |
| chr8:17913424-17913924:downstream_25853 | chr8 | 17913924 | 17942507 | - | ASAH1 | NM | 88 | -0,4 | 1 | 0 | 6,40E-01 | 8,92E-01 | 4 | 5 | 4 | 7 | 174 | 161 | 175 | 175 | -0,4 | -0,1 | -0,3 |
| chr19:57933307-57933807:downstream_14696 | chr19 | 57922528 | 57933307 | + | ZNF17 | NM | 16 | -0,5 | 1 | 0 | 6,40E-01 | NA | 12 | 9 | 4 | 11 | 28 | 29 | 15 | 16 | 0,4 | 0,9 | -0,5 |
| chr5:121364295-121364795:downstream_4385 | chr5 | 121297655 | 121364295 | + | SRFBP1 | NM | 17 | 0,5 | 1 | 0 | 6,41E-01 | NA | 24 | 11 | 31 | 19 | 9 | 8 | 22 | 11 | -0,5 | -1,0 | 0,5 |
| chrX:54471731-54472231:downstream_7747 | chrX | 54466852 | 54471731 | + | TSR2 | NM | 80 | -0,3 | 1 | 0 | 6,42E-01 | 8,94E-01 | 33 | 18 | 17 | 21 | 204 | 143 | 118 | 87 | 0,5 | 0,8 | -0,3 |
| chr1:211916298-211916798:downstream_285 | chr1 | 211916798 | 212004114 | - | LPGAT1 | NM | 58 | -0,4 | 1 | 0 | 6,42E-01 | 8,94E-01 | 3 | 8 | 11 | 20 | 47 | 82 | 149 | 141 | -1,5 | -1,2 | -0,3 |
| chrX:67757127-67757627:downstream_27120 | chrX | 67718623 | 67757127 | + | YIPF6 | NM | 23 | 0,4 | 1 | 0 | 6,42E-01 | NA | 18 | 15 | 11 | 14 | 33 | 28 | 28 | 35 | 0,4 | -0,1 | 0,4 |
| chr5:40758980-40759480:downstream_24313 | chr5 | 40759480 | 40798297 | - | PRKAA1 | NM | 10 | 0,6 | 1 | 0 | 6,42E-01 | NA | 4 | 7 | 6 | 5 | 8 | 16 | 14 | 22 | 0,0 | -0,6 | 0,6 |
| chr14:102814118-102814618:downstream_11374 | chr14 | 102814618 | 102829253 | - | CINP | NM | 13 | 0,7 | 1 | 0 | 6,43E-01 | NA | 9 | 0 | 6 | 6 | 23 | 7 | 19 | 39 | -0,3 | -1,0 | 0,6 |
| chr17:27887188-27887688:downstream_12946 | chr17 | 27887688 | 27894042 | - | ABHD15 | NM | 11 | 0,6 | 1 | 0 | 6,43E-01 | NA | 4 | 13 | 4 | 5 | 7 | 22 | 17 | 15 | 0,5 | -0,1 | 0,6 |
| chr5:106712089-106712589:downstream_3973 | chr5 | 106712589 | 107006596 | - | EFNA5 | NM | 16 | 0,9 | 2 | 0 | 6,43E-01 | NA | 1 | 0 | 4 | 6 | 3 | 2 | 50 | 60 | -3,6 | -4,4 | 0,8 |
| chr1:150325704-150326204:downstream_982 | chr1 | 150293927 | 150325704 | + | PRPF3 | NM | 65 | 0,3 | 1 | 0 | 6,43E-01 | 8,95E-01 | 15 | 19 | 22 | 24 | 66 | 100 | 118 | 156 | -0,4 | -0,7 | 0,3 |
| chr16:31718745-31719245:downstream_12323 | chr16 | 31711933 | 31718745 | + | CLUHP3 | NR | 10 | 0,6 | 1 | 0 | 6,44E-01 | NA | 7 | 10 | 8 | 10 | 11 | 6 | 7 | 20 | -0,1 | -0,7 | 0,6 |
| chr7:100804557-100805057:downstream_6013 | chr7 | 100797685 | 100804557 | + | AP1S1 | NM | 11 | 0,7 | 1 | 0 | 6,44E-01 | NA | 23 | 7 | 1 | 9 | 10 | 22 | 7 | 11 | 1,5 | 0,8 | 0,7 |
| chr7:44101316-44101816:downstream_25479 | chr7 | 44084238 | 44101316 | + | DBNL | NM | 22 | 0,6 | 1 | 0 | 6,45E-01 | NA | 7 | 1 | 3 | 2 | 45 | 47 | 31 | 37 | 0,9 | 0,4 | 0,5 |
| chr6:109485115-109485615:downstream_24560 | chr6 | 109416355 | 109485115 | + | CEP57L1 | NM | 14 | -0,6 | 1 | 0 | 6,45E-01 | NA | 9 | 1 | 39 | 32 | 3 | 2 | 8 | 15 | -2,8 | -2,2 | -0,7 |
| chr11:82969634-82970134:downstream_17935 | chr11 | 82970134 | 82997450 | - | CCDC90B | NM | 19 | -0,5 | 1 | 0 | 6,45E-01 | NA | 48 | 27 | 15 | 10 | 24 | 18 | 7 | 3 | 1,6 | 2,0 | -0,5 |
| chr8:107764921-107765421:downstream_26041 | chr8 | 107282405 | 107764921 | + | OXR1 | NM | 7 | 1,0 | 2 | 0 | 6,46E-01 | NA | 8 | 0 | 8 | 0 | 2 | 12 | 11 | 13 | 0,0 | -0,9 | 0,8 |
| chr10:18966940-18967440:downstream_8435 | chr10 | 18948312 | 18966940 | + | ARL5B | NM | 37 | 0,3 | 1 | 0 | 6,46E-01 | 8,97E-01 | 38 | 38 | 21 | 21 | 49 | 55 | 36 | 37 | 0,8 | 0,5 | 0,3 |
| chr8:32622558-32623058:downstream_26033 | chr8 | 31497267 | 32622558 | + | NRG1 | NM | 54 | -0,4 | 1 | 0 | 6,46E-01 | 8,97E-01 | 9 | 24 | 107 | 109 | 9 | 21 | 72 | 80 | -2,7 | -2,3 | -0,4 |
| chr3:197598456-197598956:downstream_2789 | chr3 | 197518096 | 197598456 | + | LRCH3 | NM | 10 | -0,6 | 1 | 0 | 6,46E-01 | NA | 3 | 12 | 14 | 15 | 10 | 6 | 11 | 9 | -0,9 | -0,4 | -0,6 |
| chr11:125762879-125763379:downstream_18239 | chr11 | 125763379 | 125773116 | - | PUS3 | NM | 36 | 0,4 | 1 | 0 | 6,48E-01 | 8,98E-01 | 16 | 30 | 14 | 2 | 90 | 66 | 35 | 34 | 1,5 | 1,2 | 0,3 |
| chr14:67804080-67804580:downstream_11190 | chr14 | 67804580 | 67826720 | - | ATP6V1D | NM | 16 | 0,5 | 1 | 0 | 6,48E-01 | NA | 7 | 13 | 8 | 2 | 35 | 21 | 17 | 25 | 0,9 | 0,4 | 0,4 |
| chr9:103063856-103064356:downstream_26532 | chr9 | 103064356 | 103115259 | - | TEX10 | NM | 9 | -0,7 | 1 | 0 | 6,48E-01 | NA | 11 | 7 | 13 | 4 | 21 | 5 | 7 | 8 | 0,1 | 0,8 | -0,7 |
| chr15:44826202-44826702:downstream_19539 | chr15 | 44826702 | 44829121 | - | EIF3J-AS1 | NR | 12 | -0,6 | 1 | 0 | 6,48E-01 | NA | 17 | 3 | 4 | 6 | 21 | 31 | 10 | 7 | 1,0 | 1,6 | -0,6 |
| chrX:75398145-75398645:downstream_26958 | chrX | 75392763 | 75398145 | + | PBDC1 | NM | 56 | 0,3 | 1 | 0 | 6,49E-01 | 8,98E-01 | 63 | 66 | 22 | 21 | 111 | 85 | 39 | 43 | 1,6 | 1,3 | 0,3 |
| chr12:122219974-122220474:downstream_9945 | chr12 | 122150657 | 122219974 | + | TMEM120B | NM | 6 | -0,9 | 2 | 0 | 6,49E-01 | NA | 2 | 0 | 3 | 7 | 2 | 7 | 14 | 13 | -2,6 | -1,7 | -1,0 |
| chr19:49379319-49379819:downstream_14458 | chr19 | 49375648 | 49379319 | + | PPP1R15A | NM | 42 | 0,4 | 1 | 0 | 6,49E-01 | 8,98E-01 | 83 | 51 | 10 | 9 | 66 | 92 | 13 | 16 | 2,8 | 2,5 | 0,3 |
| chr16:21994668-21995168:downstream_12251 | chr16 | 21964608 | 21994668 | + | UQCRC2 | NM | 158 | -0,3 | 1 | 0 | 6,49E-01 | 8,98E-01 | 23 | 25 | 25 | 30 | 273 | 313 | 264 | 309 | -0,2 | 0,0 | -0,2 |
| chr17:17149437-17149937:downstream_20415 | chr17 | 17149937 | 17184617 | - | COPS3 | NM | 41 | 0,4 | 1 | 0 | 6,50E-01 | 8,98E-01 | 6 | 8 | 11 | 3 | 72 | 57 | 63 | 110 | -0,1 | -0,4 | 0,3 |
| chr16:71507475-71507975:downstream_12458 | chr16 | 71507975 | 71523254 | - | ZNF19 | NM | 5 | 0,8 | 2 | 0 | 6,50E-01 | NA | 7 | 3 | 3 | 6 | 5 | 3 | 4 | 7 | 0,3 | -0,6 | 0,8 |
| chr1:186390503-186391003:downstream_16537 | chr1 | 186344889 | 186390503 | + | C1orf27 | NM | 27 | -0,5 | 1 | 0 | 6,50E-01 | NA | 4 | 2 | 4 | 7 | 63 | 25 | 64 | 49 | -0,8 | -0,4 | -0,5 |
| chr16:47114930-47115430:downstream_20139 | chr16 | 47115430 | 47177936 | - | NETO2 | NM | 18 | 0,5 | 1 | 0 | 6,51E-01 | NA | 3 | 8 | 7 | 12 | 15 | 17 | 40 | 42 | -0,8 | -1,4 | 0,5 |
| chr2:74687683-74688183:downstream_22084 | chr2 | 74688183 | 74692537 | - | MOGS | NM | 20 | 0,5 | 1 | 0 | 6,51E-01 | NA | 9 | 8 | 18 | 10 | 22 | 15 | 36 | 45 | -0,7 | -1,2 | 0,4 |
| chr22:42908566-42909066:downstream_15656 | chr22 | 42896584 | 42908566 | + | SERHL | NR | 9 | -0,9 | 2 | 0 | 6,52E-01 | NA | 4 | 24 | 1 | 3 | 4 | 28 | 1 | 2 | 2,6 | 3,4 | -0,8 |
| chr2:29406679-29407179:downstream_21836 | chr2 | 29320541 | 29406679 | + | CLIP4 | NM | 9 | 0,7 | 2 | 0 | 6,52E-01 | NA | 2 | 8 | 14 | 6 | 2 | 7 | 8 | 22 | -1,1 | -1,7 | 0,7 |
| chr5:151121882-151122382:downstream_4111 | chr5 | 151122382 | 151138210 | - | ATOX1 | NM | 5 | 0,9 | 2 | 0 | 6,52E-01 | NA | 8 | 1 | 1 | 3 | 7 | 7 | 7 | 6 | 1,0 | 0,1 | 0,9 |
| chr16:81413803-81414303:downstream_12030 | chr16 | 81348570 | 81413803 | + | GAN | NM | 26 | -0,4 | 1 | 0 | 6,53E-01 | NA | 23 | 23 | 54 | 34 | 14 | 15 | 22 | 21 | -0,9 | -0,5 | -0,4 |
| chr10:44113352-44113852:downstream_8490 | chr10 | 44101854 | 44113352 | + | ZNF485 | NM | 9 | -0,7 | 2 | 0 | 6,53E-01 | NA | 5 | 4 | 1 | 2 | 28 | 20 | 6 | 8 | 1,2 | 1,8 | -0,6 |
| chr21:40556903-40557403:downstream_22627 | chr21 | 40557403 | 40685712 | - | BRWD1 | NM | 11 | -0,6 | 1 | 0 | 6,53E-01 | NA | 2 | 1 | 19 | 11 | 5 | 4 | 25 | 23 | -3,1 | -2,5 | -0,7 |
| chr10:70968854-70969354:downstream_17781 | chr10 | 70939959 | 70968854 | + | SUPV3L1 | NM | 62 | 0,4 | 1 | 0 | 6,53E-01 | 8,99E-01 | 5 | 8 | 6 | 3 | 135 | 114 | 120 | 103 | 0,5 | 0,2 | 0,3 |
| chr15:25296719-25297219:downstream_11560 | chr15 | 25296622 | 25296719 | + | SNORD116-1 | NR | 5 | -0,9 | 2 | 0 | 6,53E-01 | NA | 2 | 0 | 4 | 7 | 3 | 4 | 7 | 11 | -2,2 | -1,3 | -0,9 |
| chr1:75627218-75627718:downstream_16875 | chr1 | 75594118 | 75627218 | + | LHX8 | NM | 21 | -0,5 | 1 | 0 | 6,53E-01 | NA | 7 | 0 | 15 | 16 | 10 | 21 | 46 | 54 | -2,3 | -1,7 | -0,6 |
| chr17:42252841-42253341:downstream_20338 | chr17 | 42253341 | 42264085 | - | ASB16-AS1 | NR | 26 | -0,4 | 1 | 0 | 6,54E-01 | NA | 34 | 30 | 18 | 20 | 46 | 28 | 15 | 18 | 0,8 | 1,2 | -0,4 |
| chr9:27062931-27063431:downstream_26353 | chr9 | 26947036 | 27062931 | + | IFT74 | NM | 15 | 0,6 | 1 | 0 | 6,54E-01 | NA | 15 | 4 | 25 | 33 | 4 | 4 | 11 | 23 | -1,6 | -2,1 | 0,5 |
| chr3:179305754-179306254:downstream_23283 | chr3 | 179306254 | 179322434 | - | MRPL47 | NM | 35 | 0,3 | 1 | 0 | 6,54E-01 | 8,99E-01 | 33 | 33 | 13 | 16 | 58 | 61 | 32 | 33 | 1,2 | 0,9 | 0,4 |
| chr2:26360323-26360823:downstream_1590 | chr2 | 26256728 | 26360323 | + | RAB10 | NM | 120 | -0,4 | 1 | 0 | 6,54E-01 | 8,99E-01 | 1 | 7 | 7 | 21 | 118 | 103 | 323 | 382 | -1,9 | -1,7 | -0,2 |
| chr16:67181504-67182004:downstream_12408 | chr16 | 67182004 | 67184902 | - | B3GNT9 | NM | 13 | 0,6 | 1 | 0 | 6,54E-01 | NA | 5 | 3 | 6 | 7 | 19 | 7 | 26 | 33 | -0,6 | -1,2 | 0,6 |
| chr3:45590328-45590828:downstream_2615 | chr3 | 45430074 | 45590328 | + | LARS2 | NM | 13 | 0,7 | 1 | 0 | 6,54E-01 | NA | 3 | 0 | 8 | 4 | 10 | 3 | 35 | 37 | -1,9 | -2,5 | 0,6 |
| chr6:125595995-125596495:downstream_5332 | chr6 | 125596495 | 125623282 | - | HDDC2 | NM | 25 | 0,4 | 1 | 0 | 6,54E-01 | NA | 21 | 20 | 6 | 18 | 38 | 36 | 25 | 34 | 0,8 | 0,3 | 0,5 |
| chr7:102740210-102740710:downstream_25391 | chr7 | 102715327 | 102740210 | + | ARMC10 | NM | 13 | 0,6 | 1 | 0 | 6,55E-01 | NA | 23 | 3 | 17 | 11 | 9 | 11 | 19 | 12 | -0,1 | -0,7 | 0,6 |
| chr10:96361856-96362356:downstream_17594 | chr10 | 96305523 | 96361856 | + | HELLS | NM | 50 | 0,9 | 2 | 0 | 6,55E-01 | 8,99E-01 | 23 | 45 | 104 | 91 | 21 | 0 | 58 | 56 | -1,5 | -2,4 | 0,9 |
| chr1:220141441-220141941:downstream_52 | chr1 | 220141941 | 220220000 | - | EPRS | NM | 118 | 0,3 | 1 | 0 | 6,55E-01 | 8,99E-01 | 3 | 7 | 10 | 11 | 125 | 117 | 380 | 294 | -1,1 | -1,5 | 0,4 |
| chr5:64885006-64885506:downstream_24424 | chr5 | 64885506 | 64920187 | - | TRIM23 | NM | 17 | -0,5 | 1 | 0 | 6,55E-01 | NA | 7 | 20 | 32 | 21 | 15 | 9 | 11 | 21 | -0,9 | -0,4 | -0,5 |
| chr7:43670250-43670750:downstream_5806 | chr7 | 43670750 | 43769140 | - | COA1 | NR | 19 | -0,5 | 1 | 0 | 6,56E-01 | NA | 8 | 15 | 19 | 36 | 7 | 18 | 21 | 23 | -1,2 | -0,8 | -0,5 |
| chr2:96850102-96850602:downstream_1672 | chr2 | 96850602 | 96874573 | - | STARD7 | NM | 93 | 0,3 | 1 | 0 | 6,56E-01 | 8,99E-01 | 11 | 11 | 15 | 8 | 170 | 138 | 180 | 211 | -0,1 | -0,3 | 0,2 |
| chr12:57909870-57910370:downstream_18536 | chr12 | 57910370 | 57914300 | - | DDIT3 | NM | 40 | 0,4 | 1 | 0 | 6,56E-01 | 8,99E-01 | 63 | 79 | 11 | 15 | 81 | 44 | 18 | 12 | 2,5 | 2,0 | 0,4 |
| chr12:12875316-12875816:downstream_10069 | chr12 | 12870203 | 12875316 | + | CDKN1B | NM | 19 | 0,5 | 1 | 0 | 6,56E-01 | NA | 5 | 4 | 7 | 5 | 16 | 29 | 43 | 41 | -0,4 | -0,9 | 0,5 |
| chrX:135594503-135595003:downstream_26857 | chrX | 135579237 | 135594503 | + | HTATSF1 | NM | 54 | 0,9 | 2 | 0 | 6,56E-01 | 8,99E-01 | 0 | 11 | 1 | 5 | 91 | 104 | 100 | 124 | 0,8 | -0,2 | 1,0 |
| chr5:158713048-158713548:downstream_4547 | chr5 | 158690088 | 158713048 | + | UBLCP1 | NM | 39 | 0,4 | 1 | 0 | 6,57E-01 | 8,99E-01 | 16 | 12 | 11 | 11 | 60 | 70 | 64 | 70 | 0,3 | 0,0 | 0,3 |
| chr15:56382230-56382730:downstream_11473 | chr15 | 56382730 | 56535483 | - | RFX7 | NM | 9 | -1,0 | 2 | 0 | 6,57E-01 | NA | 0 | 0 | 4 | 14 | 2 | 0 | 18 | 30 | -Inf | -4,9 | -Inf |
| chr17:2225481-2225981:downstream_12586 | chr17 | 2225981 | 2240678 | - | TSR1 | NM | 49 | 0,3 | 1 | 0 | 6,57E-01 | 8,99E-01 | 60 | 50 | 40 | 38 | 61 | 48 | 50 | 45 | 0,5 | 0,2 | 0,3 |
| chr20:34288902-34289402:downstream_14997 | chr20 | 34287231 | 34288902 | + | ROMO1 | NM | 33 | 0,5 | 1 | 0 | 6,57E-01 | 8,99E-01 | 6 | 5 | 3 | 2 | 110 | 50 | 39 | 52 | 1,3 | 0,8 | 0,4 |
| chr20:57569741-57570241:downstream_15114 | chr20 | 57570241 | 57582309 | - | CTSZ | NM | 199 | -0,4 | 1 | 0 | 6,58E-01 | 8,99E-01 | 3 | 6 | 1 | 7 | 424 | 459 | 330 | 363 | 0,2 | 0,3 | -0,2 |
| chr15:85349663-85350163:downstream_11519 | chr15 | 85291817 | 85349663 | + | ZNF592 | NM | 7 | 0,8 | 2 | 0 | 6,58E-01 | NA | 1 | 10 | 4 | 12 | 2 | 7 | 10 | 13 | -0,7 | -1,4 | 0,8 |
| chr10:95427139-95427639:downstream_8646 | chr10 | 95427639 | 95462329 | - | FRA10AC1 | NM | 8 | 0,7 | 2 | 0 | 6,58E-01 | NA | 16 | 5 | 13 | 9 | 7 | 0 | 6 | 6 | 0,0 | -0,8 | 0,8 |
| chr8:121063157-121063657:downstream_25905 | chr8 | 120885894 | 121063157 | + | DEPTOR | NM | 5 | -1,1 | 2 | 0 | 6,58E-01 | NA | 0 | 0 | 8 | 3 | 2 | 0 | 11 | 16 | -Inf | -4,1 | -Inf |
| chr9:127908351-127908851:downstream_26453 | chr9 | 127908851 | 127952218 | - | PPP6C | NM | 5 | -0,9 | 2 | 0 | 6,58E-01 | NA | 1 | 1 | 4 | 7 | 6 | 1 | 8 | 14 | -2,6 | -1,7 | -0,8 |
| chr8:71484952-71485452:downstream_6595 | chr8 | 71485452 | 71520694 | - | TRAM1 | NM | 97 | -0,3 | 1 | 0 | 6,59E-01 | 9,00E-01 | 12 | 14 | 17 | 19 | 187 | 146 | 163 | 216 | -0,5 | -0,2 | -0,3 |
| chr9:91931618-91932118:downstream_7175 | chr9 | 91926109 | 91931618 | + | CKS2 | NM | 28 | 0,4 | 1 | 0 | 6,59E-01 | NA | 45 | 55 | 10 | 15 | 41 | 33 | 11 | 13 | 2,0 | 1,6 | 0,4 |
| chr20:31438211-31438711:downstream_14965 | chr20 | 31407698 | 31438211 | + | MAPRE1 | NM | 50 | 0,5 | 1 | 0 | 6,60E-01 | 9,00E-01 | 9 | 0 | 4 | 3 | 62 | 122 | 85 | 113 | 0,3 | -0,1 | 0,4 |
| chr9:97064111-97064611:downstream_26581 | chr9 | 97021547 | 97064111 | + | ZNF169 | NM | 6 | 0,7 | 2 | 0 | 6,60E-01 | NA | 7 | 5 | 8 | 5 | 3 | 5 | 13 | 2 | -0,1 | -0,9 | 0,8 |
| chr21:38122218-38122718:downstream_22708 | chr21 | 38071432 | 38122218 | + | SIM2 | NM | 6 | -0,9 | 2 | 0 | 6,60E-01 | NA | 5 | 0 | 11 | 11 | 7 | 0 | 1 | 14 | -2,2 | -1,2 | -1,0 |
| chr12:48176536-48177036:downstream_10135 | chr12 | 48166966 | 48176536 | + | SLC48A1 | NM | 10 | -0,6 | 1 | 0 | 6,61E-01 | NA | 10 | 17 | 7 | 5 | 15 | 14 | 4 | 4 | 1,2 | 1,8 | -0,6 |
| chr5:43287071-43287571:downstream_24163 | chr5 | 43287571 | 43313614 | - | HMGCS1 | NM | 16 | 0,6 | 1 | 0 | 6,61E-01 | NA | 38 | 31 | 6 | 25 | 13 | 5 | 1 | 10 | 1,2 | 0,7 | 0,5 |
| chr5:122952738-122953238:downstream_24076 | chr5 | 122847792 | 122952738 | + | CSNK1G3 | NM | 8 | -0,7 | 2 | 0 | 6,61E-01 | NA | 2 | 1 | 10 | 11 | 7 | 2 | 15 | 19 | -2,6 | -1,9 | -0,7 |
| chr12:58019177-58019677:downstream_18450 | chr12 | 58019677 | 58027022 | - | B4GALNT1 | NM | 9 | -0,6 | 1 | 0 | 6,61E-01 | NA | 5 | 4 | 10 | 8 | 11 | 9 | 14 | 12 | -1,0 | -0,4 | -0,6 |
| chr10:106027593-106028093:downstream_8711 | chr10 | 106028093 | 106028163 | - | MIR4482 | NR | 33 | -0,4 | 1 | 0 | 6,61E-01 | 9,00E-01 | 54 | 43 | 7 | 14 | 56 | 72 | 13 | 10 | 2,2 | 2,5 | -0,3 |
| chr6:41757634-41758134:downstream_5131 | chr6 | 41755180 | 41757634 | + | TOMM6 | NM | 76 | 0,3 | 1 | 0 | 6,61E-01 | 9,00E-01 | 25 | 26 | 8 | 11 | 178 | 189 | 72 | 98 | 1,4 | 1,1 | 0,3 |
| chr20:47729375-47729875:downstream_22576 | chr20 | 47729875 | 47804904 | - | STAU1 | NM | 48 | 0,9 | 2 | 0 | 6,62E-01 | 9,00E-01 | 5 | 2 | 14 | 0 | 38 | 41 | 146 | 136 | -1,0 | -1,8 | 0,8 |
| chr9:35753264-35753764:downstream_7058 | chr9 | 35749276 | 35753264 | + | RGP1 | NM | 24 | 0,4 | 1 | 0 | 6,62E-01 | NA | 15 | 42 | 32 | 38 | 14 | 12 | 15 | 26 | -0,3 | -0,7 | 0,4 |
| chr14:37147011-37147511:downstream_11112 | chr14 | 37126772 | 37147011 | + | PAX9 | NM | 17 | -0,5 | 1 | 0 | 6,62E-01 | NA | 20 | 22 | 10 | 10 | 20 | 36 | 3 | 15 | 1,1 | 1,7 | -0,6 |
| chr1:100729797-100730297:downstream_17203 | chr1 | 100730297 | 100731730 | - | RTCA-AS1 | NR | 5 | 0,8 | 2 | 0 | 6,62E-01 | NA | 7 | 6 | 6 | 5 | 8 | 0 | 10 | 2 | 0,3 | -0,6 | 0,9 |
| chr19:11257330-11257830:downstream_14005 | chr19 | 11257830 | 11266484 | - | SPC24 | NM | 4 | 1,0 | 2 | 0 | 6,62E-01 | NA | 2 | 2 | 6 | 8 | 0 | 2 | 1 | 10 | -1,7 | -2,5 | 0,9 |
| chrX:153194779-153195279:downstream_26937 | chrX | 153195279 | 153200607 | - | NAA10 | NM | 18 | 0,5 | 1 | 0 | 6,63E-01 | NA | 7 | 6 | 4 | 5 | 42 | 18 | 28 | 36 | 0,4 | -0,1 | 0,5 |
| chr15:65293349-65293849:downstream_11764 | chr15 | 65293849 | 65321977 | - | MTFMT | NM | 3 | 1,0 | 2 | 0 | 6,63E-01 | NA | 8 | 4 | 0 | 2 | 7 | 3 | 1 | 2 | 2,9 | 1,6 | 1,2 |
| chr4:165996729-165997229:downstream_3898 | chr4 | 165997229 | 166034024 | - | TMEM192 | NM | 12 | -0,6 | 1 | 0 | 6,64E-01 | NA | 2 | 5 | 17 | 7 | 18 | 3 | 25 | 21 | -1,7 | -1,1 | -0,6 |
| chr6:131894605-131895105:downstream_25010 | chr6 | 131895105 | 131949379 | - | MED23 | NM | 6 | 0,8 | 2 | 0 | 6,64E-01 | NA | 7 | 14 | 4 | 2 | 3 | 13 | 4 | 3 | 1,8 | 1,1 | 0,7 |
| chr1:112264185-112264685:downstream_16718 | chr1 | 112264685 | 112298419 | - | FAM212B | NM | 8 | -0,8 | 2 | 0 | 6,64E-01 | NA | 18 | 8 | 3 | 1 | 20 | 10 | 1 | 1 | 2,8 | 3,7 | -0,9 |
| chr13:28710479-28710979:downstream_10685 | chr13 | 28710979 | 28713311 | - | PAN3-AS1 | NR | 18 | 0,5 | 1 | 0 | 6,65E-01 | NA | 35 | 26 | 11 | 7 | 28 | 17 | 6 | 12 | 1,7 | 1,3 | 0,4 |
| chr22:41220038-41220538:downstream_22926 | chr22 | 41220538 | 41253012 | - | ST13 | NM | 7 | -0,6 | 1 | 0 | 6,65E-01 | NA | 3 | 5 | 6 | 9 | 8 | 8 | 7 | 11 | -0,9 | -0,2 | -0,6 |
| chr2:99934986-99935486:downstream_2140 | chr2 | 99935486 | 99952860 | - | TXNDC9 | NM | 44 | -0,4 | 1 | 0 | 6,65E-01 | 9,02E-01 | 10 | 16 | 21 | 9 | 90 | 68 | 74 | 63 | -0,2 | 0,2 | -0,4 |
| chr11:17398868-17399368:downstream_9160 | chr11 | 17373308 | 17398868 | + | NCR3LG1 | NM | 15 | 0,5 | 1 | 0 | 6,65E-01 | NA | 38 | 26 | 10 | 10 | 15 | 8 | 4 | 6 | 1,7 | 1,2 | 0,5 |
| chr12:110971736-110972236:downstream_9925 | chr12 | 110972236 | 111021064 | - | PPTC7 | NM | 16 | -0,5 | 1 | 0 | 6,65E-01 | NA | 15 | 9 | 32 | 23 | 6 | 13 | 13 | 16 | -1,2 | -0,7 | -0,5 |
| chr11:64011607-64012107:downstream_18022 | chr11 | 64008412 | 64011607 | + | FKBP2 | NM | 80 | 0,4 | 1 | 0 | 6,65E-01 | 9,02E-01 | 4 | 19 | 13 | 2 | 220 | 130 | 135 | 121 | 0,7 | 0,4 | 0,3 |
| chr4:53738650-53739150:downstream_3439 | chr4 | 53739150 | 54232242 | - | SCFD2 | NM | 4 | 1,0 | 2 | 0 | 6,65E-01 | NA | 0 | 8 | 7 | 8 | 2 | 0 | 4 | 6 | -1,0 | -2,0 | 1,1 |
| chrX:140269430-140269930:downstream_7969 | chrX | 140269930 | 140271399 | - | LDOC1 | NM | 37 | 0,4 | 1 | 0 | 6,65E-01 | 9,02E-01 | 6 | 13 | 7 | 4 | 79 | 71 | 61 | 55 | 0,7 | 0,4 | 0,4 |
| chr1:153618782-153619282:downstream_16600 | chr1 | 153606457 | 153618782 | + | CHTOP | NM | 44 | 0,4 | 1 | 0 | 6,66E-01 | 9,02E-01 | 30 | 10 | 17 | 16 | 68 | 69 | 71 | 69 | 0,3 | 0,0 | 0,3 |
| chr11:62326572-62327072:downstream_9394 | chr11 | 62327072 | 62341460 | - | EEF1G | NM | 240 | 0,3 | 1 | 0 | 6,66E-01 | 9,02E-01 | 24 | 5 | 11 | 5 | 616 | 534 | 406 | 322 | 0,9 | 0,7 | 0,2 |
| chr7:143004789-143005289:downstream_25428 | chr7 | 142985307 | 143004789 | + | CASP2 | NM | 10 | 0,6 | 1 | 0 | 6,66E-01 | NA | 5 | 3 | 8 | 3 | 10 | 11 | 17 | 26 | -0,6 | -1,1 | 0,5 |
| chr10:105148822-105149322:downstream_8704 | chr10 | 105127723 | 105148822 | + | TAF5 | NM | 11 | 0,6 | 1 | 0 | 6,66E-01 | NA | 6 | 23 | 13 | 10 | 7 | 12 | 6 | 15 | 0,4 | -0,2 | 0,5 |
| chr12:46119002-46119502:downstream_10131 | chr12 | 46119502 | 46121704 | - | LINC00938 | NR | 19 | 0,5 | 1 | 0 | 6,67E-01 | NA | 33 | 34 | 8 | 2 | 33 | 26 | 7 | 6 | 2,6 | 2,2 | 0,4 |
| chr8:144361824-144362324:downstream_25984 | chr8 | 144362324 | 144363870 | - | LINC01604 | NR | 28 | -0,6 | 1 | 0 | 6,68E-01 | NA | 6 | 4 | 3 | 1 | 54 | 118 | 18 | 21 | 1,4 | 2,2 | -0,7 |
| chr6:116262192-116262692:downstream_5311 | chr6 | 116262692 | 116381921 | - | FRK | NM | 36 | -0,4 | 1 | 0 | 6,68E-01 | 9,04E-01 | 5 | 6 | 43 | 16 | 27 | 16 | 92 | 85 | -2,5 | -2,0 | -0,4 |
| chr15:69564544-69565044:downstream_11497 | chr15 | 69452972 | 69564544 | + | GLCE | NM | 15 | -0,9 | 2 | 0 | 6,68E-01 | NA | 1 | 0 | 3 | 7 | 14 | 0 | 38 | 55 | -3,6 | -2,7 | -0,9 |
| chr2:27548215-27548715:downstream_21955 | chr2 | 27548715 | 27579901 | - | GTF3C2 | NM | 24 | 0,5 | 1 | 0 | 6,68E-01 | NA | 5 | 5 | 6 | 3 | 31 | 49 | 53 | 43 | 0,1 | -0,3 | 0,4 |
| chrX:18909915-18910415:downstream_7620 | chrX | 18910415 | 19002480 | - | PHKA2 | NM | 5 | -1,0 | 2 | 0 | 6,69E-01 | NA | 0 | 1 | 0 | 9 | 4 | 1 | 13 | 14 | -3,2 | -2,4 | -0,8 |
| chr17:1324146-1324646:downstream_20419 | chr17 | 1324646 | 1359561 | - | CRK | NM | 77 | 0,3 | 1 | 0 | 6,69E-01 | 9,05E-01 | 25 | 17 | 14 | 32 | 96 | 125 | 185 | 118 | -0,1 | -0,5 | 0,4 |
| chr1:86049648-86050148:downstream_810 | chr1 | 86046443 | 86049648 | + | CYR61 | NM | 9 | 0,7 | 2 | 0 | 6,69E-01 | NA | 6 | 3 | 0 | 2 | 28 | 16 | 10 | 7 | 2,4 | 1,4 | 1,0 |

| | | | | | | | | | | | | | | | | | | | | | | | |
|---|---|---|---|---|---|---|---|---|---|---|---|---|---|---|---|---|---|---|---|---|---|---|---|
| chr19:55740646-55741146:downstream_14656 | chr19 | 55741146 | 55770038 | - | PPP6R1 | NM | 13 | 0,5 | 1 | 0 | 6,69E-01 | NA | 6 | 6 | 3 | 4 | 23 | 24 | 17 | 25 | 0,7 | 0,2 | 0,6 |
| chr15:99926514-99927014:downstream_19607 | chr15 | 99791566 | 99926514 | + | LRRC28 | NM | 5 | 0,9 | 2 | 0 | 6,70E-01 | NA | 2 | 3 | 17 | 11 | 0 | 1 | 6 | 5 | -2,6 | -3,4 | 0,8 |
| chrX:2852172-2852672:downstream_26627 | chrX | 2852672 | 2886351 | - | ARSE | NM | 30 | -0,4 | 1 | 0 | 6,70E-01 | 9,05E-01 | 20 | 16 | 26 | 33 | 45 | 20 | 28 | 52 | -0,7 | -0,3 | -0,4 |
| chr8:96256640-96257140:downstream_6651 | chr8 | 96257140 | 96281462 | - | C8orf37 | NM | 5 | -0,9 | 2 | 0 | 6,70E-01 | NA | 2 | 0 | 13 | 9 | 2 | 0 | 6 | 6 | -3,1 | -2,2 | -0,9 |
| chr11:93467775-93468275:downstream_9579 | chr11 | 93468275 | 93468402 | - | SNORA40 | NR | 261 | 0,2 | 0 | 0 | 6,71E-01 | 9,05E-01 | 196 | 137 | 124 | 121 | 416 | 401 | 342 | 348 | 0,4 | 0,2 | 0,2 |
| chr7:65994791-65995291:downstream_25543 | chr7 | 65995291 | 66057394 | - | GS1-124K5.11 | NR | 5 | 0,8 | 2 | 0 | 6,71E-01 | NA | 3 | 7 | 0 | 6 | 6 | 8 | 3 | 11 | 0,8 | -0,1 | 0,9 |
| chr1:159996961-159997461:downstream_1154 | chr1 | 159997461 | 160001783 | - | PIGM | NM | 30 | 0,4 | 1 | 0 | 6,71E-01 | 9,05E-01 | 27 | 16 | 13 | 8 | 58 | 53 | 29 | 37 | 1,1 | 0,7 | 0,3 |
| chr11:64883374-64883874:downstream_9437 | chr11 | 64883874 | 64885210 | - | ZNHIT2 | NM | 17 | -0,5 | 1 | 0 | 6,71E-01 | NA | 13 | 25 | 6 | 9 | 42 | 21 | 7 | 11 | 1,4 | 1,9 | -0,5 |
| chr4:124324915-124325415:downstream_23873 | chr4 | 124317949 | 124324915 | + | SPRY1 | NM | 16 | 0,5 | 1 | 0 | 6,71E-01 | NA | 14 | 11 | 6 | 1 | 30 | 45 | 10 | 15 | 1,9 | 1,6 | 0,3 |
| chr9:91093622-91094122:downstream_7172 | chr9 | 91003296 | 91093622 | + | SPIN1 | NM | 9 | -0,7 | 2 | 0 | 6,71E-01 | NA | 2 | 0 | 6 | 6 | 9 | 6 | 21 | 20 | -2,2 | -1,5 | -0,8 |
| chr10:74035797-74036297:downstream_8560 | chr10 | 74033676 | 74035797 | + | DDIT4 | NM | 104 | 0,3 | 1 | 0 | 6,72E-01 | 9,06E-01 | 32 | 23 | 25 | 16 | 167 | 230 | 171 | 171 | 0,4 | 0,2 | 0,2 |
| chr19:56156989-56157489:downstream_14668 | chr19 | 56154985 | 56156989 | + | ZNF581 | NM | 5 | 0,8 | 2 | 0 | 6,72E-01 | NA | 7 | 9 | 3 | 2 | 8 | 2 | 3 | 3 | 1,5 | 0,7 | 0,8 |
| chr8:90803292-90803792:downstream_6630 | chr8 | 90769974 | 90803292 | + | RIPK2 | NM | 16 | -0,5 | 1 | 0 | 6,73E-01 | NA | 4 | 1 | 14 | 10 | 10 | 14 | 35 | 39 | -2,2 | -1,7 | -0,6 |
| chr6:39156246-39156746:downstream_4728 | chr6 | 39156746 | 39197251 | - | KCNK5 | NM | 13 | 0,5 | 1 | 0 | 6,73E-01 | NA | 6 | 5 | 21 | 13 | 7 | 4 | 24 | 24 | -1,7 | -2,2 | 0,5 |
| chr9:96941202-96941702:downstream_7196 | chr9 | 96941115 | 96941202 | + | MIRLET7D | NR | 17 | 0,4 | 1 | 0 | 6,73E-01 | NA | 15 | 11 | 13 | 12 | 25 | 13 | 31 | 19 | 0,1 | -0,4 | 0,5 |
| chr6:12297427-12297927:downstream_24741 | chr6 | 12290528 | 12297427 | + | EDN1 | NM | 161 | 0,2 | 1 | 0 | 6,74E-01 | 9,06E-01 | 22 | 43 | 40 | 40 | 189 | 269 | 334 | 349 | -0,3 | -0,6 | 0,3 |
| chr1:157093958-157094458:downstream_16707 | chr1 | 157094458 | 157108383 | - | ETV3 | NM | 51 | -0,3 | 1 | 0 | 6,74E-01 | 9,06E-01 | 85 | 78 | 53 | 47 | 37 | 58 | 26 | 21 | 0,7 | 1,0 | -0,3 |
| chr4:53456626-53457126:downstream_23992 | chr4 | 53457126 | 53525502 | - | USP46 | NM | 9 | -0,7 | 2 | 0 | 6,75E-01 | NA | 0 | 5 | 24 | 20 | 2 | 2 | 10 | 11 | -3,2 | -2,5 | -0,6 |
| chr12:62997549-62998049:downstream_10288 | chr12 | 62997465 | 62997549 | + | MIRLET7I | NR | 133 | -0,2 | 1 | 0 | 6,75E-01 | 9,06E-01 | 82 | 108 | 298 | 363 | 28 | 25 | 90 | 66 | -1,8 | -1,6 | -0,2 |
| chr7:17502568-17503068:downstream_5515 | chr7 | 17503068 | 17598533 | - | LOC101927630 | NR | 11 | -0,5 | 1 | 0 | 6,75E-01 | NA | 11 | 4 | 26 | 19 | 5 | 5 | 7 | 13 | -1,6 | -1,0 | -0,6 |
| chr1:210030910-210031410:downstream_1350 | chr1 | 210001311 | 210030910 | + | DIEXF | NM | 19 | 0,6 | 1 | 0 | 6,76E-01 | NA | 11 | 5 | 0 | 2 | 45 | 69 | 10 | 12 | 3,2 | 2,4 | 0,9 |
| chr6:26022278-26022778:downstream_4908 | chr6 | 26021906 | 26022278 | + | HIST1H4A | NM | 18 | 0,5 | 1 | 0 | 6,76E-01 | NA | 7 | 24 | 8 | 16 | 25 | 19 | 22 | 25 | 0,4 | -0,1 | 0,5 |
| chr15:44096232-44096732:downstream_11696 | chr15 | 44096732 | 44116951 | - | MFAP1 | NM | 12 | 0,6 | 1 | 0 | 6,76E-01 | NA | 17 | 13 | 1 | 10 | 23 | 11 | 14 | 7 | 1,4 | 0,7 | 0,7 |
| chr2:74186088-74186588:downstream_21867 | chr2 | 74153952 | 74186088 | + | DGUOK | NM | 20 | 0,5 | 1 | 0 | 6,76E-01 | NA | 11 | 4 | 6 | 4 | 29 | 45 | 22 | 40 | 0,7 | 0,3 | 0,4 |
| chrX:71491952-71492452:downstream_7792 | chrX | 71492452 | 71497141 | - | RPS4X | NM | 487 | -0,2 | 1 | 0 | 6,76E-01 | 9,06E-01 | 60 | 44 | 31 | 47 | 988 | 1258 | 710 | 756 | 0,4 | 0,6 | -0,2 |
| chr17:76792464-76792964:downstream_12701 | chr17 | 76792964 | 76836969 | - | USP36 | NM | 26 | -0,4 | 1 | 0 | 6,76E-01 | NA | 32 | 36 | 31 | 25 | 19 | 34 | 15 | 18 | 0,3 | 0,7 | -0,4 |
| chr18:46445722-46446222:downstream_20945 | chr18 | 46446222 | 46477081 | - | SMAD7 | NM | 9 | -0,7 | 2 | 0 | 6,77E-01 | NA | 4 | 6 | 0 | 3 | 21 | 23 | 7 | 4 | 1,6 | 2,0 | -0,4 |
| chr12:120936298-120936798:downstream_18547 | chr12 | 120907659 | 120936298 | + | DYNLL1 | NM | 207 | 0,3 | 1 | 0 | 6,77E-01 | 9,06E-01 | 11 | 9 | 7 | 5 | 480 | 477 | 330 | 336 | 0,8 | 0,5 | 0,2 |
| chr2:70106727-70107227:downstream_2012 | chr2 | 70056817 | 70106727 | + | GMCL1 | NM | 30 | -0,4 | 1 | 0 | 6,78E-01 | 9,06E-01 | 13 | 10 | 72 | 63 | 7 | 7 | 35 | 30 | -2,6 | -2,2 | -0,4 |
| chr1:68154021-68154521:downstream_16758 | chr1 | 68150859 | 68154021 | + | GADD45A | NM | 165 | -0,3 | 1 | 0 | 6,78E-01 | 9,06E-01 | 150 | 82 | 14 | 11 | 555 | 429 | 39 | 45 | 3,2 | 3,5 | -0,3 |
| chr10:101611949-101612449:downstream_8331 | chr10 | 101542354 | 101611949 | + | ABCC2 | NM | 254 | -0,2 | 1 | 0 | 6,78E-01 | 9,06E-01 | 101 | 145 | 256 | 336 | 163 | 224 | 347 | 458 | -1,3 | -1,1 | -0,2 |
| chr1:155533735-155534235:downstream_1098 | chr1 | 155531771 | 155533735 | + | ASH1L-AS1 | NR | 31 | -0,4 | 1 | 0 | 6,78E-01 | 9,06E-01 | 28 | 20 | 8 | 5 | 91 | 65 | 17 | 15 | 1,9 | 2,3 | -0,4 |
| chr5:140078903-140079403:downstream_24158 | chr5 | 140071010 | 140078903 | + | HARS2 | NM | 59 | -0,4 | 1 | 0 | 6,78E-01 | 9,06E-01 | 5 | 2 | 7 | 11 | 67 | 75 | 160 | 143 | -1,4 | -1,1 | -0,4 |
| chr17:1952701-1953201:downstream_12733 | chr17 | 1953201 | 1953302 | - | MIR132 | NR | 9 | 0,7 | 2 | 0 | 6,79E-01 | NA | 15 | 27 | 3 | 1 | 11 | 15 | 3 | 1 | 3,6 | 2,9 | 0,7 |
| chr2:37431886-37432386:downstream_21813 | chr2 | 37423634 | 37431886 | + | CEBPZOS | NR | 23 | -0,5 | 1 | 0 | 6,79E-01 | NA | 21 | 7 | 3 | 10 | 54 | 49 | 24 | 13 | 1,1 | 1,5 | -0,4 |
| chr8:82568650-82569150:downstream_25965 | chr8 | 82569150 | 82598589 | - | IMPA1 | NM | 8 | 0,6 | 2 | 0 | 6,79E-01 | NA | 12 | 17 | 10 | 6 | 9 | 2 | 3 | 6 | 0,9 | 0,4 | 0,6 |
| chr3:53900593-53901093:downstream_3009 | chr3 | 53901093 | 53916229 | - | ACTR8 | NM | 18 | 0,5 | 1 | 0 | 6,79E-01 | NA | 11 | 13 | 11 | 17 | 7 | 29 | 33 | 27 | -0,3 | -0,8 | 0,5 |
| chr3:33189265-33189765:downstream_2600 | chr3 | 33155449 | 33189265 | + | CRTAP | NM | 39 | 0,3 | 1 | 0 | 6,79E-01 | 9,06E-01 | 15 | 29 | 21 | 21 | 52 | 53 | 61 | 62 | 0,1 | -0,2 | 0,3 |
| chrX:45707008-45707508:downstream_7687 | chrX | 45707508 | 45710920 | - | LOC401585 | NR | 34 | 0,4 | 1 | 0 | 6,79E-01 | 9,06E-01 | 7 | 23 | 6 | 10 | 54 | 78 | 40 | 53 | 0,9 | 0,5 | 0,5 |
| chr3:107879158-107879658:downstream_3090 | chr3 | 107879658 | 107941417 | - | IFT57 | NM | 4 | -0,8 | 2 | 0 | 6,80E-01 | NA | 4 | 8 | 6 | 4 | 2 | 7 | 3 | 2 | 0,3 | 1,1 | -0,8 |
| chr14:21967321-21967821:downstream_19424 | chr14 | 21945334 | 21967321 | + | TOX4 | NM | 26 | 0,4 | 1 | 0 | 6,80E-01 | NA | 28 | 23 | 22 | 29 | 33 | 15 | 35 | 26 | 0,0 | -0,4 | 0,4 |
| chr6:53425586-53426086:downstream_5201 | chr6 | 53426086 | 53438073 | - | LOC101927136 | NR | 14 | 0,5 | 1 | 0 | 6,80E-01 | NA | 2 | 10 | 6 | 7 | 14 | 20 | 33 | 21 | -0,1 | -0,7 | 0,6 |
| chr6:26016759-26017259:downstream_4906 | chr6 | 26017259 | 26018040 | - | HIST1H1A | NM | 7 | -1,0 | 3 | 0 | 6,81E-01 | NA | 7 | 1 | 0 | 0 | 30 | 18 | 1 | 0 | Inf | 5,1 | Inf |
| chr7:7647110-7647610:downstream_5709 | chr7 | 7606615 | 7647110 | + | MIOS | NM | 16 | 0,5 | 1 | 0 | 6,81E-01 | NA | 3 | 14 | 4 | 7 | 22 | 29 | 19 | 28 | 0,6 | 0,1 | 0,5 |
| chr10:89512374-89512874:downstream_8314 | chr10 | 89512874 | 89577917 | - | ATAD1 | NM | 166 | 0,3 | 1 | 0 | 6,81E-01 | 9,07E-01 | 11 | 5 | 10 | 14 | 246 | 201 | 449 | 395 | -0,6 | -0,9 | 0,3 |
| chr7:105096459-105096959:downstream_5601 | chr7 | 105096959 | 105162685 | - | PUS7 | NM | 9 | 0,6 | 1 | 0 | 6,81E-01 | NA | 2 | 8 | 7 | 11 | 10 | 3 | 15 | 19 | -0,8 | -1,4 | 0,6 |
| chr16:74340186-74340686:downstream_12469 | chr16 | 74330672 | 74340186 | + | PSMD7 | NM | 95 | 0,3 | 1 | 0 | 6,82E-01 | 9,07E-01 | 38 | 20 | 17 | 13 | 236 | 181 | 124 | 127 | 1,0 | 0,7 | 0,2 |
| chr4:140210570-140211070:downstream_23767 | chr4 | 140211070 | 140223705 | - | NDUFC1 | NM | 14 | -0,5 | 1 | 0 | 6,82E-01 | NA | 10 | 15 | 3 | 4 | 38 | 25 | 11 | 2 | 1,8 | 2,2 | -0,4 |
| chr7:155101945-155102445:downstream_25574 | chr7 | 155089485 | 155101945 | + | INSIG1 | NM | 37 | -0,5 | 1 | 0 | 6,82E-01 | 9,07E-01 | 2 | 0 | 13 | 5 | 15 | 36 | 100 | 126 | -2,8 | -2,1 | -0,7 |
| chr12:54058443-54058943:downstream_18449 | chr12 | 54058943 | 54070512 | - | ATP5G2 | NM | 159 | 0,3 | 1 | 0 | 6,83E-01 | 9,07E-01 | 29 | 13 | 19 | 13 | 301 | 328 | 225 | 344 | 0,4 | 0,1 | 0,2 |
| chr6:47594996-47595496:downstream_4736 | chr6 | 47445524 | 47594996 | + | CD2AP | NM | 11 | -0,6 | 1 | 0 | 6,83E-01 | NA | 2 | 3 | 19 | 16 | 3 | 5 | 22 | 15 | -2,7 | -2,2 | -0,6 |
| chr5:157166772-157167272:downstream_4545 | chr5 | 157158322 | 157166772 | + | THG1L | NM | 26 | 0,4 | 1 | 0 | 6,83E-01 | NA | 10 | 24 | 24 | 17 | 24 | 29 | 47 | 36 | -0,3 | -0,6 | 0,4 |
| chr20:25658708-25659208:downstream_22614 | chr20 | 25604680 | 25658708 | + | ZNF337-AS1 | NR | 11 | 0,5 | 1 | 0 | 6,83E-01 | NA | 12 | 11 | 8 | 8 | 9 | 14 | 15 | 8 | 0,5 | -0,1 | 0,5 |
| chr13:52336171-52336671:downstream_10601 | chr13 | 52158483 | 52336171 | + | WDFY2 | NM | 5 | 0,9 | 2 | 0 | 6,83E-01 | NA | 3 | 0 | 10 | 8 | 2 | 0 | 3 | 14 | -2,5 | -3,4 | 0,9 |
| chr17:58027786-58028286:downstream_20686 | chr17 | 57970406 | 58027786 | + | RPS6KB1 | NM | 16 | 0,5 | 1 | 0 | 6,84E-01 | NA | 13 | 0 | 11 | 8 | 20 | 10 | 32 | 31 | -0,6 | -1,1 | 0,5 |
| chr4:148558435-148558935:downstream_23809 | chr4 | 148558935 | 148605381 | - | PRMT9 | NM | 10 | -0,5 | 1 | 0 | 6,84E-01 | NA | 8 | 13 | 11 | 15 | 7 | 10 | 6 | 9 | -0,3 | 0,2 | -0,5 |
| chr19:57135544-57136044:downstream_14684 | chr19 | 57106663 | 57135544 | + | ZNF71 | NM | 5 | 0,7 | 2 | 0 | 6,84E-01 | NA | 5 | 8 | 6 | 5 | 4 | 5 | 4 | 7 | 0,3 | -0,4 | 0,6 |
| chr8:55061074-55061574:downstream_6297 | chr8 | 55047780 | 55061074 | + | MRPL15 | NM | 13 | 0,5 | 1 | 0 | 6,84E-01 | NA | 21 | 15 | 7 | 1 | 22 | 26 | 7 | 7 | 2,2 | 1,8 | 0,4 |
| chrX:57931363-57931863:downstream_7509 | chrX | 57931863 | 57937067 | - | ZXDA | NM | 6 | 0,8 | 2 | 0 | 6,84E-01 | NA | 17 | 6 | 3 | 1 | 10 | 8 | 3 | 2 | 2,7 | 2,0 | 0,7 |
| chr12:110477235-110477735:downstream_10419 | chr12 | 110437234 | 110477235 | + | ANKRD13A | NM | 9 | 0,6 | 1 | 0 | 6,84E-01 | NA | 8 | 3 | 7 | 3 | 7 | 15 | 8 | 21 | 0,1 | -0,4 | 0,5 |
| chr1:24086371-24086871:downstream_548 | chr1 | 24086871 | 24104787 | - | TCEB3-AS1 | NR | 6 | 0,7 | 2 | 0 | 6,85E-01 | NA | 2 | 6 | 4 | 14 | 2 | 2 | 8 | 10 | -1,3 | -2,1 | 0,8 |
| chr4:1381837-1382337:downstream_3571 | chr4 | 1341103 | 1381837 | + | UVSSA | NM | 7 | 0,7 | 2 | 0 | 6,86E-01 | NA | 8 | 5 | 1 | 7 | 11 | 4 | 6 | 11 | 0,6 | -0,2 | 0,7 |
| chr3:160212282-160212782:downstream_3261 | chr3 | 160212782 | 160283376 | - | KPNA4 | NM | 17 | 0,5 | 1 | 0 | 6,86E-01 | NA | 15 | 1 | 7 | 15 | 20 | 13 | 32 | 30 | -0,4 | -0,9 | 0,5 |
| chr12:123848810-123849310:downstream_10485 | chr12 | 123849310 | 123849390 | - | MIR8072 | NR | 8 | 0,6 | 1 | 0 | 6,86E-01 | NA | 11 | 7 | 18 | 8 | 5 | 2 | 8 | 7 | -0,5 | -1,1 | 0,6 |
| chr19:10420386-10420886:downstream_13992 | chr19 | 10420886 | 10426691 | - | FDX1L | NM | 21 | 0,5 | 1 | 0 | 6,86E-01 | NA | 19 | 2 | 7 | 3 | 52 | 32 | 31 | 25 | 1,0 | 0,6 | 0,4 |
| chr3:9428475-9428975:downstream_23469 | chr3 | 9404716 | 9428475 | + | THUMPD3 | NM | 20 | 0,4 | 1 | 0 | 6,86E-01 | NA | 14 | 8 | 7 | 7 | 25 | 40 | 32 | 25 | 0,6 | 0,2 | 0,4 |
| chr14:74946142-74946642:downstream_11227 | chr14 | 74946642 | 74960084 | - | NPC2 | NM | 20 | 0,5 | 1 | 0 | 6,87E-01 | NA | 2 | 12 | 3 | 2 | 37 | 62 | 13 | 30 | 1,7 | 1,2 | 0,5 |
| chr19:54972477-54972977:downstream_21229 | chr19 | 54972977 | 54974894 | - | LENG9 | NM | 18 | 0,4 | 1 | 0 | 6,87E-01 | NA | 19 | 37 | 21 | 24 | 6 | 14 | 11 | 10 | 0,3 | -0,1 | 0,4 |
| chr5:66296945-66297445:downstream_4280 | chr5 | 66297445 | 66299982 | - | LOC101928794 | NR | 10 | 0,6 | 2 | 0 | 6,87E-01 | NA | 15 | 6 | 17 | 5 | 14 | 2 | 21 | 5 | -0,1 | -0,7 | 0,6 |
| chr17:78108512-78109012:downstream_12702 | chr17 | 78109012 | 78120982 | - | EIF4A3 | NM | 21 | -0,4 | 1 | 0 | 6,87E-01 | NA | 14 | 10 | 6 | 9 | 38 | 51 | 10 | 31 | 0,7 | 1,1 | -0,4 |
| chrX:129262837-129263337:downstream_26604 | chrX | 129263337 | 129299861 | - | AIFM1 | NM | 10 | -0,6 | 1 | 0 | 6,87E-01 | NA | 3 | 8 | 4 | 4 | 10 | 28 | 11 | 8 | 0,4 | 1,0 | -0,6 |
| chr21:34732128-34732628:downstream_15295 | chr21 | 34697213 | 34732128 | + | IFNAR1 | NM | 16 | -0,5 | 1 | 0 | 6,87E-01 | NA | 10 | 4 | 21 | 15 | 22 | 6 | 11 | 39 | -1,4 | -0,8 | -0,5 |
| chr10:115672265-115672765:downstream_8728 | chr10 | 115614390 | 115672265 | + | NHLRC2 | NM | 63 | -0,3 | 1 | 0 | 6,87E-01 | 9,12E-01 | 47 | 40 | 170 | 116 | 17 | 19 | 43 | 55 | -1,7 | -1,4 | -0,3 |
| chr19:39423659-39424159:downstream_21291 | chr19 | 39421347 | 39423659 | + | MRPS12 | NM | 36 | 0,4 | 1 | 0 | 6,88E-01 | 9,12E-01 | 15 | 28 | 8 | 4 | 92 | 81 | 33 | 29 | 1,8 | 1,5 | 0,3 |
| chr12:123110947-123111447:downstream_9947 | chr12 | 123011808 | 123110947 | + | KNTC1 | NM | 7 | 0,7 | 2 | 0 | 6,88E-01 | NA | 1 | 2 | 8 | 10 | 1 | 2 | 10 | 21 | -2,7 | -3,5 | 0,7 |
| chr16:1752073-1752573:downstream_12093 | chr16 | 1728277 | 1752073 | + | HN1L | NM | 19 | 0,5 | 1 | 0 | 6,88E-01 | NA | 8 | 3 | 1 | 6 | 45 | 21 | 35 | 30 | 0,6 | 0,0 | 0,6 |
| chr9:96938715-96939215:downstream_7197 | chr9 | 96938628 | 96938715 | + | MIRLET7F1 | NR | 92 | -0,3 | 1 | 0 | 6,88E-01 | 9,12E-01 | 103 | 54 | 83 | 74 | 146 | 87 | 81 | 112 | 0,0 | 0,3 | -0,3 |
| chr3:48485537-48486037:downstream_2938 | chr3 | 48481685 | 48485537 | + | TMA7 | NM | 12 | 0,6 | 1 | 0 | 6,88E-01 | NA | 12 | 24 | 0 | 7 | 16 | 20 | 3 | 9 | 2,3 | 1,6 | 0,7 |
| chr11:58293843-58294343:downstream_18110 | chr11 | 58294343 | 58345712 | - | LPXN | NM | 4 | 0,9 | 2 | 0 | 6,89E-01 | NA | 4 | 5 | 1 | 0 | 4 | 16 | 1 | 3 | 2,7 | 2,1 | 0,5 |
| chr1:40759856-40760356:downstream_661 | chr1 | 40723721 | 40759856 | + | ZMPSTE24 | NM | 20 | 0,5 | 1 | 0 | 6,89E-01 | NA | 3 | 7 | 0 | 7 | 37 | 28 | 31 | 47 | 0,4 | -0,2 | 0,7 |
| chr12:89981325-89981825:downstream_18448 | chr12 | 89981825 | 90049844 | - | ATP2B1 | NM | 14 | 0,6 | 1 | 0 | 6,89E-01 | NA | 0 | 4 | 3 | 9 | 7 | 9 | 40 | 40 | -1,6 | -2,3 | 0,7 |
| chr17:40707232-40707732:downstream_13123 | chr17 | 40703983 | 40707232 | + | HSD17B1 | NM | 4 | 0,8 | 2 | 0 | 6,89E-01 | NA | 5 | 5 | 1 | 2 | 4 | 9 | 3 | 4 | 1,7 | 0,9 | 0,8 |
| chr17:77761449-77761949:downstream_20379 | chr17 | 77751976 | 77761449 | + | CBX2 | NM | 5 | -0,8 | 2 | 0 | 6,90E-01 | NA | 1 | 2 | 11 | 5 | 5 | 0 | 11 | 6 | -2,5 | -1,8 | -0,8 |
| chr8:92099323-92099823:downstream_26039 | chr8 | 92082423 | 92099323 | + | OTUD6B | NM | 23 | 0,4 | 1 | 0 | 6,90E-01 | NA | 15 | 8 | 8 | 7 | 29 | 46 | 26 | 42 | 0,5 | 0,1 | 0,4 |
| chr1:66840262-66840762:downstream_17061 | chr1 | 66258192 | 66840262 | + | PDE4B | NM | 17 | -0,6 | 1 | 0 | 6,90E-01 | NA | 1 | 2 | 19 | 11 | 8 | 3 | 51 | 36 | -3,5 | -3,0 | -0,5 |
| chr14:24908471-24908971:downstream_19375 | chr14 | 24908971 | 24912111 | - | SDR39U1 | NM | 13 | 0,5 | 1 | 0 | 6,90E-01 | NA | 23 | 14 | 7 | 5 | 23 | 16 | 8 | 9 | 1,6 | 1,2 | 0,4 |
| chr6:25987557-25988057:downstream_4902 | chr6 | 25962916 | 25987557 | + | TRIM38 | NM | 10 | 0,5 | 1 | 0 | 6,91E-01 | NA | 12 | 9 | 13 | 9 | 12 | 4 | 7 | 16 | 0,0 | -0,5 | 0,5 |
| chr18:57571538-57572038:downstream_13649 | chr18 | 57567191 | 57571538 | + | PMAIP1 | NM | 102 | 0,4 | 1 | 0 | 6,91E-01 | 9,14E-01 | 30 | 42 | 1 | 3 | 328 | 353 | 28 | 34 | 3,9 | 3,5 | 0,5 |
| chr5:176733960-176734460:downstream_24311 | chr5 | 176730762 | 176733960 | + | PRELID1 | NM | 23 | 0,4 | 1 | 0 | 6,91E-01 | NA | 8 | 16 | 3 | 3 | 44 | 71 | 17 | 22 | 2,0 | 1,6 | 0,5 |
| chr2:96078879-96079379:downstream_1669 | chr2 | 96068447 | 96078879 | + | FAHD2A | NM | 8 | 0,6 | 1 | 0 | 6,91E-01 | NA | 14 | 12 | 10 | 6 | 6 | 7 | 4 | 7 | 0,7 | 0,2 | 0,5 |
| chr17:76169659-76170159:downstream_13396 | chr17 | 76170159 | 76183285 | - | TK1 | NM | 39 | -0,4 | 1 | 0 | 6,92E-01 | 9,14E-01 | 4 | 5 | 7 | 4 | 84 | 71 | 49 | 91 | -0,3 | 0,1 | -0,5 |
| chr4:75254477-75254977:downstream_3459 | chr4 | 75230859 | 75254477 | + | EREG | NM | 30 | 0,3 | 1 | 0 | 6,92E-01 | 9,14E-01 | 24 | 11 | 11 | 10 | 47 | 57 | 39 | 41 | 0,7 | 0,4 | 0,3 |
| chr8:23048469-23048969:downstream_6229 | chr8 | 23048969 | 23082680 | - | TNFRSF10A | NM | 334 | 0,2 | 0 | 0 | 6,92E-01 | 9,14E-01 | 672 | 647 | 159 | 168 | 440 | 360 | 139 | 89 | 2,0 | 1,8 | 0,2 |
| chr2:86440477-86440977:downstream_22086 | chr2 | 86426555 | 86440477 | + | MRPL35 | NM | 12 | 0,5 | 1 | 0 | 6,92E-01 | NA | 20 | 16 | 11 | 9 | 12 | 12 | 11 | 7 | 0,8 | 0,4 | 0,5 |
| chr3:101367782-101368282:downstream_3081 | chr3 | 101368282 | 101395988 | - | ZBTB11 | NM | 18 | 0,4 | 1 | 0 | 6,92E-01 | NA | 7 | 10 | 3 | 7 | 23 | 43 | 21 | 30 | 0,9 | 0,4 | 0,5 |
| chr11:20530879-20531379:downstream_18231 | chr11 | 20409075 | 20530879 | + | PRMT3 | NM | 14 | -0,5 | 1 | 0 | 6,93E-01 | NA | 9 | 2 | 11 | 7 | 17 | 24 | 22 | 24 | -0,7 | -0,2 | -0,5 |
| chr1:166823709-166824209:downstream_1196 | chr1 | 166808723 | 166823709 | + | POGK | NM | 54 | 0,3 | 1 | 0 | 6,93E-01 | 9,14E-01 | 24 | 21 | 7 | 23 | 88 | 104 | 90 | 76 | 0,6 | 0,2 | 0,4 |
| chr1:213446808-213447308:downstream_17198 | chr1 | 213224574 | 213446808 | + | RPS6KC1 | NM | 15 | 0,6 | 2 | 0 | 6,93E-01 | NA | 0 | 2 | 10 | 14 | 3 | 1 | 47 | 43 | -3,6 | -4,4 | 0,8 |
| chr9:15170341-15170841:downstream_26558 | chr9 | 15170841 | 15307358 | - | TTC39B | NM | 8 | -0,7 | 2 | 0 | 6,93E-01 | NA | 3 | 0 | 18 | 9 | 6 | 0 | 7 | 21 | -3,1 | -2,3 | -0,8 |
| chr19:9405485-9405985:downstream_13976 | chr19 | 9405985 | 9415795 | - | ZNF699 | NM | 22 | 0,4 | 1 | 0 | 6,93E-01 | NA | 33 | 24 | 18 | 12 | 35 | 16 | 24 | 12 | 0,9 | 0,5 | 0,4 |
| chr12:108913380-108913880:downstream_10404 | chr12 | 108909050 | 108913380 | + | FICD | NM | 14 | 0,5 | 1 | 0 | 6,93E-01 | NA | 24 | 7 | 22 | 21 | 7 | 5 | 6 | 17 | -0,5 | -0,9 | 0,5 |
| chr3:122130199-122130699:downstream_3139 | chr3 | 122130699 | 122134882 | - | WDR5B | NM | 20 | -0,4 | 1 | 0 | 6,93E-01 | NA | 13 | 8 | 7 | 2 | 43 | 62 | 17 | 12 | 1,3 | 1,9 | -0,6 |
| chr5:167982127-167982627:downstream_4564 | chr5 | 167982627 | 168006614 | - | PANK3 | NM | 76 | -0,3 | 1 | 0 | 6,94E-01 | 9,15E-01 | 44 | 17 | 68 | 66 | 63 | 82 | 146 | 117 | -1,1 | -0,9 | -0,3 |
| chr1:207216693-207217193:downstream_17411 | chr1 | 207217193 | 207226325 | - | YOD1 | NM | 30 | 0,4 | 1 | 0 | 6,95E-01 | 9,16E-01 | 56 | 24 | 11 | 9 | 47 | 56 | 22 | 11 | 2,0 | 1,6 | 0,4 |
| chr1:38275126-38275626:downstream_16526 | chr1 | 38273472 | 38275126 | + | C1orf122 | NM | 23 | 0,5 | 1 | 0 | 6,95E-01 | NA | 13 | 0 | 3 | 3 | 44 | 54 | 26 | 37 | 1,1 | 0,6 | 0,5 |
| chr4:4236768-4237268:downstream_23888 | chr4 | 4237268 | 4249969 | - | TMEM128 | NM | 9 | 0,6 | 2 | 0 | 6,96E-01 | NA | 1 | 10 | 3 | 3 | 18 | 12 | 11 | 16 | 0,8 | 0,1 | 0,6 |
| chr4:113196281-113196781:downstream_3814 | chr4 | 113196781 | 113207059 | - | TIFA | NM | 8 | 0,6 | 2 | 0 | 6,96E-01 | NA | 11 | 4 | 14 | 11 | 7 | 0 | 8 | 11 | -0,7 | -1,4 | 0,6 |
| chr17:18176734-18177234:downstream_12876 | chr17 | 18177234 | 18218321 | - | TOP3A | NM | 22 | 0,4 | 1 | 0 | 6,97E-01 | NA | 20 | 26 | 24 | 25 | 14 | 21 | 26 | 21 | -0,1 | -0,4 | 0,4 |
| chr11:105878128-105878628:downstream_9624 | chr11 | 105878628 | 105892954 | - | MSANTD4 | NM | 12 | 0,5 | 1 | 0 | 6,97E-01 | NA | 29 | 8 | 8 | 4 | 15 | 17 | 8 | 7 | 1,6 | 1,1 | 0,5 |
| chr12:124105069-124105569:downstream_10490 | chr12 | 124105569 | 124118323 | - | EIF2B1 | NM | 57 | 0,3 | 1 | 0 | 6,97E-01 | 9,17E-01 | 14 | 15 | 4 | 8 | 136 | 130 | 60 | 91 | 1,2 | 0,8 | 0,4 |
| chr8:37700897-37701397:downstream_6527 | chr8 | 37701397 | 37707431 | - | BRF2 | NM | 23 | -0,4 | 1 | 0 | 6,97E-01 | NA | 8 | 15 | 7 | 7 | 51 | 51 | 25 | 21 | 0,8 | 1,2 | -0,4 |
| chr9:34609981-34610481:downstream_26485 | chr9 | 34610481 | 34612110 | - | RPP25L | NM | 36 | 0,4 | 1 | 0 | 6,97E-01 | 9,17E-01 | 9 | 26 | 7 | 7 | 71 | 92 | 36 | 44 | 1,4 | 1,0 | 0,4 |
| chr15:66841822-66842322:downstream_19853 | chr15 | 66797420 | 66841822 | + | ZWILCH | NM | 26 | 0,4 | 1 | 0 | 6,97E-01 | NA | 7 | 0 | 8 | 7 | 21 | 32 | 67 | 70 | -1,0 | -1,4 | 0,3 |
| chr2:158167913-158168413:downstream_2282 | chr2 | 158114339 | 158167913 | + | GALNT5 | NM | 23 | 0,4 | 1 | 0 | 6,98E-01 | NA | 23 | 33 | 43 | 38 | 6 | 10 | 17 | 12 | -0,5 | -0,9 | 0,4 |
| chr17:905390-905890:downstream_12722 | chr17 | 900356 | 905390 | + | TIMM22 | NM | 12 | 0,5 | 1 | 0 | 6,98E-01 | NA | 13 | 17 | 13 | 4 | 15 | 15 | 11 | 11 | 0,9 | 0,5 | 0,4 |
| chr8:48974454-48974954:downstream_6550 | chr8 | 48920994 | 48974454 | + | UBE2V2 | NM | 12 | 0,5 | 1 | 0 | 6,98E-01 | NA | 7 | 12 | 7 | 7 | 12 | 18 | 11 | 22 | 0,3 | -0,1 | 0,5 |
| chrX:118599495-118599995:downstream_7898 | chrX | 118599995 | 118603083 | - | SLC25A5-AS1 | NR | 5 | -0,8 | 2 | 0 | 6,98E-01 | NA | 5 | 9 | 1 | 2 | 11 | 10 | 4 | 0 | 1,8 | 2,3 | -0,5 |
| chr1:153746555-153747055:downstream_1068 | chr1 | 153700566 | 153746555 | + | INTS3 | NM | 13 | -0,6 | 2 | 0 | 6,98E-01 | NA | 2 | 0 | 4 | 7 | 13 | 5 | 35 | 39 | -2,7 | -2,1 | -0,7 |
| chr16:68054676-68055176:downstream_12430 | chr16 | 68055176 | 68057770 | - | DDX28 | NM | 19 | 0,4 | 1 | 0 | 6,99E-01 | NA | 19 | 15 | 10 | 15 | 29 | 17 | 17 | 29 | 0,4 | 0,0 | 0,4 |
| chr1:65522937-65523437:downstream_777 | chr1 | 65523437 | 65523525 | - | MIR3671 | NR | 10 | -0,5 | 1 | 0 | 6,99E-01 | NA | 7 | 5 | 22 | 10 | 7 | 7 | 7 | 16 | -1,4 | -0,8 | -0,6 |
| chr3:142719186-142719686:downstream_3215 | chr3 | 142719686 | 142720309 | - | LOC100289361 | NR | 30 | -0,4 | 1 | 0 | 7,00E-01 | 9,18E-01 | 30 | 21 | 11 | 7 | 66 | 69 | 11 | 23 | 1,5 | 2,0 | -0,4 |
| chr2:211295472-211295972:downstream_22017 | chr2 | 211295972 | 211341499 | - | LANCL1 | NM | 110 | -0,3 | 1 | 0 | 7,00E-01 | 9,18E-01 | 5 | 1 | 8 | 16 | 89 | 106 | 347 | 312 | -2,1 | -1,8 | -0,3 |
| chr1:18812480-18812980:downstream_506 | chr1 | 18807423 | 18812480 | + | KLHDC7A | NM | 10 | -0,6 | 1 | 0 | 7,00E-01 | NA | 7 | 2 | 4 | 2 | 32 | 13 | 10 | 11 | 0,5 | 1,1 | -0,6 |
| chr4:15692070-15692570:downstream_3629 | chr4 | 15683351 | 15692070 | + | FAM200B | NM | 14 | 0,6 | 1 | 0 | 7,00E-01 | NA | 5 | 8 | 3 | 2 | 15 | 47 | 6 | 25 | 1,5 | 1,0 | 0,5 |
| chr11:784284-784784:downstream_18107 | chr11 | 777577 | 784284 | + | LOC171391 | NR | 11 | 0,6 | 2 | 0 | 7,00E-01 | NA | 1 | 11 | 4 | 2 | 24 | 16 | 13 | 17 | 1,0 | 0,5 | 0,5 |
| chr3:5026865-5027365:downstream_2797 | chr3 | 5021096 | 5026865 | + | BHLHE40 | NM | 44 | 0,3 | 1 | 0 | 7,01E-01 | 9,18E-01 | 6 | 9 | 6 | 6 | 67 | 98 | 77 | 89 | 0,3 | 0,0 | 0,4 |
| chr12:69054385-69054885:downstream_18786 | chr12 | 69004618 | 69054385 | + | RAP1B | NM | 13 | -0,5 | 1 | 0 | 7,01E-01 | NA | 15 | 30 | 10 | 12 | 8 | 17 | 3 | 6 | 1,0 | 1,6 | -0,5 |
| chr20:43708593-43709093:downstream_14800 | chr20 | 43595119 | 43708593 | + | STK4 | NM | 16 | -0,5 | 1 | 0 | 7,01E-01 | NA | 4 | 3 | 10 | 8 | 12 | 24 | 26 | 40 | -1,4 | -0,9 | -0,5 |
| chr7:141354209-141354709:downstream_5645 | chr7 | 141251077 | 141354209 | + | AGK | NM | 11 | 0,6 | 2 | 0 | 7,01E-01 | NA | 5 | 1 | 17 | 16 | 5 | 0 | 25 | 16 | -2,5 | -3,1 | 0,6 |
| chr3:119387871-119388371:downstream_3129 | chr3 | 119388371 | 119396243 | - | COX17 | NM | 65 | -0,3 | 1 | 0 | 7,02E-01 | 9,18E-01 | 24 | 13 | 8 | 13 | 160 | 151 | 71 | 80 | 0,8 | 1,0 | -0,3 |
| chr22:39414825-39415325:downstream_15615 | chr22 | 39410264 | 39414825 | + | APOBEC3C | NM | 32 | 0,3 | 1 | 0 | 7,02E-01 | 9,18E-01 | 36 | 32 | 19 | 11 | 48 | 51 | 25 | 30 | 1,1 | 0,9 | 0,3 |
| chr4:39455244-39455744:downstream_23831 | chr4 | 39455744 | 39460568 | - | RPL9 | NM | 37 | -0,3 | 1 | 0 | 7,02E-01 | 9,18E-01 | 37 | 52 | 24 | 14 | 54 | 72 | 14 | 27 | 1,2 | 1,6 | -0,4 |
| chr14:67853199-67853699:downstream_11191 | chr14 | 67853699 | 67878828 | - | PLEK2 | NM | 8 | 0,7 | 2 | 0 | 7,02E-01 | NA | 18 | 7 | 3 | 1 | 6 | 23 | 3 | 3 | 2,8 | 2,3 | 0,5 |
| chr20:44516610-44517110:downstream_22484 | chr20 | 44517110 | 44519926 | - | NEURL2 | NM | 4 | -0,8 | 2 | 0 | 7,03E-01 | NA | 3 | 0 | 4 | 6 | 3 | 3 | 6 | 5 | -1,6 | -0,8 | -0,8 |
| chr17:42927154-42927654:downstream_20460 | chr17 | 42927654 | 42976993 | - | EFTUD2 | NM | 7 | -0,6 | 2 | 0 | 7,03E-01 | NA | 6 | 9 | 8 | 5 | 9 | 7 | 6 | 4 | 0,1 | 0,7 | -0,6 |
| chr2:133404169-133404669:downstream_1717 | chr2 | 133174146 | 133404169 | + | GPR39 | NM | 14 | -0,7 | 2 | 0 | 7,03E-01 | NA | 0 | 0 | 4 | 6 | 3 | 2 | 46 | 52 | -Inf | -4,2 | -Inf |
| chr8:15624158-15624658:downstream_26170 | chr8 | 15397595 | 15624158 | + | TUSC3 | NM | 7 | -0,7 | 2 | 0 | 7,03E-01 | NA | 2 | 0 | 7 | 7 | 9 | 0 | 15 | 16 | -2,6 | -1,8 | -0,8 |
| chr14:51185980-51186480:downstream_19321 | chr14 | 51186480 | 51297839 | - | NIN | NM | 22 | 0,5 | 1 | 0 | 7,04E-01 | NA | 2 | 1 | 7 | 7 | 7 | 19 | 56 | 81 | -2,0 | -2,4 | 0,4 |
| chr11:62441162-62441662:downstream_9395 | chr11 | 62439125 | 62441162 | + | UQCC3 | NM | 10 | 0,5 | 1 | 0 | 7,04E-01 | NA | 8 | 7 | 1 | 3 | 21 | 17 | 13 | 7 | 1,7 | 1,0 | 0,7 |
| chr9:32540041-32540541:downstream_26546 | chr9 | 32540541 | 32552626 | - | TOPORS | NM | 13 | -0,5 | 1 | 0 | 7,05E-01 | NA | 6 | 3 | 3 | 6 | 19 | 32 | 18 | 19 | 0,0 | 0,5 | -0,4 |
| chr6:26123194-26123694:downstream_4916 | chr6 | 26123694 | 26124132 | - | HIST1H2BC | NM | 209 | -0,2 | 1 | 0 | 7,05E-01 | 9,21E-01 | 12 | 4 | 35 | 36 | 207 | 136 | 555 | 686 | -2,1 | -1,9 | -0,3 |
| chr7:130146138-130146638:downstream_25630 | chr7 | 130126015 | 130146138 | + | MEST | NM | 34 | 0,4 | 1 | 0 | 7,05E-01 | 9,21E-01 | 7 | 4 | 7 | 2 | 60 | 68 | 61 | 65 | 0,2 | 0,0 | 0,2 |
| chr17:25950718-25951218:downstream_12619 | chr17 | 25799035 | 25950718 | + | KSR1 | NM | 17 | -0,4 | 1 | 0 | 7,05E-01 | NA | 3 | 4 | 22 | 13 | 11 | 9 | 36 | 34 | -2,3 | -1,9 | -0,4 |
| chr18:28645439-28645939:downstream_20857 | chr18 | 28645939 | 28682395 | - | DSC2 | NM | 9 | 0,7 | 2 | 0 | 7,06E-01 | NA | 2 | 0 | 6 | 5 | 6 | 0 | 33 | 25 | -2,7 | -3,3 | 0,7 |
| chr2:169721849-169722349:downstream_22121 | chr2 | 169643048 | 169721849 | + | NOSTRIN | NM | 11 | -0,6 | 2 | 0 | 7,06E-01 | NA | 11 | 0 | 21 | 9 | 15 | 2 | 15 | 14 | -1,4 | -0,8 | -0,6 |
| chr14:105565833-105566333:downstream_11394 | chr14 | 105560483 | 105565833 | + | LOC102723354 | NR | 7 | 0,7 | 2 | 0 | 7,07E-01 | NA | 10 | 2 | 3 | 2 | 14 | 11 | 3 | 11 | 1,4 | 0,9 | 0,5 |
| chr7:45022126-45022626:downstream_5817 | chr7 | 45022626 | 45026259 | - | SNHG15 | NR | 18 | 0,5 | 1 | 0 | 7,07E-01 | NA | 16 | 7 | 3 | 2 | 40 | 50 | 13 | 15 | 2,1 | 1,7 | 0,4 |
| chr4:73940001-73940501:downstream_23556 | chr4 | 73940501 | 74124502 | - | ANKRD17 | NM | 9 | 0,7 | 2 | 0 | 7,07E-01 | NA | 1 | 1 | 6 | 14 | 2 | 0 | 24 | 24 | -3,5 | -4,3 | 0,8 |
| chr16:57505921-57506421:downstream_12387 | chr16 | 57496550 | 57505921 | + | POLR2C | NM | 115 | 0,3 | 1 | 0 | 7,07E-01 | 9,23E-01 | 21 | 10 | 19 | 11 | 200 | 206 | 237 | 213 | 0,0 | -0,1 | 0,2 |
| chr1:155289750-155290250:downstream_1095 | chr1 | 155290250 | 155293938 | - | RUSC1-AS1 | NM | 7 | 0,6 | 2 | 0 | 7,08E-01 | NA | 8 | 4 | 7 | 8 | 7 | 2 | 13 | 6 | -0,3 | -1,0 | 0,6 |
| chr12:9811010-9811510:downstream_18658 | chr12 | 9769879 | 9811010 | + | LOC374443 | NR | 11 | 0,5 | 1 | 0 | 7,08E-01 | NA | 4 | 5 | 6 | 11 | 12 | 6 | 26 | 21 | -0,9 | -1,4 | 0,5 |
| chr17:29225500-29226000:downstream_12623 | chr17 | 29226000 | 29233286 | - | TEFM | NM | 22 | 0,4 | 1 | 0 | 7,08E-01 | NA | 8 | 7 | 8 | 5 | 22 | 48 | 43 | 36 | 0,2 | -0,2 | 0,3 |
| chrX:6966460-6966960:downstream_26850 | chrX | 6966960 | 7066231 | - | HDHD1 | NM | 15 | 0,5 | 1 | 0 | 7,08E-01 | NA | 0 | 7 | 19 | 12 | 9 | 1 | 33 | 38 | -2,2 | -2,8 | 0,6 |
| chr20:32579793-32580293:downstream_22526 | chr20 | 32580293 | 32582173 | - | RALY-AS1 | NR | 11 | 0,5 | 1 | 0 | 7,09E-01 | NA | 24 | 11 | 7 | 6 | 9 | 19 | 1 | 12 | 1,5 | 1,0 | 0,4 |
| chr1:242053241-242053741:downstream_16710 | chr1 | 242011492 | 242053241 | + | EXO1 | NM | 14 | 0,5 | 1 | 0 | 7,09E-01 | NA | 13 | 0 | 21 | 24 | 4 | 5 | 18 | 25 | -1,8 | -2,3 | 0,5 |
| chr1:89317820-89318320:downstream_815 | chr1 | 89318320 | 89357301 | - | GTF2B | NM | 17 | -0,4 | 1 | 0 | 7,09E-01 | NA | 9 | 11 | 10 | 4 | 27 | 44 | 17 | 19 | 0,5 | 1,0 | -0,5 |

| | | | | | | | | | | | | | | | | | | | | | | | |
|---|---|---|---|---|---|---|---|---|---|---|---|---|---|---|---|---|---|---|---|---|---|---|---|
| chr16:19091417-19091917:downstream_19951 | chr16 | 19078916 | 19091417 | + | COQ7 | NM | 10 | 0,6 | 2 | 0 | 7,09E-01 | NA | 7 | 2 | 6 | 8 | 15 | 1 | 21 | 16 | -0,6 | -1,2 | 0,6 |
| chr12:49081740-49082240:downstream_18487 | chr12 | 49082240 | 49110781 | - | CCNT1 | NM | 68 | 0,2 | 1 | 0 | 7,09E-01 | 9,24E-01 | 31 | 22 | 19 | 15 | 128 | 134 | 99 | 98 | 0,6 | 0,4 | 0,2 |
| chr2:3591981-3592481:downstream_22191 | chr2 | 3592481 | 3605961 | - | RNASEH1 | NM | 52 | 0,3 | 1 | 0 | 7,10E-01 | 9,24E-01 | 57 | 98 | 38 | 34 | 59 | 65 | 35 | 34 | 1,1 | 0,9 | 0,3 |
| chr11:57408759-57409259:downstream_9318 | chr11 | 57408670 | 57408759 | + | MIR130A | NR | 18 | -0,4 | 1 | 0 | 7,10E-01 | NA | 18 | 7 | 22 | 25 | 26 | 5 | 25 | 20 | -1,0 | -0,5 | -0,4 |
| chr18:213739-214239:downstream_20969 | chr18 | 158482 | 213739 | + | USP14 | NM | 35 | -0,3 | 1 | 0 | 7,10E-01 | 9,24E-01 | 28 | 20 | 53 | 59 | 30 | 11 | 28 | 47 | -1,2 | -0,9 | -0,3 |
| chr17:4574178-4574678:downstream_20627 | chr17 | 4574678 | 4607632 | - | PELP1 | NM | 11 | 0,5 | 1 | 0 | 7,11E-01 | NA | 5 | 15 | 6 | 3 | 17 | 22 | 8 | 15 | 1,2 | 0,8 | 0,4 |
| chr17:36891858-36892358:downstream_13017 | chr17 | 36886509 | 36891858 | + | CISD3 | NM | 6 | -0,6 | 2 | 0 | 7,11E-01 | NA | 5 | 6 | 8 | 7 | 4 | 6 | 6 | 4 | -0,6 | 0,0 | -0,6 |
| chr9:37503694-37504194:downstream_26450 | chr9 | 37485931 | 37503694 | + | POLR1E | NM | 65 | 0,2 | 1 | 0 | 7,11E-01 | 9,24E-01 | 71 | 44 | 46 | 30 | 103 | 82 | 72 | 72 | 0,6 | 0,4 | 0,2 |
| chr15:99188594-99189094:downstream_11914 | chr15 | 99189094 | 99194450 | - | IRAIN | NR | 21 | 0,4 | 1 | 0 | 7,11E-01 | NA | 42 | 34 | 7 | 5 | 35 | 32 | 11 | 3 | 2,7 | 2,2 | 0,5 |
| chr9:88902147-88902647:downstream_26577 | chr9 | 88902647 | 88969402 | - | ZCCHC6 | NM | 18 | 0,5 | 1 | 0 | 7,12E-01 | NA | 5 | 1 | 13 | 2 | 17 | 15 | 50 | 45 | -1,3 | -1,6 | 0,3 |
| chr5:61683011-61683511:downstream_24187 | chr5 | 61601988 | 61683011 | + | KIF2A | NM | 48 | 0,3 | 1 | 0 | 7,12E-01 | 9,24E-01 | 20 | 40 | 141 | 53 | 13 | 14 | 54 | 52 | -1,7 | -2,0 | 0,3 |
| chr12:95703760-95704260:downstream_10362 | chr12 | 95703698 | 95703760 | + | MIR3685 | NR | 31 | 0,4 | 1 | 0 | 7,12E-01 | 9,24E-01 | 10 | 1 | 19 | 33 | 7 | 19 | 82 | 78 | -2,3 | -2,6 | 0,3 |
| chr10:27292544-27293044:downstream_17470 | chr10 | 27293044 | 27389427 | - | ANKRD26 | NM | 4 | -0,9 | 2 | 0 | 7,13E-01 | NA | 2 | 0 | 0 | 10 | 5 | 1 | 6 | 7 | -2,0 | -1,1 | -0,9 |
| chr8:419875-420375:downstream_25939 | chr8 | 356807 | 419875 | + | FBXO25 | NM | 9 | -0,5 | 1 | 0 | 7,13E-01 | NA | 4 | 8 | 10 | 14 | 7 | 10 | 11 | 12 | -1,0 | -0,5 | -0,5 |
| chr14:23569665-23570165:downstream_11059 | chr14 | 23564682 | 23569665 | + | C14orf119 | NM | 11 | 0,5 | 1 | 0 | 7,13E-01 | NA | 20 | 17 | 11 | 9 | 7 | 9 | 7 | 5 | 0,9 | 0,4 | 0,5 |
| chr5:127356743-127357243:downstream_4083 | chr5 | 127357243 | 127418766 | - | LINC01184 | NR | 17 | 0,4 | 1 | 0 | 7,13E-01 | NA | 6 | 6 | 11 | 21 | 9 | 11 | 36 | 36 | -1,5 | -1,9 | 0,4 |
| chr5:170837888-170838388:downstream_24254 | chr5 | 170814707 | 170837888 | + | NPM1 | NM | 43 | 0,3 | 1 | 0 | 7,13E-01 | 9,24E-01 | 7 | 10 | 4 | 6 | 71 | 106 | 68 | 73 | 0,7 | 0,3 | 0,4 |
| chr2:91824208-91824708:downstream_2106 | chr2 | 91824708 | 91847975 | - | LOC654342 | NR | 5 | 0,7 | 2 | 0 | 7,13E-01 | NA | 2 | 7 | 3 | 4 | 9 | 1 | 4 | 8 | 0,4 | -0,3 | 0,7 |
| chr10:60591197-60591697:downstream_8283 | chr10 | 60272903 | 60591197 | + | BICC1 | NM | 56 | -0,4 | 1 | 0 | 7,14E-01 | 9,24E-01 | 1 | 4 | 11 | 10 | 54 | 38 | 178 | 155 | -2,2 | -1,9 | -0,3 |
| chr2:153507606-153508106:downstream_2278 | chr2 | 153508106 | 153573975 | - | PRPF40A | NM | 11 | 0,5 | 1 | 0 | 7,14E-01 | NA | 8 | 21 | 11 | 13 | 5 | 11 | 10 | 8 | 0,3 | -0,2 | 0,5 |
| chr2:58387055-58387555:downstream_22318 | chr2 | 58134785 | 58387055 | + | VRK2 | NM | 34 | 0,3 | 1 | 0 | 7,14E-01 | 9,24E-01 | 11 | 15 | 39 | 36 | 19 | 20 | 75 | 59 | -1,5 | -1,8 | 0,3 |
| chr3:133540336-133540836:downstream_3192 | chr3 | 133502876 | 133540336 | + | SRPRB | NM | 21 | -0,4 | 1 | 0 | 7,14E-01 | NA | 6 | 4 | 1 | 6 | 40 | 53 | 28 | 29 | 0,4 | 0,7 | -0,3 |
| chr5:2745778-2746278:downstream_24176 | chr5 | 2746278 | 2751769 | - | IRX2 | NM | 5 | -0,7 | 2 | 0 | 7,14E-01 | NA | 6 | 11 | 1 | 4 | 12 | 5 | 0 | 3 | 1,6 | 2,4 | -0,8 |
| chr22:29655343-29655843:downstream_15548 | chr22 | 29655843 | 29663914 | - | RHBDD3 | NM | 8 | 0,7 | 2 | 0 | 7,15E-01 | NA | 2 | 15 | 6 | 0 | 6 | 23 | 10 | 4 | 1,6 | 1,1 | 0,5 |
| chr20:25593072-25593572:downstream_14933 | chr20 | 25593572 | 25604648 | - | NANP | NM | 14 | 0,5 | 1 | 0 | 7,15E-01 | NA | 31 | 12 | 6 | 2 | 28 | 22 | 3 | 9 | 2,4 | 2,1 | 0,3 |
| chr10:31604956-31605456:downstream_8469 | chr10 | 31605456 | 31608024 | - | ZEB1-AS1 | NR | 7 | 0,6 | 2 | 0 | 7,15E-01 | NA | 2 | 10 | 3 | 2 | 17 | 6 | 11 | 6 | 1,1 | 0,4 | 0,7 |
| chr7:97595407-97595907:downstream_5958 | chr7 | 97595907 | 97601638 | - | MGC72080 | NR | 13 | 0,5 | 1 | 0 | 7,15E-01 | NA | 16 | 3 | 22 | 15 | 11 | 2 | 17 | 17 | -1,0 | -1,4 | 0,5 |
| chr5:470124-470624:downstream_4149 | chr5 | 470624 | 473080 | - | PP7080 | NR | 9 | -0,5 | 1 | 0 | 7,15E-01 | NA | 9 | 1 | 3 | 4 | 23 | 17 | 8 | 11 | 0,5 | 1,1 | -0,6 |
| chr3:156390459-156390959:downstream_3250 | chr3 | 156390959 | 156393502 | - | TIPARP-AS1 | NR | 21 | -0,4 | 1 | 0 | 7,15E-01 | NA | 40 | 27 | 6 | 7 | 41 | 40 | 0 | 10 | 2,5 | 3,0 | -0,6 |
| chrX:74272506-74273006:downstream_26599 | chrX | 74273006 | 74376175 | - | ABCB7 | NM | 23 | 0,4 | 1 | 0 | 7,16E-01 | NA | 4 | 4 | 8 | 6 | 26 | 23 | 54 | 57 | -0,8 | -1,2 | 0,3 |
| chr11:44266980-44267480:downstream_18010 | chr11 | 44117098 | 44266980 | + | EXT2 | NM | 43 | 0,3 | 1 | 0 | 7,16E-01 | 9,26E-01 | 25 | 22 | 86 | 54 | 25 | 9 | 61 | 62 | -1,6 | -1,9 | 0,3 |
| chr2:85831875-85832375:downstream_2080 | chr2 | 85832375 | 85839179 | - | C2orf68 | NM | 21 | -0,4 | 1 | 0 | 7,16E-01 | NA | 9 | 4 | 8 | 7 | 34 | 42 | 38 | 29 | -0,2 | 0,2 | -0,4 |
| chr16:85840607-85841107:downstream_12511 | chr16 | 85833172 | 85840607 | + | COX4I1 | NM | 243 | 0,3 | 1 | 0 | 7,16E-01 | 9,26E-01 | 2 | 9 | 3 | 3 | 516 | 566 | 431 | 416 | 0,8 | 0,4 | 0,4 |
| chrX:56844004-56844504:downstream_7760 | chrX | 56755717 | 56844004 | + | LINC01420 | NR | 48 | -0,3 | 1 | 0 | 7,17E-01 | 9,26E-01 | 21 | 29 | 15 | 20 | 82 | 104 | 57 | 53 | 0,5 | 0,8 | -0,2 |
| chr11:63341443-63341943:downstream_18212 | chr11 | 63341943 | 63381941 | - | PLA2G16 | NM | 27 | -0,4 | 1 | 0 | 7,17E-01 | NA | 7 | 2 | 4 | 7 | 42 | 58 | 46 | 52 | -0,3 | 0,0 | -0,4 |
| chr16:4511177-4511677:downstream_20144 | chr16 | 4511677 | 4524896 | - | NMRAL1 | NM | 12 | 0,5 | 1 | 0 | 7,18E-01 | NA | 14 | 1 | 31 | 11 | 7 | 2 | 22 | 12 | -1,5 | -2,0 | 0,5 |
| chr12:69136473-69136973:downstream_10310 | chr12 | 69080730 | 69136473 | + | NUP107 | NM | 12 | -0,6 | 2 | 0 | 7,18E-01 | NA | 2 | 0 | 14 | 12 | 6 | 3 | 13 | 46 | -3,4 | -2,8 | -0,7 |
| chr2:191367041-191367541:downstream_1762 | chr2 | 191273080 | 191367041 | + | MFSD6 | NM | 11 | -0,7 | 2 | 0 | 7,19E-01 | NA | 1 | 0 | 6 | 7 | 7 | 1 | 28 | 37 | -3,9 | -3,1 | -0,8 |
| chr11:74553458-74553958:downstream_9533 | chr11 | 74459912 | 74553458 | + | RNF169 | NM | 13 | 0,5 | 1 | 0 | 7,19E-01 | NA | 1 | 5 | 7 | 9 | 7 | 9 | 32 | 34 | -1,5 | -2,0 | 0,5 |
| chr17:61896677-61897177:downstream_20444 | chr17 | 61851548 | 61896677 | + | DDX42 | NM | 54 | -0,3 | 1 | 0 | 7,19E-01 | 9,28E-01 | 2 | 12 | 8 | 18 | 68 | 81 | 107 | 135 | -0,9 | -0,7 | -0,2 |
| chr17:3566397-3566897:downstream_20435 | chr17 | 3539761 | 3566397 | + | CTNS | NM | 7 | -0,7 | 2 | 0 | 7,20E-01 | NA | 7 | 6 | 3 | 1 | 7 | 26 | 4 | 2 | 1,8 | 2,5 | -0,7 |
| chr9:68415388-68415888:downstream_7118 | chr9 | 68415307 | 68415388 | + | MIR4477B | NR | 11 | -0,7 | 2 | 0 | 7,20E-01 | NA | 33 | 30 | 6 | 2 | 7 | 8 | 1 | 0 | 3,0 | 3,4 | -0,4 |
| chr2:43449040-43449540:downstream_1953 | chr2 | 43449540 | 43453745 | - | ZFP36L2 | NM | 46 | 0,3 | 1 | 0 | 7,20E-01 | 9,28E-01 | 4 | 15 | 15 | 12 | 61 | 50 | 114 | 93 | -0,6 | -0,9 | 0,3 |
| chr12:132610885-132611385:downstream_10521 | chr12 | 132568827 | 132610885 | + | EP400NL | NR | 5 | -0,9 | 2 | 0 | 7,20E-01 | NA | 11 | 0 | 7 | 2 | 2 | 10 | 6 | 0 | 0,3 | 1,1 | -0,8 |
| chr1:15756839-15757339:downstream_477 | chr1 | 15736390 | 15756839 | + | EFHD2 | NM | 15 | -0,4 | 1 | 0 | 7,20E-01 | NA | 2 | 4 | 11 | 5 | 20 | 15 | 39 | 24 | -1,3 | -0,9 | -0,5 |
| chr10:54077417-54077917:downstream_8525 | chr10 | 54074040 | 54077417 | + | DKK1 | NM | 484 | -0,2 | 0 | 0 | 7,20E-01 | 9,28E-01 | 349 | 261 | 344 | 301 | 658 | 692 | 743 | 527 | -0,1 | 0,1 | -0,2 |
| chr16:21263750-21264250:downstream_12242 | chr16 | 21245015 | 21263750 | + | ANKS4B | NM | 21 | -0,4 | 1 | 0 | 7,21E-01 | NA | 2 | 2 | 25 | 22 | 5 | 8 | 57 | 43 | -3,4 | -3,0 | -0,4 |
| chr6:139692891-139693391:downstream_24563 | chr6 | 139693391 | 139695787 | - | CITED2 | NM | 11 | 0,4 | 1 | 0 | 7,21E-01 | NA | 12 | 15 | 7 | 7 | 17 | 11 | 11 | 8 | 1,0 | 0,5 | 0,5 |
| chr2:128461407-128461907:downstream_2228 | chr2 | 128458596 | 128461407 | + | SFT2D3 | NM | 38 | -0,3 | 1 | 0 | 7,21E-01 | 9,28E-01 | 28 | 18 | 17 | 16 | 60 | 83 | 47 | 35 | 0,5 | 0,8 | -0,3 |
| chr4:38703129-38703629:downstream_3421 | chr4 | 38665789 | 38703129 | + | KLF3 | NM | 8 | 0,5 | 1 | 0 | 7,21E-01 | NA | 10 | 7 | 4 | 14 | 7 | 4 | 7 | 10 | -0,1 | -0,7 | 0,6 |
| chr1:3696283-3696783:downstream_395 | chr1 | 3696783 | 3713068 | - | LRRC47 | NM | 19 | 0,4 | 1 | 0 | 7,22E-01 | NA | 8 | 14 | 6 | 5 | 47 | 29 | 14 | 34 | 1,0 | 0,7 | 0,4 |
| chr15:78887611-78888111:downstream_19508 | chr15 | 78857861 | 78887611 | + | CHRNA5 | NM | 4 | -0,8 | 2 | 0 | 7,22E-01 | NA | 3 | 0 | 7 | 4 | 5 | 1 | 6 | 6 | -1,8 | -1,0 | -0,8 |
| chr18:3256235-3256735:downstream_20900 | chr18 | 3247479 | 3256235 | + | MYL12A | NM | 233 | 0,3 | 1 | 0 | 7,22E-01 | 9,28E-01 | 8 | 7 | 7 | 2 | 574 | 518 | 320 | 430 | 0,7 | 0,5 | 0,1 |
| chr12:75890918-75891418:downstream_10325 | chr12 | 75891418 | 75905418 | - | KRR1 | NM | 89 | -0,2 | 1 | 0 | 7,22E-01 | 9,28E-01 | 142 | 103 | 110 | 95 | 97 | 56 | 61 | 49 | 0,3 | 0,5 | -0,2 |
| chr20:6020697-6021197:downstream_22388 | chr20 | 5986738 | 6020697 | + | CRLS1 | NM | 12 | 0,5 | 1 | 0 | 7,22E-01 | NA | 14 | 8 | 3 | 7 | 28 | 10 | 10 | 14 | 1,2 | 0,7 | 0,5 |
| chr1:45809650-45810150:downstream_703 | chr1 | 45805341 | 45809650 | + | TOE1 | NM | 16 | 0,4 | 1 | 0 | 7,23E-01 | NA | 8 | 4 | 4 | 10 | 21 | 18 | 39 | 25 | -0,2 | -0,7 | 0,5 |
| chr16:46835458-46835958:downstream_12340 | chr16 | 46835958 | 46865074 | - | C16orf87 | NM | 19 | 0,4 | 1 | 0 | 7,23E-01 | NA | 8 | 5 | 3 | 8 | 24 | 37 | 49 | 23 | 0,2 | -0,2 | 0,5 |
| chr4:99365012-99365512:downstream_23815 | chr4 | 99182526 | 99365012 | + | RAP1GDS1 | NM | 7 | -0,6 | 2 | 0 | 7,23E-01 | NA | 2 | 0 | 14 | 4 | 4 | 3 | 15 | 16 | -2,9 | -2,1 | -0,7 |
| chr16:56511407-56511907:downstream_12002 | chr16 | 56485423 | 56511407 | + | OGFOD1 | NM | 26 | 0,3 | 1 | 0 | 7,24E-01 | NA | 38 | 42 | 15 | 10 | 39 | 34 | 18 | 11 | 1,7 | 1,3 | 0,3 |
| chr9:140509283-140509783:downstream_26218 | chr9 | 140509783 | 140513347 | - | ARRDC1-AS1 | NR | 18 | 0,4 | 1 | 0 | 7,24E-01 | NA | 46 | 42 | 13 | 14 | 13 | 7 | 3 | 5 | 1,7 | 1,4 | 0,4 |
| chr7:6061377-6061877:downstream_25496 | chr7 | 6061877 | 6098860 | - | EIF2AK1 | NM | 7 | 0,6 | 2 | 0 | 7,24E-01 | NA | 6 | 9 | 13 | 7 | 2 | 5 | 3 | 11 | -0,4 | -0,9 | 0,5 |
| chr18:45358965-45359465:downstream_20944 | chr18 | 45359465 | 45457517 | - | SMAD2 | NM | 19 | -0,4 | 1 | 0 | 7,25E-01 | NA | 7 | 6 | 31 | 37 | 12 | 2 | 26 | 34 | -2,5 | -2,1 | -0,4 |
| chr3:51697612-51698112:downstream_2628 | chr3 | 51575595 | 51697612 | + | RAD54L2 | NM | 6 | -0,7 | 2 | 0 | 7,25E-01 | NA | 0 | 4 | 7 | 8 | 4 | 3 | 8 | 10 | -2,0 | -1,4 | -0,6 |
| chr5:149372185-149372685:downstream_24412 | chr5 | 149372685 | 149380730 | - | TIGD6 | NM | 14 | 0,5 | 1 | 0 | 7,25E-01 | NA | 16 | 8 | 1 | 2 | 37 | 39 | 7 | 7 | 3,0 | 2,5 | 0,5 |
| chr11:6620643-6621143:downstream_9102 | chr11 | 6621143 | 6624880 | - | RRP8 | NM | 19 | -0,4 | 1 | 0 | 7,25E-01 | NA | 22 | 12 | 7 | 11 | 41 | 31 | 13 | 17 | 0,9 | 1,3 | -0,4 |
| chr3:111837073-111837573:downstream_23035 | chr3 | 111805181 | 111837073 | + | C3orf52 | NM | 51 | 0,3 | 1 | 0 | 7,25E-01 | 9,29E-01 | 19 | 22 | 6 | 3 | 121 | 164 | 50 | 25 | 2,2 | 1,9 | 0,3 |
| chr16:30361586-30362086:downstream_19914 | chr16 | 30362086 | 30366682 | - | CD2BP2 | NM | 18 | 0,4 | 1 | 0 | 7,25E-01 | NA | 5 | 8 | 4 | 2 | 45 | 35 | 24 | 22 | 1,1 | 0,8 | 0,3 |
| chr19:10596295-10596795:downstream_21203 | chr19 | 10596795 | 10614054 | - | KEAP1 | NM | 10 | 0,5 | 1 | 0 | 7,25E-01 | NA | 4 | 5 | 4 | 5 | 16 | 11 | 15 | 23 | 0,0 | -0,5 | 0,5 |
| chrX:102193228-102193728:downstream_7855 | chrX | 102192199 | 102193228 | + | RAB40AL | NM | 6 | 0,7 | 2 | 0 | 7,26E-01 | NA | 7 | 0 | 17 | 13 | 2 | 0 | 4 | 8 | -2,2 | -2,9 | 0,7 |
| chr9:79014954-79015454:downstream_7145 | chr9 | 79013514 | 79014954 | + | RPSAP9 | NR | 11 | -0,6 | 2 | 0 | 7,26E-01 | NA | 22 | 26 | 8 | 2 | 20 | 5 | 4 | 0 | 2,2 | 2,5 | -0,4 |
| chr11:6505911-6506411:downstream_9099 | chr11 | 6502676 | 6505911 | + | TIMM10B | NM | 23 | 0,3 | 1 | 0 | 7,26E-01 | NA | 36 | 32 | 18 | 16 | 16 | 32 | 14 | 16 | 1,0 | 0,7 | 0,3 |
| chr7:91740962-91741462:downstream_25476 | chr7 | 91741462 | 91764059 | - | CYP51A1 | NM | 167 | 0,3 | 1 | 0 | 7,26E-01 | 9,29E-01 | 6 | 9 | 11 | 8 | 207 | 298 | 404 | 392 | -0,4 | -0,7 | 0,2 |
| chr19:15347800-15348300:downstream_21027 | chr19 | 15348300 | 15391262 | - | BRD4 | NM | 10 | 0,5 | 2 | 0 | 7,26E-01 | NA | 2 | 9 | 8 | 17 | 8 | 1 | 13 | 19 | -1,2 | -1,8 | 0,6 |
| chr4:170644338-170644838:downstream_23592 | chr4 | 170541671 | 170644338 | + | CLCN3 | NM | 8 | -0,6 | 2 | 0 | 7,26E-01 | NA | 2 | 0 | 11 | 3 | 3 | 8 | 21 | 18 | -2,6 | -1,8 | -0,7 |
| chr14:31089267-31089767:downstream_19234 | chr14 | 31028328 | 31089267 | + | G2E3 | NM | 11 | 0,6 | 2 | 0 | 7,26E-01 | NA | 16 | 0 | 14 | 16 | 6 | 5 | 8 | 20 | -0,9 | -1,4 | 0,5 |
| chr7:100031749-100032249:downstream_25629 | chr7 | 100026412 | 100031749 | + | MEPCE | NM | 20 | 0,5 | 1 | 0 | 7,26E-01 | NA | 7 | 2 | 1 | 2 | 35 | 57 | 22 | 36 | 1,1 | 0,7 | 0,5 |
| chr2:241503431-241503931:downstream_2513 | chr2 | 241499470 | 241503431 | + | DUSP28 | NM | 9 | 0,5 | 2 | 0 | 7,27E-01 | NA | 9 | 3 | 1 | 2 | 20 | 19 | 8 | 11 | 1,6 | 1,1 | 0,6 |
| chr22:22867560-22868060:downstream_15504 | chr22 | 22868060 | 22874624 | - | ZNF280A | NM | 10 | 0,6 | 2 | 0 | 7,27E-01 | NA | 5 | 10 | 4 | 0 | 32 | 12 | 8 | 7 | 1,8 | 1,5 | 0,3 |
| chr12:56516280-56516780:downstream_18922 | chr12 | 56512003 | 56516280 | + | ZC3H10 | NM | 15 | 0,4 | 1 | 0 | 7,27E-01 | NA | 30 | 18 | 13 | 13 | 16 | 12 | 8 | 11 | 0,9 | 0,6 | 0,4 |
| chrX:47001114-47001614:downstream_26938 | chrX | 47001614 | 47004609 | - | NDUFB11 | NM | 95 | 0,4 | 1 | 0 | 7,28E-01 | 9,29E-01 | 6 | 3 | 4 | 1 | 275 | 185 | 129 | 157 | 0,8 | 0,7 | 0,1 |
| chr17:37884908-37885408:downstream_13035 | chr17 | 37885408 | 37886788 | - | MIEN1 | NM | 26 | -0,3 | 1 | 0 | 7,28E-01 | NA | 12 | 6 | 6 | 5 | 70 | 56 | 31 | 26 | 0,8 | 1,1 | -0,4 |
| chr5:133491581-133492081:downstream_24359 | chr5 | 133492081 | 133512724 | - | SKP1 | NM | 25 | 0,3 | 1 | 0 | 7,28E-01 | NA | 32 | 28 | 17 | 5 | 37 | 45 | 15 | 20 | 1,5 | 1,2 | 0,2 |
| chr10:115881473-115881973:downstream_8730 | chr10 | 115881973 | 115934364 | - | CCDC186 | NM | 42 | -0,3 | 1 | 0 | 7,28E-01 | 9,29E-01 | 33 | 16 | 60 | 56 | 34 | 23 | 64 | 47 | -1,2 | -0,9 | -0,3 |
| chr14:89079853-89080353:downstream_19443 | chr14 | 89029252 | 89079853 | + | ZC3H14 | NM | 34 | -0,3 | 1 | 0 | 7,29E-01 | 9,29E-01 | 20 | 10 | 57 | 62 | 24 | 6 | 42 | 54 | -2,0 | -1,7 | -0,3 |
| chr4:32157028-32157528:downstream_3416 | chr4 | 31999000 | 32157028 | + | LOC102723828 | NR | 5 | -0,7 | 2 | 0 | 7,29E-01 | NA | 1 | 3 | 10 | 10 | 2 | 1 | 7 | 5 | -2,4 | -1,8 | -0,6 |
| chr11:82784754-82785254:downstream_9554 | chr11 | 82783107 | 82784754 | + | RAB30-AS1 | NR | 77 | -0,2 | 1 | 0 | 7,29E-01 | 9,29E-01 | 85 | 64 | 68 | 56 | 110 | 92 | 77 | 69 | 0,3 | 0,5 | -0,2 |
| chr18:44389522-44390022:downstream_20910 | chr18 | 44390022 | 44497495 | - | PIAS2 | NM | 8 | 0,5 | 2 | 0 | 7,30E-01 | NA | 2 | 9 | 6 | 13 | 2 | 9 | 14 | 14 | -0,9 | -1,4 | 0,6 |
| chr1:40538321-40538821:downstream_16550 | chr1 | 40506254 | 40538321 | + | CAP1 | NM | 91 | -0,2 | 1 | 0 | 7,30E-01 | 9,29E-01 | 15 | 33 | 24 | 25 | 160 | 175 | 120 | 173 | 0,0 | 0,2 | -0,2 |
| chr9:131070895-131071395:downstream_6826 | chr9 | 131071395 | 131084697 | - | TRUB2 | NM | 70 | 0,2 | 1 | 0 | 7,30E-01 | 9,29E-01 | 23 | 28 | 21 | 24 | 130 | 97 | 125 | 111 | 0,2 | -0,1 | 0,2 |
| chr19:40021672-40022172:downstream_14287 | chr19 | 40022172 | 40023507 | - | EID2B | NM | 79 | 0,2 | 1 | 0 | 7,30E-01 | 9,29E-01 | 103 | 77 | 11 | 20 | 190 | 161 | 43 | 30 | 2,6 | 2,3 | 0,3 |
| chr5:16451127-16451627:downstream_4199 | chr5 | 16451627 | 16465894 | - | ZNF622 | NM | 67 | 0,2 | 1 | 0 | 7,31E-01 | 9,29E-01 | 54 | 45 | 11 | 15 | 147 | 166 | 43 | 54 | 1,9 | 1,7 | 0,2 |
| chr20:54943718-54944218:downstream_15096 | chr20 | 54933982 | 54943718 | + | FAM210B | NM | 35 | 0,4 | 1 | 0 | 7,31E-01 | 9,29E-01 | 9 | 6 | 7 | 1 | 81 | 85 | 57 | 39 | 0,9 | 0,8 | 0,1 |
| chr5:61683850-61684350:downstream_4271 | chr5 | 61684350 | 61699728 | - | DIMT1 | NM | 35 | 0,3 | 1 | 0 | 7,32E-01 | 9,29E-01 | 16 | 16 | 14 | 6 | 73 | 60 | 51 | 42 | 0,7 | 0,5 | 0,2 |
| chr10:37521495-37521995:downstream_8268 | chr10 | 37414784 | 37521495 | + | ANKRD30A | NM | 26 | -0,3 | 1 | 0 | 7,32E-01 | NA | 7 | 4 | 38 | 39 | 8 | 10 | 61 | 42 | -2,9 | -2,5 | -0,3 |
| chr17:57917952-57918452:downstream_12669 | chr17 | 57784862 | 57917952 | + | VMP1 | NM | 2445 | -0,1 | 0 | 0 | 7,32E-01 | 9,29E-01 | 1322 | 1397 | 3178 | 3417 | 1445 | 1747 | 3504 | 3546 | -1,3 | -1,1 | -0,1 |
| chr14:45367610-45368110:downstream_11123 | chr14 | 45368110 | 45381120 | - | LOC101927418 | NR | 7 | -0,7 | 2 | 0 | 7,32E-01 | NA | 3 | 26 | 4 | 4 | 2 | 15 | 0 | 2 | 1,8 | 2,7 | -0,9 |
| chr5:87484949-87485449:downstream_24416 | chr5 | 87485449 | 87564696 | - | TMEM161B | NM | 10 | -0,5 | 2 | 0 | 7,33E-01 | NA | 6 | 1 | 17 | 3 | 13 | 4 | 10 | 23 | -1,6 | -1,0 | -0,6 |
| chrX:56593443-56593943:downstream_7758 | chrX | 56590025 | 56593443 | + | UBQLN2 | NM | 29 | -0,3 | 1 | 0 | 7,33E-01 | 9,29E-01 | 5 | 19 | 4 | 13 | 49 | 72 | 31 | 41 | 0,5 | 0,7 | -0,3 |
| chr14:59929739-59930239:downstream_11163 | chr14 | 59930239 | 59932059 | - | GPR135 | NM | 26 | 0,3 | 1 | 0 | 7,33E-01 | NA | 28 | 33 | 18 | 26 | 25 | 30 | 33 | 17 | 0,5 | 0,1 | 0,3 |
| chr2:175202268-175202768:downstream_2315 | chr2 | 175199820 | 175202268 | + | SP9 | NM | 33 | 0,3 | 1 | 0 | 7,33E-01 | 9,29E-01 | 31 | 39 | 53 | 56 | 25 | 2 | 28 | 27 | -0,6 | -1,0 | 0,4 |
| chr17:55757299-55757799:downstream_20592 | chr17 | 55333930 | 55757299 | + | MSI2 | NM | 55 | 0,2 | 1 | 0 | 7,34E-01 | 9,29E-01 | 44 | 49 | 122 | 112 | 11 | 16 | 53 | 29 | -1,3 | -1,6 | 0,3 |
| chr4:169908241-169908741:downstream_3901 | chr4 | 169908741 | 169931468 | - | CBR4 | NM | 25 | 0,3 | 1 | 0 | 7,34E-01 | NA | 11 | 23 | 11 | 20 | 36 | 27 | 38 | 36 | 0,1 | -0,2 | 0,4 |
| chr11:122927699-122928199:downstream_18053 | chr11 | 122928199 | 122933043 | - | HSPA8 | NM | 664 | 0,2 | 0 | 0 | 7,34E-01 | 9,29E-01 | 28 | 34 | 38 | 34 | 1244 | 993 | 1456 | 1488 | -0,2 | -0,4 | 0,2 |
| chr1:114228545-114229045:downstream_16932 | chr1 | 113933474 | 114228545 | + | MAGI3 | NM | 29 | 0,3 | 1 | 0 | 7,35E-01 | 9,29E-01 | 8 | 9 | 53 | 34 | 5 | 13 | 77 | 35 | -2,4 | -2,7 | 0,3 |
| chr1:6244579-6245079:downstream_402 | chr1 | 6245079 | 6259679 | - | RPL22 | NM | 26 | 0,3 | 1 | 0 | 7,35E-01 | NA | 12 | 6 | 6 | 4 | 46 | 65 | 36 | 36 | 0,9 | 0,6 | 0,3 |
| chr2:667472-667972:downstream_1825 | chr2 | 667972 | 677439 | - | TMEM18 | NM | 9 | -0,5 | 1 | 0 | 7,35E-01 | NA | 5 | 2 | 7 | 5 | 11 | 12 | 11 | 16 | -0,8 | -0,3 | -0,5 |
| chrX:48769958-48770458:downstream_7713 | chrX | 48770458 | 48776413 | - | PIM2 | NM | 18 | 0,5 | 1 | 0 | 7,35E-01 | NA | 4 | 14 | 0 | 2 | 33 | 65 | 10 | 13 | 2,8 | 2,1 | 0,7 |
| chr12:9094060-9094560:downstream_10025 | chr12 | 9067315 | 9094060 | + | PHC1 | NM | 54 | 0,3 | 1 | 0 | 7,36E-01 | 9,29E-01 | 48 | 147 | 60 | 79 | 19 | 36 | 15 | 31 | 0,5 | 0,2 | 0,3 |
| chr14:96967484-96967984:downstream_11301 | chr14 | 96967984 | 96968663 | - | LOC730202 | NR | 10 | 0,6 | 2 | 0 | 7,36E-01 | NA | 16 | 20 | 4 | 0 | 18 | 15 | 0 | 4 | 3,1 | 3,0 | 0,1 |
| chr10:28971868-28972368:downstream_8263 | chr10 | 28966423 | 28971868 | + | BAMBI | NM | 70 | -0,3 | 1 | 0 | 7,36E-01 | 9,29E-01 | 7 | 7 | 18 | 10 | 95 | 104 | 163 | 154 | -1,0 | -0,7 | -0,3 |
| chr14:24778332-24778832:downstream_19323 | chr14 | 24769059 | 24778332 | + | NOP9 | NM | 16 | 0,4 | 1 | 0 | 7,37E-01 | NA | 3 | 11 | 7 | 3 | 24 | 28 | 18 | 30 | 0,4 | 0,1 | 0,3 |
| chr20:10037410-10037910:downstream_22349 | chr20 | 10015646 | 10037410 | + | ANKEF1 | NM | 40 | -0,2 | 1 | 0 | 7,37E-01 | 9,29E-01 | 24 | 22 | 21 | 23 | 67 | 59 | 51 | 49 | 0,1 | 0,3 | -0,2 |
| chr17:28796675-28797175:downstream_20417 | chr17 | 28705941 | 28796675 | + | CPD | NM | 14 | 0,4 | 1 | 0 | 7,37E-01 | NA | 2 | 4 | 4 | 11 | 11 | 8 | 42 | 33 | -1,4 | -2,0 | 0,5 |
| chr1:35648700-35649200:downstream_15 | chr1 | 35649200 | 35658743 | - | SFPQ | NM | 104 | 0,2 | 1 | 0 | 7,37E-01 | 9,29E-01 | 74 | 63 | 75 | 55 | 151 | 120 | 159 | 134 | 0,1 | -0,1 | 0,2 |
| chr9:106903700-106904200:downstream_26507 | chr9 | 106856540 | 106903700 | + | SMC2 | NM | 15 | -0,4 | 1 | 0 | 7,37E-01 | NA | 4 | 11 | 32 | 29 | 7 | 4 | 18 | 18 | -2,0 | -1,7 | -0,4 |
| chr9:37437599-37438099:downstream_7076 | chr9 | 37438099 | 37465407 | - | ZBTB5 | NM | 21 | 0,4 | 1 | 0 | 7,37E-01 | NA | 20 | 18 | 8 | 2 | 42 | 50 | 10 | 18 | 1,9 | 1,7 | 0,2 |
| chr15:38779911-38780411:downstream_11657 | chr15 | 38746327 | 38779911 | + | FAM98B | NM | 8 | -0,5 | 1 | 0 | 7,38E-01 | NA | 6 | 4 | 3 | 5 | 18 | 12 | 8 | 9 | 0,3 | 0,8 | -0,5 |
| chr5:141369856-141370356:downstream_24337 | chr5 | 141346401 | 141369856 | + | RNF14 | NM | 11 | -0,4 | 1 | 0 | 7,38E-01 | NA | 19 | 17 | 14 | 7 | 10 | 14 | 1 | 7 | 0,8 | 1,4 | -0,6 |
| chr17:27051374-27051874:downstream_12937 | chr17 | 27046999 | 27051374 | + | RPL23A | NM | 62 | 0,2 | 1 | 0 | 7,38E-01 | 9,29E-01 | 111 | 133 | 63 | 31 | 57 | 55 | 22 | 27 | 1,4 | 1,2 | 0,2 |
| chr8:27141902-27142402:downstream_26164 | chr8 | 27142402 | 27168836 | - | TRIM35 | NM | 10 | 0,6 | 2 | 0 | 7,38E-01 | NA | 4 | 23 | 6 | 1 | 6 | 27 | 3 | 7 | 2,1 | 1,7 | 0,4 |
| chr1:27327377-27327877:downstream_581 | chr1 | 27320194 | 27327377 | + | TRNP1 | NM | 78 | -0,2 | 1 | 0 | 7,38E-01 | 9,29E-01 | 18 | 37 | 46 | 35 | 98 | 115 | 142 | 130 | -0,6 | -0,4 | -0,2 |
| chr8:17080241-17080741:downstream_6264 | chr8 | 17013835 | 17080241 | + | ZDHHC2 | NM | 10 | 0,5 | 2 | 0 | 7,38E-01 | NA | 11 | 0 | 11 | 13 | 8 | 3 | 15 | 18 | -1,1 | -1,6 | 0,5 |
| chr4:38967865-38968365:downstream_23890 | chr4 | 38968365 | 39034041 | - | TMEM156 | NM | 23 | -0,4 | 1 | 0 | 7,39E-01 | NA | 7 | 2 | 13 | 16 | 28 | 14 | 53 | 49 | -1,6 | -1,3 | -0,4 |
| chr1:156691182-156691682:downstream_1116 | chr1 | 156691682 | 156698231 | - | ISG20L2 | NM | 13 | -0,4 | 1 | 0 | 7,39E-01 | NA | 22 | 13 | 7 | 5 | 15 | 31 | 7 | 5 | 1,5 | 2,0 | -0,4 |
| chr12:32531141-32531641:downstream_18457 | chr12 | 32260184 | 32531141 | + | BICD1 | NM | 8 | 0,6 | 2 | 0 | 7,39E-01 | NA | 1 | 5 | 10 | 19 | 3 | 0 | 15 | 11 | -2,3 | -3,0 | 0,7 |
| chr5:172461900-172462400:downstream_4582 | chr5 | 172410762 | 172461900 | + | ATP6V0E1 | NM | 415 | -0,4 | 1 | 0 | 7,39E-01 | 9,29E-01 | 23 | 3 | 19 | 7 | 949 | 915 | 646 | 756 | -0,1 | 0,4 | -0,5 |
| chr17:37408396-37408896:downstream_20478 | chr17 | 37408896 | 37557909 | - | FBXL20 | NM | 11 | -0,7 | 2 | 0 | 7,40E-01 | NA | 0 | 0 | 7 | 4 | 1 | 3 | 31 | 43 | -Inf | -4,3 | -Inf |
| chr7:127949935-127950435:downstream_25705 | chr7 | 127950435 | 127983962 | - | RBM28 | NM | 20 | -0,3 | 1 | 0 | 7,40E-01 | NA | 17 | 8 | 26 | 21 | 21 | 14 | 26 | 26 | -0,9 | -0,6 | -0,3 |
| chr15:72117860-72118360:downstream_11418 | chr15 | 72118360 | 72410440 | - | MYO9A | NM | 32 | -0,3 | 1 | 0 | 7,40E-01 | 9,29E-01 | 7 | 17 | 60 | 38 | 13 | 19 | 61 | 43 | -2,0 | -1,7 | -0,3 |
| chr17:17103808-17104308:downstream_12866 | chr17 | 17104308 | 17109646 | - | PLD6 | NM | 16 | -0,4 | 1 | 0 | 7,40E-01 | NA | 7 | 5 | 6 | 1 | 46 | 36 | 19 | 9 | 0,9 | 1,5 | -0,6 |
| chr11:6701115-6701615:downstream_9106 | chr11 | 6701615 | 6704632 | - | MRPL17 | NM | 31 | 0,3 | 1 | 0 | 7,41E-01 | 9,29E-01 | 27 | 39 | 10 | 13 | 62 | 51 | 22 | 26 | 1,5 | 1,2 | 0,3 |
| chr4:75670947-75671447:downstream_3743 | chr4 | 75671447 | 75719882 | - | BTC | NM | 6 | 0,7 | 2 | 0 | 7,41E-01 | NA | 8 | 0 | 7 | 6 | 7 | 0 | 8 | 10 | -0,6 | -1,3 | 0,7 |
| chr1:110174677-110175177:downstream_16470 | chr1 | 110162434 | 110174677 | + | AMPD2 | NM | 13 | 0,4 | 1 | 0 | 7,41E-01 | NA | 3 | 8 | 7 | 1 | 20 | 24 | 24 | 16 | 0,5 | 0,2 | 0,3 |
| chrX:45605920-45606420:downstream_7685 | chrX | 45606420 | 45606530 | - | MIR222 | NR | 69 | 0,3 | 1 | 0 | 7,41E-01 | 9,29E-01 | 197 | 154 | 39 | 38 | 54 | 45 | 3 | 21 | 2,2 | 2,1 | 0,1 |
| chr1:186650578-186651078:downstream_1270 | chr1 | 186649785 | 186650578 | + | PACERR | NR | 40 | 0,3 | 1 | 0 | 7,41E-01 | 9,29E-01 | 22 | 64 | 64 | 58 | 24 | 18 | 38 | 35 | -0,5 | -0,8 | 0,3 |
| chr1:145590166-145590666:downstream_17097 | chr1 | 145590666 | 145611044 | - | POLR3C | NM | 45 | -0,3 | 1 | 0 | 7,42E-01 | 9,29E-01 | 41 | 29 | 28 | 16 | 84 | 78 | 56 | 27 | 0,7 | 1,0 | -0,3 |
| chr19:58984945-58985445:downstream_13809 | chr19 | 58978411 | 58984945 | + | ZNF324 | NM | 19 | 0,3 | 1 | 0 | 7,42E-01 | NA | 47 | 29 | 19 | 13 | 15 | 15 | 10 | 7 | 1,2 | 0,9 | 0,3 |
| chr1:242251188-242251688:downstream_17091 | chr1 | 242251688 | 242687998 | - | PLD5 | NM | 7 | -0,6 | 2 | 0 | 7,42E-01 | NA | 2 | 0 | 13 | 3 | 4 | 3 | 15 | 12 | -2,7 | -2,0 | -0,7 |
| chr20:54979582-54980082:downstream_22396 | chr20 | 54967426 | 54979582 | + | CSTF1 | NM | 19 | -0,3 | 1 | 0 | 7,42E-01 | NA | 7 | 4 | 8 | 8 | 24 | 33 | 36 | 29 | -0,6 | -0,2 | -0,4 |
| chr17:43100205-43100705:downstream_20442 | chr17 | 43100705 | 43138477 | - | DCAKD | NM | 30 | 0,3 | 1 | 0 | 7,42E-01 | 9,29E-01 | 9 | 5 | 1 | 5 | 72 | 62 | 46 | 39 | 1,1 | 0,7 | 0,5 |
| chr16:70611571-70612071:downstream_12454 | chr16 | 70557690 | 70611571 | + | SF3B3 | NM | 23 | -0,3 | 1 | 0 | 7,43E-01 | NA | 6 | 6 | 8 | 14 | 32 | 27 | 56 | 39 | -1,0 | -0,7 | -0,3 |
| chr11:57510485-57510985:downstream_9320 | chr11 | 57510985 | 57519253 | - | BTBD18 | NM | 5 | 0,8 | 2 | 0 | 7,43E-01 | NA | 2 | 16 | 0 | 2 | 3 | 11 | 3 | 0 | 3,5 | 2,3 | 1,2 |
| chr2:3483342-3483842:downstream_1564 | chr2 | 3383445 | 3483342 | + | TRAPPC12 | NM | 11 | -0,5 | 2 | 0 | 7,44E-01 | NA | 2 | 0 | 11 | 16 | 4 | 3 | 24 | 30 | -3,5 | -2,9 | -0,6 |
| chr10:60130513-60131013:downstream_17807 | chr10 | 60094738 | 60130513 | + | UBE2D1 | NM | 14 | -0,4 | 1 | 0 | 7,44E-01 | NA | 17 | 21 | 6 | 5 | 34 | 20 | 6 | 6 | 1,9 | 2,3 | -0,4 |
| chr9:119187003-119187503:downstream_26222 | chr9 | 119187503 | 120177317 | - | ASTN2 | NM | 5 | -0,7 | 2 | 0 | 7,44E-01 | NA | 1 | 4 | 8 | 8 | 5 | 0 | 3 | 8 | -1,8 | -1,2 | -0,7 |
| chr19:11032943-11033443:downstream_14004 | chr19 | 11033443 | 11039688 | - | YIPF2 | NM | 25 | -0,3 | 1 | 0 | 7,44E-01 | NA | 15 | 16 | 19 | 20 | 26 | 38 | 38 | 29 | -0,3 | -0,1 | -0,3 |
| chr18:48476162-48476662:downstream_20894 | chr18 | 48405431 | 48476162 | + | ME2 | NM | 6 | 0,6 | 2 | 0 | 7,44E-01 | NA | 4 | 3 | 11 | 17 | 2 | 0 | 7 | 4 | -2,0 | -2,8 | 0,7 |
| chr5:156455923-156456423:downstream_24159 | chr5 | 156456423 | 156486130 | - | HAVCR1 | NM | 334 | -0,2 | 0 | 0 | 7,44E-01 | 9,30E-01 | 262 | 293 | 441 | 510 | 189 | 269 | 303 | 400 | -0,8 | -0,6 | -0,2 |
| chr2:85824831-85825331:downstream_2079 | chr2 | 85822836 | 85824831 | + | RNF181 | NM | 29 | 0,3 | 1 | 0 | 7,44E-01 | 9,30E-01 | 23 | 30 | 8 | 15 | 46 | 56 | 28 | 28 | 1,2 | 0,9 | 0,3 |
| chrX:15261608-15262108:downstream_26630 | chrX | 15262108 | 15288589 | - | ASB9 | NM | 11 | -0,5 | 2 | 0 | 7,44E-01 | NA | 2 | 2 | 3 | 10 | 19 | 2 | 24 | 30 | -1,8 | -1,4 | -0,4 |
| chr2:100016728-100017228:downstream_1679 | chr2 | 99953833 | 100016728 | + | EIF5B | NM | 21 | -0,3 | 1 | 0 | 7,45E-01 | NA | 9 | 8 | 33 | 17 | 12 | 16 | 36 | 32 | -1,6 | -1,3 | -0,4 |
| chr15:83423616-83424116:downstream_19553 | chr15 | 83424116 | 83474822 | - | FSD2 | NM | 18 | -0,5 | 2 | 0 | 7,45E-01 | NA | 32 | 76 | 3 | 9 | 8 | 16 | 0 | 2 | 3,2 | 3,9 | -0,7 |
| chr17:7260538-7261038:downstream_12798 | chr17 | 7258496 | 7260538 | + | TMEM95 | NM | 6 | -0,9 | 3 | 0 | 7,46E-01 | NA | 12 | 4 | 0 | 0 | 19 | 11 | 0 | 0 | Inf | Inf | NaN |
| chr5:131980313-131980813:downstream_4406 | chr5 | 131892615 | 131980313 | + | RAD50 | NM | 15 | -0,4 | 1 | 0 | 7,46E-01 | NA | 8 | 0 | 26 | 26 | 9 | 2 | 22 | 29 | -2,7 | -2,2 | -0,5 |
| chr7:107561643-107562143:downstream_25484 | chr7 | 107531551 | 107561643 | + | DLD | NM | 20 | -0,3 | 1 | 0 | 7,46E-01 | NA | 5 | 6 | 11 | 11 | 18 | 29 | 43 | 36 | -1,1 | -0,8 | -0,3 |
| chr10:94113721-94114221:downstream_8317 | chr10 | 94050919 | 94113721 | + | MARCH5 | NM | 13 | 0,4 | 1 | 0 | 7,46E-01 | NA | 8 | 2 | 8 | 7 | 11 | 16 | 18 | 33 | -0,6 | -0,9 | 0,3 |

| | | | | | | | | | | | | | | | | | | | | | | | |
|---|---|---|---|---|---|---|---|---|---|---|---|---|---|---|---|---|---|---|---|---|---|---|---|
| chr2:68356780-68357280:downstream_22323 | chr2 | 68357280 | 68384692 | - | WDR92 | NM | 8 | 0,5 | 2 | 0 | 7,46E-01 | NA | 12 | 5 | 13 | 14 | 2 | 4 | 11 | 2 | -0,6 | -1,2 | 0,6 |
| chr13:48649363-48649863:downstream_19025 | chr13 | 48649863 | 48669277 | - | MED4 | NM | 10 | -0,5 | 1 | 0 | 7,47E-01 | NA | 6 | 15 | 3 | 2 | 18 | 27 | 3 | 5 | 2,0 | 2,5 | -0,5 |
| chr18:71017124-71017624:downstream_13670 | chr18 | 70992175 | 71017124 | + | LOC100505817 | NR | 67 | -0,2 | 1 | 0 | 7,47E-01 | 9,30E-01 | 31 | 20 | 78 | 73 | 39 | 55 | 127 | 112 | -1,6 | -1,3 | -0,2 |
| chr2:24223693-24224193:downstream_1886 | chr2 | 24163375 | 24223693 | + | UBXN2A | NM | 10 | 0,4 | 1 | 0 | 7,47E-01 | NA | 4 | 9 | 10 | 16 | 3 | 9 | 17 | 16 | -1,0 | -1,4 | 0,4 |
| chr5:74666427-74666927:downstream_24069 | chr5 | 74666927 | 74807806 | - | COL4A3BP | NM | 13 | -0,5 | 2 | 0 | 7,47E-01 | NA | 0 | 2 | 11 | 5 | 4 | 9 | 33 | 43 | -3,1 | -2,6 | -0,5 |
| chr2:178994382-178994882:downstream_2338 | chr2 | 178977150 | 178994382 | + | RBM45 | NM | 14 | -0,4 | 1 | 0 | 7,47E-01 | NA | 11 | 7 | 11 | 4 | 20 | 26 | 19 | 11 | 0,2 | 0,6 | -0,4 |
| chr11:839220-839720:downstream_8986 | chr11 | 839720 | 842529 | - | POLR2L | NM | 14 | 0,4 | 1 | 0 | 7,48E-01 | NA | 9 | 19 | 6 | 5 | 29 | 19 | 11 | 12 | 1,4 | 1,1 | 0,4 |
| chr9:100436029-100436529:downstream_6898 | chr9 | 100395704 | 100436029 | + | NCBP1 | NM | 41 | -0,3 | 1 | 0 | 7,48E-01 | 9,30E-01 | 7 | 19 | 25 | 18 | 59 | 50 | 75 | 72 | -0,7 | -0,4 | -0,3 |
| chr5:34925787-34926287:downstream_4215 | chr5 | 34915819 | 34925787 | + | BRIX1 | NM | 88 | 0,2 | 1 | 0 | 7,48E-01 | 9,30E-01 | 49 | 17 | 15 | 17 | 211 | 176 | 107 | 115 | 1,0 | 0,8 | 0,2 |
| chr2:32449181-32449681:downstream_22223 | chr2 | 32390909 | 32449181 | + | SLC30A6 | NM | 25 | 0,3 | 1 | 0 | 7,48E-01 | NA | 15 | 15 | 39 | 30 | 11 | 15 | 40 | 31 | -1,2 | -1,5 | 0,3 |
| chr3:44155802-44156302:downstream_2903 | chr3 | 44155703 | 44155802 | + | MIR138-1 | NR | 19 | 0,3 | 1 | 0 | 7,48E-01 | NA | 9 | 5 | 36 | 25 | 3 | 9 | 32 | 34 | -2,2 | -2,5 | 0,3 |
| chr19:13064469-13064969:downstream_14041 | chr19 | 13064969 | 13068068 | - | GADD45GIP1 | NM | 39 | -0,3 | 1 | 0 | 7,48E-01 | 9,30E-01 | 6 | 4 | 6 | 3 | 105 | 70 | 46 | 75 | 0,1 | 0,5 | -0,4 |
| chr13:20248391-20248891:downstream_19074 | chr13 | 20248891 | 20357159 | - | PSPC1 | NR | 7 | -0,5 | 2 | 0 | 7,48E-01 | NA | 2 | 8 | 1 | 9 | 8 | 12 | 4 | 11 | -0,2 | 0,3 | -0,5 |
| chr12:64842463-64842963:downstream_10292 | chr12 | 64798152 | 64842463 | + | XPOT | NM | 79 | -0,3 | 1 | 0 | 7,48E-01 | 9,30E-01 | 3 | 6 | 13 | 14 | 89 | 80 | 241 | 185 | -1,5 | -1,3 | -0,2 |
| chr17:34895150-34895650:downstream_13002 | chr17 | 34891402 | 34895150 | + | PIGW | NM | 14 | 0,4 | 1 | 0 | 7,49E-01 | NA | 20 | 23 | 3 | 7 | 24 | 17 | 7 | 7 | 2,1 | 1,6 | 0,5 |
| chr8:94743230-94743730:downstream_6640 | chr8 | 94743730 | 94753224 | - | RBM12B | NM | 43 | 0,2 | 1 | 0 | 7,49E-01 | 9,30E-01 | 43 | 81 | 33 | 28 | 43 | 58 | 32 | 27 | 1,0 | 0,8 | 0,2 |
| chr4:41702061-41702561:downstream_23722 | chr4 | 41361623 | 41702061 | + | LIMCH1 | NM | 44 | 0,3 | 1 | 0 | 7,49E-01 | 9,30E-01 | 0 | 8 | 11 | 20 | 24 | 26 | 128 | 138 | -2,0 | -2,4 | 0,4 |
| chr17:40725221-40725721:downstream_20587 | chr17 | 40719077 | 40725221 | + | MLX | NM | 8 | -0,5 | 2 | 0 | 7,49E-01 | NA | 12 | 8 | 13 | 5 | 9 | 6 | 7 | 2 | 0,2 | 0,6 | -0,5 |
| chr12:12798042-12798542:downstream_10068 | chr12 | 12764766 | 12798042 | + | CREBL2 | NM | 23 | -0,3 | 1 | 0 | 7,49E-01 | NA | 3 | 6 | 3 | 7 | 37 | 48 | 50 | 31 | -0,2 | 0,1 | -0,2 |
| chr5:111064499-111064999:downstream_24258 | chr5 | 111064999 | 111312628 | - | NREP | NM | 3 | 1,0 | 3 | 0 | 7,49E-01 | NA | 0 | 0 | 1 | 7 | 0 | 0 | 7 | 11 | -Inf | -Inf | NaN |
| chr3:73115011-73115511:downstream_3038 | chr3 | 73046118 | 73115011 | + | PPP4R2 | NM | 72 | 0,2 | 1 | 0 | 7,49E-01 | 9,30E-01 | 34 | 54 | 72 | 77 | 53 | 61 | 116 | 107 | -0,8 | -1,0 | 0,2 |
| chr6:41902170-41902670:downstream_24551 | chr6 | 41902670 | 42016632 | - | CCND3 | NM | 16 | 0,4 | 1 | 0 | 7,50E-01 | NA | 8 | 6 | 10 | 11 | 25 | 6 | 26 | 35 | -0,6 | -1,0 | 0,4 |
| chrX:54842448-54842948:downstream_26901 | chrX | 54834031 | 54842448 | + | MAGED2 | NM | 59 | 0,3 | 1 | 0 | 7,51E-01 | 9,31E-01 | 1 | 15 | 6 | 5 | 94 | 148 | 110 | 89 | 0,6 | 0,3 | 0,4 |
| chr3:149691029-149691529:downstream_3230 | chr3 | 149689065 | 149691029 | + | LOC646903 | NR | 4 | 0,6 | 2 | 0 | 7,51E-01 | NA | 8 | 8 | 4 | 1 | 7 | 3 | 3 | 2 | 1,7 | 1,1 | 0,6 |
| chr4:52783003-52783503:downstream_23616 | chr4 | 52709165 | 52783003 | + | DCUN1D4 | NM | 5 | -0,6 | 2 | 0 | 7,51E-01 | NA | 2 | 0 | 7 | 5 | 2 | 6 | 7 | 15 | -2,3 | -1,5 | -0,7 |
| chr5:132204536-132205036:downstream_4410 | chr5 | 132202318 | 132204536 | + | UQCRQ | NM | 49 | 0,2 | 1 | 0 | 7,52E-01 | 9,31E-01 | 54 | 54 | 14 | 23 | 85 | 90 | 40 | 31 | 1,6 | 1,3 | 0,3 |
| chr12:49747467-49747967:downstream_18544 | chr12 | 49740699 | 49747467 | + | DNAJC22 | NM | 68 | -0,2 | 1 | 0 | 7,52E-01 | 9,31E-01 | 28 | 54 | 49 | 60 | 95 | 68 | 99 | 90 | -0,4 | -0,2 | -0,2 |
| chr15:49937333-49937833:downstream_19532 | chr15 | 49913225 | 49937333 | + | DTWD1 | NM | 9 | -0,4 | 1 | 0 | 7,53E-01 | NA | 14 | 8 | 18 | 5 | 8 | 9 | 4 | 8 | -0,1 | 0,4 | -0,5 |
| chr8:67342968-67343468:downstream_6582 | chr8 | 67341262 | 67342968 | + | RRS1 | NM | 70 | -0,2 | 1 | 0 | 7,53E-01 | 9,31E-01 | 15 | 44 | 14 | 10 | 146 | 216 | 56 | 63 | 1,3 | 1,6 | -0,3 |
| chr17:48458093-48458593:downstream_13228 | chr17 | 48458593 | 48474914 | - | LRRC59 | NM | 209 | 0,2 | 1 | 0 | 7,53E-01 | 9,31E-01 | 11 | 3 | 3 | 9 | 392 | 416 | 426 | 412 | 0,3 | -0,1 | 0,3 |
| chr4:76567452-76567952:downstream_23660 | chr4 | 76567952 | 76598667 | - | G3BP2 | NM | 55 | -0,2 | 1 | 0 | 7,53E-01 | 9,31E-01 | 20 | 21 | 36 | 17 | 59 | 108 | 67 | 108 | -0,4 | -0,1 | -0,3 |
| chr9:88637217-88637717:downstream_6884 | chr9 | 88556056 | 88637217 | + | NAA35 | NM | 20 | -0,3 | 1 | 0 | 7,53E-01 | NA | 7 | 11 | 26 | 30 | 5 | 19 | 24 | 35 | -1,6 | -1,3 | -0,4 |
| chr19:53915262-53915762:downstream_14578 | chr19 | 53898396 | 53915262 | + | ZNF765 | NM | 27 | 0,3 | 1 | 0 | 7,54E-01 | NA | 37 | 20 | 25 | 14 | 26 | 38 | 39 | 15 | 0,5 | 0,2 | 0,3 |
| chr7:131181398-131181898:downstream_25632 | chr7 | 130794854 | 131181398 | + | MKLN1 | NM | 27 | -0,4 | 1 | 0 | 7,54E-01 | NA | 2 | 1 | 18 | 21 | 4 | 57 | 11 | 103 | -3,9 | -3,4 | -0,4 |
| chr4:22328489-22328989:downstream_3639 | chr4 | 22328989 | 22341289 | - | LOC100505912 | NR | 4 | 0,7 | 2 | 0 | 7,54E-01 | NA | 0 | 4 | 3 | 8 | 2 | 0 | 7 | 6 | -1,5 | -2,4 | 0,9 |
| chr9:37867665-37868165:downstream_26276 | chr9 | 37800550 | 37867665 | + | DCAF10 | NM | 14 | 0,4 | 1 | 0 | 7,54E-01 | NA | 4 | 3 | 15 | 7 | 7 | 8 | 45 | 19 | -1,7 | -2,1 | 0,4 |
| chr6:24171482-24171982:downstream_24720 | chr6 | 24171982 | 24383520 | - | DCDC2 | NM | 30 | 0,3 | 1 | 0 | 7,54E-01 | 9,31E-01 | 6 | 12 | 19 | 24 | 34 | 10 | 75 | 64 | -1,3 | -1,7 | 0,3 |
| chr3:40353915-40354415:downstream_2884 | chr3 | 40351172 | 40353915 | + | EIF1B | NM | 18 | -0,3 | 1 | 0 | 7,54E-01 | NA | 15 | 7 | 8 | 7 | 41 | 32 | 18 | 20 | 0,6 | 0,9 | -0,4 |
| chr2:233412179-233412679:downstream_2484 | chr2 | 233412679 | 233415300 | - | TIGD1 | NM | 23 | 0,3 | 1 | 0 | 7,55E-01 | NA | 12 | 18 | 7 | 4 | 51 | 49 | 22 | 21 | 1,5 | 1,2 | 0,3 |
| chr2:122288585-122289085:downstream_2222 | chr2 | 122288455 | 122288585 | + | RNU4ATAC | NR | 16 | 0,4 | 1 | 0 | 7,55E-01 | NA | 15 | 4 | 7 | 1 | 27 | 41 | 15 | 16 | 1,3 | 1,1 | 0,2 |
| chr2:206987302-206987802:downstream_22109 | chr2 | 206987802 | 207024243 | - | NDUFS1 | NM | 28 | -0,3 | 1 | 0 | 7,55E-01 | 9,32E-01 | 11 | 16 | 19 | 29 | 25 | 36 | 38 | 51 | -0,8 | -0,5 | -0,3 |
| chr5:102614361-102614861:downstream_4355 | chr5 | 102594441 | 102614361 | + | C5orf30 | NM | 10 | -0,4 | 1 | 0 | 7,55E-01 | NA | 5 | 5 | 10 | 7 | 10 | 7 | 13 | 21 | -0,8 | -0,3 | -0,5 |
| chr2:217084915-217085415:downstream_2407 | chr2 | 217081611 | 217084915 | + | PKI55 | NR | 12 | -0,4 | 1 | 0 | 7,55E-01 | NA | 5 | 5 | 13 | 13 | 11 | 11 | 17 | 26 | -1,4 | -1,0 | -0,4 |
| chr5:81551216-81551716:downstream_24019 | chr5 | 81267843 | 81551216 | + | ATG10 | NM | 5 | -0,6 | 2 | 0 | 7,56E-01 | NA | 2 | 2 | 10 | 11 | 2 | 2 | 7 | 7 | -2,6 | -2,0 | -0,6 |
| chr3:14239869-14240369:downstream_2830 | chr3 | 14220227 | 14239869 | + | LSM3 | NM | 79 | 0,3 | 1 | 0 | 7,56E-01 | 9,32E-01 | 5 | 8 | 4 | 2 | 189 | 188 | 135 | 98 | 0,9 | 0,7 | 0,2 |
| chr11:28041662-28042162:downstream_8862 | chr11 | 28042162 | 28129746 | - | KIF18A | NM | 13 | 0,5 | 2 | 0 | 7,56E-01 | NA | 2 | 2 | 17 | 9 | 7 | 0 | 35 | 32 | -2,9 | -3,4 | 0,5 |
| chr12:56078394-56078894:downstream_10242 | chr12 | 56075329 | 56078394 | + | METTL7B | NM | 32 | 0,3 | 1 | 0 | 7,57E-01 | 9,32E-01 | 5 | 6 | 9 | 6 | 5 | 52 | 68 | 46 | 67 | 0,4 | 0,1 | 0,3 |
| chr14:34392920-34393420:downstream_19217 | chr14 | 34393420 | 34420284 | - | EGLN3 | NM | 6 | 0,5 | 2 | 0 | 7,58E-01 | NA | 11 | 10 | 1 | 7 | 9 | 3 | 3 | 5 | 1,2 | 0,6 | 0,6 |
| chr10:81115089-81115589:downstream_8587 | chr10 | 81107219 | 81115089 | + | PPIF | NM | 9 | 0,5 | 2 | 0 | 7,58E-01 | NA | 7 | 9 | 7 | 0 | 19 | 15 | 3 | 16 | 1,1 | 0,8 | 0,3 |
| chr17:48939086-48939586:downstream_20785 | chr17 | 48939586 | 48945339 | - | TOB1 | NM | 90 | -0,2 | 1 | 0 | 7,58E-01 | 9,32E-01 | 71 | 47 | 64 | 59 | 108 | 142 | 95 | 132 | -0,1 | 0,1 | -0,2 |
| chr6:21598849-21599349:downstream_4886 | chr6 | 21593971 | 21598849 | + | SOX4 | NM | 59 | -0,2 | 1 | 0 | 7,58E-01 | 9,32E-01 | 15 | 3 | 21 | 25 | 55 | 74 | 125 | 155 | -1,4 | -1,1 | -0,3 |
| chr11:13033653-13034153:downstream_9150 | chr11 | 13030969 | 13033653 | + | RASSF10 | NM | 19 | 0,3 | 1 | 0 | 7,58E-01 | NA | 11 | 15 | 29 | 20 | 8 | 15 | 25 | 29 | -1,0 | -1,2 | 0,3 |
| chr14:59938905-59939405:downstream_11162 | chr14 | 59939405 | 59951073 | - | L3HYPDH | NM | 18 | 0,3 | 1 | 0 | 7,58E-01 | NA | 34 | 15 | 29 | 28 | 9 | 6 | 17 | 6 | -0,2 | -0,6 | 0,4 |
| chr3:160788817-160789317:downstream_2738 | chr3 | 160473995 | 160788817 | + | PPM1L | NM | 6 | -0,6 | 2 | 0 | 7,59E-01 | NA | 2 | 0 | 10 | 7 | 1 | 3 | 8 | 15 | -3,4 | -2,6 | -0,8 |
| chr15:43698911-43699411:downstream_19816 | chr15 | 43699411 | 43802707 | - | TP53BP1 | NM | 16 | 0,4 | 1 | 0 | 7,59E-01 | NA | 5 | 0 | 17 | 19 | 8 | 0 | 35 | 45 | -2,9 | -3,3 | 0,4 |
| chr5:43572786-43573286:downstream_4241 | chr5 | 43573286 | 43603332 | - | NNT-AS1 | NR | 5 | 0,7 | 2 | 0 | 7,59E-01 | NA | 3 | 0 | 6 | 5 | 4 | 0 | 10 | 11 | -1,7 | -2,3 | 0,6 |
| chr20:49201300-49201800:downstream_22519 | chr20 | 49126857 | 49201300 | + | PTPN1 | NM | 6 | 0,5 | 2 | 0 | 7,59E-01 | NA | 6 | 3 | 4 | 5 | 4 | 7 | 7 | 13 | -0,5 | -0,9 | 0,4 |
| chrX:13683527-13684027:downstream_27070 | chrX | 13671224 | 13683527 | + | TCEANC | NM | 6 | 0,6 | 2 | 0 | 7,59E-01 | NA | 10 | 9 | 3 | 0 | 15 | 8 | 3 | 2 | 2,7 | 2,3 | 0,4 |
| chr1:179889212-179889712:downstream_17345 | chr1 | 179851176 | 179889212 | + | TOR1AIP1 | NM | 45 | -0,2 | 1 | 0 | 7,60E-01 | 9,33E-01 | 22 | 13 | 36 | 31 | 46 | 51 | 83 | 76 | -1,0 | -0,7 | -0,2 |
| chr19:45574257-45574757:downstream_14375 | chr19 | 45574757 | 45579688 | - | ZNF296 | NM | 3 | 0,9 | 3 | 0 | 7,60E-01 | NA | 7 | 10 | 0 | 0 | 7 | 1 | 0 | 0 | Inf | Inf | NaN |
| chr11:9611313-9611813:downstream_18391 | chr11 | 9595227 | 9611313 | + | WEE1 | NM | 12 | 0,4 | 1 | 0 | 7,60E-01 | NA | 7 | 5 | 4 | 7 | 11 | 22 | 21 | 21 | 0,1 | -0,3 | 0,4 |
| chr1:51739119-51739619:downstream_729 | chr1 | 51701944 | 51739119 | + | RNF11 | NM | 12 | 0,4 | 1 | 0 | 7,60E-01 | NA | 15 | 35 | 7 | 5 | 12 | 13 | 1 | 6 | 2,1 | 1,8 | 0,3 |
| chr15:88419487-88419987:downstream_19653 | chr15 | 88419987 | 88799962 | - | NTRK3 | NM | 12 | 0,4 | 1 | 0 | 7,60E-01 | NA | 5 | 15 | 17 | 11 | 11 | 8 | 24 | 11 | -0,5 | -0,9 | 0,4 |
| chr22:20098888-20099388:downstream_22951 | chr22 | 20099388 | 20104818 | - | TRMT2A | NM | 9 | -0,5 | 1 | 0 | 7,60E-01 | NA | 7 | 6 | 8 | 2 | 10 | 22 | 11 | 6 | 0,4 | 0,9 | -0,5 |
| chr17:34965407-34965907:downstream_13001 | chr17 | 34958024 | 34965407 | + | MRM1 | NM | 15 | 0,4 | 1 | 0 | 7,60E-01 | NA | 6 | 3 | 1 | 5 | 30 | 20 | 26 | 25 | 0,4 | 0,0 | 0,5 |
| chr12:58155616-58156116:downstream_10281 | chr12 | 58156116 | 58160976 | - | CYP27B1 | NM | 2 | 1,0 | 3 | 0 | 7,60E-01 | NA | 7 | 1 | 0 | 0 | 10 | 1 | 0 | 1 | Inf | 3,7 | Inf |
| chr20:47656499-47656999:downstream_22391 | chr20 | 47656999 | 47662593 | - | CSE1L-AS1 | NR | 5 | -0,6 | 2 | 0 | 7,61E-01 | NA | 9 | 14 | 4 | 2 | 8 | 3 | 1 | 1 | 1,8 | 2,3 | -0,6 |
| chr10:100175454-100175954:downstream_17600 | chr10 | 100175954 | 100206704 | - | HPS1 | NM | 5 | -0,7 | 2 | 0 | 7,61E-01 | NA | 8 | 0 | 3 | 7 | 14 | 0 | 4 | 6 | -0,2 | 0,5 | -0,7 |
| chr7:17385775-17386275:downstream_5720 | chr7 | 17338275 | 17385775 | + | AHR | NM | 14 | 0,4 | 1 | 0 | 7,61E-01 | NA | 7 | 3 | 11 | 11 | 12 | 9 | 33 | 25 | -1,1 | -1,5 | 0,3 |
| chr11:71807837-71808337:downstream_9517 | chr11 | 71808337 | 71814433 | - | LAMTOR1 | NM | 32 | 0,3 | 1 | 0 | 7,62E-01 | 9,35E-01 | 17 | 5 | 15 | 7 | 44 | 55 | 57 | 57 | 0,0 | -0,2 | 0,2 |
| chr9:5233967-5234467:downstream_6975 | chr9 | 5231418 | 5233967 | + | INSL4 | NM | 518 | -0,1 | 0 | 0 | 7,62E-01 | 9,35E-01 | 342 | 282 | 182 | 234 | 971 | 961 | 543 | 633 | 0,6 | 0,7 | -0,1 |
| chr16:28977767-28978267:downstream_11991 | chr16 | 28962317 | 28977767 | + | NFATC2IP | NM | 23 | -0,3 | 1 | 0 | 7,62E-01 | NA | 9 | 14 | 31 | 19 | 14 | 25 | 38 | 32 | -1,1 | -0,8 | -0,3 |
| chr3:97660160-97660660:downstream_23274 | chr3 | 97660660 | 97691295 | - | MINA | NM | 19 | 0,3 | 1 | 0 | 7,63E-01 | NA | 22 | 15 | 11 | 16 | 24 | 20 | 21 | 19 | 0,5 | 0,2 | 0,3 |
| chr3:53258222-53258722:downstream_23471 | chr3 | 53258722 | 53290130 | - | TKT | NM | 13 | 0,3 | 1 | 0 | 7,63E-01 | NA | 11 | 7 | 7 | 11 | 11 | 19 | 22 | 18 | 0,0 | -0,4 | 0,4 |
| chr9:33460850-33461350:downstream_26413 | chr9 | 33461350 | 33473941 | - | NOL6 | NM | 25 | -0,3 | 1 | 0 | 7,63E-01 | NA | 18 | 20 | 7 | 4 | 47 | 78 | 13 | 16 | 1,8 | 2,2 | -0,4 |
| chr10:5903188-5903688:downstream_17469 | chr10 | 5903688 | 5931860 | - | ANKRD16 | NM | 7 | 0,5 | 2 | 0 | 7,63E-01 | NA | 5 | 15 | 6 | 9 | 4 | 5 | 9 | 4 | 0,4 | -0,1 | 0,5 |
| chr8:61100922-61101422:downstream_6569 | chr8 | 61101422 | 61193954 | - | CA8 | NM | 11 | -0,4 | 1 | 0 | 7,64E-01 | NA | 6 | 0 | 14 | 10 | 8 | 7 | 19 | 25 | -2,1 | -1,6 | -0,5 |
| chr9:132573560-132574060:downstream_7357 | chr9 | 132565431 | 132573560 | + | TOR1B | NM | 39 | 0,3 | 1 | 0 | 7,64E-01 | 9,37E-01 | 15 | 11 | 10 | 5 | 80 | 82 | 57 | 49 | 0,8 | 0,6 | 0,2 |
| chrX:23704514-23705014:downstream_7631 | chrX | 23685644 | 23704514 | + | PRDX4 | NM | 14 | -0,4 | 1 | 0 | 7,65E-01 | NA | 3 | 10 | 10 | 8 | 26 | 14 | 17 | 26 | -0,5 | -0,1 | -0,4 |
| chr1:35581459-35581959:downstream_17426 | chr1 | 35525386 | 35581459 | + | ZMYM1 | NM | 26 | 0,3 | 1 | 0 | 7,65E-01 | NA | 18 | 17 | 57 | 30 | 11 | 10 | 32 | 30 | -1,3 | -1,6 | 0,3 |
| chr6:149732749-149733249:downstream_25290 | chr6 | 149639059 | 149732749 | + | TAB2 | NM | 11 | -0,5 | 2 | 0 | 7,65E-01 | NA | 2 | 0 | 7 | 7 | 11 | 2 | 28 | 26 | -2,6 | -2,0 | -0,5 |
| chr19:9758837-9759337:downstream_21673 | chr19 | 9759337 | 9785776 | - | ZNF562 | NM | 23 | -0,3 | 1 | 0 | 7,65E-01 | NA | 6 | 3 | 4 | 4 | 59 | 38 | 42 | 32 | 0,1 | 0,4 | -0,3 |
| chr1:39471737-39472237:downstream_16462 | chr1 | 39456915 | 39471737 | + | AKIRIN1 | NM | 17 | 0,3 | 1 | 0 | 7,66E-01 | NA | 17 | 18 | 17 | 7 | 20 | 24 | 13 | 21 | 0,6 | 0,4 | 0,2 |
| chr19:1021627-1022127:downstream_21379 | chr19 | 1021520 | 1021627 | + | RNU6-2 | NR | 8 | 0,4 | 2 | 0 | 7,67E-01 | NA | 11 | 16 | 6 | 2 | 10 | 11 | 6 | 2 | 1,8 | 1,3 | 0,4 |
| chr7:141922124-141922624:downstream_5647 | chr7 | 141811548 | 141922124 | + | MGAM2 | NM | 11 | -0,4 | 1 | 0 | 7,67E-01 | NA | 2 | 12 | 18 | 8 | 7 | 13 | 15 | 13 | -1,0 | -0,6 | -0,4 |
| chr19:44612479-44612979:downstream_14362 | chr19 | 44598481 | 44612479 | + | ZNF224 | NM | 65 | -0,2 | 1 | 0 | 7,67E-01 | 9,38E-01 | 33 | 24 | 39 | 32 | 99 | 89 | 111 | 94 | -0,3 | -0,1 | -0,2 |
| chr8:19253729-19254229:downstream_26114 | chr8 | 19171080 | 19253729 | + | SH2D4A | NM | 31 | -0,3 | 1 | 0 | 7,67E-01 | 9,38E-01 | 8 | 1 | 29 | 25 | 18 | 15 | 93 | 58 | -2,6 | -2,2 | -0,3 |
| chrX:53457705-53458205:downstream_26856 | chrX | 53458205 | 53461323 | - | HSD17B10 | NM | 29 | 0,3 | 1 | 0 | 7,67E-01 | 9,38E-01 | 13 | 7 | 6 | 7 | 54 | 58 | 39 | 39 | 0,7 | 0,4 | 0,3 |
| chr11:74202422-74202922:downstream_9532 | chr11 | 74202922 | 74204755 | - | LIPT2 | NM | 18 | 0,3 | 1 | 0 | 7,67E-01 | NA | 9 | 3 | 7 | 8 | 24 | 21 | 40 | 31 | -0,4 | -0,7 | 0,3 |
| chr19:59024396-59024896:downstream_14730 | chr19 | 59024896 | 59030921 | - | ZBTB45 | NM | 8 | -0,5 | 2 | 0 | 7,68E-01 | NA | 11 | 8 | 3 | 10 | 1 | 22 | 4 | 7 | 0,6 | 1,1 | -0,5 |
| chr4:139967093-139967593:downstream_3847 | chr4 | 139936912 | 139967093 | + | CCRN4L | NM | 19 | 0,3 | 1 | 0 | 7,68E-01 | NA | 24 | 10 | 7 | 7 | 36 | 33 | 13 | 21 | 1,3 | 1,0 | 0,3 |
| chr5:167946309-167946809:downstream_4565 | chr5 | 167913462 | 167946309 | + | RARS | NM | 26 | 0,3 | 1 | 0 | 7,68E-01 | NA | 7 | 3 | 6 | 5 | 47 | 37 | 47 | 56 | 0,0 | -0,3 | 0,3 |
| chr1:32696760-32697260:downstream_613 | chr1 | 32697260 | 32707311 | - | MTMR9LP | NR | 8 | -0,5 | 2 | 0 | 7,68E-01 | NA | 7 | 3 | 19 | 19 | 0 | 5 | 3 | 9 | -1,9 | -1,3 | -0,6 |
| chr6:143661480-143661980:downstream_24469 | chr6 | 143380499 | 143661480 | + | AIG1 | NM | 27 | 0,3 | 1 | 0 | 7,68E-01 | NA | 2 | 9 | 0 | 5 | 60 | 58 | 31 | 48 | 1,2 | 0,6 | 0,6 |
| chr20:30922811-30923311:downstream_14962 | chr20 | 30865453 | 30922811 | + | KIF3B | NM | 11 | -0,4 | 1 | 0 | 7,69E-01 | NA | 5 | 4 | 6 | 12 | 11 | 12 | 14 | 21 | -1,0 | -0,7 | -0,4 |
| chr5:153800543-153801043:downstream_4114 | chr5 | 153570294 | 153800543 | + | GALNT10 | NM | 39 | -0,3 | 1 | 0 | 7,69E-01 | 9,38E-01 | 7 | 0 | 18 | 16 | 40 | 19 | 89 | 121 | -2,2 | -1,8 | -0,4 |
| chr4:153241909-153242409:downstream_23649 | chr4 | 153242409 | 153456393 | - | FBXW7 | NM | 25 | 0,3 | 1 | 0 | 7,69E-01 | NA | 36 | 56 | 6 | 8 | 44 | 34 | 4 | 10 | 2,7 | 2,5 | 0,3 |
| chr14:68117366-68117866:downstream_11196 | chr14 | 68117866 | 68141602 | - | VTI1B | NM | 37 | -0,2 | 1 | 0 | 7,70E-01 | 9,38E-01 | 37 | 17 | 33 | 25 | 50 | 47 | 31 | 57 | -0,1 | 0,2 | -0,3 |
| chr10:121588415-121588915:downstream_17647 | chr10 | 121588915 | 121633140 | - | MCMBP | NM | 9 | 0,4 | 1 | 0 | 7,70E-01 | NA | 15 | 3 | 7 | 10 | 4 | 8 | 11 | 0,4 | -0,1 | -0,5 | 0,4 |
| chr15:44855001-44855501:downstream_19538 | chr15 | 44829265 | 44855001 | + | EIF3J | NM | 31 | -0,3 | 1 | 0 | 7,70E-01 | 9,38E-01 | 26 | 22 | 10 | 21 | 35 | 72 | 29 | 30 | 0,6 | 0,9 | -0,2 |
| chr2:65498390-65498890:downstream_21720 | chr2 | 65454828 | 65498390 | + | ACTR2 | NM | 76 | 0,3 | 1 | 0 | 7,70E-01 | 9,38E-01 | 4 | 3 | 10 | 7 | 95 | 56 | 228 | 203 | -1,3 | -1,5 | 0,2 |
| chr8:29190078-29190578:downstream_25912 | chr8 | 29190578 | 29208267 | - | DUSP4 | NM | 54 | -0,2 | 1 | 0 | 7,71E-01 | 9,38E-01 | 24 | 9 | 32 | 27 | 65 | 71 | 88 | 116 | -0,8 | -0,6 | -0,3 |
| chr11:35017675-35018175:downstream_18196 | chr11 | 34937676 | 35017675 | + | PDHX | NM | 20 | 0,3 | 1 | 0 | 7,71E-01 | NA | 5 | 4 | 18 | 11 | 56 | 8 | 54 | 43 | -1,7 | -2,0 | 0,3 |
| chr1:153590775-153591275:downstream_17209 | chr1 | 153591275 | 153606568 | - | S100A13 | NM | 104 | -0,6 | 2 | 0 | 7,71E-01 | 9,38E-01 | 0 | 11 | 1 | 4 | 348 | 251 | 111 | 110 | 1,0 | 1,4 | -0,5 |
| chr6:34213656-34214156:downstream_24535 | chr6 | 34214156 | 34216885 | - | C6orf1 | NM | 18 | 0,3 | 1 | 0 | 7,71E-01 | NA | 26 | 30 | 22 | 19 | 10 | 15 | 14 | 9 | 0,4 | 0,1 | 0,3 |
| chr1:9789172-9789672:downstream_425 | chr1 | 9711789 | 9789172 | + | PIK3CD | NM | 8 | 0,5 | 2 | 0 | 7,71E-01 | NA | 1 | 7 | 6 | 9 | 4 | 2 | 6 | 11 | -0,9 | -1,5 | 0,6 |
| chr1:231136479-231136979:downstream_1418 | chr1 | 231114822 | 231136479 | + | ARV1 | NM | 63 | 0,3 | 1 | 0 | 7,72E-01 | 9,38E-01 | 7 | 6 | 1 | 7 | 118 | 150 | 107 | 109 | 0,7 | 0,3 | 0,4 |
| chr7:149243744-149244244:downstream_25830 | chr7 | 149244244 | 149321881 | - | ZNF767P | NR | 8 | -0,4 | 2 | 0 | 7,72E-01 | NA | 9 | 2 | 14 | 14 | 7 | 2 | 11 | 7 | -1,4 | -0,9 | -0,4 |
| chr12:116974318-116974818:downstream_18637 | chr12 | 116971226 | 116974318 | + | LINC00173 | NR | 10 | -0,5 | 2 | 0 | 7,72E-01 | NA | 2 | 0 | 7 | 9 | 7 | 4 | 26 | 21 | -2,7 | -2,2 | -0,5 |
| chr11:76493124-76493624:downstream_18100 | chr11 | 76493624 | 76494106 | - | LOC101928837 | NR | 62 | 0,2 | 1 | 0 | 7,72E-01 | 9,38E-01 | 22 | 33 | 43 | 16 | 101 | 71 | 125 | 87 | -0,1 | -0,3 | 0,2 |
| chr20:61579827-61580327:downstream_15143 | chr20 | 61569440 | 61579827 | + | GID8 | NM | 8 | -0,5 | 2 | 0 | 7,72E-01 | NA | 11 | 0 | 6 | 2 | 14 | 16 | 13 | 3 | 0,4 | 0,9 | -0,5 |
| chr1:23607301-23607801:downstream_542 | chr1 | 23607801 | 23612011 | - | LINC01355 | NR | 9 | 0,5 | 2 | 0 | 7,72E-01 | NA | 0 | 3 | 10 | 2 | 6 | 2 | 17 | 30 | -2,1 | -2,6 | 0,5 |
| chrX:134167575-134168075:downstream_7939 | chrX | 134166332 | 134167575 | + | FAM127A | NM | 45 | -0,2 | 1 | 0 | 7,72E-01 | 9,38E-01 | 29 | 11 | 19 | 9 | 91 | 93 | 47 | 58 | 0,5 | 0,8 | -0,3 |
| chr14:23356889-23357389:downstream_11058 | chr14 | 23352431 | 23356889 | + | REM2 | NM | 14 | -0,4 | 1 | 0 | 7,73E-01 | NA | 11 | 2 | 3 | 2 | 24 | 49 | 10 | 11 | 1,3 | 1,8 | -0,5 |
| chr21:34949358-34949858:downstream_15301 | chr21 | 34949858 | 34961014 | - | DONSON | NM | 8 | 0,4 | 2 | 0 | 7,73E-01 | NA | 5 | 6 | 3 | 8 | 13 | 3 | 11 | 12 | 0,0 | -0,6 | 0,5 |
| chr21:18965897-18966397:downstream_22645 | chr21 | 18885223 | 18965897 | + | CXADR | NM | 13 | 0,4 | 1 | 0 | 7,74E-01 | NA | 7 | 7 | 11 | 2 | 24 | 12 | 15 | 25 | 0,1 | -0,2 | 0,3 |
| chr15:66628507-66629007:downstream_19808 | chr15 | 66629007 | 66649054 | - | TIPIN | NM | 32 | 0,2 | 1 | 0 | 7,74E-01 | 9,38E-01 | 30 | 46 | 31 | 26 | 39 | 25 | 29 | 27 | 0,4 | 0,2 | 0,2 |
| chr10:60049019-60049519:downstream_8284 | chr10 | 60028861 | 60049019 | + | CISD1 | NM | 9 | 0,4 | 1 | 0 | 7,74E-01 | NA | 11 | 21 | 7 | 11 | 9 | 6 | 4 | 7 | 0,9 | 0,5 | 0,4 |
| chr2:42559688-42560188:downstream_21898 | chr2 | 42396477 | 42559688 | + | EML4 | NM | 159 | -0,3 | 1 | 0 | 7,74E-01 | 9,38E-01 | 2 | 0 | 10 | 13 | 50 | 59 | 655 | 482 | -3,8 | -3,4 | -0,4 |
| chr17:33332088-33332588:downstream_20540 | chr17 | 33307516 | 33332088 | + | LIG3 | NM | 28 | -0,3 | 1 | 0 | 7,75E-01 | 9,38E-01 | 38 | 28 | 42 | 23 | 18 | 35 | 18 | 24 | 0,0 | 0,3 | -0,3 |
| chr15:49172380-49172880:downstream_11716 | chr15 | 49170289 | 49172380 | + | EID1 | NM | 17 | 0,3 | 1 | 0 | 7,75E-01 | NA | 8 | 16 | 3 | 6 | 28 | 44 | 11 | 21 | 1,5 | 1,1 | 0,4 |
| chr5:76361058-76361558:downstream_4310 | chr5 | 76326209 | 76361058 | + | AGGF1 | NM | 8 | -0,4 | 1 | 0 | 7,75E-01 | NA | 5 | 5 | 8 | 8 | 11 | 6 | 15 | 7 | -0,8 | -0,4 | -0,4 |
| chr7:99689822-99690322:downstream_5981 | chr7 | 99686582 | 99689822 | + | COPS6 | NM | 39 | 0,3 | 1 | 0 | 7,75E-01 | 9,38E-01 | 19 | 9 | 6 | 10 | 56 | 104 | 46 | 64 | 0,8 | 0,5 | 0,3 |
| chr12:125510349-125510849:downstream_10499 | chr12 | 125478193 | 125510349 | + | BRI3BP | NM | 89 | 0,2 | 1 | 0 | 7,76E-01 | 9,38E-01 | 47 | 49 | 102 | 93 | 72 | 58 | 157 | 135 | -1,0 | -1,2 | 0,2 |
| chr12:117150624-117151124:downstream_10435 | chr12 | 117151124 | 117175875 | - | C12orf49 | NM | 9 | 0,4 | 1 | 0 | 7,77E-01 | NA | 11 | 3 | 8 | 3 | 7 | 16 | 13 | 13 | 0,2 | -0,1 | 0,3 |
| chr11:82967141-82967641:downstream_17863 | chr11 | 82904780 | 82967141 | + | ANKRD42 | NM | 4 | 0,7 | 3 | 0 | 7,77E-01 | NA | 5 | 9 | 0 | 1 | 2 | 15 | 1 | 1 | 4,0 | 3,0 | 1,1 |
| chr3:183911795-183912295:downstream_3297 | chr3 | 183903862 | 183911795 | + | ABCF3 | NM | 7 | 0,5 | 2 | 0 | 7,77E-01 | NA | 10 | 0 | 8 | 7 | 3 | 6 | 8 | 11 | -0,7 | -1,1 | 0,4 |
| chr19:51228981-51229481:downstream_14513 | chr19 | 51226604 | 51228981 | + | CLEC11A | NM | 8 | 0,4 | 2 | 0 | 7,77E-01 | NA | 17 | 8 | 3 | 2 | 17 | 9 | 4 | 3 | 2,2 | 1,8 | 0,5 |
| chr7:130561005-130561505:downstream_6094 | chr7 | 130561505 | 130561569 | - | MIR29A | NR | 26 | 0,3 | 1 | 0 | 7,78E-01 | NA | 12 | 10 | 24 | 23 | 14 | 26 | 56 | 45 | -1,1 | -1,3 | 0,2 |
| chr1:110043063-110043563:downstream_16637 | chr1 | 110036657 | 110043063 | + | CYB561D1 | NM | 7 | -0,5 | 2 | 0 | 7,78E-01 | NA | 7 | 1 | 7 | 2 | 11 | 11 | 7 | 8 | -0,1 | 0,5 | -0,6 |
| chr15:63615229-63615729:downstream_19489 | chr15 | 63615729 | 63674309 | - | CA12 | NM | 1365 | -0,1 | 0 | 0 | 7,78E-01 | 9,39E-01 | 73 | 99 | 145 | 91 | 2196 | 2496 | 2796 | 3027 | -0,5 | -0,3 | -0,1 |
| chr19:1981336-1981836:downstream_13710 | chr19 | 1941160 | 1981336 | + | CSNK1G2 | NM | 6 | -0,5 | 2 | 0 | 7,78E-01 | NA | 4 | 5 | 8 | 7 | 7 | 2 | 7 | 4 | -0,8 | -0,4 | -0,4 |
| chr13:110405683-110406183:downstream_10874 | chr13 | 110406183 | 110438914 | - | IRS2 | NM | 32 | -0,3 | 1 | 0 | 7,79E-01 | 9,39E-01 | 2 | 0 | 7 | 7 | 16 | 32 | 90 | 103 | -2,5 | -2,0 | -0,5 |
| chr12:125400205-125400705:downstream_10497 | chr12 | 125400092 | 125400205 | + | MIR5188 | NR | 26 | -0,3 | 1 | 0 | 7,79E-01 | NA | 19 | 31 | 3 | 16 | 50 | 55 | 28 | 8 | 1,4 | 1,5 | -0,2 |
| chr20:13694468-13694968:downstream_22422 | chr20 | 13694968 | 13765579 | - | ESF1 | NM | 29 | -0,3 | 1 | 0 | 7,79E-01 | 9,39E-01 | 9 | 6 | 24 | 9 | 30 | 36 | 77 | 40 | -1,1 | -0,8 | -0,3 |
| chrX:100263833-100264333:downstream_27085 | chrX | 100264333 | 100307105 | - | TRMT2B | NM | 23 | 0,3 | 1 | 0 | 7,79E-01 | NA | 20 | 10 | 47 | 30 | 6 | 14 | 38 | 23 | -1,4 | -1,7 | 0,3 |
| chr1:92853732-92854232:downstream_178 | chr1 | 92764521 | 92853732 | + | RPAP2 | NM | 13 | 0,3 | 1 | 0 | 7,80E-01 | NA | 7 | 14 | 10 | 15 | 15 | 8 | 25 | 12 | -0,3 | -0,7 | 0,4 |
| chr4:78873944-78874444:downstream_3752 | chr4 | 78783804 | 78873944 | + | MRPL1 | NM | 9 | 0,4 | 1 | 0 | 7,81E-01 | NA | 2 | 2 | 3 | 7 | 6 | 9 | 24 | 19 | -1,1 | -1,6 | 0,5 |
| chr5:131527803-131528303:downstream_24267 | chr5 | 131528303 | 131563556 | - | P4HA2 | NM | 35 | -0,2 | 1 | 0 | 7,81E-01 | 9,40E-01 | 11 | 25 | 19 | 11 | 59 | 65 | 47 | 40 | 0,2 | 0,5 | -0,3 |
| chr10:73575554-73576054:downstream_17732 | chr10 | 73576054 | 73611082 | - | PSAP | NM | 507 | 0,2 | 1 | 0 | 7,81E-01 | 9,40E-01 | 8 | 11 | 3 | 10 | 948 | 1217 | 916 | 942 | 0,6 | 0,2 | 0,4 |
| chr17:53469473-53469973:downstream_12572 | chr17 | 53469973 | 53499341 | - | MMD | NM | 61 | 0,3 | 1 | 0 | 7,81E-01 | 9,40E-01 | 6 | 0 | 10 | 7 | 52 | 55 | 168 | 186 | -1,6 | -1,7 | 0,1 |
| chr8:122624770-122625270:downstream_6696 | chr8 | 122625270 | 122653630 | - | HAS2 | NM | 3 | -0,9 | 3 | 0 | 7,81E-01 | NA | 0 | 0 | 6 | 11 | 0 | 0 | 1 | 7 | -Inf | -Inf | NaN |
| chr2:85764089-85764589:downstream_2082 | chr2 | 85764589 | 85766009 | - | PARTICL | NR | 17 | 0,3 | 1 | 0 | 7,81E-01 | NA | 11 | 15 | 7 | 5 | 31 | 31 | 24 | 11 | 1,1 | 0,8 | 0,3 |
| chr19:58082433-58082933:downstream_14702 | chr19 | 58082933 | 58090243 | - | ZNF416 | NM | 10 | 0,4 | 1 | 0 | 7,81E-01 | NA | 6 | 8 | 4 | 2 | 20 | 15 | 11 | 11 | 1,0 | 0,7 | 0,3 |
| chr10:12209072-12209572:downstream_8417 | chr10 | 12209572 | 12238143 | - | NUDT5 | NM | 107 | 0,2 | 1 | 0 | 7,82E-01 | 9,40E-01 | 59 | 44 | 47 | 39 | 158 | 187 | 164 | 153 | 0,3 | 0,1 | 0,1 |
| chr15:79190081-79190581:downstream_19635 | chr15 | 79165122 | 79190081 | + | MORF4L1 | NM | 18 | 0,3 | 1 | 0 | 7,83E-01 | NA | 15 | 15 | 22 | 22 | 14 | 13 | 17 | 31 | -0,6 | -0,9 | 0,3 |
| chrX:46407910-46408410:downstream_7690 | chrX | 46404924 | 46407910 | + | ZNF674-AS1 | NR | 9 | 0,5 | 2 | 0 | 7,84E-01 | NA | 16 | 2 | 0 | 5 | 13 | 23 | 6 | 9 | 1,9 | 1,3 | 0,6 |
| chr12:123956909-123957409:downstream_10486 | chr12 | 123942650 | 123956909 | + | SNRNP35 | NR | 5 | 0,6 | 2 | 0 | 7,84E-01 | NA | 7 | 6 | 3 | 3 | 0 | 13 | 0 | 9 | 1,0 | 0,5 | 0,5 |
| chr16:71964540-71965040:downstream_20031 | chr16 | 71928310 | 71964540 | + | IST1 | NM | 26 | 0,3 | 1 | 0 | 7,84E-01 | NA | 16 | 14 | 10 | 2 | 67 | 49 | 29 | 24 | 1,3 | 1,1 | 0,1 |
| chr6:159397765-159398265:downstream_5421 | chr6 | 159398265 | 159421198 | - | RSPH3 | NM | 14 | -0,4 | 1 | 0 | 7,84E-01 | NA | 13 | 0 | 13 | 7 | 23 | 15 | 18 | 22 | -0,6 | -0,1 | -0,4 |
| chr1:196763203-196763703:downstream_16593 | chr1 | 196743929 | 196763203 | + | CFHR3 | NM | 7 | -0,4 | 2 | 0 | 7,85E-01 | NA | 5 | 5 | 4 | 2 | 6 | 21 | 6 | 7 | 0,6 | 1,1 | -0,5 |
| chr6:163833596-163834096:downstream_5431 | chr6 | 163834096 | 163834982 | - | CAHM | NR | 9 | -0,4 | 2 | 0 | 7,85E-01 | NA | 1 | 10 | 3 | 7 | 11 | 18 | 11 | 10 | 0,2 | 0,5 | -0,3 |
| chr7:92166823-92167323:downstream_5943 | chr7 | 92158086 | 92166823 | + | RBM48 | NM | 18 | -0,3 | 1 | 0 | 7,86E-01 | NA | 11 | 10 | 1 | 7 | 41 | 44 | 18 | 14 | 1,3 | 1,4 | -0,2 |
| chr1:111982011-111982511:downstream_900 | chr1 | 111982511 | 111991830 | - | WDR77 | NM | 34 | -0,2 | 1 | 0 | 7,86E-01 | 9,44E-01 | 61 | 77 | 22 | 11 | 44 | 43 | 14 | 4 | 2,1 | 2,3 | -0,2 |
| chr14:75476294-75476794:downstream_11230 | chr14 | 75469611 | 75476294 | + | EIF2B2 | NM | 118 | -0,2 | 1 | 0 | 7,86E-01 | 9,44E-01 | 15 | 10 | 8 | 4 | 295 | 345 | 116 | 148 | 1,0 | 1,3 | -0,3 |
| chr1:169761169-169761669:downstream_1219 | chr1 | 169761669 | 169764061 | - | METTL18 | NM | 23 | 0,3 | 1 | 0 | 7,87E-01 | NA | 10 | 3 | 0 | 3 | 66 | 53 | 21 | 25 | 1,9 | 1,4 | 0,6 |

| | | | | | | | | | | | | | | | | | | | | | | | |
|---|---|---|---|---|---|---|---|---|---|---|---|---|---|---|---|---|---|---|---|---|---|---|---|
| chr1:67464514-67465014:downstream_152 | chr1 | 67465014 | 67520080 | - | SLC35D1 | NM | 8 | -0,4 | 1 | 0 | 7,87E-01 | NA | 12 | 6 | 13 | 4 | 7 | 12 | 4 | 8 | 0,1 | 0,6 | -0,5 |
| chr8:38070819-38071319:downstream_25861 | chr8 | 38034105 | 38070819 | + | BAG4 | NM | 12 | 0,4 | 1 | 0 | 7,87E-01 | NA | 6 | 1 | 10 | 7 | 9 | 10 | 21 | 34 | -1,3 | -1,6 | 0,3 |
| chr18:21741564-21742064:downstream_20839 | chr18 | 21718919 | 21741564 | + | CABYR | NM | 29 | 0,3 | 1 | 0 | 7,87E-01 | 9,44E-01 | 46 | 35 | 4 | 7 | 64 | 56 | 14 | 8 | 2,8 | 2,4 | 0,4 |
| chr7:102992676-102993176:downstream_25727 | chr7 | 102993176 | 103086624 | - | SLC26A5 | NM | 7 | -0,6 | 2 | 0 | 7,87E-01 | NA | 4 | 23 | 1 | 2 | 3 | 22 | 0 | 2 | 3,2 | 4,0 | -0,8 |
| chr1:118068320-118068820:downstream_202 | chr1 | 117910084 | 118068320 | + | MAN1A2 | NM | 25 | -0,3 | 1 | 0 | 7,87E-01 | NA | 5 | 9 | 47 | 48 | 12 | 1 | 40 | 38 | -2,8 | -2,6 | -0,2 |
| chr3:65339405-65339905:downstream_23259 | chr3 | 65339905 | 66024509 | - | MAGI1 | NM | 11 | -0,6 | 2 | 0 | 7,88E-01 | NA | 1 | 0 | 18 | 6 | 3 | 0 | 36 | 25 | -4,9 | -4,2 | -0,6 |
| chr11:35642421-35642921:downstream_9205 | chr11 | 35639734 | 35642421 | + | FJX1 | NM | 23 | 0,3 | 1 | 0 | 7,89E-01 | NA | 9 | 11 | 0 | 3 | 68 | 62 | 13 | 20 | 2,6 | 2,0 | 0,6 |
| chr9:125085742-125086242:downstream_26400 | chr9 | 125026881 | 125085742 | + | MRRF | NM | 18 | -0,3 | 1 | 0 | 7,89E-01 | NA | 8 | 5 | 4 | 14 | 30 | 23 | 22 | 40 | -0,5 | -0,2 | -0,3 |
| chr19:47340914-47341414:downstream_21000 | chr19 | 47341414 | 47354252 | - | AP2S1 | NM | 7 | -0,4 | 2 | 0 | 7,89E-01 | NA | 3 | 2 | 6 | 7 | 11 | 3 | 8 | 18 | -1,3 | -0,9 | -0,4 |
| chr4:178284092-178284592:downstream_3396 | chr4 | 178230990 | 178284092 | + | NEIL3 | NM | 12 | -0,8 | 3 | 0 | 7,89E-01 | NA | 3 | 0 | 3 | 16 | 17 | 0 | 33 | 21 | -2,5 | -1,7 | -0,8 |
| chr1:162499419-162499919:downstream_17382 | chr1 | 162466963 | 162499419 | + | UHMK1 | NM | 18 | 0,3 | 1 | 0 | 7,90E-01 | NA | 6 | 12 | 14 | 2 | 36 | 19 | 29 | 29 | 0,2 | -0,1 | 0,3 |
| chr12:58161850-58162350:downstream_18677 | chr12 | 58162350 | 58165914 | - | METTL1 | NM | 12 | 0,3 | 1 | 0 | 7,90E-01 | NA | 15 | 10 | 10 | 2 | 19 | 16 | 8 | 12 | 1,0 | 0,8 | 0,2 |
| chr5:76372031-76372531:downstream_4311 | chr5 | 76372531 | 76383030 | - | ZBED3 | NM | 9 | 0,4 | 1 | 0 | 7,90E-01 | NA | 7 | 11 | 10 | 11 | 6 | 7 | 11 | 8 | -0,3 | -0,6 | 0,4 |
| chr11:61214239-61214739:downstream_8890 | chr11 | 61197596 | 61214239 | + | SDHAF2 | NM | 22 | -0,2 | 1 | 0 | 7,90E-01 | NA | 27 | 27 | 26 | 26 | 23 | 15 | 21 | 11 | 0,0 | 0,2 | -0,2 |
| chr6:133822889-133823389:downstream_4800 | chr6 | 133823389 | 134210144 | - | TARID | NR | 16 | 0,3 | 1 | 0 | 7,90E-01 | NA | 3 | 8 | 24 | 25 | 6 | 5 | 28 | 31 | -2,2 | -2,5 | 0,3 |
| chr4:71680998-71681498:downstream_23679 | chr4 | 71681498 | 71705627 | - | GRSF1 | NM | 11 | 0,3 | 1 | 0 | 7,91E-01 | NA | 18 | 11 | 13 | 13 | 10 | 8 | 7 | 12 | 0,2 | -0,1 | 0,3 |
| chr6:3268300-3268800:downstream_25084 | chr6 | 3259161 | 3268300 | + | PSMG4 | NM | 16 | 0,3 | 1 | 0 | 7,91E-01 | NA | 11 | 5 | 8 | 2 | 35 | 25 | 24 | 21 | 0,6 | 0,4 | 0,2 |
| chr1:153946244-153946744:downstream_1076 | chr1 | 153946744 | 153950451 | - | JTB | NM | 160 | -0,2 | 1 | 0 | 7,91E-01 | 9,48E-01 | 5 | 2 | 3 | 7 | 289 | 280 | 360 | 339 | -0,5 | -0,3 | -0,2 |
| chr12:15772574-15773074:downstream_9830 | chr12 | 15773074 | 15942510 | - | EPS8 | NM | 67 | -0,3 | 1 | 0 | 7,91E-01 | 9,48E-01 | 2 | 1 | 8 | 9 | 36 | 44 | 255 | 179 | -2,7 | -2,4 | -0,3 |
| chr20:43356452-43356952:downstream_15027 | chr20 | 43343884 | 43356452 | + | WISP2 | NM | 7 | -0,4 | 2 | 0 | 7,92E-01 | NA | 2 | 7 | 3 | 2 | 17 | 15 | 6 | 6 | 1,1 | 1,5 | -0,5 |
| chr6:142767403-142767903:downstream_24463 | chr6 | 142623055 | 142767403 | + | ADGRG6 | NM | 69 | -0,3 | 1 | 0 | 7,94E-01 | 9,49E-01 | 2 | 1 | 8 | 11 | 46 | 30 | 231 | 224 | -2,9 | -2,6 | -0,3 |
| chr17:60019465-60019965:downstream_12670 | chr17 | 60019965 | 60142643 | - | MED13 | NM | 17 | -0,3 | 1 | 0 | 7,94E-01 | NA | 8 | 0 | 13 | 16 | 12 | 15 | 36 | 38 | -1,8 | -1,5 | -0,4 |
| chr3:72423243-72423743:downstream_2650 | chr3 | 72423743 | 72495774 | - | RYBP | NM | 12 | -0,3 | 1 | 0 | 7,94E-01 | NA | 4 | 2 | 6 | 7 | 11 | 18 | 31 | 19 | -1,1 | -0,8 | -0,3 |
| chr4:38824828-38825328:downstream_3655 | chr4 | 38825328 | 38858438 | - | TLR6 | NM | 12 | -0,3 | 1 | 0 | 7,96E-01 | NA | 11 | 9 | 19 | 25 | 5 | 8 | 18 | 6 | -1,2 | -0,9 | -0,3 |
| chr12:120878545-120879045:downstream_10448 | chr12 | 120875892 | 120878545 | + | COX6A1 | NM | 115 | -0,3 | 1 | 0 | 7,96E-01 | 9,49E-01 | 3 | 6 | 4 | 2 | 265 | 341 | 125 | 175 | 0,6 | 1,0 | -0,4 |
| chr16:11927554-11928054:downstream_12195 | chr16 | 11928054 | 11945442 | - | RSL1D1 | NM | 40 | 0,2 | 1 | 0 | 7,96E-01 | 9,49E-01 | 24 | 45 | 4 | 7 | 99 | 98 | 22 | 18 | 2,6 | 2,3 | 0,3 |
| chr9:2803654-2804154:downstream_6967 | chr9 | 2804154 | 2844130 | - | KIAA0020 | NM | 53 | 0,2 | 1 | 0 | 7,96E-01 | 9,49E-01 | 52 | 35 | 51 | 31 | 68 | 53 | 70 | 60 | 0,1 | -0,1 | 0,2 |
| chr1:173855774-173856274:downstream_17416 | chr1 | 173837492 | 173855774 | + | ZBTB37 | NM | 26 | -0,2 | 1 | 0 | 7,96E-01 | NA | 10 | 19 | 51 | 43 | 14 | 9 | 31 | 32 | -1,7 | -1,5 | -0,2 |
| chr3:145786727-145787227:downstream_23347 | chr3 | 145787227 | 145879282 | - | PLOD2 | NM | 62 | -0,2 | 1 | 0 | 7,96E-01 | 9,49E-01 | 7 | 12 | 15 | 18 | 74 | 100 | 127 | 146 | -0,8 | -0,7 | -0,2 |
| chr1:94350255-94350755:downstream_16763 | chr1 | 94350755 | 94375154 | - | GCLM | NM | 30 | 0,2 | 1 | 0 | 7,97E-01 | 9,49E-01 | 10 | 8 | 18 | 13 | 28 | 35 | 67 | 61 | -0,8 | -1,0 | 0,2 |
| chr2:75196859-75197359:downstream_2054 | chr2 | 75185774 | 75196859 | + | POLE4 | NM | 14 | 0,3 | 1 | 0 | 7,97E-01 | NA | 9 | 15 | 3 | 7 | 31 | 19 | 8 | 16 | 1,4 | 1,0 | 0,4 |
| chr19:2478257-2478757:downstream_13872 | chr19 | 2476122 | 2478257 | + | GADD45B | NM | 57 | -0,2 | 1 | 0 | 7,97E-01 | 9,49E-01 | 8 | 10 | 6 | 8 | 124 | 128 | 92 | 84 | 0,4 | 0,5 | -0,2 |
| chr2:206979796-206980296:downstream_2384 | chr2 | 206980296 | 206981296 | - | GCSHP3 | NR | 27 | -0,2 | 1 | 0 | 7,97E-01 | NA | 7 | 16 | 43 | 24 | 12 | 25 | 47 | 41 | -1,5 | -1,2 | -0,3 |
| chr14:23369353-23369853:downstream_19361 | chr14 | 23369853 | 23388396 | - | RBM23 | NM | 13 | -0,3 | 1 | 0 | 7,97E-01 | NA | 2 | 15 | 15 | 16 | 19 | 5 | 17 | 18 | -0,9 | -0,6 | -0,3 |
| chr6:43608688-43609188:downstream_24877 | chr6 | 43597278 | 43608688 | + | MAD2L1BP | NM | 28 | -0,3 | 1 | 0 | 7,98E-01 | NA | 4 | 6 | 4 | 1 | 71 | 82 | 31 | 25 | 1,0 | 1,5 | -0,5 |
| chr12:90105729-90106229:downstream_10346 | chr12 | 90102731 | 90105729 | + | LINC00936 | NR | 8 | -0,4 | 2 | 0 | 7,98E-01 | NA | 15 | 9 | 7 | 3 | 12 | 8 | 4 | 2 | 1,2 | 1,6 | -0,4 |
| chr12:27848497-27848997:downstream_18753 | chr12 | 27677044 | 27848497 | + | PPFIBP1 | NM | 6 | 0,5 | 2 | 0 | 7,98E-01 | NA | 2 | 0 | 10 | 8 | 2 | 0 | 10 | 16 | -2,9 | -3,4 | 0,5 |
| chr6:43638433-43638933:downstream_25023 | chr6 | 43638933 | 43655549 | - | MRPS18A | NM | 15 | 0,3 | 1 | 0 | 7,98E-01 | NA | 16 | 33 | 7 | 10 | 27 | 10 | 13 | 4 | 1,5 | 1,1 | 0,4 |
| chr10:13672868-13673368:downstream_8202 | chr10 | 13628938 | 13672868 | + | PRPF18 | NM | 27 | -0,2 | 1 | 0 | 7,98E-01 | NA | 45 | 27 | 29 | 39 | 33 | 11 | 22 | 13 | 0,1 | 0,3 | -0,2 |
| chr14:74429813-74430313:downstream_19187 | chr14 | 74416636 | 74429813 | + | COQ6 | NM | 5 | -0,4 | 2 | 0 | 7,98E-01 | NA | 4 | 3 | 8 | 6 | 6 | 6 | 7 | -0,5 | -0,1 | -0,4 |
| chr11:117065828-117066328:downstream_9669 | chr11 | 117066328 | 117072630 | - | LOC100652768 | NR | 8 | 0,5 | 2 | 0 | 7,99E-01 | NA | 2 | 17 | 4 | 2 | 5 | 22 | 8 | 2 | 1,8 | 1,3 | 0,4 |
| chr2:65314142-65314642:downstream_1998 | chr2 | 65283494 | 65314142 | + | CEP68 | NM | 14 | -0,3 | 1 | 0 | 8,00E-01 | NA | 7 | 15 | 18 | 10 | 11 | 17 | 18 | 11 | -0,3 | 0,0 | -0,3 |
| chr19:6751672-6752172:downstream_13931 | chr19 | 6752172 | 6767523 | - | SH2D3A | NM | 6 | 0,5 | 2 | 0 | 8,00E-01 | NA | 7 | 3 | 1 | 3 | 2 | 18 | 6 | 7 | 1,1 | 0,6 | 0,5 |
| chr19:36604110-36604610:downstream_14241 | chr19 | 36604610 | 36606206 | - | POLR2I | NM | 21 | 0,3 | 1 | 0 | 8,01E-01 | NA | 7 | 3 | 1 | 4 | 33 | 57 | 31 | 33 | 0,9 | 0,5 | 0,4 |
| chr10:1063708-1064208:downstream_8199 | chr10 | 1034348 | 1063708 | + | GTPBP4 | NM | 84 | -0,2 | 1 | 0 | 8,01E-01 | 9,53E-01 | 20 | 17 | 25 | 16 | 168 | 133 | 154 | 134 | -0,1 | 0,1 | -0,2 |
| chr17:19286857-19287357:downstream_20569 | chr17 | 19281033 | 19286857 | + | MAPK7 | NM | 6 | -0,4 | 2 | 0 | 8,01E-01 | NA | 2 | 3 | 7 | 5 | 8 | 4 | 11 | 9 | -1,2 | -0,7 | -0,4 |
| chr9:5128183-5128683:downstream_6833 | chr9 | 4985244 | 5128183 | + | JAK2 | NM | 6 | -0,5 | 2 | 0 | 8,01E-01 | NA | 2 | 0 | 8 | 5 | 4 | 13 | 19 | -2,3 | -1,8 | -0,5 |
| chr7:38763042-38763542:downstream_25802 | chr7 | 38763542 | 38948800 | - | VPS41 | NM | 10 | 0,4 | 2 | 0 | 8,02E-01 | NA | 2 | 2 | 15 | 16 | 3 | 1 | 18 | 21 | -2,8 | -3,2 | 0,4 |
| chr6:114184652-114185152:downstream_5307 | chr6 | 114178513 | 114184652 | + | MARCKS | NM | 29 | 0,3 | 1 | 0 | 8,02E-01 | 9,53E-01 | 4 | 2 | 6 | 8 | 28 | 28 | 65 | 90 | -1,2 | -1,5 | 0,3 |
| chr11:126215648-126216148:downstream_9743 | chr11 | 126173646 | 126215648 | + | DCPS | NM | 8 | -0,4 | 2 | 0 | 8,02E-01 | NA | 3 | 2 | 6 | 7 | 3 | 13 | 11 | 16 | -1,2 | -0,8 | -0,4 |
| chr10:32093825-32094325:downstream_17476 | chr10 | 32094325 | 32217804 | - | ARHGAP12 | NM | 7 | 0,4 | 2 | 0 | 8,03E-01 | NA | 2 | 4 | 10 | 13 | 1 | 4 | 13 | 10 | -1,9 | -2,3 | 0,4 |
| chr11:8017719-8018219:downstream_9129 | chr11 | 8008866 | 8017719 | + | EIF3F | NM | 30 | -0,2 | 1 | 0 | 8,03E-01 | 9,54E-01 | 43 | 27 | 31 | 9 | 49 | 40 | 17 | 24 | 0,8 | 1,1 | -0,3 |
| chr2:192698531-192699031:downstream_2359 | chr2 | 192699031 | 192712006 | - | SDPR | NM | 21 | 0,3 | 1 | 0 | 8,03E-01 | NA | 8 | 16 | 4 | 5 | 37 | 58 | 14 | 29 | 1,4 | 1,2 | 0,3 |
| chr10:30750762-30751262:downstream_17642 | chr10 | 30722949 | 30750762 | + | MAP3K8 | NM | 6 | 0,5 | 2 | 0 | 8,03E-01 | NA | 1 | 1 | 15 | 5 | 2 | 0 | 18 | 10 | -3,5 | -4,1 | 0,6 |
| chrX:133674870-133675370:downstream_7935 | chrX | 133675370 | 133675467 | - | MIR542 | NR | 4 | -0,5 | 2 | 0 | 8,03E-01 | NA | 4 | 5 | 4 | 1 | 7 | 7 | 3 | 2 | 0,8 | 1,4 | -0,6 |
| chr19:44471752-44472252:downstream_21616 | chr19 | 44455374 | 44471752 | + | ZNF221 | NM | 6 | 0,4 | 2 | 0 | 8,04E-01 | NA | 2 | 3 | 8 | 4 | 2 | 5 | 13 | 7 | -1,2 | -1,6 | 0,4 |
| chr10:90683259-90683759:downstream_8625 | chr10 | 90639943 | 90683259 | + | STAMBPL1 | NM | 12 | 0,3 | 1 | 0 | 8,04E-01 | NA | 11 | 13 | 11 | 4 | 11 | 21 | 6 | 19 | 0,6 | 0,4 | 0,2 |
| chr7:30412410-30412910:downstream_5774 | chr7 | 30409665 | 30412410 | + | DKFZP586I1420 | NR | 9 | 0,4 | 2 | 0 | 8,05E-01 | NA | 7 | 0 | 7 | 10 | 6 | 5 | 17 | 18 | -1,4 | -1,7 | 0,4 |
| chrX:102842664-102843164:downstream_27068 | chrX | 102831158 | 102842664 | + | TCEAL4 | NM | 26 | 0,2 | 1 | 0 | 8,05E-01 | NA | 33 | 17 | 25 | 15 | 35 | 25 | 25 | 29 | 0,4 | 0,2 | 0,2 |
| chr4:140637045-140637545:downstream_3512 | chr4 | 140637545 | 141075233 | - | MAML3 | NM | 6 | -0,5 | 2 | 0 | 8,05E-01 | NA | 2 | 0 | 7 | 10 | 4 | 0 | 14 | 16 | -3,4 | -2,9 | -0,5 |
| chr8:33355526-33356026:downstream_26169 | chr8 | 33356026 | 33370703 | - | TTI2 | NM | 12 | 0,3 | 1 | 0 | 8,06E-01 | NA | 8 | 15 | 8 | 2 | 14 | 24 | 10 | 11 | 1,1 | 0,9 | 0,2 |
| chr1:84963505-84964005:downstream_802 | chr1 | 84964005 | 84972262 | - | GNG5 | NM | 19 | -0,3 | 1 | 0 | 8,07E-01 | NA | 6 | 6 | 8 | 7 | 32 | 34 | 25 | 30 | 0,0 | 0,3 | -0,3 |
| chr14:73960105-73960605:downstream_11211 | chr14 | 73957638 | 73960105 | + | C14orf169 | NM | 26 | 0,2 | 1 | 0 | 8,07E-01 | NA | 4 | 9 | 6 | 7 | 51 | 30 | 56 | 46 | 0,0 | -0,3 | 0,3 |
| chr7:128142978-128143478:downstream_6075 | chr7 | 128116782 | 128142978 | + | METTL2B | NM | 24 | -0,2 | 1 | 0 | 8,07E-01 | NA | 15 | 3 | 29 | 22 | 24 | 14 | 47 | 39 | -1,5 | -1,2 | -0,3 |
| chr2:96915445-96915945:downstream_22285 | chr2 | 96915945 | 96931751 | - | TMEM127 | NM | 72 | 0,1 | 1 | 0 | 8,07E-01 | 9,58E-01 | 89 | 84 | 45 | 43 | 108 | 95 | 64 | 50 | 1,0 | 0,8 | 0,2 |
| chr21:16333055-16333555:downstream_15182 | chr21 | 16333555 | 16437126 | - | NRIP1 | NM | 34 | -0,2 | 1 | 0 | 8,08E-01 | 9,58E-01 | 10 | 7 | 24 | 34 | 21 | 28 | 71 | 77 | -1,8 | -1,6 | -0,2 |
| chr16:732768-733268:downstream_20227 | chr16 | 730110 | 732768 | + | STUB1 | NM | 39 | 0,2 | 1 | 0 | 8,08E-01 | 9,58E-01 | 10 | 15 | 8 | 9 | 72 | 79 | 60 | 62 | 0,5 | 0,3 | 0,2 |
| chr17:73261809-73262309:downstream_20581 | chr17 | 73262309 | 73267311 | - | MIF4GD | NM | 23 | 0,3 | 1 | 0 | 8,08E-01 | NA | 3 | 8 | 4 | 9 | 24 | 40 | 36 | 61 | -0,3 | -0,6 | 0,3 |
| chr6:136662918-136663418:downstream_24939 | chr6 | 136663418 | 136871957 | - | MAP7 | NM | 17 | 0,3 | 1 | 0 | 8,09E-01 | NA | 7 | 5 | 32 | 36 | 5 | 2 | 25 | 22 | -2,5 | -2,8 | 0,3 |
| chr1:46160108-46160608:downstream_707 | chr1 | 46153846 | 46160108 | + | TMEM69 | NM | 14 | 0,3 | 1 | 0 | 8,09E-01 | NA | 17 | 3 | 4 | 10 | 24 | 17 | 21 | 16 | 0,5 | 0,2 | 0,3 |
| chr15:31230643-31231143:downstream_11631 | chr15 | 31231143 | 31283807 | - | MTMR10 | NM | 12 | 0,4 | 2 | 0 | 8,09E-01 | NA | 2 | 0 | 7 | 9 | 4 | 2 | 29 | 45 | -3,3 | -3,6 | 0,3 |
| chr10:16555744-16556244:downstream_17735 | chr10 | 16478941 | 16555744 | + | PTER | NM | 13 | 0,8 | 3 | 0 | 8,09E-01 | NA | 22 | 0 | 14 | 46 | 4 | 0 | 13 | 7 | -1,4 | -2,2 | 0,8 |
| chr9:4665272-4665772:downstream_6972 | chr9 | 4662293 | 4665272 | + | PPAPDC2 | NM | 41 | -0,2 | 1 | 0 | 8,09E-01 | 9,59E-01 | 21 | 19 | 15 | 20 | 62 | 82 | 46 | 64 | 0,2 | 0,4 | -0,2 |
| chr6:71298606-71299106:downstream_5218 | chr6 | 71276624 | 71298606 | + | SDHAF4 | NM | 24 | 0,3 | 1 | 0 | 8,10E-01 | NA | 6 | 8 | 4 | 2 | 37 | 78 | 35 | 22 | 1,2 | 1,0 | 0,2 |
| chr15:31514470-31514970:downstream_11632 | chr15 | 31514970 | 31523050 | - | LOC283710 | NM | 5 | -0,5 | 2 | 0 | 8,10E-01 | NA | 15 | 2 | 4 | 7 | 2 | 7 | 3 | 2 | 0,6 | 1,1 | -0,4 |
| chr8:125322658-125323158:downstream_6704 | chr8 | 125323158 | 125384940 | - | TMEM65 | NM | 21 | -0,2 | 1 | 0 | 8,10E-01 | NA | 15 | 15 | 38 | 24 | 11 | 15 | 24 | 21 | -1,0 | -0,7 | -0,2 |
| chr14:60601532-60602032:downstream_11165 | chr14 | 60558628 | 60601532 | + | PCNXL4 | NM | 21 | 0,2 | 1 | 0 | 8,11E-01 | NA | 6 | 7 | 18 | 11 | 17 | 15 | 36 | 53 | -1,2 | -1,5 | 0,2 |
| chr7:99704803-99705303:downstream_5985 | chr7 | 99699129 | 99704803 | + | AP4M1 | NM | 9 | -0,3 | 1 | 0 | 8,11E-01 | NA | 3 | 7 | 6 | 5 | 8 | 19 | 14 | 9 | -0,1 | 0,3 | -0,3 |
| chr15:59427667-59428167:downstream_11482 | chr15 | 59428167 | 59665071 | - | MYO1E | NM | 28 | 0,3 | 1 | 0 | 8,11E-01 | NA | 11 | 2 | 33 | 38 | 14 | 4 | 56 | 63 | -2,5 | -2,7 | 0,2 |
| chr10:3178997-3179497:downstream_17706 | chr10 | 3109711 | 3178997 | + | PFKP | NM | 20 | 0,3 | 1 | 0 | 8,11E-01 | NA | 5 | 5 | 3 | 9 | 34 | 20 | 43 | 42 | -0,3 | -0,6 | 0,4 |
| chr11:11980872-11981372:downstream_18386 | chr11 | 11862969 | 11980872 | + | USP47 | NM | 11 | -0,3 | 1 | 0 | 8,12E-01 | NA | 3 | 1 | 11 | 9 | 7 | 6 | 35 | 14 | -2,3 | -1,9 | -0,4 |
| chr19:49103356-49103856:downstream_14448 | chr19 | 49103856 | 49116694 | - | FAM83E | NM | 4 | 0,5 | 2 | 0 | 8,12E-01 | NA | 6 | 11 | 0 | 4 | 2 | 8 | 1 | 2 | 2,0 | 1,4 | 0,6 |
| chr3:196661772-196662272:downstream_23296 | chr3 | 196662272 | 196669464 | - | NCBP2 | NM | 41 | -0,2 | 1 | 0 | 8,12E-01 | 9,61E-01 | 10 | 10 | 17 | 4 | 83 | 72 | 63 | 70 | -0,1 | 0,2 | -0,3 |
| chr18:67997434-67997934:downstream_13665 | chr18 | 67956136 | 67997434 | + | SOCS6 | NM | 31 | 0,2 | 1 | 0 | 8,13E-01 | 9,62E-01 | 13 | 15 | 56 | 54 | 11 | 10 | 39 | 53 | -2,0 | -2,2 | 0,2 |
| chr3:135867259-135867759:downstream_23284 | chr3 | 135867759 | 135914688 | - | MSL2 | NM | 17 | -0,2 | 1 | 0 | 8,13E-01 | NA | 24 | 27 | 18 | 20 | 16 | 15 | 8 | 11 | 0,4 | 0,7 | -0,2 |
| chr19:41891775-41892275:downstream_14317 | chr19 | 41892275 | 41903256 | - | EXOSC5 | NM | 23 | 0,2 | 1 | 0 | 8,14E-01 | NA | 9 | 18 | 8 | 8 | 37 | 43 | 32 | 25 | 0,7 | 0,5 | 0,2 |
| chr20:52836492-52836992:downstream_15091 | chr20 | 52824501 | 52836492 | + | PFDN4 | NM | 11 | 0,4 | 2 | 0 | 8,15E-01 | NA | 8 | 0 | 3 | 2 | 22 | 18 | 14 | 18 | 0,6 | 0,3 | 0,3 |
| chr17:34174246-34174746:downstream_20758 | chr17 | 34136458 | 34174246 | + | TAF15 | NM | 9 | 0,3 | 1 | 0 | 8,15E-01 | NA | 18 | 10 | 10 | 6 | 7 | 9 | 6 | 5 | 0,8 | 0,5 | 0,3 |
| chr14:102972697-102973197:downstream_11375 | chr14 | 102973197 | 102976128 | - | ANKRD9 | NM | 6 | 0,4 | 2 | 0 | 8,16E-01 | NA | 2 | 9 | 4 | 11 | 7 | 0 | 8 | 5 | -0,5 | -1,0 | 0,5 |
| chr1:54386733-54387233:downstream_136 | chr1 | 54387233 | 54411288 | - | HSPB11 | NM | 25 | -0,3 | 1 | 0 | 8,16E-01 | NA | 6 | 1 | 8 | 3 | 33 | 47 | 42 | 63 | -0,8 | -0,4 | -0,4 |
| chr1:32371521-32372021:downstream_17140 | chr1 | 32372021 | 32403988 | - | PTP4A2 | NM | 16 | 0,3 | 1 | 0 | 8,16E-01 | NA | 9 | 1 | 14 | 11 | 11 | 12 | 42 | 25 | -1,4 | -1,6 | 0,2 |
| chr15:75191798-75192298:downstream_19636 | chr15 | 75182351 | 75191798 | + | MPI | NM | 6 | 0,4 | 2 | 0 | 8,17E-01 | NA | 6 | 8 | 7 | 0 | 9 | 7 | 7 | 3 | 0,9 | 0,6 | 0,3 |
| chr19:45685058-45685558:downstream_14378 | chr19 | 45682002 | 45685058 | + | BLOC1S3 | NM | 13 | 0,3 | 1 | 0 | 8,17E-01 | NA | 10 | 5 | 1 | 5 | 22 | 31 | 15 | 15 | 1,2 | 0,8 | 0,4 |
| chr15:65952456-65952956:downstream_19525 | chr15 | 65952956 | 66084631 | - | DENND4A | NM | 15 | -0,3 | 1 | 0 | 8,17E-01 | NA | 7 | 1 | 42 | 28 | 5 | 1 | 18 | 21 | -3,1 | -2,8 | -0,3 |
| chr18:46569671-46570171:downstream_13510 | chr18 | 46570171 | 46987079 | - | DYM | NM | 15 | -0,3 | 1 | 0 | 8,17E-01 | NA | 10 | 0 | 10 | 21 | 10 | 12 | 31 | 24 | -1,7 | -1,3 | -0,3 |
| chr5:122344902-122345402:downstream_4076 | chr5 | 122181159 | 122344902 | + | SNX24 | NM | 10 | -0,4 | 2 | 0 | 8,18E-01 | NA | 2 | 0 | 10 | 8 | 7 | 0 | 33 | 19 | -3,5 | -3,0 | -0,5 |
| chr6:170884159-170884659:downstream_25050 | chr6 | 170884659 | 170893780 | - | PDCD2 | NM | 14 | 0,3 | 1 | 0 | 8,18E-01 | NA | 7 | 24 | 10 | 3 | 24 | 21 | 14 | 8 | 1,3 | 1,0 | 0,3 |
| chr20:20014273-20014773:downstream_22477 | chr20 | 19997933 | 20014273 | + | NAA20 | NM | 37 | -0,2 | 1 | 0 | 8,18E-01 | 9,65E-01 | 17 | 4 | 14 | 7 | 71 | 72 | 67 | 47 | 0,0 | 0,3 | -0,3 |
| chr4:122737098-122737598:downstream_3835 | chr4 | 122737598 | 122745088 | - | CCNA2 | NM | 34 | 0,2 | 1 | 0 | 8,18E-01 | 9,65E-01 | 6 | 2 | 14 | 13 | 23 | 24 | 90 | 97 | -1,8 | -2,0 | 0,2 |
| chr3:129033113-129033613:downstream_3176 | chr3 | 129033613 | 129035120 | - | H1FX | NM | 15 | -0,3 | 1 | 0 | 8,18E-01 | NA | 3 | 11 | 6 | 11 | 16 | 27 | 29 | 15 | -0,2 | 0,0 | -0,2 |
| chr6:76629254-76629754:downstream_25031 | chr6 | 76458892 | 76629254 | + | MYO6 | NM | 6 | 0,4 | 2 | 0 | 8,18E-01 | NA | 3 | 0 | 7 | 9 | 2 | 7 | 14 | 9 | -2,3 | -2,7 | 0,4 |
| chr5:90664040-90664540:downstream_4342 | chr5 | 90664540 | 90679149 | - | ARRDC3 | NM | 51 | -0,2 | 1 | 0 | 8,19E-01 | 9,65E-01 | 20 | 25 | 47 | 57 | 31 | 53 | 100 | 77 | -1,2 | -1,1 | -0,2 |
| chr7:93514208-93514708:downstream_25776 | chr7 | 93514708 | 93520303 | - | TFPI2 | NM | 88 | -0,1 | 1 | 0 | 8,19E-01 | 9,65E-01 | 33 | 21 | 19 | 17 | 218 | 163 | 116 | 116 | 0,6 | 0,7 | -0,2 |
| chr1:156218514-156219014:downstream_39 | chr1 | 156219014 | 156252620 | - | SMG5 | NM | 24 | 0,2 | 1 | 0 | 8,19E-01 | NA | 9 | 5 | 3 | 8 | 39 | 44 | 49 | 34 | 0,3 | 0,0 | 0,3 |
| chr7:23543900-23544400:downstream_25787 | chr7 | 23544400 | 23571660 | - | TRA2A | NM | 27 | 0,2 | 1 | 0 | 8,19E-01 | NA | 4 | 11 | 3 | 6 | 54 | 58 | 39 | 43 | 0,8 | 0,5 | 0,3 |
| chr17:19240366-19240866:downstream_20352 | chr17 | 19240866 | 19281495 | - | B9D1 | NM | 4 | -0,4 | 2 | 0 | 8,19E-01 | NA | 5 | 5 | 1 | 8 | 2 | 8 | 3 | 4 | 0,0 | 0,4 | -0,4 |
| chr20:2461965-2462465:downstream_22616 | chr20 | 2462465 | 2505168 | - | ZNF343 | NM | 18 | 0,3 | 1 | 0 | 8,20E-01 | NA | 7 | 8 | 3 | 7 | 24 | 41 | 35 | 21 | 0,6 | 0,2 | 0,4 |
| chr9:35055564-35056064:downstream_7049 | chr9 | 35056064 | 35072739 | - | VCP | NM | 58 | 0,1 | 1 | 0 | 8,20E-01 | 9,65E-01 | 84 | 68 | 49 | 48 | 62 | 65 | 39 | 50 | 0,6 | 0,5 | 0,1 |
| chr6:97336686-97337186:downstream_5274 | chr6 | 97337186 | 97345767 | - | NDUFAF4 | NM | 7 | 0,4 | 2 | 0 | 8,20E-01 | NA | 7 | 5 | 7 | 1 | 11 | 23 | 3 | 5 | 2,5 | 2,2 | 0,3 |
| chr11:62534187-62534687:downstream_9405 | chr11 | 62529010 | 62534187 | + | POLR2G | NM | 17 | 0,2 | 1 | 0 | 8,20E-01 | NA | 14 | 15 | 7 | 8 | 25 | 33 | 13 | 24 | 0,9 | 0,7 | 0,2 |
| chr2:242026008-242026508:downstream_22090 | chr2 | 242026508 | 242041747 | - | MTERF4 | NM | 8 | -0,3 | 1 | 0 | 8,20E-01 | NA | 7 | 7 | 4 | 6 | 15 | 13 | 11 | 5 | 0,5 | 0,8 | -0,3 |
| chr14:21667737-21668237:downstream_19280 | chr14 | 21668237 | 21675059 | - | LINC00641 | NR | 11 | -0,3 | 1 | 0 | 8,21E-01 | NA | 11 | 8 | 13 | 16 | 8 | 10 | 14 | 9 | -0,6 | -0,4 | -0,3 |
| chr5:10493138-10493638:downstream_4195 | chr5 | 10493638 | 10502840 | - | LOC101929412 | NR | 6 | -0,4 | 2 | 0 | 8,21E-01 | NA | 2 | 7 | 4 | 10 | 7 | 5 | 10 | 6 | -0,7 | -0,4 | -0,3 |
| chrX:84531894-84532394:downstream_26986 | chrX | 84532394 | 84634748 | - | POF1B | NM | 66 | -0,2 | 1 | 0 | 8,22E-01 | 9,65E-01 | 3 | 1 | 4 | 6 | 101 | 70 | 188 | 152 | -1,2 | -1,0 | -0,2 |
| chr17:48458820-48459320:downstream_20465 | chr17 | 48450580 | 48458820 | + | EME1 | NM | 32 | 0,2 | 1 | 0 | 8,22E-01 | 9,65E-01 | 27 | 47 | 25 | 30 | 25 | 43 | 32 | 26 | 0,4 | 0,2 | 0,2 |
| chr4:39552045-39552545:downstream_3422 | chr4 | 39552545 | 39640481 | - | SMIM14 | NM | 68 | 0,1 | 1 | 0 | 8,22E-01 | 9,65E-01 | 65 | 54 | 135 | 121 | 32 | 17 | 58 | 59 | -1,1 | -1,3 | 0,2 |
| chr1:231490773-231491273:downstream_17273 | chr1 | 231473681 | 231490773 | + | SPRTN | NM | 31 | 0,2 | 1 | 0 | 8,22E-01 | 9,65E-01 | 50 | 22 | 26 | 23 | 28 | 45 | 32 | 25 | 0,6 | 0,4 | 0,2 |
| chr17:57774317-57774817:downstream_20412 | chr17 | 57697049 | 57774317 | + | CLTC | NM | 18 | 0,2 | 1 | 0 | 8,23E-01 | NA | 10 | 19 | 17 | 18 | 9 | 22 | 32 | 13 | -0,3 | -0,5 | 0,3 |
| chr2:220073987-220074487:downstream_2445 | chr2 | 220074487 | 220083712 | - | ABCB6 | NM | 25 | 0,3 | 1 | 0 | 8,23E-01 | NA | 11 | 3 | 4 | 2 | 65 | 52 | 25 | 35 | 1,1 | 1,0 | 0,1 |
| chr6:36822105-36822605:downstream_4723 | chr6 | 36822605 | 36842800 | - | PPIL1 | NM | 43 | -0,2 | 1 | 0 | 8,23E-01 | 9,65E-01 | 8 | 4 | 10 | 7 | 85 | 64 | 75 | 94 | -0,4 | -0,2 | -0,3 |
| chr1:55222070-55222570:downstream_749 | chr1 | 55222570 | 55230226 | - | PARS2 | NM | 10 | 0,3 | 1 | 0 | 8,24E-01 | NA | 5 | 13 | 7 | 2 | 18 | 13 | 13 | 7 | 0,9 | 0,6 | 0,3 |
| chr2:61368226-61368726:downstream_1984 | chr2 | 61368726 | 61372110 | - | LOC339803 | NR | 7 | -0,4 | 2 | 0 | 8,24E-01 | NA | 5 | 0 | 6 | 6 | 2 | 13 | 17 | 7 | -1,2 | -0,8 | -0,5 |
| chr8:25276273-25276773:downstream_25951 | chr8 | 25276773 | 25282556 | - | GNRH1 | NM | 5 | -0,4 | 2 | 0 | 8,24E-01 | NA | 5 | 1 | 3 | 7 | 11 | 1 | 3 | 11 | -0,7 | -0,2 | -0,5 |
| chr17:1962981-1963481:downstream_20512 | chr17 | 1958392 | 1962981 | + | HIC1 | NM | 5 | -0,6 | 3 | 0 | 8,24E-01 | NA | 11 | 4 | 0 | 0 | 20 | 3 | 0 | 0 | Inf | Inf | NaN |
| chr1:153506575-153507075:downstream_1066 | chr1 | 153507075 | 153508717 | - | S100A6 | NM | 5636 | 0,1 | 1 | 0 | 8,24E-01 | 9,65E-01 | 19 | 24 | 14 | 13 | 14011 | 12675 | 10203 | 8129 | 0,7 | 0,5 | 0,1 |
| chr1:36920861-36921361:downstream_640 | chr1 | 36921361 | 36930040 | - | MRPS15 | NM | 40 | 0,2 | 1 | 0 | 8,24E-01 | 9,65E-01 | 21 | 9 | 15 | 16 | 61 | 60 | 77 | 64 | 0,0 | -0,2 | 0,2 |
| chr15:35148051-35148551:downstream_11441 | chr15 | 35148551 | 35261995 | - | AQR | NM | 48 | 0,2 | 1 | 0 | 8,24E-01 | 9,65E-01 | 7 | 5 | 32 | 22 | 38 | 12 | 152 | 120 | -2,3 | -2,4 | 0,2 |
| chr2:3628509-3629009:downstream_1828 | chr2 | 3622852 | 3628509 | + | RPS7 | NM | 44 | 0,2 | 1 | 0 | 8,25E-01 | 9,65E-01 | 28 | 6 | 10 | 7 | 92 | 101 | 65 | 45 | 1,0 | 0,8 | 0,2 |
| chr8:8174757-8175257:downstream_6430 | chr8 | 8175257 | 8239344 | - | SGK223 | NM | 8 | -0,3 | 1 | 0 | 8,25E-01 | NA | 3 | 4 | 8 | 10 | 6 | 6 | 17 | 7 | -1,4 | -1,1 | -0,3 |
| chr1:155720479-155720979:downstream_16989 | chr1 | 155715558 | 155720479 | + | MSTO2P | NR | 11 | 0,3 | 1 | 0 | 8,25E-01 | NA | 8 | 21 | 13 | 25 | 3 | 5 | 4 | 8 | -0,3 | -0,6 | 0,3 |
| chr22:19427909-19428409:downstream_22761 | chr22 | 19428409 | 19435755 | - | C22orf39 | NM | 7 | 0,3 | 2 | 0 | 8,26E-01 | NA | 4 | 6 | 3 | 3 | 11 | 11 | 7 | 7 | 0,7 | 0,3 | 0,4 |
| chr3:15294423-15294923:downstream_2839 | chr3 | 15247732 | 15294423 | + | CAPN7 | NM | 27 | 0,2 | 1 | 0 | 8,26E-01 | NA | 33 | 24 | 49 | 43 | 19 | 4 | 15 | 26 | -0,7 | -0,9 | 0,2 |
| chr4:140311935-140312435:downstream_3511 | chr4 | 140222675 | 140311935 | + | NAA15 | NM | 8 | 0,4 | 2 | 0 | 8,26E-01 | NA | 3 | 11 | 17 | 15 | 5 | 0 | 6 | 9 | -1,2 | -1,6 | 0,4 |
| chr12:25357949-25358449:downstream_10105 | chr12 | 25348149 | 25357949 | + | LYRM5 | NM | 10 | -0,3 | 1 | 0 | 8,27E-01 | NA | 8 | 3 | 1 | 5 | 19 | 25 | 7 | 14 | 0,8 | 1,1 | -0,3 |
| chr19:42751212-42751712:downstream_21122 | chr19 | 42751712 | 42759316 | - | ERF | NM | 24 | 0,2 | 1 | 0 | 8,27E-01 | NA | 10 | 12 | 6 | 3 | 24 | 84 | 22 | 29 | 1,3 | 1,1 | 0,2 |
| chr3:12875943-12876443:downstream_23394 | chr3 | 12876443 | 12883081 | - | RPL32 | NM | 19 | 0,2 | 1 | 0 | 8,28E-01 | NA | 23 | 20 | 8 | 11 | 41 | 11 | 11 | 20 | 1,1 | 0,9 | 0,2 |
| chr5:65479444-65479944:downstream_24394 | chr5 | 65440045 | 65479444 | + | SREK1 | NM | 11 | -0,3 | 2 | 0 | 8,28E-01 | NA | 0 | 4 | 7 | 6 | 4 | 15 | 35 | 18 | -1,7 | -1,4 | -0,3 |
| chr12:121862155-121862655:downstream_18803 | chr12 | 121837885 | 121862155 | + | RNF34 | NM | 9 | -0,3 | 2 | 0 | 8,29E-01 | NA | 5 | 0 | 7 | 5 | 11 | 8 | 18 | 16 | -1,3 | -0,9 | -0,4 |
| chr10:43880564-43881064:downstream_17597 | chr10 | 43881064 | 43904696 | - | HNRNPF | NM | 52 | -0,2 | 1 | 0 | 8,29E-01 | 9,68E-01 | 15 | 1 | 15 | 7 | 91 | 88 | 117 | 86 | -0,5 | -0,2 | -0,3 |
| chr14:37642998-37643498:downstream_11113 | chr14 | 37641230 | 37642998 | + | SLC25A21-AS1 | NR | 34 | -0,2 | 1 | 0 | 8,29E-01 | 9,68E-01 | 37 | 42 | 0 | 18 | 70 | 72 | 17 | 14 | 2,1 | 2,2 | -0,1 |
| chr17:41478504-41479004:downstream_13149 | chr17 | 41476352 | 41478504 | + | ARL4D | NM | 4 | 0,5 | 2 | 0 | 8,29E-01 | NA | 5 | 5 | 1 | 2 | 1 | 13 | 4 | 3 | 1,3 | 0,8 | 0,5 |
| chr17:26655711-26656211:downstream_12921 | chr17 | 26646120 | 26655711 | + | TMEM97 | NM | 57 | 0,2 | 1 | 0 | 8,30E-01 | 9,68E-01 | 18 | 53 | 38 | 53 | 70 | 50 | 70 | 105 | -0,4 | -0,5 | 0,2 |
| chr20:43537175-43537675:downstream_22606 | chr20 | 43514239 | 43537175 | + | YWHAB | NM | 79 | 0,2 | 1 | 0 | 8,30E-01 | 9,68E-01 | 12 | 15 | 14 | 11 | 180 | 115 | 148 | 140 | 0,2 | 0,0 | 0,1 |
| chrX:47519510-47520010:downstream_7698 | chrX | 47518231 | 47519510 | + | UXT-AS1 | NR | 6 | 0,4 | 2 | 0 | 8,31E-01 | NA | 10 | 4 | 4 | 2 | 8 | 11 | 10 | 1 | 1,2 | 0,8 | 0,4 |
| chr9:127115243-127115743:downstream_6935 | chr9 | 127115743 | 127177752 | - | PSMB7 | NM | 100 | -0,2 | 1 | 0 | 8,32E-01 | 9,69E-01 | 6 | 6 | 1 | 10 | 211 | 183 | 189 | 198 | 0,0 | 0,0 | 0,0 |
| chr7:66786513-66787013:downstream_25751 | chr7 | 66767624 | 66786513 | + | STAG3L4 | NR | 13 | 0,3 | 2 | 0 | 8,32E-01 | NA | 11 | 2 | 29 | 7 | 11 | 0 | 24 | 18 | -1,5 | -1,9 | 0,3 |
| chr16:89633237-89633737:downstream_20197 | chr16 | 89627064 | 89633237 | + | RPL13 | NM | 15 | 0,2 | 1 | 0 | 8,32E-01 | NA | 20 | 20 | 4 | 5 | 28 | 25 | 7 | 7 | 2,2 | 1,9 | 0,3 |
| chr3:31701816-31702316:downstream_23320 | chr3 | 31702316 | 32023342 | - | OSBPL10 | NM | 6 | -0,5 | 2 | 0 | 8,32E-01 | NA | 1 | 0 | 3 | 9 | 3 | 0 | 22 | 11 | -3,9 | -3,4 | -0,5 |
| chr5:139499001-139499501:downstream_4453 | chr5 | 139493707 | 139499001 | + | PURA | NM | 33 | 0,2 | 1 | 0 | 8,33E-01 | 9,69E-01 | 37 | 44 | 32 | 21 | 41 | 34 | 24 | 30 | 0,6 | 0,5 | 0,1 |
| chr7:108202087-108202587:downstream_25778 | chr7 | 108202587 | 108210212 | - | THAP5 | NM | 48 | 0,2 | 1 | 0 | 8,33E-01 | 9,69E-01 | 45 | 41 | 29 | 32 | 66 | 64 | 74 | 32 | 0,5 | 0,3 | 0,2 |
| chr2:71305998-71306498:downstream_1521 | chr2 | 71295407 | 71305998 | + | NAGK | NM | 14 | 0,2 | 1 | 0 | 8,33E-01 | NA | 11 | 25 | 7 | 12 | 13 | 22 | 10 | 13 | 0,9 | 0,6 | 0,3 |
| chr7:22851750-22852250:downstream_5733 | chr7 | 22852250 | 22862471 | - | TOMM7 | NM | 28 | -0,2 | 1 | 0 | 8,33E-01 | 9,69E-01 | 15 | 10 | 6 | 14 | 58 | 47 | 38 | 39 | 0,3 | 0,5 | -0,1 |
| chr7:33053224-33053724:downstream_25653 | chr7 | 33053724 | 33102409 | - | NT5C3A | NM | 57 | 0,2 | 1 | 0 | 8,34E-01 | 9,69E-01 | 1 | 10 | 6 | 10 | 67 | 84 | 135 | 141 | -0,6 | -0,9 | 0,3 |
| chr1:53679869-53680369:downstream_740 | chr1 | 53662100 | 53679869 | + | CPT2 | NM | 19 | 0,2 | 1 | 0 | 8,34E-01 | NA | 11 | 6 | 8 | 5 | 29 | 38 | 24 | 34 | 0,4 | 0,2 | 0,1 |
| chr12:4723054-4723554:downstream_18549 | chr12 | 4699237 | 4723054 | + | DYRK4 | NM | 19 | 0,2 | 1 | 0 | 8,35E-01 | NA | 15 | 10 | 10 | 6 | 38 | 30 | 22 | 25 | 0,7 | 0,5 | 0,2 |
| chr19:5910263-5910763:downstream_13918 | chr19 | 5904851 | 5910263 | + | VMAC | NM | 5 | 0,4 | 2 | 0 | 8,35E-01 | NA | 2 | 2 | 6 | 10 | 2 | 0 | 4 | 11 | -2,1 | -2,6 | 0,5 |

| | | | | | | | | | | | | | | | | | | | | | | | |
|---|---|---|---|---|---|---|---|---|---|---|---|---|---|---|---|---|---|---|---|---|---|---|---|
| chr5:90598302-90598802:downstream_24212 | chr5 | 90598802 | 90610219 | - | LUCAT1 | NR | 17 | -0,2 | 1 | 0 | 8,35E-01 | NA | 6 | 6 | 24 | 29 | 6 | 10 | 39 | 21 | -2,2 | -2,0 | -0,2 |
| chr9:99403032-99403532:downstream_7208 | chr9 | 99403532 | 99417599 | - | AAED1 | NM | 22 | -0,2 | 1 | 0 | 8,35E-01 | NA | 10 | 8 | 19 | 20 | 28 | 14 | 40 | 40 | -1,2 | -1,0 | -0,2 |
| chr1:114934898-114935398:downstream_17355 | chr1 | 114935398 | 115053781 | - | TRIM33 | NM | 8 | -0,4 | 2 | 0 | 8,36E-01 | NA | 0 | 3 | 6 | 7 | 2 | 8 | 24 | 14 | -2,2 | -1,9 | -0,3 |
| chr2:63273621-63274121:downstream_21893 | chr2 | 62900985 | 63273621 | + | EHBP1 | NM | 53 | -0,3 | 1 | 0 | 8,37E-01 | 9,71E-01 | 2 | 0 | 4 | 11 | 29 | 16 | 177 | 186 | -3,3 | -3,0 | -0,3 |
| chr1:46501796-46502296:downstream_127 | chr1 | 46269284 | 46501796 | + | MAST2 | NM | 8 | -0,4 | 2 | 0 | 8,37E-01 | NA | 2 | 1 | 0 | 8 | 6 | 11 | 13 | 20 | -1,3 | -1,0 | -0,3 |
| chr16:11347773-11348273:downstream_12186 | chr16 | 11348273 | 11350039 | - | SOCS1 | NM | 4 | 0,6 | 3 | 0 | 8,37E-01 | NA | 6 | 14 | 0 | 0 | 10 | 3 | 0 | 0 | Inf | Inf | NaN |
| chr7:73248420-73248920:downstream_5897 | chr7 | 73248920 | 73256855 | - | WBSCR27 | NM | 9 | 0,3 | 1 | 0 | 8,37E-01 | NA | 11 | 11 | 13 | 7 | 10 | 7 | 10 | 7 | 0,2 | -0,1 | 0,3 |
| chr18:72258509-72259009:downstream_13675 | chr18 | 72259009 | 72265071 | - | LINC00909 | NR | 9 | -0,3 | 2 | 0 | 8,38E-01 | NA | 15 | 3 | 7 | 1 | 17 | 19 | 10 | 2 | 1,2 | 1,6 | -0,4 |
| chr11:93465893-93466393:downstream_9580 | chr11 | 93466393 | 93466466 | - | SNORD5 | NR | 159 | -0,1 | 1 | 0 | 8,38E-01 | 9,72E-01 | 231 | 199 | 198 | 201 | 143 | 95 | 104 | 101 | 0,1 | 0,2 | -0,1 |
| chr9:123362695-123363195:downstream_6930 | chr9 | 123363195 | 123476765 | - | MEGF9 | NM | 11 | -0,4 | 2 | 0 | 8,38E-01 | NA | 0 | 0 | 6 | 7 | 2 | 0 | 26 | 50 | -Inf | -5,0 | -Inf |
| chr17:62473401-62473901:downstream_13309 | chr17 | 62473901 | 62493184 | - | POLG2 | NM | 20 | -0,2 | 1 | 0 | 8,38E-01 | NA | 29 | 27 | 14 | 9 | 26 | 35 | 8 | 12 | 1,3 | 1,6 | -0,3 |
| chr15:77777946-77778446:downstream_19570 | chr15 | 77712992 | 77777946 | + | HMG20A | NM | 12 | 0,3 | 2 | 0 | 8,38E-01 | NA | 2 | 0 | 8 | 9 | 8 | 0 | 32 | 39 | -2,8 | -3,1 | 0,3 |
| chr19:52467092-52467592:downstream_14548 | chr19 | 52467592 | 52490079 | - | ZNF350 | NM | 12 | 0,3 | 1 | 0 | 8,38E-01 | NA | 7 | 10 | 4 | 3 | 16 | 31 | 13 | 13 | 1,1 | 0,9 | 0,2 |
| chr19:52092991-52093491:downstream_14537 | chr19 | 52074530 | 52092991 | + | ZNF175 | NM | 9 | 0,3 | 2 | 0 | 8,38E-01 | NA | 4 | 7 | 10 | 1 | 12 | 11 | 21 | 6 | 0,0 | -0,2 | 0,3 |
| chr12:85695561-85696061:downstream_10338 | chr12 | 85674035 | 85695561 | + | ALX1 | NM | 97 | -0,1 | 1 | 0 | 8,38E-01 | 9,72E-01 | 9 | 15 | 85 | 88 | 34 | 41 | 270 | 239 | -2,9 | -2,8 | -0,1 |
| chr18:649119-649619:downstream_13477 | chr18 | 649619 | 658340 | - | TYMSOS | NM | 19 | -0,2 | 1 | 0 | 8,39E-01 | NA | 25 | 19 | 3 | 5 | 28 | 56 | 8 | 5 | 2,5 | 2,7 | -0,1 |
| chr19:50986783-50987283:downstream_21112 | chr19 | 50979733 | 50986783 | + | EMC10 | NM | 179 | -0,1 | 1 | 0 | 8,39E-01 | 9,72E-01 | 19 | 25 | 13 | 23 | 380 | 379 | 301 | 290 | 0,3 | 0,4 | -0,1 |
| chr12:98944157-98944657:downstream_18884 | chr12 | 98909350 | 98944157 | + | TMPO | NM | 83 | -0,2 | 1 | 0 | 8,40E-01 | 9,72E-01 | 0 | 6 | 8 | 9 | 67 | 100 | 253 | 220 | -1,6 | -1,5 | -0,1 |
| chr8:106792137-106792637:downstream_26191 | chr8 | 106792637 | 107072731 | - | ZFPM2-AS1 | NR | 6 | -0,4 | 2 | 0 | 8,41E-01 | NA | 0 | 2 | 4 | 7 | 2 | 3 | 14 | 12 | -2,6 | -2,3 | -0,3 |
| chr7:148503963-148504463:downstream_25507 | chr7 | 148504463 | 148581441 | - | EZH2 | NM | 4 | 0,4 | 2 | 0 | 8,41E-01 | NA | 0 | 5 | 8 | 3 | 2 | 1 | 8 | 3 | -1,3 | -1,8 | 0,5 |
| chr1:32799224-32799724:downstream_104 | chr1 | 32757707 | 32799224 | + | HDAC1 | NM | 26 | 0,2 | 1 | 0 | 8,41E-01 | NA | 10 | 15 | 17 | 16 | 35 | 25 | 43 | 48 | -0,4 | -0,6 | 0,2 |
| chr18:19210206-19210706:downstream_20947 | chr18 | 19192229 | 19210206 | + | SNRPD1 | NM | 26 | 0,2 | 1 | 0 | 8,41E-01 | NA | 15 | 21 | 19 | 27 | 15 | 35 | 38 | 37 | -0,4 | -0,6 | 0,2 |
| chr19:9717501-9718001:downstream_13980 | chr19 | 9718001 | 9731916 | - | ZNF561 | NM | 5 | -0,4 | 2 | 0 | 8,42E-01 | NA | 7 | 9 | 0 | 2 | 13 | 9 | 0 | 2 | 3,3 | 3,7 | -0,4 |
| chr2:43455994-43456494:downstream_1954 | chr2 | 43454349 | 43455994 | + | LINC01126 | NR | 6 | 0,4 | 2 | 0 | 8,42E-01 | NA | 9 | 2 | 7 | 7 | 9 | 1 | 11 | 5 | -0,3 | -0,7 | 0,4 |
| chr5:76727568-76728068:downstream_4312 | chr5 | 76728068 | 76788332 | - | WDR41 | NM | 28 | 0,2 | 1 | 0 | 8,42E-01 | 9,73E-01 | 17 | 16 | 25 | 25 | 40 | 14 | 46 | 44 | -0,6 | -0,8 | 0,2 |
| chr8:71023766-71024266:downstream_6239 | chr8 | 71024266 | 71316020 | - | NCOA2 | NM | 31 | 0,2 | 1 | 0 | 8,42E-01 | 9,73E-01 | 10 | 11 | 65 | 66 | 6 | 6 | 42 | 42 | -2,7 | -2,9 | 0,2 |
| chr17:46138907-46139407:downstream_12569 | chr17 | 46125685 | 46138907 | + | NFE2L1 | NM | 54 | -0,2 | 1 | 0 | 8,43E-01 | 9,73E-01 | 4 | 12 | 14 | 16 | 81 | 59 | 109 | 135 | -0,9 | -0,8 | -0,1 |
| chr22:28373501-28374001:downstream_15415 | chr22 | 28374001 | 29075853 | - | TTC28 | NM | 4 | -0,6 | 3 | 0 | 8,43E-01 | NA | 0 | 0 | 4 | 6 | 1 | 0 | 8 | 10 | -Inf | -4,5 | -Inf |
| chr1:35446626-35447126:downstream_633 | chr1 | 35447126 | 35450948 | - | ZMYM6NB | NM | 49 | 0,2 | 1 | 0 | 8,43E-01 | 9,73E-01 | 17 | 11 | 13 | 7 | 98 | 101 | 67 | 77 | 0,5 | 0,5 | 0,1 |
| chr19:13265718-13266218:downstream_14048 | chr19 | 13261281 | 13265718 | + | IER2 | NM | 21 | -0,2 | 1 | 0 | 8,44E-01 | NA | 6 | 4 | 4 | 2 | 35 | 68 | 26 | 22 | 0,7 | 1,1 | -0,4 |
| chr2:27873175-27873675:downstream_22268 | chr2 | 27873675 | 27886707 | - | SUPT7L | NM | 24 | -0,2 | 1 | 0 | 8,44E-01 | NA | 45 | 29 | 26 | 35 | 17 | 18 | 4 | 21 | 0,3 | 0,5 | -0,3 |
| chr15:81296345-81296845:downstream_11849 | chr15 | 81293294 | 81296345 | + | MESDC1 | NM | 39 | 0,2 | 1 | 0 | 8,44E-01 | 9,73E-01 | 28 | 13 | 3 | 7 | 86 | 117 | 33 | 24 | 2,1 | 1,8 | 0,3 |
| chr3:53846492-53846992:downstream_23037 | chr3 | 53529075 | 53846492 | + | CACNA1D | NM | 10 | -0,3 | 1 | 0 | 8,44E-01 | NA | 6 | 1 | 8 | 4 | 7 | 18 | 21 | 16 | -0,9 | -0,5 | -0,4 |
| chr1:234508052-234508552:downstream_16901 | chr1 | 234508552 | 234509339 | - | LOC101927765 | NR | 9 | 0,3 | 2 | 0 | 8,45E-01 | NA | 1 | 13 | 6 | 5 | 11 | 15 | 11 | 14 | 0,4 | 0,0 | 0,3 |
| chr19:58291984-58292484:downstream_21685 | chr19 | 58281019 | 58291984 | + | ZNF586 | NM | 21 | 0,2 | 1 | 0 | 8,45E-01 | NA | 25 | 28 | 22 | 11 | 23 | 24 | 11 | 21 | 0,7 | 0,5 | 0,1 |
| chr19:44416275-44416775:downstream_13779 | chr19 | 44416775 | 44439411 | - | ZNF45 | NM | 20 | -0,2 | 1 | 0 | 8,45E-01 | NA | 2 | 1 | 6 | 6 | 23 | 16 | 61 | 47 | -1,7 | -1,5 | -0,3 |
| chr4:367691-368191:downstream_3559 | chr4 | 331595 | 367691 | + | ZNF141 | NM | 26 | 0,2 | 1 | 0 | 8,45E-01 | NA | 32 | 17 | 35 | 40 | 19 | 13 | 25 | 29 | -0,6 | -0,8 | 0,2 |
| chr17:61679-62179:downstream_20684 | chr17 | 62179 | 202633 | - | RPH3AL | NM | 15 | 0,2 | 1 | 0 | 8,45E-01 | NA | 14 | 11 | 24 | 17 | 15 | 5 | 24 | 15 | -0,7 | -1,0 | 0,2 |
| chr19:46386364-46386864:downstream_14389 | chr19 | 46386864 | 46389428 | - | IRF2BP1 | NM | 20 | -0,2 | 1 | 0 | 8,45E-01 | NA | 7 | 10 | 7 | 4 | 37 | 46 | 26 | 20 | 0,6 | 0,9 | -0,2 |
| chr5:14699842-14700342:downstream_3955 | chr5 | 14664782 | 14699842 | + | OTULIN | NM | 10 | -0,3 | 1 | 0 | 8,45E-01 | NA | 7 | 20 | 11 | 4 | 7 | 16 | 6 | 5 | 0,9 | 1,2 | -0,3 |
| chr4:159045231-159045731:downstream_23644 | chr4 | 159045731 | 159094202 | - | FAM198B | NM | 10 | 0,3 | 1 | 0 | 8,46E-01 | NA | 2 | 1 | 6 | 6 | 5 | 8 | 21 | 29 | -1,7 | -2,0 | 0,2 |
| chr12:11328619-11329119:downstream_10058 | chr12 | 11323779 | 11328619 | + | SMIM10L1 | NM | 14 | -0,3 | 1 | 0 | 8,46E-01 | NA | 33 | 21 | 11 | 5 | 24 | 11 | 8 | 1 | 1,7 | 1,9 | -0,2 |
| chr20:33301237-33301737:downstream_14786 | chr20 | 33292147 | 33301237 | + | TP53INP2 | NM | 8 | -0,3 | 1 | 0 | 8,46E-01 | NA | 4 | 5 | 6 | 3 | 17 | 11 | 13 | 10 | 0,0 | 0,3 | -0,3 |
| chr16:50352428-50352928:downstream_19896 | chr16 | 50352928 | 50402845 | - | BRD7 | NM | 29 | -0,2 | 1 | 0 | 8,46E-01 | 9,73E-01 | 41 | 38 | 49 | 33 | 20 | 15 | 15 | 17 | 0,0 | 0,1 | -0,2 |
| chr2:48033558-48034058:downstream_21920 | chr2 | 48034058 | 48132932 | - | FBXO11 | NM | 15 | 0,3 | 1 | 0 | 8,46E-01 | NA | 5 | 0 | 15 | 11 | 11 | 2 | 40 | 39 | -2,5 | -2,7 | 0,2 |
| chr8:62414204-62414704:downstream_6303 | chr8 | 62200524 | 62414204 | + | CLVS1 | NM | 81 | 0,2 | 1 | 0 | 8,47E-01 | 9,73E-01 | 52 | 193 | 202 | 130 | 6 | 21 | 21 | 20 | -0,4 | -0,6 | 0,2 |
| chr1:211747880-211748380:downstream_1354 | chr1 | 211748380 | 211752099 | - | SLC30A1 | NM | 655 | 0,1 | 0 | 0 | 8,47E-01 | 9,73E-01 | 1578 | 1524 | 253 | 208 | 768 | 682 | 138 | 93 | 2,8 | 2,7 | 0,1 |
| chr10:119038941-119039441:downstream_8350 | chr10 | 119000583 | 119038941 | + | SLC18A2 | NM | 11 | 0,3 | 2 | 0 | 8,48E-01 | NA | 2 | 4 | 8 | 5 | 0 | 16 | 32 | 17 | -1,3 | -1,6 | 0,3 |
| chr21:46224631-46225131:downstream_15368 | chr21 | 46222155 | 46224631 | + | LINC01424 | NR | 7 | 0,3 | 2 | 0 | 8,48E-01 | NA | 9 | 16 | 8 | 2 | 2 | 10 | 1 | 4 | 1,2 | 1,0 | 0,2 |
| chr8:144735900-144736400:downstream_26197 | chr8 | 144718182 | 144735900 | + | ZNF623 | NM | 36 | 0,1 | 1 | 0 | 8,48E-01 | 9,73E-01 | 37 | 41 | 31 | 36 | 36 | 36 | 36 | 32 | 0,2 | 0,1 | 0,2 |
| chr5:145668784-145669284:downstream_4101 | chr5 | 145583162 | 145668784 | + | RBM27 | NM | 15 | -0,3 | 1 | 0 | 8,48E-01 | NA | 12 | 1 | 29 | 28 | 10 | 2 | 14 | 28 | -2,1 | -1,8 | -0,3 |
| chr17:8371495-8371995:downstream_20607 | chr17 | 8339169 | 8371495 | + | NDEL1 | NM | 7 | 0,4 | 2 | 0 | 8,49E-01 | NA | 8 | 10 | 8 | 3 | 13 | 0 | 1 | 9 | 0,6 | 0,3 | 0,3 |
| chr2:160174989-160175489:downstream_21765 | chr2 | 160175489 | 160473112 | - | BAZ2B | NM | 16 | 0,3 | 2 | 0 | 8,49E-01 | NA | 2 | 0 | 6 | 8 | 9 | 1 | 43 | 58 | -3,1 | -3,4 | 0,3 |
| chr2:158592457-158592957:downstream_21722 | chr2 | 158592957 | 158732374 | - | ACVR1 | NM | 12 | 0,3 | 2 | 0 | 8,49E-01 | NA | 2 | 0 | 6 | 5 | 5 | 10 | 39 | 32 | -2,1 | -2,3 | 0,2 |
| chr1:206822542-206823042:downstream_16674 | chr1 | 206808880 | 206822542 | + | DYRK3 | NM | 27 | -0,2 | 1 | 0 | 8,49E-01 | NA | 37 | 26 | 15 | 11 | 40 | 52 | 14 | 19 | 1,3 | 1,5 | -0,2 |
| chr5:173033647-173034147:downstream_24025 | chr5 | 173034147 | 173043666 | - | BOD1 | NM | 23 | -0,2 | 1 | 0 | 8,49E-01 | NA | 18 | 21 | 11 | 14 | 39 | 36 | 24 | 20 | 0,6 | 0,8 | -0,1 |
| chr10:135234174-135234674:downstream_8802 | chr10 | 135207620 | 135234174 | + | MTG1 | NM | 9 | 0,3 | 1 | 0 | 8,50E-01 | NA | 10 | 2 | 7 | 3 | 13 | 13 | 13 | 13 | 0,2 | 0,0 | 0,2 |
| chr5:98190407-98190907:downstream_4349 | chr5 | 98190907 | 98262238 | - | CHD1 | NM | 34 | -0,2 | 1 | 0 | 8,50E-01 | 9,73E-01 | 7 | 0 | 38 | 24 | 11 | 12 | 88 | 90 | -3,2 | -3,0 | -0,3 |
| chr17:48127212-48127712:downstream_13221 | chr17 | 48127712 | 48133103 | - | LOC284080 | NR | 4 | -0,5 | 3 | 0 | 8,50E-01 | NA | 7 | 14 | 1 | 0 | 6 | 5 | 0 | 0 | 3,9 | Inf | -Inf |
| chr3:156464631-156465131:downstream_2732 | chr3 | 156465131 | 156534851 | - | LINC00886 | NR | 5 | -0,4 | 2 | 0 | 8,51E-01 | NA | 5 | 0 | 10 | 7 | 6 | 0 | 4 | 11 | -1,8 | -1,4 | -0,4 |
| chr5:65376851-65377351:downstream_24107 | chr5 | 65222381 | 65376851 | + | ERBB2IP | NM | 8 | -0,3 | 2 | 0 | 8,51E-01 | NA | 3 | 0 | 10 | 23 | 3 | 0 | 14 | 12 | -3,3 | -3,0 | -0,3 |
| chr6:143815448-143815948:downstream_5382 | chr6 | 143815948 | 143833020 | - | FUCA2 | NM | 36 | 0,1 | 1 | 0 | 8,51E-01 | 9,73E-01 | 46 | 57 | 49 | 28 | 25 | 37 | 28 | 23 | 0,4 | 0,3 | 0,1 |
| chr1:149821258-149821758:downstream_978 | chr1 | 149821758 | 149822340 | - | HIST2H2BC | NR | 44 | -0,2 | 1 | 0 | 8,51E-01 | 9,73E-01 | 133 | 101 | 42 | 33 | 6 | 27 | 3 | 6 | 1,6 | 1,9 | -0,3 |
| chr10:90823167-90823667:downstream_8627 | chr10 | 90823092 | 90823167 | + | MIR4679-1 | NR | 7 | -0,5 | 2 | 0 | 8,52E-01 | NA | 9 | 30 | 1 | 0 | 11 | 9 | 0 | 0 | 4,8 | Inf | -Inf |
| chr10:131978646-131979146:downstream_17588 | chr10 | 131934638 | 131978646 | + | GLRX3 | NM | 6 | 0,3 | 2 | 0 | 8,52E-01 | NA | 9 | 11 | 3 | 3 | 3 | 9 | 7 | 3 | 1,7 | 1,4 | 0,3 |
| chr17:7765600-7766100:downstream_12815 | chr17 | 7761063 | 7765600 | + | CYB5D1 | NM | 32 | 0,2 | 1 | 0 | 8,52E-01 | 9,73E-01 | 12 | 3 | 3 | 4 | 73 | 78 | 46 | 34 | 1,1 | 0,9 | 0,2 |
| chr20:42934696-42935196:downstream_15022 | chr20 | 42935196 | 42939889 | - | FITM2 | NM | 30 | 0,2 | 1 | 0 | 8,52E-01 | 9,73E-01 | 34 | 12 | 7 | 7 | 70 | 67 | 17 | 28 | 1,8 | 1,6 | 0,1 |
| chr5:86689578-86690078:downstream_24049 | chr5 | 86690078 | 86708850 | - | CCNH | NM | 43 | -0,1 | 1 | 0 | 8,52E-01 | 9,73E-01 | 15 | 24 | 21 | 18 | 58 | 80 | 68 | 57 | 0,0 | 0,1 | -0,1 |
| chr18:12329824-12330324:downstream_20965 | chr18 | 12307667 | 12329824 | + | TUBB6 | NM | 13 | -0,2 | 1 | 0 | 8,53E-01 | NA | 6 | 4 | 6 | 2 | 32 | 24 | 15 | 17 | 0,4 | 0,8 | -0,4 |
| chr17:37005820-37006320:downstream_13023 | chr17 | 37006320 | 37010053 | - | RPL23 | NM | 404 | -0,1 | 0 | 0 | 8,53E-01 | 9,73E-01 | 110 | 94 | 100 | 63 | 870 | 772 | 610 | 612 | 0,3 | 0,4 | -0,1 |
| chr7:150773207-150773707:downstream_25516 | chr7 | 150773707 | 150777970 | - | FASTK | NM | 13 | -0,2 | 1 | 0 | 8,53E-01 | NA | 12 | 26 | 11 | 18 | 11 | 10 | 6 | 8 | 0,4 | 0,6 | -0,2 |
| chr17:66132070-66132570:downstream_13318 | chr17 | 66097695 | 66132070 | + | LINC00674 | NR | 12 | 0,2 | 1 | 0 | 8,53E-01 | NA | 6 | 6 | 10 | 2 | 13 | 21 | 19 | 21 | -0,1 | -0,2 | 0,1 |
| chr12:89741101-89741601:downstream_18546 | chr12 | 89741601 | 89746636 | - | DUSP6 | NM | 19 | -0,2 | 1 | 0 | 8,54E-01 | NA | 14 | 13 | 6 | 4 | 39 | 50 | 14 | 14 | 1,4 | 1,7 | -0,2 |
| chr5:55299475-55299975:downstream_4260 | chr5 | 55290994 | 55299475 | + | FLJ31104 | NR | 5 | -0,4 | 2 | 0 | 8,54E-01 | NA | 5 | 6 | 1 | 0 | 11 | 16 | 1 | 1 | 2,9 | 3,7 | -0,7 |
| chr7_gl000195_random:42437-42937:downstream_16434 | chr7 | gl000195 | random | LOC389831 | 42937 | 86719 | 84 | -0,1 | 1 | 0 | 8,54E-01 | 9,73E-01 | 19 | 19 | 35 | 46 | 102 | 85 | 181 | 189 | -1,1 | -1,0 | -0,1 |
| chr10:32556143-32556643:downstream_17557 | chr10 | 32556643 | 32667726 | - | EPC1 | NM | 20 | 0,2 | 1 | 0 | 8,55E-01 | NA | 22 | 33 | 22 | 29 | 12 | 15 | 13 | 16 | 0,1 | -0,1 | 0,2 |
| chr3:49394108-49394608:downstream_23179 | chr3 | 49394608 | 49395791 | - | GPX1 | NM | 22 | -0,2 | 1 | 0 | 8,55E-01 | NA | 15 | 7 | 0 | 2 | 81 | 58 | 7 | 9 | 3,1 | 3,1 | 0,0 |
| chr17:29097068-29097568:downstream_12955 | chr17 | 29036625 | 29097068 | + | SUZ12P1 | NR | 27 | -0,2 | 1 | 0 | 8,55E-01 | NA | 15 | 15 | 19 | 34 | 24 | 25 | 40 | 42 | -0,9 | -0,7 | -0,1 |
| chr17:33457840-33458340:downstream_20610 | chr17 | 33458340 | 33469334 | - | NLE1 | NM | 19 | 0,2 | 1 | 0 | 8,55E-01 | NA | 20 | 15 | 0 | 5 | 42 | 53 | 10 | 8 | 2,9 | 2,4 | 0,5 |
| chr1:178518024-178518524:downstream_1244 | chr1 | 178511930 | 178518024 | + | C1orf220 | NR | 8 | -0,3 | 2 | 0 | 8,55E-01 | NA | 7 | 0 | 11 | 11 | 3 | 7 | 8 | 15 | -1,6 | -1,2 | -0,4 |
| chr19:15771570-15772070:downstream_21089 | chr19 | 15751706 | 15771570 | + | CYP4F3 | NM | 26 | 0,2 | 1 | 0 | 8,55E-01 | NA | 10 | 8 | 8 | 17 | 26 | 35 | 57 | 46 | -0,5 | -0,8 | 0,2 |
| chr6:26171576-26172076:downstream_24796 | chr6 | 26158348 | 26171576 | + | HIST1H2BD | NM | 96 | 0,1 | 1 | 0 | 8,56E-01 | 9,73E-01 | 75 | 92 | 29 | 28 | 248 | 151 | 67 | 80 | 1,5 | 1,4 | 0,1 |
| chr2:152656979-152657479:downstream_21749 | chr2 | 152657479 | 152685009 | - | ARL5A | NM | 40 | 0,1 | 1 | 0 | 8,57E-01 | 9,73E-01 | 30 | 38 | 57 | 48 | 24 | 30 | 46 | 44 | -0,6 | -0,8 | 0,1 |
| chr4:54875457-54875957:downstream_3440 | chr4 | 54875957 | 54930815 | - | CHIC2 | NM | 18 | 0,2 | 1 | 0 | 8,57E-01 | NA | 23 | 14 | 22 | 21 | 17 | 9 | 15 | 20 | -0,3 | -0,4 | 0,2 |
| chr12:755044-755544:downstream_9965 | chr12 | 740056 | 755044 | + | LOC100049716 | NR | 8 | 0,3 | 1 | 0 | 8,57E-01 | NA | 3 | 7 | 10 | 11 | 5 | 5 | 13 | 11 | -1,0 | -1,3 | 0,3 |
| chr2:65313487-65313987:downstream_22173 | chr2 | 65313987 | 65357435 | - | RAB1A | NM | 38 | 0,2 | 1 | 0 | 8,57E-01 | 9,73E-01 | 19 | 10 | 7 | 11 | 52 | 94 | 72 | 37 | 0,6 | 0,4 | 0,2 |
| chr8:56738005-56738505:downstream_6556 | chr8 | 56685790 | 56738005 | + | TGS1 | NM | 26 | 0,2 | 1 | 0 | 8,57E-01 | NA | 13 | 22 | 15 | 5 | 50 | 45 | 31 | 28 | 0,8 | 0,7 | 0,1 |
| chr3:119540301-119540801:downstream_23184 | chr3 | 119540801 | 119813264 | - | GSK3B | NM | 9 | -0,3 | 2 | 0 | 8,57E-01 | NA | 0 | 5 | 14 | 7 | 2 | 8 | 14 | 20 | -2,1 | -1,8 | -0,3 |
| chr3:185225069-185225569:downstream_3304 | chr3 | 185225569 | 185270369 | - | LIPH | NM | 47 | 0,2 | 1 | 0 | 8,57E-01 | 9,73E-01 | 4 | 10 | 22 | 50 | 21 | 19 | 134 | 113 | -2,4 | -2,6 | 0,2 |
| chr4:139084747-139085247:downstream_3846 | chr4 | 139085247 | 139163503 | - | SLC7A11 | NM | 53 | 0,1 | 1 | 0 | 8,58E-01 | 9,73E-01 | 3 | 7 | 18 | 22 | 37 | 30 | 128 | 180 | -2,0 | -2,2 | 0,2 |
| chrX:49030650-49031150:downstream_26991 | chrX | 49031150 | 49042853 | - | PRICKLE3 | NM | 70 | 0,1 | 1 | 0 | 8,58E-01 | 9,73E-01 | 71 | 227 | 78 | 71 | 27 | 47 | 17 | 24 | 1,0 | 0,9 | 0,1 |
| chr10:104847273-104847773:downstream_17679 | chr10 | 104847773 | 104953063 | - | NT5C2 | NM | 42 | 0,2 | 1 | 0 | 8,58E-01 | 9,73E-01 | 7 | 5 | 24 | 14 | 30 | 32 | 120 | 102 | -1,7 | -1,8 | 0,1 |
| chr15:66782165-66782665:downstream_11772 | chr15 | 66782665 | 66790146 | - | SNAPC5 | NM | 65 | -0,1 | 1 | 0 | 8,58E-01 | 9,73E-01 | 90 | 57 | 18 | 11 | 176 | 116 | 28 | 25 | 2,3 | 2,5 | -0,2 |
| chr5:151148853-151149353:downstream_4528 | chr5 | 151149353 | 151151069 | - | LOC100652758 | NM | 9 | -0,2 | 1 | 0 | 8,58E-01 | NA | 11 | 15 | 10 | 6 | 8 | 14 | 3 | 7 | 0,8 | 1,1 | -0,3 |
| chr5:118730299-118730799:downstream_24420 | chr5 | 118604386 | 118730299 | + | TNFAIP8 | NM | 14 | -0,2 | 1 | 0 | 8,59E-01 | NA | 2 | 6 | 7 | 14 | 16 | 8 | 33 | 28 | -1,5 | -1,4 | -0,1 |
| chr14:50090199-50090699:downstream_11129 | chr14 | 50087488 | 50090199 | + | MGAT2 | NM | 18 | -0,2 | 1 | 0 | 8,59E-01 | NA | 10 | 7 | 6 | 6 | 44 | 31 | 18 | 26 | 0,5 | 0,8 | -0,2 |
| chr1:52883992-52884492:downstream_733 | chr1 | 52870218 | 52883992 | + | PRPF38A | NM | 25 | 0,2 | 1 | 0 | 8,59E-01 | NA | 28 | 32 | 13 | 17 | 46 | 26 | 33 | 9 | 1,0 | 0,8 | 0,2 |
| chr15:76508128-76508628:downstream_19542 | chr15 | 76508628 | 76603810 | - | ETFA | NM | 172 | -0,1 | 1 | 0 | 8,59E-01 | 9,73E-01 | 20 | 3 | 15 | 14 | 370 | 245 | 391 | 317 | -0,4 | -0,2 | -0,2 |
| chr12:49315241-49315741:downstream_18568 | chr12 | 49315741 | 49319330 | - | FKBP11 | NM | 48 | 0,1 | 1 | 0 | 8,59E-01 | 9,73E-01 | 7 | 22 | 13 | 16 | 69 | 85 | 83 | 88 | 0,0 | -0,2 | 0,2 |
| chr7:92169079-92169579:downstream_5941 | chr7 | 92167788 | 92169079 | + | MGC16142 | NR | 4 | -0,3 | 2 | 0 | 8,59E-01 | NA | 2 | 7 | 1 | 6 | 7 | 5 | 4 | 3 | 0,4 | 0,6 | -0,2 |
| chr1:45285015-45285515:downstream_17136 | chr1 | 45285515 | 45308616 | - | PTCH2 | NM | 5 | 0,3 | 2 | 0 | 8,59E-01 | NA | 1 | 7 | 4 | 6 | 4 | 4 | 4 | 9 | -0,4 | -0,7 | 0,3 |
| chr6:42997337-42997837:downstream_4658 | chr6 | 42989384 | 42997337 | + | RRP36 | NM | 39 | 0,1 | 1 | 0 | 8,60E-01 | 9,73E-01 | 13 | 15 | 19 | 22 | 56 | 35 | 99 | 56 | -0,6 | -0,8 | 0,2 |
| chr11:22843097-22843597:downstream_9186 | chr11 | 22843597 | 22851382 | - | SVIP | NM | 12 | 0,2 | 1 | 0 | 8,60E-01 | NA | 13 | 18 | 11 | 11 | 9 | 14 | 10 | 9 | 0,5 | 0,3 | 0,2 |
| chr6:7881961-7882461:downstream_4687 | chr6 | 7727010 | 7881961 | + | BMP6 | NM | 84 | 0,1 | 1 | 0 | 8,60E-01 | 9,73E-01 | 23 | 74 | 67 | 43 | 49 | 158 | 109 | 150 | -0,2 | -0,3 | 0,1 |
| chr6:108844251-108844751:downstream_4779 | chr6 | 108616097 | 108844251 | + | LACE1 | NM | 5 | 0,3 | 2 | 0 | 8,60E-01 | NA | 2 | 3 | 6 | 7 | 6 | 0 | 7 | 10 | -1,2 | -1,6 | 0,4 |
| chr11:4619401-4619901:downstream_18369 | chr11 | 4619901 | 4629489 | - | TRIM68 | NM | 12 | -0,2 | 1 | 0 | 8,60E-01 | NA | 3 | 6 | 1 | 7 | 14 | 29 | 17 | 21 | 0,0 | 0,2 | -0,1 |
| chr5:137273141-137273641:downstream_24114 | chr5 | 137273641 | 137368802 | - | FAM13B | NM | 7 | 0,3 | 2 | 0 | 8,61E-01 | NA | 4 | 1 | 8 | 11 | 3 | 2 | 13 | 12 | -2,0 | -2,3 | 0,3 |
| chr8:95383687-95384187:downstream_26078 | chr8 | 95384187 | 95487343 | - | RAD54B | NM | 7 | 0,3 | 2 | 0 | 8,61E-01 | NA | 3 | 5 | 4 | 16 | 7 | 0 | 8 | 12 | -1,3 | -1,7 | 0,4 |
| chr16:14397906-14398406:downstream_12201 | chr16 | 14397823 | 14397906 | + | MIR193B | NR | 12 | -0,2 | 1 | 0 | 8,62E-01 | NA | 4 | 9 | 13 | 10 | 10 | 15 | 22 | 15 | -0,8 | -0,6 | -0,2 |
| chr19:51600362-51600862:downstream_14523 | chr19 | 51600862 | 51611647 | - | CTU1 | NM | 9 | -0,2 | 1 | 0 | 8,62E-01 | NA | 19 | 14 | 14 | 5 | 7 | 10 | 6 | 2 | 0,8 | 1,0 | -0,2 |
| chr15:25581895-25582395:downstream_19827 | chr15 | 25582395 | 25684175 | - | UBE3A | NM | 26 | 0,2 | 1 | 0 | 8,62E-01 | NA | 4 | 2 | 21 | 21 | 11 | 8 | 81 | 61 | -2,8 | -2,9 | 0,2 |
| chr12:101780397-101780897:downstream_9794 | chr12 | 101673904 | 101780397 | + | UTP20 | NM | 12 | 0,3 | 2 | 0 | 8,63E-01 | NA | 0 | 2 | 10 | 6 | 3 | 4 | 43 | 30 | -3,0 | -3,4 | 0,4 |
| chr17:57774166-57774666:downstream_20659 | chr17 | 57774666 | 57784959 | - | PTRH2 | NM | 63 | -0,1 | 1 | 0 | 8,63E-01 | 9,73E-01 | 12 | 20 | 18 | 16 | 127 | 99 | 96 | 118 | -0,1 | 0,1 | -0,1 |
| chr3:108268217-108268717:downstream_3091 | chr3 | 108268717 | 108308491 | - | KIAA1524 | NM | 13 | -0,2 | 1 | 0 | 8,63E-01 | NA | 5 | 5 | 18 | 22 | 2 | 9 | 19 | 20 | -2,1 | -1,8 | -0,2 |
| chr7:104650488-104650988:downstream_6032 | chr7 | 104650988 | 104654588 | - | KMT2E-AS1 | NR | 58 | 0,1 | 1 | 0 | 8,64E-01 | 9,73E-01 | 98 | 105 | 26 | 45 | 68 | 69 | 22 | 30 | 1,5 | 1,4 | 0,1 |
| chr7:156473069-156473569:downstream_5663 | chr7 | 156473569 | 156685902 | - | LMBR1 | NM | 37 | 0,1 | 1 | 0 | 8,64E-01 | 9,73E-01 | 6 | 9 | 28 | 34 | 20 | 17 | 106 | 75 | -2,1 | -2,3 | 0,2 |
| chr14:73741347-73741847:downstream_11210 | chr14 | 73704204 | 73741347 | + | PAPLN | NM | 12 | -0,2 | 1 | 0 | 8,64E-01 | NA | 4 | 2 | 4 | 8 | 13 | 15 | 29 | 21 | -1,0 | -0,9 | -0,2 |
| chr2:220113932-220114432:downstream_22306 | chr2 | 220114432 | 220119330 | - | TUBA4A | NM | 6 | 0,3 | 2 | 0 | 8,64E-01 | NA | 10 | 16 | 1 | 2 | 7 | 10 | 1 | 2 | 2,8 | 2,4 | 0,4 |
| chr9:97849500-97850000:downstream_26238 | chr9 | 97488950 | 97849500 | + | C9orf3 | NM | 90 | 0,1 | 1 | 0 | 8,64E-01 | 9,73E-01 | 4 | 1 | 17 | 19 | 41 | 38 | 287 | 312 | -2,8 | -2,9 | 0,1 |
| chr15:73851843-73852343:downstream_19649 | chr15 | 73852343 | 73925753 | - | NPTN | NM | 104 | 0,1 | 1 | 0 | 8,64E-01 | 9,73E-01 | 12 | 6 | 7 | 17 | 117 | 197 | 228 | 250 | -0,4 | -0,6 | 0,2 |
| chr9:4709056-4709556:downstream_26212 | chr9 | 4709556 | 4742043 | - | AK3 | NM | 10 | 0,2 | 1 | 0 | 8,64E-01 | NA | 7 | 13 | 4 | 4 | 20 | 15 | 8 | 9 | 1,2 | 1,0 | 0,2 |
| chr17:41297125-41297625:downstream_12645 | chr17 | 41277599 | 41297125 | + | NBR2 | NR | 10 | 0,2 | 1 | 0 | 8,64E-01 | NA | 2 | 8 | 7 | 8 | 15 | 5 | 25 | 14 | -0,7 | -1,0 | 0,3 |
| chr7:89866992-89867492:downstream_25754 | chr7 | 89840999 | 89866992 | + | STEAP2 | NM | 35 | 0,1 | 1 | 0 | 8,65E-01 | 9,73E-01 | 8 | 6 | 25 | 11 | 33 | 28 | 81 | 89 | -1,4 | -1,5 | 0,1 |
| chr6:3019889-3020389:downstream_4845 | chr6 | 3020389 | 3025005 | - | HTATSF1P2 | NR | 36 | -0,1 | 1 | 0 | 8,65E-01 | 9,73E-01 | 40 | 39 | 54 | 37 | 35 | 24 | 21 | 40 | -0,2 | 0,0 | -0,2 |
| chr1:229795946-229796446:downstream_308 | chr1 | 229761980 | 229795946 | + | URB2 | NM | 12 | 0,2 | 1 | 0 | 8,66E-01 | NA | 13 | 12 | 25 | 22 | 6 | 1 | 7 | 8 | -0,9 | -1,2 | 0,2 |
| chr8:23286164-23286664:downstream_25920 | chr8 | 23286664 | 23315244 | - | ENTPD4 | NM | 28 | 0,2 | 1 | 0 | 8,66E-01 | NA | 12 | 5 | 15 | 27 | 13 | 30 | 54 | 65 | -1,3 | -1,5 | 0,2 |
| chr10:95351092-95351592:downstream_8645 | chr10 | 95351592 | 95360993 | - | RBP4 | NM | 26 | -0,2 | 1 | 0 | 8,66E-01 | NA | 11 | 11 | 13 | 6 | 50 | 47 | 33 | 39 | 0,2 | 0,4 | -0,2 |
| chr7:7676074-7676574:downstream_5507 | chr7 | 7676574 | 7758238 | - | RPA3 | NM | 38 | 0,1 | 1 | 0 | 8,66E-01 | 9,73E-01 | 35 | 41 | 43 | 41 | 37 | 27 | 36 | 41 | -0,2 | -0,3 | 0,1 |
| chr3:132396944-132397444:downstream_23496 | chr3 | 132373289 | 132396944 | + | UBA5 | NM | 19 | 0,2 | 1 | 0 | 8,66E-01 | NA | 22 | 26 | 25 | 28 | 13 | 9 | 8 | 18 | -0,1 | -0,3 | 0,1 |
| chr9:116853417-116853917:downstream_7272 | chr9 | 116853917 | 116861337 | - | KIF12 | NM | 8 | 0,3 | 2 | 0 | 8,67E-01 | NA | 4 | 1 | 8 | 7 | 2 | 8 | 18 | 14 | -1,6 | -1,8 | 0,2 |
| chr3:149086304-149086804:downstream_3225 | chr3 | 149086804 | 149095568 | - | TM4SF1 | NM | 2668 | -0,2 | 1 | 0 | 8,67E-01 | 9,73E-01 | 20 | 2 | 38 | 21 | 3604 | 3013 | 8280 | 6365 | -1,4 | -1,1 | -0,2 |
| chr17:8271470-8271970:downstream_20535 | chr17 | 8271970 | 8280029 | - | KRBA2 | NM | 34 | -0,1 | 1 | 0 | 8,67E-01 | 9,73E-01 | 41 | 47 | 21 | 14 | 54 | 54 | 25 | 14 | 1,4 | 1,5 | -0,1 |
| chr15:59954561-59955061:downstream_11741 | chr15 | 59955061 | 59981642 | - | BNIP2 | NM | 45 | 0,1 | 1 | 0 | 8,67E-01 | 9,73E-01 | 40 | 32 | 43 | 48 | 41 | 43 | 61 | 54 | -0,3 | -0,5 | 0,1 |
| chr9:33039062-33039562:downstream_6854 | chr9 | 33025208 | 33039062 | + | DNAJA1 | NM | 311 | -0,1 | 0 | 0 | 8,68E-01 | 9,73E-01 | 208 | 217 | 223 | 225 | 488 | 321 | 411 | 397 | -0,1 | 0,0 | -0,1 |
| chr6:109765122-109765622:downstream_5296 | chr6 | 109761930 | 109765122 | + | SMPD2 | NM | 4 | 0,3 | 2 | 0 | 8,68E-01 | NA | 5 | 5 | 3 | 4 | 6 | 3 | 0 | 7 | 0,5 | 0,2 | 0,3 |
| chr7:6500211-6500711:downstream_25585 | chr7 | 6500711 | 6523849 | - | KDELR2 | NM | 185 | -0,1 | 1 | 0 | 8,68E-01 | 9,73E-01 | 7 | 3 | 8 | 6 | 340 | 326 | 402 | 387 | -0,5 | -0,2 | -0,2 |
| chr22:17539682-17540182:downstream_15449 | chr22 | 17517459 | 17539682 | + | CECR7 | NR | 5 | 0,3 | 2 | 0 | 8,68E-01 | NA | 9 | 6 | 3 | 7 | 9 | 0 | 1 | 6 | 0,7 | 0,3 | 0,3 |
| chr7:8382911-8383411:downstream_5710 | chr7 | 8301854 | 8382911 | + | LOC100505938 | NR | 8 | -0,3 | 2 | 0 | 8,69E-01 | NA | 2 | 5 | 3 | 13 | 11 | 3 | 11 | 15 | -1,1 | -0,9 | -0,2 |
| chr14:93306306-93306806:downstream_11274 | chr14 | 93260575 | 93306306 | + | GOLGA5 | NM | 24 | -0,2 | 1 | 0 | 8,69E-01 | NA | 7 | 1 | 13 | 6 | 22 | 34 | 45 | 66 | -1,3 | -1,0 | -0,3 |
| chrX:46334074-46334574:downstream_26875 | chrX | 46306623 | 46334074 | + | KRBOX4 | NM | 9 | -0,3 | 2 | 0 | 8,69E-01 | NA | 7 | 9 | 6 | 2 | 3 | 32 | 6 | 9 | 0,9 | 1,3 | -0,3 |
| chr7:134850967-134851467:downstream_6100 | chr7 | 134832765 | 134850967 | + | TMEM140 | NM | 9 | -0,2 | 1 | 0 | 8,70E-01 | NA | 4 | 5 | 10 | 11 | 6 | 9 | 14 | 16 | -1,3 | -1,0 | -0,2 |
| chr1:78482995-78483495:downstream_797 | chr1 | 78470635 | 78482995 | + | DNAJB4 | NM | 7 | -0,3 | 2 | 0 | 8,70E-01 | NA | 7 | 2 | 6 | 8 | 10 | 5 | 8 | 10 | -0,6 | -0,3 | -0,3 |
| chr20:34820721-34821221:downstream_22420 | chr20 | 34679425 | 34820721 | + | EPB41L1 | NM | 12 | -0,2 | 1 | 0 | 8,70E-01 | NA | 5 | 3 | 7 | 9 | 17 | 9 | 25 | 21 | -1,0 | -0,8 | -0,2 |
| chr12:81190670-81191170:downstream_9890 | chr12 | 81191170 | 81331694 | - | LIN7A | NM | 7 | 0,3 | 2 | 0 | 8,70E-01 | NA | 2 | 2 | 18 | 11 | 2 | 0 | 14 | 7 | -2,8 | -3,1 | 0,4 |
| chr3:9798528-9799028:downstream_2807 | chr3 | 9799028 | 9811668 | - | CAMK1 | NM | 6 | 0,3 | 2 | 0 | 8,70E-01 | NA | 1 | 10 | 4 | 5 | 6 | 7 | 10 | 4 | 0,2 | -0,2 | 0,4 |
| chr8:23119512-23120012:downstream_6494 | chr8 | 23101149 | 23119512 | + | CHMP7 | NM | 13 | 0,2 | 1 | 0 | 8,70E-01 | NA | 7 | 4 | 7 | 7 | 13 | 19 | 26 | 18 | -0,3 | -0,5 | 0,2 |
| chr20:36152090-36152590:downstream_22487 | chr20 | 36149606 | 36152090 | + | NNAT | NM | 7 | -0,4 | 2 | 0 | 8,70E-01 | NA | 6 | 25 | 3 | 0 | 2 | 16 | 1 | 0 | 3,5 | 3,8 | -0,3 |
| chr12:6571488-6571988:downstream_9992 | chr12 | 6561176 | 6571488 | + | TAPBPL | NM | 15 | 0,2 | 1 | 0 | 8,70E-01 | NA | 10 | 3 | 11 | 4 | 22 | 19 | 24 | 30 | -0,3 | -0,4 | 0,1 |
| chr11:14525921-14526421:downstream_18234 | chr11 | 14526421 | 14665180 | - | PSMA1 | NM | 106 | -0,1 | 1 | 0 | 8,71E-01 | 9,76E-01 | 5 | 7 | 3 | 11 | 176 | 192 | 205 | 252 | -0,3 | -0,3 | 0,0 |
| chr13:79998468-79998968:downstream_10811 | chr13 | 79980443 | 79998468 | + | RBM26-AS1 | NR | 6 | -0,4 | 2 | 0 | 8,71E-01 | NA | 16 | 4 | 1 | 0 | 11 | 15 | 1 | 0 | 3,9 | 4,3 | -0,4 |
| chr12:118855840-118856340:downstream_10440 | chr12 | 118814357 | 118855840 | + | SUDS3 | NM | 17 | 0,2 | 1 | 0 | 8,72E-01 | NA | 2 | 5 | 21 | 25 | 7 | 3 | 40 | 30 | -2,6 | -2,9 | 0,2 |
| chr19:9638180-9638680:downstream_21642 | chr19 | 9638680 | 9649321 | - | ZNF426 | NM | 16 | -0,2 | 1 | 0 | 8,72E-01 | NA | 21 | 18 | 15 | 17 | 17 | 16 | 15 | 10 | 0,3 | 0,4 | -0,1 |
| chr8:117656554-117657054:downstream_6343 | chr8 | 117657054 | 117768062 | - | EIF3H | NM | 76 | 0,1 | 1 | 0 | 8,73E-01 | 9,76E-01 | 12 | 3 | 8 | 7 | 143 | 141 | 127 | 166 | 0,0 | 0,0 | 0,1 |
| chr12:56214959-56215459:downstream_10245 | chr12 | 56211805 | 56214959 | + | ORMDL2 | NM | 72 | 0,1 | 1 | 0 | 8,73E-01 | 9,76E-01 | 54 | 44 | 42 | 33 | 138 | 84 | 109 | 73 | 0,4 | 0,3 | 0,1 |
| chr2:37868524-37869024:downstream_21810 | chr2 | 37869024 | 37899678 | - | CDC42EP3 | NM | 40 | 0,1 | 1 | 0 | 8,73E-01 | 9,76E-01 | 29 | 35 | 3 | 13 | 93 | 91 | 21 | 33 | 2,0 | 1,8 | 0,2 |
| chr20:60777810-60778310:downstream_15126 | chr20 | 60758080 | 60777810 | + | MTG2 | NM | 11 | -0,2 | 1 | 0 | 8,73E-01 | NA | 13 | 16 | 10 | 8 | 15 | 11 | 8 | 6 | 0,7 | 0,9 | -0,2 |

| | | | | | | | | | | | | | | | | | | | | | | | |
|---|---|---|---|---|---|---|---|---|---|---|---|---|---|---|---|---|---|---|---|---|---|---|---|
| chr20:33863635-33864135:downstream_22473 | chr20 | 33864135 | 33865960 | - | MMP24-AS1 | NR | 9 | -0,2 | 1 | 0 | 8,73E-01 | NA | 10 | 5 | 4 | 6 | 20 | 11 | 8 | 10 | 0,6 | 0,8 | -0,2 |
| chr20:36145318-36145818:downstream_22364 | chr20 | 36145818 | 36156333 | - | BLCAP | NM | 19 | 0,2 | 1 | 0 | 8,74E-01 | NA | 4 | 15 | 3 | 2 | 49 | 46 | 13 | 18 | 1,8 | 1,6 | 0,2 |
| chr18:48514490-48514990:downstream_13513 | chr18 | 48494386 | 48514490 | + | ELAC1 | NM | 5 | 0,3 | 2 | 0 | 8,74E-01 | NA | 10 | 8 | 3 | 1 | 3 | 10 | 1 | 2 | 2,3 | 2,1 | 0,2 |
| chr19:42732353-42732853:downstream_13778 | chr19 | 42724491 | 42732353 | + | ZNF526 | NM | 28 | -0,1 | 1 | 0 | 8,74E-01 | 9,76E-01 | 26 | 20 | 31 | 18 | 45 | 23 | 46 | 19 | -0,1 | 0,1 | -0,1 |
| chr19:12740676-12741176:downstream_14033 | chr19 | 12721731 | 12740676 | + | ZNF791 | NM | 26 | -0,1 | 1 | 0 | 8,74E-01 | NA | 42 | 35 | 35 | 28 | 22 | 17 | 21 | 9 | 0,3 | 0,4 | -0,1 |
| chr16:767480-767980:downstream_12068 | chr16 | 765172 | 767480 | + | METRN | NM | 20 | 0,1 | 1 | 0 | 8,75E-01 | NA | 20 | 21 | 19 | 22 | 19 | 16 | 25 | 16 | 0,0 | -0,2 | 0,2 |
| chr19:37734574-37735074:downstream_14249 | chr19 | 37717365 | 37734574 | + | ZNF383 | NM | 17 | 0,2 | 1 | 0 | 8,75E-01 | NA | 19 | 30 | 13 | 11 | 24 | 15 | 14 | 8 | 1,0 | 0,8 | 0,2 |
| chr7:99660969-99661469:downstream_25824 | chr7 | 99661469 | 99680171 | - | ZNF3 | NM | 12 | -0,2 | 1 | 0 | 8,76E-01 | NA | 4 | 9 | 6 | 4 | 29 | 16 | 15 | 15 | 0,4 | 0,6 | -0,2 |
| chr9:139618545-139619045:downstream_26508 | chr9 | 139619045 | 139622636 | - | SNHG7 | NR | 52 | -0,1 | 1 | 0 | 8,76E-01 | 9,76E-01 | 6 | 13 | 11 | 12 | 88 | 82 | 85 | 119 | -0,4 | -0,3 | -0,1 |
| chr7:128697293-128697793:downstream_6080 | chr7 | 128695276 | 128697293 | + | TPI1P2 | NR | 4 | 0,4 | 2 | 0 | 8,76E-01 | NA | 5 | 11 | 0 | 2 | 6 | 8 | 1 | 1 | 3,2 | 2,6 | 0,6 |
| chr17:73772014-73772514:downstream_13365 | chr17 | 73772514 | 73776016 | - | H3F3B | NM | 41 | 0,1 | 1 | 0 | 8,76E-01 | 9,76E-01 | 7 | 9 | 4 | 4 | 102 | 90 | 67 | 44 | 1,0 | 0,8 | 0,2 |
| chr1:19628701-19629201:downstream_512 | chr1 | 19629201 | 19638640 | - | AKR7A2 | NM | 4 | -0,3 | 2 | 0 | 8,76E-01 | NA | 3 | 1 | 7 | 7 | 5 | 0 | 7 | 6 | -1,7 | -1,4 | -0,3 |
| chr4:83841284-83841784:downstream_3762 | chr4 | 83821836 | 83841284 | + | THAP9 | NM | 7 | 0,2 | 2 | 0 | 8,78E-01 | NA | 3 | 7 | 3 | 8 | 7 | 7 | 8 | 11 | -0,1 | -0,4 | 0,3 |
| chr5:68576018-68576518:downstream_24046 | chr5 | 68576518 | 68628620 | - | CCDC125 | NM | 10 | 0,2 | 1 | 0 | 8,78E-01 | NA | 1 | 8 | 10 | 11 | 5 | 9 | 22 | 18 | -1,3 | -1,6 | 0,3 |
| chr9:123939914-123940414:downstream_7288 | chr9 | 123940414 | 123964365 | - | RAB14 | NM | 28 | 0,2 | 1 | 0 | 8,78E-01 | 9,76E-01 | 43 | 62 | 36 | 11 | 17 | 30 | 19 | 5 | 1,1 | 0,9 | 0,2 |
| chr13:114545814-114546314:downstream_10897 | chr13 | 114518582 | 114545814 | + | GAS6-AS1 | NR | 6 | -0,2 | 2 | 0 | 8,78E-01 | NA | 9 | 10 | 8 | 6 | 4 | 7 | 6 | 2 | 0,4 | 0,6 | -0,2 |
| chr5:85916583-85917083:downstream_4333 | chr5 | 85913783 | 85916583 | + | COX7C | NM | 42 | 0,1 | 1 | 0 | 8,78E-01 | 9,76E-01 | 59 | 48 | 32 | 47 | 46 | 39 | 32 | 36 | 0,4 | 0,3 | 0,1 |
| chr16:14360636-14361136:downstream_20119 | chr16 | 14165169 | 14360636 | + | MKL2 | NM | 12 | 0,3 | 2 | 0 | 8,78E-01 | NA | 2 | 0 | 8 | 9 | 0 | 6 | 35 | 33 | -3,4 | -3,5 | 0,1 |
| chr1:234739514-234740014:downstream_16830 | chr1 | 234740014 | 234745271 | - | IRF2BP2 | NM | 128 | 0,1 | 1 | 0 | 8,78E-01 | 9,76E-01 | 105 | 56 | 173 | 123 | 89 | 104 | 185 | 189 | -0,9 | -1,0 | 0,1 |
| chr12:133696054-133696554:downstream_10534 | chr12 | 133696554 | 133707059 | - | ZNF891 | NM | 6 | 0,3 | 2 | 0 | 8,79E-01 | NA | 7 | 6 | 0 | 7 | 2 | 15 | 6 | 7 | 0,8 | 0,5 | 0,4 |
| chr18:33291798-33292298:downstream_13607 | chr18 | 33234532 | 33291798 | + | GALNT1 | NM | 30 | -0,1 | 1 | 0 | 8,79E-01 | 9,76E-01 | 3 | 7 | 14 | 12 | 34 | 26 | 64 | 80 | -1,4 | -1,3 | -0,1 |
| chr7:156430559-156431059:downstream_6212 | chr7 | 156431059 | 156433348 | - | C7orf13 | NR | 8 | 0,3 | 2 | 0 | 8,79E-01 | NA | 24 | 14 | 1 | 5 | 12 | 3 | 3 | 1 | 2,6 | 2,1 | 0,5 |
| chr16:69377013-69377513:downstream_20142 | chr16 | 69373414 | 69377013 | + | NIP7 | NM | 65 | 0,2 | 1 | 0 | 8,80E-01 | 9,76E-01 | 24 | 1 | 6 | 6 | 148 | 179 | 81 | 78 | 1,2 | 1,0 | 0,1 |
| chr5:204374-204874:downstream_4145 | chr5 | 204874 | 218297 | - | CCDC127 | NM | 35 | 0,1 | 1 | 0 | 8,80E-01 | 9,76E-01 | 11 | 14 | 6 | 7 | 59 | 95 | 42 | 51 | 0,9 | 0,7 | 0,2 |
| chr15:34633416-34633916:downstream_11642 | chr15 | 34633916 | 34635362 | - | NOP10 | NM | 137 | -0,1 | 1 | 0 | 8,81E-01 | 9,76E-01 | 21 | 22 | 11 | 14 | 394 | 267 | 173 | 191 | 0,8 | 0,9 | -0,1 |
| chr3:134196045-134196545:downstream_23000 | chr3 | 134196545 | 134204865 | - | ANAPC13 | NM | 41 | -0,1 | 1 | 0 | 8,81E-01 | 9,76E-01 | 5 | 7 | 3 | 3 | 97 | 113 | 50 | 51 | 0,9 | 1,1 | -0,1 |
| chr10:86278277-86278777:downstream_17514 | chr10 | 86088344 | 86278277 | + | CCSER2 | NM | 9 | 0,4 | 2 | 0 | 8,82E-01 | NA | 0 | 0 | 3 | 12 | 1 | 0 | 29 | 25 | -Inf | -6,1 | -Inf |
| chr8:145141119-145141619:downstream_6779 | chr8 | 145137523 | 145141119 | + | GPAA1 | NM | 12 | 0,2 | 1 | 0 | 8,82E-01 | NA | 18 | 9 | 3 | 5 | 21 | 23 | 10 | 6 | 1,8 | 1,5 | 0,3 |
| chr5:10266524-10267024:downstream_24053 | chr5 | 10250032 | 10266524 | + | CCT5 | NM | 12 | 0,2 | 1 | 0 | 8,82E-01 | NA | 6 | 10 | 8 | 2 | 20 | 18 | 17 | 12 | 0,5 | 0,4 | 0,1 |
| chr1:203278729-203279229:downstream_1309 | chr1 | 203274663 | 203278729 | + | BTG2 | NM | 102 | 0,1 | 1 | 0 | 8,82E-01 | 9,76E-01 | 89 | 102 | 4 | 2 | 295 | 305 | 7 | 11 | 5,0 | 5,1 | -0,1 |
| chr12:52471279-52471779:downstream_18445 | chr12 | 52463757 | 52471279 | + | ATG101 | NM | 60 | 0,1 | 1 | 0 | 8,82E-01 | 9,76E-01 | 11 | 8 | 6 | 6 | 151 | 118 | 102 | 80 | 0,7 | 0,6 | 0,1 |
| chr20:23608033-23608533:downstream_22394 | chr20 | 23608533 | 23618685 | - | CST3 | NM | 53 | 0,1 | 1 | 0 | 8,82E-01 | 9,76E-01 | 97 | 44 | 18 | 4 | 146 | 76 | 18 | 18 | 2,7 | 2,6 | 0,0 |
| chr4:184938875-184939375:downstream_3922 | chr4 | 184826508 | 184938875 | + | STOX2 | NM | 8 | 0,2 | 1 | 0 | 8,84E-01 | NA | 4 | 11 | 7 | 11 | 5 | 10 | 10 | 11 | -0,3 | -0,5 | 0,2 |
| chr8:31031277-31031777:downstream_6283 | chr8 | 30890777 | 31031277 | + | WRN | NM | 6 | 0,3 | 2 | 0 | 8,84E-01 | NA | 4 | 0 | 7 | 7 | 3 | 4 | 14 | 13 | -1,7 | -1,9 | 0,2 |
| chr8:22398657-22399157:downstream_26064 | chr8 | 22298482 | 22398657 | + | PPP3CC | NM | 12 | 0,2 | 1 | 0 | 8,84E-01 | NA | 1 | 4 | 6 | 5 | 8 | 14 | 29 | 28 | -1,2 | -1,4 | 0,2 |
| chr9:100846134-100846634:downstream_26552 | chr9 | 100846634 | 100881635 | - | TRIM14 | NM | 7 | -0,2 | 2 | 0 | 8,84E-01 | NA | 8 | 8 | 6 | 16 | 6 | 4 | 7 | 5 | -0,5 | -0,3 | -0,2 |
| chr2:68268831-68269331:downstream_21776 | chr2 | 68269331 | 68290159 | - | C1D | NM | 13 | -0,2 | 1 | 0 | 8,84E-01 | NA | 10 | 13 | 7 | 4 | 26 | 25 | 8 | 13 | 1,0 | 1,3 | -0,2 |
| chr5:140090958-140091458:downstream_4466 | chr5 | 140090860 | 140090958 | + | VTRNA1-1 | NR | 23 | 0,2 | 1 | 0 | 8,85E-01 | NA | 58 | 52 | 3 | 2 | 37 | 33 | 1 | 2 | 4,6 | 4,5 | 0,1 |
| chr5:89706486-89706986:downstream_4340 | chr5 | 89705816 | 89706486 | + | LOC731157 | NR | 32 | 0,1 | 1 | 0 | 8,85E-01 | 9,79E-01 | 66 | 61 | 31 | 20 | 31 | 22 | 11 | 11 | 1,3 | 1,2 | 0,1 |
| chr14:65211064-65211564:downstream_11183 | chr14 | 65171125 | 65211064 | + | PLEKHG3 | NM | 10 | 0,2 | 2 | 0 | 8,85E-01 | NA | 1 | 4 | 13 | 2 | 4 | 10 | 25 | 25 | -1,7 | -1,9 | 0,2 |
| chr11:73009670-73010170:downstream_18186 | chr11 | 72975549 | 73009670 | + | P2RY6 | NM | 10 | -0,2 | 1 | 0 | 8,86E-01 | NA | 2 | 9 | 15 | 7 | 5 | 13 | 18 | 11 | -1,0 | -0,8 | -0,2 |
| chr16:89784573-89785073:downstream_12039 | chr16 | 89778263 | 89784573 | + | VPS9D1-AS1 | NR | 7 | 0,2 | 2 | 0 | 8,86E-01 | NA | 15 | 15 | 8 | 2 | 10 | 3 | 1 | 3 | 1,5 | 1,4 | 0,1 |
| chrX:119010629-119011129:downstream_7901 | chrX | 119005733 | 119010629 | + | NDUFA1 | NM | 55 | 0,1 | 1 | 0 | 8,87E-01 | 9,79E-01 | 5 | 4 | 3 | 4 | 111 | 116 | 89 | 112 | 0,3 | 0,2 | 0,2 |
| chr3:191116459-191116959:downstream_23048 | chr3 | 191046873 | 191116459 | + | CCDC50 | NM | 6 | 0,2 | 2 | 0 | 8,87E-01 | NA | 7 | 4 | 6 | 10 | 5 | 2 | 6 | 6 | -0,5 | -0,7 | 0,3 |
| chr7:152343086-152343586:downstream_6201 | chr7 | 152343586 | 152373250 | - | XRCC2 | NM | 57 | 0,1 | 1 | 0 | 8,88E-01 | 9,79E-01 | 57 | 59 | 114 | 121 | 20 | 12 | 49 | 23 | -1,0 | -1,2 | 0,1 |
| chr5:180682885-180683385:downstream_4638 | chr5 | 180683385 | 180688119 | - | TRIM52 | NM | 31 | -0,1 | 1 | 0 | 8,88E-01 | 9,79E-01 | 44 | 51 | 39 | 48 | 20 | 16 | 19 | 12 | 0,1 | 0,2 | -0,1 |
| chr14:77857587-77858087:downstream_10971 | chr14 | 77843761 | 77857587 | + | SAMD15 | NM | 5 | -0,2 | 2 | 0 | 8,89E-01 | NA | 3 | 4 | 4 | 7 | 3 | 6 | 7 | 6 | -0,7 | -0,5 | -0,2 |
| chr1:35887545-35888045:downstream_109 | chr1 | 35734567 | 35887545 | + | ZMYM4 | NM | 11 | -0,3 | 2 | 0 | 8,89E-01 | NA | 0 | | 8 | 7 | 0 | 2 | 29 | 41 | -Inf | -5,2 | -Inf |
| chr8:42978323-42978823:downstream_26058 | chr8 | 42948648 | 42978323 | + | POMK | NM | 31 | 0,1 | 1 | 0 | 8,90E-01 | 9,79E-01 | 26 | 27 | 39 | 33 | 20 | 29 | 39 | 33 | -0,4 | -0,5 | 0,1 |
| chr11:67818366-67818866:downstream_18334 | chr11 | 67806461 | 67818366 | + | TCIRG1 | NM | 15 | 0,2 | 1 | 0 | 8,90E-01 | NA | 9 | 10 | 11 | 12 | 23 | 9 | 18 | 26 | -0,3 | -0,5 | 0,2 |
| chr12:6600815-6601315:downstream_9993 | chr12 | 6601315 | 6602471 | - | MRPL51 | NM | 196 | -0,1 | 1 | 0 | 8,90E-01 | 9,79E-01 | 15 | 7 | 8 | 6 | 566 | 403 | 216 | 348 | 0,6 | 0,8 | -0,2 |
| chr1:24863510-24864010:downstream_17166 | chr1 | 24828840 | 24863510 | + | RCAN3 | NM | 20 | -0,1 | 1 | 0 | 8,90E-01 | NA | 7 | 9 | 35 | 21 | 7 | 14 | 38 | 29 | -1,9 | -1,7 | -0,2 |
| chr3:120405027-120405527:downstream_2676 | chr3 | 120405527 | 120461384 | - | RABL3 | NM | 10 | 0,2 | 2 | 0 | 8,91E-01 | NA | 5 | 0 | 13 | 11 | 7 | 1 | 21 | 20 | -2,2 | -2,4 | 0,2 |
| chr10:106027222-106027722:downstream_17590 | chr10 | 106013951 | 106027222 | + | GSTO1 | NM | 12 | 0,2 | 1 | 0 | 8,91E-01 | NA | 6 | 3 | 4 | 8 | 15 | 12 | 26 | 19 | -0,5 | -0,7 | 0,2 |
| chr4:142948681-142949181:downstream_23705 | chr4 | 142949181 | 143767604 | - | INPP4B | NM | 14 | 0,2 | 1 | 0 | 8,91E-01 | NA | 5 | 5 | 19 | 25 | 2 | 7 | 28 | 20 | -2,2 | -2,4 | 0,2 |
| chr6:99981059-99981559:downstream_5279 | chr6 | 99968869 | 99981059 | + | TSTD3 | NM | 14 | 0,2 | 1 | 0 | 8,91E-01 | NA | 13 | 11 | 8 | 11 | 23 | 12 | 13 | 20 | 0,3 | 0,1 | 0,2 |
| chr13:45005778-45006278:downstream_19120 | chr13 | 45006278 | 45150701 | - | TSC22D1 | NM | 10 | -0,2 | 1 | 0 | 8,91E-01 | NA | 13 | 6 | 8 | 5 | 14 | 16 | 11 | 7 | 0,5 | 0,7 | -0,2 |
| chr5:154256352-154256852:downstream_24068 | chr5 | 154237808 | 154256352 | + | CNOT8 | NM | 24 | -0,2 | 1 | 0 | 8,92E-01 | NA | 1 | 3 | 11 | 4 | 28 | 10 | 58 | 79 | -2,0 | -1,8 | -0,2 |
| chr10:5806685-5807185:downstream_17586 | chr10 | 5807185 | 5855512 | - | GDI2 | NM | 255 | 0,1 | 1 | 0 | 8,92E-01 | 9,79E-01 | 20 | 10 | 18 | 23 | 424 | 381 | 591 | 573 | -0,5 | -0,5 | 0,1 |
| chr5:43705668-43706168:downstream_24252 | chr5 | 43602790 | 43705668 | + | NNT | NM | 28 | 0,2 | 1 | 0 | 8,92E-01 | 9,79E-01 | 2 | 0 | 8 | 5 | 20 | 13 | 106 | 73 | -2,4 | -2,5 | 0,0 |
| chrX:47781655-47782155:downstream_7493 | chrX | 47696300 | 47781655 | + | ZNF81 | NM | 6 | -0,2 | 2 | 0 | 8,92E-01 | NA | 3 | 0 | 4 | 12 | 3 | 3 | 8 | 17 | -2,3 | -2,1 | -0,3 |
| chr1:153964667-153965167:downstream_17040 | chr1 | 153965167 | 154127592 | - | NUP210L | NM | 11 | 0,2 | 2 | 0 | 8,93E-01 | NA | 2 | 8 | 3 | 1 | 12 | 40 | 13 | 7 | 1,5 | 1,4 | 0,1 |
| chr17:46146913-46147413:downstream_20378 | chr17 | 46147413 | 46178883 | - | CBX1 | NM | 43 | -0,1 | 1 | 0 | 8,93E-01 | 9,79E-01 | 2 | 4 | 4 | 6 | 50 | 56 | 121 | 103 | -1,2 | -1,1 | -0,1 |
| chr10:119039499-119039999:downstream_8741 | chr10 | 119039999 | 119134978 | - | PDZD8 | NM | 268 | 0,1 | 1 | 0 | 8,93E-01 | 9,79E-01 | 117 | 75 | 507 | 331 | 94 | 107 | 483 | 431 | -2,1 | -2,2 | 0,1 |
| chr10:43978007-43978507:downstream_8489 | chr10 | 43932573 | 43978007 | + | ZNF487 | NR | 7 | -0,2 | 2 | 0 | 8,94E-01 | NA | 3 | 0 | 10 | 11 | 1 | 4 | 8 | 16 | -2,7 | -2,4 | -0,4 |
| chr11:18501357-18501857:downstream_9170 | chr11 | 18501857 | 18548503 | - | TSG101 | NM | 22 | 0,1 | 1 | 0 | 8,94E-01 | NA | 7 | 2 | 7 | 6 | 28 | 35 | 46 | 45 | -0,5 | -0,5 | 0,1 |
| chr6:83878190-83878690:downstream_24730 | chr6 | 83777384 | 83878190 | + | DOPEY1 | NM | 8 | 0,2 | 2 | 0 | 8,94E-01 | NA | 1 | 3 | 6 | 8 | 3 | 6 | 22 | 18 | -1,9 | -2,2 | 0,3 |
| chr20:32867570-32868070:downstream_22348 | chr20 | 32868070 | 32899608 | - | AHCY | NM | 290 | -0,1 | 1 | 0 | 8,94E-01 | 9,79E-01 | 11 | 23 | 21 | 12 | 718 | 463 | 550 | 523 | 0,0 | 0,1 | -0,1 |
| chr1:78411666-78412166:downstream_796 | chr1 | 78412166 | 78444889 | - | FUBP1 | NR | 46 | 0,1 | 1 | 0 | 8,94E-01 | 9,79E-01 | 72 | 58 | 46 | 50 | 34 | 44 | 35 | 27 | 0,4 | 0,3 | 0,1 |
| chr10:97923517-97924017:downstream_17839 | chr10 | 97889471 | 97923517 | + | ZNF518A | NM | 15 | 0,2 | 1 | 0 | 8,94E-01 | NA | 17 | 3 | 24 | 27 | 6 | 7 | 19 | 16 | -1,3 | -1,5 | 0,1 |
| chr4:15733796-15734296:downstream_3370 | chr4 | 15704572 | 15733796 | + | BST1 | NM | 14 | 0,2 | 1 | 0 | 8,94E-01 | NA | 18 | 1 | 7 | 8 | 22 | 19 | 14 | 23 | 0,3 | 0,2 | 0,2 |
| chr12:54679030-54679530:downstream_18596 | chr12 | 54674487 | 54679030 | + | HNRNPA1 | NM | 104 | 0,1 | 1 | 0 | 8,94E-01 | 9,79E-01 | 148 | 127 | 198 | 202 | 41 | 20 | 45 | 51 | -0,5 | -0,6 | 0,1 |
| chr18:70820792-70821292:downstream_13529 | chr18 | 70821292 | 70931733 | - | LOC400655 | NR | 8 | -0,3 | 2 | 0 | 8,95E-01 | NA | 2 | 0 | 19 | 13 | 2 | 0 | 11 | 16 | -4,3 | -4,0 | -0,3 |
| chr1:82458422-82458922:downstream_16449 | chr1 | 82165454 | 82458422 | + | ADGRL2 | NM | 4 | -0,4 | 3 | 0 | 8,95E-01 | NA | 0 | 0 | 10 | 10 | 0 | 0 | 7 | 8 | -Inf | -Inf | NaN |
| chr12:106740792-106741292:downstream_18867 | chr12 | 106696568 | 106740792 | + | TCP11L2 | NM | 54 | 0,1 | 1 | 0 | 8,95E-01 | 9,79E-01 | 3 | 10 | 49 | 62 | 17 | 13 | 157 | 126 | -3,1 | -3,3 | 0,2 |
| chr18:33171770-33172270:downstream_13608 | chr18 | 33171700 | 33171770 | + | MIR3975 | NR | 32 | 0,1 | 1 | 0 | 8,95E-01 | 9,79E-01 | 52 | 36 | 6 | 14 | 50 | 66 | 13 | 16 | 2,2 | 2,0 | 0,2 |
| chr15:91506355-91506855:downstream_19695 | chr15 | 91498105 | 91506355 | + | RCCD1 | NM | 7 | -0,2 | 2 | 0 | 8,95E-01 | NA | 11 | 5 | 4 | 4 | 9 | 11 | 7 | 2 | 0,9 | 1,1 | -0,2 |
| chr3:101042532-101043032:downstream_23412 | chr3 | 101043032 | 101232085 | - | SENP7 | NM | 3 | 0,4 | 3 | 0 | 8,95E-01 | NA | 0 | 0 | 8 | 3 | 0 | 0 | 4 | 11 | -Inf | -Inf | NaN |
| chr15:68588203-68588703:downstream_11782 | chr15 | 68570140 | 68588203 | + | FEM1B | NM | 15 | 0,2 | 1 | 0 | 8,95E-01 | NA | 7 | 2 | 4 | 2 | 18 | 43 | 18 | 25 | 0,5 | 0,5 | 0,0 |
| chr3:169484210-169484710:downstream_3268 | chr3 | 169484710 | 169487683 | - | ACTRT3 | NM | 7 | -0,2 | 2 | 0 | 8,95E-01 | NA | 7 | 8 | 10 | 4 | 12 | 4 | 10 | 3 | 0,1 | 0,3 | -0,2 |
| chr2:71336305-71336805:downstream_2025 | chr2 | 71336805 | 71357394 | - | MCEE | NM | 9 | 0,2 | 2 | 0 | 8,96E-01 | NA | 10 | 1 | 3 | 5 | 13 | 15 | 7 | 16 | 0,5 | 0,3 | 0,2 |
| chr15:49416970-49417470:downstream_19516 | chr15 | 49417470 | 49447854 | - | COPS2 | NM | 21 | 0,1 | 1 | 0 | 8,96E-01 | NA | 20 | 58 | 17 | 19 | 21 | 13 | 11 | 7 | 1,1 | 0,9 | 0,2 |
| chr2:62451866-62452366:downstream_1987 | chr2 | 62423261 | 62451866 | + | B3GNT2 | NM | 31 | 0,1 | 1 | 0 | 8,96E-01 | 9,80E-01 | 2 | 3 | 13 | 5 | 35 | 16 | 97 | 80 | -1,7 | -1,8 | 0,1 |
| chr19:40487671-40488171:downstream_21360 | chr19 | 40476911 | 40487671 | + | PSMC4 | NM | 502 | 0,1 | 0 | 0 | 8,97E-01 | 9,80E-01 | 122 | 120 | 60 | 61 | 1063 | 1339 | 694 | 555 | 1,0 | 0,9 | 0,1 |
| chr3:23933131-23933631:downstream_23497 | chr3 | 23847383 | 23933131 | + | UBE2E1 | NM | 20 | -0,1 | 1 | 0 | 8,97E-01 | NA | 2 | 5 | 4 | 8 | 28 | 27 | 39 | 50 | -0,8 | -0,7 | -0,1 |
| chr17:1421390-1421890:downstream_12727 | chr17 | 1420212 | 1421390 | + | PITPNA-AS1 | NR | 16 | 0,2 | 1 | 0 | 8,97E-01 | NA | 8 | 4 | 3 | 6 | 33 | 24 | 25 | 22 | 0,5 | 0,3 | 0,2 |
| chr10:102313681-102314181:downstream_8677 | chr10 | 102295640 | 102313681 | + | HIF1AN | NM | 17 | 0,2 | 1 | 0 | 8,98E-01 | NA | 14 | 0 | 10 | 6 | 25 | 21 | 33 | 23 | -0,2 | -0,3 | 0,1 |
| chr13:76180156-76180656:downstream_19124 | chr13 | 76123615 | 76180156 | + | UCHL3 | NM | 10 | -0,2 | 1 | 0 | 8,98E-01 | NA | 20 | 4 | 13 | 10 | 7 | 10 | 3 | 11 | 0,1 | 0,3 | -0,3 |
| chrX:139854841-139855341:downstream_26880 | chrX | 139791923 | 139854841 | + | LINC00632 | NR | 3 | -0,3 | 2 | 0 | 8,98E-01 | NA | 2 | 0 | 1 | 7 | 2 | 1 | 4 | 6 | -1,8 | -1,5 | -0,3 |
| chr22:31830172-31830672:downstream_15565 | chr22 | 31795538 | 31830172 | + | DRG1 | NM | 35 | -0,1 | 1 | 0 | 8,98E-01 | 9,81E-01 | 16 | 23 | 36 | 13 | 36 | 56 | 47 | 56 | -0,3 | -0,2 | -0,2 |
| chr3:182655445-182655945:downstream_23110 | chr3 | 182655945 | 182703741 | - | DCUN1D1 | NM | 8 | -0,2 | 2 | 0 | 8,99E-01 | NA | 8 | 4 | 8 | 5 | 10 | 9 | 14 | 4 | -0,1 | 0,0 | -0,2 |
| chr1:160964500-160965000:downstream_1169 | chr1 | 160965000 | 160991133 | - | F11R | NM | 9 | 0,2 | 1 | 0 | 8,99E-01 | NA | 8 | 1 | 7 | 7 | 12 | 7 | 17 | 15 | -0,6 | -0,7 | 0,2 |
| chr14:20946165-20946665:downstream_11029 | chr14 | 20937537 | 20946165 | + | PNP | NM | 45 | 0,1 | 1 | 0 | 9,00E-01 | 9,81E-01 | 73 | 63 | 7 | 16 | 76 | 93 | 17 | 15 | 2,6 | 2,4 | 0,2 |
| chr4:128754526-128755026:downstream_3504 | chr4 | 128703452 | 128754526 | + | HSPA4L | NM | 50 | 0,1 | 1 | 0 | 9,00E-01 | 9,81E-01 | 65 | 32 | 42 | 37 | 61 | 59 | 53 | 50 | 0,3 | 0,2 | 0,1 |
| chr3:48509044-48509544:downstream_23488 | chr3 | 48506918 | 48509044 | + | TREX1 | NM | 15 | 0,1 | 1 | 0 | 9,00E-01 | NA | 15 | 26 | 26 | 15 | 9 | 7 | 14 | 4 | 0,0 | -0,2 | 0,2 |
| chr11:118964084-118964584:downstream_9686 | chr11 | 118964584 | 118966177 | - | H2AFX | NM | 22 | 0,1 | 1 | 0 | 9,01E-01 | NA | 2 | 7 | 6 | 7 | 26 | 35 | 40 | 50 | -0,4 | -0,6 | 0,2 |
| chr7:12276890-12277390:downstream_25780 | chr7 | 12250847 | 12276890 | + | TMEM106B | NM | 42 | 0,1 | 1 | 0 | 9,01E-01 | 9,81E-01 | 5 | 6 | 8 | 8 | 51 | 64 | 93 | 99 | -0,6 | -0,7 | 0,1 |
| chr5:140098598-140099098:downstream_4460 | chr5 | 140098510 | 140098598 | + | VTRNA1-2 | NR | 32 | -0,2 | 1 | 0 | 9,02E-01 | 9,81E-01 | 76 | 94 | 1 | 3 | 50 | 26 | 0 | 2 | 5,2 | 5,5 | -0,4 |
| chr4:109588978-109589478:downstream_23783 | chr4 | 109571740 | 109588978 | + | OSTC | NM | 114 | -0,1 | 1 | 0 | 9,02E-01 | 9,81E-01 | 7 | 4 | 6 | 4 | 266 | 241 | 186 | 202 | 0,2 | 0,4 | -0,2 |
| chr2:143799885-143800385:downstream_22016 | chr2 | 143635194 | 143799885 | + | KYNU | NM | 34 | -0,1 | 1 | 0 | 9,02E-01 | 9,81E-01 | 5 | 0 | 7 | 8 | 48 | 20 | 89 | 96 | -1,6 | -1,4 | -0,2 |
| chr9:100778224-100778724:downstream_7220 | chr9 | 100745488 | 100778224 | + | ANP32B | NM | 22 | -0,1 | 1 | 0 | 9,03E-01 | NA | 4 | 11 | 21 | 7 | 24 | 28 | 43 | 43 | -0,9 | -0,7 | -0,2 |
| chrX:114885179-114885679:downstream_26977 | chrX | 114795176 | 114885179 | + | PLS3 | NM | 47 | 0,1 | 1 | 0 | 9,03E-01 | 9,81E-01 | 3 | 4 | 8 | 7 | 45 | 64 | 103 | 139 | -1,1 | -1,2 | 0,1 |
| chr19:15463831-15464331:downstream_13738 | chr19 | 15464331 | 15490612 | - | AKAP8 | NM | 19 | -0,2 | 1 | 0 | 9,03E-01 | NA | 1 | 14 | 4 | 3 | 29 | 61 | 21 | 21 | 0,9 | 1,1 | -0,2 |
| chr11:65336190-65336690:downstream_9443 | chr11 | 65336690 | 65337884 | - | SSSCA1-AS1 | NR | 22 | -0,1 | 1 | 0 | 9,03E-01 | NA | 16 | 55 | 6 | 12 | 32 | 36 | 10 | 7 | 2,0 | 2,0 | -0,1 |
| chr17:54964769-54965269:downstream_13250 | chr17 | 54965269 | 54991409 | - | TRIM25 | NM | 5 | 0,2 | 2 | 0 | 9,04E-01 | NA | 2 | 9 | 3 | 2 | 9 | 7 | 6 | 3 | 1,1 | 0,8 | 0,3 |
| chr3:48787592-48788092:downstream_2945 | chr3 | 48788092 | 48885270 | - | PRKAR2A | NM | 23 | 0,1 | 1 | 0 | 9,04E-01 | NA | 8 | 10 | 29 | 17 | 23 | 8 | 60 | 28 | -1,4 | -1,5 | 0,1 |
| chr5:114856107-114856607:downstream_4369 | chr5 | 114856607 | 114880591 | - | FEM1C | NM | 22 | 0,1 | 1 | 0 | 9,04E-01 | NA | 21 | 21 | 3 | 7 | 33 | 69 | 13 | 13 | 2,2 | 2,0 | 0,2 |
| chr19:44110875-44111375:downstream_14349 | chr19 | 44111375 | 44124014 | - | ZNF428 | NM | 34 | -0,1 | 1 | 0 | 9,04E-01 | 9,81E-01 | 4 | 9 | 3 | 5 | 55 | 105 | 49 | 46 | 0,7 | 0,8 | 0,0 |
| chr6:42854731-42855231:downstream_5143 | chr6 | 42847670 | 42854731 | + | RPL7L1 | NM | 77 | -0,1 | 1 | 0 | 9,04E-01 | 9,81E-01 | 112 | 107 | 79 | 57 | 76 | 89 | 61 | 36 | 0,7 | 0,8 | -0,1 |
| chr20:46285621-46286121:downstream_22479 | chr20 | 46130600 | 46285621 | + | NCOA3 | NM | 44 | 0,2 | 1 | 0 | 9,05E-01 | 9,81E-01 | 2 | 0 | 7 | 7 | 13 | 22 | 156 | 144 | -3,1 | -3,1 | 0,0 |
| chr5:52782304-52782804:downstream_24132 | chr5 | 52776263 | 52782304 | + | FST | NM | 73 | 0,1 | 1 | 0 | 9,05E-01 | 9,81E-01 | 60 | 81 | 60 | 55 | 71 | 106 | 89 | 63 | 0,3 | 0,2 | 0,1 |
| chr1:109471629-109472129:downstream_16602 | chr1 | 109472129 | 109506121 | - | CLCC1 | NM | 16 | 0,1 | 1 | 0 | 9,05E-01 | NA | 8 | 6 | 13 | 13 | 20 | 8 | 26 | 30 | -0,9 | -1,0 | 0,1 |
| chr15:25353499-25353999:downstream_11586 | chr15 | 25353414 | 25353499 | + | SNORD116-30 | NR | 7 | 0,3 | 3 | 0 | 9,06E-01 | NA | 0 | 0 | 8 | 18 | 0 | 0 | 21 | 12 | -Inf | -Inf | NaN |
| chr7:32907277-32907777:downstream_5788 | chr7 | 32907777 | 32931468 | - | KBTBD2 | NM | 22 | -0,1 | 1 | 0 | 9,06E-01 | NA | 4 | 19 | 13 | 20 | 11 | 40 | 38 | 28 | -0,5 | -0,4 | -0,1 |
| chr1:214837914-214838414:downstream_288 | chr1 | 214776531 | 214837914 | + | CENPF | NM | 8 | -0,3 | 2 | 0 | 9,06E-01 | NA | 0 | 0 | 4 | 7 | 0 | 2 | 21 | 30 | -Inf | -4,7 | -Inf |
| chr9:14081341-14081841:downstream_26408 | chr9 | 14081841 | 14398982 | - | NFIB | NM | 16 | 0,2 | 2 | 0 | 9,07E-01 | NA | 0 | 2 | 7 | 8 | 5 | 6 | 40 | 62 | -3,0 | -3,3 | 0,3 |
| chr4:99799106-99799606:downstream_23624 | chr4 | 99799606 | 99851786 | - | EIF4E | NM | 13 | -0,2 | 1 | 0 | 9,07E-01 | NA | 25 | 26 | 18 | 16 | 7 | 6 | 0 | 7 | 0,6 | 0,9 | -0,3 |
| chr14:21676795-21677295:downstream_19254 | chr14 | 21677295 | 21737638 | - | HNRNPC | NM | 87 | 0,1 | 1 | 0 | 9,07E-01 | 9,82E-01 | 99 | 78 | 125 | 85 | 81 | 57 | 83 | 88 | -0,2 | -0,3 | 0,1 |
| chr12:54624230-54624730:downstream_18481 | chr12 | 54624730 | 54673915 | - | CBX5 | NM | 10 | 0,2 | 2 | 0 | 9,08E-01 | NA | 0 | | 3 | 7 | 7 | 6 | 19 | 38 | -2,1 | -2,2 | 0,2 |
| chr19:56114504-56115004:downstream_14669 | chr19 | 56111705 | 56114504 | + | ZNF524 | NM | 5 | 0,2 | 2 | 0 | 9,08E-01 | NA | 4 | 7 | 3 | 3 | 7 | 10 | 3 | 7 | 0,8 | 0,7 | 0,2 |
| chr5:157212250-157212750:downstream_24066 | chr5 | 157212750 | 157286183 | - | CLINT1 | NM | 4 | 0,2 | 2 | 0 | 9,08E-01 | NA | 2 | 3 | 7 | 4 | 2 | 3 | 7 | 6 | -1,0 | -1,2 | 0,2 |
| chr16:18793776-18794276:downstream_20198 | chr16 | 18794276 | 18801656 | - | RPS15A | NM | 26 | -0,1 | 1 | 0 | 9,08E-01 | NA | 18 | 19 | 15 | 4 | 48 | 57 | 29 | 19 | 0,9 | 1,1 | -0,2 |
| chr11:66012218-66012718:downstream_8901 | chr11 | 65837823 | 66012218 | + | PACS1 | NM | 5 | -0,2 | 2 | 0 | 9,09E-01 | NA | 2 | 0 | 6 | 6 | 3 | 3 | 11 | 12 | -2,2 | -1,9 | -0,3 |
| chr3:62280573-62281073:downstream_2549 | chr3 | 61547242 | 62280573 | + | PTPRG | NM | 26 | 0,1 | 1 | 0 | 9,09E-01 | NA | 12 | 8 | 53 | 38 | 13 | 3 | 43 | 36 | -2,2 | -2,3 | 0,1 |
| chr15:93441977-93442477:downstream_19598 | chr15 | 93426072 | 93441977 | + | LINC01578 | NR | 61 | 0,1 | 1 | 0 | 9,09E-01 | 9,83E-01 | 124 | 66 | 88 | 58 | 43 | 44 | 28 | 41 | 0,4 | 0,3 | 0,1 |
| chr16:68334017-68334517:downstream_12433 | chr16 | 68334517 | 68344868 | - | SLC7A6OS | NM | 70 | 0,1 | 1 | 0 | 9,09E-01 | 9,83E-01 | 94 | 112 | 51 | 67 | 94 | 50 | 47 | 41 | 0,8 | 0,7 | 0,1 |
| chr5:140105831-140106331:downstream_4464 | chr5 | 140105743 | 140105831 | + | VTRNA1-3 | NR | 89 | 0,1 | 1 | 0 | 9,10E-01 | 9,83E-01 | 144 | 86 | 19 | 17 | 243 | 138 | 38 | 27 | 2,6 | 2,6 | 0,1 |
| chr5:178507691-178508191:downstream_4622 | chr5 | 178487606 | 178507691 | + | ZNF354C | NM | 10 | -0,2 | 1 | 0 | 9,10E-01 | NA | 8 | 10 | 26 | 8 | 5 | 6 | 11 | 7 | -1,0 | -0,8 | -0,2 |
| chr6:149825130-149825630:downstream_5393 | chr6 | 149825630 | 149867238 | - | PPIL4 | NM | 10 | 0,2 | 1 | 0 | 9,10E-01 | NA | 8 | 11 | 15 | 7 | 5 | 14 | 10 | 13 | -0,2 | -0,3 | 0,1 |
| chr1:249104150-249104650:downstream_1501 | chr1 | 249104650 | 249120154 | - | SH3BP5L | NM | 6 | 0,2 | 2 | 0 | 9,11E-01 | NA | 3 | 6 | 8 | 3 | 9 | 4 | 8 | 10 | -0,4 | -0,5 | 0,1 |
| chr20:30195043-30195543:downstream_14951 | chr20 | 30194988 | 30195043 | + | MIR3193 | NR | 8 | -0,2 | 2 | 0 | 9,11E-01 | NA | 7 | 7 | 17 | 9 | 11 | 0 | 8 | 10 | -1,0 | -0,8 | -0,2 |
| chr13:28009846-28010346:downstream_10677 | chr13 | 27998680 | 28009846 | + | GTF3A | NM | 38 | -0,1 | 1 | 0 | 9,12E-01 | 9,83E-01 | 1 | 16 | 6 | 8 | 58 | 96 | 75 | 43 | 0,3 | 0,4 | 0,0 |
| chr18:65173318-65173818:downstream_13662 | chr18 | 65173818 | 65183967 | - | DSEL | NM | 5 | 0,3 | 3 | 0 | 9,12E-01 | NA | 5 | 6 | 0 | 0 | 3 | 21 | 1 | 0 | Inf | 4,1 | Inf |
| chr7:66805012-66805512:downstream_5875 | chr7 | 66800962 | 66805012 | + | LINC01372 | NR | 6 | -0,2 | 2 | 0 | 9,13E-01 | NA | 7 | 0 | 8 | 12 | 6 | 0 | 8 | 7 | -1,7 | -1,5 | -0,2 |
| chr1:53692063-53692563:downstream_739 | chr1 | 53692563 | 53704282 | - | MAGOH | NM | 13 | 0,1 | 1 | 0 | 9,13E-01 | NA | 8 | 1 | 4 | 6 | 24 | 16 | 32 | 16 | -0,1 | -0,3 | 0,2 |
| chr10:94415152-94415652:downstream_8215 | chr10 | 94352824 | 94415152 | + | KIF11 | NM | 25 | -0,1 | 1 | 0 | 9,14E-01 | NA | 4 | 0 | 13 | 18 | 16 | 6 | 77 | 71 | -2,9 | -2,7 | -0,2 |
| chr19:863610-864110:downstream_13833 | chr19 | 859664 | 863610 | + | CFD | NM | 190 | 0,1 | 1 | 0 | 9,14E-01 | 9,83E-01 | 9 | 8 | 6 | 3 | 489 | 488 | 270 | 247 | 0,9 | 0,9 | 0,0 |
| chr3:10168874-10169374:downstream_2819 | chr3 | 10157332 | 10168874 | + | BRK1 | NM | 19 | -0,1 | 1 | 0 | 9,15E-01 | NA | 4 | 5 | 3 | 2 | 39 | 52 | 25 | 24 | 0,8 | 0,9 | -0,1 |
| chrX:119369808-119370308:downstream_7909 | chrX | 119370308 | 119379122 | - | NKAPP1 | NR | 11 | 0,2 | 1 | 0 | 9,15E-01 | NA | 7 | 10 | 28 | 20 | 0 | 6 | 14 | 4 | -1,5 | -1,6 | 0,2 |
| chr18:32890730-32891230:downstream_20976 | chr18 | 32870235 | 32890730 | + | ZNF271P | NR | 4 | 0,3 | 2 | 0 | 9,15E-01 | NA | 8 | 7 | 3 | 2 | 9 | 0 | 4 | 0 | 1,5 | 1,1 | 0,4 |
| chr4:174298683-174299183:downstream_3905 | chr4 | 174292092 | 174298683 | + | SAP30 | NM | 5 | 0,2 | 2 | 0 | 9,15E-01 | NA | 6 | 3 | 6 | 7 | 4 | 2 | 6 | 5 | -0,6 | -0,8 | 0,2 |
| chr4:773436-773936:downstream_23733 | chr4 | 773936 | 775636 | - | LOC100129917 | NR | 40 | 0,1 | 1 | 0 | 9,15E-01 | 9,83E-01 | 37 | 29 | 58 | 49 | 32 | 20 | 56 | 34 | -0,7 | -0,8 | 0,1 |
| chr3:9430036-9430536:downstream_2537 | chr3 | 9430536 | 9439174 | - | THUMPD3-AS1 | NR | 78 | -0,1 | 1 | 0 | 9,15E-01 | 9,83E-01 | 87 | 107 | 38 | 39 | 107 | 151 | 47 | 50 | 1,3 | 1,4 | -0,1 |
| chr14:55614333-55614833:downstream_19209 | chr14 | 55614833 | 55658396 | - | DLGAP5 | NM | 7 | -0,2 | 2 | 0 | 9,15E-01 | NA | 0 | 1 | 7 | 7 | 2 | 1 | 15 | 21 | -3,9 | -3,8 | -0,1 |
| chr7:86824977-86825477:downstream_5928 | chr7 | 86825477 | 86849031 | - | TMEM243 | NM | 34 | 0,1 | 1 | 0 | 9,15E-01 | 9,83E-01 | 30 | 23 | 26 | 23 | 50 | 37 | 36 | 48 | 0,1 | 0,0 | 0,1 |
| chr2:61414686-61415186:downstream_1985 | chr2 | 61404554 | 61414686 | + | AHSA2 | NM | 55 | -0,1 | 1 | 0 | 9,16E-01 | 9,83E-01 | 101 | 42 | 54 | 38 | 84 | 43 | 31 | 45 | 0,6 | 0,7 | -0,1 |
| chr20:20014504-20015004:downstream_22389 | chr20 | 20015004 | 20036690 | - | CRNKL1 | NM | 55 | 0,1 | 1 | 0 | 9,16E-01 | 9,83E-01 | 73 | 64 | 81 | 91 | 34 | 21 | 46 | 29 | -0,3 | -0,4 | 0,1 |
| chr12:120881263-120881763:downstream_10453 | chr12 | 120881763 | 120884215 | - | TRIAP1 | NM | 276 | -0,1 | 1 | 0 | 9,16E-01 | 9,83E-01 | 47 | 40 | 13 | 5 | 863 | 929 | 168 | 148 | 2,3 | 2,5 | -0,2 |
| chr20:44526758-44527258:downstream_22510 | chr20 | 44527258 | 44541003 | - | PLTP | NM | 6 | 0,2 | 2 | 0 | 9,16E-01 | NA | 3 | 9 | 4 | 4 | 4 | 10 | 10 | 2 | 0,5 | 0,3 | 0,3 |
| chr3:63819045-63819545:downstream_23466 | chr3 | 63819545 | 63849599 | - | THOC7 | NM | 42 | -0,1 | 1 | 0 | 9,16E-01 | 9,83E-01 | 11 | 0 | 11 | 5 | 76 | 58 | 81 | 93 | -0,6 | -0,4 | -0,2 |
| chr12:49578077-49578577:downstream_18895 | chr12 | 49578577 | 49583107 | - | TUBA1A | NM | 136 | -0,1 | 1 | 0 | 9,16E-01 | 9,83E-01 | 2 | 10 | 8 | 2 | 290 | 290 | 253 | 235 | 0,1 | 0,3 | -0,2 |
| chr3:53849823-53850323:downstream_2631 | chr3 | 53850323 | 53880420 | - | CHDH | NM | 6 | 0,2 | 2 | 0 | 9,17E-01 | NA | 1 | 4 | 6 | 7 | 2 | 7 | 8 | 16 | -1,4 | -1,6 | 0,2 |
| chr14:32544124-32544624:downstream_11100 | chr14 | 32544624 | 32545905 | - | ARHGAP5-AS1 | NR | 16 | 0,1 | 1 | 0 | 9,17E-01 | NA | 8 | 3 | 11 | 3 | 21 | 23 | 33 | 25 | -0,4 | -0,4 | 0,0 |
| chr1:85018303-85018803:downstream_803 | chr1 | 85018803 | 85040163 | - | CTBS | NM | 7 | 0,2 | 2 | 0 | 9,17E-01 | NA | 7 | 2 | 8 | 5 | 7 | 7 | 11 | 11 | -0,5 | -0,6 | 0,1 |
| chr6:26055467-26055967:downstream_4903 | chr6 | 26055967 | 26056699 | - | HIST1H1C | NM | 513 | 0,1 | 1 | 0 | 9,17E-01 | 9,83E-01 | 15 | 7 | 39 | 33 | 557 | 351 | 1706 | 1398 | -1,7 | -1,8 | 0,0 |
| chr9:98868442-98868942:downstream_7205 | chr9 | 98868942 | 98878693 | - | LOC158434 | NM | 6 | -0,2 | 2 | 0 | 9,17E-01 | NA | 2 | 4 | 8 | 7 | 2 | 5 | 13 | 4 | -1,5 | -1,4 | -0,2 |

| | | | | | | | | | | | | | | | | | | | | | | | |
|---|---|---|---|---|---|---|---|---|---|---|---|---|---|---|---|---|---|---|---|---|---|---|---|
| chr16:71315702-71316202:downstream_19944 | chr16 | 71316202 | 71323509 | - | CMTR2 | NM | 21 | 0,1 | 1 | 0 | 9,18E-01 | NA | 8 | 25 | 18 | 6 | 37 | 27 | 19 | 28 | 0,5 | 0,4 | 0,1 |
| chr6:7612200-7612700:downstream_4688 | chr6 | 7590431 | 7612200 | + | SNRNP48 | NM | 14 | -0,1 | 1 | 0 | 9,19E-01 | NA | 8 | 13 | 7 | 13 | 14 | 22 | 19 | 14 | 0,0 | 0,1 | -0,1 |
| chr12:76438662-76439162:downstream_18699 | chr12 | 76439162 | 76478813 | - | NAP1L1 | NM | 31 | 0,1 | 1 | 0 | 9,19E-01 | 9,84E-01 | 18 | 30 | 18 | 21 | 54 | 34 | 33 | 43 | 0,3 | 0,2 | 0,1 |
| chr9:35737005-35737505:downstream_7061 | chr9 | 35732316 | 35737005 | + | CREB3 | NM | 31 | -0,1 | 1 | 0 | 9,19E-01 | 9,84E-01 | 26 | 16 | 31 | 21 | 38 | 35 | 38 | 45 | -0,3 | -0,2 | -0,1 |
| chr12:54573641-54574141:downstream_18844 | chr12 | 54574141 | 54582778 | - | SMUG1 | NM | 16 | 0,1 | 1 | 0 | 9,19E-01 | NA | 19 | 27 | 15 | 21 | 15 | 10 | 17 | 6 | 0,4 | 0,2 | 0,2 |
| chr5:125877032-125877532:downstream_24011 | chr5 | 125877532 | 125931082 | - | ALDH7A1 | NM | 6 | -0,2 | 2 | 0 | 9,20E-01 | NA | 6 | 0 | 10 | 6 | 5 | [illegible] | 11 | 5 | -1,4 | -1,2 | -0,2 |
| chr5:80172634-80173134:downstream_4042 | chr5 | 79950466 | 80172634 | + | MSH3 | NM | 11 | 0,1 | 1 | 0 | 9,20E-01 | NA | 3 | 6 | 8 | 17 | 7 | 6 | 22 | 17 | -1,5 | -1,7 | 0,2 |
| chr11:116648775-116649275:downstream_8949 | chr11 | 116649275 | 116658739 | - | ZPR1 | NM | 80 | -0,1 | 1 | 0 | 9,20E-01 | 9,85E-01 | 57 | 28 | 17 | 16 | 192 | 191 | 81 | 57 | 1,4 | 1,5 | -0,1 |
| chr3:49199467-49199967:downstream_2951 | chr3 | 49199967 | 49203785 | - | CCDC71 | NM | 6 | 0,2 | 2 | 0 | 9,21E-01 | NA | 11 | 0 | 4 | 0 | 12 | 12 | 8 | 2 | 1,3 | 1,3 | 0,1 |
| chr1:91380356-91380856:downstream_17433 | chr1 | 91380856 | 91487812 | - | ZNF644 | NM | 8 | 0,2 | 2 | 0 | 9,21E-01 | NA | 3 | 1 | 3 | 10 | 5 | 7 | 15 | 24 | -1,6 | -1,7 | 0,2 |
| chr12:119632551-119633051:downstream_10441 | chr12 | 119616594 | 119632551 | + | HSPB8 | NM | 100 | -0,1 | 1 | 0 | 9,21E-01 | 9,85E-01 | 279 | 162 | 178 | 114 | 25 | 15 | 13 | 6 | 0,6 | 0,7 | -0,1 |
| chr1:40537881-40538381:downstream_17109 | chr1 | 40538381 | 40563142 | - | PPT1 | NM | 85 | 0,1 | 1 | 0 | 9,22E-01 | 9,85E-01 | 6 | 15 | 6 | 10 | 207 | 148 | 131 | 159 | 0,5 | 0,3 | 0,2 |
| chr3:47891686-47892186:downstream_23113 | chr3 | 47844398 | 47891686 | + | DHX30 | NM | 6 | 0,2 | 2 | 0 | 9,22E-01 | NA | 6 | 11 | 7 | 2 | 8 | 4 | 6 | 2 | 0,9 | 0,7 | 0,2 |
| chr17:7362184-7362684:downstream_20814 | chr17 | 7362684 | 7387568 | - | ZBTB4 | NM | 36 | 0,1 | 1 | 0 | 9,23E-01 | 9,85E-01 | 5 | 4 | 6 | 6 | 41 | 72 | 85 | 71 | -0,4 | -0,5 | 0,1 |
| chr3:10058736-10059236:downstream_2813 | chr3 | 10059236 | 10067820 | - | CIDECP | NR | 13 | 0,1 | 1 | 0 | 9,23E-01 | NA | 17 | 15 | 1 | 7 | 17 | 29 | 1 | 12 | 1,9 | 1,8 | 0,1 |
| chr1:168212088-168212588:downstream_1205 | chr1 | 168195254 | 168212088 | + | SFT2D2 | NM | 140 | 0,0 | 1 | 0 | 9,23E-01 | 9,85E-01 | 109 | 114 | 180 | 209 | 93 | 88 | 139 | 186 | -0,8 | -0,8 | 0,0 |
| chr15:52204331-52204831:downstream_11469 | chr15 | 52121824 | 52204331 | + | TMOD3 | NM | 83 | -0,1 | 1 | 0 | 9,24E-01 | 9,85E-01 | 2 | 3 | 10 | 11 | 72 | 47 | 306 | 213 | -2,2 | -2,1 | 0,0 |
| chr15:59170743-59171243:downstream_19726 | chr15 | 59171243 | 59225852 | - | SLTM | NM | 14 | 0,1 | 1 | 0 | 9,24E-01 | NA | 3 | 3 | 6 | 6 | 15 | 17 | 26 | 37 | -0,9 | -1,0 | 0,1 |
| chr1:151556059-151556559:downstream_17374 | chr1 | 151512780 | 151556059 | + | TUFT1 | NM | 7 | 0,2 | 2 | 0 | 9,24E-01 | NA | 1 | 14 | 0 | 5 | 10 | 15 | 4 | 7 | 1,5 | 1,2 | 0,3 |
| chr15:25228937-25229437:downstream_11562 | chr15 | 25227140 | 25228937 | + | PWARSN | NR | 49 | -0,1 | 1 | 0 | 9,25E-01 | 9,85E-01 | 35 | 51 | 57 | 43 | 58 | 40 | 51 | 57 | -0,2 | -0,2 | -0,1 |
| chr8:145747260-145747760:downstream_6254 | chr8 | 145747760 | 145752416 | - | LRRC24 | NM | 8 | 0,1 | 2 | 0 | 9,25E-01 | NA | 9 | 16 | 14 | 6 | 5 | 4 | 4 | 3 | 0,4 | 0,2 | 0,1 |
| chr19:36260077-36260577:downstream_14231 | chr19 | 36249043 | 36260077 | + | PROSER3 | NM | 10 | -0,1 | 1 | 0 | 9,25E-01 | NA | 15 | 9 | 6 | 16 | 9 | 8 | 8 | 7 | 0,1 | 0,2 | -0,1 |
| chr3:49060926-49061426:downstream_23301 | chr3 | 49057907 | 49060926 | + | NDUFAF3 | NM | 25 | -0,1 | 1 | 0 | 9,25E-01 | NA | 31 | 16 | 10 | 7 | 48 | 57 | 19 | 14 | 1,5 | 1,7 | -0,1 |
| chr6:28270326-28270826:downstream_25053 | chr6 | 28249313 | 28270326 | + | PGBD1 | NM | 6 | 0,1 | 2 | 0 | 9,25E-01 | NA | 6 | 7 | 4 | 4 | 10 | 7 | 6 | 7 | 0,6 | 0,4 | 0,1 |
| chr6:88223595-88224095:downstream_5257 | chr6 | 88224095 | 88299735 | - | RARS2 | NM | 9 | -0,1 | 1 | 0 | 9,26E-01 | NA | 9 | 10 | 7 | 16 | 3 | 13 | 6 | 12 | -0,3 | -0,2 | -0,2 |
| chr9:115448290-115448790:downstream_6921 | chr9 | 115448790 | 115480387 | - | INIP | NM | 16 | 0,1 | 1 | 0 | 9,27E-01 | NA | 14 | 21 | 32 | 31 | 7 | 3 | 13 | 7 | -0,8 | -1,0 | 0,2 |
| chr17:8090322-8090822:downstream_12822 | chr17 | 8090262 | 8090322 | + | MIR4521 | NR | 32 | 0,1 | 1 | 0 | 9,27E-01 | 9,86E-01 | 36 | 44 | 7 | 6 | 73 | 68 | 11 | 12 | 2,6 | 2,6 | 0,1 |
| chr8:74202373-74202873:downstream_6601 | chr8 | 74202873 | 74205869 | - | RPL7 | NM | 66 | -0,1 | 1 | 0 | 9,27E-01 | 9,86E-01 | 155 | 138 | 40 | 47 | 72 | 44 | 22 | 12 | 1,7 | 1,8 | 0,0 |
| chr16:53524451-53524951:downstream_19869 | chr16 | 53524951 | 53537216 | - | AKTIP | NM | 10 | 0,1 | 1 | 0 | 9,27E-01 | NA | 7 | 7 | 18 | 14 | 3 | 8 | 13 | 14 | -1,2 | -1,3 | 0,1 |
| chr7:32769595-32770095:downstream_5535 | chr7 | 32767561 | 32769595 | + | ZNRF2P1 | NR | 8 | -0,1 | 2 | 0 | 9,27E-01 | NA | 11 | 2 | 11 | 6 | 7 | 7 | 8 | 7 | -0,3 | -0,2 | -0,2 |
| chr12:120634002-120634502:downstream_18809 | chr12 | 120634502 | 120639014 | - | RPLP0 | NM | 20 | -0,1 | 1 | 0 | 9,27E-01 | NA | 7 | 7 | 7 | 3 | 46 | 37 | 19 | 33 | 0,5 | 0,7 | -0,2 |
| chr7:27139877-27140377:downstream_25561 | chr7 | 27135712 | 27139877 | + | HOTAIRM1 | NR | 18 | 0,1 | 1 | 0 | 9,28E-01 | NA | 28 | 0 | 14 | 11 | 25 | 23 | 21 | 24 | 0,2 | 0,1 | 0,1 |
| chr12:108106257-108106757:downstream_10401 | chr12 | 108079589 | 108106257 | + | PWP1 | NM | 50 | 0,1 | 1 | 0 | 9,28E-01 | 9,86E-01 | 31 | 24 | 38 | 42 | 59 | 45 | 65 | 92 | -0,5 | -0,6 | 0,1 |
| chr15:65409216-65409716:downstream_11766 | chr15 | 65409716 | 65426174 | - | PDCD7 | NM | 13 | 0,1 | 1 | 0 | 9,29E-01 | NA | 6 | 15 | 13 | 7 | 12 | 17 | 17 | 13 | 0,1 | 0,0 | 0,1 |
| chrX:70278596-70279096:downstream_27031 | chrX | 70279096 | 70293276 | - | SNX12 | NM | 47 | 0,1 | 1 | 0 | 9,29E-01 | 9,86E-01 | 21 | 22 | 35 | 25 | 57 | 54 | 67 | 92 | -0,5 | -0,5 | 0,0 |
| chr2:223808119-223808619:downstream_21718 | chr2 | 223725731 | 223808119 | + | ACSL3 | NM | 295 | 0,0 | 1 | 0 | 9,29E-01 | 9,86E-01 | 107 | 90 | 433 | 332 | 149 | 130 | 674 | 445 | -2,0 | -2,0 | 0,0 |
| chr22:26887904-26888404:downstream_15539 | chr22 | 26879849 | 26887904 | + | SRRD | NM | 7 | -0,1 | 1 | 0 | 9,29E-01 | NA | 7 | 6 | 4 | 7 | 7 | 10 | 8 | 6 | 0,2 | 0,3 | -0,1 |
| chr17:58254190-58254690:downstream_12668 | chr17 | 58254690 | 58469586 | - | USP32 | NM | 7 | 0,2 | 3 | 0 | 9,29E-01 | NA | 0 | 0 | 7 | 6 | 1 | 0 | 21 | 19 | -Inf | -5,6 | -Inf |
| chr3:133875477-133875977:downstream_23400 | chr3 | 133875977 | 133969586 | - | RYK | NM | 27 | 0,1 | 1 | 0 | 9,29E-01 | NA | 15 | 0 | 29 | 25 | 16 | 15 | 70 | 48 | -1,8 | -1,9 | 0,1 |
| chr5:137523404-137523904:downstream_4442 | chr5 | 137514416 | 137523404 | + | KIF20A | NM | 54 | 0,1 | 1 | 0 | 9,30E-01 | 9,86E-01 | 9 | 10 | 11 | 18 | 71 | 72 | 122 | 118 | -0,6 | -0,8 | 0,1 |
| chr1:26803133-26803633:downstream_574 | chr1 | 26798901 | 26803133 | + | HMGN2 | NM | 7 | 0,1 | 2 | 0 | 9,30E-01 | NA | 7 | 9 | 8 | 7 | 3 | 3 | [illegible] | 8 | -0,1 | -0,2 | 0,1 |
| chr19:58133636-58134136:downstream_14701 | chr19 | 58125829 | 58133636 | + | ZNF134 | NM | 45 | -0,1 | 1 | 0 | 9,30E-01 | 9,86E-01 | 70 | 60 | 42 | 38 | 43 | 52 | 36 | 21 | 0,7 | 0,7 | 0,0 |
| chr10:6627323-6627823:downstream_17728 | chr10 | 6622386 | 6627323 | + | PRKCQ-AS1 | NR | 15 | 0,1 | 1 | 0 | 9,30E-01 | NA | 23 | 12 | 8 | 2 | 30 | 25 | 3 | 14 | 1,7 | 1,7 | -0,1 |
| chr19:16756458-16756958:downstream_21413 | chr19 | 16756958 | 16770968 | - | SMIM7 | NM | 27 | 0,1 | 1 | 0 | 9,31E-01 | NA | 10 | 8 | 13 | 16 | 39 | 23 | 45 | 65 | -0,7 | -0,8 | 0,1 |
| chr19:40946247-40946747:downstream_21395 | chr19 | 40946747 | 40950282 | - | SERTAD3 | NM | 31 | 0,1 | 1 | 0 | 9,33E-01 | 9,87E-01 | 8 | 3 | 11 | 8 | 34 | 45 | 75 | 66 | -0,8 | -0,8 | 0,0 |
| chr7:94988683-94989183:downstream_5951 | chr7 | 94989183 | 95025687 | - | PON3 | NM | 96 | -0,1 | 1 | 0 | 9,33E-01 | 9,87E-01 | 21 | 20 | 8 | 25 | 246 | 134 | 161 | 151 | 0,3 | 0,3 | 0,0 |
| chr14:75304013-75304513:downstream_10964 | chr14 | 75230068 | 75304013 | + | YLPM1 | NM | 9 | 0,1 | 2 | 0 | 9,33E-01 | NA | 7 | 1 | 19 | 16 | 5 | 0 | 10 | 13 | -2,1 | -2,2 | 0,1 |
| chr15:75136696-75137196:downstream_11806 | chr15 | 75137196 | 75165670 | - | SCAMP2 | NM | 14 | -0,1 | 1 | 0 | 9,33E-01 | NA | 9 | 3 | 7 | 14 | 27 | 2 | 24 | 24 | -0,8 | -0,7 | -0,1 |
| chr15:34396591-34397091:downstream_11641 | chr15 | 34394273 | 34396591 | + | PGBD4 | NM | 35 | -0,1 | 1 | 0 | 9,33E-01 | 9,87E-01 | 28 | 39 | 14 | 20 | 61 | 61 | 25 | 34 | 1,0 | 1,0 | -0,1 |
| chr14:104003410-104003910:downstream_11381 | chr14 | 103995508 | 104003410 | + | TRMT61A | NM | 16 | -0,1 | 1 | 0 | 9,33E-01 | NA | 18 | 39 | 4 | 4 | 28 | 27 | 6 | 2 | 2,8 | 2,8 | 0,0 |
| chr2:192575195-192575695:downstream_2358 | chr2 | 192554906 | 192575195 | + | LOC105747689 | NR | 6 | -0,2 | 2 | 0 | 9,34E-01 | NA | 7 | 15 | 0 | 1 | 16 | 9 | 0 | 1 | 4,7 | 4,9 | -0,2 |
| chr11:72547289-72547789:downstream_8916 | chr11 | 72547789 | 72853143 | - | FCHSD2 | NM | 5 | 0,2 | 2 | 0 | 9,34E-01 | NA | 1 | 0 | 8 | 4 | 2 | 0 | 13 | 15 | -3,9 | -4,1 | 0,1 |
| chr3:99897476-99897976:downstream_23087 | chr3 | 99536677 | 99897476 | + | CMSS1 | NM | 17 | -0,1 | 1 | 0 | 9,35E-01 | NA | 3 | 9 | 7 | 7 | 17 | 33 | 17 | 39 | -0,3 | -0,2 | -0,1 |
| chr14:101418269-101418769:downstream_11331 | chr14 | 101418192 | 101418269 | + | SNORD114-2 | NR | 5 | -0,2 | 2 | 0 | 9,36E-01 | NA | 5 | 2 | 0 | 10 | 7 | 2 | 1 | 11 | -0,5 | -0,4 | -0,1 |
| chr5:60453035-60453535:downstream_4267 | chr5 | 60453535 | 60458302 | - | SMIM15 | NM | 17 | 0,1 | 1 | 0 | 9,36E-01 | NA | 16 | 14 | 7 | 7 | 39 | 25 | 26 | 6 | 1,1 | 1,0 | 0,1 |
| chr6:76005082-76005582:downstream_24768 | chr6 | 76005582 | 76203545 | - | FILIP1 | NM | 10 | -0,1 | 2 | 0 | 9,36E-01 | NA | 7 | 2 | 28 | 22 | 2 | 1 | 13 | 5 | -2,4 | -2,4 | -0,1 |
| chr19:39220331-39220831:downstream_14267 | chr19 | 39220831 | 39235114 | - | CAPN12 | NM | 38 | 0,1 | 1 | 0 | 9,37E-01 | 9,89E-01 | 25 | 62 | 49 | 40 | 21 | 41 | 35 | 31 | 0,0 | -0,1 | 0,1 |
| chr19:44830205-44830705:downstream_21602 | chr19 | 44830705 | 44860856 | - | ZNF112 | NM | 11 | -0,1 | 1 | 0 | 9,37E-01 | NA | 17 | 16 | 13 | 14 | 13 | 5 | 10 | 4 | 0,3 | 0,4 | 0,0 |
| chr15:25330624-25331124:downstream_11575 | chr15 | 25330530 | 25330624 | + | SNORD116-18 | NR | 430 | 0,0 | 0 | 0 | 9,37E-01 | 9,89E-01 | 1016 | 874 | 501 | 524 | 160 | 176 | 93 | 94 | 0,9 | 0,8 | 0,0 |
| chr20:13147411-13147911:downstream_14767 | chr20 | 12989626 | 13147411 | + | SPTLC3 | NM | 5 | -0,2 | 3 | 0 | 9,38E-01 | NA | 0 | 0 | 4 | 7 | 1 | 0 | 14 | 11 | -Inf | -5,0 | -Inf |
| chrX:48885737-48886237:downstream_27075 | chrX | 48886237 | 48901043 | - | TFE3 | NM | 31 | -0,1 | 1 | 0 | 9,39E-01 | 9,89E-01 | 8 | 0 | 3 | 7 | 60 | 48 | 67 | 52 | -0,2 | -0,1 | -0,1 |
| chr19:23835207-23835707:downstream_13754 | chr19 | 23835707 | 23870017 | - | ZNF675 | NM | 16 | 0,1 | 1 | 0 | 9,39E-01 | NA | 7 | 3 | 8 | 6 | 20 | 22 | 31 | 34 | -0,6 | -0,6 | 0,0 |
| chr1:45244412-45244912:downstream_698 | chr1 | 45241245 | 45244412 | + | RPS8 | NM | 28 | 0,1 | 1 | 0 | 9,39E-01 | NA | 30 | 60 | 25 | 12 | 45 | 22 | 14 | 15 | 1,3 | 1,2 | 0,0 |
| chr2:203168384-203168884:downstream_1776 | chr2 | 203130514 | 203168384 | + | NOP58 | NM | 27 | 0,1 | 1 | 0 | 9,39E-01 | NA | 17 | 10 | 8 | 11 | 62 | 34 | 40 | 35 | 0,4 | 0,3 | 0,1 |
| chr9:99578772-99579272:downstream_6820 | chr9 | 99579272 | 99616389 | - | ZNF782 | NM | 11 | 0,1 | 2 | 0 | 9,40E-01 | NA | 8 | 0 | 17 | 11 | 11 | 1 | 22 | 21 | -1,8 | -1,9 | 0,1 |
| chr9:139687769-139688269:downstream_7413 | chr9 | 139685776 | 139687769 | + | TMEM141 | NM | 16 | -0,1 | 1 | 0 | 9,40E-01 | NA | 14 | 1 | 13 | 4 | 27 | 21 | 26 | 21 | -0,2 | 0,0 | -0,2 |
| chr17:76169009-76169509:downstream_13397 | chr17 | 76164670 | 76169009 | + | SYNGR2 | NM | 138 | 0,1 | 1 | 0 | 9,40E-01 | 9,89E-01 | 6 | 6 | 10 | 7 | 231 | 197 | 385 | 265 | -0,6 | -0,6 | 0,0 |
| chr6:72117066-72117566:downstream_24841 | chr6 | 72117566 | 72130448 | - | LINC00472 | NR | 64 | -0,1 | 1 | 0 | 9,40E-01 | 9,89E-01 | 44 | 28 | 156 | 154 | 10 | 15 | 63 | 42 | -2,1 | -2,0 | -0,1 |
| chr1:113667824-113668324:downstream_195 | chr1 | 113615791 | 113667824 | + | LRIG2 | NM | 16 | 0,1 | 1 | 0 | 9,40E-01 | NA | 14 | 15 | 29 | 25 | 10 | 6 | 14 | 17 | -0,9 | -1,0 | 0,1 |
| chr14:96710666-96711166:downstream_11298 | chr14 | 96671134 | 96710666 | + | BDKRB2 | NM | 17 | 0,1 | 1 | 0 | 9,41E-01 | NA | 2 | 0 | 13 | 11 | 8 | 2 | 43 | 55 | -3,2 | -3,3 | 0,0 |
| chr7:44809479-44809979:downstream_25819 | chr7 | 44788164 | 44809479 | + | ZMIZ2 | NM | 13 | 0,1 | 1 | 0 | 9,41E-01 | NA | 26 | 16 | 11 | 10 | 12 | 13 | 8 | 5 | 1,0 | 0,9 | 0,1 |
| chr20:43880379-43880879:downstream_15039 | chr20 | 43880879 | 43883205 | - | SLPI | NM | 28 | -0,1 | 1 | 0 | 9,41E-01 | 9,89E-01 | 4 | 5 | 1 | 2 | 83 | 66 | 31 | 34 | 1,2 | 1,2 | 0,0 |
| chr1:245013101-245013601:downstream_16804 | chr1 | 245013601 | 245027827 | - | HNRNPU | NM | 86 | 0,0 | 1 | 0 | 9,41E-01 | 9,89E-01 | 108 | 57 | 120 | 84 | 63 | 80 | 86 | 94 | -0,3 | -0,3 | 0,0 |
| chr20:18776709-18777209:downstream_14902 | chr20 | 18774692 | 18776709 | + | LOC100270804 | NR | 6 | -0,1 | 2 | 0 | 9,41E-01 | NA | 6 | 10 | 4 | 3 | 11 | 9 | 8 | 1 | 1,0 | 1,1 | 0,0 |
| chr1:193065094-193065594:downstream_16772 | chr1 | 193065594 | 193075244 | - | GLRX2 | NM | 40 | -0,1 | 1 | 0 | 9,41E-01 | 9,89E-01 | 34 | 25 | 33 | 25 | 36 | 72 | 40 | 58 | 0,0 | 0,1 | -0,1 |
| chr11:14899055-14899555:downstream_9154 | chr11 | 14899555 | 14913751 | - | CYP2R1 | NM | 13 | 0,1 | 1 | 0 | 9,41E-01 | NA | 11 | 4 | 11 | 4 | 15 | 20 | 21 | 17 | -0,1 | -0,1 | 0,0 |
| chr7:27869386-27869886:downstream_25768 | chr7 | 27778991 | 27869386 | + | TAX1BP1 | NM | 28 | -0,1 | 1 | 0 | 9,41E-01 | 9,89E-01 | 9 | 7 | 10 | 13 | 37 | 42 | 46 | 63 | -0,5 | -0,5 | -0,1 |
| chr11:125492654-125493154:downstream_18320 | chr11 | 125462689 | 125492654 | + | STT3A | NM | 6 | 0,1 | 2 | 0 | 9,42E-01 | NA | 7 | 7 | 7 | 7 | 2 | [illegible] | 6 | 13 | -1,1 | -1,1 | 0,1 |
| chr12:58141009-58141509:downstream_10283 | chr12 | 58141509 | 58146230 | - | CDK4 | NM | 27 | -0,1 | 1 | 0 | 9,42E-01 | NA | 62 | 73 | 25 | 22 | 7 | 19 | 6 | 3 | 1,5 | 1,5 | 0,0 |
| chr2:190634492-190634992:downstream_22131 | chr2 | 190634992 | 190649097 | - | ORMDL1 | NM | 13 | -0,1 | 1 | 0 | 9,43E-01 | NA | 18 | 5 | 3 | 6 | 30 | 25 | 8 | 11 | 1,4 | 1,5 | -0,1 |
| chr14:75479966-75480466:downstream_19311 | chr14 | 75480466 | 75518235 | - | MLH3 | NM | 13 | 0,1 | 1 | 0 | 9,43E-01 | NA | 12 | 2 | 14 | 10 | 13 | 12 | 15 | 27 | -0,7 | -0,8 | 0,0 |
| chr11:27515464-27515964:downstream_8861 | chr11 | 27515964 | 27528326 | - | LIN7C | NM | 21 | 0,1 | 1 | 0 | 9,43E-01 | NA | 8 | 13 | 1 | 10 | 50 | 31 | 32 | 20 | 0,9 | 0,7 | 0,2 |
| chr10:93790080-93790580:downstream_8319 | chr10 | 93683735 | 93790080 | + | BTAF1 | NM | 31 | 0,1 | 1 | 0 | 9,44E-01 | 9,91E-01 | 9 | 6 | 32 | 19 | 8 | 33 | 71 | 73 | -1,8 | -1,8 | 0,0 |
| chr12:125430869-125431369:downstream_9950 | chr12 | 125431369 | 125473667 | - | DHX37 | NM | 10 | -0,1 | 1 | 0 | 9,45E-01 | NA | 2 | 11 | 3 | 4 | 14 | 28 | 10 | 11 | 0,9 | 1,0 | -0,1 |
| chr1:59119910-59120410:downstream_759 | chr1 | 59120410 | 59165747 | - | MYSM1 | NM | 7 | 0,1 | 2 | 0 | 9,45E-01 | NA | 9 | 11 | 4 | 6 | 8 | 7 | 4 | 4 | 1,0 | 0,8 | 0,1 |
| chr19:37212238-37212738:downstream_21676 | chr19 | 37178513 | 37212238 | + | ZNF567 | NM | 14 | -0,1 | 2 | 0 | 9,45E-01 | NA | 9 | 0 | 3 | 2 | 18 | 51 | 14 | 16 | 1,0 | 1,2 | -0,2 |
| chr13:45907105-45907605:downstream_19115 | chr13 | 45907605 | 45915419 | - | TPT1 | NM | 48 | -0,1 | 1 | 0 | 9,45E-01 | 9,92E-01 | 28 | 17 | 50 | 55 | 50 | 22 | 67 | 94 | -1,2 | -1,2 | -0,1 |
| chr10:112773425-112773925:downstream_17767 | chr10 | 112679300 | 112773425 | + | SHOC2 | NM | 20 | 0,1 | 1 | 0 | 9,45E-01 | NA | 4 | 4 | 6 | 19 | 16 | 13 | 53 | 45 | -1,6 | -1,8 | 0,1 |
| chr4:84382876-84383376:downstream_23761 | chr4 | 84377084 | 84382876 | + | MRPS18C | NM | 12 | 0,1 | 1 | 0 | 9,46E-01 | NA | 17 | 3 | 10 | 9 | 20 | 8 | 14 | 13 | 0,1 | 0,0 | 0,1 |
| chr10:14913740-14914240:downstream_17602 | chr10 | 14880158 | 14913740 | + | HSPA14 | NM | 12 | 0,1 | 1 | 0 | 9,46E-01 | NA | 7 | 18 | 4 | 2 | 26 | 24 | 8 | 6 | 1,9 | 1,8 | 0,1 |
| chr2:25045245-25045745:downstream_21815 | chr2 | 25016174 | 25045245 | + | CENPO | NM | 9 | 0,1 | 2 | 0 | 9,47E-01 | NA | 9 | 1 | 13 | 20 | 4 | 3 | 6 | 18 | -1,7 | -1,8 | 0,1 |
| chr3:156863790-156864290:downstream_23054 | chr3 | 156864290 | 156878549 | - | CCNL1 | NM | 53 | 0,0 | 1 | 0 | 9,47E-01 | 9,93E-01 | 84 | 91 | 26 | 28 | 81 | 66 | 25 | 20 | 1,7 | 1,7 | 0,0 |
| chr18:29711524-29712024:downstream_20923 | chr18 | 29671817 | 29711524 | + | RNF138 | NM | 6 | -0,1 | 2 | 0 | 9,47E-01 | NA | 2 | 9 | 3 | 9 | 5 | 8 | 6 | 8 | -0,2 | -0,1 | -0,1 |
| chr3:57556589-57557089:downstream_3017 | chr3 | 57557089 | 57583215 | - | ARF4 | NM | 186 | 0,0 | 1 | 0 | 9,47E-01 | 9,93E-01 | 6 | 13 | 7 | 13 | 298 | 378 | 429 | 341 | -0,1 | -0,2 | 0,1 |
| chr6:26600277-26600777:downstream_4934 | chr6 | 26597170 | 26600277 | + | ABT1 | NM | 16 | 0,1 | 1 | 0 | 9,47E-01 | NA | 20 | 12 | 6 | 8 | 39 | 18 | 13 | 14 | 1,2 | 1,1 | 0,1 |
| chr10:82282391-82282891:downstream_17802 | chr10 | 82214037 | 82282391 | + | TSPAN14 | NM | 18 | 0,1 | 1 | 0 | 9,47E-01 | NA | 2 | 1 | 14 | 10 | 8 | 7 | 47 | 58 | -2,8 | -2,8 | 0,0 |
| chr17:646075-646575:downstream_12718 | chr17 | 635846 | 646075 | + | FAM57A | NM | 15 | -0,1 | 1 | 0 | 9,48E-01 | NA | 3 | 14 | 3 | 9 | 19 | 33 | 14 | 22 | 0,5 | 0,5 | 0,0 |
| chr20:44527458-44527958:downstream_22399 | chr20 | 44519590 | 44527458 | + | CTSA | NM | 44 | 0,1 | 1 | 0 | 9,48E-01 | 9,93E-01 | 2 | 9 | 11 | 2 | 72 | 78 | 77 | 102 | -0,3 | -0,2 | 0,0 |
| chr12:57124863-57125363:downstream_10260 | chr12 | 57125363 | 57146146 | - | PRIM1 | NM | 14 | 0,1 | 1 | 0 | 9,49E-01 | NA | 14 | 8 | 46 | 24 | 4 | 0 | 4 | 10 | -1,7 | -1,8 | 0,1 |
| chr12:112842493-112842993:downstream_18808 | chr12 | 112842993 | 112847443 | - | RPL6 | NM | 21 | -0,1 | 1 | 0 | 9,49E-01 | NA | 17 | 19 | 8 | 11 | 44 | 28 | 19 | 19 | 0,9 | 0,9 | 0,0 |
| chr3:97663830-97664330:downstream_2655 | chr3 | 97540883 | 97663830 | + | CRYBG3 | NM | 16 | 0,1 | 1 | 0 | 9,49E-01 | NA | 2 | 6 | 7 | 10 | 21 | 8 | 43 | 30 | -1,2 | -1,3 | 0,2 |
| chr3:110913016-110913516:downstream_23369 | chr3 | 110790605 | 110913016 | + | PVRL3 | NM | 78 | 0,0 | 1 | 0 | 9,49E-01 | 9,93E-01 | 40 | 68 | 149 | 134 | 26 | 37 | 78 | 91 | -1,4 | -1,4 | 0,0 |
| chr2:128237882-128238382:downstream_2227 | chr2 | 128238382 | 128284087 | - | IWS1 | NM | 28 | 0,1 | 1 | 0 | 9,49E-01 | 9,93E-01 | 5 | 2 | 7 | 10 | 20 | 37 | 57 | 89 | -1,3 | -1,4 | 0,0 |
| chr1:151023871-151024371:downstream_990 | chr1 | 151020258 | 151023871 | + | C1orf56 | NM | 13 | -0,1 | 1 | 0 | 9,50E-01 | NA | 9 | 15 | 8 | 10 | 4 | 31 | 10 | 16 | 0,4 | 0,5 | -0,1 |
| chr11:63526863-63527363:downstream_9411 | chr11 | 63527363 | 63536113 | - | C11orf95 | NM | 16 | -0,1 | 1 | 0 | 9,50E-01 | NA | 8 | 22 | 32 | 16 | 7 | 12 | 8 | 19 | -0,6 | -0,5 | -0,1 |
| chr17:46026674-46027174:downstream_13193 | chr17 | 46018888 | 46026674 | + | PNPO | NM | 14 | 0,1 | 1 | 0 | 9,50E-01 | NA | 7 | 7 | 6 | 1 | 28 | 39 | 13 | 16 | 1,1 | 1,3 | -0,1 |
| chr4:77968676-77969176:downstream_3466 | chr4 | 77969176 | 77997125 | - | CCNI | NM | 94 | 0,0 | 1 | 0 | 9,50E-01 | 9,94E-01 | 48 | 31 | 29 | 71 | 130 | 115 | 181 | 147 | -0,3 | -0,4 | 0,1 |
| chr9:34650595-34651095:downstream_26324 | chr9 | 34646585 | 34650595 | + | GALT | NM | 9 | 0,1 | 1 | 0 | 9,51E-01 | NA | 2 | 9 | 11 | 4 | 8 | 11 | 10 | 19 | -0,6 | -0,6 | 0,0 |
| chr1:112021134-112021634:downstream_16531 | chr1 | 112016490 | 112021134 | + | C1orf162 | NM | 5 | -0,2 | 3 | 0 | 9,51E-01 | NA | 9 | 0 | [illegible] | 0 | 15 | 15 | 1 | 0 | Inf | 4,4 | Inf |
| chr14:23742343-23742843:downstream_11065 | chr14 | 23742843 | 23755309 | - | HOMEZ | NM | 8 | 0,1 | 1 | 0 | 9,51E-01 | NA | 7 | 11 | 3 | 9 | 12 | 7 | 6 | 8 | 0,6 | 0,5 | 0,1 |
| chr12:56146665-56147165:downstream_10246 | chr12 | 56137063 | 56146665 | + | GDF11 | NM | 6 | -0,1 | 2 | 0 | 9,51E-01 | NA | 5 | 11 | 10 | 5 | 6 | 5 | 4 | 5 | 0,1 | 0,2 | -0,1 |
| chr7:12730558-12731058:downstream_25390 | chr7 | 12726451 | 12730558 | + | ARL4A | NM | 21 | -0,1 | 1 | 0 | 9,51E-01 | NA | 18 | 33 | 21 | 11 | 21 | 32 | 15 | 16 | 0,7 | 0,7 | -0,1 |
| chr17:58029222-58029722:downstream_13278 | chr17 | 58029722 | 58042117 | - | RNFT1 | NM | 5 | -0,1 | 2 | 0 | 9,51E-01 | NA | 3 | 3 | 7 | 4 | 6 | 4 | 11 | 5 | -0,8 | -0,7 | -0,1 |
| chr13:41301463-41301963:downstream_10714 | chr13 | 41301963 | 41302011 | - | MIR320D1 | NR | 9 | 0,1 | 1 | 0 | 9,51E-01 | NA | 8 | 10 | 1 | 6 | 23 | 9 | 4 | 10 | 1,3 | 1,2 | 0,2 |
| chr15:70946392-70946892:downstream_19824 | chr15 | 70946892 | 71055850 | - | UACA | NM | 6 | -0,1 | 2 | 0 | 9,51E-01 | NA | 1 | 0 | 6 | 9 | 1 | 1 | 14 | 13 | -4,2 | -3,9 | -0,2 |
| chr5:133844921-133845421:downstream_4421 | chr5 | 133842242 | 133844921 | + | LOC101927934 | NR | 8 | -0,1 | 2 | 0 | 9,52E-01 | NA | 7 | 2 | 19 | 20 | 1 | 2 | 6 | 5 | -2,1 | -1,9 | -0,1 |
| chrX:43085847-43086347:downstream_7680 | chrX | 43036242 | 43085847 | + | LOC101927501 | NR | 3 | 0,2 | 3 | 0 | 9,52E-01 | NA | 10 | 2 | 0 | [illegible] | 2 | 9 | 0 | 0 | Inf | Inf | NaN |
| chr7:138709951-138710451:downstream_6107 | chr7 | 138710451 | 138720775 | - | ZC3HAV1L | NM | 25 | -0,1 | 1 | 0 | 9,52E-01 | NA | 23 | 26 | 33 | 31 | 27 | 13 | 19 | 30 | -0,4 | -0,3 | -0,1 |
| chr6:16148478-16148978:downstream_4877 | chr6 | 16129316 | 16148478 | + | MYLIP | NM | 15 | 0,1 | 1 | 0 | 9,52E-01 | NA | 9 | 12 | 15 | 6 | 23 | 15 | 21 | 18 | 0,0 | -0,1 | 0,0 |
| chr4:53580305-53580805:downstream_3674 | chr4 | 53578620 | 53580305 | + | DANCR | NR | 32 | 0,1 | 1 | 0 | 9,52E-01 | 9,94E-01 | 16 | 15 | 8 | 7 | 65 | 72 | 28 | 41 | 1,0 | 1,0 | 0,0 |
| chr17:72206019-72206519:downstream_20685 | chr17 | 72199794 | 72206019 | + | RPL38 | NM | 50 | 0,0 | 1 | 0 | 9,52E-01 | 9,94E-01 | 15 | 33 | 19 | 11 | 107 | 88 | 42 | 84 | 0,6 | 0,6 | 0,0 |
| chr16:30930139-30930639:downstream_11993 | chr16 | 30930639 | 30934590 | - | FBXL19-AS1 | NR | 16 | 0,1 | 1 | 0 | 9,52E-01 | NA | 28 | 15 | 33 | 14 | 9 | 8 | 7 | 11 | -0,1 | -0,1 | 0,0 |
| chr12:72002878-72003378:downstream_10318 | chr12 | 72003378 | 72057749 | - | ZFC3H1 | NM | 18 | 0,1 | 1 | 0 | 9,53E-01 | NA | 14 | 7 | 7 | 13 | 9 | 41 | 22 | 29 | 0,0 | 0,0 | 0,1 |
| chr14:101412056-101412556:downstream_11315 | chr14 | 101411985 | 101412056 | + | SNORD113-9 | NR | 3 | 0,2 | 3 | 0 | 9,53E-01 | NA | 2 | 12 | 1 | 0 | 5 | 6 | 0 | 1 | 3,2 | 3,7 | -0,5 |
| chr8:28910242-28910742:downstream_25960 | chr8 | 28747910 | 28910242 | + | HMBOX1 | NM | 19 | 0,1 | 1 | 0 | 9,53E-01 | NA | 9 | 5 | 18 | 14 | 20 | 11 | 43 | 30 | -1,2 | -1,3 | 0,1 |
| chr6:7418270-7418770:downstream_25168 | chr6 | 7390061 | 7418270 | + | RIOK1 | NM | 9 | -0,1 | 2 | 0 | 9,54E-01 | NA | 14 | 3 | 6 | 4 | 7 | 23 | 4 | 11 | 0,8 | 1,0 | -0,2 |
| chr13:28009275-28009775:downstream_19029 | chr13 | 28009775 | 28024739 | - | MTIF3 | NM | 21 | -0,1 | 1 | 0 | 9,54E-01 | NA | 7 | 8 | 4 | 9 | 46 | 30 | 32 | 35 | 0,2 | 0,2 | 0,0 |
| chr22:21353560-21354060:downstream_22940 | chr22 | 21354060 | 21356404 | - | THAP7 | NM | 12 | 0,1 | 1 | 0 | 9,55E-01 | NA | 7 | 0 | 6 | 5 | 13 | 16 | 35 | 14 | -0,7 | -0,7 | 0,0 |
| chr15:67492512-67493012:downstream_19450 | chr15 | 67493012 | 67547536 | - | AAGAB | NM | 18 | 0,1 | 1 | 0 | 9,55E-01 | NA | 16 | 15 | 14 | 13 | 18 | 28 | 14 | 27 | 0,2 | 0,2 | 0,0 |
| chr3:122077937-122078437:downstream_23050 | chr3 | 122078437 | 122102078 | - | CCDC58 | NM | 10 | -0,1 | 1 | 0 | 9,55E-01 | NA | 8 | 7 | 6 | 8 | 20 | 6 | 14 | 9 | 0,1 | 0,1 | 0,0 |
| chr17:45018733-45019233:downstream_20500 | chr17 | 45000485 | 45018733 | + | GOSR2 | NM | 38 | 0,0 | 1 | 0 | 9,55E-01 | 9,95E-01 | 39 | 48 | 47 | 30 | 33 | 41 | 28 | 38 | 0,2 | 0,2 | 0,0 |
| chrX:20004434-20004934:downstream_7623 | chrX | 20004934 | 20007897 | - | LOC729609 | NR | 9 | 0,1 | 2 | 0 | 9,56E-01 | NA | 16 | 25 | 4 | 7 | 13 | [illegible] | 1 | 2 | 1,9 | 1,9 | 0,1 |
| chr10:105148308-105148808:downstream_17811 | chr10 | 105148808 | 105156270 | - | USMG5 | NM | 29 | 0,1 | 1 | 0 | 9,56E-01 | 9,95E-01 | 23 | 16 | 17 | 13 | 25 | 67 | 31 | 40 | 0,4 | 0,4 | 0,0 |
| chr12:76737765-76738265:downstream_10327 | chr12 | 76738265 | 76742222 | - | BBS10 | NM | 7 | 0,1 | 2 | 0 | 9,56E-01 | NA | 7 | 5 | 6 | 2 | 20 | 4 | 10 | 7 | 0,5 | 0,4 | 0,0 |
| chr16:70195184-70195684:downstream_12448 | chr16 | 70147528 | 70195184 | + | PDPR | NM | 32 | 0,0 | 1 | 0 | 9,56E-01 | 9,95E-01 | 16 | 30 | 39 | 48 | 24 | 16 | 36 | 43 | -0,9 | -1,0 | 0,1 |
| chr1:149894708-149895208:downstream_972 | chr1 | 149895208 | 149900144 | - | SF3B4 | NM | 30 | 0,1 | 1 | 0 | 9,57E-01 | 9,95E-01 | 10 | 3 | 4 | 3 | 76 | 65 | 43 | 39 | 0,8 | 0,8 | 0,0 |
| chr7:128098472-128098972:downstream_25557 | chr7 | 128095883 | 128098472 | + | HILPDA | NM | 49 | 0,0 | 1 | 0 | 9,57E-01 | 9,95E-01 | 26 | 20 | 64 | 74 | 24 | 27 | 74 | 83 | -1,6 | -1,6 | 0,0 |
| chr19:1086077-1086577:downstream_13840 | chr19 | 1086577 | 1095391 | - | POLR2E | NM | 8 | -0,1 | 2 | 0 | 9,57E-01 | NA | 7 | 3 | 3 | 4 | 11 | 19 | 6 | 15 | 0,5 | 0,6 | -0,1 |
| chr8:146281416-146281916:downstream_6802 | chr8 | 146277823 | 146281416 | + | C8orf33 | NM | 13 | 0,1 | 2 | 0 | 9,57E-01 | NA | 50 | 5 | 10 | 7 | 19 | 9 | 0 | 8 | 1,7 | 1,7 | -0,1 |
| chr5:178311424-178311924:downstream_4620 | chr5 | 178286953 | 178311424 | + | ZNF354B | NM | 15 | -0,1 | 1 | 0 | 9,57E-01 | NA | 4 | 9 | 7 | 10 | 24 | 17 | 21 | 32 | -0,4 | -0,4 | 0,0 |
| chr5:154348971-154349471:downstream_24234 | chr5 | 154320632 | 154348971 | + | MRPL22 | NM | 4 | -0,1 | 2 | 0 | 9,57E-01 | NA | 7 | 1 | 7 | 0 | 7 | 4 | 4 | 3 | 0,3 | 0,5 | -0,2 |
| chr12:56988879-56989379:downstream_18451 | chr12 | 56989379 | 57030163 | - | BAZ2A | NM | 8 | 0,1 | 2 | 0 | 9,58E-01 | NA | 5 | 7 | 18 | 7 | 7 | 1 | 11 | 7 | -1,1 | -1,2 | 0,1 |
| chr16:50139375-50139875:downstream_12346 | chr16 | 50099880 | 50139375 | + | HEATR3 | NM | 24 | 0,1 | 1 | 0 | 9,58E-01 | NA | 20 | 20 | 74 | 58 | 2 | 3 | 7 | 8 | -1,7 | -1,7 | 0,0 |
| chr16:3175029-3175529:downstream_12144 | chr16 | 3175529 | 3184883 | - | ZNF213-AS1 | NR | 5 | -0,1 | 2 | 0 | 9,58E-01 | NA | 6 | 12 | 4 | 1 | 6 | 8 | 3 | 1 | 1,8 | 1,9 | -0,1 |
| chr2:20447952-20448452:downstream_22171 | chr2 | 20448452 | 20550611 | - | PUM2 | NM | 6 | -0,1 | 2 | 0 | 9,58E-01 | NA | 4 | 1 | 8 | 5 | 5 | 4 | 11 | 10 | -1,4 | -1,3 | -0,1 |
| chrX:70525221-70525721:downstream_26867 | chrX | 70521597 | 70525221 | + | ITGB1BP2 | NM | 11 | 0,1 | 1 | 0 | 9,58E-01 | NA | 9 | 15 | 4 | 3 | 28 | 15 | 11 | 3 | 1,7 | 1,6 | 0,1 |
| chr19:40315486-40315986:downstream_21108 | chr19 | 40315986 | 40324873 | - | DYRK1B | NM | 8 | -0,1 | 2 | 0 | 9,59E-01 | NA | 15 | 7 | 7 | 4 | 8 | 14 | 8 | 2 | 1,0 | 1,0 | -0,1 |
| chr1:100548929-100549429:downstream_187 | chr1 | 100503788 | 100548929 | + | HIAT1 | NM | 24 | -0,1 | 1 | 0 | 9,60E-01 | NA | 5 | 0 | 4 | 8 | 26 | 26 | 65 | 60 | -1,3 | -1,3 | -0,1 |
| chr19:8375683-8376183:downstream_13958 | chr19 | 8376183 | 8386280 | - | NDUFA7 | NM | 39 | 0,1 | 1 | 0 | 9,60E-01 | 9,98E-01 | 4 | 10 | 1 | 7 | 75 | 97 | 45 | 71 | 0,8 | 0,6 | 0,2 |
| chrX:100417978-100418478:downstream_7530 | chrX | 100354797 | 100417978 | + | CENPI | NM | 10 | 0,1 | 1 | 0 | 9,60E-01 | NA | 3 | 3 | 22 | 16 | 2 | 2 | 10 | 18 | -2,6 | -2,7 | 0,0 |
| chr14:75408032-75408532:downstream_19336 | chr14 | 75408532 | 75422467 | - | PGF | NM | 9 | 0,1 | 2 | 0 | 9,60E-01 | NA | 14 | 30 | 0 | 2 | 15 | 8 | 1 | 0 | 4,7 | 4,0 | 0,7 |
| chr14:52471019-52471519:downstream_11139 | chr14 | 52471519 | 52535946 | - | NID2 | NM | 11 | -0,1 | 1 | 0 | 9,60E-01 | NA | 4 | 10 | 14 | 12 | 3 | 13 | 11 | 17 | -0,9 | -0,8 | -0,1 |
| chr12:21624182-21624682:downstream_10095 | chr12 | 21590537 | 21624182 | + | PYROXD1 | NM | 6 | 0,1 | 2 | 0 | 9,60E-01 | NA | 2 | 7 | 10 | 7 | 7 | 1 | 11 | 7 | -1,0 | -1,1 | 0,1 |
| chr5:44815618-44816118:downstream_4245 | chr5 | 44809026 | 44815618 | + | MRPS30 | NM | 66 | 0,0 | 1 | 0 | 9,61E-01 | 9,98E-01 | 74 | 53 | 49 | 54 | 89 | 64 | 74 | 71 | 0,3 | 0,3 | 0,0 |
| chr7:142966222-142966722:downstream_25545 | chr7 | 142960521 | 142966222 | + | GSTK1 | NM | 78 | 0,0 | 1 | 0 | 9,62E-01 | 9,99E-01 | 10 | 8 | [illegible] | 9 | 115 | 178 | 116 | 182 | 0,0 | 0,0 | 0,0 |
| chr19:57842144-57842644:downstream_14694 | chr19 | 57831864 | 57842144 | + | ZNF543 | NM | 15 | -0,1 | 1 | 0 | 9,62E-01 | NA | 7 | 7 | 1 | 5 | 20 | 48 | 18 | 14 | 1,2 | 1,1 | 0,1 |

| | | | | | | | | | | | | | | | | | | | | | | | |
|---|---|---|---|---|---|---|---|---|---|---|---|---|---|---|---|---|---|---|---|---|---|---|---|
| chr6:88052046-88052546:downstream_25213 | chr6 | 88032305 | 88052046 | + | SMIM8 | NM | 9 | 0,1 | 1 | 0 | 9,62E-01 | NA | 15 | 9 | 14 | 8 | 9 | 3 | 7 | 5 | 0,1 | 0,0 | 0,1 |
| chr5:17276954-17277454:downstream_24022 | chr5 | 17216931 | 17276954 | + | BASP1 | NM | 209 | 0,0 | 1 | 0 | 9,63E-01 | 9,99E-01 | 25 | 4 | 22 | 39 | 203 | 320 | 490 | 573 | -1,1 | -1,0 | 0,0 |
| chr10:101154087-101154587:downstream_8668 | chr10 | 101088855 | 101154087 | + | CNNM1 | NM | 9 | 0,1 | 1 | 0 | 9,64E-01 | NA | 2 | 4 | 8 | 7 | 6 | 7 | 26 | 10 | -1,4 | -1,5 | 0,1 |
| chr12:6500737-6501237:downstream_18667 | chr12 | 6484533 | 6500737 | + | LTBR | NM | 95 | 0,0 | 1 | 0 | 9,64E-01 | 9,99E-01 | 13 | 29 | 51 | 30 | 100 | 116 | 214 | 208 | -0,9 | -1,0 | 0,0 |
| chr7:100032411-100032911:downstream_5995 | chr7 | 100032911 | 100034094 | - | PPP1R35 | NM | 12 | 0,1 | 1 | 0 | 9,64E-01 | NA | 7 | 6 | 8 | 10 | 15 | 10 | 14 | 22 | -0,5 | -0,5 | 0,1 |
| chr17_gl000205_random:119732-120232:downstream_16435 | chr17 | gl000205 | random | MGC70870 | 116622 | 119732 | 8 | 0,1 | 1 | 0 | 9,64E-01 | NA | 6 | 9 | 15 | 11 | 7 | 2 | 4 | 11 | -0,9 | -0,9 | 0,0 |
| chr3:113434806-113435306:downstream_23295 | chr3 | 113435306 | 113465146 | - | NAA50 | NM | 23 | 0,0 | 1 | 0 | 9,65E-01 | NA | 24 | 18 | 28 | 24 | 20 | 18 | 17 | 31 | -0,3 | -0,3 | 0,0 |
| chr8:66622798-66623298:downstream_26019 | chr8 | 66556887 | 66622798 | + | MTFR1 | NM | 23 | 0,0 | 1 | 0 | 9,65E-01 | NA | 7 | 1 | 10 | 11 | 24 | 20 | 56 | 57 | -1,4 | -1,4 | 0,0 |
| chr2:210886984-210887484:downstream_22194 | chr2 | 210867288 | 210886984 | + | RPE | NM | 38 | 0,0 | 1 | 0 | 9,65E-01 | 1,00E+00 | 16 | 21 | 22 | 14 | 32 | 89 | 53 | 58 | 0,1 | 0,1 | -0,1 |
| chr7:35946715-35947215:downstream_25722 | chr7 | 35840595 | 35946715 | + | sept-07 | NM | 8 | 0,1 | 1 | 0 | 9,66E-01 | NA | 4 | 12 | 7 | 10 | 7 | 9 | 14 | 5 | -0,1 | -0,2 | 0,1 |
| chr15:83806303-83806803:downstream_11859 | chr15 | 83806803 | 83876770 | - | HDGFRP3 | NM | 33 | 0,0 | 1 | 0 | 9,66E-01 | 1,00E+00 | 24 | 16 | 56 | 36 | 21 | 19 | 54 | 40 | -1,2 | -1,2 | 0,0 |
| chr11:214529-215029:downstream_18289 | chr11 | 215029 | 236362 | - | SIRT3 | NM | 8 | -0,1 | 1 | 0 | 9,66E-01 | NA | 2 | 7 | 15 | 13 | 2 | 6 | 11 | 11 | -1,6 | -1,6 | -0,1 |
| chr3:12625210-12625710:downstream_23276 | chr3 | 12598512 | 12625210 | + | MKRN2 | NM | 18 | -0,1 | 1 | 0 | 9,67E-01 | NA | 9 | 1 | 13 | 4 | 20 | 27 | 32 | 38 | -0,7 | -0,6 | -0,2 |
| chr15:52014760-52015260:downstream_19610 | chr15 | 52015260 | 52043650 | - | LYSMD2 | NM | 14 | 0,1 | 2 | 0 | 9,67E-01 | NA | 1 | 20 | 0 | 5 | 16 | 53 | 8 | 11 | 2,1 | 1,9 | 0,2 |
| chr2:187374087-187374587:downstream_2347 | chr2 | 187350884 | 187374087 | + | ZC3H15 | NM | 67 | 0,0 | 1 | 0 | 9,67E-01 | 1,00E+00 | 7 | 5 | 8 | 4 | 115 | 146 | 129 | 125 | 0,0 | 0,0 | -0,1 |
| chr13:88452797-88453297:downstream_10820 | chr13 | 88453297 | 88462775 | - | LINC00397 | NR | 8 | -0,1 | 2 | 0 | 9,67E-01 | NA | 3 | 12 | 17 | 9 | 7 | 1 | 11 | 3 | -0,8 | -0,8 | 0,0 |
| chr2:38789827-38790327:downstream_21968 | chr2 | 38790327 | 38830178 | - | HNRNPLL | NM | 24 | 0,0 | 1 | 0 | 9,67E-01 | NA | 25 | 16 | 21 | 17 | 19 | 39 | 25 | 28 | 0,1 | 0,1 | 0,0 |
| chr18:3458406-3458906:downstream_20960 | chr18 | 3411924 | 3458406 | + | TGIF1 | NM | 41 | 0,0 | 1 | 0 | 9,68E-01 | 1,00E+00 | 6 | 6 | 21 | 17 | 25 | 40 | 106 | 112 | -1,7 | -1,7 | 0,0 |
| chr7:105205079-105205579:downstream_6035 | chr7 | 105205579 | 105221976 | - | EFCAB10 | NR | 4 | 0,1 | 2 | 0 | 9,68E-01 | NA | 6 | 4 | 1 | 1 | 12 | 2 | 3 | 1 | 2,1 | 2,0 | 0,1 |
| chr12:86197830-86198330:downstream_10339 | chr12 | 86198330 | 86230318 | - | RASSF9 | NM | 30 | 0,0 | 1 | 0 | 9,68E-01 | 1,00E+00 | 7 | 3 | 60 | 53 | 10 | 0 | 53 | 54 | -3,5 | -3,5 | 0,0 |
| chr12:77414525-77415025:downstream_10328 | chr12 | 77415025 | 77459360 | - | E2F7 | NM | 6 | -0,1 | 2 | 0 | 9,68E-01 | NA | 15 | 0 | 0 | 4 | 18 | 8 | 4 | 3 | 1,8 | 1,8 | 0,1 |
| chrX:14939459-14939959:downstream_26932 | chrX | 14891526 | 14939459 | + | MOSPD2 | NM | 14 | 0,0 | 1 | 0 | 9,70E-01 | NA | 11 | 10 | 26 | 19 | 6 | 9 | 18 | 15 | -1,2 | -1,2 | 0,0 |
| chr11:202423-202923:downstream_17894 | chr11 | 202923 | 207422 | - | BET1L | NM | 20 | 0,0 | 1 | 0 | 9,70E-01 | NA | 20 | 12 | 7 | 17 | 36 | 21 | 28 | 17 | 0,4 | 0,3 | 0,0 |
| chr5:180684587-180685087:downstream_4637 | chr5 | 180673540 | 180684587 | + | CTC-338M12.4 | NR | 11 | 0,0 | 1 | 0 | 9,71E-01 | NA | 6 | 15 | 8 | 7 | 14 | 17 | 18 | 5 | 0,4 | 0,4 | 0,0 |
| chr14:92630543-92631043:downstream_11272 | chr14 | 92588297 | 92630543 | + | CPSF2 | NM | 21 | 0,0 | 1 | 0 | 9,71E-01 | NA | 11 | 6 | 25 | 28 | 10 | 13 | 29 | 43 | -1,7 | -1,7 | 0,0 |
| chr20:34844863-34845363:downstream_22340 | chr20 | 34824338 | 34844863 | + | AAR2 | NM | 7 | 0,1 | 2 | 0 | 9,71E-01 | NA | 7 | 0 | 3 | 10 | 11 | 1 | 10 | 11 | -0,8 | -0,9 | 0,1 |
| chr20:29634007-29634507:downstream_14780 | chr20 | 29611878 | 29634007 | + | FRG1B | NR | 7 | -0,1 | 2 | 0 | 9,71E-01 | NA | 6 | 9 | 8 | 7 | 7 | 7 | 3 | 11 | -0,1 | 0,1 | -0,1 |
| chr1:33961995-33962495:downstream_107 | chr1 | 33938231 | 33961995 | + | ZSCAN20 | NM | 8 | -0,1 | 2 | 0 | 9,72E-01 | NA | 6 | 0 | 1 | 7 | 18 | 2 | 15 | 15 | -0,6 | -0,6 | 0,0 |
| chr16:61089110-61089610:downstream_12396 | chr16 | 61089610 | 61089670 | - | MIR4426 | NR | 256 | 0,0 | 1 | 0 | 9,72E-01 | 1,00E+00 | 143 | 96 | 49 | 52 | 720 | 485 | 264 | 236 | 1,3 | 1,3 | 0,0 |
| chr9:113005591-113006091:downstream_26561 | chr9 | 113006091 | 113018920 | - | TXN | NM | 77 | 0,0 | 1 | 0 | 9,72E-01 | 1,00E+00 | 49 | 63 | 45 | 40 | 111 | 127 | 74 | 107 | 0,4 | 0,4 | 0,0 |
| chr2:39475906-39476406:downstream_22068 | chr2 | 39476406 | 39664453 | - | MAP4K3 | NM | 9 | -0,1 | 2 | 0 | 9,72E-01 | NA | 1 | 0 | 10 | 13 | 2 | 0 | 18 | 25 | -4,8 | -4,7 | -0,1 |
| chr2:74382710-74383210:downstream_2041 | chr2 | 74383210 | 74405995 | - | MOB1A | NM | 198 | 0,0 | 1 | 0 | 9,72E-01 | 1,00E+00 | 172 | 117 | 218 | 162 | 211 | 189 | 259 | 258 | -0,4 | -0,4 | 0,0 |
| chr3:10005135-10005635:downstream_2814 | chr3 | 10005635 | 10028522 | - | EMC3 | NM | 98 | 0,0 | 1 | 0 | 9,72E-01 | 1,00E+00 | 23 | 13 | 17 | 15 | 187 | 200 | 146 | 186 | 0,2 | 0,2 | -0,1 |
| chr9:102740962-102741462:downstream_6822 | chr9 | 102741462 | 102861334 | - | ERP44 | NM | 14 | 0,0 | 1 | 0 | 9,72E-01 | NA | 4 | 16 | 18 | 19 | 4 | 15 | 24 | 13 | -0,8 | -0,9 | 0,1 |
| chr18:33513550-33514050:downstream_13609 | chr18 | 33514050 | 33514105 | - | MIR3929 | NR | 4 | 0,1 | 3 | 0 | 9,73E-01 | NA | 4 | 11 | 0 | 0 | 4 | 10 | 0 | 0 | Inf | Inf | NaN |
| chr2:190535557-190536057:downstream_2352 | chr2 | 190526124 | 190535557 | + | ASNSD1 | NM | 21 | 0,0 | 1 | 0 | 9,74E-01 | NA | 22 | 10 | 4 | 14 | 25 | 47 | 18 | 24 | 0,8 | 0,8 | 0,0 |
| chr5:133531647-133532147:downstream_4415 | chr5 | 133532147 | 133561950 | - | PPP2CA | NM | 20 | 0,0 | 1 | 0 | 9,74E-01 | NA | 7 | 16 | 11 | 21 | 29 | 14 | 18 | 43 | -0,5 | -0,5 | 0,1 |
| chr15:34375723-34376223:downstream_11640 | chr15 | 34376223 | 34394053 | - | EMC7 | NM | 141 | 0,0 | 1 | 0 | 9,74E-01 | 1,00E+00 | 12 | 15 | 11 | 9 | 304 | 328 | 198 | 249 | 0,5 | 0,5 | 0,0 |
| chr5:179263765-179264265:downstream_24034 | chr5 | 179264265 | 179285840 | - | C5orf45 | NM | 23 | 0,0 | 1 | 0 | 9,74E-01 | NA | 29 | 36 | 19 | 34 | 17 | 20 | 18 | 13 | 0,3 | 0,3 | 0,0 |
| chr1:212535205-212535705:downstream_17108 | chr1 | 212458878 | 212535205 | + | PPP2R5A | NM | 28 | 0,0 | 1 | 0 | 9,75E-01 | NA | 2 | 1 | 4 | 7 | 13 | 26 | 92 | 76 | -2,0 | -2,1 | 0,1 |
| chr14:38058256-38058756:downstream_11114 | chr14 | 38058756 | 38064325 | - | FOXA1 | NM | 7 | 0,1 | 2 | 0 | 9,75E-01 | NA | 3 | 0 | 10 | 3 | 2 | 6 | 15 | 13 | -2,0 | -1,9 | -0,1 |
| chr3:195964752-195965252:downstream_3347 | chr3 | 195965252 | 196014584 | - | PCYT1A | NM | 21 | 0,0 | 1 | 0 | 9,75E-01 | NA | 15 | 11 | 18 | 26 | 14 | 22 | 21 | 41 | -0,8 | -0,8 | 0,0 |
| chr16:3077756-3078256:downstream_20240 | chr16 | 3074031 | 3077756 | + | THOC6 | NM | 9 | 0,1 | 2 | 0 | 9,75E-01 | NA | 11 | 1 | 0 | 0 | 27 | 32 | 0 | 2 | Inf | 5,2 | Inf |
| chr5:77654087-77654587:downstream_4314 | chr5 | 77654587 | 77656365 | - | SCAMP1-AS1 | NR | 20 | 0,0 | 1 | 0 | 9,75E-01 | NA | 9 | 4 | 8 | 4 | 34 | 35 | 40 | 22 | 0,0 | 0,1 | -0,1 |
| chr2:9628591-9629091:downstream_1849 | chr2 | 9614669 | 9628591 | + | IAH1 | NM | 60 | 0,0 | 1 | 0 | 9,75E-01 | 1,00E+00 | 27 | 23 | 21 | 25 | 110 | 95 | 82 | 102 | 0,1 | 0,2 | 0,0 |
| chr17:48444727-48445227:downstream_13227 | chr17 | 48445227 | 48450562 | - | MRPL27 | NM | 61 | 0,0 | 1 | 0 | 9,76E-01 | 1,00E+00 | 14 | 10 | 10 | 8 | 131 | 126 | 65 | 122 | 0,4 | 0,5 | -0,1 |
| chr8:25270619-25271119:downstream_6220 | chr8 | 25042286 | 25270619 | + | DOCK5 | NM | 128 | 0,0 | 1 | 0 | 9,76E-01 | 1,00E+00 | 92 | 39 | 291 | 330 | 23 | 24 | 124 | 102 | -2,2 | -2,3 | 0,0 |
| chr7:6063465-6063965:downstream_5700 | chr7 | 6048881 | 6063465 | + | AIMP2 | NM | 113 | 0,0 | 1 | 0 | 9,76E-01 | 1,00E+00 | 99 | 202 | 181 | 153 | 55 | 73 | 63 | 76 | -0,1 | -0,1 | 0,0 |
| chr7:138270332-138270832:downstream_25788 | chr7 | 138145078 | 138270332 | + | TRIM24 | NM | 25 | 0,0 | 1 | 0 | 9,76E-01 | NA | 5 | 0 | 10 | 15 | 20 | 8 | 83 | 57 | -2,3 | -2,3 | 0,0 |
| chr5:154266475-154266975:downstream_24142 | chr5 | 154266975 | 154317776 | - | GEMIN5 | NM | 16 | 0,0 | 1 | 0 | 9,77E-01 | NA | 11 | 4 | 18 | 16 | 11 | 14 | 25 | 33 | -1,2 | -1,2 | 0,0 |
| chr11:20405329-20405829:downstream_18054 | chr11 | 20385230 | 20405329 | + | HTATIP2 | NM | 55 | 0,1 | 2 | 0 | 9,77E-01 | 1,00E+00 | 11 | 0 | 10 | 2 | 107 | 95 | 88 | 128 | -0,1 | -0,1 | 0,0 |
| chr2:48606434-48606934:downstream_1970 | chr2 | 48541794 | 48606434 | + | FOXN2 | NM | 18 | 0,0 | 1 | 0 | 9,77E-01 | NA | 9 | 0 | 11 | 3 | 17 | 32 | 42 | 33 | -0,7 | -0,6 | -0,1 |
| chr20:23335408-23335908:downstream_14918 | chr20 | 23331372 | 23335408 | + | NXT1 | NM | 11 | 0,0 | 1 | 0 | 9,77E-01 | NA | 7 | 7 | 6 | 5 | 20 | 16 | 14 | 14 | 0,4 | 0,4 | 0,0 |
| chr6:42174038-42174538:downstream_5138 | chr6 | 42174538 | 42185633 | - | MRPS10 | NM | 7 | -0,1 | 2 | 0 | 9,77E-01 | NA | 9 | 0 | 1 | 3 | 17 | 11 | 4 | 10 | 0,9 | 1,0 | 0,0 |
| chr12:118469996-118470496:downstream_18917 | chr12 | 118470496 | 118499979 | - | WSB2 | NM | 11 | 0,0 | 1 | 0 | 9,78E-01 | NA | 9 | 11 | 15 | 5 | 11 | 12 | 11 | 11 | 0,0 | 0,0 | 0,0 |
| chr17:77767675-77768175:downstream_13409 | chr17 | 77768175 | 77770915 | - | CBX8 | NM | 18 | 0,0 | 1 | 0 | 9,78E-01 | NA | 7 | 3 | 4 | 7 | 30 | 23 | 32 | 36 | -0,3 | -0,4 | 0,1 |
| chr6:76001580-76002080:downstream_5235 | chr6 | 75994729 | 76001580 | + | LOC100506804 | NR | 14 | 0,0 | 1 | 0 | 9,78E-01 | NA | 4 | 19 | 7 | 6 | 32 | 17 | 15 | 12 | 0,9 | 0,8 | 0,1 |
| chr1:212800120-212800620:downstream_1359 | chr1 | 212797788 | 212800120 | + | FAM71A | NM | 3 | -0,2 | 7 | 0 | 9,78E-01 | NA | 7 | 5 | 0 | 0 | 14 | 0 | 0 | 0 | Inf | Inf | NaN |
| chr12:29493078-29493578:downstream_10116 | chr12 | 29493578 | 29534143 | - | ERGIC2 | NM | 219 | 0,0 | 1 | 0 | 9,78E-01 | 1,00E+00 | 307 | 215 | 419 | 282 | 123 | 104 | 159 | 142 | -0,4 | -0,4 | 0,0 |
| chr16:2949383-2949883:downstream_19992 | chr16 | 2933195 | 2949383 | + | FLYWCH2 | NM | 7 | 0,0 | 2 | 0 | 9,78E-01 | NA | 8 | 6 | 13 | 5 | 12 | 0 | 8 | 7 | -0,3 | -0,4 | 0,0 |
| chr8:95891952-95892452:downstream_6645 | chr8 | 95892452 | 95907482 | - | CCNE2 | NM | 5 | 0,1 | 2 | 0 | 9,79E-01 | NA | 5 | 0 | 10 | 11 | 3 | 0 | 4 | 10 | -2,1 | -2,1 | 0,0 |
| chr12:97347469-97347969:downstream_18704 | chr12 | 97301000 | 97347469 | + | NEDD1 | NM | 16 | 0,0 | 1 | 0 | 9,79E-01 | NA | 13 | 2 | 14 | 9 | 20 | 16 | 25 | 27 | -0,6 | -0,5 | -0,1 |
| chr2:100016437-100016937:downstream_22185 | chr2 | 100016937 | 100106480 | - | REV1 | NM | 11 | 0,0 | 1 | 0 | 9,79E-01 | NA | 8 | 3 | 22 | 14 | 4 | 5 | 13 | 16 | -1,7 | -1,7 | 0,0 |
| chr2:129022553-129023053:downstream_2232 | chr2 | 129023053 | 129076171 | - | HS6ST1 | NM | 11 | 0,0 | 1 | 0 | 9,79E-01 | NA | 2 | 5 | 8 | 5 | 9 | 14 | 32 | 16 | -1,0 | -1,1 | 0,0 |
| chr12:114181881-114182381:downstream_18639 | chr12 | 114182381 | 114211488 | - | LINC01234 | NR | 15 | 0,0 | 1 | 0 | 9,79E-01 | NA | 6 | 7 | 11 | 9 | 11 | 23 | 21 | 33 | -0,7 | -0,6 | -0,1 |
| chrX:119392251-119392751:downstream_27121 | chrX | 119384606 | 119392251 | + | ZBTB33 | NM | 37 | 0,0 | 1 | 0 | 9,79E-01 | 1,00E+00 | 6 | 3 | 1 | 4 | 62 | 104 | 53 | 63 | 0,6 | 0,5 | 0,1 |
| chr9:116055185-116055685:downstream_26454 | chr9 | 116037913 | 116055185 | + | PRPF4 | NM | 19 | 0,0 | 1 | 0 | 9,79E-01 | NA | 20 | 17 | 10 | 13 | 27 | 29 | 10 | 23 | 0,7 | 0,8 | 0,0 |
| chr9:123939886-123940386:downstream_6931 | chr9 | 123850573 | 123939886 | + | CNTRL | NM | 4 | 0,1 | 2 | 0 | 9,79E-01 | NA | 1 | 4 | 0 | 8 | 3 | 4 | 8 | 6 | -0,8 | -1,0 | 0,2 |
| chr1:22419436-22419936:downstream_16577 | chr1 | 22379119 | 22419436 | + | CDC42 | NM | 93 | 0,0 | 1 | 0 | 9,80E-01 | 1,00E+00 | 15 | 10 | 19 | 15 | 143 | 153 | 170 | 217 | -0,4 | -0,4 | -0,1 |
| chr5:94031573-94032073:downstream_4051 | chr5 | 93954390 | 94031573 | + | ANKRD32 | NM | 4 | -0,1 | 3 | 0 | 9,80E-01 | NA | 1 | 0 | 8 | 8 | 1 | 0 | 11 | 5 | -4,3 | -4,3 | 0,0 |
| chr5:173387994-173388494:downstream_24071 | chr5 | 173315330 | 173387994 | + | CPEB4 | NM | 5 | 0,0 | 2 | 0 | 9,80E-01 | NA | 4 | 6 | 4 | 6 | 8 | 2 | 1 | 8 | 0,0 | 0,1 | -0,1 |
| chr9:131395439-131395939:downstream_7342 | chr9 | 131395939 | 131419129 | - | WDR34 | NM | 32 | 0,0 | 1 | 0 | 9,81E-01 | 1,00E+00 | 37 | 25 | 38 | 39 | 35 | 16 | 21 | 42 | -0,3 | -0,3 | 0,0 |
| chr19:42385439-42385939:downstream_21055 | chr19 | 42381189 | 42385439 | + | CD79A | NM | 6 | -0,1 | 3 | 0 | 9,81E-01 | NA | 6 | 8 | 0 | 0 | 10 | 27 | 0 | 1 | Inf | 5,5 | Inf |
| chr6:27774756-27775256:downstream_4950 | chr6 | 27775256 | 27775709 | - | HIST1H2BL | NM | 71 | 0,0 | 1 | 0 | 9,81E-01 | 1,00E+00 | 1 | 1 | 3 | 7 | 61 | 17 | 203 | 271 | -2,5 | -2,6 | 0,1 |
| chr11:73685215-73685715:downstream_9528 | chr11 | 73685715 | 73693889 | - | UCP2 | NM | 4 | 0,0 | 2 | 0 | 9,81E-01 | NA | 7 | 1 | 3 | 2 | 7 | 7 | 1 | 6 | 0,9 | 1,0 | -0,1 |
| chr17:41604164-41604664:downstream_20450 | chr17 | 41561284 | 41604164 | + | DHX8 | NM | 4 | -0,1 | 2 | 0 | 9,81E-01 | NA | 7 | 2 | 3 | 1 | 4 | 9 | 1 | 3 | 1,2 | 1,5 | -0,2 |
| chr16:18802490-18802990:downstream_12232 | chr16 | 18802990 | 18812857 | - | ARL6IP1 | NM | 616 | 0,0 | 1 | 0 | 9,82E-01 | 1,00E+00 | 7 | 5 | 3 | 6 | 1172 | 1571 | 969 | 1195 | 0,4 | 0,3 | 0,1 |
| chr12:58115340-58115840:downstream_18721 | chr12 | 58087737 | 58115340 | + | OS9 | NM | 140 | 0,0 | 1 | 0 | 9,82E-01 | 1,00E+00 | 2 | 15 | 8 | 3 | 296 | 380 | 212 | 202 | 0,6 | 0,7 | -0,1 |
| chr4:170014906-170015406:downstream_3536 | chr4 | 170015406 | 170192249 | - | SH3RF1 | NM | 7 | 0,0 | 2 | 0 | 9,82E-01 | NA | 2 | 0 | 4 | 7 | 6 | 0 | 17 | 22 | -2,8 | -2,8 | -0,1 |
| chr12:46312413-46312913:downstream_10132 | chr12 | 46312913 | 46384401 | - | SCAF11 | NM | 23 | 0,0 | 1 | 0 | 9,82E-01 | NA | 2 | 0 | 6 | 6 | 17 | 14 | 77 | 60 | -2,2 | -2,2 | -0,1 |
| chr10:89731687-89732187:downstream_17734 | chr10 | 89623194 | 89731687 | + | PTEN | NM | 6 | 0,0 | 2 | 0 | 9,83E-01 | NA | 11 | 1 | 7 | 7 | 2 | 7 | 6 | 4 | -0,1 | -0,1 | -0,1 |
| chr1:23410015-23410515:downstream_16924 | chr1 | 23410515 | 23495351 | - | LUZP1 | NM | 12 | 0,0 | 1 | 0 | 9,83E-01 | NA | 7 | 8 | 13 | 21 | 12 | 1 | 22 | 10 | -1,2 | -1,3 | 0,0 |
| chr7:155434109-155434609:downstream_6208 | chr7 | 155434609 | 155437100 | - | LOC100506302 | NR | 34 | 0,0 | 1 | 0 | 9,83E-01 | 1,00E+00 | 48 | 30 | 31 | 21 | 41 | 47 | 24 | 33 | 0,6 | 0,6 | -0,1 |
| chr10:104838344-104838844:downstream_17522 | chr10 | 104678074 | 104838344 | + | CNNM2 | NM | 4 | 0,1 | 2 | 0 | 9,83E-01 | NA | 2 | 0 | 4 | 8 | 2 | 0 | 6 | 7 | -2,9 | -3,0 | 0,1 |
| chrX:55785207-55785707:downstream_27013 | chrX | 55744109 | 55785207 | + | RRAGB | NM | 11 | 0,0 | 1 | 0 | 9,83E-01 | NA | 9 | 5 | 13 | 11 | 14 | 6 | 15 | 18 | -0,8 | -0,8 | 0,0 |
| chr12:14938911-14939411:downstream_9829 | chr12 | 14939411 | 14956401 | - | WBP11 | NM | 17 | 0,0 | 1 | 0 | 9,83E-01 | NA | 7 | 4 | 8 | 7 | 30 | 16 | 29 | 36 | -0,5 | -0,5 | 0,0 |
| chr9:114122472-114122972:downstream_6912 | chr9 | 114122972 | 114247025 | - | KIAA0368 | NM | 8 | 0,0 | 2 | 0 | 9,83E-01 | NA | 0 | 1 | 8 | 4 | 2 | 1 | 24 | 22 | -3,7 | -3,7 | 0,1 |
| chr12:12619838-12620338:downstream_18660 | chr12 | 12510012 | 12619838 | + | LOH12CR1 | NM | 9 | 0,0 | 1 | 0 | 9,84E-01 | NA | 3 | 5 | 10 | 17 | 5 | 4 | 17 | 14 | -1,7 | -1,8 | 0,1 |
| chr21:17252390-17252890:downstream_22726 | chr21 | 17102343 | 17252390 | + | USP25 | NM | 7 | 0,0 | 2 | 0 | 9,84E-01 | NA | 2 | 1 | 13 | 12 | 2 | 1 | 7 | 18 | -3,3 | -3,3 | 0,0 |
| chr12:6647541-6648041:downstream_18573 | chr12 | 6643570 | 6647541 | + | GAPDH | NM | 896 | 0,0 | 1 | 0 | 9,84E-01 | 1,00E+00 | 4 | 7 | 4 | 3 | 2274 | 1999 | 1142 | 1732 | 0,5 | 0,6 | 0,0 |
| chrX:11141204-11141704:downstream_26848 | chrX | 11129405 | 11141204 | + | HCCS | NM | 26 | 0,0 | 1 | 0 | 9,84E-01 | NA | 7 | 4 | 1 | 4 | 81 | 46 | 29 | 37 | 1,0 | 0,9 | 0,1 |
| chr5:1317499-1317999:downstream_4161 | chr5 | 1317999 | 1345002 | - | CLPTM1L | NM | 97 | 0,0 | 1 | 0 | 9,84E-01 | 1,00E+00 | 19 | 36 | 18 | 31 | 139 | 209 | 182 | 145 | 0,1 | 0,1 | 0,1 |
| chr4:48498879-48499379:downstream_3435 | chr4 | 48499379 | 48782316 | - | FRYL | NM | 4 | 0,0 | 2 | 0 | 9,85E-01 | NA | 2 | 0 | 8 | 3 | 1 | 2 | 8 | 9 | -2,8 | -2,7 | -0,2 |
| chr3:151599876-151600376:downstream_3239 | chr3 | 151591430 | 151599876 | + | SUCNR1 | NM | 76 | 0,0 | 1 | 0 | 9,85E-01 | 1,00E+00 | 24 | 10 | 96 | 89 | 36 | 24 | 189 | 136 | -2,4 | -2,4 | 0,0 |
| chr19:9675903-9676403:downstream_21604 | chr19 | 9676403 | 9695209 | - | ZNF121 | NM | 86 | 0,0 | 1 | 0 | 9,86E-01 | 1,00E+00 | 37 | 52 | 19 | 14 | 229 | 184 | 65 | 86 | 1,4 | 1,4 | 0,0 |
| chr13:53262433-53262933:downstream_19101 | chr13 | 53226830 | 53262433 | + | SUGT1 | NM | 31 | 0,0 | 1 | 0 | 9,86E-01 | 1,00E+00 | 19 | 10 | 32 | 31 | 32 | 17 | 61 | 49 | -1,2 | -1,2 | 0,0 |
| chr2:42285668-42286168:downstream_1947 | chr2 | 42275160 | 42285668 | + | PKDCC | NM | 35 | 0,0 | 2 | 0 | 9,86E-01 | 1,00E+00 | 1 | 0 | 22 | 17 | 5 | 0 | 121 | 113 | -5,6 | -5,6 | 0,0 |
| chr2:203634480-203634980:downstream_1780 | chr2 | 203499900 | 203634480 | + | FAM117B | NM | 4 | 0,0 | 3 | 0 | 9,87E-01 | NA | 0 | 0 | 1 | 7 | 1 | 0 | 7 | 18 | -Inf | -4,9 | -Inf |
| chr2:10074545-10075045:downstream_1851 | chr2 | 9983570 | 10074545 | + | TAF1B | NM | 15 | 0,0 | 1 | 0 | 9,88E-01 | NA | 5 | 3 | 18 | 16 | 11 | 4 | 32 | 33 | -2,1 | -2,2 | 0,0 |
| chr13:21723224-21723724:downstream_10653 | chr13 | 21714652 | 21723224 | + | SAP18 | NM | 8 | 0,0 | 1 | 0 | 9,88E-01 | NA | 7 | 8 | 6 | 5 | 8 | 14 | 10 | 6 | 0,5 | 0,5 | 0,0 |
| chr10:33188745-33189245:downstream_17613 | chr10 | 33189245 | 33247293 | - | ITGB1 | NM | 150 | 0,0 | 1 | 0 | 9,89E-01 | 1,00E+00 | 136 | 156 | 277 | 262 | 74 | 55 | 138 | 104 | -0,9 | -0,9 | 0,0 |
| chr13:21753220-21753720:downstream_10654 | chr13 | 21750371 | 21753220 | + | MRPL57 | NM | 20 | 0,0 | 1 | 0 | 9,89E-01 | NA | 13 | 22 | 7 | 9 | 34 | 40 | 15 | 19 | 1,1 | 1,1 | 0,0 |
| chr17:42472144-42472644:downstream_20501 | chr17 | 42472644 | 42580970 | - | GPATCH8 | NM | 8 | 0,0 | 2 | 0 | 9,89E-01 | NA | 3 | 0 | 6 | 5 | 4 | 9 | 22 | 17 | -1,7 | -1,6 | -0,1 |
| chr1:23883920-23884420:downstream_544 | chr1 | 23884420 | 23886285 | - | ID3 | NM | 23 | 0,0 | 1 | 0 | 9,89E-01 | NA | 16 | 15 | 18 | 43 | 16 | 13 | 33 | 25 | -1,0 | -1,0 | 0,1 |
| chr15:50650503-50651003:downstream_11719 | chr15 | 50646370 | 50650503 | + | GABPB1-AS1 | NR | 66 | 0,0 | 1 | 0 | 9,90E-01 | 1,00E+00 | 54 | 104 | 68 | 39 | 100 | 56 | 56 | 51 | 0,6 | 0,6 | 0,0 |
| chr7:127231759-127232259:downstream_6067 | chr7 | 127228405 | 127231759 | + | ARF5 | NM | 16 | 0,0 | 3 | 0 | 9,90E-01 | NA | 0 | 12 | 8 | 0 | 26 | 40 | 19 | 25 | 0,5 | 0,5 | -0,1 |
| chr16:3589051-3589551:downstream_19938 | chr16 | 3550944 | 3589051 | + | CLUAP1 | NM | 7 | 0,0 | 2 | 0 | 9,90E-01 | NA | 10 | 0 | 7 | 6 | 8 | 7 | 7 | 11 | -0,4 | -0,3 | -0,1 |
| chr8:109213471-109213971:downstream_6679 | chr8 | 109213971 | 109260959 | - | EIF3E | NM | 29 | 0,0 | 1 | 0 | 9,91E-01 | 1,00E+00 | 21 | 12 | 25 | 30 | 37 | 15 | 53 | 37 | -0,8 | -0,8 | 0,0 |
| chr1:245003439-245003939:downstream_1454 | chr1 | 245003939 | 245010243 | - | HNRNPU-AS1 | NR | 58 | 0,0 | 1 | 0 | 9,91E-01 | 1,00E+00 | 22 | 16 | 29 | 20 | 114 | 55 | 114 | 97 | -0,3 | -0,3 | 0,0 |
| chr20:3019315-3019815:downstream_22520 | chr20 | 2844840 | 3019315 | + | PTPRA | NM | 15 | 0,0 | 2 | 0 | 9,91E-01 | NA | 2 | 0 | 4 | 7 | 9 | 5 | 45 | 52 | -2,8 | -2,8 | 0,0 |
| chr3:193920304-193920804:downstream_23244 | chr3 | 193920804 | 193967942 | - | LOC100505920 | NR | 2 | -0,1 | 7 | 0 | 9,92E-01 | NA | 2 | 7 | 0 | 0 | 0 | 10 | 0 | 0 | Inf | Inf | NaN |
| chr13:45858239-45858739:downstream_10593 | chr13 | 45694630 | 45858239 | + | GTF2F2 | NM | 21 | 0,0 | 1 | 0 | 9,92E-01 | NA | 11 | 1 | 22 | 13 | 25 | 5 | 56 | 35 | -1,6 | -1,6 | 0,0 |
| chr21:43406439-43406939:downstream_22730 | chr21 | 43406939 | 43430496 | - | ZBTB21 | NM | 7 | 0,0 | 2 | 0 | 9,93E-01 | NA | 8 | 0 | 4 | 0 | 19 | 13 | 8 | 6 | 1,0 | 1,2 | -0,2 |
| chr14:73741417-73741917:downstream_19329 | chr14 | 73741917 | 73925286 | - | NUMB | NM | 9 | 0,0 | 2 | 0 | 9,93E-01 | NA | 1 | 0 | 7 | 4 | 2 | 2 | 28 | 25 | -3,8 | -3,6 | -0,2 |
| chr20:52769487-52769987:downstream_22400 | chr20 | 52769987 | 52790516 | - | CYP24A1 | NM | 208 | 0,0 | 1 | 0 | 9,93E-01 | 1,00E+00 | 3 | 0 | 14 | 16 | 64 | 104 | 655 | 808 | -3,2 | -3,1 | -0,1 |
| chr1:226107076-226107576:downstream_17148 | chr1 | 226107576 | 226112040 | - | PYCR2 | NM | 75 | 0,0 | 1 | 0 | 9,93E-01 | 1,00E+00 | 11 | 16 | 11 | 6 | 155 | 192 | 90 | 116 | 0,7 | 0,7 | -0,1 |
| chr8:77891993-77892493:downstream_26048 | chr8 | 77892493 | 77913280 | - | PEX2 | NM | 13 | 0,0 | 1 | 0 | 9,93E-01 | NA | 25 | 18 | 15 | 7 | 11 | 15 | 6 | 7 | 1,0 | 1,0 | -0,1 |
| chr10:75001213-75001713:downstream_8565 | chr10 | 75001713 | 75007089 | - | DNAJC9 | NM | 31 | 0,0 | 1 | 0 | 9,93E-01 | 1,00E+00 | 39 | 49 | 24 | 16 | 41 | 42 | 18 | 18 | 1,2 | 1,2 | 0,0 |
| chr7:99751542-99752042:downstream_25414 | chr7 | 99752042 | 99756344 | - | C7orf43 | NM | 15 | 0,0 | 1 | 0 | 9,93E-01 | NA | 12 | 20 | 4 | 5 | 19 | 41 | 10 | 7 | 1,8 | 1,8 | 0,0 |
| chr19:39875537-39876037:downstream_21391 | chr19 | 39833098 | 39875537 | + | SAMD4B | NM | 19 | 0,0 | 1 | 0 | 9,93E-01 | NA | 15 | 11 | 11 | 11 | 21 | 35 | 24 | 23 | 0,3 | 0,3 | 0,0 |
| chr14:77253085-77253585:downstream_11242 | chr14 | 77253585 | 77279283 | - | ANGEL1 | NM | 7 | 0,0 | 2 | 0 | 9,93E-01 | NA | 3 | 3 | 7 | 6 | 11 | 1 | 17 | 9 | -1,0 | -1,1 | 0,0 |
| chr17:38248536-38249036:downstream_13039 | chr17 | 38249036 | 38256978 | - | NR1D1 | NM | 4 | 0,0 | 3 | 0 | 9,93E-01 | NA | 5 | 4 | 1 | 0 | 7 | 11 | 1 | 1 | 2,7 | 3,0 | -0,3 |
| chr10:103866824-103867324:downstream_17624 | chr10 | 103867324 | 103880210 | - | LDB1 | NM | 34 | 0,0 | 1 | 0 | 9,94E-01 | 1,00E+00 | 41 | 35 | 10 | 21 | 54 | 61 | 10 | 37 | 1,3 | 1,3 | 0,0 |
| chr6:89794879-89795379:downstream_5261 | chr6 | 89790428 | 89794879 | + | PNRC1 | NM | 37 | 0,0 | 1 | 0 | 9,94E-01 | 1,00E+00 | 13 | 8 | 17 | 12 | 55 | 49 | 72 | 70 | -0,5 | -0,4 | 0,0 |
| chr9:133452236-133452736:downstream_7362 | chr9 | 133452736 | 133454881 | - | LOC100272217 | NR | 6 | 0,0 | 2 | 0 | 9,94E-01 | NA | 3 | 9 | 1 | 2 | 9 | 15 | 7 | 2 | 1,6 | 1,5 | 0,1 |
| chr2:70888715-70889215:downstream_21725 | chr2 | 70889215 | 70995375 | - | ADD2 | NM | 5 | 0,0 | 2 | 0 | 9,94E-01 | NA | 2 | 0 | 3 | 7 | 7 | 0 | 14 | 11 | -1,9 | -2,0 | 0,0 |
| chr5:114546026-114546526:downstream_4071 | chr5 | 114546526 | 114598569 | - | PGGT1B | NM | 39 | 0,0 | 1 | 0 | 9,94E-01 | 1,00E+00 | 37 | 24 | 24 | 30 | 54 | 50 | 35 | 57 | 0,2 | 0,2 | 0,0 |
| chr8:126450647-126451147:downstream_26163 | chr8 | 126442562 | 126450647 | + | TRIB1 | NM | 38 | 0,0 | 1 | 0 | 9,95E-01 | 1,00E+00 | 24 | 26 | 7 | 17 | 77 | 75 | 47 | 30 | 1,1 | 1,0 | 0,1 |
| chrX:54593720-54594220:downstream_26839 | chrX | 54556643 | 54593720 | + | GNL3L | NM | 10 | 0,0 | 1 | 0 | 9,95E-01 | NA | 5 | 2 | 7 | 5 | 4 | 18 | 17 | 21 | -0,8 | -0,7 | -0,1 |
| chr2:85771477-85771977:downstream_21937 | chr2 | 85771977 | 85788657 | - | GGCX | NM | 30 | 0,0 | 1 | 0 | 9,95E-01 | 1,00E+00 | 31 | 47 | 45 | 34 | 22 | 19 | 18 | 23 | 0,0 | 0,0 | 0,0 |
| chr11:62618959-62619459:downstream_9403 | chr11 | 62619459 | 62623360 | - | SNHG1 | NR | 83 | 0,0 | 1 | 0 | 9,95E-01 | 1,00E+00 | 25 | 16 | 13 | 20 | 137 | 192 | 121 | 138 | 0,4 | 0,3 | 0,0 |
| chr9:33282067-33282567:downstream_26262 | chr9 | 33264876 | 33282067 | + | CHMP5 | NM | 9 | 0,0 | 2 | 0 | 9,96E-01 | NA | 23 | 17 | 3 | 7 | 15 | 2 | 3 | 2 | 2,1 | 2,0 | 0,1 |
| chr4:76455236-76455736:downstream_3744 | chr4 | 76439653 | 76455236 | + | THAP6 | NM | 11 | 0,0 | 1 | 0 | 9,96E-01 | NA | 14 | 9 | 17 | 10 | 9 | 11 | 11 | 11 | -0,2 | -0,2 | 0,0 |
| chr2:178478525-178479025:downstream_2336 | chr2 | 178479025 | 178483694 | - | TTC30A | NM | 18 | 0,0 | 1 | 0 | 9,96E-01 | NA | 10 | 4 | 11 | 11 | 19 | 24 | 35 | 33 | -0,7 | -0,7 | 0,0 |
| chr12:95364603-95365103:downstream_18703 | chr12 | 95365103 | 95397489 | - | NDUFA12 | NM | 35 | 0,0 | 1 | 0 | 9,97E-01 | 1,00E+00 | 7 | 8 | 7 | 2 | 89 | 77 | 43 | 46 | 0,7 | 0,9 | -0,2 |
| chr12:70615469-70615969:downstream_18640 | chr12 | 70615969 | 70637140 | - | LINC01481 | NR | 4 | 0,0 | 3 | 0 | 9,97E-01 | NA | 12 | 5 | 0 | 0 | 7 | 10 | 0 | 0 | Inf | Inf | NaN |
| chr15:78841563-78842063:downstream_19688 | chr15 | 78832746 | 78841563 | + | PSMA4 | NM | 478 | 0,0 | 1 | 0 | 9,97E-01 | 1,00E+00 | 43 | 17 | 32 | 13 | 1327 | 848 | 736 | 807 | 0,4 | 0,5 | -0,1 |
| chr16:69500167-69500667:downstream_12019 | chr16 | 69458497 | 69500167 | + | CYB5B | NM | 29 | 0,0 | 1 | 0 | 9,97E-01 | 1,00E+00 | 12 | 23 | 39 | 29 | 28 | 17 | 35 | 51 | -0,9 | -0,9 | 0,0 |
| chr1:247364768-247365268:downstream_1464 | chr1 | 247365268 | 247365362 | - | MIR3916 | NR | 28 | 0,0 | 1 | 0 | 9,98E-01 | 1,00E+00 | 13 | 14 | 32 | 23 | 40 | 7 | 47 | 49 | -1,0 | -1,1 | 0,0 |
| chr8:40012827-40013327:downstream_6538 | chr8 | 40010986 | 40012827 | + | C8orf4 | NM | 45 | 0,0 | 1 | 0 | 9,98E-01 | 1,00E+00 | 12 | 29 | 125 | 101 | 11 | 4 | 47 | 34 | -2,5 | -2,5 | 0,0 |
| chr2:172606668-172607168:downstream_21885 | chr2 | 172543918 | 172606668 | + | DYNC1I2 | NM | 7 | 0,0 | 2 | 0 | 9,98E-01 | NA | 10 | 8 | 0 | 13 | 10 | 4 | 7 | 4 | 0,4 | 0,3 | 0,1 |
| chr11:27387007-27387507:downstream_8860 | chr11 | 27387507 | 27494334 | - | LGR4 | NM | 93 | 0,0 | 1 | 0 | 9,98E-01 | 1,00E+00 | 2 | 4 | 22 | 20 | 38 | 41 | 298 | 317 | -2,9 | -3,0 | 0,0 |
| chr10:74766479-74766979:downstream_17689 | chr10 | 74766979 | 74856732 | - | P4HA1 | NM | 112 | 0,0 | 1 | 0 | 9,98E-01 | 1,00E+00 | 8 | 0 | 6 | 9 | 163 | 152 | 333 | 223 | -0,8 | -0,8 | 0,0 |
| chr19:33873592-33874092:downstream_21064 | chr19 | 33864574 | 33873592 | + | CEBPG | NM | 51 | 0,0 | 1 | 0 | 9,99E-01 | 1,00E+00 | 9 | 24 | 14 | 12 | 92 | 104 | 63 | 93 | 0,3 | 0,3 | 0,0 |
| chr11:62557872-62558372:downstream_9406 | chr11 | 62554873 | 62557872 | + | TMEM179B | NM | 34 | 0,0 | 1 | 0 | 9,99E-01 | 1,00E+00 | 28 | 23 | 21 | 15 | 54 | 56 | 28 | 47 | 0,5 | 0,6 | 0,0 |
| chr19:58807254-58807754:downstream_14719 | chr19 | 58790317 | 58807254 | + | ZNF8 | NM | 17 | 0,0 | 1 | 0 | 9,99E-01 | NA | 7 | 0 | 4 | 6 | 24 | 27 | 36 | 31 | -0,4 | -0,4 | 0,0 |
| chr1:153578866-153579366:downstream_1062 | chr1 | 153579366 | 153585514 | - | S100A16 | NM | 68 | 0,0 | 1 | 0 | 9,99E-01 | 1,00E+00 | 56 | 68 | 35 | 26 | 133 | 110 | 60 | 59 | 1,0 | 1,0 | 0,0 |
| chr2:122484020-122484520:downstream_2223 | chr2 | 122484520 | 122494503 | - | NIFK | NM | 63 | 0,0 | 1 | 0 | 9,99E-01 | 1,00E+00 | 8 | 12 | 3 | 7 | 159 | 155 | 89 | 72 | 1,1 | 1,0 | 0,1 |
| chr15:83657214-83657714:downstream_19482 | chr15 | 83657714 | 83680393 | - | C15orf40 | NM | 4 | 0,0 | 3 | 0 | 1,00E+00 | NA | 7 | 8 | 0 | 0 | 2 | 14 | 0 | 0 | Inf | Inf | NaN |
| chr6:133138703-133139203:downstream_5360 | chr6 | 133135707 | 133138703 | + | RPS12 | NM | 75 | 0,0 | 1 | 0 | 1,00E+00 | 1,00E+00 | 158 | 139 | 60 | 54 | 79 | 58 | 22 | 30 | 1,4 | 1,4 | 0,0 |
| chr12:4476892-4477392:downstream_9983 | chr12 | 4477392 | 4488894 | - | FGF23 | NM | 2 | 0,0 | 7 | 0 | 1,00E+00 | NA | 0 | 10 | 0 | 0 | 0 | 10 | 0 | 0 | Inf | Inf | NaN |
| chr11:128834454-128834954:downstream_17875 | chr11 | 128834954 | 129062093 | - | ARHGAP32 | NM | 2 | 0,0 | 3 | 0 | 1,00E+00 | NA | 0 | 0 | 4 | 6 | 0 | 0 | 4 | 6 | -Inf | -Inf | NaN |
| chr19:893590-894090:downstream_21378 | chr19 | 893483 | 893590 | + | RNU6-2 | NR | 4 | 0,0 | 2 | 0 | 1,00E+00 | NA | 5 | 8 | 1 | 2 | 5 | 8 | 1 | 2 | 2,1 | 2,1 | 0,0 |
