## Supplementary Table 2 for "Uncoupling from transcription protects polyadenylation site cleavage from inhibition by DNA damage"

| Gene | siRNA sequence |
|---|---|
| *CPSF160* | GCUUUAAGAAGGUCCCUCA |
| *CPSF100* | CUCAACUUCUUGAUCAGAU |
| *CPSF73* | CCAUAUACUGGUCCCUUUA |
| *CPSF30* | GUGCCUAUAUCUGUGAUUU |
| *CstF77* | GAAGACUUAUGAACGCCUU |
| *CstF64* | GGCUUUAGUCCCGGGCAGA |
| *CstF50* | GUCGUAAGUCCGUGCACCA |
| *CFIm68* | CUGCAAUUUCUUUAAUUAA |
| *CFIm25* | CCUCUUACCAAUUAUACUU |
| *CFIm59* | CUCAUCUGCUCGUGUGGAU |
| *CLP1* | GCUUAUGUCUCCAAGGACA |
| *Fip1* | CGAAUGGGACUUGAAGUUA |
| *PCF11_1* | GUACCUUAUGGAUUCUAUU |
| *PCF11_2 (pool)* | GAUACAAAUCAGCGACUUA |
| | GUGUGCAAAUUUAACGAAA |
| | AAGUUAAGGAAGAACGAAU |
| | GGAUAAGACCGAUGGCAAA |
| *hnRNP H1* | GGUAUUCGUUUCAUCUACA |
| *hnRNP F* | GGUGUCCAUUUCAUCUACA |
| *DHX36* | GGUGUUCGGAAAAUAGUAA |

siRNA sequences for all genes tested.
