## Supplementary Table 3 for "Uncoupling from transcription protects polyadenylation site cleavage from inhibition by DNA damage"

| sgRNA | Sequence | Reverse complement | PAM |
|---|---|---|---|
| **sgRNA_A** | CTGTCCTTGCCTCTGTAGAC | GTCTACAGAGGCAAGGACAG | AGG |
| **sgRNA_B** | CGTGCTGATTAATTTGATTG | CAATCAAATTAATCAGCACG | TGG |

20-mer protospacer sequences for two sgRNA and their reverse complement for the deletion of p53 CoTC
