## Supplementary Table 4 for "Uncoupling from transcription protects polyadenylation site cleavage from inhibition by DNA damage"

**Supplementary Figure 4**

| sgRNA | Sequence | Reverse complement | PAM |
| --- | --- | --- | --- |
| **sgRNA_A** | CACCGCTGTCCTTGCCTCTGTAGAC | AAACGTCTACAGAGGCAAGGACAGC | AGG |
| **sgRNA_B** | CACCGCGTGCTGATTAATTTGATTG | AAACCAATCAAATTAATCAGCACGC | TGG |

Protospacer sequences and their reverse complements with “CACC” and “AAAC” added for cloning into the pX458/pX459 vector using BbsI restriction enzyme
