## Supplementary Table 5 for "Uncoupling from transcription protects polyadenylation site cleavage from inhibition by DNA damage"

| Gene | Primer name | Sequence |
|---|---|---|
| Primer sequences | | |
| *TP53* | Forward | AGGCGATCCACCTGTCTCA |
| | Reverse R1 | TAGCCTGCACTGGCGTTC |
| | Reverse R2 | TGGAGGCTCAGCCTTGCTAA |
| | Reverse R3 | AGTACTGAGCTCCTCAACC |
| | Reverse R4 | GAGTGTTTGGCATTCCCTAGTA |
| | Reverse R5 | GAAGCAGCACAGCACAGCAGAAATAAA |
| *AKRB10* | Forward | TCTGCCAACACTGAGGATGT |
| | Reverse R1 | TTGAGCAAGTTCCTCCTCCC |
| | Reverse R2 | GGACAAACAGAAATGTTCCAGAT |
| | Reverse R3 | ACAGGGAGAGAGGGAGAGAG |
| | Reverse R4 | TATCACTGGGCTCTGGGTTG |
| | Reverse R5 | GGTAAGTCTAGCCCTCTGGA |
| *XRCC5* | Forward | AGCACCCTCATAAGTCGTCA |
| | Reverse R1 | TGAGCACCTGTATGTCAAGTT |
| | Reverse R2 | GCACAAATAATCCTGCTGCA |
| | Reverse R3 | ACTGGCAAAGGATTAAAACCCA |
| | Reverse R4 | TGTACTCCAGCCTCGGTG |
| | Reverse R5 | CTCTGCCTCCCAAAGTGCT |
| *FOS* | Forward | TGTTTGCTTATTGTTCCAAGACA |
| | Reverse R1 | CGTCCCCAGAAGCAGTAGAA |
| | Reverse R2 | GCAGGAAGATTCTAATGCCGA |
| | Reverse R3 | ACGATCAGCCATTATTTGTGC |
| | Reverse R4 | TGAACAGCAAACAGGGATCC |
| | Reverse R5 | TTGAGGTCAGGAGTTCGAGG |
| *CPS1* | Forward | AGGGCAGCCTTTGTTACTTT |
| | Reverse R1 | AGCAAGGGAGGGACAAGAAA |
| | Reverse R2 | TGGTAATCAATTGACTGTGAGGT |
| | Reverse R3 | ATGGTGATGGTGGTTGTGGT |
| | Reverse R4 | CAGCCTGCTCACTTTTAGTCA |
| | Reverse R5 | CATTGTTCAAGAGGCTGTGGA |
| *MDM2* | Forward | AGGTAGATATCTGAAAGCACCA |
| | Reverse R1 | TGTTTCAGTACCACTCCTCTCT |
| | Reverse R2 | GGAGGTTGAGGCTGTAGTGA |
| | Reverse R3 | CTCACGCCTGTAATCCCAGT |
| | Reverse R4 | TGGGGAGGTGTGAACCAAAA |
| | Reverse R5 | CTTCAAGGTGGAGTAGGGGT |
| *Puma* | Forward | CGCTGCTGTAGATACCGGAA |
| | Reverse R1 | GCCTTTCTTCTGATGGAGCC |
| | Reverse R2 | GGCTTGATCATCGCTCACTG |
| | Reverse R3 | CGTCTCGATCTCCTGACCTC |
| | Reverse R4 | GCTCGCTGTAACCTTTATCTCC |
| | Reverse R5 | CGTACAGTGGTGCAATCTCG |
| *PTEN* | Forward | AATGCCTCATCCCAATCAGAT |
| | Reverse R1 | TTCTGAACTAGCAACAGCACT |
| | Reverse R2 | TGTTGTTGTGATGGGGAAGT |
| | Reverse R3 | AGCCACTGAATTCGAAAGGA |
| | Reverse R4 | ACGCGGTAATTTTCAGAGCT |
| | Reverse R5 | GCCTCACTTCATTCCACACA |
| *FAS* | Forward | TTTGCCCCTTGTGTTTGGAA |
| | Reverse R1 | TGTGCTGTTTGGAAGAGGTC |
| | Reverse R2 | GGAACCCTAAGCAAAGCACA |
| | Reverse R3 | CCACCACAAAGAGAACCAGG |
| | Reverse R4 | AGCAGACATAATCAACAGCAACA |
| | Reverse R5 | TCCTAAAATGCAACATACGGAGA |
| *PCF11* | Forward total | AGCCGAAAAGTCACTCATAGAC |
| | Reverse total | GCCTCTTGAGTTTTGAGCAC |
| *TP53* | Forward total | AGGCGATCCACCTGTCTCA |
| | Reverse total | CAGATGTGCTTGCAGAATGT |
| | Forward uncleaved | AGGCGATCCACCTGTCTCA |
| | Reverse uncleaved | TAGCCTGCACTGGCGTTC |
| *TBP* | Forward total | GGAAGGGGCATTATTTGTG |
| | Reverse total | GCCCAGATAGCAGCACGGTA |
| | Forward uncleaved | GCAGGACAGAATATATGTGTTAATG |
| | Reverse uncleaved | CAGTATGATCACATGACTCTTACAAGG |

| | | |
|---|---|---|
| *WDR13* | Forward total | CATGGTCATCGTCTGGAGGC |
| | Reverse total | TAAGAGGGGTGGGATGGAGG |
| | Forward uncleaved | CATTCATGCATCGACGGATTC |
| | Reverse uncleaved | TAGAACAGTTCCTGGCACAC |
| *GAPDH* | Forward total | CCAAGGAGTAAGACCCCTGG |
| | Reverse total | GTACATGACAAGGTGCGGC |
| | Forward uncleaved | TACCCTGTGCTCAACCAGTTA |
| | Reverse uncleaved | CAGCTTCCTGTAGCACTCAA |
| *ABCC3* | Forward total | AACAGAAGACAGCTGCTGGG |
| | Reverse total | AATGGATTCAGGCAGCACCC |
| | Forward uncleaved | CAGTAGTCTTTTTGCACTTGTTCAC |
| | Reverse uncleaved | GTAGAAAGTCTTCCTCTTGGCCT |
| *HMGB1* | Forward total | TCGTCCCATCACAGTGTTGTT |
| | Reverse total | CTCGGGTACACAGGACACAC |
| | Forward uncleaved | GCGCCCATGTAACACAAACT |
| | Reverse uncleaved | TCCTACAATGTCTGAGCAATGG |
| *CPS1* | Forward total | TTCCCTTAAGACGATGGATTCTG |
| | Reverse total | TGTAGAAGGAATGGTGTCCTTG |
| | Forward uncleaved | AGGGCAGCCTTTGTTACTT |
| | Reverse uncleaved | AAACCAGATTCAACTGCATTACC |
| *DDRGK1* | Forward total | TGGTGTGGCTTGGTGTG |
| | Reverse total | AACAGGACTTCACCAGCTTC |
| | Forward uncleaved | AAATAGCCTGTTGCACATTTACTC |
| | Reverse uncleaved | TTAACAGAGATGTGGCCCAAG |
| *AKR1C3* | Forward total | CTGAGTCCATAGGCCAGAAAG |
| | Reverse total | ACACTACAGAACAGAGTAGGTAAAG |
| | Forward uncleaved | CCTACTCTGTTCTGTAGTGTGTG |
| | Reverse uncleaved | CCCTGTTGAGCCAGAAGAAA |
| *AKR1B10* | Forward total | GACGAGAATCGAGGTGCTGT |
| | Reverse total | TCAAGCCATGCTTTTCTGTGAT |
| | Forward uncleaved | GCGATCGATGGTCATCCTCTT |
| | Reverse uncleaved | GAAGGCAAGCTGTGAGAGCA |
| *NGFRAP* | Forward total | CCATGTGTCAAGTGGGTCTT |
| | Reverse total | CCATGCAAATGGGTGAAACTAC |
| | Forward uncleaved | CACTAGAGTGTTAATTGGTGAACAT |
| | Reverse uncleaved | ATCTCCTGACCTCGTGATCT |
| *FOS* | Forward total | TTGTTGAGGTGGTCTGAATGT |
| | Reverse total | CTTGGAACAATAAGCAAACAATGC |
| | Forward uncleaved | AGTTGAATGCGACCAACCT |
| | Reverse uncleaved | GTCCTCTTTGATAAGGGATCAGAC |
| *XRCC5* | Forward total | TTGTGGATGGTGTCTCCTTTAC |
| | Reverse total | CACCAAAGAGGAACTGGAACT |
| | Forward uncleaved | GCTGAGAATTGAACACCCTTATC |
| | Reverse uncleaved | GATGTCCTAGAAGCCCAAAGTA |
| *FAM60A* | Forward total | GCTGCAGTATTGGTGGTAGAA |
| | Reverse total | CAGTACATCCTACAGGCAAAGAG |
| | Forward uncleaved | GTACTGTATGTAGTCATGCACTTTG |
| | Reverse uncleaved | CCTGTCAAACAAAGCCACAA |
| *EIF4EBP2* | Forward total | TGTCTCCCATGATGTGTTGTT |
| | Reverse total | CACACAGGACTGCCTCAAG |
| | Forward uncleaved | TTCTGGTGAAATCCTGCTAAGG |
| | Reverse uncleaved | AGTGTGGAGAACTGACAGATAAAG |
| *ZRANB2* | Forward total | GCTGTACTAAGCAAATGCAAGG |
| | Reverse total | TGCTTGACTCACAGGCTTTAT |
| | Forward uncleaved | ATTCCAAAGCCATTATCACTGC |
| | Reverse uncleaved | TCAGGAAGCACACTACGATATG |
| *CTNNB1* | Forward total | GTATGGGTAGGGTAAATCAGTAAGAG |
| | Reverse total | TCTCTTGAAGCATCGTATCACAG |
| | Forward uncleaved | CTGTGATACGATGCTTCAA |
| | Reverse uncleaved | ACCACCCTCACAAACCATTTA |
| *PRDX6* | Forward total | TTCCGATGATGTGTACATGAAAGA |
| | Reverse total | AAATAGCAACCCACTGCAAGA |
| | Forward uncleaved | GGGTCAGAGAATTCTGTTGTCATA |
| | Reverse uncleaved | CACGTTCTTCAGCTGTTCCT |
| *MRPL32* | Forward total | GGAAGATTCTTTATGTTGTTGTGCT |
| | Reverse total | AATCCATTGAGCCTTTGGATAAAC |
| | Forward uncleaved | CCAAAGGCTCAATGGATTATGT |
| | Reverse uncleaved | AAAGGCACTGGCAAACAAA |
| *TBK1* | Forward total | CAGAACCGCACCACTGTTA |
| | Reverse total | GGATACAAGGATAACTGGGATCTG |
| | Forward uncleaved | AGAGTTCATGTGTTTCTTTGTATCC |
| | Reverse uncleaved | TTGTCCCTAGATCCAATATTCTGAG |

| | | |
|---|---|---|
| *HSPD1* | Forward total | ACCAGTGTACTGCTTTCAACT |
| | Reverse total | AAGGCTGCTTAACTTCTCATCT |
| | Forward uncleaved | GATGAGAAGTTAAGCAGCCTTTC |
| | Reverse uncleaved | CTCCCAAGTAGCTGGGATTA |
| *ARF6* | Forward total | GAAACACAGCAGTTCTTGGTAAAG |
| | Reverse total | AGCCATCTACAGCAAGTGATAAG |
| | Forward uncleaved | ACTATGTTGCAAGTCTGTTTCATC |
| | Reverse uncleaved | CCACTGTGGGCTAAGTTTACTA |
| *MDM2* | Forward total | CGCTTTATGGGTGGATGCTG |
| | Reverse total | ATTGAAAGCTGGCTACATGGT |
| | Forward uncleaved | CACCAGCACTTGGAAGGTGT |
| | Reverse uncleaved | GAGTACAGCAATCATTTCAGATGC |
| *PUMA* | Forward total | GAGATTTTGGCTGAAGCCGC |
| | Reverse total | CAGTATCTTACAGGCTGGGC |
| | Forward uncleaved | GCTGCTGTAGATACCGGAATGA |
| | Reverse uncleaved | AGGGAAGGCAAGCAGAAAGA |
| *PTEN* | Forward total | AGCAGTGGCTCTGTGTGTAA |
| | Reverse total | CATCTGATTGGGATGAGGCA |
| | Forward uncleaved | TCTTGTCATTGTGTGGGTGT |
| | Reverse uncleaved | AGGCTTTGAAGGACAGCAGG |
| *FAS* | Forward total | AGCAGATACCTGGAACCACC |
| | Reverse total | TTATAATTCCAAACACAAGGGGC |
| | Forward uncleaved | AGCAGATACCTGGAACCACC |
| | Reverse uncleaved | GAGTACAGCAATCATTTCAGATGC |
| *NOXA* | Forward total | AGGTTGTAGTCACTTTAGATGGAA |
| | Reverse total | TACCAGATGGTAAAATAGCTGCCT |
| | Forward uncleaved | AAGTTGATACTGTGGCAGTAAAC |
| | Reverse uncleaved | GTCTGCTGATGGAAATCAGTTAAA |

Oligonucleotides: Primer sequences used for all genes tested.
